# Image-based DNA Sequencing Encoding for Detecting Low-Mosaicism Somatic Mobile Element Insertions

**DOI:** 10.1101/2024.11.07.619809

**Authors:** Miaomiao Tan, Zhinan Lin, Zhuofu Chen, Haonan Zhou, Junseok Park, Ziting He, Eunjung A. Lee, Zhipeng Gao, Xiaowei Zhu

**Affiliations:** Key Laboratory of Artificial Organs and Computational Medicine in Zhejiang Province, Institute of Translational Medicine, Zhejiang Shuren University, Hangzhou, China; Department of Neuroscience, City University of Hong Kong, Hong Kong SAR, China; Division of Genetics and Genomics, Boston Children’s Hospital, 3 Blackfan Circle, Boston, MA, USA; Department of Pediatrics, Harvard Medical School, 25 Shattuck St, Boston, MA, USA; Broad Institute of MIT and Harvard, Cambridge, MA, USA; Shanghai Institute for Advanced Study, Zhejiang University, Hangzhou, China

**Author notes:** Correspondence should be addressed to: Xiaowei Zhu. These authors contributed equally to this work (Miaomiao Tan, Zhinan Lin).

## Abstract

Active LINE-1 (L1), *Alu*, and SVA mobile elements in the human genome are capable of retrotransposition, resulting in novel mobile element insertions (MEIs) in both germline and somatic tissues. Detecting MEIs through DNA sequencing relies on supporting reads overlapping MEI junctions; however, artifacts from DNA amplification, sequencing, and alignment errors produce numerous false positives. Systematic detection of somatic MEIs, particularly those with low mosaicism, remains a significant challenge. Previous methods had required a high number of supporting reads which limits the detection sensitivity, or human inspections that are susceptible to biases. Here, we developed RetroNet, an algorithm that encodes MEI-supporting sequencing reads into images, and employs a deep neural network to identify somatic MEIs with as few as two reads. Trained on extensive and diverse datasets and benchmarked across various conditions, RetroNet surpasses previous methods and eliminates the need for extensive manual examinations. The RetroNet analysis on the Illumina sequencing of 161× or 195× of a cancer cell line achieved an average precision of 0.885 and recall of 0.579 for detecting somatic L1 insertions that are present in as few as 1.79% of the cells. Additionally, we demonstrated that RetroNet is effective for analyzing highly degraded DNA, such as circulating tumor DNA. RetroNet is applicable to the rapidly generated short-read sequencing data and has the potential to provide further insights into the functional and pathological implications of somatic retrotranspositions.

## Introduction

Mobile element (ME), or transposon, is a class of short DNA fragments capable of changing genomic locations, including cut-and-paste DNA transposon and copy-and-paste retrotransposon. About 45% of the human genomic sequence consists of ME^1^. Retrotransposons are the dominant class, including long terminal repeat (LTR) transposons such as ∼ 98,000 human endogenous retroviruses and non-LTR transposons such as ∼560,000 L1 (or LINE-1), ∼1,200,000 *Alu*, and ∼5,100 SVA (or SINE-VNTR-*Alu*s) elements. While most human MEs have accumulated with many mutations in evolution and are incapable of moving, a small number of non-LTR retrotransposons remain active, including ∼80 active L1 (L1Hs subfamily)^2^, over 800 active *Alu* (*Alu*Y subfamilies)^3^, and ∼25 active SVA (SVA-E and SVA-F subfamilies) elements per person^4^. These active retrotransposons can be copied to new genomic loci via a retrotransposition process such as target-primed reverse transcription (TPRT)^5^, creating *de novo* mobile element insertion (MEI) mutations^6^. Retrotranspositions can occur in germline and somatic cells, yielding inter-individual and intra-individual genetic diversities. More than one hundred MEIs have been linked to human diseases^7,8^, including somatic SVA insertions, found in over 75% of the cells (termed tissue allele frequency, or tAF), within the *NF1* gene in patients with Neurofibromatosis Type I^9^.

Widespread somatic retrotranspositions with various tAFs have been identified in normal tissues^10^ as well as cancer^11^. Somatic L1 insertions can occur in neural stem cells^12^ and mature neurons^13^, typically resulting in a tAF below 2% per insertion in the adult human brain^14–16^. Different L1 mutations create genomic mosaicism among various brain cells through insertional mutagenesis and alterations in transcriptional regulation^17^. In cancer, somatic insertions from *Alu* and SVA elements, along with L1, have been extensively documented, with the retrotransposition activities dependent on both the type of mobile element and the tumor’s origin^11,18,19^. While driver mutations in tumors usually have high tAFs, clinically significant mutations can still occur at low frequencies due to low tumor purity or the presence of new subclonal populations that confer resistance to treatment^20^.

The characterization of low-tAF somatic MEIs remains challenging due to weak signals and abundant noise produced by current high-throughput DNA sequencing technologies^21^. The standard whole genome sequencing (WGS) approach with short, paired-end sequencing reads can reveal evidence of *de novo* MEI mutations, including supporting reads with one read-end mapped in the human reference genome, and the other end mapped to mobile element consensus sequences. Previous studies on brain or tumor somatic MEIs have utilized various approaches, including whole genome or targeted sequencing of bulk tissue or single cells^14,16,22–24^. To capture the somatic mutations of low tAFs, these methods typically employ high sequencing depth (greater than 100×), ME-targeted PCR amplification, or enzymatic whole-genome amplification of single cells. However, these approaches can increase the likelihood of sequencing artifacts, such as PCR chimeras that link unrelated sequences. When the chimeras form around the abundant mobile elements in the human reference genome, they can mimic low-frequency, novel ME junctions, leading to a high number of false somatic MEI discoveries. Standard MEI detection algorithms generally require a minimum of four supporting reads^18^ or five supporting reads^25–28^ that cover both upstream and downstream insertion junctions. While this requirement helps reduce noise, it greatly limits the ability for detecting low-tAF MEIs that do not reach the threshold of supporting reads.

*Bona fide* ME retrotransposition exhibits hallmark features such as specific alleles^29^ and high sequence identity to the active ME consensus sequences^7^, which can be utilized to differentiate it from the randomly generated false ME junctions. RetroSom, a random-forest model, was previously designed to classify a single supporting read as true or false based on its sequence^16^. With further manual inspections for proper read positions using a visualization tool, RetroVis, RetroSom reported somatic MEIs as those with a minimum of two supporting reads. The RetroSom and RetroVis analysis, however, still have many limitations. First, it could not detect somatic SVA insertions, and the sensitivity for low-tAF L1 and *Alu* MEIs remains low. Second, the training data of RetroSom came from 11 individuals of one Caucasian family^30^ – this limited scope could lead to lower accuracies when applied to individuals of different ancestries. Lastly, the manual inspection with RetroVis, despite its effectiveness in reducing false positives, requires substantial prior knowledge and is prone to human biases. Nevertheless, the graphic design used to visualize MEIs can act as a strong foundation for deep learning techniques. These techniques have recently advanced significantly in various fields of genomics^31^, improving the detection of single nucleotide polymorphisms, short insertion-deletion mutations^32^, and DNA structural variants^33,34^ such as deletions, duplications, inversions, and complex variants.

Here, we formulated the RetroNet framework that uses image encoding to integrate both the sequence and positional features of candidate MEIs and employs a deep neural network to predict somatic L1, *Alu*, and SVA insertions in the human genome (**Figure 1)**. The training data consisted of high-coverage WGS from 549 parent-offspring trios with diverse ancestries, significantly improving the model’s generalizability^35^. Following the exclusion of low-quality reads, candidate MEIs were labeled as true or false based on the inheritance patterns. Groups of two supporting reads were encoded into images to illustrate their alignments to ME consensus sequences and their relative positions. Next, we trained three neural network classifiers based on state-of-the-art architectures for image processing: ResNet-18^36^, GoogLeNet^37^, and Vision Transformer (ViT)^38^. The performance was benchmarked in independent and diverse test datasets. This included (1) detecting germline MEIs in the Illumina Polaris Project^39^, (2) detecting simulated somatic MEIs with a wide range of tAFs in a genome mixing dataset^16^, and (3) detecting simulated MEIs across various signal-to-noise ratios. To demonstrate the accuracy in real datasets, we analyzed somatic MEIs from paired tumor-normal sequencing of a patient with pancreatic ductal adenocarcinoma, HG008^40–42^. Additionally, we detected somatic MEIs in the cell-free DNA (cfDNA) from a metastatic castration-resistant prostate cancer patient, DTB-205^43^, and benchmarked with the matching tumor tissue. All relevant datasets are summarized in **Supplementary Table 1**.

**Figure 1.**
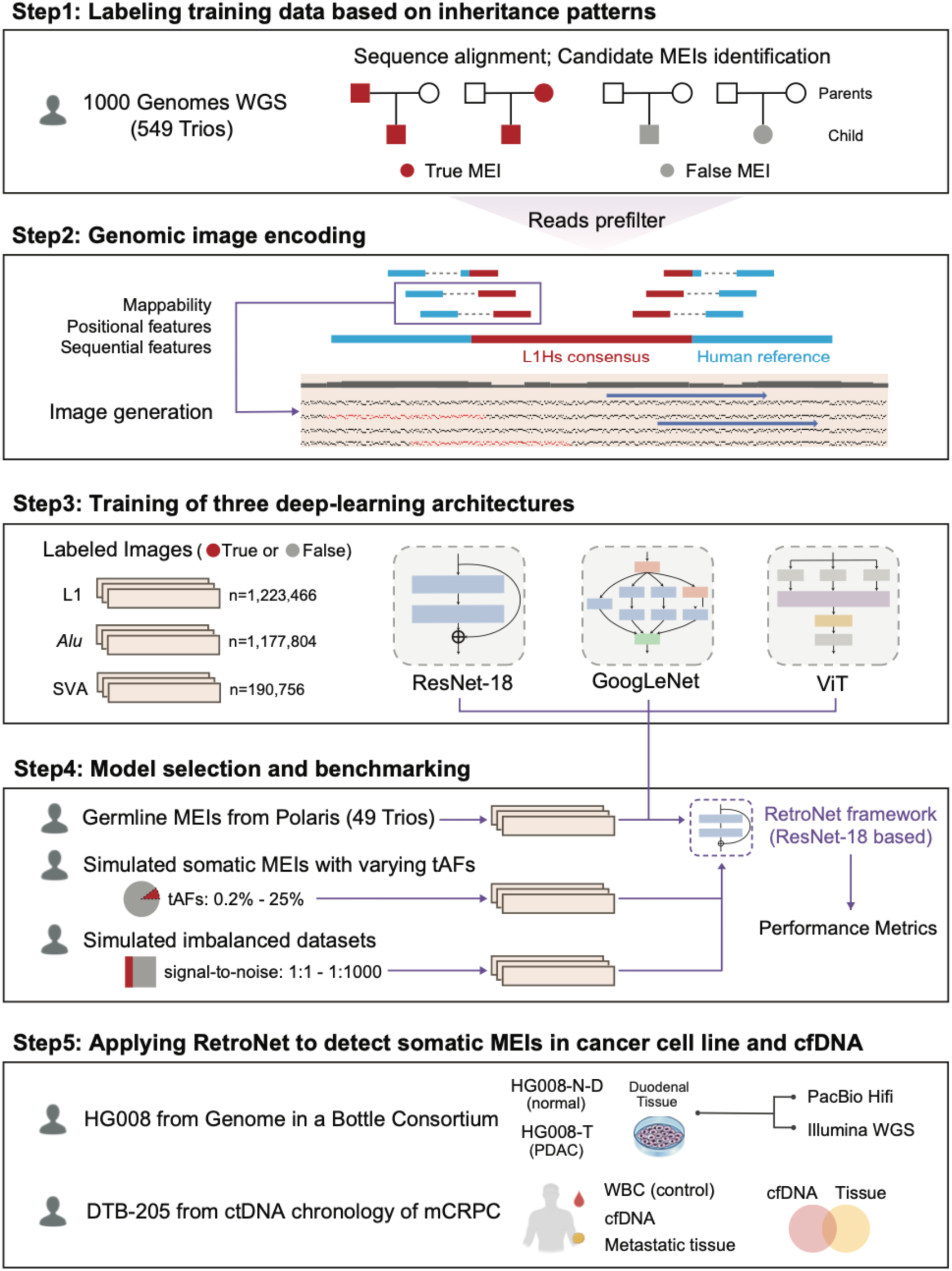
Schematic overview of the RetroNet framework. The RetroNet pipeline was developed in five major steps. First, we extrapolated candidate germline MEIs from 549 trios of the 1000 Genomes Project, and labeled true (red) and false (gray) MEIs based on the inheritance patterns. Second, we used an image-based encoding to convert groups of two MEI supporting reads, each shown as a pair of blue (flanking sequence in the human reference genome) and red (ME sequence) boxes. Two reads supporting an L1 insertion were transformed into an image formatted into nine tracks. These tracks contain both the sequence-L1 alignment represented by black and red dots, as well as the relative positions of the flanking sequences indicated by blue arrows. Third, we trained three deep-learning models based on ResNet-18, GoogLeNet, and vision transformer (ViT) architectures to classify the labeled L1, *Alu,* or SVA images. Fourth, we benchmarked the trained models in three independent datasets, including germline MEIs (for model selection), somatic MEIs with low tissue allele frequencies (tAFs), and simulated imbalanced datasets with varying signal-to-noise ratios. Finally, we applied RetroNet to detect somatic MEIs in paired tumor-normal sequencing for a pancreatic ductal adenocarcinoma (PDAC) tumor cell line HG008-T, using matched normal duodenal tissue (HG008-N-D) as a control. Furthermore, we analyzed somatic MEIs in matched cfDNA and metastatic tissue samples from a metastatic castration-resistant prostate cancer (mCRPC) patient DTB-205.

RetroNet extends the detection of somatic MEIs to SVA elements. Based on the area under the precision-recall curve (AUPR), a metric suitable for binary classification of rare events^44^, RetroNet outperforms RetroSom in detecting both germline and somatic L1 and *Alu* insertions. In detecting germline L1 elements from the Polaris dataset, RetroNet achieved an average AUPR score of 0.990 (95% confidence interval (CI): 0.988-0.993), compared to RetroSom’s 0.936 (95% CI: 0.929-0.944). Similarly, in detecting simulated somatic L1 insertions at 1% tAF with 200× WGS, RetroNet achieved a 43.4% increase in AUPR over RetroSom (0.223 vs. 0.156). With a more stringent cutoff tuned for highly imbalanced datasets, RetroNet has significantly improved precision while maintaining good recall for detecting low-tAF somatic L1 insertions in the genomic DNA of a cancer cell line, as well as in the cell-free DNA from a cancer patient. Finally, we interpreted the RetroNet neural network and confirmed that it could utilize hallmark sequencing and positional parameters from *bona fide* MEIs to guide the detection. The code and environment of RetroNet were packaged in a container (https://github.com/Czhuofu/RetroNet).

## Results

### Training RetroNet using an image-based encoding of DNA sequencing reads

Due to the limited number of *bona fide* somatic MEIs, we developed RetroNet by utilizing transfer learning from the more abundant, evolutionarily recent germline MEIs that share the same retrotransposition mechanisms. The training data were true and false MEIs detected in 549 father-mother-offspring trios from the 1000 Genomes Project high coverage (average > 30×) WGS dataset^35^, after excluding 53 trios that overlap with the benchmarking datasets (**Supplementary Table 1**). True and false L1, *Alu*, and SVA insertions were labeled in each offspring based on their inheritance patterns: true MEIs were those present in the offspring and one parent, while false insertions were found only in the offspring. MEIs that were present in all three family members were excluded, as these could represent either evolutionarily old MEIs with high population frequencies or common alignment errors. We further excluded genomic loci with repetitive sequences that are prone to alignment errors and ME junctions formed by DNA structural variations instead of retrotranspositions. Using a threshold of two or more supporting reads, we identified 287,096 true MEIs (22,629 L1, 250,874 *Alu*, and 13,593 SVA) and 1,023,785 false MEIs (834,656 L1, 97,755 *Alu*, and 91,374 SVA) (**Supplementary Table 2** and **Supplementary Data 1-3**). The class labels were validated using long-read sequencing from 11 randomly selected individuals (**Supplementary Note 1**). Most of the MEIs in the training set have a population frequency of less than 0.1 across all individuals (**Supplementary Figure 1**), and therefore the risk of over-representing subsets of MEIs was low. The supporting reads of the labeled MEIs can be classified into three groups, including (1) split reads (SRs) with one read-end map onto the MEI junction and the other to the flanking human sequence, (2) paired-end reads (PEs), with one read-end fully aligned to the ME consensus sequence and the other end to the flank, and (3) clipped paired-end reads (clipped PEs), in which one read-end maps to the ME consensus sequence and the other to the MEI junction (**Figure 2a**). Both SRs and clipped PEs could identify the exact MEI junctions, whereas PE supporting reads only outline an approximate junction window.

**Figure 2.**
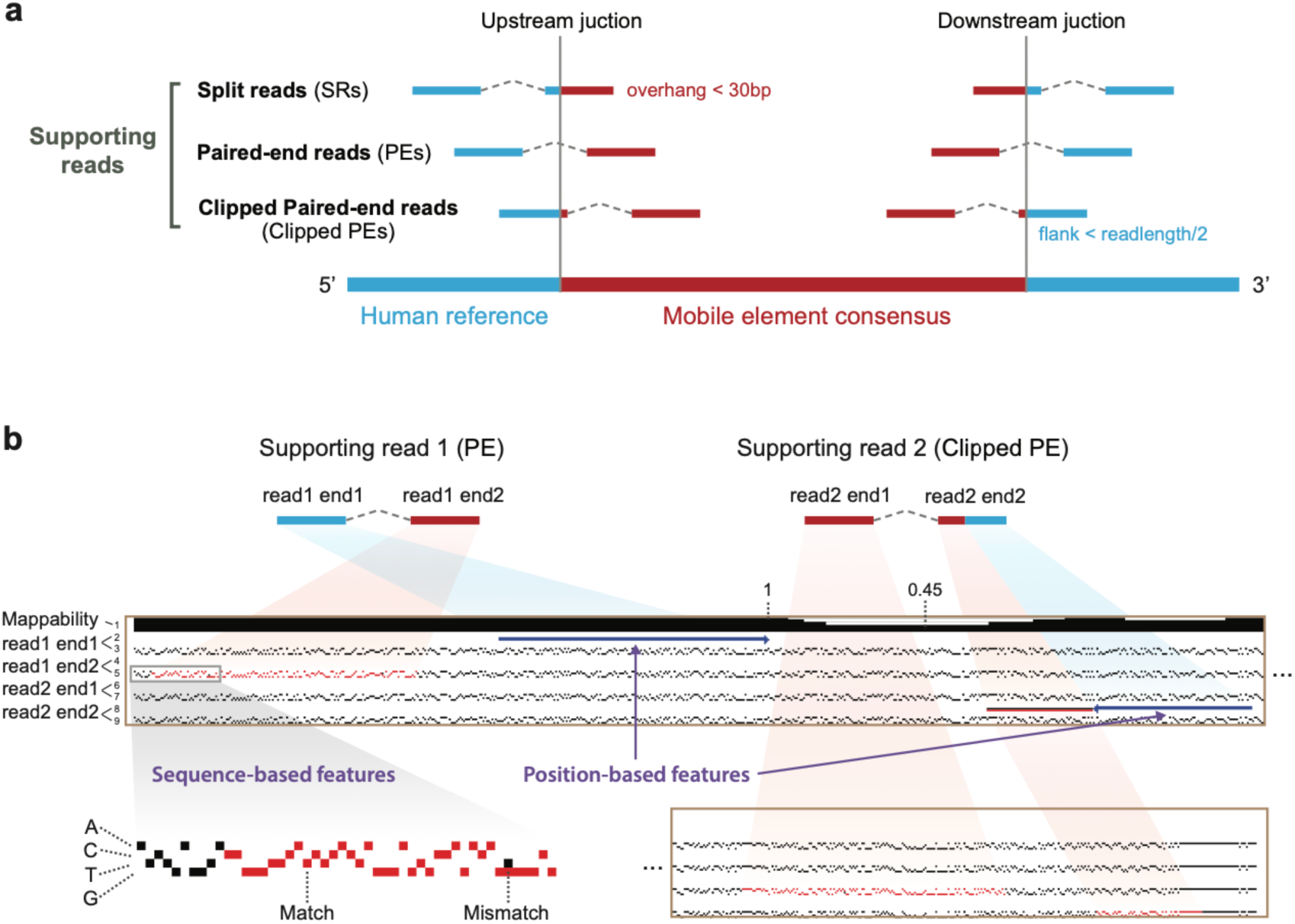
Image-based Encoding of MEI candidate supporting reads. (**a**) The types of supporting read pairs used to detect MEI include split-reads (SRs), paired-end reads (PEs) and clipped paired-end reads (clipped PEs). Blue indicates the segment of the supporting read that mapped to the flanking sequence, while red denotes the segment that mapped to the mobile element (ME) consensus sequence. For SRs, the mapped segment to the ME consensus must be greater than 30 bp, while clipped PEs require that the mapped region to the flanking sequence is higher than half of the read length to ensure mapping accuracy. (**b**) An example of encoding two supporting reads of a candidate L1 insertion into a three-channel image with nine tracks, by encapsulating both position-based features (relative read positions) and the sequence-based features (read-ME alignment). The L1 consensus sequence is divided into the 5’ end (top) and the 3’ end (bottom), with a further zoom in to the 5’ end to illustrate the encoding syntaxes. Briefly, the first track indicates the mappability of 100 bp DNA fragments from the human genomic region near the insertion. A fully black bar represents a genomic region with a mappability of 1, indicating all DNA fragments can be uniquely and properly aligned. The subsequent tracks provide a detailed depiction of the supporting reads’ position in the human genome (tracks 2, 4, 6, 8) and the alignment to the active L1 (L1Hs) consensus sequence (tracks 3, 5, 7, 9). For example, read 1 is a PE supporting read with end1 mapped in the human genome and end2 mapped in L1. We denote the end1 position with a blue arrow in track 2 and the L1 alignment of end2 by red pixels in track 5. In track 5, the L1Hs consensus sequence is denoted by a matrix where columns represent the base positions and rows represent the nucleotides A, C, T, and G, from top to bottom. The read sequence aligning to L1Hs is highlighted in red pixels. A mismatch appears as a column with both black (L1Hs) and red (supporting read) pixels. The other read, read 2, is a clipped PE supporting read. The shorter blue arrow in track 8 indicates that read2 end2 partially maps to the human genome, with the unmapped portion represented by a proportional black line. The proportional red line below the black line indicates that this portion maps to L1Hs, and its sequence is displayed as red pixels in track 9. Additional encoding syntaxes for *Alu* and SVA are shown in **Supplementary Figure 2**.

We encoded groups of two supporting reads into fixed-size images to integrate both the sequence-based features, such as hallmark alleles of the active ME subfamilies (e.g., L1Hs, *Alu*Ya5, and SVA-E)^29^, and positional features, such as the relative arrangement of the supporting reads. For L1 insertions, two supporting reads, read 1 and read 2, each with two paired sequencing ends, end1, and end2, are encoded into an image of 60×6620 pixels that can be divided into nine tracks of information (**Figure 2b**). Track 1 illustrates the mappability^45^ of the flanking sequences, chosen as a window of 500 bp upstream and downstream to the putative insertion junction. The mappability is defined as the degree of local, 100 bp short sequence fragments that could be uniquely and properly aligned, and the values range between 0 and 1. A full black bar represents a genomic region with a mappability of 1, or the DNA alignment of this region is highly reliable. Track 2 to 9 depict each of the four sequencing read ends, read1 end1, read1 end2, read2 end1, and read2 end2, that were mapped to either the flanking human genome, the active L1Hs consensus sequence^46^, or both (for SR and clipped-PE reads). Each end is encoded into two tracks, with the top track row featuring the possible alignment in the flank, shown as blue arrows to indicate the relative position. The bottom track features a one-hot encoding of the consensus L1Hs sequence – a 4×6064 matrix with white-black pixels, where the columns denote the base positions and rows represent A, C, T, or G from top to bottom. The alignment of the supporting read is denoted by overlaying a string of red pixels, representing the nucleotides from the read, to the L1Hs sequence. One sequence mismatch will, therefore, be encoded into the coexistence of a black pixel (L1 reference) and a red pixel (supporting read) in one column. Several additional encoding syntaxes are implemented to ensure a consistent and comprehensive sequence-to-image conversion (see Methods and **Supplementary Figure 2a-d**). Finally, to avoid overrepresenting MEIs with a higher number of supporting reads (e.g., homozygous over heterozygous), we chose to include a maximum of five randomly selected images per insertion. The resulting training dataset for L1 MEIs contains 135,774 true images and 1,087,692 false images (**Supplementary Table 2**). The encoding of *Alu* and SVA insertions are similar to L1, with the exception of including multiple consensus ME sequences to represent the different active ME subfamilies (**Supplementary Figure 2e**).

We adopted state-of-the-art deep learning models to solve the binary classification of the true MEI-derived images from the false ones. We included two convolutional neural networks (CNN) architecture-based models: ResNet-18 and GoogLeNet. As the encoded images are organized into horizontal tracks and may challenge the spatial locality principle of the CNN models^47^, we incorporated a third model, the Vision Transformer (ViT), which allows the partitioning of the input images into ordered tracks and then utilizes a transformer architecture^48^ for image analysis. The architectures of the three deep learning models, along with the output sizes of each layer, are presented in **Supplementary Table 3**. By the 30th epoch, the loss function values in the validation dataset, chosen as 10% of the images, had converged and remained stable (**Supplementary Figure 3a**). Ultimately, the models achieved AUPR values of 0.997 for ResNet-18, 0.998 for GoogLeNet, and 0.994 for ViT for L1 classification in the validation set. High-performance values were also observed in the models for *Alu* and SVA classifications (**Supplementary Figure 3b**).

### Comparable accuracies from three neural network models in detecting germline MEIs

To evaluate the trained ResNet-18, GoogLeNet, and ViT models, we benchmarked their performance in detecting germline MEIs in 49 family trios from the Illumina Polaris Project (**Supplementary Table 1**). Notably, the same individuals were also part of the 602 trios sequenced by the 1000 Genome Project but were excluded from model training to avoid data leakage. Using the same labeling process and a threshold of at least two supporting reads, we identified a total of 26,960 true MEIs (2,381 L1, 23,287 *Alu*, and 1,292 SVA) and 31,566 false MEIs (15,844 L1, 11,145 *Alu*, and 4,577 SVA) from the 49 offsprings (**Supplementary Table 2** and **Supplementary Data 4-6**). Following the sequence-to-image encoding, we evaluated the trained ResNet-18, GoogLeNet, and ViT models in classifying the true and false images (**Supplementary Table 4**). For L1 and *Alu* insertions, specifically, we also performed the RetroSom analysis and compared its performance with the three deep learning models.

All three deep learning models achieved similar AUPR scores for the classification of L1-derived images: ResNet-18: 0.990 (95% CI: 0.988-0.993), GoogLeNet: 0.990 (95% CI: 0.988-0.993), ViT: 0.991 (95% CI: 0.988-0.993), significantly outperforming the RetroSom model (AUPR=0.936, 95% CI: 0.929-0.944, *P* < 2e-16) (**Figure 3**). Other performance metrics showed similar results: ResNet-18 and GoogLeNet exhibited nearly equal precision (=0.970) and recall (=0.956) for L1 detection at the default classification cutoff (Probability = 0.5). In comparison, ViT exhibited a lower average precision of 0.961 (*P* = 1.3e-7) but a higher recall of 0.976 (*P* = 4.6e-8) (**Figure 3b**). Similar results were also observed for benchmarking the *Alu-*derived and SVA-derived images (**Figure 3**). The results indicate that the three deep learning models exhibit similar overall performance, which surpasses that of RetroSom. Among the three models, ResNet-18 and GoogLeNet are less computationally intensive and have fewer parameters than ViT (**Supplementary Table 3**). ResNet-18’s residual blocks are simpler, more scalable, and more interpretable than GoogLeNet’s inception modules^37^. This makes ResNet-18 more efficient in training and inference^49^, and thus we chose ResNet-18 as the default model implemented in the RetroNet framework.

**Figure 3.**
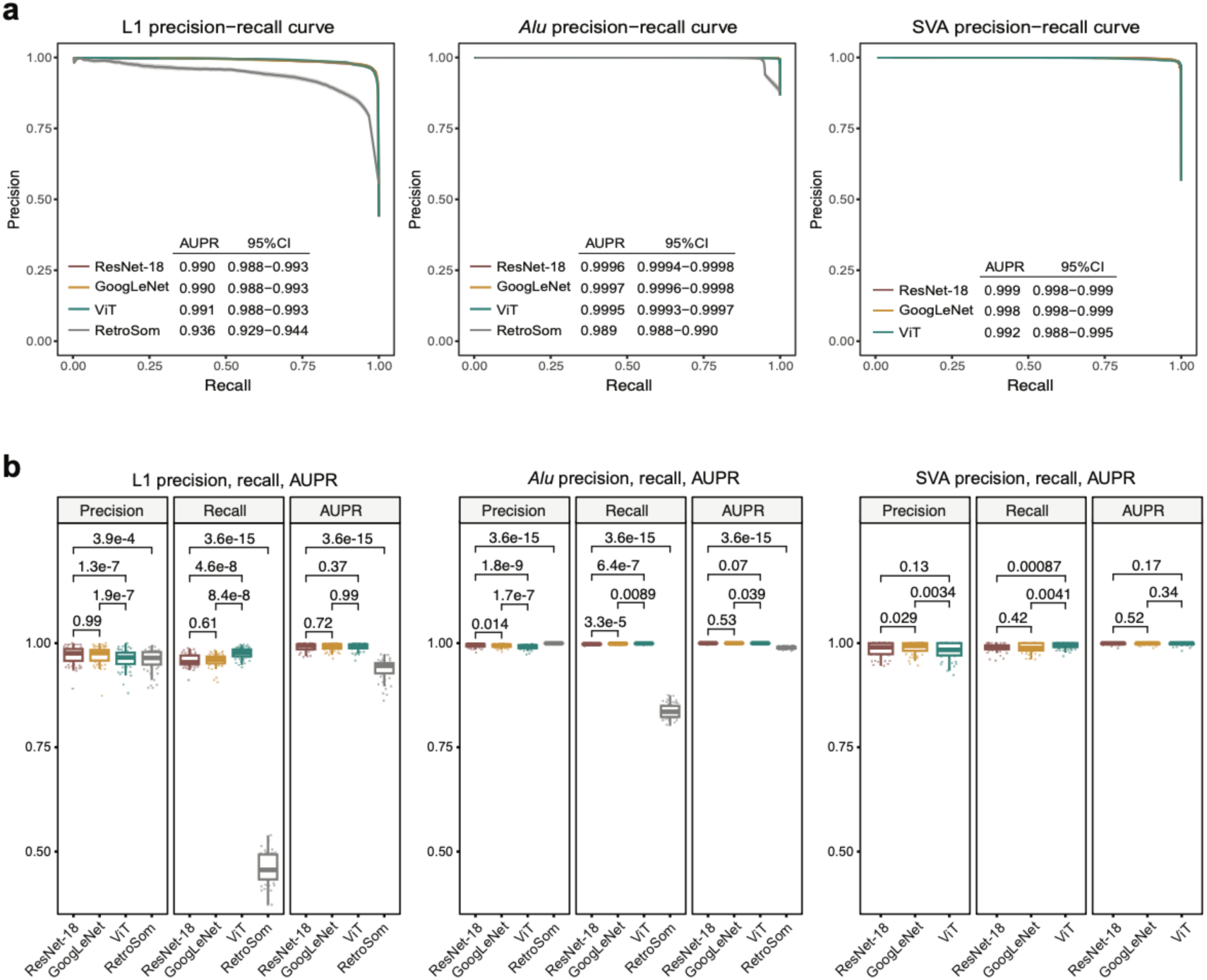
Benchmarking ResNet-18, GoogLeNet, and ViT models using independent germline MEIs. (**a**) Precision-recall curves of the trained ResNet-18, GoogLeNet, ViT models, and the RetroSom model (for L1 and *Alu*) were assessed on the labeled L1, *Alu*, and SVA image datasets extracted from 49 offspring in the trio sequencing of the Illumina Polaris Project. We labeled the average and 95% confidence intervals of the area under the precision-recall curve (AUPR) scores for each model. (**b**) Boxplots to compare the precision, recall, and AUPR scores of the ResNet-18, GoogLeNet, ViT models, and the RetroSom model, using the default threshold of probability > 0.5. The boundaries of the boxplots represent the first quartile (Q1, 25^th^ percentile) below and the third quartile (Q3, 75^th^ percentile) above, with the black line inside the box marking the median. The whiskers (vertical lines) extend above the box to the largest data point ≤ Q3 + 1.5 * IQR and below the box to the smallest data point ≥ Q1 - 1.5 * IQR, where IQR is the interquartile range (Q3 - Q1).

### RetroNet outperforms previous methods in detecting simulated somatic MEIs with low levels of mosaicism

Unlike germline MEIs that are present in all cells, somatic MEIs are mosaic with various levels of tAFs, which pose additional challenges in the detection^50^. Thus, we benchmarked RetroNet at detecting MEIs with various levels of mosaicisms using a genome mixing dataset^16^ to simulate somatic MEIs at a range of tAFs (**Supplementary Table 1**). The dataset contains 50×, 100×, 200× and 400× coverage WGS of a mixture of six genomic DNA from unrelated individuals, of which true MEIs have been defined previously^16^, into the NA12878 genomic DNA at ratios of 0.04%, 0.2%, 1%, 5%, and 25%. A heterozygous germline MEI found in only one of the six genomes thus could simulate somatic MEIs, with a tAF from 0.04% to 25%. We then identified candidate MEIs that are present in the DNA mixture but absent in NA12878, chose a maximum of 10 pairs of supporting reads per insertion, and then applied the RetroNet or RetroSom (for L1 and *Alu* only) analyses. Somatic MEIs were reported if any pair of the supporting reads was predicted to have a probability above the selected stringency cutoff. Additionally, we compared the results with a separate MEI detection method, xTea (short-read module)^51^.

At a sequencing coverage of 200×, both RetroNet and RetroSom could detect the simulated somatic L1 insertions with a tAF as low as 1%, with RetroNet outperforming RetroSom. For L1 of 1% tAF, RetroNet’s AUPR was 0.223, while RetroSom’s AUPR was 0.156. Notably, the theoretical maximum of the AUPR was 0.238, since the 200× coverage is insufficient to sample all 1% L1 insertions with two or more supporting reads. RetroNet reached an optimal F_1_ score of 0.356, defined as the harmonic mean of the recall and precision, at a probability cutoff of 0.8 (recall=0.238, precision=0.714). Comparatively, RetroSom’s optimal F_1_ was 0.250 (recall=0.143, precision=1). Similarly, at 5% tAF, RetroNet’s AUPR was 0.861, and RetroSom’s was 0.852 (maximum = 0.875), while at 25% tAF, both RetroNet and RetroSom achieved an AUPR of 1 (**Figure 4**). When considering the other sequencing coverages, from 50× to 400×, we found that RetroNet consistently outperforms RetroSom, and AUPR scores were positively correlated with sequencing coverage, which determines the number of supporting reads (**Figure 4**). The only exception was that 200× coverage outperformed the 400× coverage data when detecting L1 insertions with 5% tAF. This was due to the higher sequencing depth of the control (400× NA12878) contained additional sequencing noise that masked a true L1 insertion at chr3:43221180^52^, which is present in the genome that was mixed at 5%. RetroNet’s prediction performance showed similar patterns for *Alu* and SVA insertion, with AUPR positively correlated with sequencing depth and tAF and consistently outperforming RetroSom for *Alu* detection (**Supplementary Figures 4** and **5**). Finally, xTea achieved perfect scores in identifying MEIs with a tAF of 25% in the 400× dataset. However, its performance declined at lower tAFs or reduced sequencing depth. This aligns with the requirement of a relatively large number of supporting reads by xTea, which may explain its diminished efficacy for low tAF somatic mutations.

**Figure 4.**
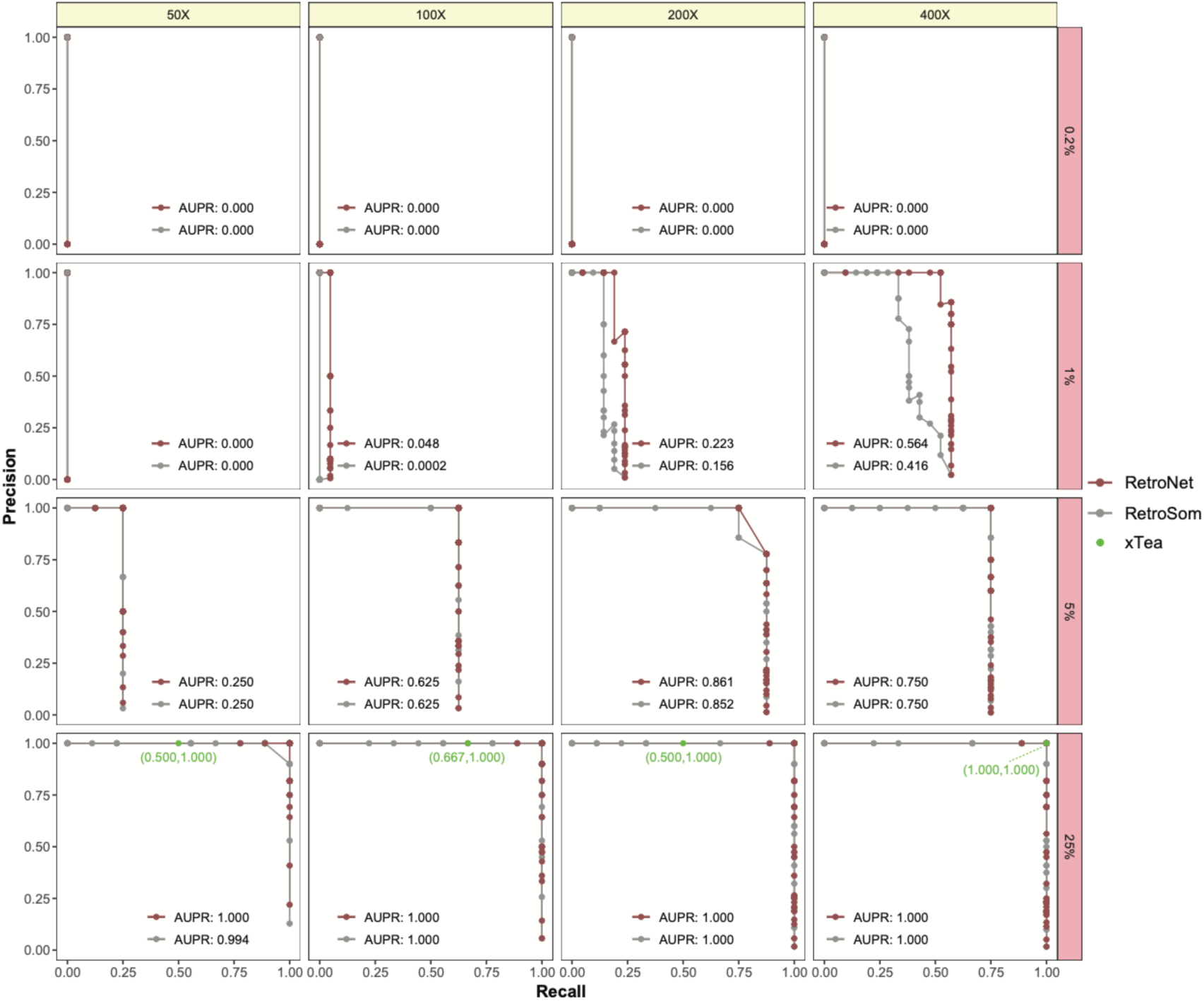
Identification of simulated somatic L1 insertions with various frequencies and sequencing depth. The precision-recall curves were evaluated for detecting somatic L1 insertions at mixing proportions of 0.2%-25% (y-axis) under sequencing depth of 50× to 400× (x-axis), using RetroNet (red), RetroSom (gray) and xTea (green). Based on the AUPR scores, RetroNet could consistently outperform RetroSom at detecting low-mosaicism L1 MEIs. In contrast, xTea only predicts MEIs with relatively high tAF and performs significantly worse than RetroNet. Similar results were also observed for *Alu* and SVA MEIs (**Supplementary Figures 4** and **5**).

### Enhancing probability threshold for tackling imbalanced datasets

The AUPR metric shows the tradeoff between precision and recall across different probability cutoffs. The optimal cutoff depends on the signal-to-noise ratio (SNR) in real-world applications. The generally fewer true somatic MEIs found in tissue samples and their low tAFs typically lead to extremely imbalanced data, where the noise could overwhelm the signal. For detecting somatic L1 insertions in 200× WGS data of bulk brain tissues, for instance, the previous estimate of the true insertions is between 1 and 10, while the false insertions at the cutoff of two or more supporting reads are ∼1000^16^, leading to a SNR at around 1:100 to 1:1000. Germline L1 insertions in the Polaris dataset, for instance, have a SNR at 1:7 (**Supplementary Table 2**). The SNRs for *Alu* insertions are likely considerably lower, as there are more reference *Alu* sequences that can mix into PCR chimeras^53^. Similarly, SVA insertions may also have reduced SNRs because the activity of the retrotransposition process is likely lower, given that SVA is a non-autonomous mobile element. The significant data imbalance in somatic MEI detection can greatly affect the performance of the classifier model.

We simulated a challenging scenario where all true MEIs had low tAFs and only two supporting reads, with one representing image. To evaluate RetroNet’s performance in classifying individual images in imbalanced conditions, we sampled true and false images from the Polaris datasets into testing data with varying SNRs, ranging from 1:1, 1:10, 1:100, to 1:1000. RetroNet performs well when positive and negative samples are balanced; however, as the SNR decreases, the AUPR steadily declines. At extremely low SNRs, predicting positive samples can become challenging. For example, at a SNR of 1:1000, RetroNet shows AUPR values of 0.264 (95% CI: 0.246-0.283) for L1, 0.426 for *Alu* (95% CI: 0.399-0.453), and 0.422 for SVA (95% CI: 0.384-0.461) (**Figure 5a**). In real scenarios with very low somatic MEI content, it is crucial to implement stricter thresholds instead of the default cutoff of probability > 0.5 to regulate false positive levels. We evaluated the relationship between the precision/recall values and the SNRs (1:1 to 1:1000) at various probability cutoffs (0.5 to 0.99) in the simulated data (**Figure 5b)**. To ensure high precision for experimental validations, we increased the default probability cutoffs to 0.95 for L1 and 0.99 for *Alu* and SVA insertions. For the simulated L1 insertions at an SNR of 1:100, for instance, the prediction of RetroNet at a cutoff of 0.95 strikes a reasonable balance between recall (=0.712) and precision (=0.478). When applied to two previously validated somatic L1 insertions with a tAF of ∼1%^16^, RetroNet predicted both with high confidence above 0.95: L1_1 had a probability score of 0.997, and L1_2 was 0.998 (**Supplementary Figure 6**). The actual optimal cutoffs depend on the ME types, including the retrotransposition activities in somatic tissues that affect the level of the signals, as well as the likelihood of forming false ME junctions (e.g., PCR chimera) that affect the noise level.

**Figure 5.**
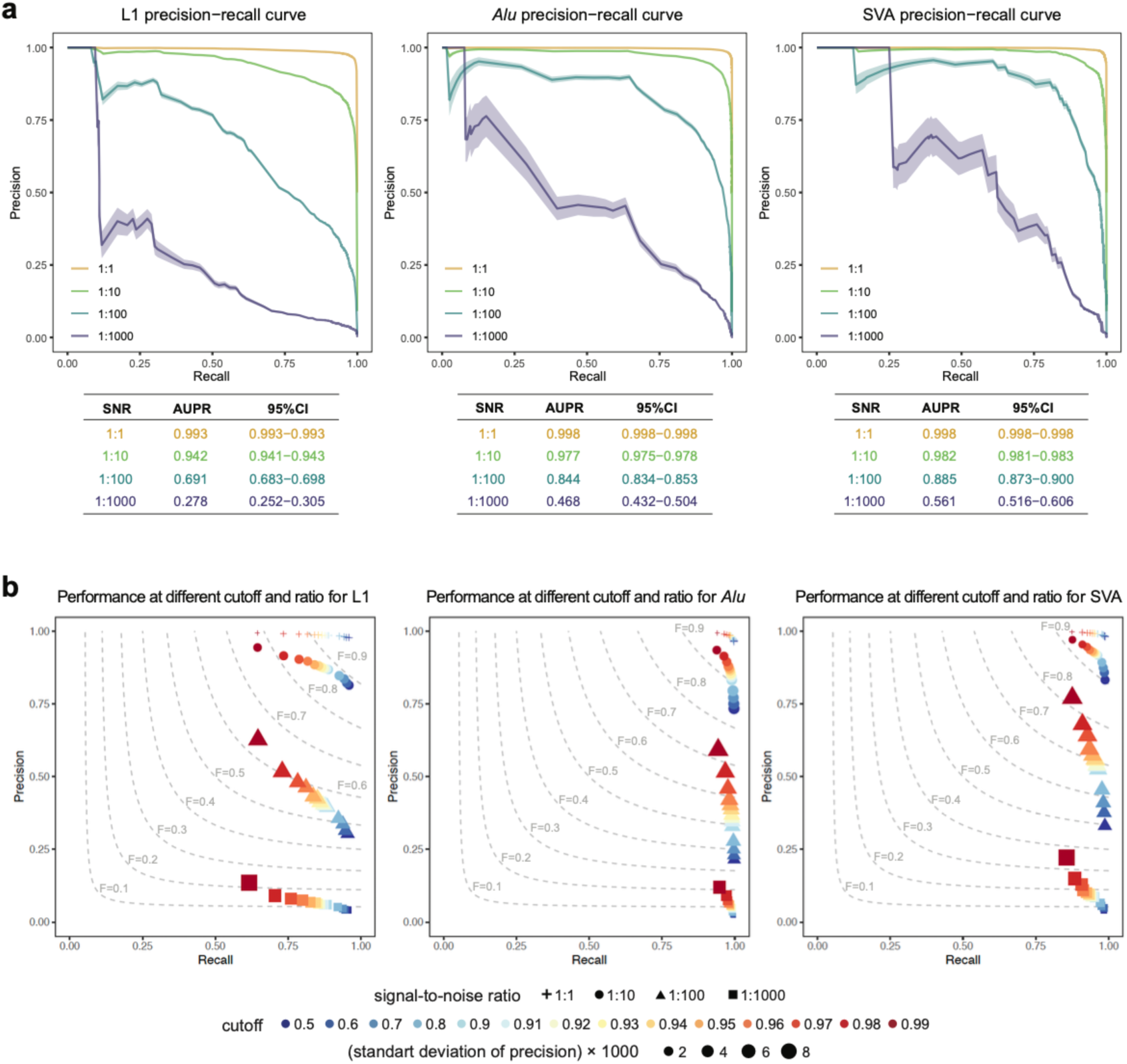
Benchmarking stringency cutoffs in simulated imbalanced datasets. (**a**) Precision-recall (PR) curves of RetroNet to identify resampled L1 insertions with various levels of noise, at signal-to-noise ratios (SNRs) of 1:1 (yellow), 1:10 (light green), 1:100 (dark green), and 1:1000 (purple). The solid line represents the average PR curve from 100 simulations, and the ribbon around the line indicates the 95% confidence interval. (**b**) The impact of more stringent probability cutoffs (from 0.5 to 0.99, blue to red) on the precision and recall of RetroNet, when applied to imbalanced datasets with an SNR of 1:1 (cross), 1:10 (circle), 1:100 (triangle), and 1:1000 (square). Despite the additional noise, RetroNet could still achieve high precision in extremely imbalanced datasets while managing reasonable levels of recall by choosing a higher stringency cutoff.

### Interpretation of the RetroNet neural network reveals known retrotransposition hallmarks

Class activation maps (CAMs)^54,55^ generated from true MEI images supported RetroNet correctly utilized both the positional features (e.g., the blue arrows) and sequence features of the supporting reads (e.g., the red pixels) for the prediction (**Supplementary Figure 7**). However, the CAM heatmaps are too coarse to precisely localize the relevant features, and thus, we carried out additional evaluations to understand the prediction behavior of RetroNet at a single-nucleotide resolution. We first investigated the supporting read positions in the L1 consensus sequence because frequent 5’-truncations and intact 3’-end are both hallmarks of true L1 retrotranspositions^56^. Among the 55,769 training images of true L1 insertions with supporting reads aligned across the 5’-junction, 23,000 images (∼40%) have intact 5’-end, and the rest supporting truncations typically between base 4,000 and 6,000 (**Figure 6a**). The distribution of the 5’-junction in true images aligns with a previous study of 20 individuals^51^ and differs significantly from that of false images. RetroNet predicted values (pre-normalized probabilities, called logits) for the true images positively correlated with the 5’-junction distribution of the true L1 insertions (beta = 0.305, *P* < 2e-16) but negatively correlated with the distribution in the false L1 insertions (beta = −0.105, *P* < 2e-16). For reads that could report L1 3’-junctions, true L1 insertions almost exclusively (∼95%) have the junctions at the L1 3’-poly(A) tail, suggesting the intact L1 3’-end (**Figure 6b**). RetroNet predicted values were also influenced by the 3’-junction locations, with a positive correlation to the distribution in true L1 insertions (beta = 0.392, *P* < 2e-16) and a weak negative correlation to the false-L1 distribution (beta = −0.027, *P* = 0.0423). 5’-truncations after L1 retrotranspositions are caused by premature termination of the reverse transcription, which is a shared mechanism between germline and somatic events^57^. Similarly, the preference for the intact L1 3’-end is led by the shared TPRT retrotransposition mechanism^5^. Thus, these results confirmed that RetroNet could learn the proper supporting read positions and implement them for predicting somatic MEIs.

**Figure 6.**
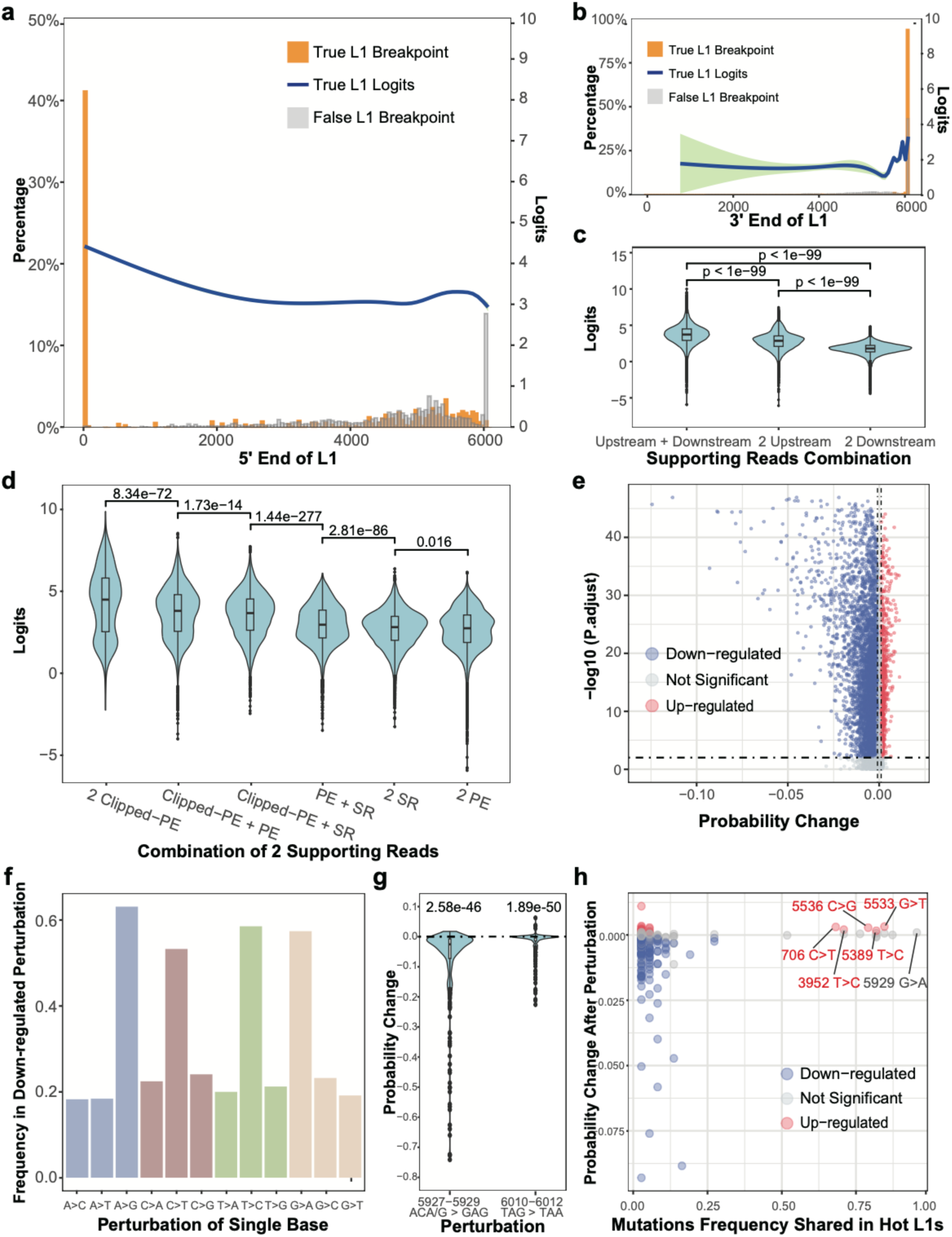
Interpretation of the RetroNet neural network reveals known L1 retrotransposition hallmarks. (**a**) RetroNet’s predicted value (blue) for L1 insertions is positively correlated with the 5’-end distribution in the true images (orange) but not the false images (grey). The predicted values are represented in pre-normalized probabilities (logits) and smoothed by the generalized additive model (GAM), showing the average in a blue line and the 95% confidence intervals in green. (**b**) RetroNet’s predicted value (blue) for L1 insertions is also positively correlated with the 3’-end distribution in the true images (orange) and not the false images (grey). Notably, since very few true L1 insertions had 3’-truncations, there is a wide 95% confidence interval for 3’-end locations near the L1 5’-end. (**c**) Boxplots of the RetroNet’s predicted values in relationship to the supporting read orientations, including upstream + downstream (left), two upstream (middle), and two downstream (right). (**d**) Boxplots of the RetroNet’s predicted values in relationship to the supporting read categories, including clipped paired-end reads (clipped PEs), paired-end reads (PEs), and split reads (SRs). (**e**) Impact of per-base perturbation of the L1Hs consensus sequence. For each base in L1Hs:3-6062, we sampled 300 training images with supporting read aligned to the base with the reference allele, permutated the supporting read allele into all three other nucleotides, and measured the changes to the RetroNet predicted probabilities. Changes to the probability is defined as significantly down-regulated (blue dot, probability change < −0.001 and adjusted *P* < 0.05), significantly up-regulated (red dot, probability change > 0.001 and adjusted *P* < 0.05), and not significant (grey dot). (**f**) Categories of perturbations with significant down-regulation in RetroNet, grouped by type of L1 consensus alleles: A (blue), C (dark brown), T (green), and G (light brown). (**g**) Impact of three-base perturbation on the L1Hs hallmark alleles, including the L1: 5927-5929 ACA/G to GAG and the L1: 6010-6012 TAG to TAA. Both perturbations had led to significant down-regulation in the RetroNet’s predicted probabilities. (**h**) Impact from permutating 328 alternative alleles identified from a group of 37 active L1 elements, in relationship to the allele frequency among the active L1s. Five alleles that are shared in over 19 of the 37 active L1s led to significantly higher prediction scores, including 706/C, 3952/C, 5389/T, 5533/T, and 5536/G. The boundaries of the boxplots indicate the first quartile (Q1, 25^th^ percentile) below and the third quartile (Q3, 75^th^ percentile) above, the black line within the box marks the median. Whiskers (vertical lines) extend above the box to the largest data point ≤ Q3 + 1.5 × IQR and below the box to the smallest data point ≥ Q1 − 1.5 × IQR, where IQR is the interquartile range (Q3 − Q1). P-values were calculated using the Wilcoxon test and adjusted using the Benjamini-Hochberg (BH) correction.

Regarding the combination of supporting reads, RetroNet prefers the orientation of one supporting read from the L1 5’-junction (called upstream) and the other from the 3’-junction (downstream) (**Figure 6c**). Candidate MEIs with both upstream and downstream supporting reads strongly support two novel junctions in the human genome — one from each end of the mobile element and thus are distinct from chimeric artifacts that only produce one novel junction^50^. The second preference is that both supporting reads come from the upstream, and lastly, the two supporting reads come from the downstream. The difference is likely due to poorer sequencing qualities near the poly(A) tails at the L1 3’ end, where Illumina sequencing is known to create errors generated by polymerase slippage on low-complexity sequences^58^. As for the types of supporting reads, RetroNet generally prefers images with at least one clipped PE, followed by the combination of PE and SR reads (**Figure 6d**). A clipped PE is preferred as it specifies not only the insertion junction, usually at the 5’-end where the sequencing quality is better, but also two segments of the L1 sequence alignments that should be arranged in proper positions (**Figure 2a**).

Active mobile elements that are capable of retrotransposition carry hallmark alleles such as ACA or ACG at L1:5927-5929 for the L1 Ta and pre-Ta subfamilies, as well as allele G at L1:6012^29,59,60^. To investigate the impact of specific sequence alleles on RetroNet, we adopted a perturbation-based test to simulate L1 mutations by permutating nucleotides of the supporting reads (i.e., the red pixels in the encoded images)^61^. Because local sequence alignment places gaps instead of mismatches near the endpoints, we omitted two nucleotides at either end of the L1Hs consensus sequence and chose bases 3 to 6062 for perturbations. For each base, we randomly selected 300 images with at least one supporting read carrying the L1Hs allele, which was then permutated to the three alternative alleles. In total, this test generated 18,180 probability change results (3 alternative DNA bases × 6060 sites), within which 5861 led to significantly down-regulated probabilities (probability change < −0.001 and adjusted p-value < 0.05) and 847 led to up-regulated probabilities (probability change > 0.001 and adjusted p-value < 0.05) (**Figure 6e**). Transition mutations, including A>G, T>C, C>T, and G>A, were common among the down-regulating perturbations (**Figure 6f**), reflecting the major difference between the active L1 sequence and the noise generated from old, inactive L1 elements that have accumulated too many mutations – a majority of them are transition mutations as they are more common than transversion mutations^62^.

The perturbation test also demonstrated that RetroNet could recognize canonical alleles of active L1.

As expected, probability changes of three-base perturbations from L1:5927-5927 ACA/G to GAG and from L1:6010-6012 TAG to TAA were significantly down-regulated (*P* =2.58e-46 and 1.89e-50, respectively) (**Figure 6g**). For single-base perturbations, we investigated 328 alternative alleles carried by 37 active L1 elements in the human genome^63^. Overall, the alleles shared among the majority of the active L1s (>50%) are more likely to cause a positive change in the RetroNet prediction than the less common alleles (*P* = 9.96e-7) (**Figure 6h)**. Five mutations that are shared in over 19 of the 37 active L1s led to significantly higher prediction scores. Among these, 706/C, 5389/T, 5533/T, and 5536/G were also recognized canonical sites of active L1s^2^. The fifth allele, 3952/C, was found in 26 of the 37 active L1 elements, including several that are considered to be highly active in *in vitro* assays^63^. Notably, the single nucleotide change of 5929 G>A did not produce significant changes since pre-Ta L1 (5927-5929:ACG) is still capable of retrotransposition, and RetroNet relies on all three nucleotides at L1:5927-5929 for the classification (**Figure 6g)**. In comparison, the majority of the rest, less common active L1 alleles generally led to down-regulation (∼47%) or insignificant (∼49%) changes in the predicted probabilities (**Figure 6h**), suggesting they were not the focus of the RetroNet’s learning.

### Detecting somatic MEIs in cancer cell line HG008

To demonstrate the efficacy of RetroNet in real bulk sequencing datasets, we characterized somatic MEIs in a pancreatic ductal adenocarcinoma tumor cell line (HG008-T) with matched normal duodenal tissue (HG008-N-D) as control, using public short and long read sequencing data from the Genome in a Bottle Consortium (**Supplementary Table 1)**^40–42^. For benchmarking, we identified a total of 19 true tumor somatic L1 insertions with an estimated tAF between 1.6% and 100%--and no *Alu* or SVA insertions from two PacBio long-read sequencing^64^ datasets with a combined average sequencing depth of 212×, using PALMER^65^, xTea_long (long-read module)^51^, and an in-house tool (**Supplementary Data 7** and **Supplementary Note 2**). We then applied RetroNet, RetroSom^16^, xTea (short-read module)^51^, and TraFiC-mem^18,19^ to two independent Illumina WGS datasets, ILMN-PCR-free-2 (I2) and ILMN-PCR-free-3 (I3), with an average depth of 195× and 161× for HG008-T, respectively. After accounting for ploidy, the average depth per haploid genome (termed number of reads per chromosome copy, or NRPCC^66^) was 49× for I2 and 40× for I3. A third dataset, ILMN-PCR-free-1, was excluded as it sequenced a different passage of HG008-T cells and used primary normal pancreatic, instead of the duodenal tissues as control.

With the requirement for at least two supporting reads and in non-repetitive genomic regions, 12 and 14 of the 19 true L1 insertions were detectable in datasets I2 and I3, respectively. Within dataset I2, RetroNet identified a total of 13 somatic L1 insertions with P > 0.95, of which 10 were true L1 retrotranspositions (recall=0.833, precision=0.769). The tAF interquartile range of the identified L1 insertions is between 5.9% and 92.3% (minimum=1.79%) (**Figure 7a** and **Supplementary Table 5**). The two false negatives include one with a 3’ transduction and 5’ inversion and therefore lacks an intact 3’-end, and a short L1 insertion near the 3’-poly(A) that caused many sequencing errors^58^. The three false positives were low-tAF calls with proper sequence and positional features, including the L1Hs hallmark alleles and both upstream and downstream supporting reads, and, therefore, were possible rare somatic events missed in the long-read sequencing. Comparatively, RetroSom’s accuracy was significantly weaker without manual inspections: recall was 0.667, and precision was 0.381 – the majority of errors were PCR duplicates or incoherent read positions (**Supplementary Figure 8**). Furthermore, xTea identified six true L1 insertions (recall=0.500, precision=0.857, tAF interquartile range: 76.9%-98.4%), while TraFiC-mem identified five true L1 insertions (recall=0.416, precision=1, tAF interquartile range: 72.8%-93.5%). Notably, xTea and TraFiC-mem generally overlooked low-tAF MEIs but could identify those that had only transduction and no mobile element sequences, known as “orphan transductions” ^67,68^ (**Figure 7a**). They represent a class of noncanonical MEIs not benchmarked here, as the accurate detection of low-tAF orphan transductions remains unresolved in long-read sequencing^51,65^. Finally, while there were no true *Alu* or SVA insertions, RetroNet misclassified 9 somatic *Alu* insertions, a significantly lower number than RetroSom’s 96 and 2 somatic SVA insertions. Contrarily, xTea misclassified 1 *Alu* insertion, while TraFiC-mem did not misclassify any *Alu* or SVA insertions (**Supplementary Table 5**).

**Figure 7.**
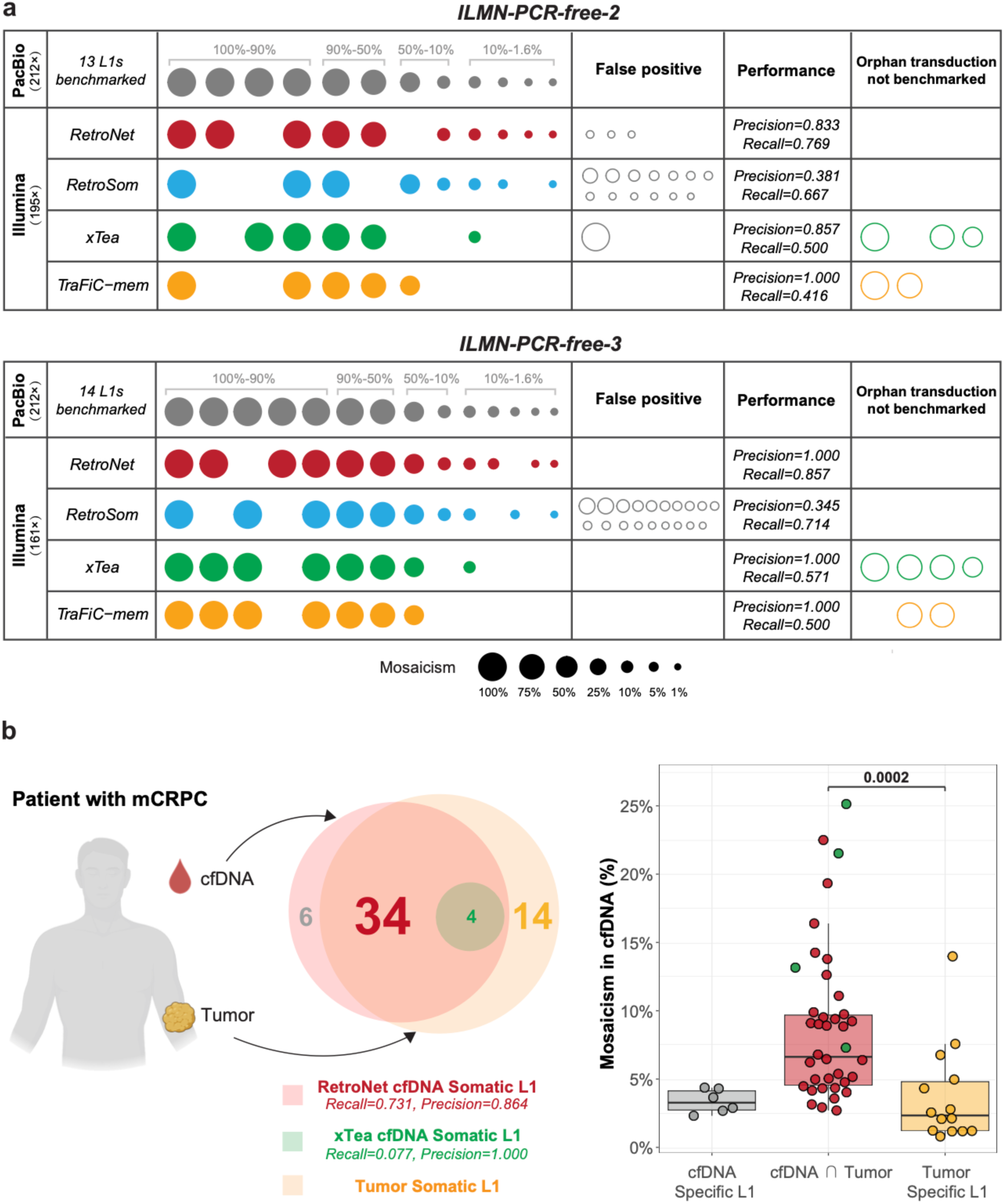
Comparison of RetroNet and other algorithms in detecting somatic L1 insertions from cancer cell line and cfDNA. (**a**) Detection of HG008-T tumor somatic L1 Insertions based on Illumina WGS data. This figure illustrates the detection of tumor somatic L1 insertions in the Illumina WGS datasets of HG008-T, ILMN-PCR-free-2 (I2, upper panel) and ILMN-PCR-free-3 (I3, lower panel) using four different algorithms. The grey ellipse represents true somatic L1 insertions benchmarked by PacBio HiFi sequencing, with at least two supporting reads in the Illumina WGS datasets as the minimum detection requirement. The colored dots indicate true positives detected by each algorithm (RetroNet: red, RetroSom: blue, xTea: green, TraFiC-mem: yellow). The grey open dots indicate false positives. The colored open dots represent orphan insertions detected by the algorithms but not included in the benchmarking list, and thus were excluded when calculating recall. (**b**) Detection of tumor-derived somatic L1 insertions in cell-free DNA from patient DTB-205 with treatment-resistant prostate cancer. The Venn diagram compares somatic L1 insertions detected in cfDNA by RetroNet (red circle) or xTea short (green circle) with those present in the tumor (yellow circle). RetroNet identified 44 L1s in cfDNA, six unique to cfDNA, and 38 that were also found in the tumor, of which four were co-detected by xTea short. Fourteen tumor specific L1s were absent from cfDNA. On this cohort, RetroNet achieved a recall = 0.731 and precision = 0.864, whereas xTea short achieved a recall = 0.077 and precision = 1.000. The right panel shows mosaicism in cfDNA for three categories: cfDNA specific insertions (grey), tumor-derived L1s detected in cfDNA by RetroNet alone (red) or co-detected by xTea (green), and tumor specific L1s (yellow). Tumor-derived L1s detected in cfDNA exhibit significantly higher mosaicism than tumor specific insertions (Wilcoxon test, *P* = 0.0002). The boundaries of the boxplots represent the first quartile (Q1, 25^th^ percentile) below and the third quartile (Q3, 75^th^ percentile) above, with the black line inside the box marking the median. The whiskers (vertical lines) extend above the box to the largest data point ≤ Q3 + 1.5 * IQR and below the box to the smallest data point ≥ Q1 - 1.5 * IQR, where IQR is the interquartile range (Q3 - Q1).

Dataset I3, compared to I2, had a lower sequencing depth (161× vs. 195×) and a higher percentage of properly paired reads (98.3% vs. 96.8%), implying it contained fewer chimeric artifacts and, therefore, a lower level of noise. For L1 analysis, RetroNet had a similar recall (=0.857) with a higher precision (=1) than I2; both outperformed RetroSom (recall=0.714, precision=0.345) (**Figure 7a** and **Supplementary Table 6**). The two false negative L1 insertions were the same L1 with 3’ transduction and 5’ inversion as found in I2, as well as a low-tAF somatic L1 insertion with only two short split-read supporting reads in I3. Both xTea and TraFiC-mem maintained a high precision score of 1, with recalls of 0.571 and 0.500, respectively (**Figure 7a**). The detected MEIs exhibited higher interquartile ranges in tAF compared to RetroNet (xTea: 64.7%-100%, TraFiC-mem: 80.9%-100% vs. RetroNet: 9.4%-93.7%). For *Alu* and SVA insertions, RetroNet and TraFiC-mem reported none, while RetroSom misclassified 6 false *Alu* insertions and xTea misclassified 6 false *Alu* insertions and 1 SVA insertion (**Supplementary Table 6)**. The performance difference between I2 and I3, as well as among various ME types, can be attributed to the differing SNRs. Specifically, the SNR for L1 insertions was 1:5475 in I2 and 1:532 in I3. For *Alu*, it was below 1:77784 in I2 and 1:731 in I3, while for SVA, it was under 1:1941 in I2 compared to 1:945 in I3.

By combining RetroNet’s filtering system with additional filters—such as the inability to resolve repetitive regions using short reads, the lack of sufficient supporting reads for low-tAF mutations, and the presence of false supports in control tissues—the average recall rate for detecting somatic L1 elements in HG008-T is 0.579 (11 out of 19). This rate is slightly lower than the previously reported recall rates for germline mobile element insertions (MEIs), which were 0.68 for short-read sequencing at 300× and 0.93 for long-read sequencing^51^. For low-tAF MEIs, RetroNet accurately detected a total of six somatic L1 insertions with fewer than five supporting reads, including four with only two supporting reads. Out of the nine true somatic L1 insertions with a tAF between 1.6% and 10%, the average recall of RetroNet recalls was 0.5 between I2 (N=4) and I3 (N=5), with few false positives (N=3 in I2; N=0 in I3). The limited recall is primarily due to insufficient supporting reads (<2) in the short-read sequencing data (N=4 in I2; N=3 in I3) (**Supplementary Data 7**). Unlike other MEI detection algorithms, which often require a substantial number of supporting reads, such as the TraFiC-mem algorithm^18,19^ that found no somatic insertions in this tAF range or xTea that found only one, our analysis using RetroNet successfully improved the detection of low mosaicism somatic MEIs.

### Benchmarking somatic MEIs in cell-free DNA and matching tumor tissues

We further assessed the use of RetroNet for short-read sequencing of fragmented cell-free DNA (cfDNA), where long-read sequencing is not applicable. The dataset includes time-matched cfDNA (average depth=185×), metastatic tumor (98×), and white blood cells (41×), from 13 metastatic castration-resistant prostate cancer (mCRPC) patients^43^. The cfDNA from patient DTB-205 has an estimated cancer fraction of 28%, with a 100% contribution from the biopsied metastatic tumor. This allows us to use the matching metastatic tissue as an independent validation to confirm the somatic MEIs identified in the cfDNA. The contributions of metastatic tumor biopsies to cfDNA for other patients ranged from 1% to 60%, and thus, were excluded from our benchmarking analysis.

Due to the significant fragmentation of cfDNA (interquartile range: 148-178 bp), we employed a more stringent cutoff of ≥3 supporting reads with ≥2 high-confidence read pairs in RetroNet. We compiled a list of 52 *true* somatic L1 insertions in the DTB-205 tumor tissue. This list includes 38 insertions identified by xTea in the tumor sequencing and an additional 14 insertions that had fewer supporting reads than the xTea threshold but were identified through manually reviewed matching RetroNet calls in cfDNA (**Supplementary Figure 9** and **Supplementary Table 7**). In addition, xTea identified no *Alu* insertions and one somatic SVA insertion in the DTB-205 tumor, indicating a low level of mosaicism from these elements. Within the cfDNA, RetroNet identified a total of 44 somatic L1 insertions, with an estimated recall of 0.731 and a precision of 0.864 (**Figure 7b**). The tAF interquartile range for the correctly identified L1 insertions was between 4.8% and 10.8%, with a minimum value of 2.9% and a maximum value of 25.1%. This range is consistent with the estimated cancer fraction of 28% in the cfDNA. The cfDNA-specific L1 insertions (N=6) may include false positives or true low-tAF mutations in the tumor that were not detected by sequencing. The tumor-specific insertions (N=14) exhibited a significantly lower tAF in cfDNA (interquartile range: 1.2%-4.8%) compared to the L1 insertions detected in cfDNA (*P* = 0.0002), including 7 cases with insufficient supporting reads (**Figure 7b** and **Supplementary Table 7)**. Attempting the same task by xTea on the cfDNA sequencing data led to the discovery of four somatic L1 in cfDNA; all were found in the tumor (recall=0.077, precision=1, tAF interquartile range: 11.7%-22.4%). Overall, the significantly improved recall for detecting somatic L1 insertions in cfDNA demonstrates RetroNet’s effectiveness in analyzing highly degraded DNA and its potential applications in scenarios where long-read sequencing is impractical.

## Discussion

Somatic mobile element insertions have increasingly been recognized for their involvement in brain development and tumorigenesis. The challenges of low tissue allele frequency and sequence repetitiveness complicate their detection, necessitating innovative strategies for precise identification. To address this, RetroNet integrates both sequence and positional features of candidate supporting reads using a deep neural network. Compared to the previous model, RetroSom, which only analyzes sequence features, RetroNet demonstrates higher sensitivity, enabling the detection of MEIs at low frequencies with improved precision, thereby reducing the need for manual validation of putative MEIs. This advancement marks a significant step towards uncovering the elusive roles of somatic MEIs in neurological diseases and cancer development.

Direct encoding of DNA sequencing data with the one-hot method has been widely used in deep learning-based models for genomics^31^. Typically, these methods involve converting the sequencing reads into a 4×M matrix, where the four rows represent the four nucleotides (i.e., A, C, T, and G), and M is the read length. Applying this encoding method to build a classifier for somatic MEIs, however, would have caused biased representations. All candidate MEI supporting reads contain flanking sequences that are largely different between the germline MEIs used for training and the targeted somatic MEIs, except for the short (∼6 bp) endonuclease cleavage sites. The insertion loci of the polymorphic germline MEIs have also been subject to genetic drift or natural selection in human evolution, which were generally absent or selected at a different level, cellular instead of species, for somatic MEIs. Furthermore, learning patterns from the MEI flanking sequences would also lead to overestimating the model’s performance, as a portion of the MEIs (e.g., those with high population frequencies) are shared across the training and the benchmarking datasets with identical flanking sequences. Finally, the conventional one-hot encoding method would also be a poor representation to capture the crucial aspects of the relative positions and orientations of the selected supporting reads, as well as the alignments with the ME consensus sequences. Hence, we leveraged insights from the prior knowledge into an unbiased and comprehensive encoding scheme, including only the relevant sequence and positional features for the sequence-to-image conversion.

Various strategies, including targeted or whole-genome sequencing and single-cell or bulk-tissue sequencing, have been applied to detect somatic MEIs. Of these methods, deep coverage WGS on bulk tissues or mixed cells, when analyzed properly, serves as a good balance between the sequencing costs, noise, and the yield of detectable somatic mutations. Targeted sequencing that captures ME junctions before sequencing is cost-effective but only detects somatic MEIs and typically requires additional PCR amplifications that could bring significant levels of chimeric artifacts. Whole genome sequencing, in comparison, enables the simultaneous identification of a broad spectrum of genetic variations, including single nucleotide variants, MEIs, and other structural variations. Furthermore, single-cell WGS allows for detecting somatic mutations with extremely low mosaicism but also requires a significant level of DNA amplification. Enzymatic whole genome amplification approaches could lead to biased sequencing depth or even artifactual chimeras, compromising the result’s accuracy. Alternatively, clonal expansion from fetal or reprogrammed cells circumvents enzymatic DNA amplification biases but is constrained by the type of suitable tissues. Given the same sequencing throughput, the sensitivity of bulk WGS for detecting low-mosaicism mutations is significantly higher than single-cell WGS. Assuming each cell is sequenced at an average depth of 33×, the cost of 200× bulk WGS equates to that of 6 single-cell WGS, and 400× bulk WGS corresponds to 12 single-cell WGS. For somatic MEIs with a tAF of 1%, the chances of sampling them in at least one of the 6 or 12 cells are 0.059 or 0.114, respectively. When using bulk WGS and >=2 supporting reads, in comparison, the theoretical maximum recall is 0.787 for 200× and 0.979 for 400× depth (**Supplementary Note 3** and **Supplementary Table 8**). The actual recall is lower after filtering for suitable supporting reads and removing false positives, RetroNet achieved the recall of 0.238 with 200× and 0.523 with 400× sequencing in a simulated benchmarking dataset, both of which are still greater than the equivalent single-cell approaches. Overall, while single-cell approaches can characterize private or extremely low-mosaicism mutations, bulk tissue-based methods remain a practical and effective strategy for MEIs with a tAF of approximately 1% or higher. Somatic mutations of even lower frequencies could, theoretically, require an unrealistic level of sequencing depth for detection using bulk tissue sequencing. In applications such as genetic testing for disease-causing mutations, however, somatic MEIs in 0.1% or lower percentage of cells likely have limited functional impacts.

The accuracy of detecting low-mosaicism MEIs using bulk sequencing depends on several intercorrelated factors, including sequencing depth, tAF of the MEI, the threshold for the number of supporting reads, and the ratio between the numbers of true and false insertions (i.e., the SNR). Among these, high sequencing depth is generally considered to be necessary for detecting MEIs at low tAF, at the cost of additional sequencing cost and specimen requirement. However, the benefit of increasing sequencing depth still has limitations. Somatic MEIs of extremely low mosaicism levels likely arise from retrotransposition events in a relatively later developmental stage, leading to narrow anatomical distributions. Sampling additional tissues for higher sequencing depth, therefore, will not guarantee the increase in the number of supporting reads if the additional tissues do not contain the somatic MEIs. Instead, the additional sequencing errors could lead to a worse signal-to-noise ratio. It is, therefore, essential to develop a tool that can reliably detect low-mosaicism MEIs without relying solely on the high sequencing depth, and thus, the tool needs to identify MEIs with as few supporting reads as possible.

Lowering the threshold of supporting read number for calling somatic MEIs, however, could lead to much higher levels of false positive insertions from sequencing or mapping artifacts that, while occurring at low probabilities, could accumulate into a large number of events. We adapted RetroNet to fit a new model for classifying MEIs using a single supporting read instead of the default two reads and demonstrated that the two-read strategy showed significantly higher precision (2 reads: 0.970; 1 read: 0.829) in classifying candidate supporting reads in the Polaris dataset (*P* = 2.46e-3) (**Supplementary Note 4** and **Supplementary Figure 10)**. Notably, in real-world scenarios, the single-read method would likely perform much worse due to the expanded imbalance between the signals and noise. A previous analysis estimated that by lowering the threshold from two supporting reads to one read and no additional filtering, the number of false positive L1 MEIs increased by approximately 80-fold^16^. When using 200× sequencing on bulk brain tissues for detecting somatic L1 insertions, for instance, we typically expect 1∼10 true insertions and 100∼1000 false positives (SNR ∼ 1:100). Using the single-read method, however, the estimated SNR is 1:10000. As demonstrated in our data imbalance simulations, the weaker SNR suggests the one supporting read strategy on bulk, short-read WGS will likely be unsuitable for real-world applications.

The RetroNet pipeline still comes with several limitations. The primary one lies in the inherent bias of transfer learning from germline MEIs that could have accumulated more mutations and tend to have shorter poly(A) tails than somatic MEIs. We attempted to mitigate this bias by choosing the polymorphic, evolutionarily young MEIs for training, and by coding the known difference between *bona fide* retrotransposition hallmarks and false MEIs in the image-based encoding. We also opt-out known features that would differ between germline and somatic MEIs, such as the flanking sequences of the MEI junction. As the number of experimentally validated somatic MEIs grows, the RetroNet framework could easily be fine-tuned using somatic MEIs only. Additional significant challenges arise from the short-read sequencing technology, including (1) complex retrotranspositions such as 5’-inversions and long or orphan transductions, (2) DNA structural variations that involve active mobile elements at the junctions and resemble retrotransposition, and (3) making identifications within repetitive genomic sequences. For instance, given RetroNet’s requirement for both supporting reads to align to the same ME strand to filter noise, it would miss low-mosaicism 5’-inversion events with <2 supporting reads covering either end (**Supplementary Figure 11a**). In addition, when transduction occurs in L1 or SVA insertions, standard short-read sequencing with a span of ∼800bp may be insufficient to capture the ME sequences junction (**Supplementary Figure 11b**).

Long-read technologies, such as PacBio HiFi sequencing^64^, allow for capturing the complete insertion sequences, which facilitates the direct identification of complex retrotranspositions, differentiation from structural variations, and improved detection in repetitive sequences^65^. However, current long-read sequencing technologies still have drawbacks, including higher sequencing costs and a notable rate of sequencing errors. These challenges are particularly significant when detecting low-tAF somatic mutations, which typically require substantial sequencing depth (e.g., >200×) and precise characterization of active mobile element hallmark alleles. Moreover, long-read sequencing is unsuitable for fragmented DNA such as ancient DNA (40-500 bp)^69^ or cell-free DNA (40-166 bp) ^70^. As demonstrated in our benchmarking of the cfDNA from a tumor patient, short-read sequencing can effectively complement long-read sequencing technologies and is capable of identifying somatic MEIs as unique cancer biomarkers. Despite the limitations associated with transfer learning and sequencing technology, RetroNet represents a meaningful advancement over previous methods. It can be effectively applied to the abundant Illumina sequencing data that is currently available or being generated for a variety of human traits and diseases. Finally, the RetroNet framework can also serve as pre-trained models for other emerging high-quality short-read sequencing technologies such as Element AVITI, Ultima UG100, and PacBio Onso^71^.

## Acknowledgments

We thank Jin Xu from Sun Yet-sen University, Chao Jiang from Zhejiang University, and members of the Somatic Mosaicism across Human Tissues (SMaHT) Network for their constructive comments on the development of the methodology. We thank Joel E. Kleinman, Thomas H. Hyde and Daniel R. Weinberger from Liber Institute for Brain Development for providing the BSMN common brain tissue, and Liana Fasching from Yale University for extracting the BSMN common brain DNA. We thank Alexander Wyatt from the University of British Columbia for sharing the Illumina sequencing datasets of the mCRPC patients. This work utilized computing resources provided by the Stanford Genetics Bioinformatics Service Center and the City University of Hong Kong high-performance computing (HPC) resources.

## Code Availability

The code for the RetroNet is available at https://github.com/Czhuofu/RetroNet.

## Author contributions

**Miaomiao Tan**: Methodology, Benchmarking, Writing - Original Draft, Visualization. **Zhinan Lin**: Methodology, Software, Neural network interpretation, Writing - Original Draft. **Zhuofu Chen**: Software, Data Curation. **Haonan Zhou**: Methodology and software development, Analysis of the cell-free DNA datasets. **Junseok Park:** Software development and performance benchmarking. **Ziting He**: Methodology and software development. **Eunjung A. Lee:** Methodology, Benchmarking, and Writing. **Zhipeng Gao**: Methodology, Writing - Review & Editing. **Xiaowei Zhu**: Conceptualization, Resources, Writing - Review & Editing, Data Curation, Funding acquisition. All authors read and approved the final manuscript.

## Supplementary information

### Methods

#### Datasets for training and benchmarking

##### High-depth 1000 Genomes Project

The expended 1000 Genomes Project comprises high sequencing depth WGS data of 3,202 individuals, including 602 parent-offspring trios. Each individual’s sample was sequenced to achieve a target depth of 30× genome coverage, utilizing PCR-free technology on Illumina NovaSeq 6000 platform (ENA accession: PRJEB55077)^35^. Notably, we excluded 53 trios in model training as they overlap with the benchmarking datasets. These include 49 trios sequenced in the Illumina Polaris dataset, and 4 individuals (NA19240, HG00733, HG00514, and NA12877) used in the genome mixing dataset. As a result, 549 family trios from the 1000 Genomes Project were used for model training.

##### Illumina Polaris Project

The Illumina Polaris Project encompasses WGS data from 49 parent-child trios, with the children forming the Kids Cohort (ENA accession: PRJEB25009) and the parents originating from the Diversity Cohort (ENA accession: PRJEB20654). All individuals were selected from the 1000 Genomes Project to represent a wide range of population diversity. Each individual’s sample was sequenced using PCR-free libraries, achieving an average depth of 30× genome coverage on the Illumina HiSeqX platform^39^.

##### Simulated Genome Mixing Dataset

DNA from six different human genomes was mixed with the HapMap sample NA12878 (whose lineage MEIs are generally established) in precise proportions ranging from 0.2% to 25%, including (1) A1S heart at 0.04%, (2) NA19240 at 0.2%, (3) HG00733 at 1%, (4) HG00514 at 1%, (5) Brain somatic mosaicism network (BSMN) common brain at 5%, and (6) NA12877 at 25%. The mixed genomic DNA and the backbone NA12878 genomic DNA were sequenced with Illumina sequencing at a coverage of 200× and resampled to 50×, 100× and 400× (NIMH Data Archive Collection 2458, Experiment 1072)^16^.

##### Simulated Imbalanced Datasets with different SNRs

We simulated datasets of somatic MEIs at various noise levels by sampling true and false images from the Polaris datasets with ratios of 1:1, 1:10, 1:100, and 1:1000, each repeated 100 times. <u>Cancer cell line HG008-T.</u> We processed four paired tumor-control sequencing datasets, including two Illumina sequencing datasets (ILMN-PCR-free-2 and ILMN-PCR-free-3) and two PacBio HiFi sequencing datasets (PB-HiFi-1 and PB-HiFi-2) from the Genome in a Bottle Consortium (https://42basepairs.com/browse/s3/giab/data_somatic/HG008/Liss_lab)^40–42^. The tumor genomic DNA was extracted from a pancreatic ductal adenocarcinoma tumor cell line (HG008-T) at passage number 23.

##### Cell-free DNA of mCRPC patient DTB-205

We analyzed sequencing data from time-matched samples, including metastatic tissue, cell-free DNA (cfDNA), and white blood cells, obtained from a metastatic castration-resistant prostate cancer (mCRPC) patient DTB-205. The dataset is publicly available through the European Genome-Phenome Archive (EGA accession: EGAS00001005783)^43^.

#### Labeling candidate MEIs in parent-offspring trios

Following the methods described by Zhu et al.^16^, candidate supporting reads were extracted using a modified RetroSeq pipeline^25^. To define a clean set of MEIs, we further removed alignments in non-primary chromosome assemblies, or within repetitive sequences that are prone to alignment errors, including telomere, centromere, and young, fixed genomic mobile elements with <10% sequence divergence^72^. Specifically for characterizing the short *Alu* insertions that are ∼300 bp in length, since the supporting read alignment positions are always close and cannot help to distinguish signal from noise, we further filtered out reference *Alu* elements of lower than 20% divergence to avoid mistaking them as somatic insertions. For human reference genome version hg38, the excluded genome regions are 11.89% for L1 and SVA insertions and 20.63% for *Alu* insertions.

We labeled candidate MEIs in the 549 offsprings of the trios from the 1000 genome project and the 49 offsprings from the Polaris project as *true* or *false* insertions based on the inheritance pattern. The true MEIs are those found in the offspring and their parent, while false MEIs are those absent in the parent. In addition, true MEIs satisfy: (1) >20 total supporting reads, including >1 split read, (2) have supporting reads for both the upstream junction and the downstream junction, with the downstream junction reads at a proportion between 10% and 90%, (3) have been previously annotated by MELT in the 1000 genome project phase III MEI list (not DNA structural variations)^73,74^, and (4) absent in one parent in one parent (not alignment errors or evolutionally old MEIs with high population frequencies). False MEIs also have >1 total supporting reads and <6 paired-end reads, or when there are ≥ 6 paired-end reads, the downstream junction reads are below 10% or above 90% (not true *de novo* MEIs).

#### The extra syntax used for encoding candidate MEI-supporting reads into images

Several additional encoding syntaxes were implemented to ensure a consistent sequence-to-image conversion. First, a maximum of five images of randomly selected pairs of supporting reads for each training MEI were generated to mitigate the risk of overrepresenting those with more supporting reads (e.g., homozygous over heterozygous MEIs). Second, the one-hot encoding of the ME consensus has its 5’-end on the left and the 3’-end on the right, consistent with the MEIs on the plus strand but opposite to the minus strand insertions. To ensure a consistent representation, we rotated the encoding of the minus-strand insertions, including the mappability track and the flank sequence arrows, by 180 degrees. Third, for any pairs of supporting reads, read 1 is selected as the one with the flanking read more on the left, and read 2 as the one with the flanking site more to the right on the image, after adjusting to the insertion’s strandness. Fourth, we also denoted the possible unmappable segments in the supporting reads (**Supplementary Figure 2**). The clipped segment in a split read or clipped-PE read is represented by a proportional black line, with the segment mappable to ME consensus denoted as a red line below. Unmappable segments in the read end that map to the ME consensus in PE supporting reads are displayed as a proportional black line above the ME sequence (**Supplementary Figure 2a**). Other insertion gaps in the read-ME alignment are marked with blue pixels above the ME sequence at the corresponding insertion loci (**Supplementary Figure 2b**). MEI with inversions, which typically occur in L1 insertions, may have one L1 segment in clipped-PE reads mapped to the L1 opposite strand, which is denoted by purple instead of red pixels in the ME consensus track (**Supplementary Figure 2c**). Finally, we denote any unmappable segments supporting *Alu* insertion carrying 5’- or 3’-transduction as green lines. Since *bona fide Alu* retrotranspositions do not carry additional sequences from the source elements, these green lines can be learned as signatures of false junctions (**Supplementary Figure 2d**).

The image encoding of the *Alu* and SVA insertion is similar to L1, except there are several active *Alu* and SVA subfamilies in the human genome (**Supplementary Figure 2e**). For *Alu* insertions, we aligned the supporting reads to the consensus sequences of four active *Alu* subfamilies, *Alu*Ya, *Alu*Yb, *Alu*Yc, and *Alu*Yk^57^, and employed four tracks to portray the sequence alignments to each *Alu* subfamily. The encoded image for *Alu* comprises 21 tracks, where track 1 represents mappability, and tracks 2, 7, 12, 17 denote the relative positions and orientations if the corresponding read end is aligned in the flanking sequences. Tracks 3-6, 8-11, 13-16, and 18-21 represent the alignments of the four read ends to the four *Alu* consensus sequences. Similarly, we used SVA_E and SVA_F consensus sequences for SVA insertions, resulting in images of 13 tracks.

#### Neural Network training and validation

The images produced in the process above served as direct inputs for the CNN architectures ResNet-18 and GoogLeNet. To train the neural networks, we randomly divided the labeled images from the 1000 genome project datasets into the training and validation datasets at a 9:1 ratio. We then trained binary classifiers based on the ResNet-18 and GoogLeNet architectures (**Supplementary Table 3**), using the cross entropy between predicted probabilities and true class labels as the loss function. To select an optimal learning rate, we chose an initial value of 0.001 and used the reduce-on-plateau method to dynamically adjust the learning rate. Specifically, if the training losses remain stable for 5 epochs, the learning rate undergoes exponential decay by a factor of 0.1. The final output layer of the CNN model is a two-class Softmax layer. Each model was trained for 30 epochs by which the training loss converged, and we chose the epoch exhibiting the lowest validation loss as the finalized model.

In the third network architecture, ViT, the input images were initially divided into nine horizontal segments, each representing the mappability, or the position and ME alignment of one of the four read ends. Each partition was resized to an identical shape by padding blank spaces when necessary. The remaining model training process for the ViT network was the same as that used for ResNet-18 and GoogLeNet. All these processes were executed in the PyTorch environment (v1.13.1).

#### Benchmarking RetroNet in independent datasets

We benchmarked RetroNet in three datasets, including (1) germline MEIs generated from 49 parent-offspring trios in the Illumina Polaris cohort^49^, labeled in the same way as the training data; (2) simulated somatic MEIs with tAF between 0.2% and 25% in a genome mixing dataset in which the true MEIs were established previously^16^; and (3) simulated somatic MEIs in datasets in a range of signal-to-noise image ratios, from 1:1, 1:10, 1:100, to 1:1000. The model performance was evaluated using the following metrics: precision = true positive/(true positive + false positive), recall = true positive/(true positive + false negative), F1 score = 2 × precision × recall / (precision + recall), and area under the precision-recall curve (AUPR). For dataset 1 with 49 individuals and dataset 3 with 100 times of resampling, we also reported the 95% confidence intervals of the AUPR metric.

#### Neural network interpretation

We generated the class activation maps using the Grad-CAM algorithm^67^. The association between RetroNet’s predicted values (logit) and the alignment positions was evaluated in a generalized linear model: 𝐿𝑜𝑔𝑖𝑡(𝑥) = 𝑤_0_ + 𝑤_1_𝐿1_𝑡𝑟𝑢𝑒_(𝑥) + 𝑤_2_𝐿1_𝑓𝑎𝑙𝑠𝑒_(𝑥), in which 𝑥 was the 5’- or 3’-junction position in the L1 sequence and 𝐿1_𝑡𝑟𝑢𝑒_ and 𝐿1_𝑓𝑎𝑙𝑠𝑒_ were the density values in histograms (bin=50bp) of the true or false L1 insertions, respectively. For each base of L1Hs:3-6062 in the perturbation test to assess mutation impacts, we randomly sampled 300 training images with one or both reads aligned across the targeted base, and then permuted the aligned base (i.e., the red pixels) to all three alternative bases. The three base permutations, including L1Hs: 5927-5929 ACA/G to GAG and L1Hs: 6010-6012 TAG to TAA, were carried out in the same process as the single-base permutations.

#### Detecting somatic MEIs in the PacBio HiFi sequencing

We extracted sequencing reads with insertion mutations from the PB-HiFi-1 and PB-HiFi-2 datasets (combined average sequencing depth=212×, combined NRPCC=56×) for the pancreatic cancer cell line HG008-T^42^. Candidate MEI supporting reads were chosen as those with insert sequences that could be aligned to consensus sequences of L1 (L1Hs), *Alu* (*Alu*Ya5*, Alu*Yk13*, Alu*Yb8*, Alu*Yb9*, Alu*Yc1 *and Alu*Ya5a2), and SVA (SVA-E and SVA-F)^57^. The ME alignment was performed with minimap v2.17-r941 (*minimap2 -a ME.fa insert.fa*)^73,75^. Each candidate insertion was then inspected and annotated for the presence of mobile element sequence, inversion, 3′ transduction, and other retrotransposition features. We called somatic MEIs based on the criteria as described below, using a merged BAM file of the tumor tissues (HG008-T) from PB-HiFi-1 and PB-HiFi-2, and a merged BAM file of the normal control tissues. Tumor somatic MEIs were identified as those with a minimum of two supporting reads in HG008-T, and no supporting reads in the control tissues. The identified somatic MEIs were further compared with PALMER (v2.0.1)^65^ and xTea_long (long-read module of xTea, v0.1.0)^51^ (**Supplementary Note 2**).

For L1 insertions, the supporting reads satisfy: (1) mapping quality ≥ 20; (2) mapping identity to L1Hs > 90% with an alignment length ≥ 50 bp; (3) the presence of at least one hallmark allele: ACA/G at L1Hs:5927-5929 or G at L1Hs:6012; (4) if transduction sequences were detected, their origin must be traceable to the flanking regions of active full-length reference or germline L1 elements; and (5) the presence of target site duplications (≥4 bp) and a polyA tail (≥10 bp), unless the alignment identity exceeded 95% and both hallmark alleles are present. For *Alu* insertions: (1) mapping quality ≥ 20; (2) mapping identity > 90% with alignment length ≥ 50 bp; and (3) exclusion of reads containing additional non-*Alu* sequences, as *Alu* elements do not undergo transduction. For SVA insertions: (1) mapping quality ≥ 20; (2) mapping identity > 90% with alignment length ≥ 50 bp; and (3) if transduction sequences were observed, their origin had to be traceable, as with L1.

#### Detecting somatic MEIs from Illumina sequencing data by xTea and TraFiC-mem

We benchmarked RetroNet with xTea (short-read module, v0.1.9)^51^ in the analysis of the simulated genome mixing dataset, the pancreatic cancer cell line tumor sample HG008-T, and the metastatic castration-resistant prostate cancer patient DTB-205. In each dataset, the xTea analysis was performed in the case-control mode to identify somatic MEIs using the default parameters. For the genome mixing dataset, the control was chosen as the pure NA12878 genomic DNA sequencing with the corresponding depth of 50×, 100×, 200×, and 400×. For HG008-T, the sequencing of the normal tissue was chosen as the control. For DTB-205, we compared both the matching tumor and control white blood cells (WBCs) to identify the true tumor somatic MEIs, and compared the cfDNA to WBCs to evaluate the performance of xTea in cfDNA sequencing. We further included the tumor somatic MEIs, possibly missed by xTea due to insufficient supporting reads, by including those with supporting reads in the tumor that are compatible with those identified by RetroNet in cfDNA. These additional somatic MEIs were assessed using RetroVis^16^, with details listed at https://github.com/Czhuofu/RetroNet.

In addition, we benchmarked another short-read-based MEI algorithm, TraFiC-mem (multispecies branch)^18,19^, which is compatible with sequencing alignments using the hg19 reference but not with the newer hg38 human reference genome. Among the various benchmarking datasets, only the HG008-T sequencing has hg19 alignments and has therefore been benchmarked with TraFiC-mem using the default parameters.

#### Statistical Analysis

The Wilcoxon test was used to estimate the difference between two groups of continuously distributed variables. P-values were adjusted for multiple comparisons using the Benjamini-Hochberg procedure (BH-adjusted *P*). The interquartile range, calculated as the 25^th^ and 75^th^ percentiles of the data, is used to describe the data dispersion.

**Supplementary Note 1:** Validating the Illumina sequencing-generated labels by PacBio HiFi sequencing

To validate the labels, we obtained PacBio HiFi sequencing from 11 randomly selected subjects from the Human Pangenome Reference Consortium (https://www.ncbi.nlm.nih.gov/bioproject/PRJNA701308)^76^ release 2 (https://github.com/human-pangenomics/hprc_intermediate_assembly/blob/main/data_tables/sequencing_data/README.md), of which 9 subjects were in our training dataset and 2 were included in the Polaris benchmarking dataset. We analyzed the MEIs from these long-read sequencing datasets using xTea_long (xTea version for the long reads, v0.1.0)^51^.

Among the 11 subjects, the average percentages of the *true* MEIs that were also detected in the PacBio data, within a window of 500bp, were 94.03% for L1, 97.35% for *Alu*, and 72.78% for SVA insertions. In contrast, the false labels had extremely low overlap with the PacBio calls: 0.09% for L1, 0.12% for *Alu*, and 0.20% for SVA (**Supplementary Table 9**). Thus the previous L1 and *Alu* labels, as defined by RetroSeq^25^ and MELT^73,74^ from the short read sequencing data, are in good concordance with the xTea_long analysis from the corresponding long read data.

Previous observations by Chu et al. (2023)^4^ have also documented a relatively low concordance for SVA calls between short-read and long-read sequencing data. Out of 716 SVA insertions identified from short-read sequencing of 39 subjects in their study, only 515 (71.9%) were found in the corresponding long-read sequencing. The additional SVA calls were validated by Sniffles2^77^ -- a long-read structural variant caller, or PCR validations^4^. This discrepancy is likely due to the limited sensitivity of current long-read tools in detecting SVA insertions. When compared to the more complete polymorphic SVA insertions as curated by Chu et al. (2023), an average of 97.22% of our *true* SVA insertions have matching calls.

Furthermore, we manually examined all discordant MEIs in the PacBio HiFi sequencing datasets to rule out true insertions possibly missed by xTea_long. The full reports for all 11 subjects are in the https://github.com/Czhuofu/RetroNet. After manual inspection, we confirmed that all discordant MEIs were true insertion mutations in the long-read data but were missed by the MEI analysis, resulting in a final validation ratio of 100% for all of the tested true L1, *Alu*, and SVA labels.

**Supplementary Note 2:** Comparison of the PacBio HiFi sequencing analysis in HG008-T with PALMER and xTea_long

We further compared the HG008-T somatic MEIs from PacBio HiFi sequencing datasets with existing long-read tools such as PALMER (v2.0.1)^65^ and xTea_long (long-read module of xTea, v0.1.0)^51^. Somatic tumor MEIs were defined as those that are present in HG008-T but absent in control tissues, using BEDTools^78^ with a window size of 500bp. Notably, the control tissues had substantially lower sequencing depths (65× and 35×), and both were considered during the filtering of germline MEIs. We used the default threshold for the number of supporting reads for xTea_long. For PALMER, we applied the recommended cutoff which is ≥ 1 high-confidence supporting reads and ≥ 10 all supporting reads (∼10% of the average sequencing depth).

PALMER identified 10 and 9 tumor somatic L1 insertions from PB-HiFi-1 and PB-HiFi-2, respectively, all of which were included in the L1 benchmark list. In contrast, xTea_long identified 11 tumor somatic L1 insertions from PB-HiFi-1, with 9 and 8 overlapping the benchmark list in PB-HiFi-1 and PB-HiFi-2, respectively. For the remaining xTea_long-reported insertions not present in the benchmark list, manual inspection using IGV revealed that the same insertions were also present in the normal tissues, although they were not called by xTea_long. Due to the relatively high supporting read thresholds required by PALMER and xTea_long, both tools demonstrated limited sensitivity in detecting low-frequency insertions. Among the nine L1 insertions with mosaicism below 10%, PALMER identified 3 in PB-HiFi-1 and 2 in PB-HiFi-2, while xTea_long identified only 1 in PB-HiFi-1 and none in PB-HiFi-2 (**Supplementary Table 10**).

PALMER also identified a total of 8 new *Alu* insertions, and 0 SVA insertions, while xTea_long discovered a different set of 15 *Alu* insertions, and 3 SVA insertions. These additional somatic MEIs, however, have also proven to be false, as manual inspection revealed that the same insertions were present in the matched normal datasets, or that the reported sites corresponded to structural variants rather than genuine MEIs (**Supplementary Table 10**). The full reports for the manual assessments for all L1, *Alu* and SVA insertions are in the https://github.com/Czhuofu/RetroNet.

**Supplementary Note 3:** Estimating the theoretical sensitivity in bulk and single-cell whole-genome Illumina sequencing

Using the methods outlined by Zhu et al.^16^, we simulated 200× and 400× bulk Illumina sequencing reads of a 10 kb DNA fragment containing an MEI located between 4500 and 5500 bp. The sequencing read length is 2×150bp, with an insert in between the paired reads following a normal distribution (mean=300bp, standard deviation=100bp). The MEI is a heterozygous mutation with a tissue allele frequency of 1%, equivalent to a frequency of 0.005 in bulk sequencing. In each of the 100,000 simulations, we sampled from a Poisson distribution (λ = 0.005) to determine the total number of reads covering the cells with the somatic MEI. We then estimated the likelihood of having more than two supporting reads covering the MEI junctions, assuming the sequencing reads are uniformly distributed across the 10 kb DNA fragment (**Figure 2a**). The simulation code can be found at https://github.com/Czhuofu/RetroNet.

For single-cell sequencing, the chance of detection is calculated as

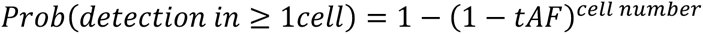

Simulation results for other tAFs, including 0.2%, 5%, and 25%, were shown in **Supplementary Table 8**.

**Supplementary Note 4:** Extending RetroNet for somatic MEIs with one supporting read

To test the possibility of identifying somatic MEIs with even lower tAFs, we adapted RetroNet to fit a new model for classifying MEIs using a single supporting read instead of the default two reads. Using the same training datasets, we included additional MEIs with only one supporting read, all labeled as false insertions. We then encoded each individual supporting read into images using a similar approach as **Figure 2**, with modifications to include only one supporting read instead of two, and trained a classifier model based on the ResNet-18. Compared to the two-read model, the one-read model had notably lower accuracies: L1 (two-read AUPR: 0.990, 95% CI: 0.988-0.993; one-read AUPR: 0.937, 95% CI: 0.932-0.942), *Alu* (two-read AUPR: 0.9996, 95% CI: 0.9994-0.9998; one-read AUPR: 0.9864, 95% CI: 0.9851-0.9876), and SVA insertions (two-read AUPR: 0.999, 95% CI: 0.998-0.999; one-read AUPR: 0.978, 95% CI: 0.975-0.982) (**Supplementary Figure 10a**). Other performance metrics, including precision and recall, also showed similar differences between the one-read and two-read models (**Supplementary Figure 10b**).

Utilizing just one supporting read for MEI detection tends to introduce considerably more noise — approximately 80 times greater than using two reads, based on a previous estimate^16^. Consequently, the SNR from the one-read model could be significantly worse in real-world applications. To investigate this, we evaluated the one-read model on a series of images sampled from the Polaris Project dataset with the SNRs set at 1:100, 1:1000, and 1:10000. As expected, the precision declined considerably as the SNRs decreased, even when employing very high cutoff values (**Supplementary Figure 10c** and 10d). This suggests that the single supporting read approach may not be well-suited for practical applications due to its limited effectiveness in highly imbalanced datasets.

