## Supplementary Figures for "Image-based DNA Sequencing Encoding for Detecting Low-Mosaicism Somatic Mobile Element Insertions"

**Supplementary Figure 1. The frequency distribution of true MEIs across all subjects in the training data.** The violin plots illustrate the frequency distributions of three major MEI types: L1, *Alu,* and SVA. The width of each plot reflects the data density across different frequency levels. The white boxes within the plots denote the interquartile range, while the dots represent individual data points.

**Supplementary Figure 2. The additional syntaxes used for encoding candidate MEI-supporting reads.** Mobile element insertion (MEI) example with a 5' inversion and an unmappable segment insertion at the 3' end. The blue, red, and magenta lines represent the human reference, mobile element consensus, and mobile element reverse complement, respectively. The black box indicates a possible unmappable segment within the ME consensus. (**a**) When the human anchor (split or clipped PE read) contains a possible unmappable segment, the clipped segment is shown as a proportional black line. The segment mappable to the ME consensus is represented by a red line below, with its sequence encoded by red pixels. (**b**) At the end of the ME end (PE or clipped PE read), an unmappable segment is displayed as a proportional black line above the ME sequence. Other insertion gaps in the ME end are marked with blue pixels above the ME sequence at the corresponding insertion loci. (**c**) MEI with inversions: clipped-PE reads mapped to the L1 opposite strand are denoted by magenta instead of red pixels in the ME consensus track. (**d**) Green lines indicate unmappable segments supporting *Alu* insertion carrying transduction. (**e**) Image encoding of the *Alu* and SVA insertion sequences: For *Alu* insertions, four tracks portray the sequence alignments to each *Alu* subfamily (*Alu*Ya, *Alu*Yb, *Alu*Yc, and *Alu*Yk). Similarly, two tracks portray the sequence alignments to each SVA subfamily (SVA_E and SVA_F).

**Supplementary Figure 3. The training progress of the three deep learning models.** (**a**) The updates of the loss function value during the training process for both the training and validation data of L1, *Alu* and SVA. (**b**) Precision, recall and AUPR of the final model, selected at the epoch with the lowest validation loss.

**Supplementary Figure 4. Precision-recall curves of RetroNet, RetroSom, and xTea for predicting simulated low-tAF Alu insertions.** The precision-recall curves were evaluated for detecting somatic *Alu* insertions at mixing proportions of 0.2%-25% (y-axis) under sequencing depth of 50× to 400× (x-axis), using RetroNet (red), RetroSom (gray) and xTea (green). Based on the AUPR scores, RetroNet could consistently outperform RetroSom at detecting low-mosaicism *Alu* insertions. In contrast, xTea only predicts *Alus* with relatively high tAF and performs significantly worse than RetroNet.

**Supplementary Figure 5. Precision-recall curves of RetroNet and xTea for predicting simulated low-tAF SVA insertions.** The precision-recall curves were evaluated for detecting somatic SVA insertions at mixing proportions of 0.2%-25% (y-axis) under sequencing depth of 50× to 400× (x-axis), using RetroNet (red) and xTea (green). RetroNet demonstrated the ability to detect SVA insertions even at low tAF and low sequencing depths, whereas xTea primarily identified SVAs with high tAF and sequencing depths.

**Supplementary Figure 6. Encoded images and the RetroNet probabilities for two validated somatic L1 insertions L1_1 and L1_2.** RetroNet was applied to the DNA whole-genome-sequencing data of the neurons of schizophrenia brain 12004. Identified two previously experimental validated L1 insertions: (**a**) L1_1 at the chr10:102989221 has 1 image with a probability exceeding 0.99, and (**b**) L1_2 at the chr10:14204877 has 5 images with a probability exceeding 0.99.

**Supplementary Figure 7. Class activation maps for predicting L1, *Alu* and SVA insertions.** Gradient-weighted Class Activation Mapping (Grad-CAM) visualizations for RetroNet predicting mobile elements, including (**a**) L1, (**b**) *Alu*, and (**c**) SVA. These activation maps highlight that RetroNet correctly focuses on relevant regions during true mobile element insertion predictions, such as positional information (indicated by blue arrows) and ME sequence information (indicated by red pixels).

**Supplementary Figure 8. Representative false positive L1 insertions predicted by RetroSom.** RetroVis visualizes all supporting reads for an L1 insertion, connecting each human anchor with the L1 end of the read using a black line. In each plot generated by RetroVis, the top axis represents the insertion region on the human reference genome, where human anchors (blue arrows) align, while the bottom axis represents the mapped region of the L1HS consensus sequence, where L1 ends (red arrows) align. (**a**) The close proximity of human anchors suggests an insertion near chr18:22690568. The two supporting reads at the top indicate that the 3’ end of this insertion is located upstream of L1Hs:5783, while the last supporting read’s human anchor is further from the insertion site, with its L1 end mapped near L1Hs:4867. The last supporting read exhibits a gap of over 1000 bp between human anchor and L1 end, which is unlikely for Illumina sequencing. (**b**) The two identical supporting reads may result from PCR duplication. (**c**) Two supporting reads display different positions for human anchors but identical L1 ends. (**d**) Two supporting reads show varied L1 end positions but identical short human anchors.

**Supplementary Figure 9. Patient DTB-205 cfDNA somatic L1 insertions identified by RetroNet with low mosaicism in the tumor.** The enhanced RetroVis plots depict two low-mosaicism somatic L1 insertions that were detected by RetroNet in circulating cell-free DNA (cfDNA) but not by xTea short owing to their low mosaicism in the tumor. A continuous blue line at the top denotes the genomic coordinates in the human reference genome, whereas a continuous red (L1Hs+) or purple (L1Hs-) line at the bottom represents the corresponding region of the L1Hs consensus sequence on the positive or negative strand, respectively. The upper red rectangle contains supporting reads from cfDNA, and the lower red rectangle contains supporting reads from the tumor. Paired-end (PE) supporting reads are illustrated as a blue arrow (human anchor) joined to a red or purple arrow (L1 end) by a continuous black line. Split-read (SR) supporting reads are displayed as a blue arrow (reference segment) linked to an empty rectangle (L1 segment), and a red or purple arrow beneath. All SRs are outlined by a blue rectangle for clarity. A vertical blue dashed line marks the insertion breakpoint in the genome, and a pair of red dashed lines delineate the insertion range of the L1. (**a**) chr1:49608812-49608854. All PE human anchors in cfDNA and tumor point to approximately chr1:49608800, and their L1 ends align to the positive strand. The insertion range of L1Hs:4176-6064 indicates a 5′-truncated insertion on the human positive strand. (**b**) chr4:54692604-54692686. Two SRs in cfDNA refine the breakpoint to chr4:54692600-54692700. Both cfDNA and tumor PE human anchors coincide with this SR-defined breakpoint, and their L1 ends align to L1Hs:6049-5500, indicating this 5'-truncated L1 on the human negative strand.

**Supplementary Figure 10. Performance benchmarking for the classification model using one supporting read.** (**a**) Precision-recall curves comparing RetroNet models using one versus two pairs of supporting reads, assessed on labeled L1, *Alu*, and SVA image datasets extracted from 49 individuals in the Illumina Polaris Project. (**b**) Statistical comparisons of precision, recall, and AUPR for the RetroNet models using one versus two pairs of supporting reads. (**c**) Precision-recall curves for the one-read RetroNet model in identifying MEIs within simulated imbalanced datasets, with signal-to-noise ratios (SNRs) of 1:100 (light green), 1:1000 (dark green), and 1:10000 (purple). Solid lines indicate average PR curves from 50 simulations, while ribbons show the 95% confidence interval. (**d**) Effect of varying probability cutoffs (ranging from 0.5 to 0.99, depicting by different colors) on the precision and recall of the one-read RetroNet model when applied to imbalanced datasets with SNRs of 1:100 (circle), 1:1000 (triangle), and 1:10000 (square).

**Supplementary Figure 11. Limitation for detecting complex retrotransposition with short read sequencing.** (**a**) Illustration of mobile element (ME) retrotransposition featuring 5'-inversion, highlighting the opposing orientations of the 5'-end and 3'-end. The long blue, red, and magenta lines represent the human reference, ME consensus, and ME reverse complement, respectively. The short magenta line indicates the read end mapped to the ME reverse complement, which is classified by the program as a reverse end. Consequently, this forward-reverse paired read is discarded. This configuration poses a challenge for RetroNet, as both supporting reads must align to the same strand of the ME consensus sequence. As a result, low-mosaicism events with insufficient supporting reads covering both ends may be missed. (**b**) The downward arrow illustrates that the 3' end of the L1 insertion (locus 2 of the human reference) carries a flanking DNA segment (green box) from the source L1 (locus 1 of the human reference). Therefore, the two ends of a read may map to either locus 2 and L1Hs, in which case they would be retained; locus 1 and L1Hs, where they could be considered a fixed L1 embedded in the human genome and thus discarded; or locus 1 and locus 2, which would also result in discarding. The upward arrow demonstrates that, in orphan transduction, the flanking DNA segment is transduced and inserted without the accompanying L1 element, making it unidentifiable.

**Supplementary Figure 1. The frequency distribution of true MEIs across all subjects in the training data**

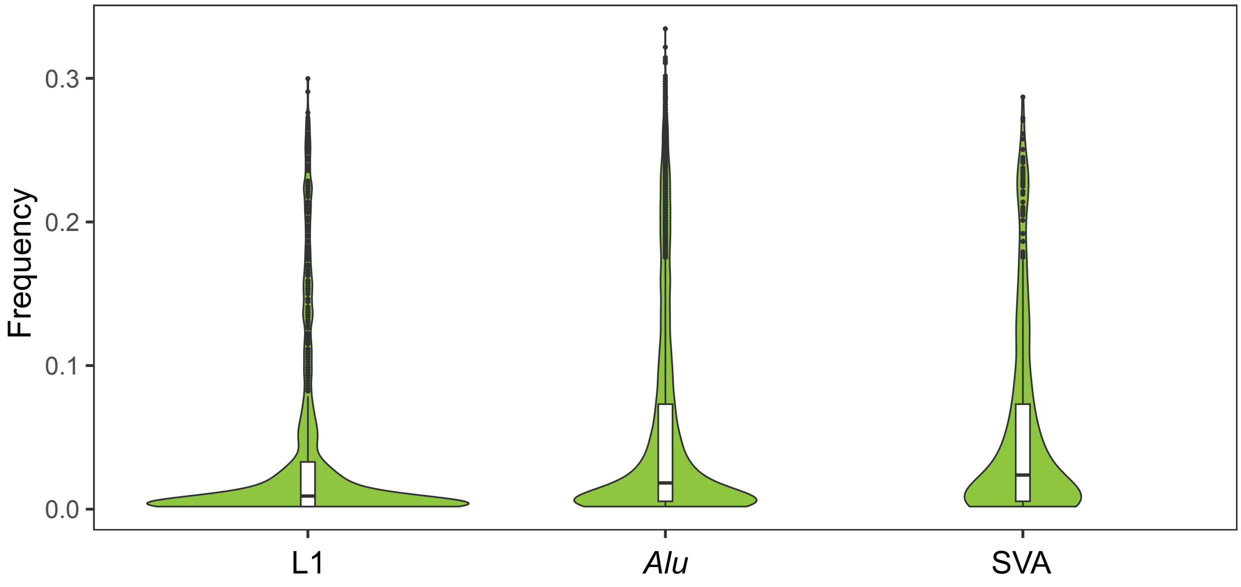

#### Supplementary Figure 2. The additional syntaxes used for encoding candidate MEI-supporting reads

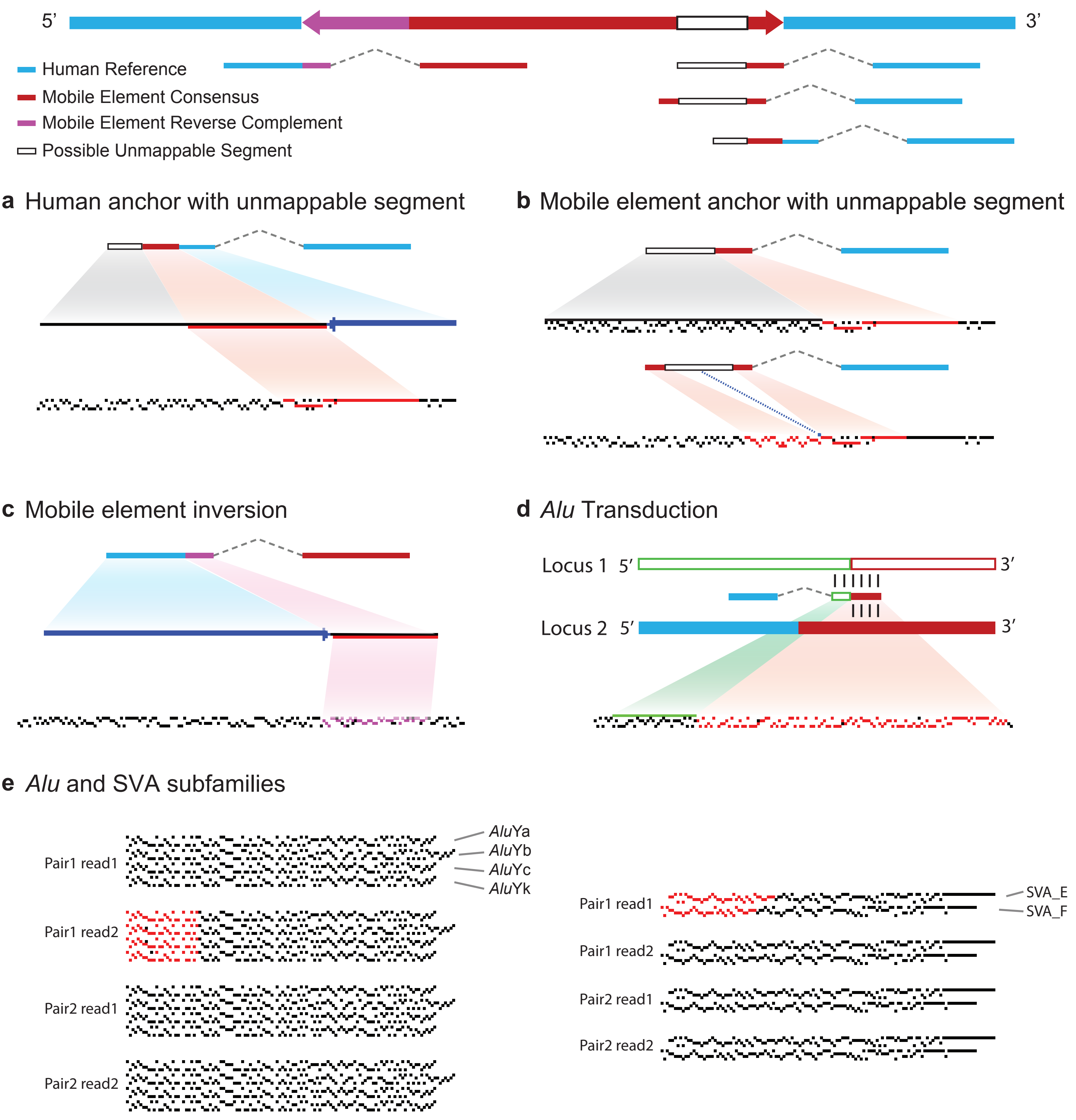

**Supplementary Figure 3. The training progress of the three deep learning models**

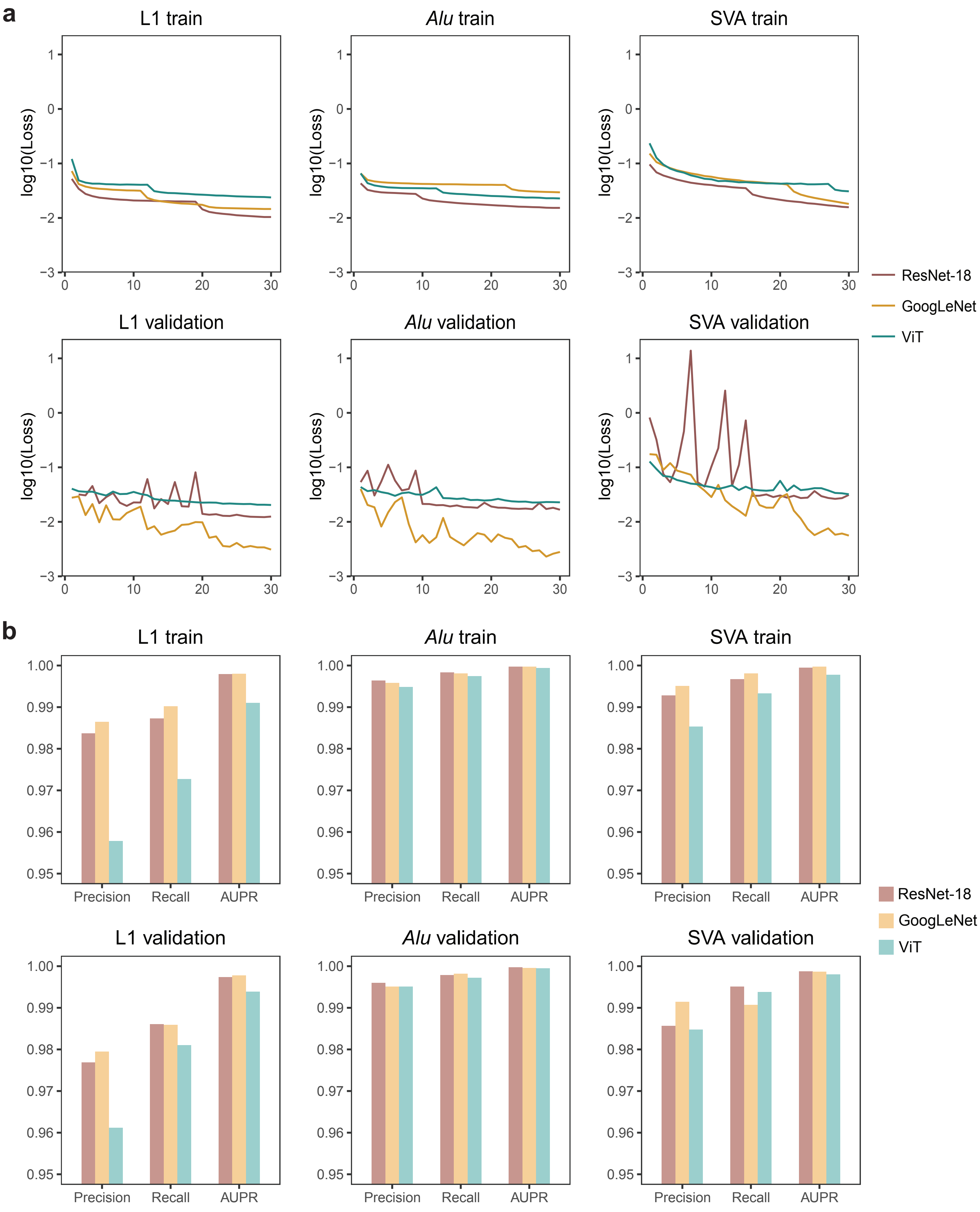

**Supplementary Figure 4. Precision-recall curves of RetroNet, RetroSom, and xTea for predicting simulated low-tAF *Alu* insertions**

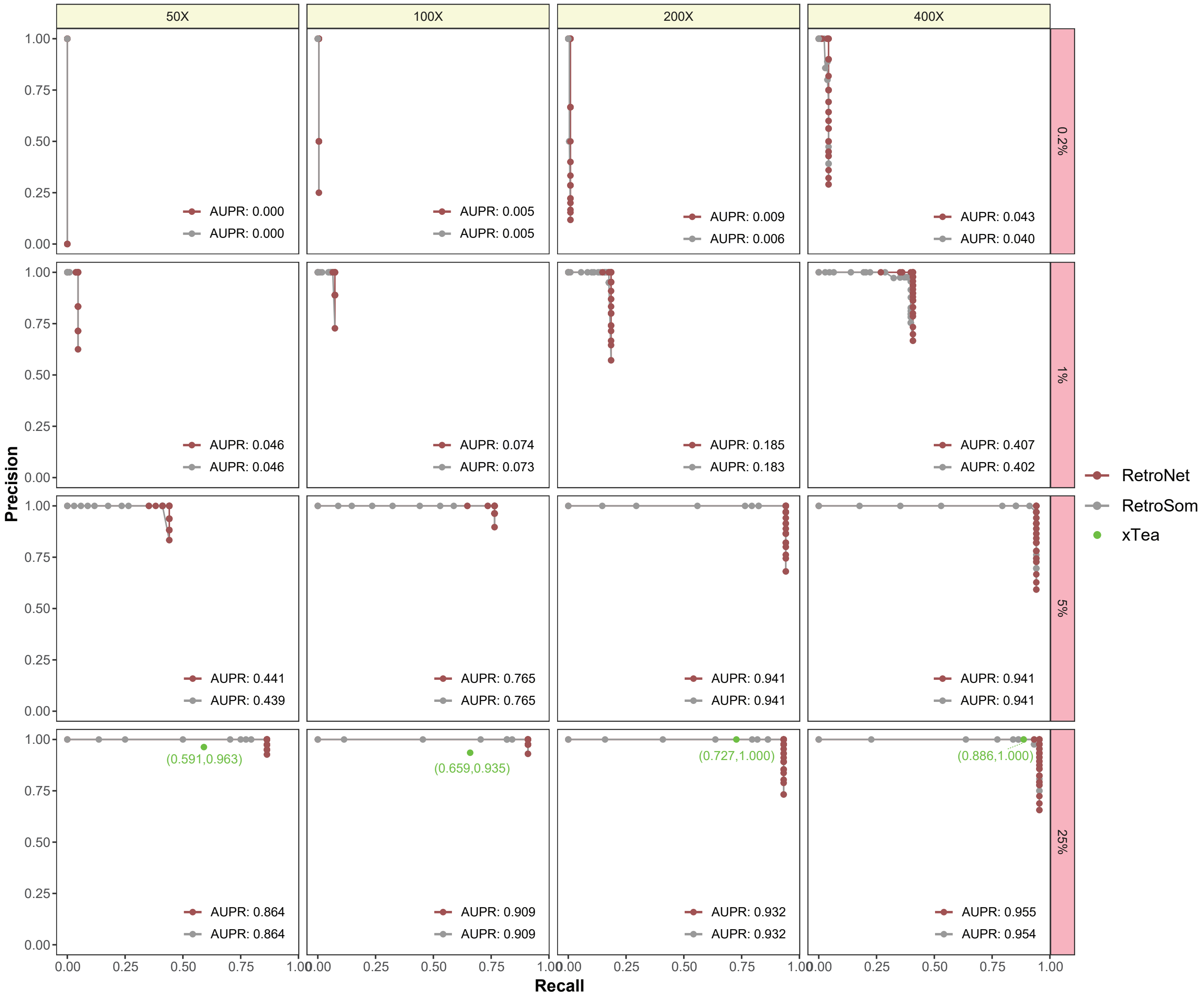

### Supplementary Figure 5. Precision-recall curves of RetroNet and xTea for predicting simulated low-tAF SVA insertions

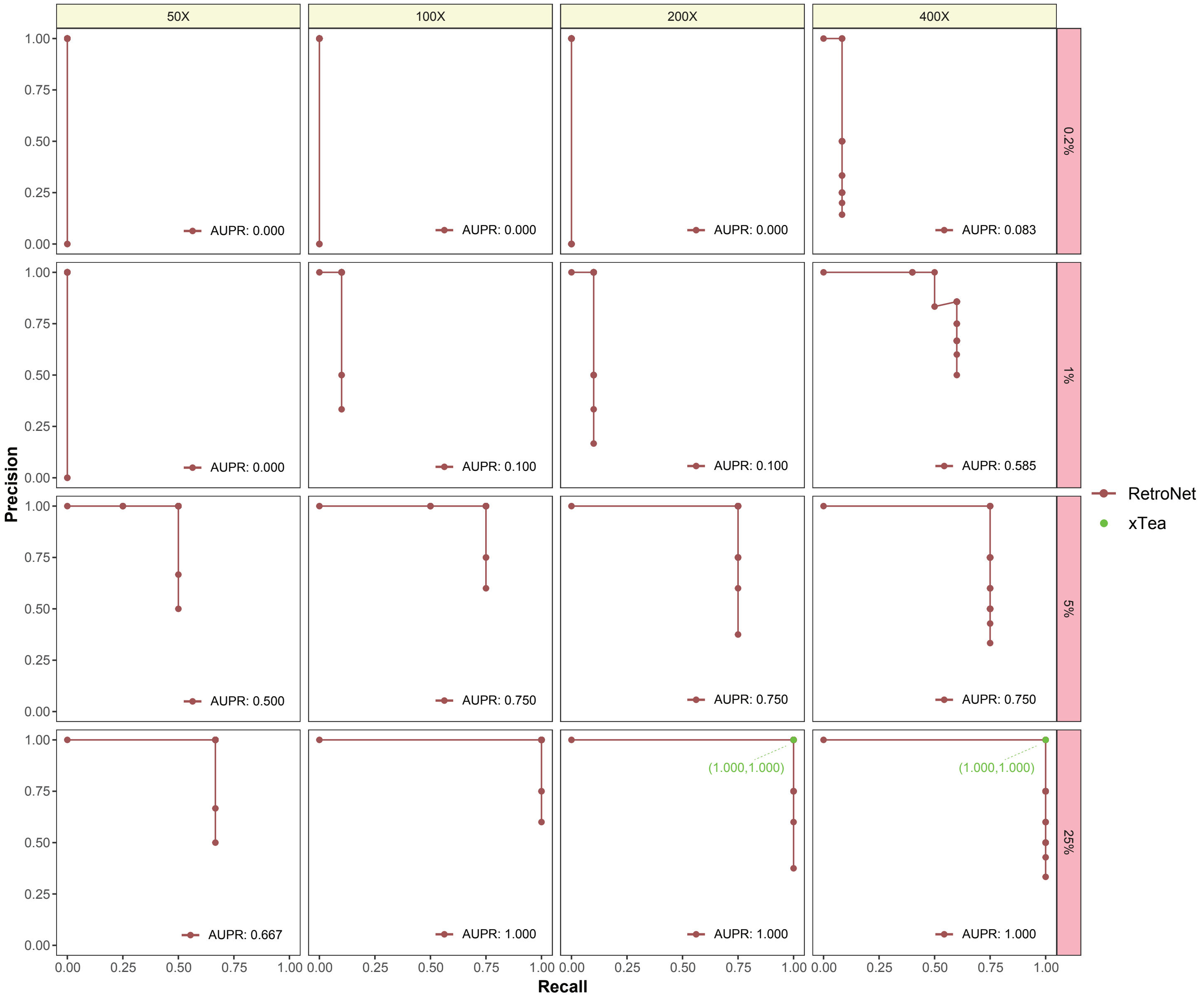

**Supplementary Figure 6. Encoded images and the RetroNet probabilities for two validated somatic L1 insertions L1_1 and L1_2**

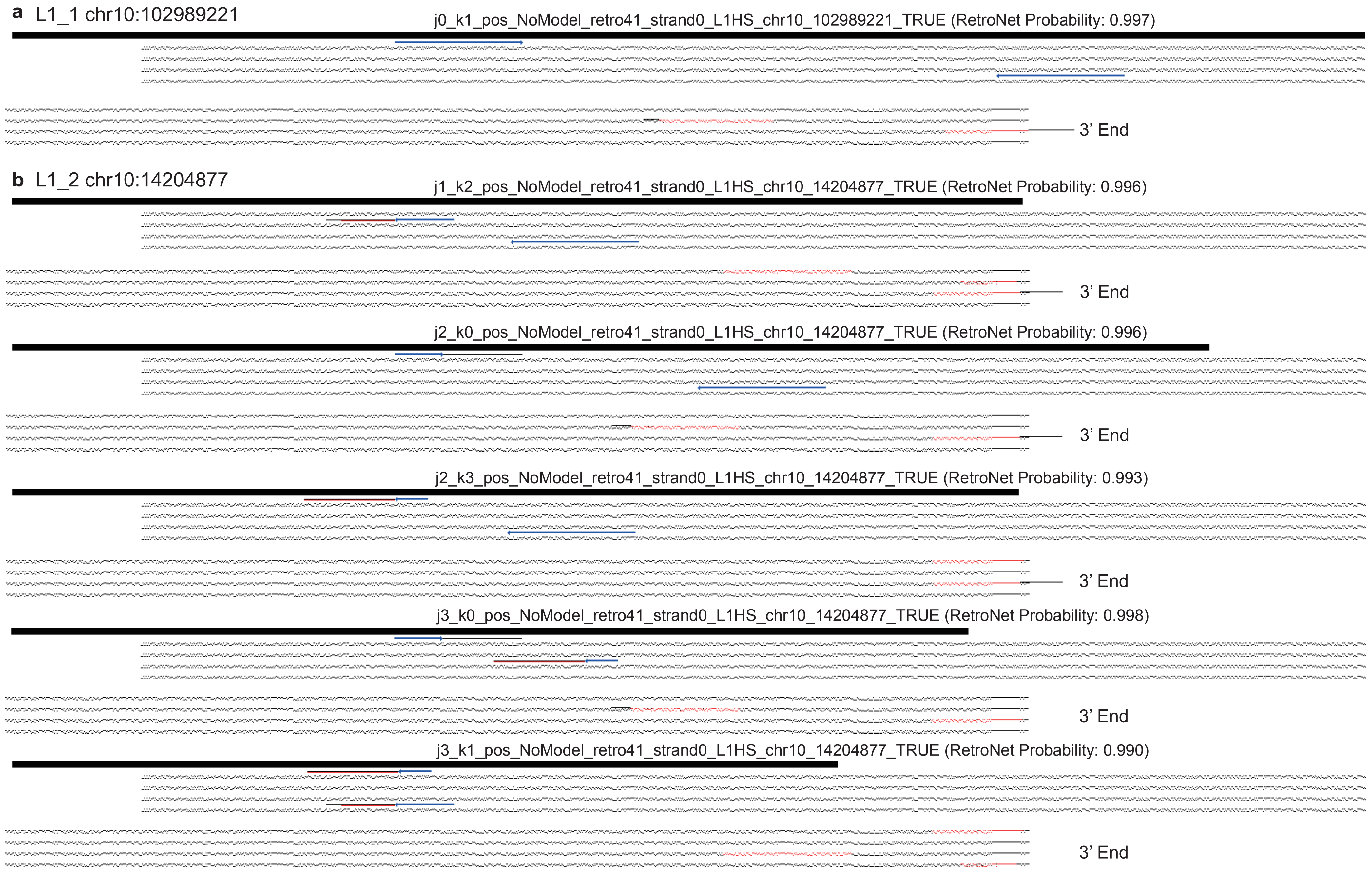

**Supplementary Figure 7. Class activation maps for predicting L1, *Alu* and SVA insertions**

**a** Activation Map for L1 Prediction

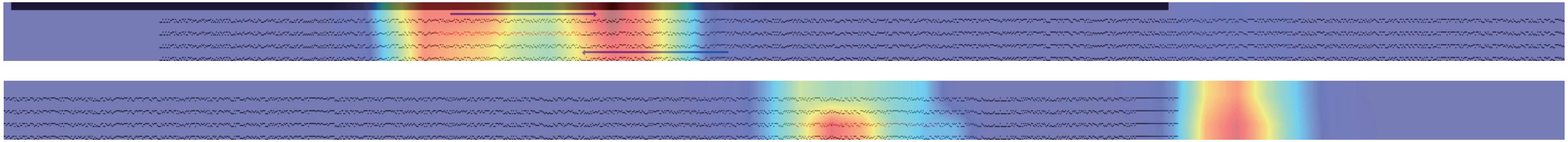

**b** Activation Map for *Alu* Prediction

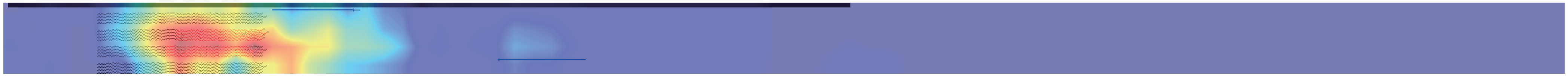

**c** Activation Map for SVA Prediction

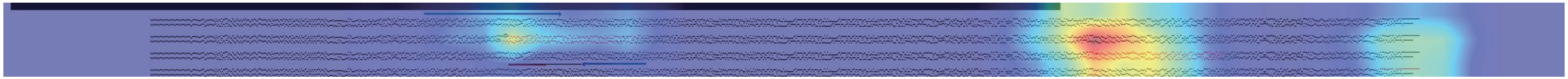

**Supplementary Figure 8. Representative false positive L1 insertions predicted by RetroSom**

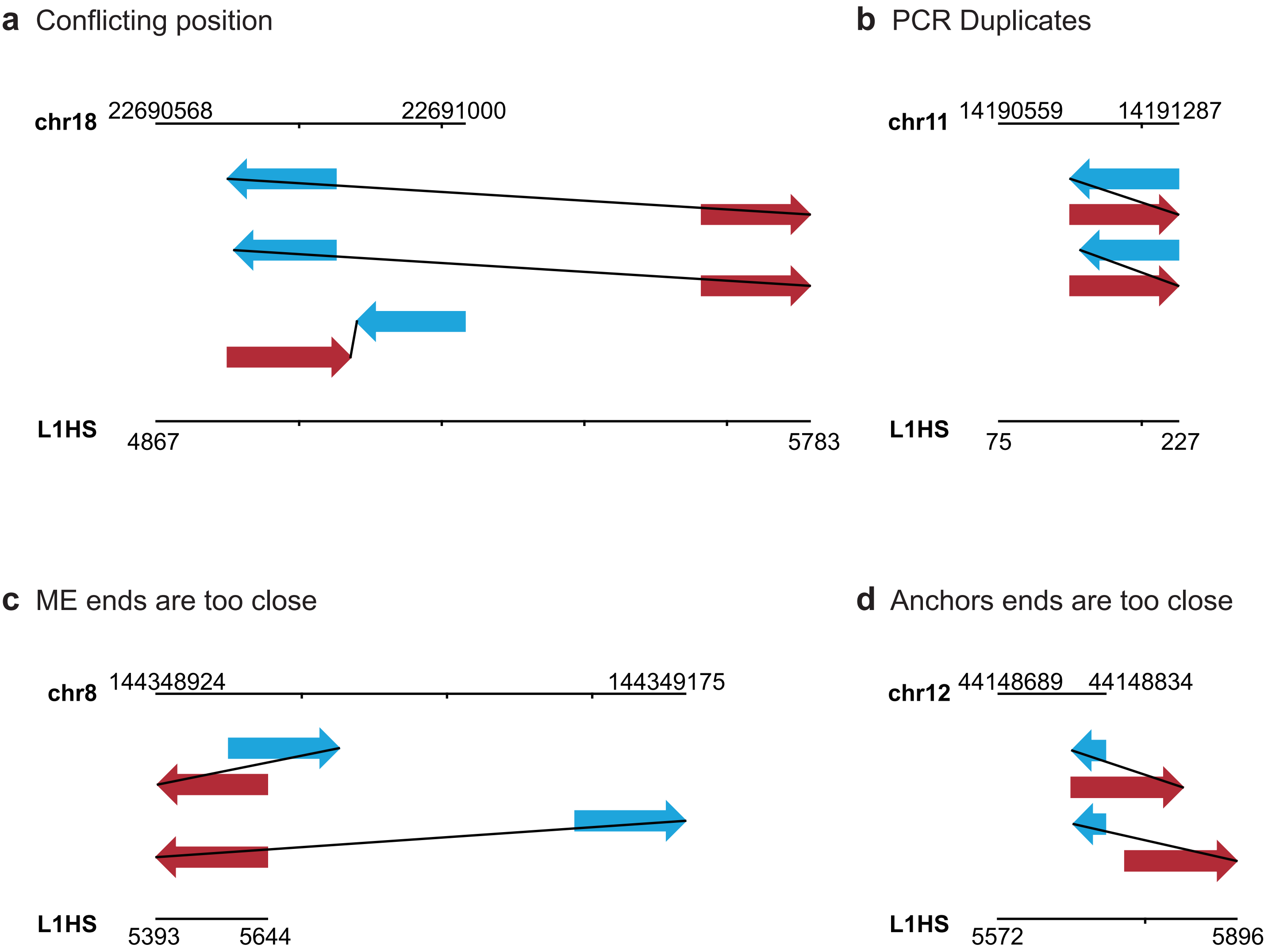

**Supplementary Figure 9. Patient DTB-205 cfDNA somatic L1 insertions identified by RetroNet with low mosaicism in the tumor**

**a** chr1:49608812-49608854

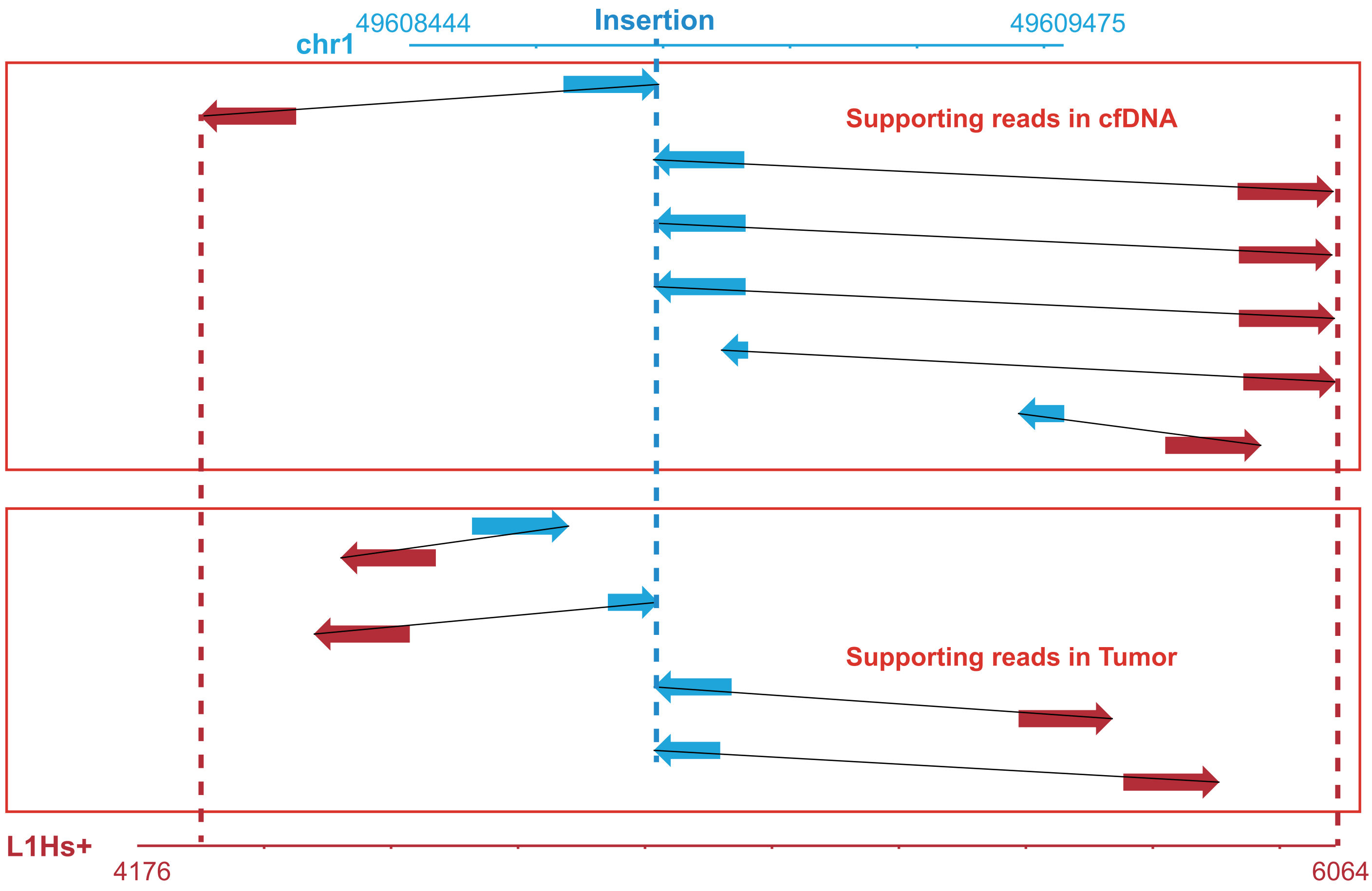


**b** chr4:54692604-54692686

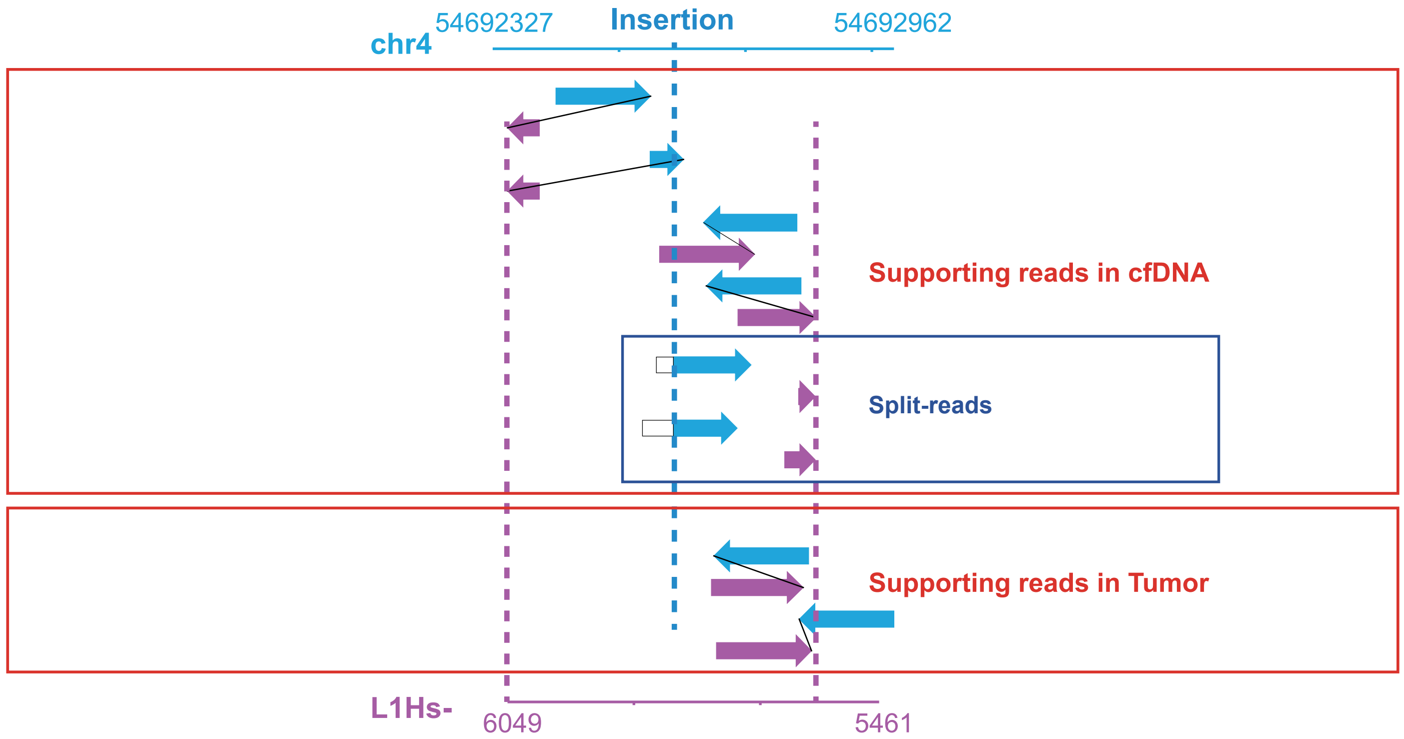

### Supplementary Figure 10. Performance benchmarking for the classification model using one supporting read

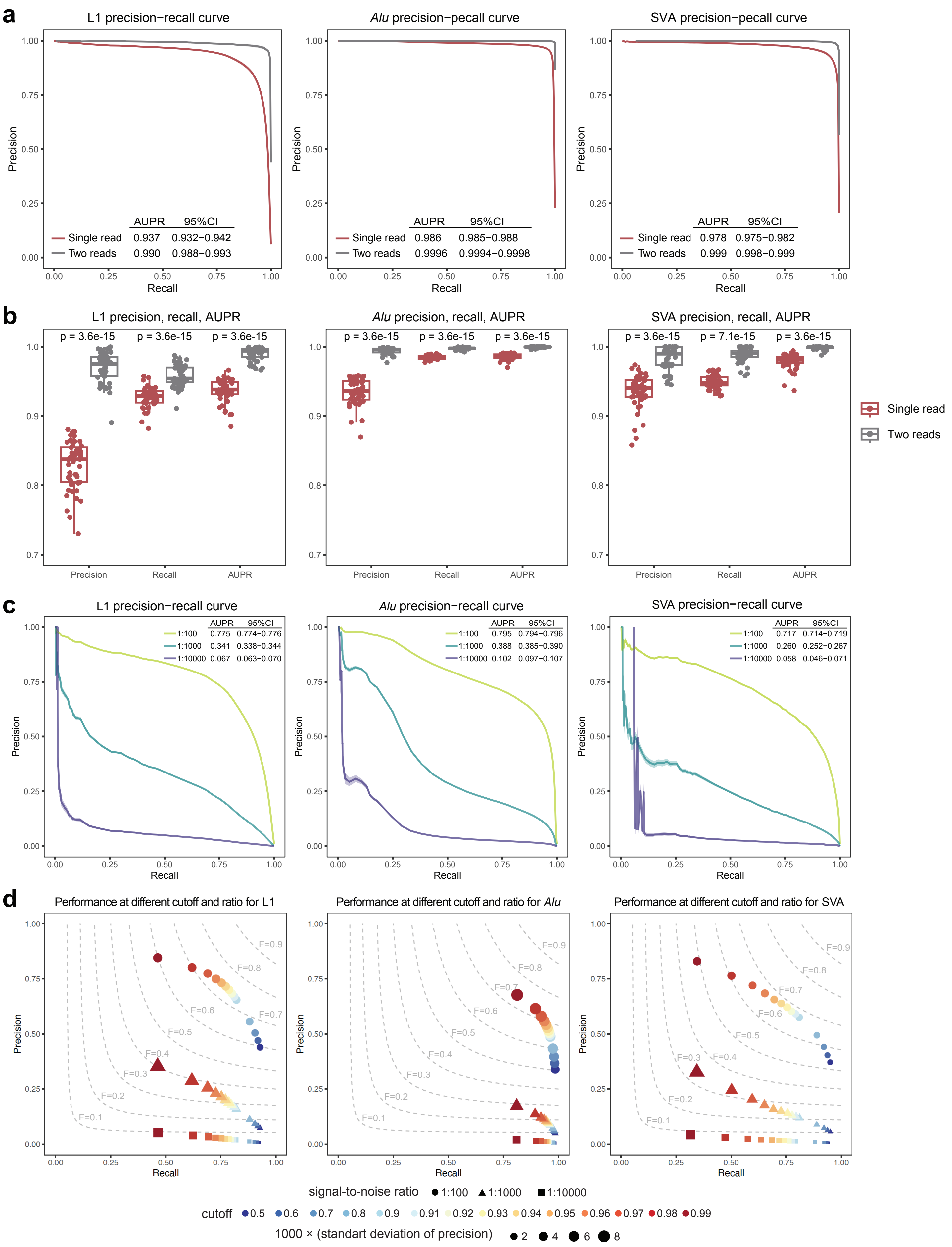

#### Supplementary Figure 11. Limitation for detecting complex retrotransposition with short read sequencing

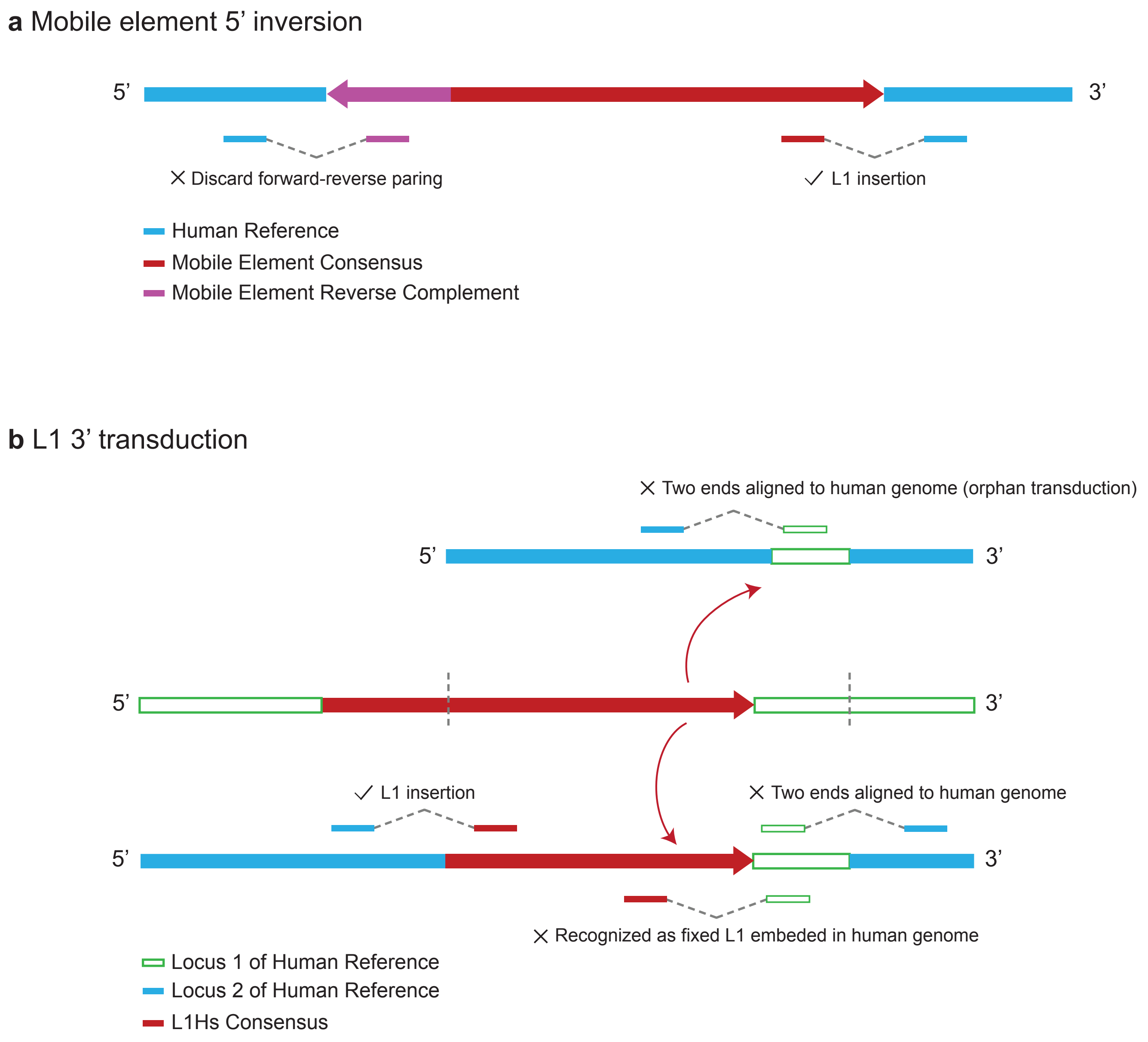
