## Extended Files for "Image-based DNA Sequencing Encoding for Detecting Low-Mosaicism Somatic Mobile Element Insertions": Extended Files 1.pdf

### Manual check of L1 label that were not identified by Pangenome xTea_long PacBio calls

(Additional explanation for Supplementary Table 8)

| ID | Chr | Coordinate | Manual Inspection |
|---|---|---|---|
| HG00438 | chr5 | 57983715 | TRUE |
| | chr5 | 161008649 | TRUE |
| | chr6 | 16950579 | TRUE |
| | chr9 | 70162482 | TRUE |
| | chr18 | 69865422 | TRUE |
| HG00621 | chr5 | 57983715 | TRUE |
| | chr6 | 13190184 | TRUE |
| | chr20 | 17880269 | TRUE |
| HG02630 | chr2 | 141016817 | TRUE |
| | chr7 | 106327425 | TRUE |
| | chr12 | 63175375 | TRUE |
| | chr16 | 15686361 | TRUE |
| HG03492 | chr7 | 106327425 | TRUE |
| HG03516 | chr3 | 103981000 | TRUE |
| | chr5 | 57983715 | TRUE |
| | chr6 | 103423568 | TRUE |
| | chr7 | 106327425 | TRUE |
| | chr9 | 118632302 | TRUE |
| | chr12 | 126318373 | TRUE |
| HG03540 | chr1 | 48963060 | TRUE |
| | chr7 | 106327425 | TRUE |
| | chr20 | 17880269 | TRUE |
| NA18906 | chr1 | 226583194 | TRUE |
| | chr5 | 19180266 | TRUE |
| | chr7 | 106327425 | TRUE |
| NA20129 | chr5 | 57983715 | TRUE |
| | chr6 | 16950580 | TRUE |
| | chr17 | 35945138 | TRUE |
| ERX2355869 (HG03579) | chr3 | 166374422 | TRUE |
| | chr4 | 68787387 | TRUE |
| | chr5 | 161008649 | TRUE |
| | chr12 | 126318373 | TRUE |

### HG00438 chr5:57983715

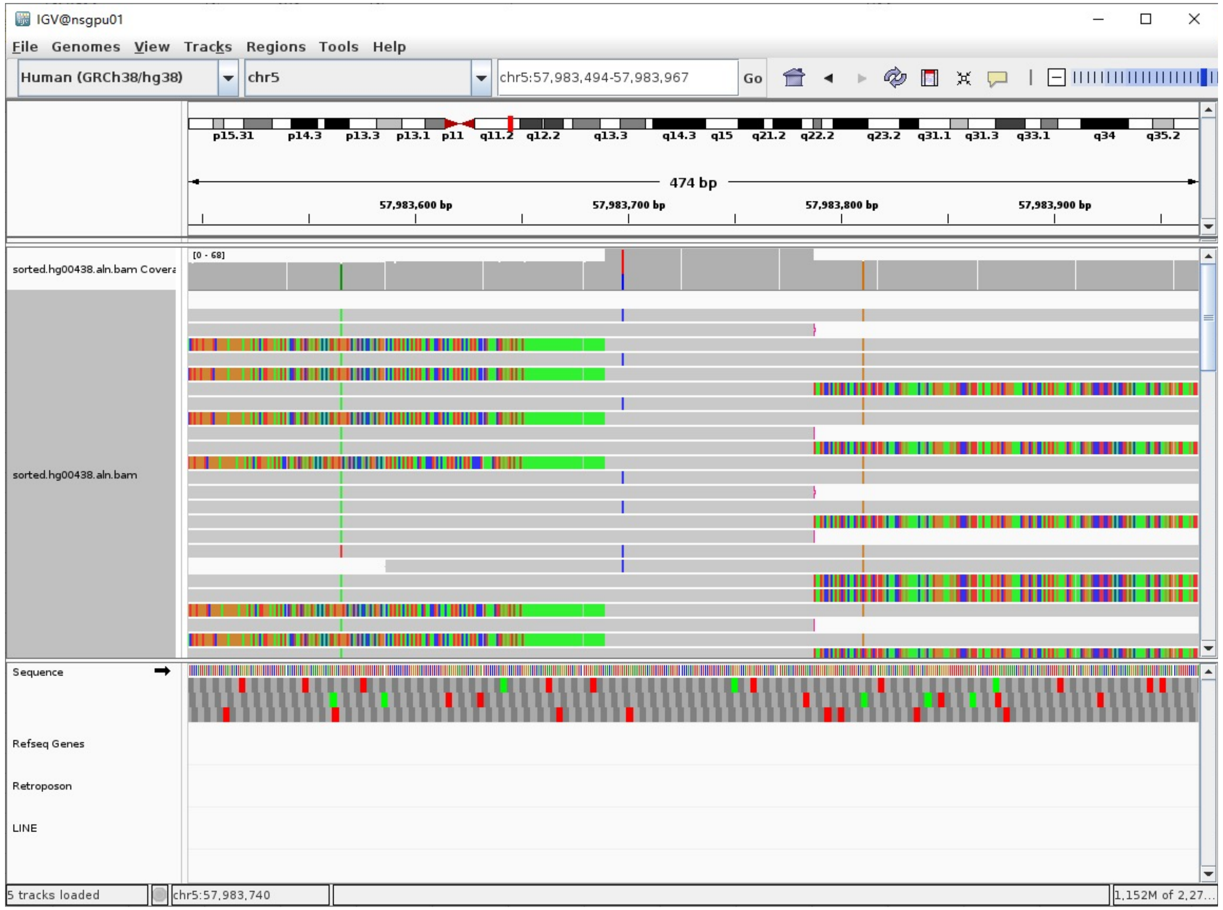


Left-clipped read (9949bp)

- Clip:1-8597 aligns to chr5:57975184-57983787

AACATTTTACTTGGCTGCCTGGAAAAAGCTGAGATGTGATTCTGATTACAAGTTTGGAAGCAAT
TAAAAATGGTGGGATTGGGAGAGGAGAAGATTTGACACTGTACTGCAAAGAAGTCATCGCAGTA
TCTTCAAGCTAGTCAGAACTGGCCT…

- **94bp TSD**: clip:8504-8597 aligns to chr5:57983694-57983787

TTTGTATAGAAATATACATTCTTTCCATTGTCCCCACATTGCTCCTTCTCTATCCATGGGCCAGTGATTT
GCATCCAGTATACACTTATTAACT

- Clip:8598-9949 aligns to **L1Hs**:4706-6058, 99.33% identity, **ACA TAG**

TAATGCCGCATATCTACAACTATCTGATCTTTGACAAACCTGAGAAAAACAAGCAATGGGAAAGGATTC
CCTATTTAATAAATGGTGCTGGGAAAACTGGCTAGCCATATGTAGAAAGCTGAAACTGGATCCCTTCCTT
ACACCTTATACAAAAATCAATTCAAGATGGATTAAAGATTTAAACGTTAAACCTAAAACCATAAAAACCC
TAGAAGAAAACCTAGGCATTACCATTCAGGACATAGGCGTGGGCAAGGACTTCATGTCCAAAACACCAAA
AGCAATGGCAACAAAAGACAAAATTGACAAATGGGATCTAATTAAACTAAAGAGCTTCTGCACAGCAAAA
GAAACTACCATCAGAGTGAACAGGCAACCTACAACATGGGAGAAAATTTTTGCAACCTACTCATCTGACA
AAGGGCTAATATCCAGAATCTACAATGAACTCAAACAAATTTACAAGAAAAAAACAAACAACCCCATCAA
AAAGTGGGCGAAGGACATGAACAGACACTTCTCAAAAGAAGACATTTATGCAGCCAAAAAACACATGAAG
AAATGCTCATCATCACTGGCCATCAGAGAAATGCAAATCAAAACCACTATGAGATATCATCTCACACCAG
TTAGAATGGCAATCATTAAAAAGTCAGGAAACAACAGGTGCTGGAGAGGATGCGGAGAAATAGGAACACT
TTTACACTGTTGGTGGGACTGTAAACTAGTTCAACCATTGTGGAAGTCAGTGTGGCGATTCCTCAGGGAT
CTAGAACTAGAAATACCATTTGACCCAGCCATCCCATTACTGGGTATATACCCAAATGAGTATAAATCAT
GCTGCTATAAAGACACATGCACACGTATGTTTATTGCGGCACTATTCACAATAGCAAAGACTTGGAACCA
ACCCAAATGTCCAACAATGATAGACTGGATTAAGAAAATGTGGCACATATACACCATGGAATACTATGCA
GCCATAAAAAATGATGAGTTCATATCCTTTGTAGGGACATGGATGAAATTGGAAACCATCATTCTCAGTA
AACTATCGCAAGAACAAAAAACCAAACACCGCATATTCTCACTCATAGGTGGGAATTGAACAATGAGATC
ACATGGACCCAGGAAGGGGAATATCACACTCTGGGGACTGTGGTGGGGTCGGGGGAGGGGGAGGGATAG
CATTGGGAGATATACCTAATGCTAGATGACACATTAGTGGGTGCAGCGCACCAGCATGGCACATGTATAC
ATATGTAACTAACCTGCACAATGTGCACATGTACCCTAAAACTTAGAGTATAATAAAAAAAAAAAAAAA
AAAAAAAAAAAAAAAAAAAAAA

- Illustration:

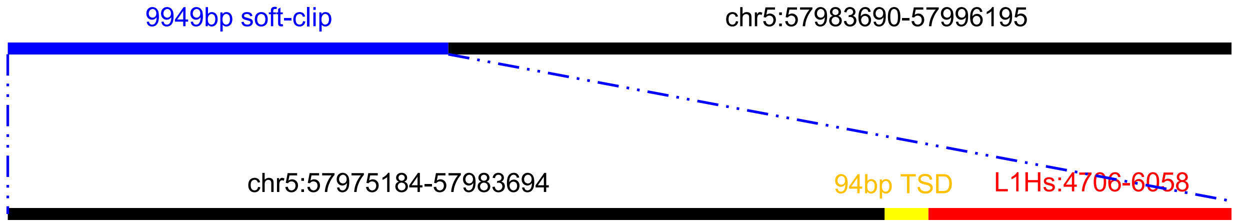

### HG00438 chr5:161008649

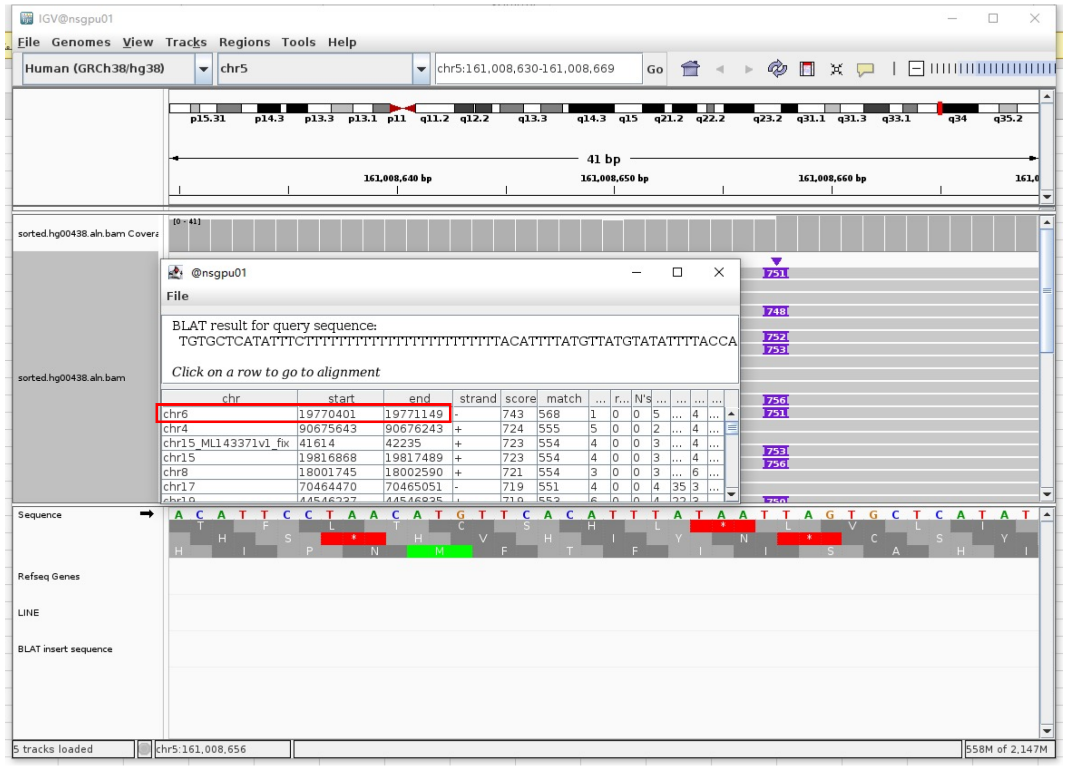


Insertion sequence (746bp)

- Insertion:1-14

GTGCTCATATTTCT

- Insertion:15-37, PolyT

TTTTTTTTTTTTTTTTTTTTTTT

- Insertion:38-168 aligns to chr3:173031214-173031344

| QUERY | SCORE | START | END | QSIZE | IDENTITY | CHROM | STRAND | START | END | SPAN |
|---|---|---|---|---|---|---|---|---|---|---|
| YourSeq | 131 | 38 | 168 | 202 | 100.0% | chr3 | + | 173031214 | 173031344 | 131 |

- Insertion:203-746 aligns to L1Hs:6064-5520, 96.86% identity, ACA

TTTTTTTTTTTTGTTTTTTTTTTTTTTTTCTTTTTTTTTTTTTTTTATTATACTCTCAGTTTT
AGGGTACATGTGCACATTGTGCAGGTTAGTTACATATGTATACATGTGCCATGCTGGTGCG
CTGCACCCACTAATGTGTCATCTAGCATTAGGTATATCTCCCAATGCTATCCCTCCCCCCT
CCCCCGACCCCACCACAGTCCCCAGAGTGTGATATTCCCCTTCCTGTGTCCATGTGATCTC
ATTGTTCAATTCCCACCTATGAGTGAGAATATGCGGTGTTTGGTTTTTTGTTCTTGCGATA
GTTTACTGAGAATGATGGTTTCCAATTTCATCCATGTCCCCTACAAAGGATATGAACTCAT
CATTTTTTATGGCTGCATAGTATTCCATGGTGTATATGTGCCACATTTTCTTAATCCAGAC
TATCATTGTTGGACATTTGGGTTGGTTCAAGTCTTTGCTATTGTGAATAGTGCCGCAATAA
ACATACGTGTGCATGTGTCTTTATAGCAGCATGGGTATATACCCAAAGGACTATA

- Illustration (L1 with transduction):

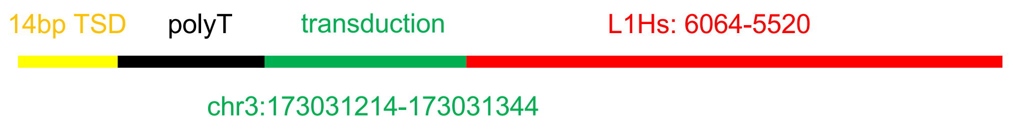

### HG00438 chr6:16950579

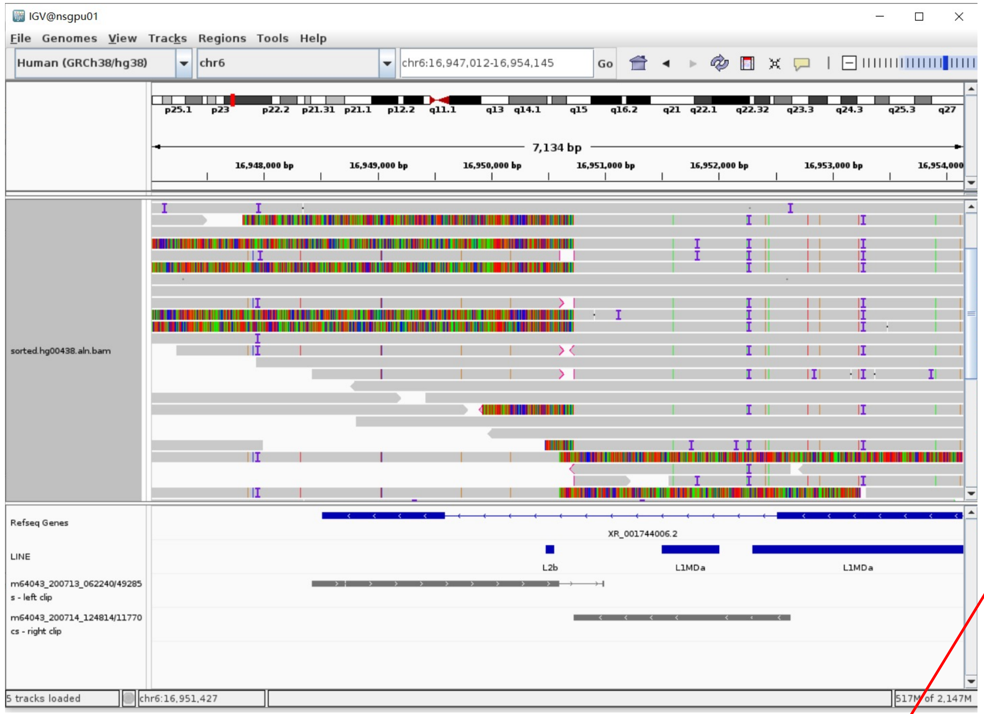

| ACTIONS | | | QUERY | SCORE | START | END | QSIZE | IDENTITY | CHROM | STRAND | START | END | SPAN |
|---|---|---|---|---|---|---|---|---|---|---|---|---|---|
| browser | new tab | details | YourSeq | 661 | 1 | 671 | 671 | 99.5% | chr1 | - | 65778104 | 65778774 | 671 |
| browser | new tab | details | YourSeq | 642 | 19 | 671 | 671 | 99.0% | chrX | + | 58133251 | 58133902 | 652 |
| browser | new tab | details | YourSeq | 640 | 20 | 671 | 671 | 98.7% | chrX | - | 125043544 | 125044192 | 649 |

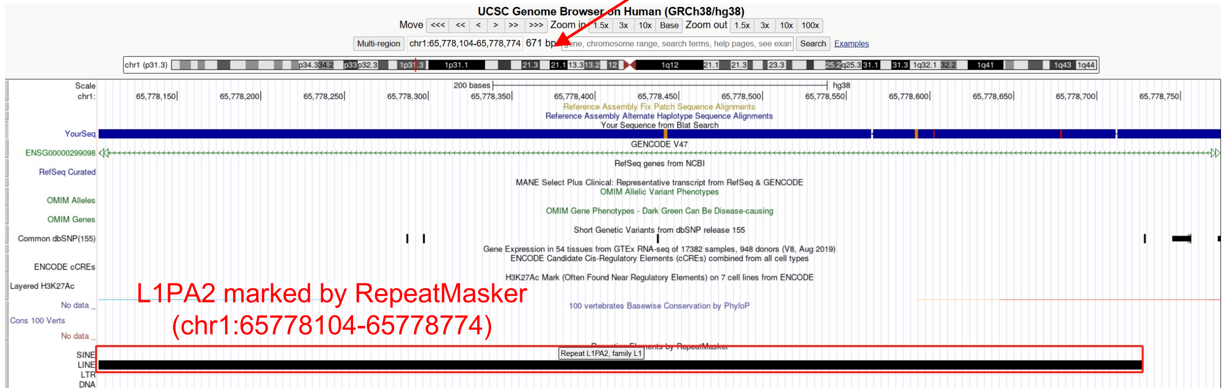


Right-clipped read (18758bp)

- Clip:1-64, repeat sequence

ATATATATATATATATATATATATATATATATATATATATATATATATTTTGTGTGTGT
GTGTGT

- Clip:65-734 aligns to chr1:65778104-65778774, which is annotated as L1PA2 (divergence: 0.7%, active in some populations) by RepeatMasker

- The sequence aligns to L1PA2 reference with 99.5% identity, ACG TAA

TTCTTGACTTCTCCATTCTTTTTTTTTTTCTTTTTTAATTATTATTATTATTATACTTT
AAGTTTTAGGGTACATGTGCACATTGTGCAGGTTACTTACATATGTATACATGTGCCAT
GCTGGTGCGCTGCACCCACTAACGTGTCATCTAGCATTAGGTATATCTCCCAATGCTAT
CCCTCCCCCCCTCCCCCGACCCCACCACAGTCCCAGAGTGTGATATTCCCCTTCCTGTG
TCCATGTGATCTCATTGTTCAATTCCCACCTATGAGTGAGAATATGCGGTGTTTGGTTT
TTTGTTCTTGCGATAGTTTACTGAGAATGATGGTTTCCCAATTTCATCCATGTCCCTAC
AAAGGACATGAACTCATCATTTTTTATGGCTGCATAGTATTCCATGGTGTATATGTGCC
ACATTTTCTTAATCCAGTCTATCATTGTTGGACATTTGGGTTGGTTCCAAGTCTTTGCT
ATTGTGAATAATGCCGCAATAAACATACGTGTGCATGTGTCTTTATAGCAGCATGATTT
ATAGTCATTTGGGTATATACCCAGTAATGGGATGGCTGGGTCAAATGGTATTTCTAGTT
CTAGATCCCTGAGGAATCGCCACACTGACTTCCACAATGGTTGAACTAGTTTACAGTCC
CACCAACAGTGTAAAAGTGTTC

- Clip:735-18758 aligns to chr6:16950726-16968747

- Illustration:

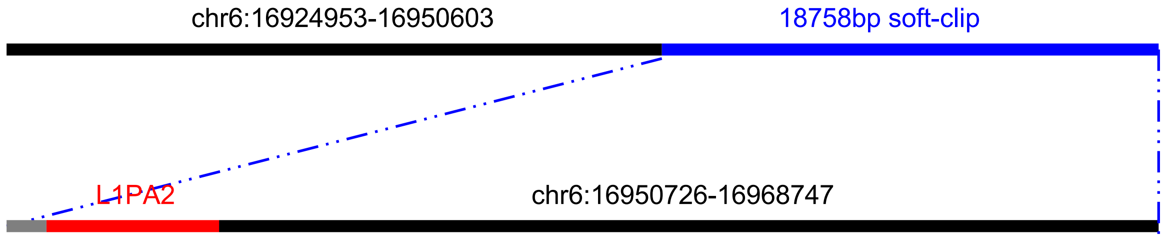

### HG00438 chr9:70162482

➢ Mapping quality is 0, indicating that this region may have alignment issues.

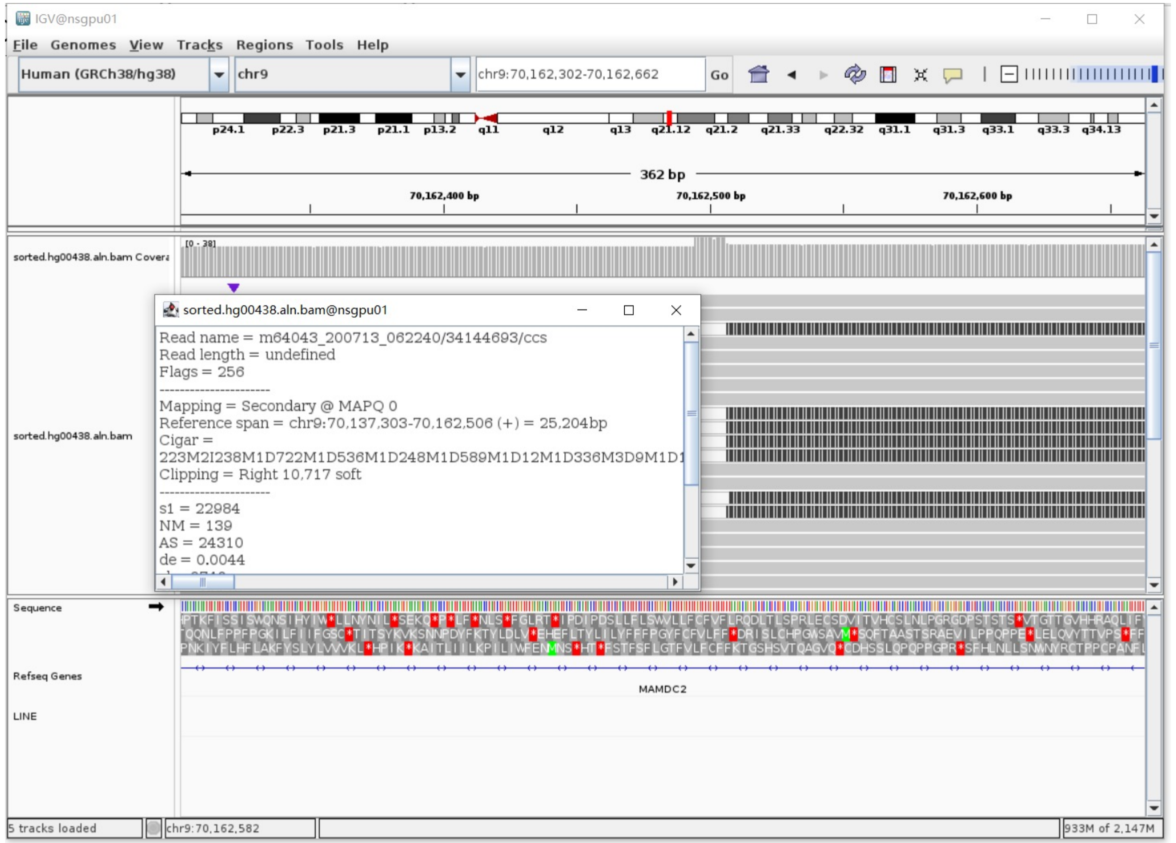


Annotation: For the read in white and black,

- White is the secondary alignment
- Black is the soft clip of the secondary alignment

➢ Fetch one read (m64043_200710_174426/657052/ccs, 32337 bp) and BLAT it to the hg38 genome.

The best alignment is to chr9_GL383541v1_alt (an alternative sequence of chr9) , which contains a L1Hs element.

| ACTIONS | | | QUERY | SCORE | START | END | QSIZE | IDENTITY | CHROM | STRAND | START | END | SPAN |
|---|---|---|---|---|---|---|---|---|---|---|---|---|---|
| browser | new tab | details | YourSeq | 29142 | 1 | 32333 | 32336 | 99.8% | chr9_GL383541v1_alt | - | 133677 | 165994 | 32318 |
| browser | new tab | details | YourSeq | 18978 | 1 | 19154 | 32336 | 99.6% | chr9 | - | 70164895 | 70184026 | 19132 |

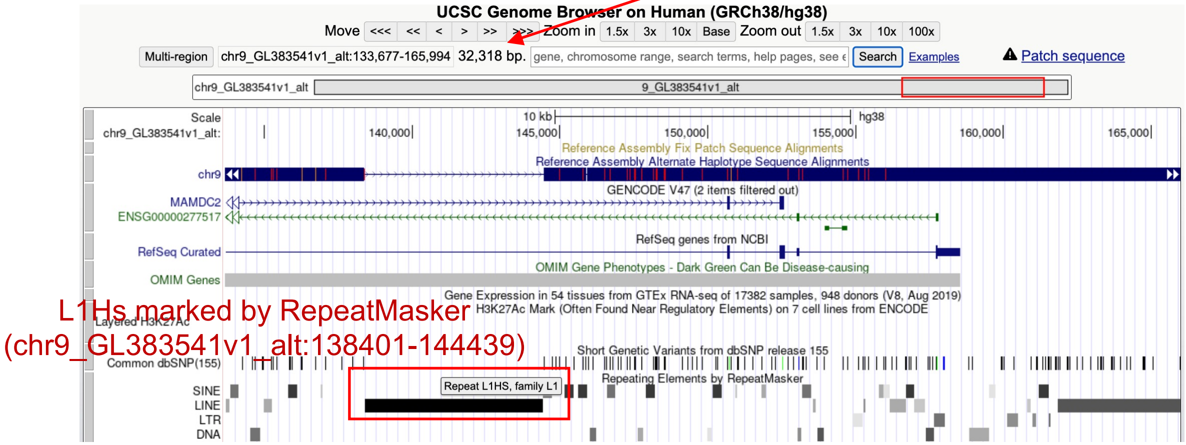


➢ Read:21568-27653 aligns to L1Hs:1-6064, 99.44% identity, ACA TAG

GGGGGGAGGAGCCAAGATGGCCGAATAGGAACAGCTCCGGTCTACAGCTCCCAGCGTGAGCGACGCAG……………A
ATGAGATCACATGGACACAGGAAGGGGAATATCACACTCTGGGGACTGTGGTGGGGTCGGGGGAGGGGGGGAGG
GATAGCATTGGGAGATATACCTAATGCTAGATGACACATTAGTGGGTGCAGCGCACCAGCATGGCACATGTATA
CATATGTAACTAACCTGCACAATGTGCACATGTACCCTAAATCTTAGAGTATAATAAAATAAAAAAAAAAAAAA
AAAAAAGAGCCATGACAAAAAAAAAAAACAAAA

- Illustration (L1Hs in the chromosome chr9_GL383541v1_alt):

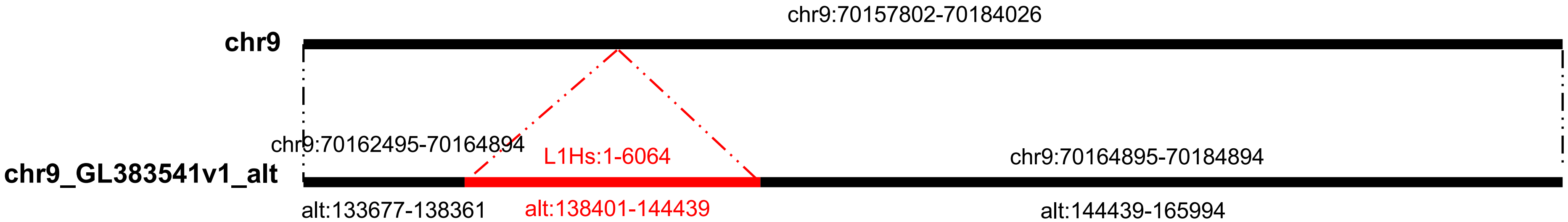

### HG00438 chr18:69865422

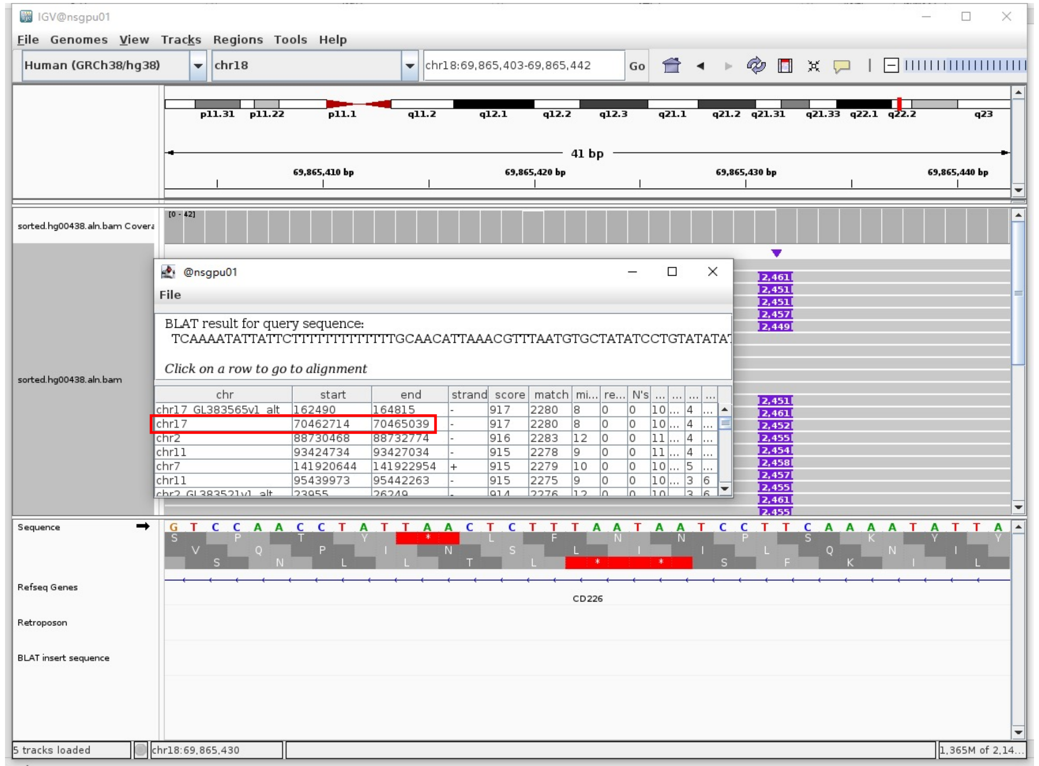


Insertion sequence (2459bp)

- Insertion:1-15, **15bp TSD**

TCAAAATATTATTCT

- Insertion:16-27, PolyT

TTTTTTTTTTTT

- Insertion:28-158 aligns to chr18:282355262-28355407

| QUERY | SCORE | START | END | QSIZE | IDENTITY | CHROM | STRAND | START | END | SPAN |
|---|---|---|---|---|---|---|---|---|---|---|
| YourSeq | 132 | 1 | 132 | 132 | 100.0% | chr18 | + | 28355262 | 28355393 | 132 |

- Inserion:159-2459 aligns to L1Hs:6053-3760, 99.13% identity, ACA TAG

TTGATTCTTTTGCATTTTTCTTTTTTTTTTTTTTATTATACTCTAAGTTTTAGGGTACATGTGCACAT
TGTGCAGGTTAGTTACATATGTATACATGTGCCATGCTGGTGCGCTGCACCCACTAATGTGTCATCT
GAGCATTAGGTATATCTCCCAATGCTATCCCCTCCCCCCTCCCCCGACCCCACCACAGTCCCCAGAG
TGTGATAGTCCCCTTCCTGTGTCCATGTGATCTCATTGTTCAATTCCCACCTATGAGTGAGAATATG
CGGTGTTTTGGTTTTTTGTTCTTGCGATAGTTTACTGAGAATGATGGTTTCCAATTTCATCCATGTC
CCTACAAAGGATATGAACTCATCATTTTTTATGGCTGCATAGTATTCCATGGTGTATATGTGCCACA
TTTTCTTAATCCAGTCTATCATTGTTGGACATTTGGGTTGGTTCCAAGTCTTTGCTATTGTGAATAG
TGCCGCAATAAACATACGTGTGCATGTGTCTTTATAGCAGCATGATTTATACTCATTTGGGTATATA
CCCAGTAATGGGATGCTGGGTCAAATGGTATTTCTAGTTCTAGATCCCTGAGGAATCGCCACACTGA
CTTCCACAATGGTTGAACTAGTTTACAGTCCCACCAACAGTGTAAAAGTGTTCCTATTTCTCCGCAT
.........CACATCCCTTGTAAGTTGGATTCCTAGGTATTTTTATTCTCTTTGAAGCAATTGTGAATGGGAG
TTCACCCATGATTTGGCTCTCTGTTTGTCTGTTGTTGGTGTATAAGAATGCTTGTGATTTTTGTACA
TTGATTTTGTATCCTGAGACTTTGCTGAAGTTGCTTATCAGCTTAAGGAGATTTTGGGCTGAGACGA
TGGGGTTTTCTAGATAAACAATCATGTCGTCTGCAAACAGGGACAATTTGACTTCCTCTTTTCCTAA
TTGAATACCCTTTATTTCCTTCTCCTGCCTGATTGCCCTGGCCAGAACTTCCAACACTATGTTGAAT
AGGAGTGGTGAGAGAGGGCATCCCTGTCTTGTGCCAGTTTTCAAAGGGAATGCTTCCAGTTTTTGCC
CATTCAGTATGATATTGGCTGTGGGTTTGTCATAGATAGCTCTTATTATTTTGAAATACGTCCCATC
AATACCTAATTTATTGAGAGTTTTTAGCATGAAGGGTTGTTGAATTTTGTCAAAGGCTTTTTCTGCA
TCTATTGAGATAATCATGTGGTTTTT

- Illustration (L1 with transduction):

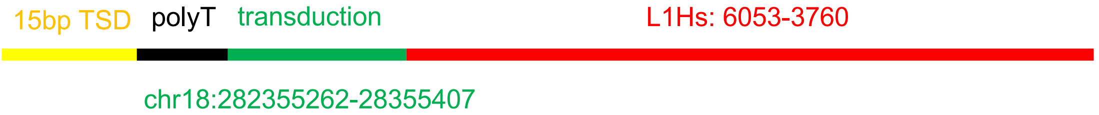

### HG00621 chr5:57983715

- The same genomic coordinate as in HG00438 (page 2)

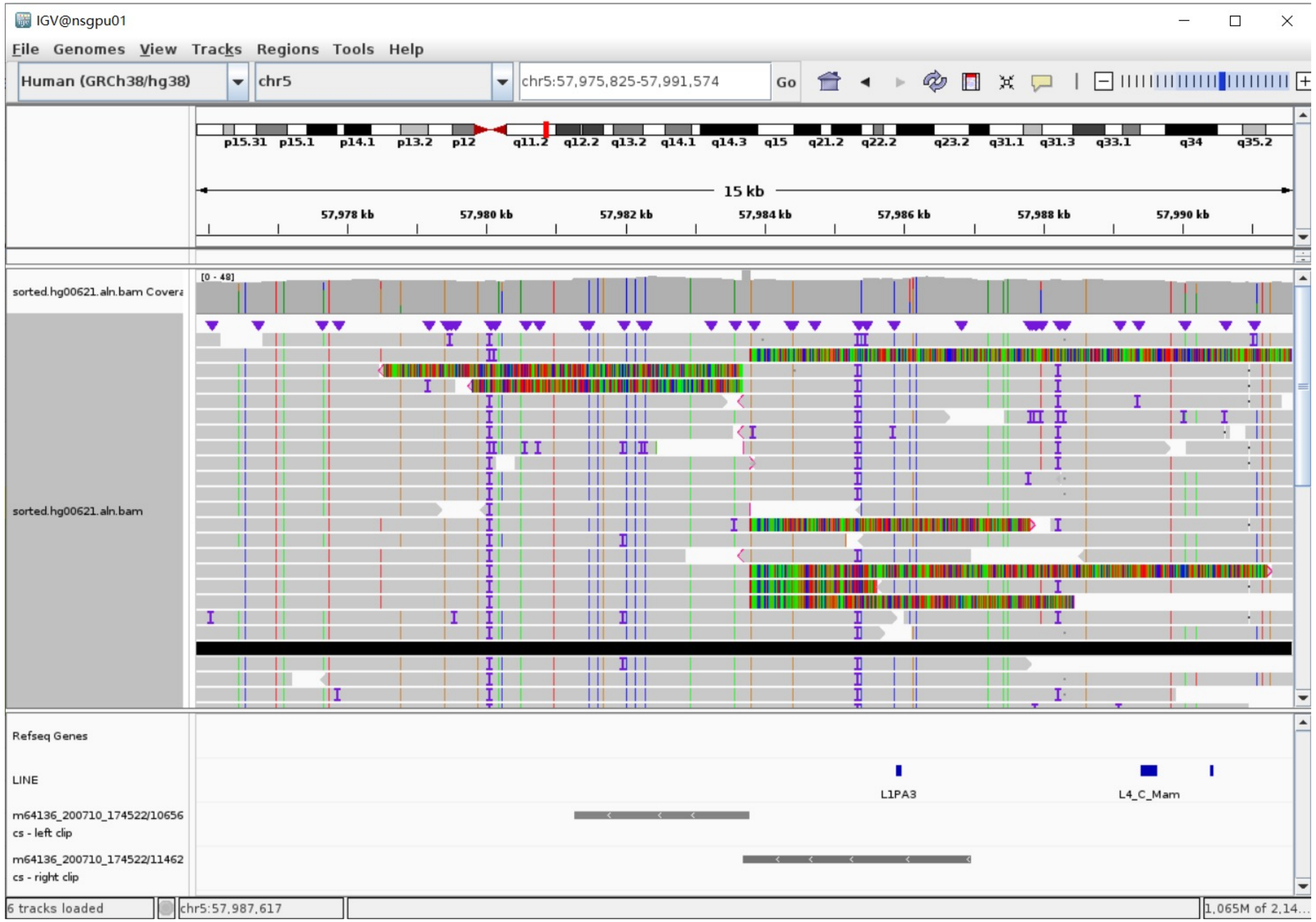

### HG00621 chr6:13190184

- Using the built-in BLAT tool in IGV to check where the inserted sequence aligned.
- The annotation at the aligned location indicated it as L1Hs.

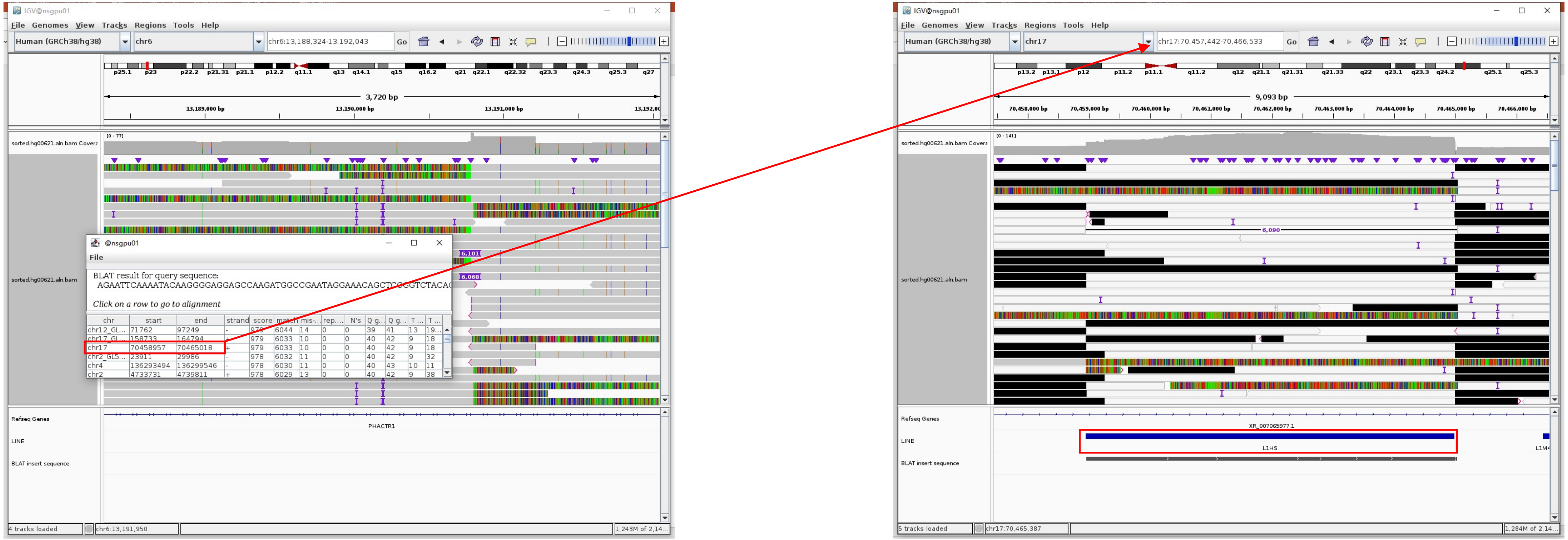


Insertion sequence (6068bp)

- 18bp TSD

AGAATTCAAAATACAAGGGG

- Insertion:16-6060 aligns to L1Hs:2-6051， 99.67% identity， ACA TAG

GGGGAGGAGCCAAGATGGCCGAATAGGAACAGCTCCGGTCTACAGCTCCCAGCGTGAGCGACGCAGAAGACGGTGATTTCTGCATTTCCATCTGAGGTACCGGGTTCATCTCACTAGGGAGTGCCAGACAGTGGGCGCAGGCCAGTGTGTGTG
CGCACCGTGCGCGAGCCGAAGCAGGGCGAGGCATTGCCTCACCTGGGAAGCGCAAGGGGTCAGGGAGTTCCCTTTCCGAGTCAAAGAAAGGGGTGACGGACGCACCTGGAAAATCGGGTCACTCCCACCCGAATATTGCGCTTTTCAGACCGG
CTTAAGAAACGGCGCACCACGAGACTATATCCCACACCTGGCTCGGAGGGTCCTACGCCCACGGAATCTCGCTGATTGCTAGCACAGCAGTCTGAGATCAAACTG............GGGAGAAATTTTTGCAACCTACTCATCTGACAAAGGGCTAATAT
CCAGAATCTACAATGAACTCAAACAAATTTACAAGAAAAAAACAAACAACCCCATCAAAAAGTGGGCGAAGGACATGAACAGACACTTCTCAAAAGAAGACATTTATGCAGCCAAAAAACACATGAAGAAATGCTCATCATCACTGGCCATCA
GAGAAATGCAAATCAAAACCACTATGAGATATCATCTCACACCAGTTAGAATGGCAATCATTAAAAAGTCAGGAAACAACAGGTGCTGGAGAGGATGCGGAGAAATAGGAACACTTTTACACTGTTGGTGGGACTGTAAACTAGTTCAACCAT
TGTGGAAGTCAGTGTGGCGATTCCTCAGGGATCTAGAACTAGAAATACCATTTGACCCAGCCATCCCATTACTGGGTATATACCCAAATGAGTATAAATCATGCTGCTATAAAGACACATGCACACGTATGTTTATTGCGGCACTATTCACAA
TAGCAAAGACTTGGAACCAACCCAAATGTCCAACAATGATAGACTGGATTAAGAAAATGTGGCACATATACACCATGGAATACTATGCAGCCATAAAAAATGATGAGTTCATATCCTTTGTAGGGACATGGATGAAATTGGAAACCATCATTC
TCAGTAAACTATCGCAAGAACAAAAAACCAAACACCGCATATTCTCACTCATAGGTGGGAATTGAACAATGAGATCACATGGACCCAGGAAGGGGAATATCACACTCTGGGGACTGTGGTGGGGTCGGGGGAGGGGGGGAGGGATAGCATTGG
GAGATATACCTAATGCTAGATGACACATTAGTGGGTGCAGTGCACCAGCATGGCACATGTATACATATGTAACTAACCTGCACAATGTGCACATGTACCCTAAAACTTAGAGTATAATAAAAAAAAAAAAAAAAAAAAAAAAAAAAAAA

### HG00621 chr20:17880269

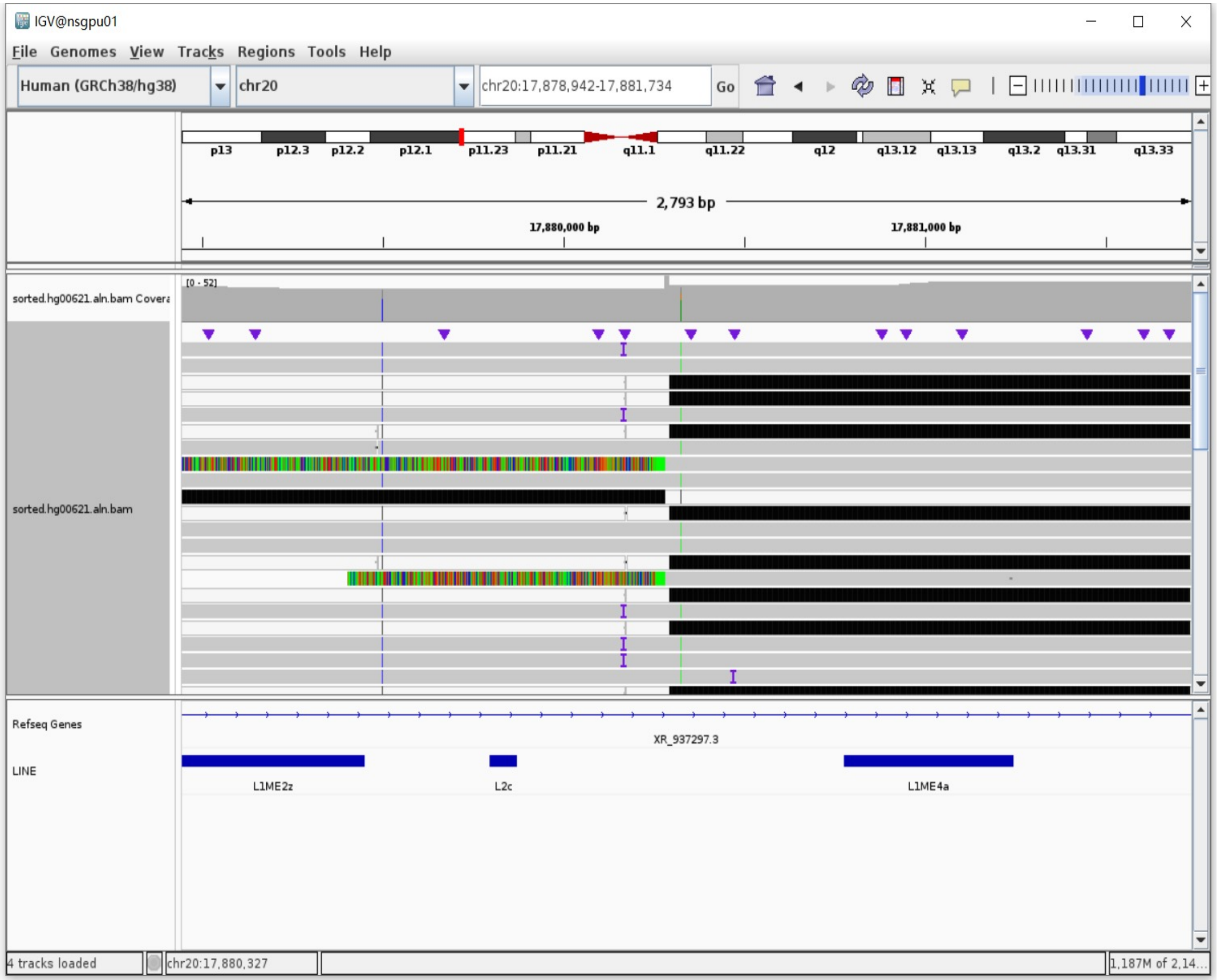


Left-clipped read (5471bp)

- Clip:1-5471 aligns to L1Hs:6045-581, 99.63% identity, ACA TAA

TTTTTTTTTTTTTTTTTTTTTTTTTATTATACTTTAAGTTTTAGGGTACATGTGCACATTGTGCAGG
TTAGTTACATATGTATACATGTGCCATGCTGGTGCGCTGCACCCACTAATGTGTCATCTAGCATTAG
GTATATCTCCCAATGCTATCCCTCCCCCCTCCCCCGACCCCATCACAGTCCCCAGAGTGTGATATTC
CCCTTCCTGTGTCCATGTGATCTCATTGTTCAATTCCCACCTATGAGTGAGAATATGTGGTGTTTGG
TTTTTTGTTCTTGCGATAGTTTACTGAGAATGATGGTTTCCAATTTCATCCATGTCCCTACAAAGGA
TATGAACTCATCATTTTTTATGGCTGCATAGTATTCCATGGTGTATATGTGCCACATTTTCTTAATC
CAGTCTATCATTGTTGGACATTTGGGTTGGTTCCAAGTCTTTGCTATTGTGAATAGTGCCGCAATAA
ACATACGTGTGCGTGTGT............TTCTCCATCCAGCTTTGTTCCGTTGCTGGTGAGGAACTGCGTTCC
TTTGGAGGAGGAGAGGCGCTCTGCGTTTTAGAGTTTCCAGTTTTTCTGTTCTGTTTTTCCCCATCTT
TGTGGTTTTATCTACTTTTGGTCTTTGATGATGGTGATGTACAGATGGGTTTTCGGTGTAGATGTCC
TTTCTGTTTGTTAGTTTTCCTTCTAACAGACAGGACCCTCAGCTGCAGGTCTGTTGGAATACCCTGC
CGTGTGAGGTGTCAGTGTGCCCCTGCTGGGGGGTGCCTCCCAGTTAGGCTGCTCGGGGGTCAGGGGT
CAGGGACCCACTTGAGGAGGCAGTCTGCCCGTTCTCAGATCTCCAGCTGCGTGCTGGGAGAACCACT
GCTCTCTTCAAAGCTGTCAGACAGGGACACTTAAGTCTGCAGAGGTTACTGCTGT

### HG02630 chr2:141016817

- Using the built-in BLAT tool in IGV to check where the inserted sequence aligned.
- The annotation at the aligned location indicated it as L1Hs.

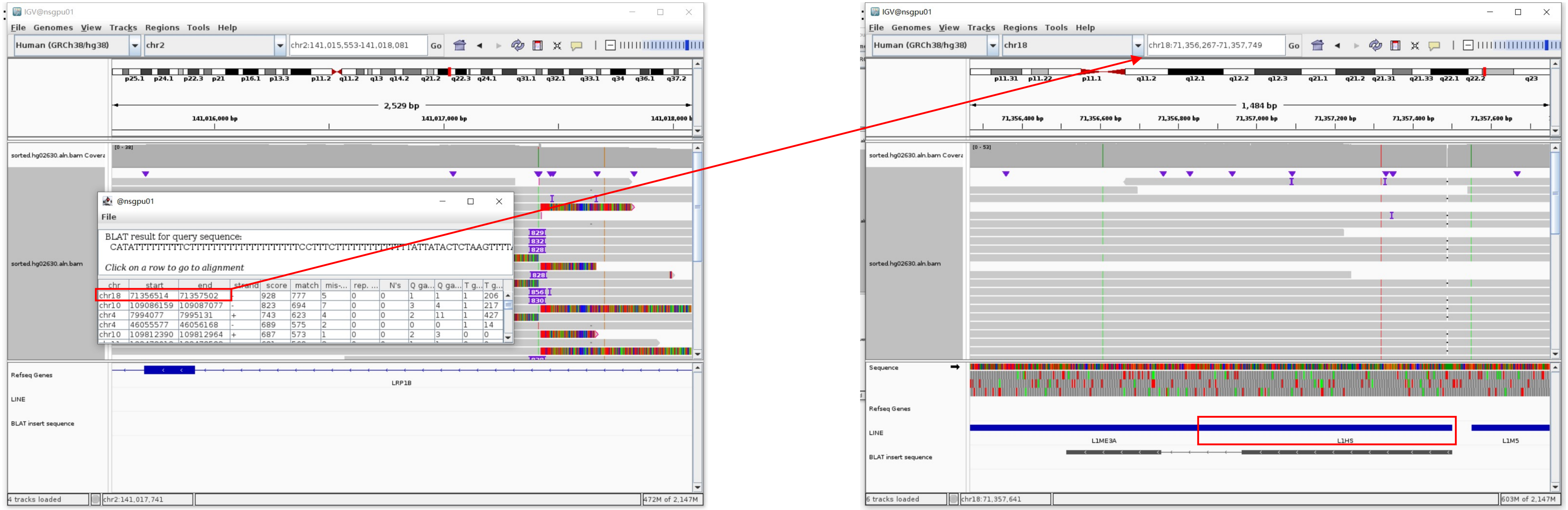


Insertion sequence (828bp)

- 13bp TSD

CATATTTTTTTTT

- Insertion:8-580 aligns to L1Hs:6064-5494, 98.77% identity, ACA TAG

TTTTTCTTTTTTTTTTTTTTTTTTTCCTTTCTTTTTTTTTTTTTTTTATTATACTCTAAGTTTTAGGGTACATGTGCACATTGTGCAGGTTAGTTACATATGTATACATGTGCCATGCTGGTGCACTGCACCCACTAATGTGTCATCT
AGCATTAGGTATATCTCCCAATGCTATCCCTCCCCCCTCCCCCCGACCCCACCACAGTCCCCAGAGTGTGATATTCCCCTTCCTGTGTCCATGTGATCTCATTGTTCAATTCCCACCTATGAGTGAGATTATGCGGTGTTTGGTTT
TTTGTTCTTGCGATAGTTTACTGAGAATGATGGTTTCCAATTTCATCCATGTCCCTACAAAGGATATGAACTCATCATTTTTTATGGCTGCATAGTATTCCATGGTGTATATGTGCCACATTTTCTTAATCCAGTCTATCATTGTT
GGACATTTGGGTTGGTTCCAAGTCTTTGCTATTGTGAATAGTGCCGCAATAAACATACGTGTGCATGTGTCTTTATAGCAGCATGATTTATAGTCCTTTGGGTATATACCCAGTAATGGATGGCTGGGTCAAA

- Insertion:577-828 aligns to L1Hs:5254-5505, 99.60% identity, 5’ inversion

AAACACATGAAGAAATGCTCATCATCACTGGCCATCAGAGAAATGCAAATCAAAACCACTATGAGATATCATCTCACACCAGTTAGAATGGCAATCATTAAAAAGTCAGGAAACAACAGGTGCTGGAGAGGATGTGGAGAAATAGG
AACACTTTTACACTGTTGGTGGGACTGTAAACTAGTTCAACCATTGTGGAAGTCAGAGTGGCGATTCCTCAGGGATCTAGAACTAGAAATACCATTTGACCCAGC

### HG02630 chr7:106327425

- Using the built-in BLAT tool in IGV to check where the inserted sequence aligned.

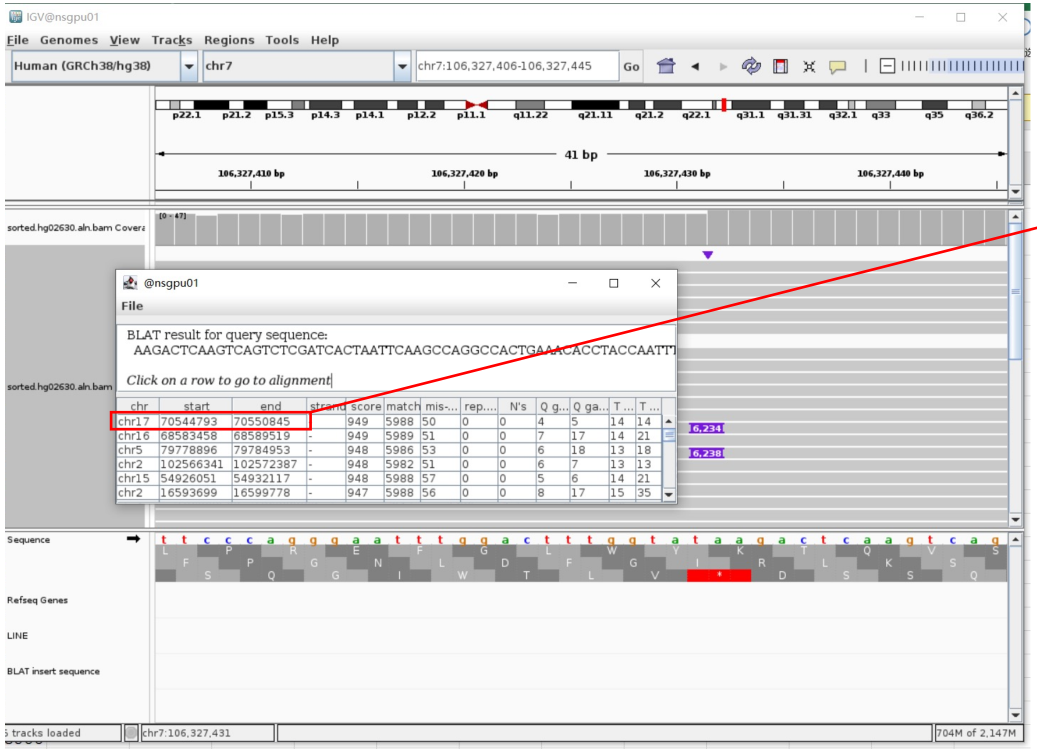


- The annotation at the aligned location indicated it as L1Hs.

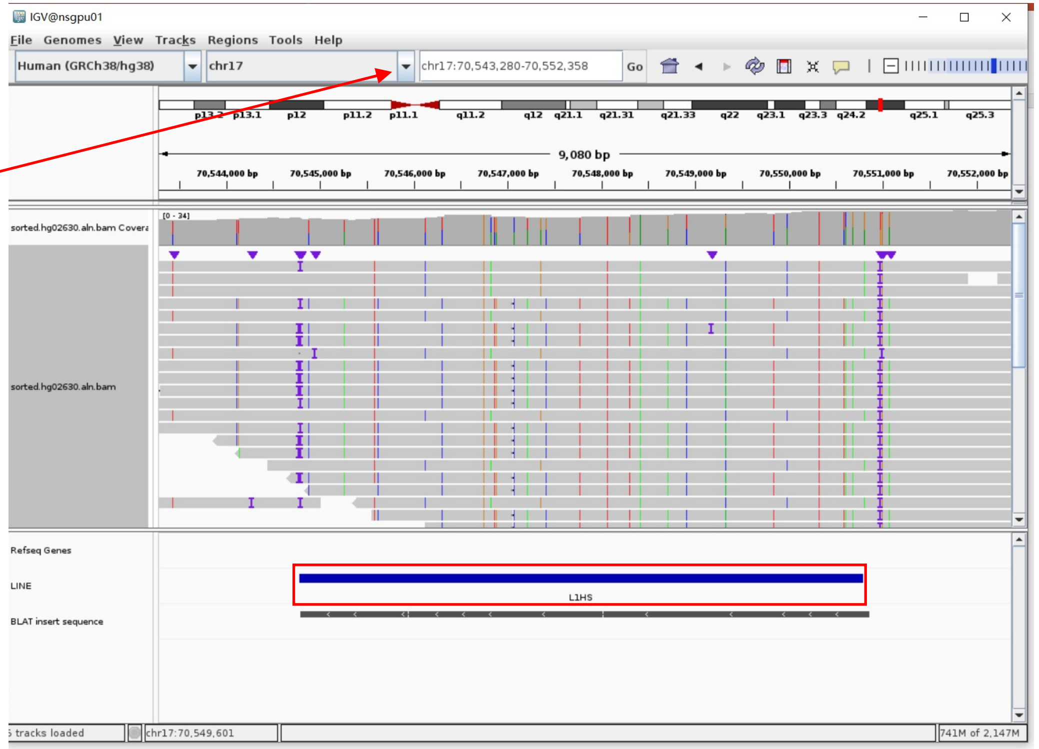


Insertion sequence (6234bp)

- 18bp TSD

AAGACTCAAGTCAGTCTC

- Insertion:191-6234 aligns to L1Hs:0-6051, 99.29% identity, ACA TAG

GAGAGGAGGAGCCAAGATGGCCGAATAGGAACAGCTCCGGTCTACAGCTCCCAGCGTGAGCGACGCAGAAGACGGGTGATTTCTGCATTTCCATCTGAGGTACCGGGTTCATCTCACTAGGAGTGCCAGACAGTGGGCGCAGGC
CAGTGGGTGCGCGCACCGTGCGTGAGCCGAAGCAGGGCGAGGCATTGCCTCACCTGGGAAGCGCAAGGGGTCAGGGAGTTCCCTTTCCGAGTCAAAGAAAAGGGGTGACGGACGCACCTGGAAAATCGGGTCACTCCCACCCGA
ATATTGCGCTTTTCAGACCGGCTTAAGAAACGGCGCACCACGAGACTATATCCCACACCTGGCTCGGAGGGTCCTACGCCCACGGAATCTCGCTGATTGCTAGCAC.........TGTCAACTAGTTCAACCATTGTGGAAGTCAGTGTG
GCGATTCCTCAGGGATCTAGAACTAGAAATACCATTTGACCCAGCCATCCCATTACTGGGTATATACCCAAAGGACTATAAATCATGCTGCTATAAAGACACATGCACACGTATGTTTATTGCGGCACTATTCACAATAGCAAA
GACTTGGAACCAACCCAAATGTCCAACAATGATAGACTGGATTAAGAAAATGTGGCACATATACACCATGGAATACTATGCAGCCATAAAAAACGATGAGTTCATGTCCTTTGTAGGGACATGGATGAAATTGGAAACCATCAT
TCTCAGTAAACTATCGCAAGAACAAAAAACCAAACACCGCATATTCTCACTCATAGGTGGGAATTGAACAATGAGATCACTTGGACACAGGAAGGGGAATATCACACTCTGGGGACTGTGGTGGGGTCGGGAGAGTGGGGAGGG
ATAGCATTGGGAGATATACCTAATGGTAGATGACACGTTAGTGGGTGCAGCGCACCAGCATGGCACATGTATACATATGTAACTAACCTGCACAATGCGCACATGTACCCTAAAACTTAGAGTATAATAAAAAAAAAAAAAATTT
AAAAAAAAAAAAAAA

### HG02630 chr12:63175375

➢ Using the built-in BLAT tool in IGV to check where the inserted sequence aligned.

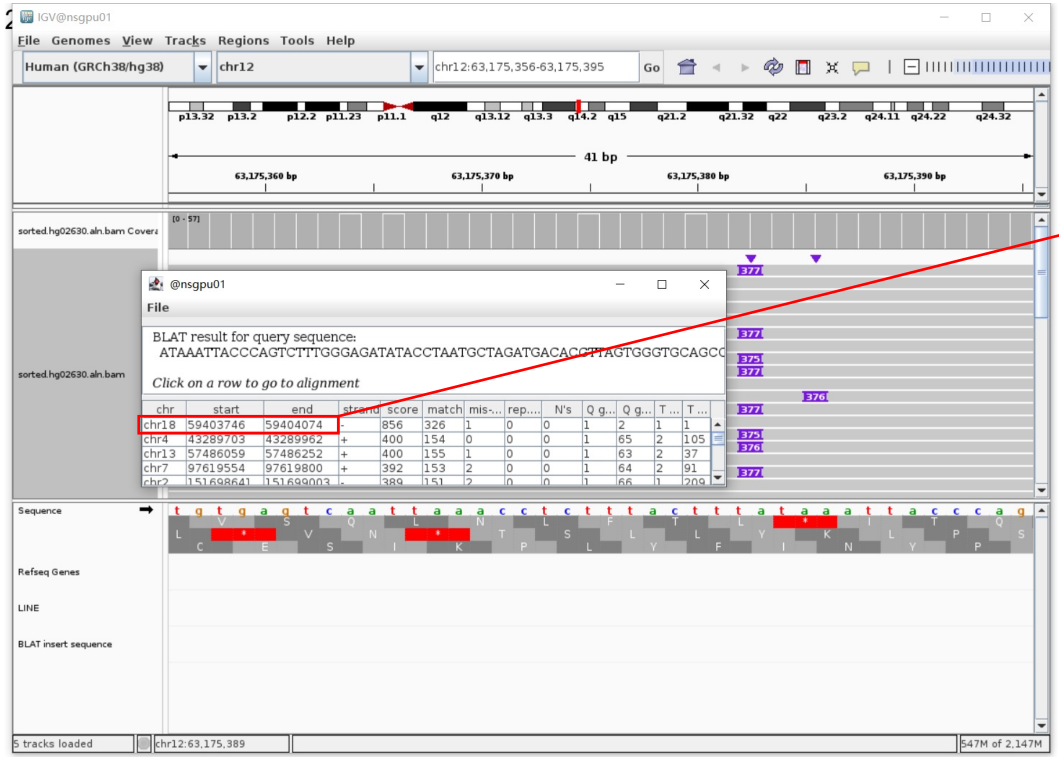


➢ The annotation at the aligned location indicated it as L1Hs.

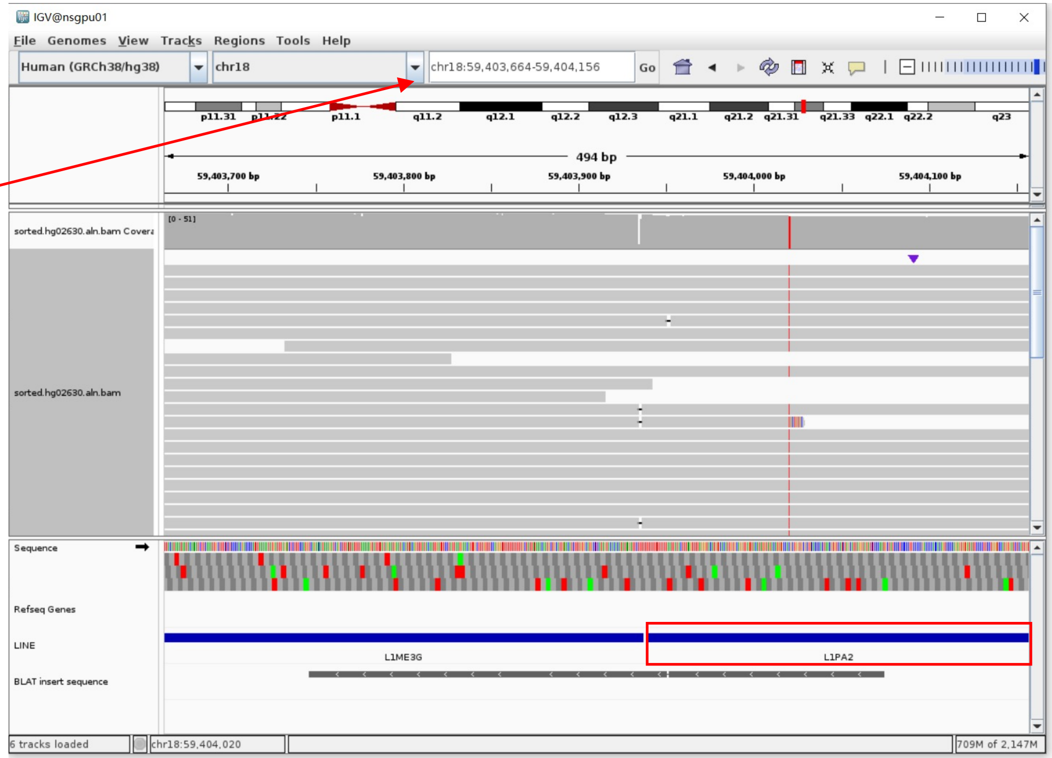


**Name:** L1PA2 (link requires registration)
**Family:** L1
**Class:** LINE
**SW Score:** 29325
**Divergence:** 0.8%
**Deletions:** 0.0%
**Insertions:** 0.0%
**Begin in repeat:** 125
**End in repeat:** 6155
**Left in repeat:** 0
**Position:** chr18:59403940-59409970
**Band:** 18q21.32
**Genomic Size:** 6031
**Strand:** -
View DNA for this feature (hg38/Human)

Insertion sequence (375bp)

➢ 17bp TSD

ATAAATTACCCAGTCTT

➢ Insertion:14-158 aligns to L1PA2 (divergence 0.8%, active in some populations), ACG TAA

CTTTGGGAGATATACCTAATGCTAGATGACACGTTAGTGGGTGCAGCGCACCAGCATGGCACATGTATACATATGTAACTAACCTGC
ACAATGTGCACATGTACCCTAAAACTTAAAGTATAATTAAAAAAAAAAAAAAAAAGAA

### HG02630 chr16:15686361

- Using the built-in BLAT tool in IGV to check where the inserted sequence aligned.

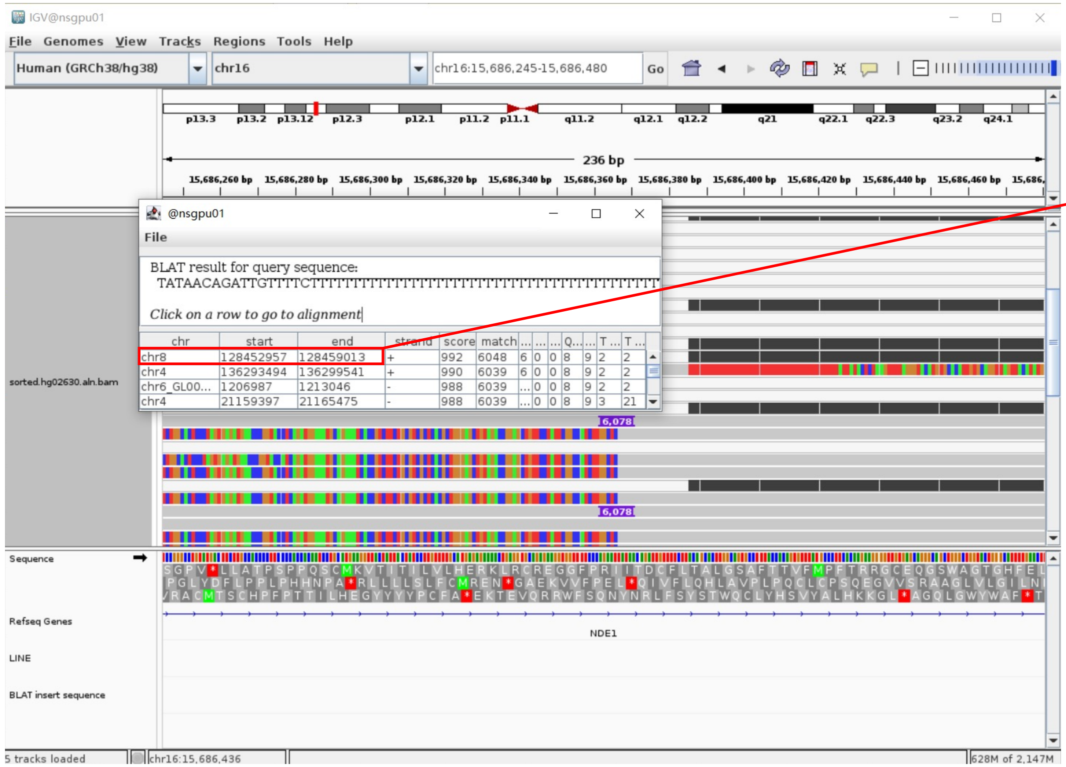


| chr | start | end | strand | score | match |
|---|---|---|---|---|---|
| chr8 | 128452957 | 128459013 | + | 992 | 6048 |
| chr4 | 136293494 | 136299541 | + | 990 | 6039 |
| chr6_GL00... | 1206987 | 1213046 | - | 988 | 6039 |
| chr4 | 21159397 | 21165475 | - | 988 | 6039 |

- The annotation at the aligned location indicated it as L1Hs.

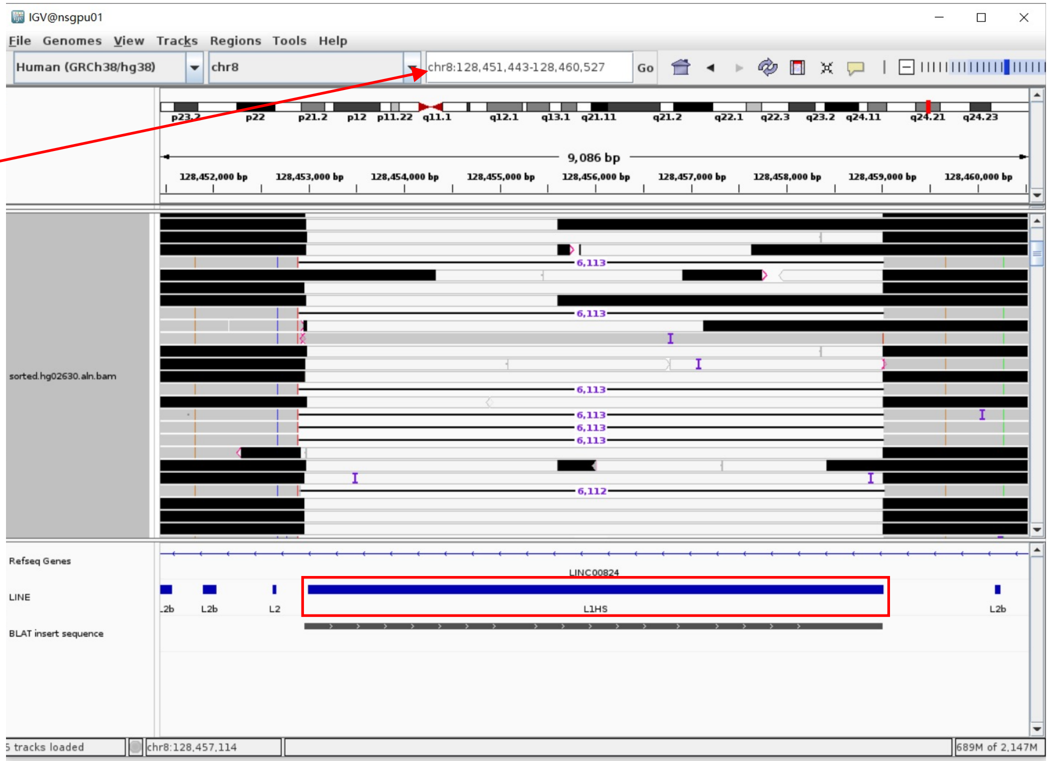


Insertion sequence (6078bp)

- 19bp TSD

TATAACAGATTGTTTTCTT

- Insertion:17-6078 aligns to L1Hs:6063-8, 99.60% identity, ACA TAG

TTTTTTTTTTTTTTTTTTTTTTTTTTTTTTTTTTTTTTTTTTTTATTATACTCTAAGTTTTAGGGTACATGTGCACATTGTGCAGGTTAGTTACATATGTATACATGTGCCATGCTGGTGCGCTGCACCCACTAATGTGTCA
TCTAGCATTAGGTATATCTCCCAATGCTATCCCTCCCCCCTCCCCCGACCCCACCACAGTCCCCAGAGTGTGATATTCCCCTTCCTGTGTCCATGTGATCTCATTGTTCAATTCCCACCTATGAGTGAGAATATGCGGTGT
TTGGTTTTTTGTTCTTGCGATAGTTTACTGAGAATGATGGTTTCCAATTTCATCCATGTCCCTACAAAGGATATGAACTCATCATTTTTTATGGCTGCATAGTATTCCATGGTGTATATGTG……………ATCTCAGACTGCTG
TGCTAGCAATCAGCGAGATTCCGTGGGGCGTAGGACCCTCCGAGCCAGGTGTGTGGGATATAGTCTCATGGTGCGCCGTTTCTTAAGCCGGTCTGAAAAGCGCAATATTCGGGTGGGAGTGACCCGATTTTTCCAGGTGCG
TCCGTCACCCCTTTCTTTGACTCGGAAAGGGAACTCCCTGACCCCTTGCGCTTCCCAGGTGAGGCAATGCCTCGCCCTGCTTCGGCTCGCGCACGGTGCGCACACACACTGGCCTGCGCCCACTGTCTGGCACTCCCTAGT
GAGATGAACCCGGTACCTCAGATGGAAATGCAGAAATCACCGTCTTCTGCGTCGCTCACGCTGGGAGCTGTAGACCGGAGCTGTTCCTATTCGGCCATCTTGGCTC

### HG03492 chr7:106327425

- ➢ The same genomic coordinate as in HG02630 (page 11)

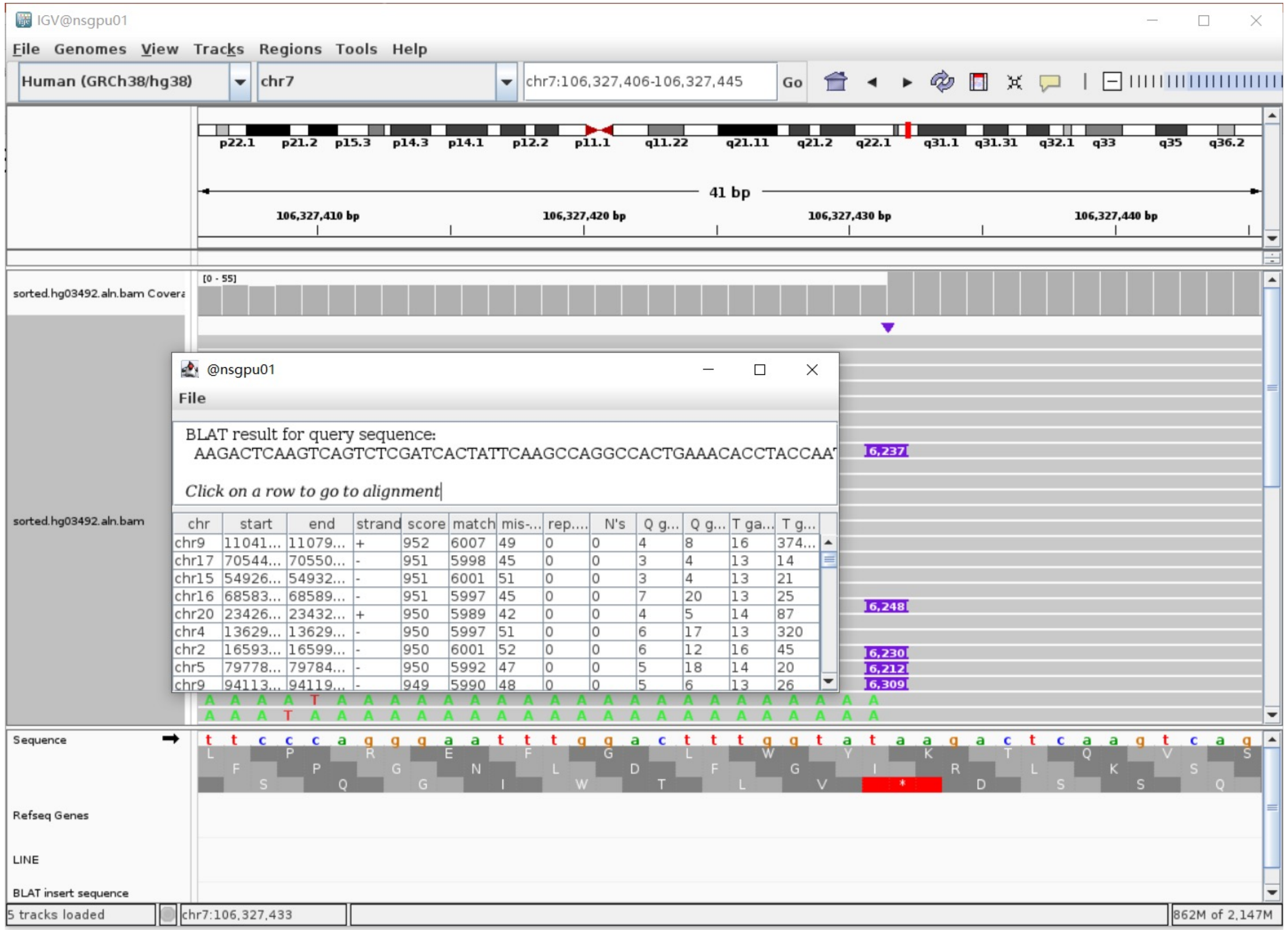

### HG03516 chr3:103981000

- ➢ Using the built-in BLAT tool in IGV to check where the inserted sequence aligned.
- ➢ The annotation at the aligned location indicated it as L1Hs.

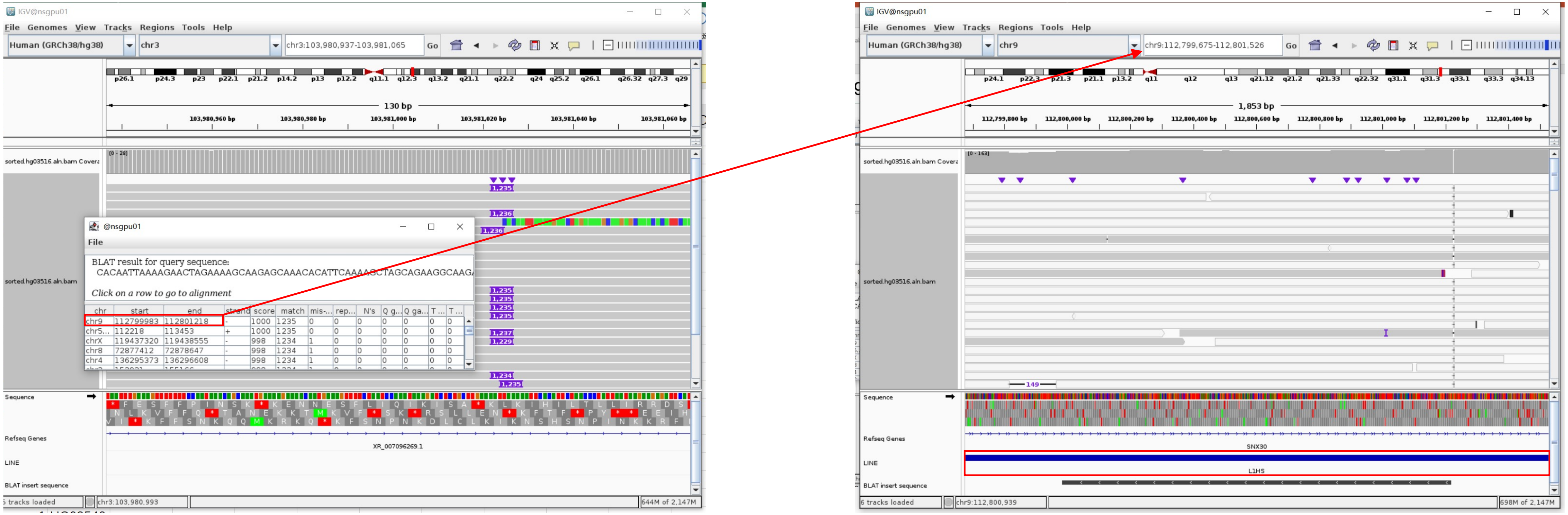

Insertion sequence (1235bp)

- ➢ Insertion:1-1235 aligns to L1Hs:2942-4177, 99.84% identity

CACAATTAAAAGAACTAGAAAAGCAAGAGCAAACACATTCAAAAGCTAGCAGAAGGCAAGAAATAACTAAAATCAGAGCAGAACTGAAGGAAATAGAGACACAAAAAACCCTTCAAAAAATCAATGAATCCAGGAGCTGGTTT
TTTGAAAGGATCAACAAAATTGATAGACCGCTAGCAAGACTAATAAAGAAAAAAAGAGAGAAGAATCAAATAGACACAATAAAAAATGATAAAGGGGATATCACCACCGATCCCACAGAAATACAAACTACCATCAGAGAATA
CTACAAACACCTCTACGCAAATAAACTAGAAAATCTAGAAGAAATGGATACATTCCTCGACACATACACTCTCCCAAGACTAAACCAGGAAGAAGTTGAATCTCTGAATAGACCAATAACAGGCTCTGAAATTGTGGCAATAA
TCAATAGTTTACCAACCAAAAAGAGTCCAGGACCAGATGGATTCACAGCCGAATTCTACCAGAGGTACATGGAGGAACTGGTACCATTCCTTCTGAAACTATTCCAATCAATAGAAAAAGAGGGAATCCTCCCTAACTCATTT
TATGAGGCCAGCATCATTCTGATACCAAAGCCGGGCAGAGACACAACCAAAAAAGAGAATTTTAGACCAATATCCTTGATGAACATTGATGCAAAAATCCTCAATAAAATACTGGCAAACCGAATCCAGCAGCACATCAAAAA
GCTTATCCACCATGATCAAGTGGGCTTCATCCCTGGGATGCAAGGCTGGTTCAATATACGCAAATCAATAAATGTAATCCAGCATATAAACAGAGCCAAAGACAAAAACCACATGATTATCTCAATAGATGCAGAAAAAGCCT
TTGACAAAATTCAACAACCCTTCATGCTAAAAACTCTCAATAAATTAGGTATTGATGGGACGTATTTCAAAATAATAAGAGCTATCTATGACAAACCCACAGCCAATATCATACTGAATGGGCAAAAACTGGAAGCATTCCCT
TTGAAAACCGGCACAAGACAGGGATGCCCTCTCTCACCGCTCCTATTCAACATAGTGTTGGAAGTTCTGGCCAGGGCAATCAGGCAGGAGAAGGAAATAAAGGGTATTCAATTAGGAAAAGAGGAAGTCAAATTGTCCCTGTT
TGCAGACGACATGATTGTTTATCTAGAAAACCCCATCGTCTCAGCCCAAAATCTCCTTAAGCTGATAAGCAACTTCAGCAAAGTCTCAGGA

### HG03516 chr5:57983715

- The same genomic coordinate as in HG00438 (page 2)

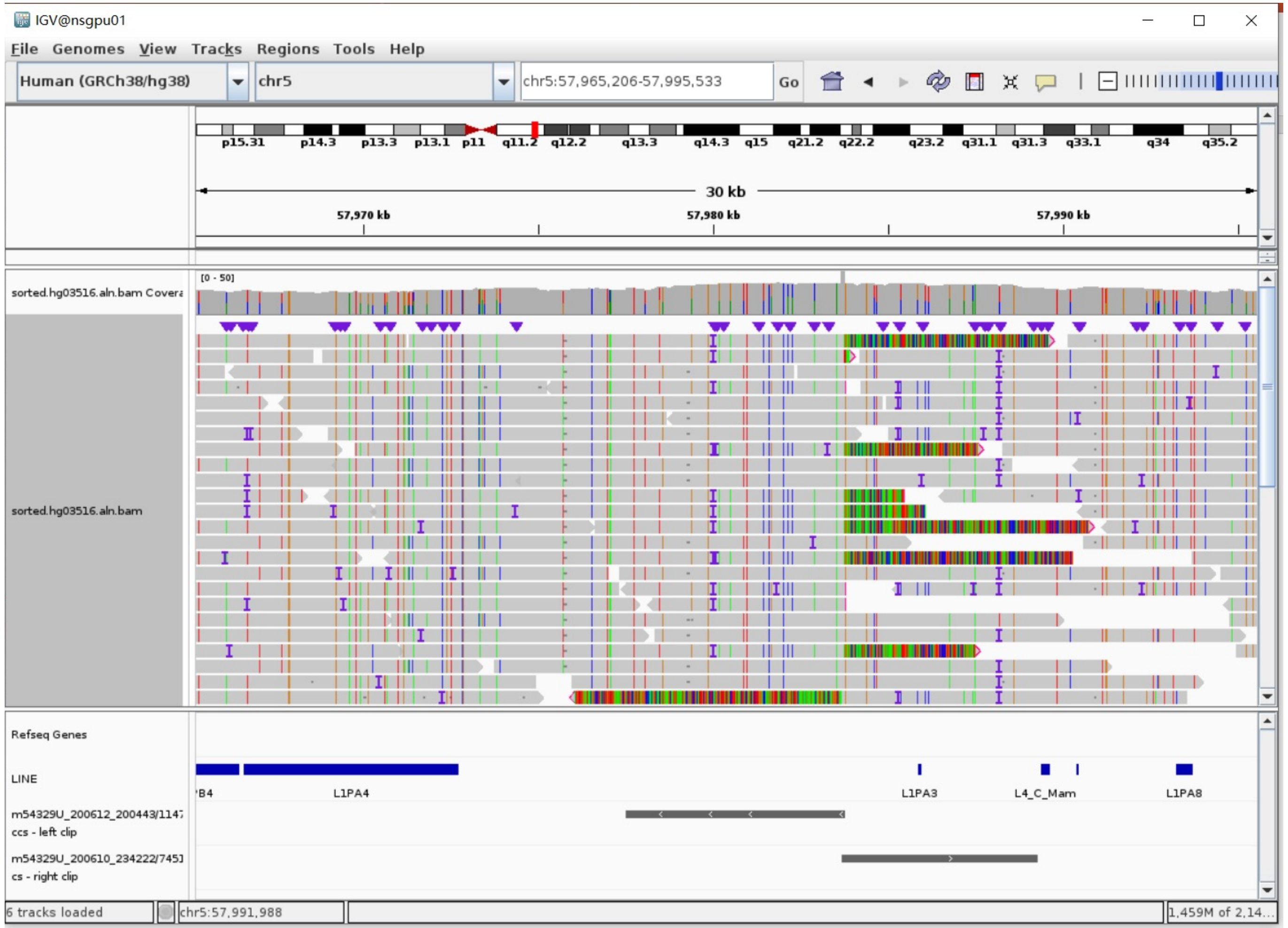

### HG03516 chr6:103423568

- ➢ Using the built-in BLAT tool in IGV to check where the inserted sequence aligned.

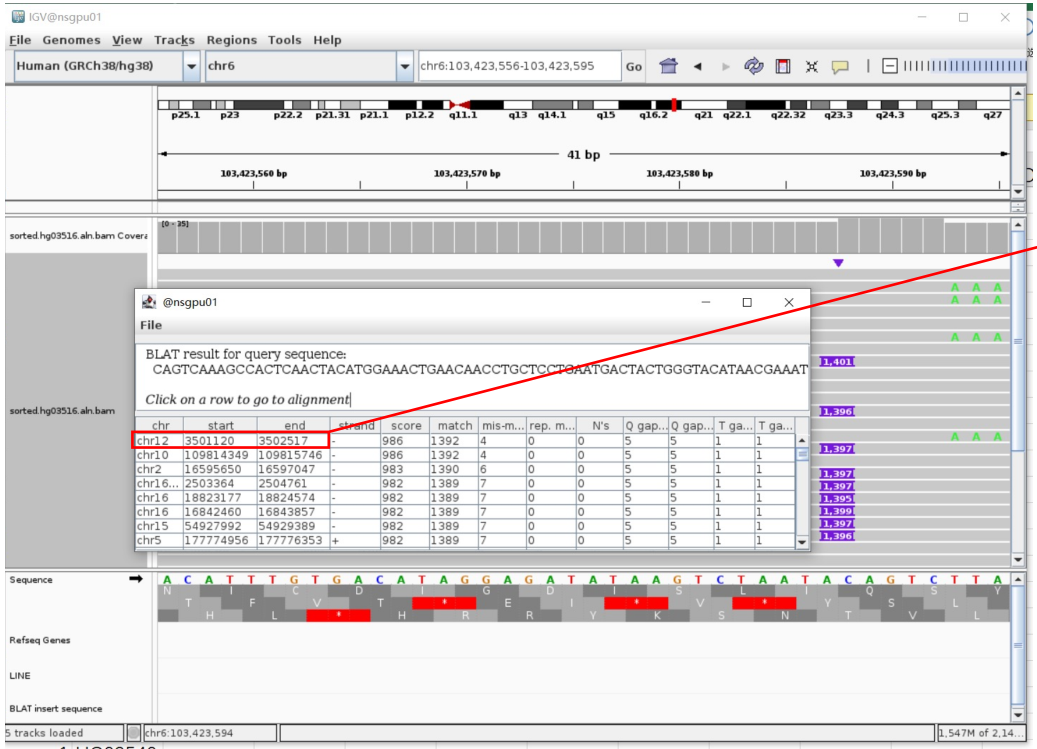


- ➢ The annotation at the aligned location indicated it as L1Hs.

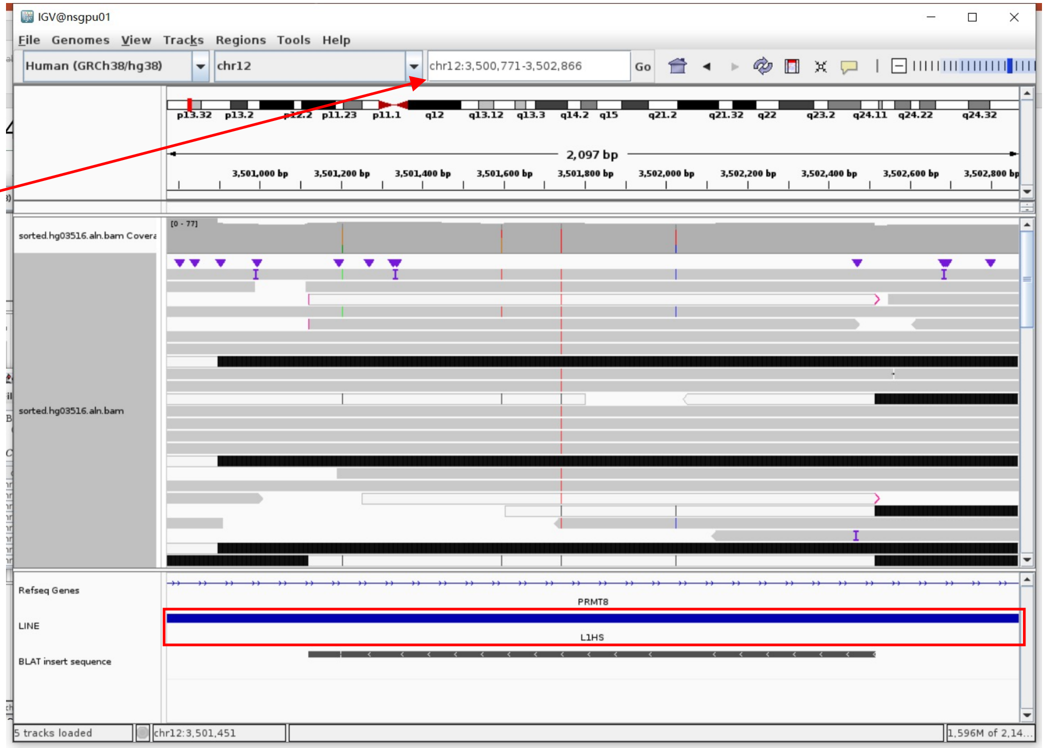


Insertion sequence (1395bp)

- ➢ 5bp TSD

CAGTC

- ➢ Insertion:0-1235 aligns to L1Hs:2712-4109, 99.50% identity

CAGTCAAAGCCACTCAACTACATGGAAACTGAACAACCTGCTCCTGAATGACTACTGGGTACATAACGAAATGAAGGCAGAAATAAAGATGTTCTTTGAAACCAACGAGAACAAAGACACCACATACCAGAATCTCTGGGACGCATTCAAA
GCAGTGTGTAGAGGGAAATTTATAGCACTAAATGCCTACAAGAGAAAGCAGGAAAGATCCAAAATTGACAGCCTAACATCACAATTAAAAGATCTAGAAAAGCAAGAGCAAACACATTCAAAAGCTAGCAGAAGGCAAGAAATAACTAAAA
TCAGAGCAGAACTGAAGGAAATAGAGACACAAAAAACCCTTCAAAAAATCAATGAATCCAGGAGCTGGTTTTTTGAAAGGATCAACAAAATTGATAGACCACTAGCAAGACTAATAAAGAAAAAAAGAGAGAAGAATCAAATAGACACAAT
AAAAAATGATAAAGGGGATATCACCACCGATCCCACAGAAATACAAACTACCATCAGGAGAATACTACAAACACCTCTACGCAAATAAACTAGAAAATCTAGAAGAAATGGATACATTCCTCGACACATACACTCTCCCAAGACTAAACC
AGGAAGAAGTTGAATCTCTGAATGAGACCAATAACAGGCTCTGAAATTGTGGCAATAATCAATAGTTTACCAACCAAAAAGAGTCCAGGACCAGATGGATTCACAGCCGAATTCTACCAGAGGTACAAGGAGGAACTGGTACCATTCCTTC
TGAAACTATTCCAATCAATAGAAAAAGAGGGAATCCTCCCTAACTCATTTTATGAGGCCAGCATCATTCTGACACCAAAGCCGGGCAGAGACACAACCAAAAAAGAGAATTTTAGACCAATATCCTTGATGAACATTGATGCAAAAATCCT
CAATAAAATACTGGCAAACCGAATCCAGCAGCACATCAAAAAGCTTATCCACCATGATCAAGTGGGCTTCATCCCTGGGATGCAAGGCTGGTTCAATATACGCAAATCAATAAATGTAATCCAGCATATAAACAGAGCCAAAGACAAAAAC
CACATGATTATCTCAATAGATGCAGAAAAAGCCTTTGACAAAATTCAACAACCCTTCATGCTAAAAACTCTCAATAAATTAGGTATTGATGGGACGTATTTCAAAATAATAAGAGCTATCTATGACAAACCCACAGCCAATATCATACTGA
ATGGGCAAAAACTGGAAGCATTACCTTTGAAAAACTGGCACAAGACAGGGATGCCCTCTCTCACCGCTCCTATTCAACATAGTGTTGGAAGTTCTGGCCAGGGCAATCAGGCAGAGAAGGAAATAAAGGGTATTCAATTAGGAAAAGAGGA
AGTCAAATTGTCCCCTGTTTGCAGACGACATGATTGTTTATC

### HG03516 chr7:106327425

- The same genomic coordinate as in HG02630 (page 11)

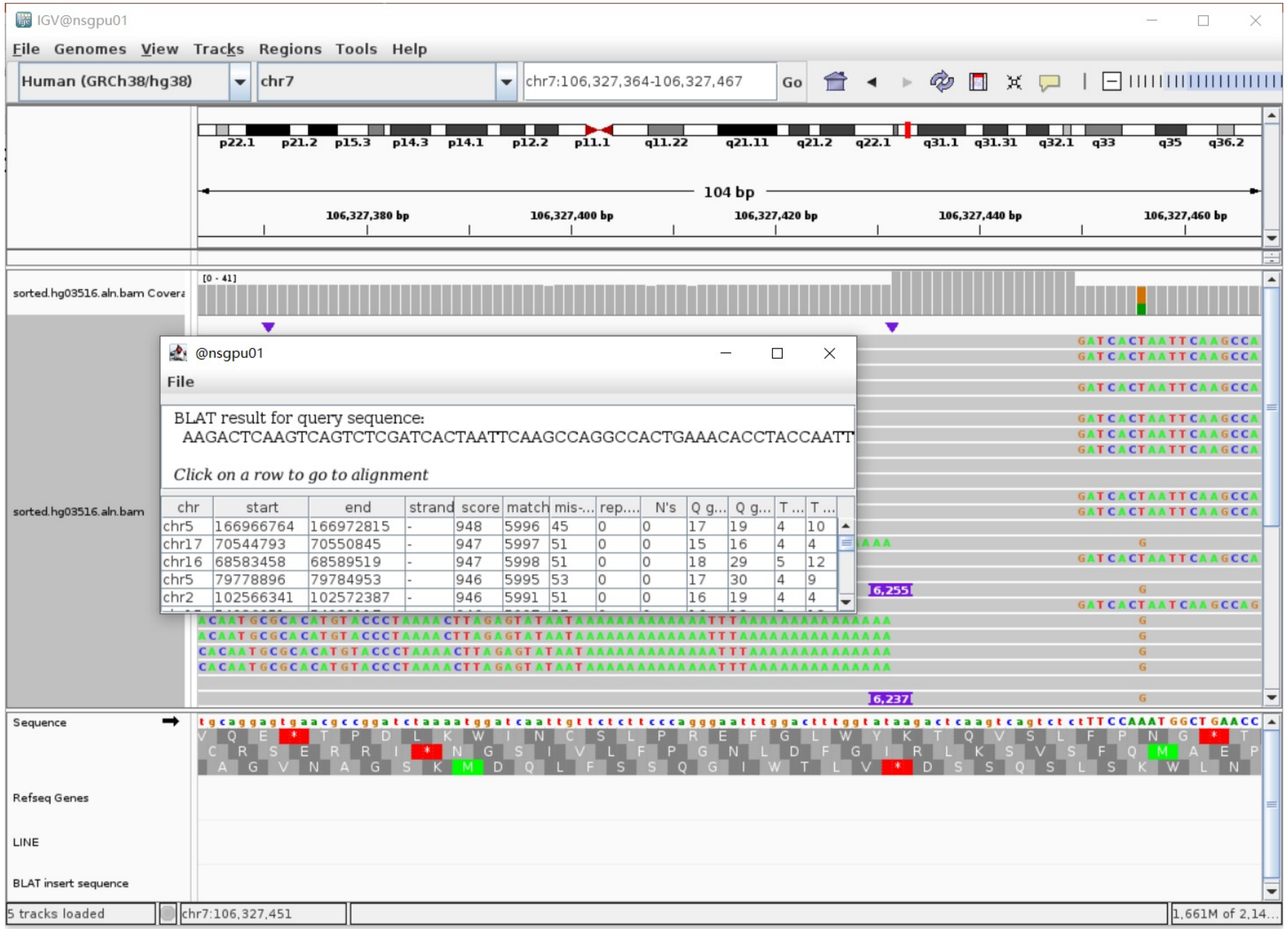

### HG03516 chr9:118632302

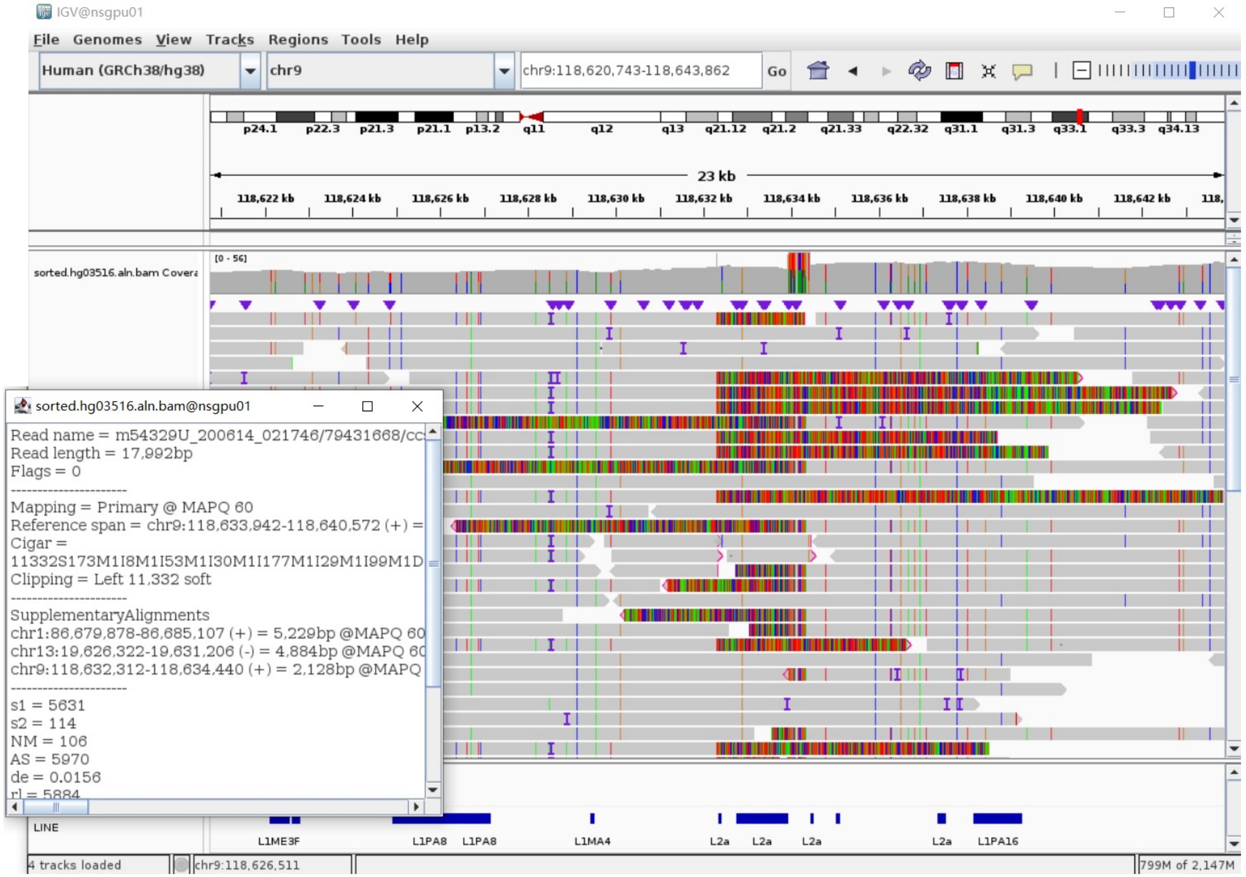


Right-clipped read

- Clip:1-6012 aligns to L1Hs:6042-7, 99.50% identity, ACA TAG

TTTTTTTTTTTTTTTTTTTTTTATTATACTCTAAGTTTTAGGGTACATGTGCACATTGTGCAGGTTAGT
TACATATGTATACATGTGCCATGCTGGTGCGCTGCACCCAACTAATGTGTCATCTAGCATTAGGTATAT
CTCCCAATGCTATCCCTCCCCCTCCCCCGACCCCACCACAGTCCCCAGAGTGTGATATTTCCCTTCCTG
TGTCCATGTGATCTCATTGTTCAATTCCCACCTATGAGTGAGAATATGCGGTGTTTGGTTTTTTGTTCT
TGCGATAGTTTACTGAGAATGATGGTTTCCAATTTCATCCATGTCCCTACAAAGGATATGAACTCATCA
TTT...........CACCAGTTCGAGCTTCCCGGCTGCTTTGTTTACCTAAGCAAGCCTGGGCAATGGCGGGCGCC
CCTCCCCCAGCCTCGTTGCCGCCTTGCAGTTTGATCTCAGACTGCTGTGCTAGCAATCAGCGAGATTCC
GTGGGCGTAGGACCCTCTGAGCCAGGTGTGGGATATAGTCTCATGGTGCGCCGTTTCTTAAGCCGGTCT
GAAAAGCGCAATATTCGGGTGGGGAGTGACCCGATTTTCCAGGTGCGTCCATCACCCCTTTCTTTGACT
CGGAAAGGGAACTCCCTGACCCCTTGCGCTTCCCAGGTGAGGCAATGCCTCGCCCTGCTTCGGCTCGCG
CACGGTGCGCACACACACTGGCCTGCGCCACTGTCTGGCACTCCCTAGTGAGATGAACCCGGTACCTCA
GATGGAAATGCAGAAATCACCCGTCTTCTGCGTCGCTCACGCTGGGAGCTGTAGACCGGAGCTGTTCCT
ATTCGGCCATCTTGGCTCCC

- Clip:6013-7642 aligns to chr9:118632313-118633943,  99% identity

### HG03516 chr12:126318373

➢ The same genomic coordinate as in ERX2355869 (page 33)

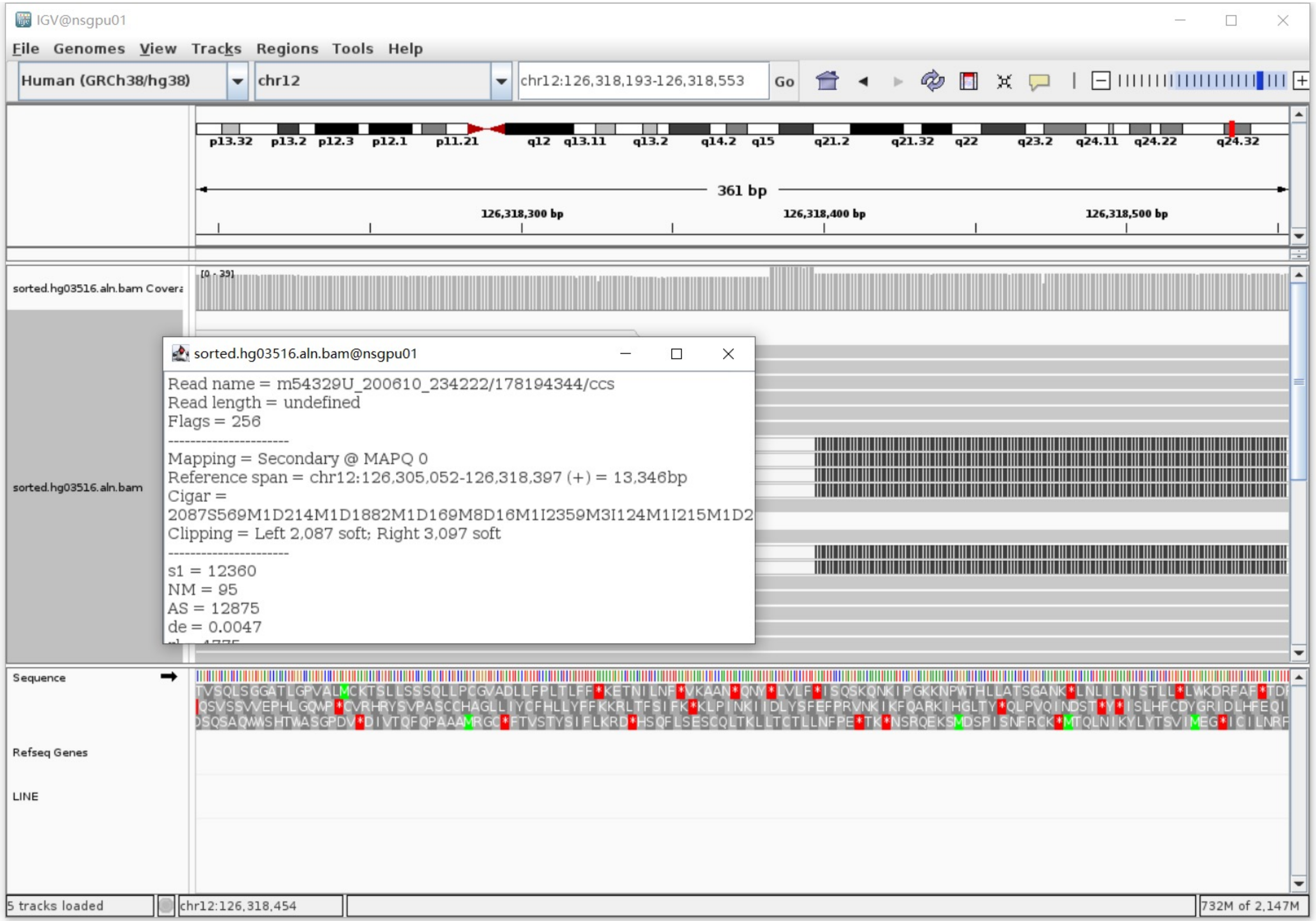


Annotation: For the read in white and black,

- White is the secondary alignment
- Black is the soft clip of the secondary alignment

### HG03540 chr1:48963060

- ➢ Using the built-in BLAT tool in IGV to check where the inserted sequence aligned.
- ➢ The annotation at the aligned location indicated it as L1Hs.

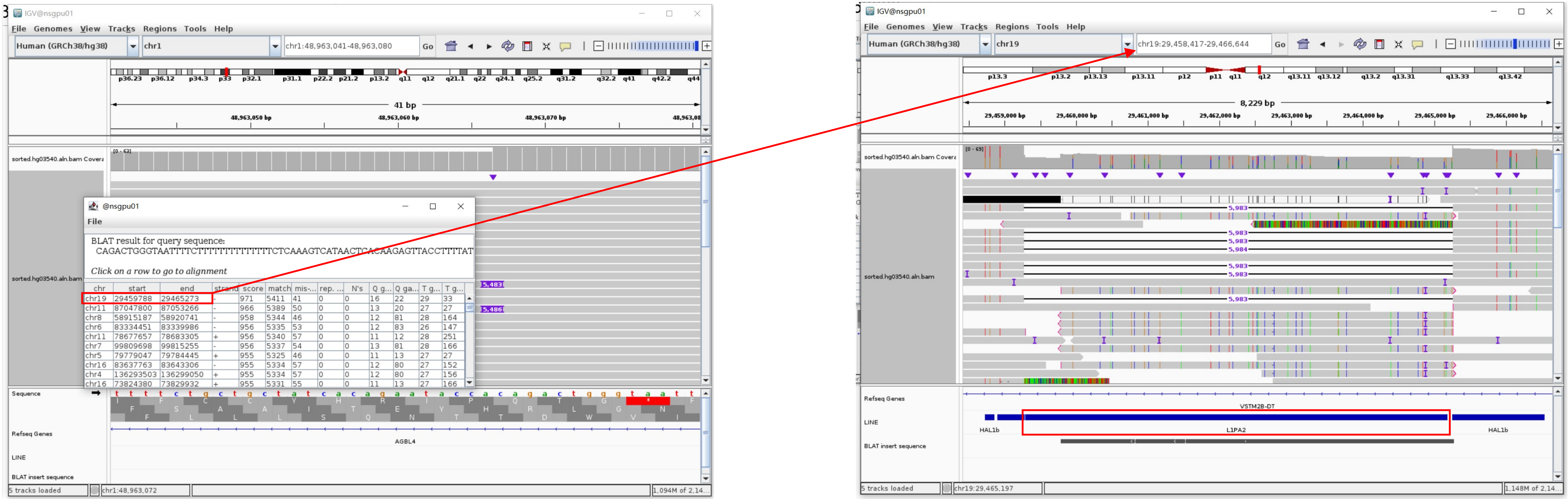


Insertion sequence (5483bp)

- ➢ 18bp TSD

CAGACTGGGTAATTTTCT

- ➢ Insertion:99-5483 aligns to L1Hs:5898-500, 99.16% identity

TGCTATCCCTCCCCCCTTCCCCGACCCCACCACAGTCCCCAGACTGTGATATTCCCCTTCCTGTGTCCATGTGATCTCATTGTTCAATTCCCACCTATGAGTGAGAATATGCGGTGTTTGGTTTTTTGTTCTTGCGATAGTT
ACTGAGAATGATGGTTTCCAATTTCATCCATGTCCCTACAAAGGACATGAACTCATCATTTTTTATGGCTGCATAGTATTCCATGGTGTATATGTGCCACATTTTCTTAATCCAGTCTATCATTGTTGGACATTTGGGTTGG
TTTCCAAGTCTTTGCTATTGTGAATAATGCCGCAATAAACATACGTGTGCATGTGTCTTTATAACAGCATGATTTATAGTCATTTGGGTATATACCCAGTAATGGAATGGCTGGGTCAAATGGTATTGCTAGTTCTAGATCC
CTGAGGAATCGCCCCACTGACTTCCACAATGGTT..............CGGCTCTGAGGCTTCTGCATTCTTCACGTAGTTCTTGAGCCTTGGTTTTCAGCTCCATCAGCTCCTTTAAGCACTTCTCTGTATTGGTTATTCTAGTTATACA
TTCTTCTAAATTTTTTTCAAAGTTTTCAACTTCTTTGCCTTTGGTTTGAATGTCCTCCCGTAGCTCAGAGTATTTGATCGTCTGAAGCCTTCTTCTCTCAGCTCGTCAAAAATCATTCTCCATCCAGCTTTGTTCCATTGCT
GGTGAGGAACTGCGTTCCTTTGGAGGAGGAGAGGCGCTCTGCGTTTTAGAGTTTCCAGTTTTTCTGTTCTGTTTTTTCCCCATCTTTGTGGTTTTATCTACTTTTGGTCTTTGATGATGGTGATGTACAGATAGGTTTTCGG
TGTAGATGTCCTTCTGGTTGTTAGTTTTCCTTCTAACAGACAGGACCCTCAGCTGCAGGTCTGTTGGATAACCTGCCGTGTGAGGTGTCAGTGTGCCCCTGCTGGGGGTGCCTCCCAGTTAGGCTGCTCGGGGGTCAGGGGT
CAGGGACCCACTTGAGGAGGCAGTCTGCCCCTTCTCAGATCTCCAGCTGCGTGCTGGGAGAACCACTGCTCTCTTCAAAGCTGTCAGACAGGGACACTTAAGTCTGCAGAGGTTACTGCTGTCTTTTTGTTTGACTGTGCCC
TGCCCCCAGAGGTGGAGCCTACAGAGGCAGGCAGGCCTCCTTGAGCTGTGGTGGGCTCCAC

### HG03540 chr7:106327425

- ➢ The same genomic coordinate as in HG02630 (page 11)

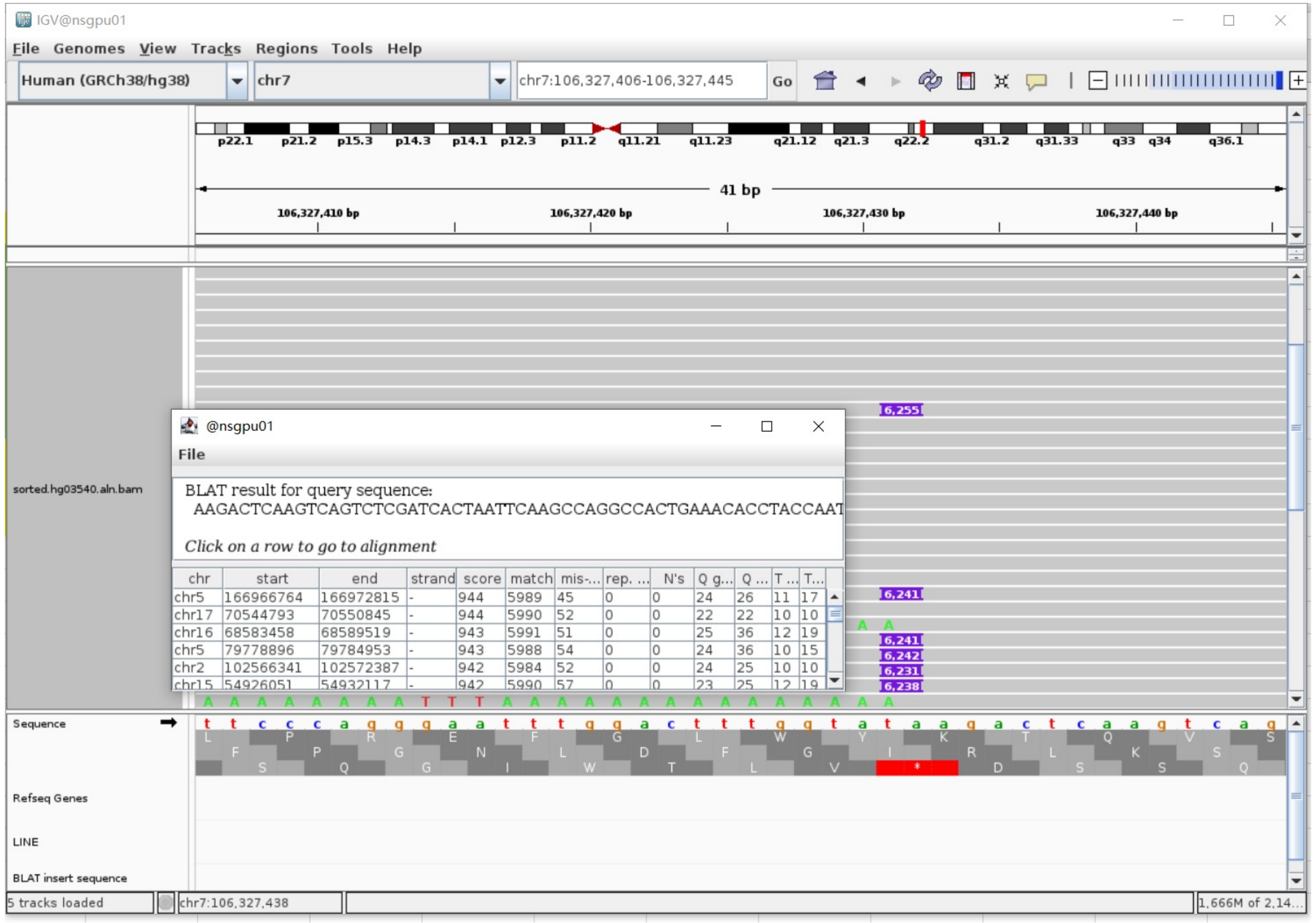

### HG03540 chr20:17880269

- Mapping quality is 0, indicating that this region may have alignment issues.

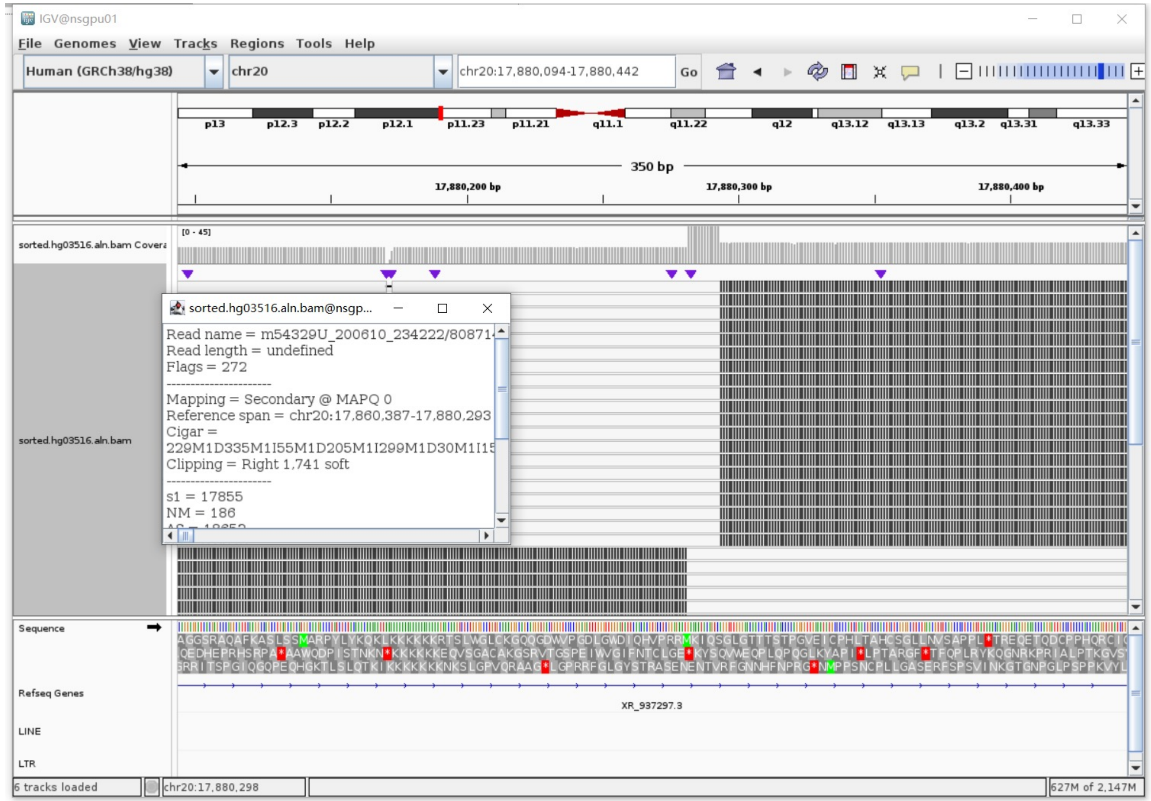


Annotation: For the read in white and black,

- White is the secondary alignment
- Black is the soft clip of the secondary alignment

- Fetch one read (m64043_200522_232930/21039823/ccs, 22763 bp) and BLAT it to the hg38 genome.

  The best alignment is to chr20_GL383577v2_alt (an alternative sequence of chr20) , which contains a L1Hs element.

| ACTIONS | | | QUERY | SCORE | START | END | QSIZE | IDENTITY | CHROM | STRAND | START | END | SPAN |
|---|---|---|---|---|---|---|---|---|---|---|---|---|---|
| browser | new tab | details | YourSeq | 21715 | 936 | 22763 | 22763 | 99.8% | chr20_GL383577v2_alt | + | 94221 | 116060 | 21840 |
| browser | new tab | details | YourSeq | 14814 | 936 | 15853 | 22763 | 99.8% | chr20 | + | 17865356 | 17880293 | 14938 |

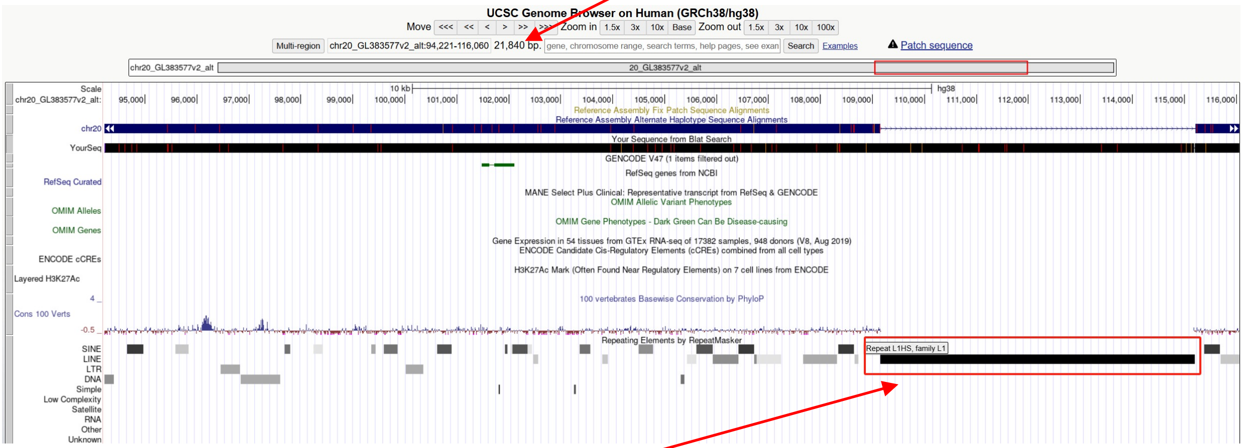


chr20_GL383577v2_alt:109154-115202 (L1Hs marked by RepeatMasker)

- Read:15855-21907 aligns to L1Hs:5-6053, 99.54% identity, ACA TAA

GAGGAGCCAAGATGGCCGAATAGGAACAGCTCCGGTCTACAGCTCCCAG...........CACACTCTGGGGACTG
TGATGGGGTCGGGGGGAGGGGGGAGGGATAGCATTGGGAGATATACCTAATGCTAGATGACACATTAGT
GGGTGCAGCGCACCAGCATGGCACATGTATACATATGTAACTAAACCTGCACAATGTGCACATGTACCC
TAAAACTTAAAGTATAATAAAAAAAAAAAAAAAAAAAAAAAAAAGAAAATA

- It’s a L1Hs in the chromosome chr20_GL383577v2_alt

### NA18906 chr1:226583194

- ➢ Using the built-in BLAT tool in IGV to check where the inserted sequence aligned.
- ➢ The annotation at the aligned location indicated it as L1Hs.

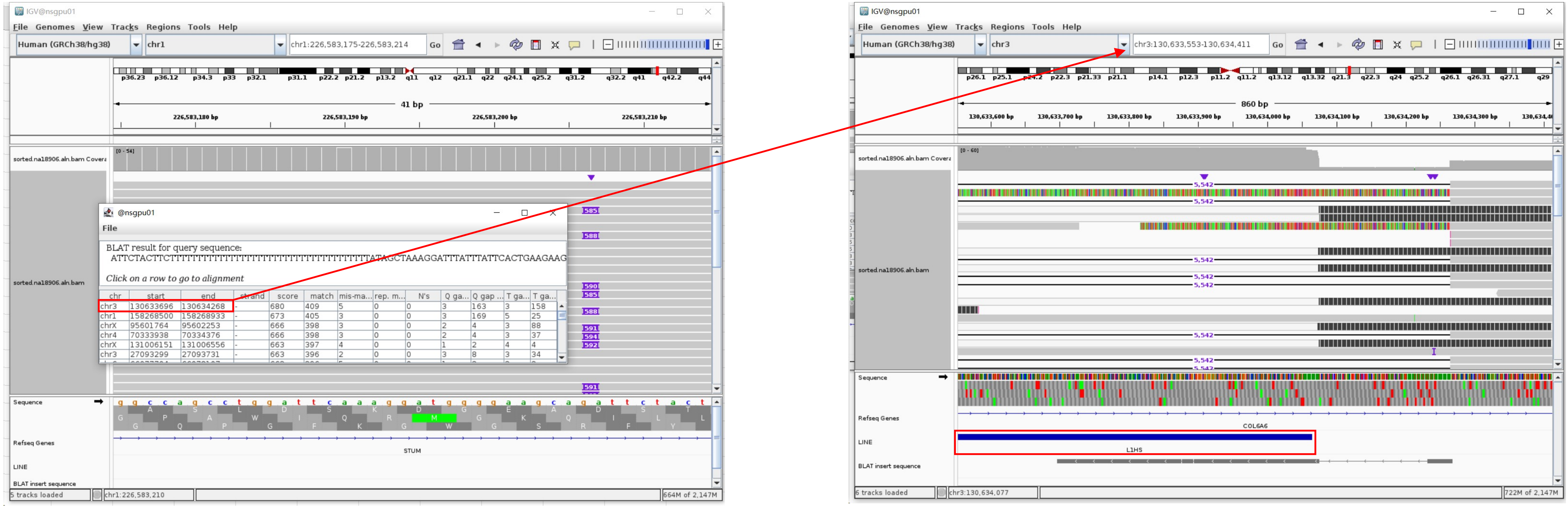


Insertion sequence (585bp)

- ➢ 12bp TSD

ATTCTACTTCTT

- ➢ Insertion:183-585 aligns to L1Hs:6064-5664, 97.49% identity, ACA

TTTTTTTTTTTTCATTATTTTTATTTTATTTTATTTTTTTTTTTTTATTATACTCGATAAGTTTTAGGGTACATGTGCACATTGTGCAGGTTAGTTACATATGTATACATGTGCCATGCTGGTGCGCTGCACCCA
CTAATGTGTCATCTAGCATTAGGTATATCTCCCAATACTATCCCTACCCCCTCCCCCAACCCCACCACAGTCCCAGAGTGTGATATTCCCTTCCTGTGTCCATGTGATCTCATTGTTCAATTCCCACCTATGAGT
GAGAATATGCGGTGTTTGGTTTTTTGTTCTTGCGATAGTTTACTGAGAATGATGGTTTCCAATTTCATCCATGTCCCTACAAAGGATATGAACTCATCATTTTTTATGGCTGCATAGTATTCCATGGTGTAT

### NA18906 chr5:19180266

- ➢ Using the built-in BLAT tool in IGV to check where the inserted sequence aligned.

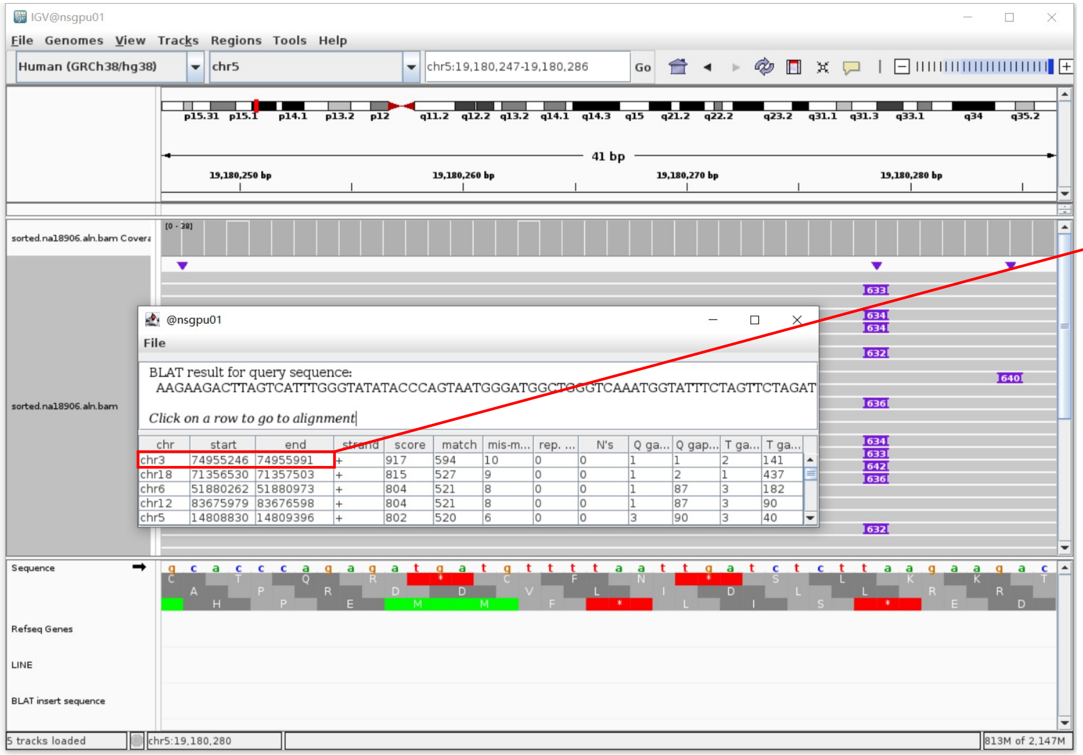


- ➢ The annotation at the aligned location indicated it as L1Hs.

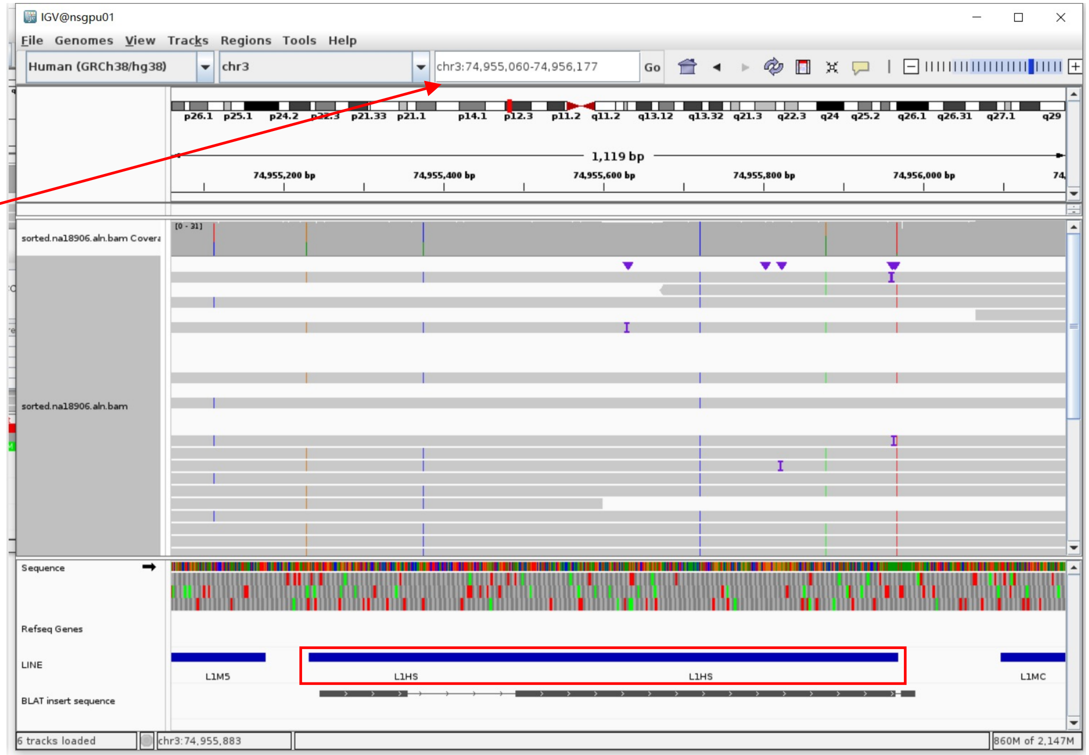


Insertion sequence

- ➢ 12bp TSD

AAGAAGACTTAG

- ➢ Insertion:112-631 aligns to L1Hs:5547-6064, 98.45% identity, ACG TAG

CTAGTTTAAAGACACATGCACACGTATGTTTATTGCGGCACTATTCACAATAGCAAAGACTTGGAACCAACCCAAATGTCCAACAATGATAGACTGGATTAAGAAAATGTGGCACATATACACCATGGAATACTATGCAGCCA
TAAAAAATGATGAGTTCATGTCCTTTGTAGGGACATGGATGAAATTGGAAACCATCATTCTCAGTAAACTATCGCAAGAACAAAAACAACCACACCGCATATTCTCACTCATAGGTGGGAATTGAACAATGAGATCACATGGA
CACAGGAAGGGGAATACCACACTCTGGGGACTGTGGTGGGGTCAGGGGAGGGGGGAGGGATAGCATTGGGAGATATACCTAATGCTAGATGACACGTTAGTGGGTGCAGCGCACCAGCATGGCACATGTATACATATGTAACT
AACCTGCACAATGTGCACATGTACCCTAAAACTTAGAGTATAATAAAAAAAAAAATTAAAAAAAAAAAAAAAAAAAAAAAAAAAAAAAAAA

### NA18906 chr7:106327425

- ➢ The same genomic coordinate as in HG02630 (page 11)

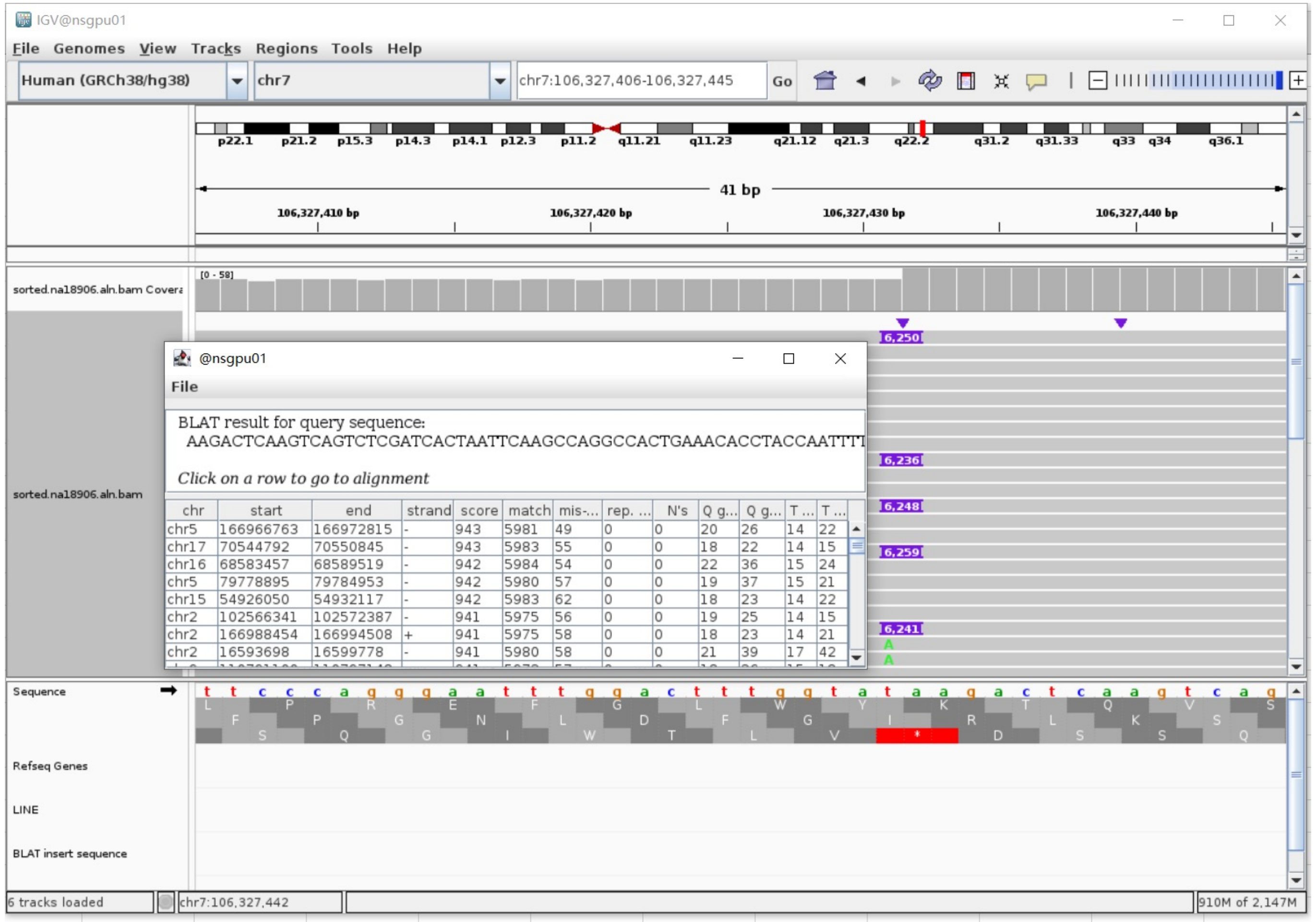

### NA20129 chr5:57983715

- The same genomic coordinate as in HG00438 (page 2)

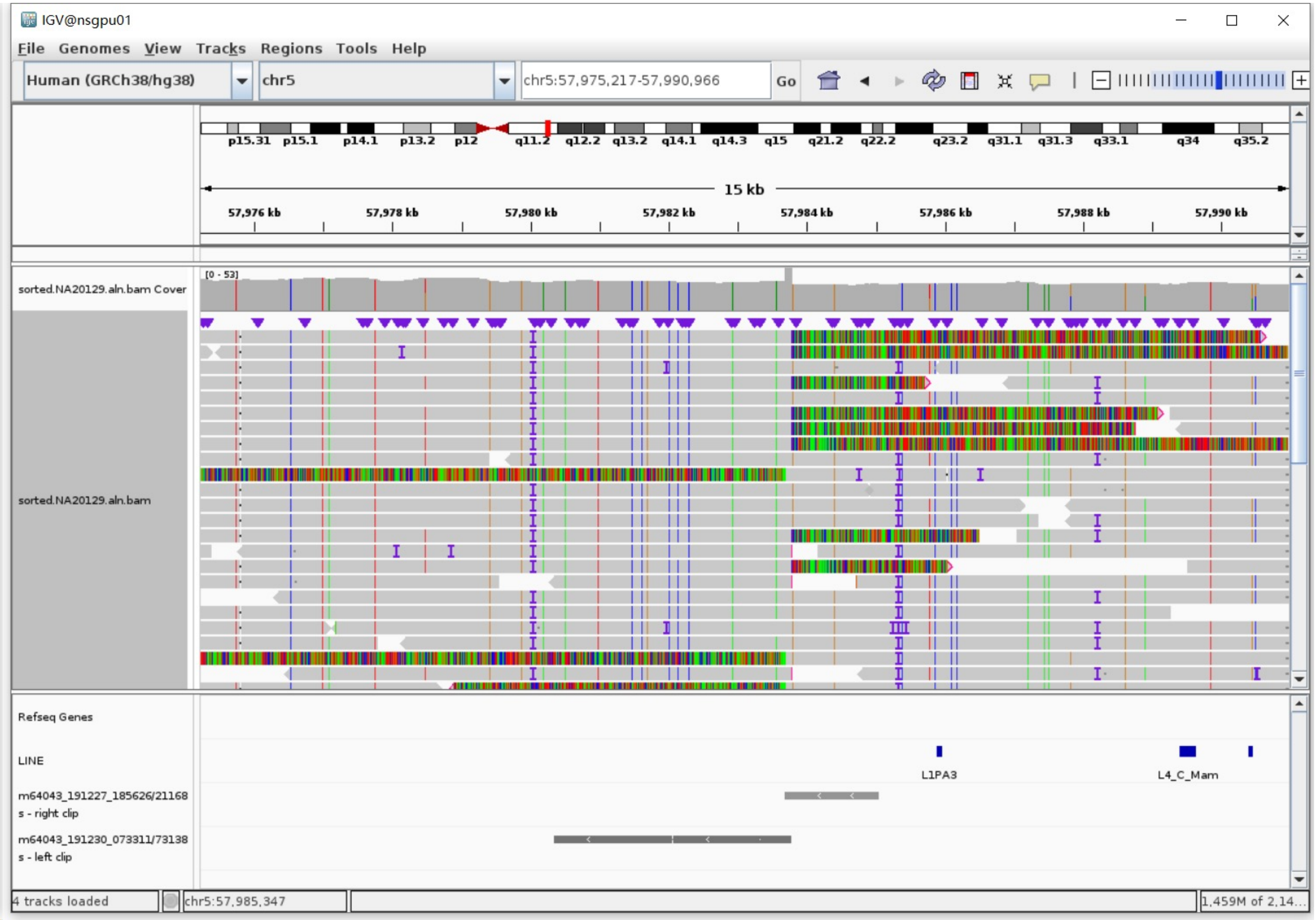

### NA20129 chr6:16950580

- The same genomic coordinate as in HG00438 (page 2)

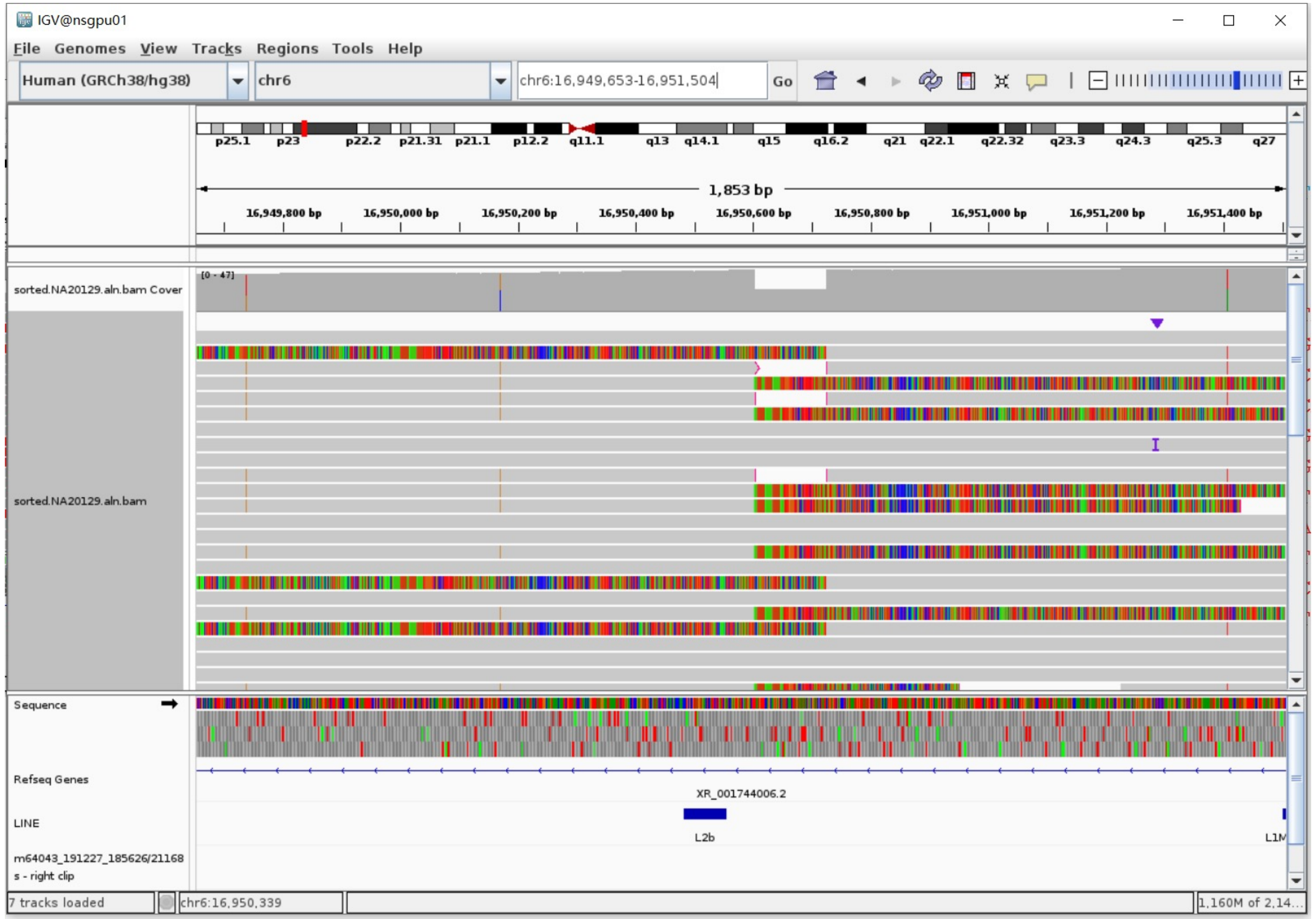

### NA20129 chr17:35945138

- ➢ Using the built-in BLAT tool in IGV to check where the inserted sequence aligned.
- ➢ The annotation at the aligned location indicated it as L1Hs.

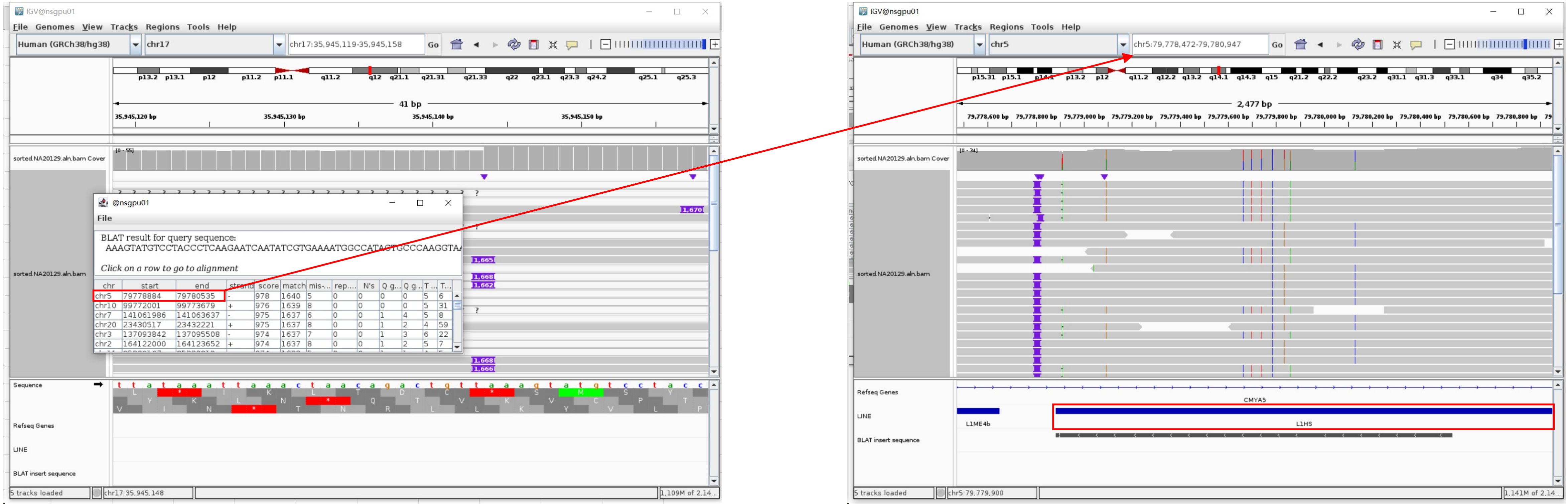


#### Insertion sequence

- ➢ Insertion:4-1659 aligns to L1Hs:4410-6064, 99.52% identity, ACG TAG

AAGAATCAATATCGTGAAAATGGCCATACTGCCCAAGGTAATTTACAGATTCAATGCCATCCCCATCAAGCTACCAATGACTTCTTTCTTCACAGAATTGGAAAAAACTACTTTAAAGTTCATATGGAACCAAAAAAGAGCCCGCA
TCGCCAAGTCAATCCTAAGCCAAAAGAACAAAGCTGGAGGCATCACACTACCTGACTTCAAACTATACTACAAGGCTACAGTAACCAAAACAGCATGGTACTGGTACCAAAACAGAGATATAGATCAATGGAACAGAACAGAGCCC
TCAGAAATAATGCCACATATCTACAACTATCTGATCTTTGACAAACCTGAGAAAAACAAGCAATGGGGAAAGGATTCCCTATTTAATAAATGGTGCTGGGAAAACTGGCTAGCCATATGTAGAAAGCTGAAACTGGATCCCTTCCT
TACACCTTATACAAAAATCAATTCAAGATGGATTAAAGATTTAAACGTTAGACCTAAACCATAAAAACCCTAGAAGAAAACCTAGGCATTACCATTCAGGACATAGGCGTGGGCAAGGACTTCATGTCCAAAACACCAAAAGCAAT
GGCAACAAAAGCCAAAATTGACAAATGGGATCTAATTAAACTAAAGAGCTTCTGCACAGCAAAAGAAACTACCATCAGAGTGAACAGGCAACCTACAAAATGGGAGAAAATTTTCGCAACCTACTCATCTGACAAAGGGCTAATAT
CCAGAATCTACAATGAACTCAAACAAATTTACAAGAAAAAAACAAACAACCCCATCAAAAAGTGGGCGAAGGACATGAACAGACACTTCTCAAAAGAAGACATTTATGCAGCCAAAAAACACATGAAGAATGCTCATCATCACTGG
CCATCAGAGAAATGCAAATCAAAACCACTATGAGATATCATCTCACACCAGTTAGAATGGGCAATCATTAAAAAGTCAGGAAACAACAGGTGCTGGAGAGGATGTGGAGAAATAGGAACACTTTTACACTGTTGGTGGGACTGTAA
ACTAGTTCAACCATTGTGGAAGTCAGTGTGGCGATTCCTCAGGGATCTAGAACTAGAAATACCATTTGACCCAGCCATCCCATTACTGGGTATATACCCAAAGGACTATAAATCATGCTGCTATAAAGACACATGCACACGTATG
TTTATTGCGGCACTATTCACAATAGCAAAGACTTGGAACCAACCCAAATGTCCAACAATGATAGACTGGATTAAGAAAATGTGGCACATATACACCATGGAATACTATGCAGCCATAAAAAATGATGAGTTCATGTCCTTTGTAGG
GACATGGATGAAACTGGAAACCATCATTCTCAGTAAACTATCGCAAGAACAAAAACCAAACACCGCATATTCTCACTCATAGGTGGGAATTGAACAATGAGATCACATGGACACAGGAAGGGGAATATCACACTCTGGGGACTGT
GGTGGGGTCGGGGAGGGGGGAGGGATAGCATTGGGAGATATACCTAATGCTAGATGACACGTTAGTGGGTGCAGCGCACCAGCATGGCACATGTATACATATGTAACTAACCTGCACAATGTGCACATGTACCCTAAAACTTAGAG
TATAATAAAAAAAAAAAATTAAAAAAAAAAATAAAAAAAAATAAAAAAAAA

### ERX2355869 chr3:166374422

- Using the built-in BLAT tool in IGV to check where the inserted sequence aligned.

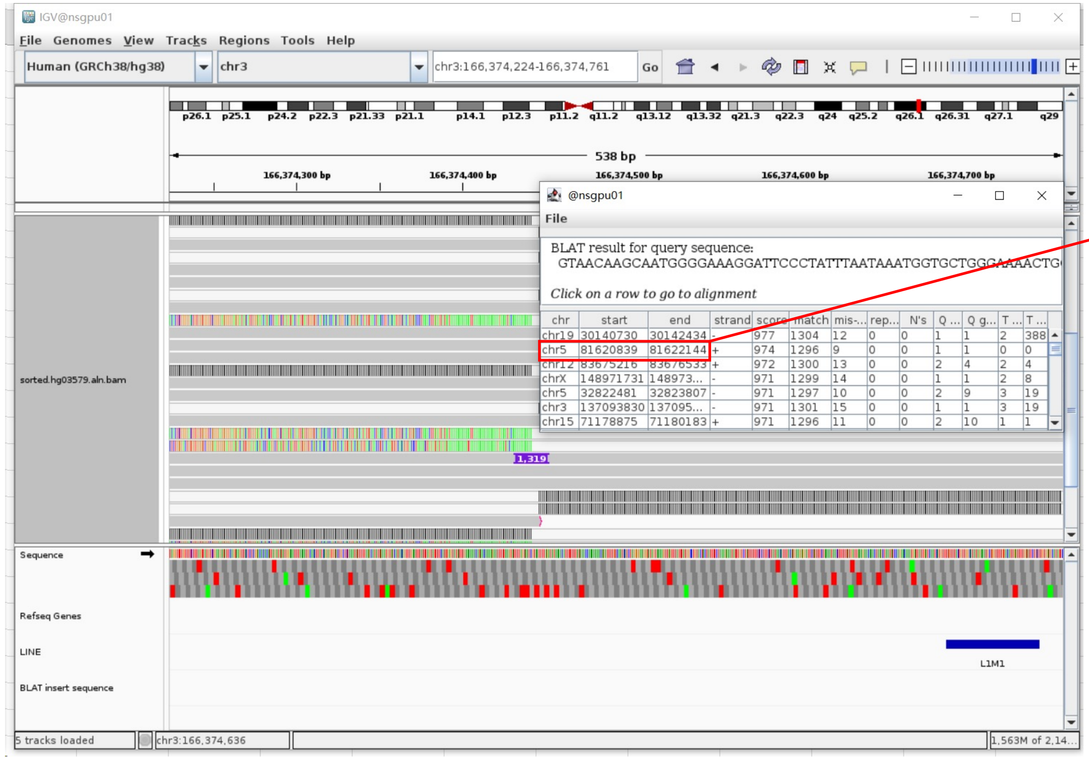


- The annotation at the aligned location indicated it as L1Hs.

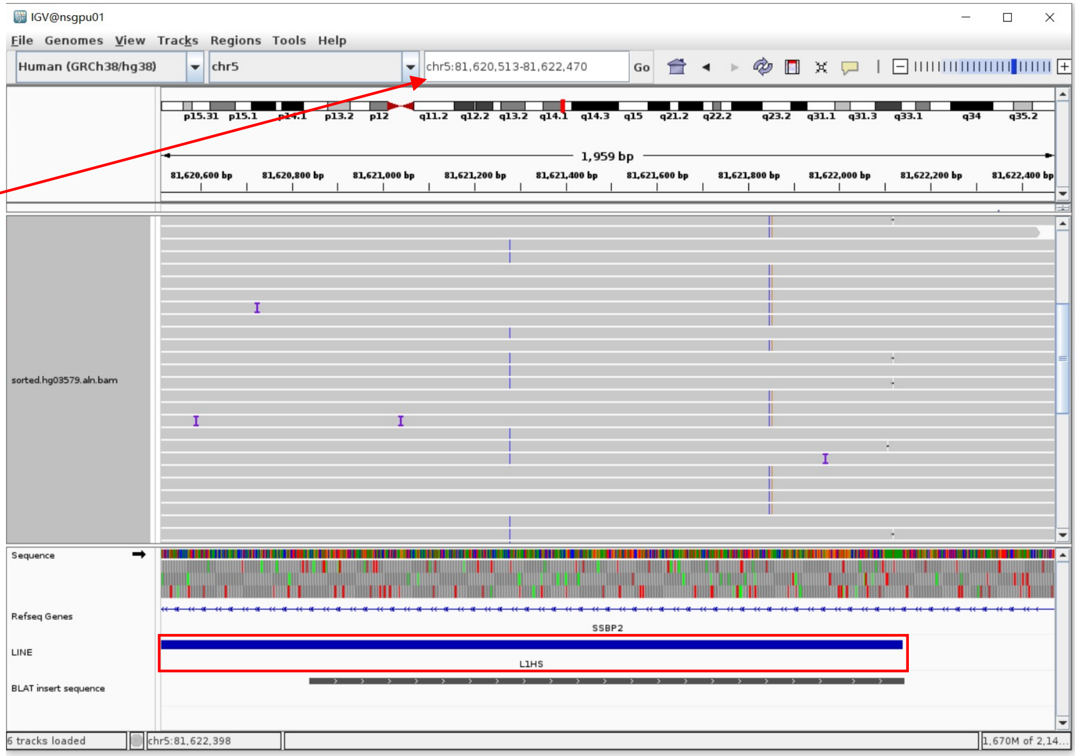


Insertion sequence (1319bp)

- 12bp TSD

AAGAAGACTTAG

- Insertion:2-1314 aligns to L1Hs:4753-6064, 98.78% identity, GAG(active in a small population) TAG

AACAAGCAATGGGGAAAGGATTCCCTATTTAATAAATGGTGCTGGGAAAACTGGCTAGCCATATGTAGAAAGCTGAAACTGGATCCCTTCCTTACACCTTATACAAAAATCAATTCAAGATGGATTAAAGATTTAAACGTTAGACC
TAAAACCATAAAAACCCTAGAAGAAAACCTAGGCATTACCATTCAGGACATAGGCGTGGGCAAGGACTTCATGTCCAAAACACCAAAAGCAATGGCAACAAAAGCCAAAATTGACAAATGGGATCTAATTAAACTCAAGAGCTTCT
GCACAGCAAAAGAAACTACCATCAGAGTGAACAGGCAACCTACAACATGGGAGAAAATTTTCGCAACCTACTCATCTGACAAAGGGCTAATATCCAGAATCTACAATGAACTCAAACAAATTTACAAGAAAAAAACAAACAACCCA
ATCAAAAAGTGGGCGAAGGACATGAACAGACACTTCTCAAAAGAAGACATTTATGCAGCCAAAAAACACATGAAGAAATGCTCATCATCACTGGCCATCAGAGAAATGCAAATCAAAACCACTATGAGATATCATCTCACACCAGT
TAGAATGGCAATCATTAAAAAGTCAGGAAACAACAGGTGCTGGAGAGGATGTGGAGAAATAGGAACACTTTTACACTGTTGGTGGGACTGTAAACTAGTTCAACCATTGTGGAAGTCAGTGTGGCGATTCCTCAGGGATCTAGAAC
TAGAAATACCATTTGACCCAGCCATCCCATTACTGGGTATATACCCAAAGGACTATAAATCATGCTGCTATAAAGACACATGCACACGTATGTTTATTGCGGCACTATTCACAATAGCAAAGACTTGGAACCAACCCAAATGTCCA
ACAATGATAGACTGGATTAAGAAAATGTGGCACATATACACCTTGGAATACTATGCAGCCATAAAAAATGATGAGTTCATGTCCTTTGTAGGGACATGGATGAAATTGGAAACCATCATTCTCAGTAAACTATTGCAAGAACAAAA
AACCAAACACCGCATGTTCTCACTCATAGGTGGGAATTGAACAATGAGATCACATGGACACAGGAAGGGGAATATCACACTCTGGGGACTGTGGTGGGGTCGGGGGAGGGGGGGAGGGATAGCATTGGGAGACATACCTAAGGCTA
GATGACGAGTTAGTGGGTGCAGCACACCAGCATGGCACATGTATACATATGTAACTAACCTGCACAATGTGCACATGTACCCTAAAACTTAGAGTATAATAAAAAAAAAAAAATTAAAAAAAAAAAAAAAGAAAATACAAAAAAA

### ERX2355869 chr4:68787387

- Using the built-in BLAT tool in IGV to check where the inserted sequence aligned.

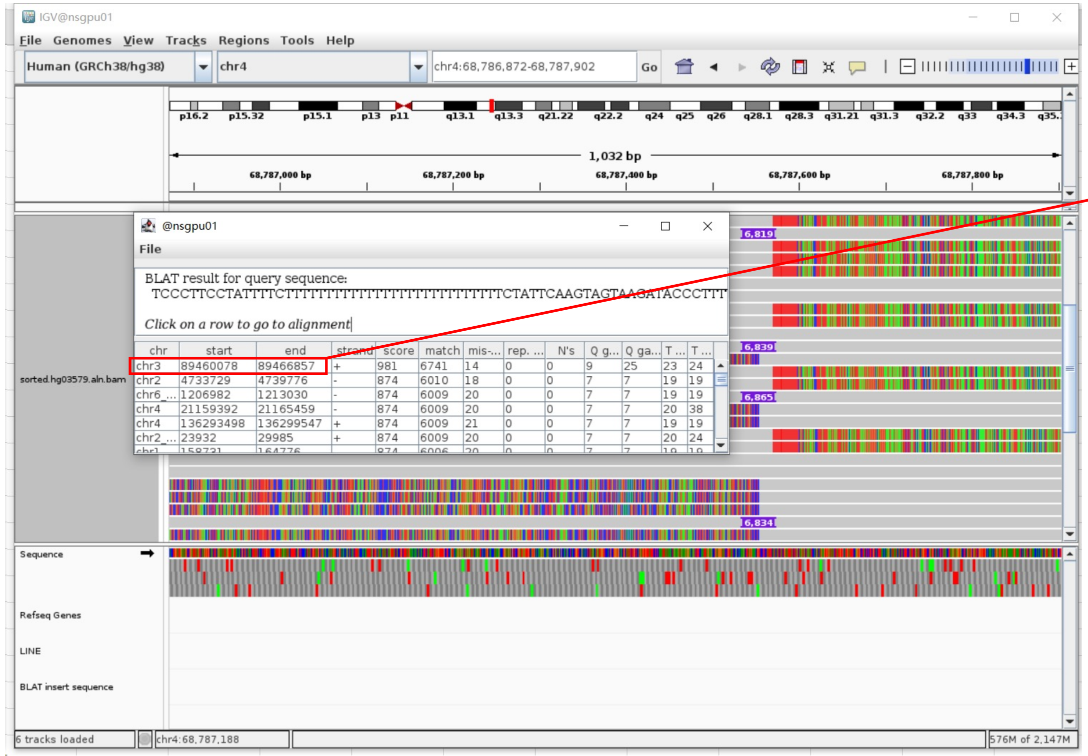


- The annotation at the aligned location indicated it as L1Hs.

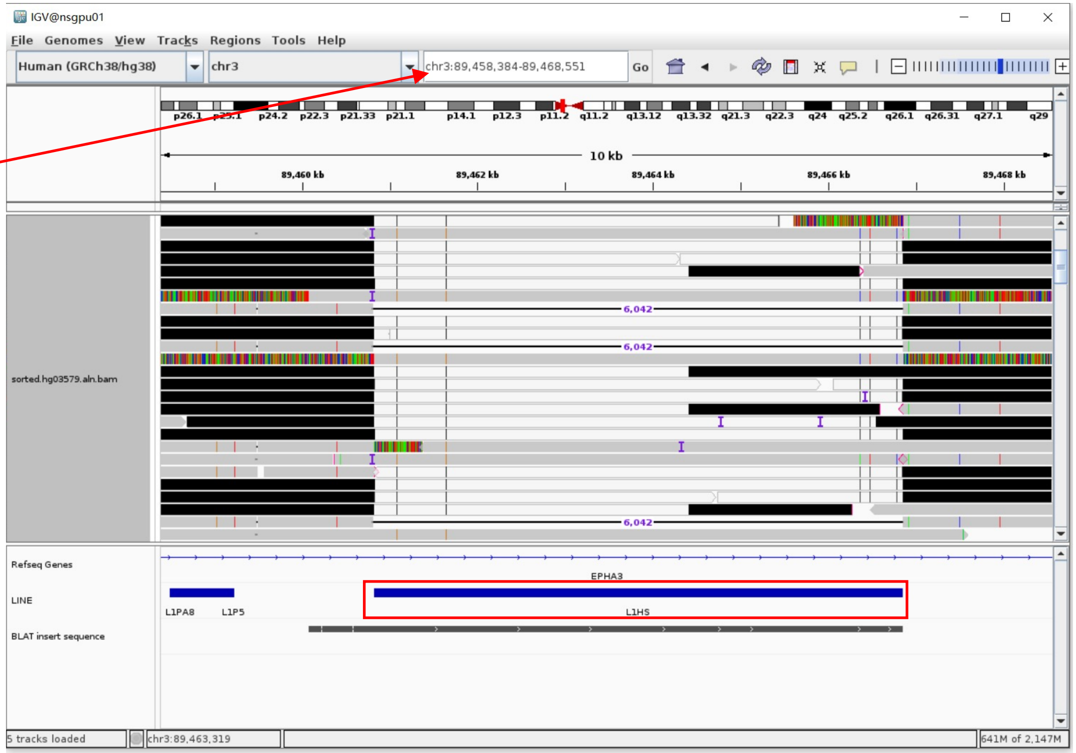


Insertion sequence (6819bp)

- 12bp TSD

AAGAAGACTTAG

- Insert:768-6819 aligns to L1Hs:6064-1, 99.49% identity, ACA TAG

TCTCTGTCTCTCTTTTTTTTTTTTTTTTTTTTTTTTTTTTTTTTTTTTATTATACTCTAAGTTTTAGGGTACATGTGCACATTGTGCAGGTTAGTTACATATGTATACATGTGCCATGCTGGTGCGCTGCACCCACTAATGTGT
CATCTAGCATTAGGTATATCTCCCAATGCTATCCCTCCCCCCCTCCCCCGACCCACCACAGTCCCCAGAGTGTGATATTCCCCTTCCTGTGTCCATGTGATCTCATTGTTCAATTCCCACCTATGAGTGAGAATATGCGG
TGTTTGGTTTTTTGTTCTTGCGATAGTTTACTGAGAATGATGGTTTCCAATTTCATCCATGTCCCTACAAAGGATATGAACTCATCATTTTTTATGGCTGCATAGTATTCCATGGTGTATATGTGCCACATTTTCTTAAT
CCAGTCTATCATTGTTGGACATTGGGTTGGTTCCAAGTCTTTGCTA……………CCTGCCCCCAGAGGTGGAGCCTACAGAGGCAGGCAGGCCTCCTTGAGCTGTGGTGGGCTCCACCCAGTTCGAGCTTCCCGCTGCTTTGT
TTACCTAAGCAAGCCTGGCAATGGGCGGCGCCCCTCCCCCAGCCTCGTTGCCGCCTTGCAGTTTGATCTCAGACTGCTGTGCTAGCAATCAGCGAGATTCCGTGGGCGTAGGACCCTCCGAGCCAGGTGTGGGATATAGT
CTTGTGGTGCGCCGTTTCTTAAGCCGGTCTGAAAAGCGCAATATTCGGGTGGGAGTGACCCGATTTTCCAGGTGCGTCCGTCACCCCTTTCTTTGACTCGGAAAGGGAACTCCCTGACCCCTTGCGCTTCCCAGGTGAGG
CAATGCCTCGCCCTGCTTCGGCTCGCGCACGGTGCGCACACACACTGGCCTGCGCCCACTGTCTGGCACTCCCTAGTGAGATGAACCCGGTACCTCAGATGGAAATGCAGAAATCACCGTCTTCTGCGTCGCTCACGCTG
GGAGCTGTAGACCGGAGCTGTTCCTATTCGGCCATCTTGGCTCCTCCCTCC

### ERX2355869 chr5:161008649

- ➢ The same genomic coordinate as in HG00438 (page 2)

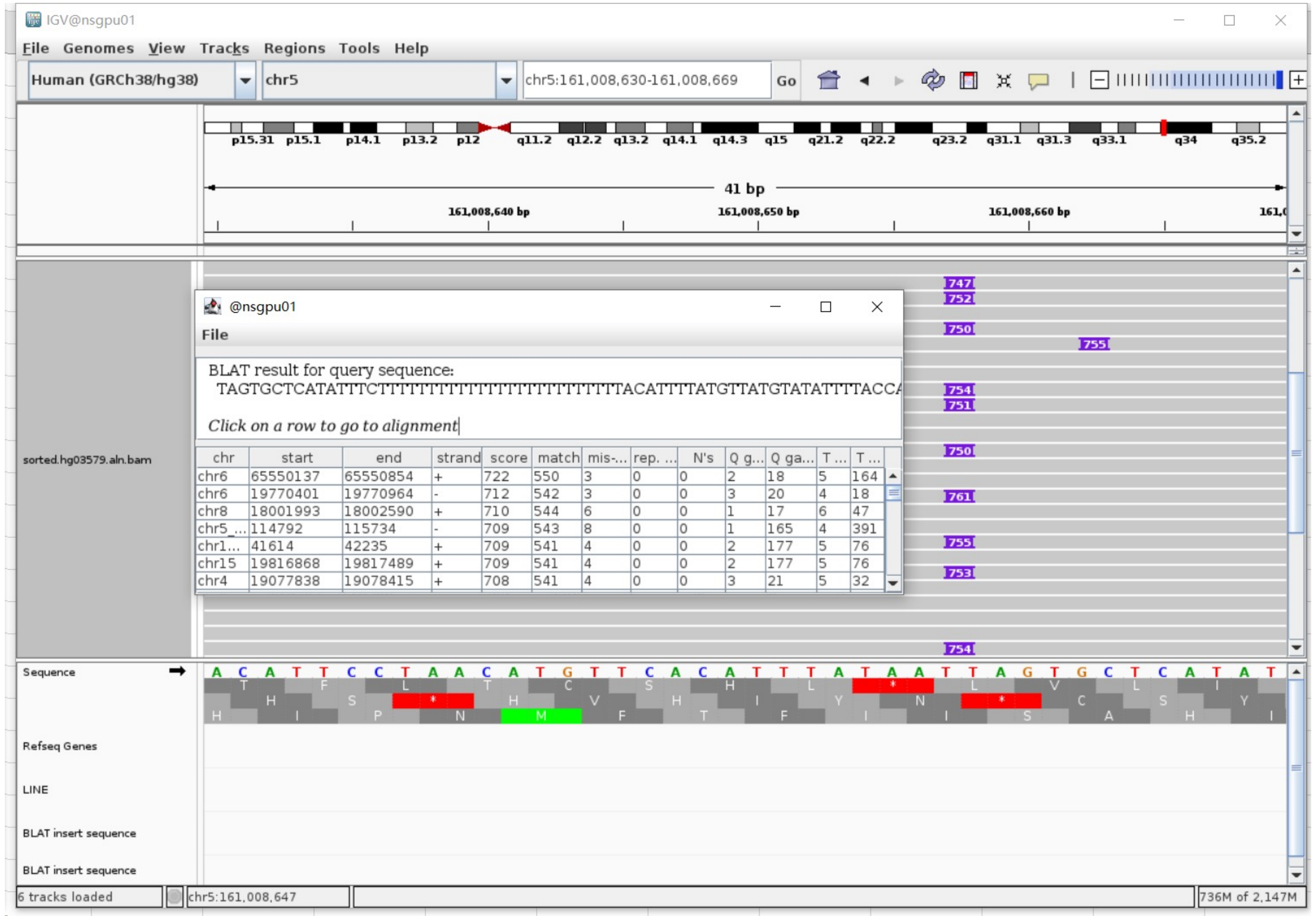

### ERX2355869 chr12:126318373

➢ Mapping quality is 0, indicating that this region may have alignment issues.

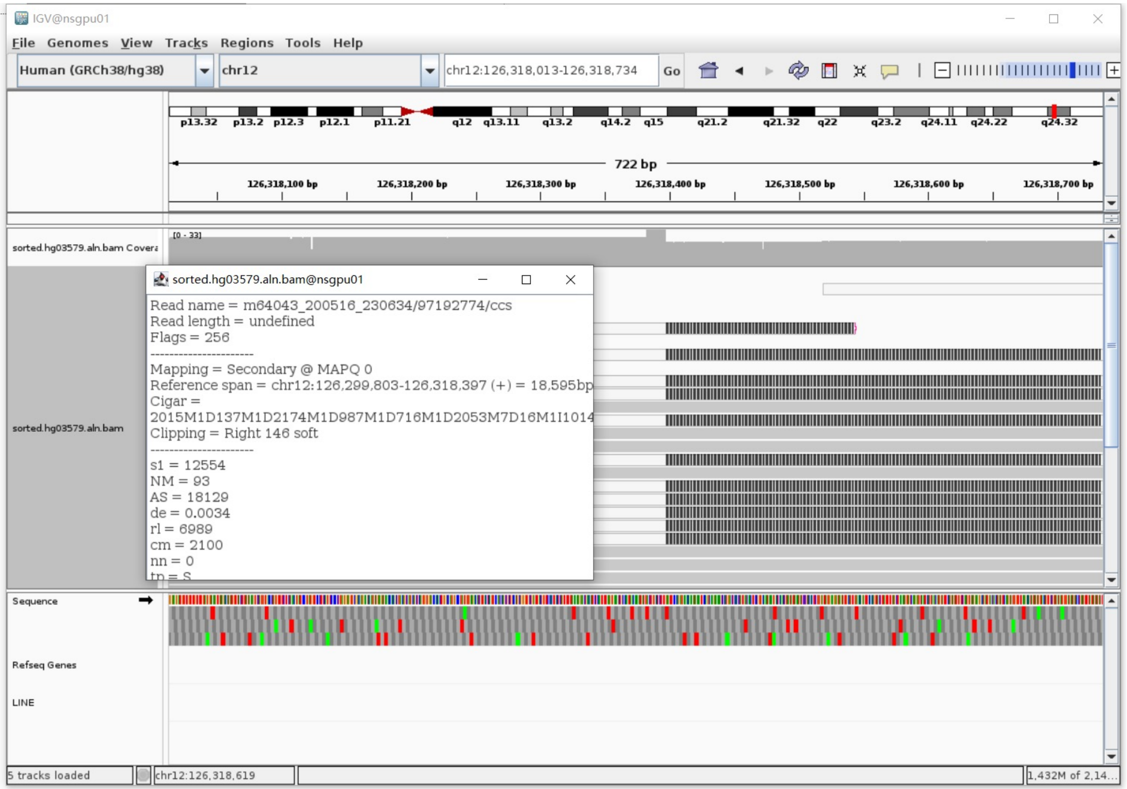


Annotation: For the read in white and black,

- White is the secondary alignment
- Black is the soft clip of the secondary alignment

➢ Fetch one read (m64043_200516_230634/5112335/ccs, 19045 bp) and BLAT it to the hg38 genome.

- Read:1-13498

The best alignment is to chr12_GL383551v1_alt (an alternative sequence of chr12), which contains a L1Hs element.

| ACTIONS | | | QUERY | SCORE | START | END | QSIZE | IDENTITY | CHROM | STRAND | START | END | SPAN |
|---|---|---|---|---|---|---|---|---|---|---|---|---|---|
| browser | new tab | details | YourSeq | 13402 | 1 | 13498 | 19045 | 99.7% | chr12_GL383551v1_alt | - | 85576 | 99074 | 13499 |
| browser | new tab | details | YourSeq | 6148 | 12844 | 19045 | 19045 | 99.7% | chr12 | - | 126307224 | 126313446 | 6223 |

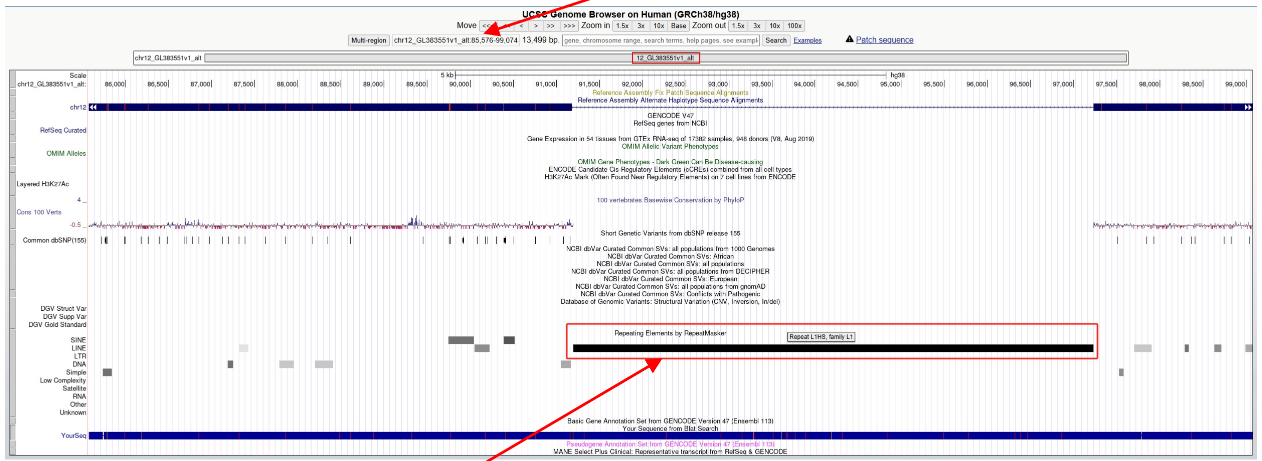


chr12_GL383551v1_alt:91197-97228 (L1HS marked by RepeatMasker)

- Read:13499-19045

The best alignment is also to chr12_GL383551v1_alt

| ACTIONS | | | QUERY | SCORE | START | END | QSIZE | IDENTITY | CHROM | STRAND | START | END | SPAN |
|---|---|---|---|---|---|---|---|---|---|---|---|---|---|
| browser | new tab | details | YourSeq | 5514 | 1 | 5547 | 5547 | 99.8% | chr12_GL383551v1_alt | - | 80017 | 85575 | 5559 |
| browser | new tab | details | YourSeq | 5499 | 1 | 5547 | 5547 | 99.7% | chr12 | - | 126307224 | 126312778 | 5555 |

- Read:1844-7906 aligns to L1Hs:1-6062, 99.50% identity, ACA TAG

GGGGGGAGGAGCCAAGATGGCCG..............TAATGCTAGATGACACATTAGTGGGTGCAGCGCACCAGCATGGCAC
ATGTATACATATGTAACTAACCTGCACAATGTGCACATGTACCCTAAAACTTAGAGTATAATAAAAAAAAAAAA
AAAAAAAAAAAAAGAGTACAAGTCAATAA

➢ It's a L1Hs in the chromosome chr12_GL383551v1_alt
