## Extended Files for "Image-based DNA Sequencing Encoding for Detecting Low-Mosaicism Somatic Mobile Element Insertions": Extended Files 2.pdf

### Manual check of *Alu* label that were not identified by Pangenome xTea_long PacBio calls

(Additional explanation for Supplementary Table 8)

| ID | Chr | Coordinate | Manual Inspection |
|---|---|---|---|
| HG00438 | chr6 | 29851071 | TRUE |
| | chr6 | 31471215 | TRUE |
| | chr6 | 32805641 | TRUE |
| | chr6 | 104041958 | TRUE |
| | chr14 | 93364572 | TRUE |
| | chr15 | 32821084 | TRUE |
| | chr18 | 51774994 | TRUE |
| HG00621 | chr4 | 21532942 | TRUE |
| | chr6 | 29851067 | TRUE |
| | chr12 | 11026415 | TRUE |
| | chr13 | 112339461 | TRUE |
| | chr15 | 32662471 | TRUE |
| | chr17 | 37561902 | TRUE |
| | chr18 | 51774994 | TRUE |
| | chr19 | 52384792 | TRUE |
| | chr21 | 22207227 | TRUE |
| HG01106 | chr1 | 82577503 | TRUE |
| | chr6 | 31282439 | TRUE |
| | chr10 | 52920858 | TRUE |
| | chr17 | 227341 | TRUE |
| | chr19 | 52384793 | TRUE |
| HG02630 | chr1 | 82577503 | TRUE |
| | chr2 | 1467932 | TRUE |
| | chr2 | 36290645 | TRUE |
| | chr6 | 29416032 | TRUE |
| | chr6 | 29851071 | TRUE |
| | chr6 | 31471215 | TRUE |
| | chr6 | 79370701 | TRUE |
| | chr7 | 155318862 | TRUE |
| | chr12 | 11026415 | TRUE |
| | chr12 | 58064416 | TRUE |
| HG02630 | chr12 | 58961589 | TRUE |
| | chr14 | 91780893 | TRUE |
| | chr15 | 30856126 | TRUE |
| | chr15 | 73690233 | TRUE |
| | chr16 | 72351770 | TRUE |
| | chr17 | 37509872 | TRUE |
| | chr17 | 37561902 | TRUE |
| | chr17 | 37660775 | TRUE |
| | chr17 | 37883970 | TRUE |
| | chr17 | 38528853 | TRUE |
| HG03492 | chr1 | 198500196 | TRUE |
| | chr3 | 40200107 | TRUE |
| | chr4 | 21604842 | TRUE |
| | chr6 | 30030695 | TRUE |
| | chr6 | 32720794 | TRUE |
| | chr6 | 79377018 | TRUE |
| | chr6 | 104041958 | TRUE |
| | chr8 | 1362327 | TRUE |
| | chr8 | 39822492 | TRUE |
| | chr11 | 134789637 | TRUE |
| | chr13 | 89749568 | TRUE |
| | chr14 | 93364571 | TRUE |
| | chr17 | 41020755 | TRUE |
| | chr18 | 29602115 | TRUE |
| | chr19 | 52384796 | TRUE |
| HG03516 | chr5 | 43541665 | TRUE |
| | chr5 | 69513668 | TRUE |
| | chr6 | 30030695 | TRUE |
| | chr6 | 32740195 | TRUE |
| | chr7 | 142976545 | TRUE |
| | chr8 | 39842932 | TRUE |
| | chr10 | 517814 | TRUE |
| HG03516 | chr12 | 58064416 | TRUE |
| | chr12 | 58961589 | TRUE |
| | chr14 | 39716321 | TRUE |
| | chr14 | 91658700 | TRUE |
| | chr14 | 106651070 | TRUE |
| | chr17 | 37561902 | TRUE |
| HG03540 | chr1 | 82577503 | TRUE |
| | chr1 | 170913134 | TRUE |
| | chr1 | 170913303 | TRUE |
| | chr1 | 238293249 | TRUE |
| | chr4 | 63331066 | TRUE |
| | chr4 | 68078452 | TRUE |
| | chr4 | 68329636 | TRUE |
| | chr6 | 29925071 | TRUE |
| | chr6 | 30254196 | TRUE |
| | chr6 | 131694946 | TRUE |
| | chr8 | 39842808 | TRUE |
| | chr9 | 101285278 | TRUE |
| | chr13 | 89749560 | TRUE |
| | chr14 | 91780893 | TRUE |
| | chr14 | 93364571 | TRUE |
| | chr14 | 93400345 | TRUE |
| | chr17 | 37561902 | TRUE |
| | chr17 | 70457036 | TRUE |
| | chr18 | 76926745 | TRUE |
| | chr19 | 52384793 | TRUE |
| | chr22 | 23928251 | TRUE |
| NA18906 | chr1 | 246243393 | TRUE |
| | chr2 | 36249528 | TRUE |
| | chr6 | 30030695 | TRUE |
| | chr6 | 31471215 | TRUE |
| | chr6 | 32719367 | TRUE |

| ID | Chr | Coordinate | Manual Inspection |
|---|---|---|---|
| NA18906 | chr6 | 32805641 | TRUE |
| | chr7 | 155318862 | TRUE |
| | chr8 | 1362348 | TRUE |
| | chr12 | 57965263 | TRUE |
| | chr12 | 58064417 | TRUE |
| | chr12 | 58987593 | TRUE |
| | chr13 | 62014795 | TRUE |
| | chr14 | 91780873 | TRUE |
| | chr14 | 106651070 | TRUE |
| | chr17 | 17831055 | TRUE |
| | chr17 | 37561902 | TRUE |
| | chr18 | 29602115 | TRUE |
| | chr22 | 23928251 | TRUE |
| NA20129 | chr1 | 82577498 | TRUE |
| | chr4 | 63331066 | TRUE |
| | chr4 | 189727059 | TRUE |
| | chr5 | 34717915 | TRUE |
| | chr6 | 28546544 | TRUE |
| | chr6 | 29851072 | TRUE |
| | chr6 | 29925071 | TRUE |
| | chr6 | 31282422 | TRUE |
| | chr6 | 31336214 | TRUE |
| | chr6 | 32719367 | TRUE |
| | chr6 | 32720794 | TRUE |
| | chr6 | 32805636 | TRUE |
| | chr8 | 1362269 | TRUE |
| | chr8 | 39842932 | TRUE |
| | chr12 | 31253691 | TRUE |
| | chr14 | 91780857 | TRUE |
| | chr14 | 106651070 | TRUE |
| | chr17 | 38594618 | TRUE |
| NA20129 | chr18 | 51774994 | TRUE |

| | | | |
|---|---|---|---|
| ERX2355888 (HG03453) | chr1 | 82577503 | TRUE |
| | chr2 | 36249528 | TRUE |
| | chr6 | 30030695 | TRUE |
| | chr6 | 32805636 | TRUE |
| | chr6 | 33235423 | TRUE |
| | chr6 | 79370732 | TRUE |
| | chr6 | 104041958 | TRUE |
| | chr7 | 142906943 | TRUE |
| | chr9 | 111517769 | TRUE |
| | chr9 | 111518090 | TRUE |
| | chr12 | 11026415 | TRUE |
| | chr12 | 31253677 | TRUE |
| | chr12 | 57965264 | TRUE |
| | chr14 | 93364571 | TRUE |
| | chr17 | 37561901 | TRUE |
| | chr18 | 67471609 | TRUE |
| | chr22 | 42237075 | TRUE |
| ERX2355869 (HG03579) | chr1 | 82577503 | TRUE |
| | chr1 | 246243395 | TRUE |
| | chr2 | 1467935 | TRUE |
| | chr6 | 31282440 | TRUE |
| | chr6 | 99022883 | TRUE |
| | chr6 | 104042021 | TRUE |
| | chr8 | 1362348 | TRUE |
| | chr8 | 85019904 | TRUE |
| | chr12 | 58064363 | TRUE |
| | chr14 | 78516322 | TRUE |
| | chr15 | 30774821 | TRUE |
| | chr17 | 37883970 | TRUE |
| | chr17 | 54885771 | TRUE |

### HG00438 chr6:29851071

- Mapping quality is 0, indicating that this region may have alignment issues.

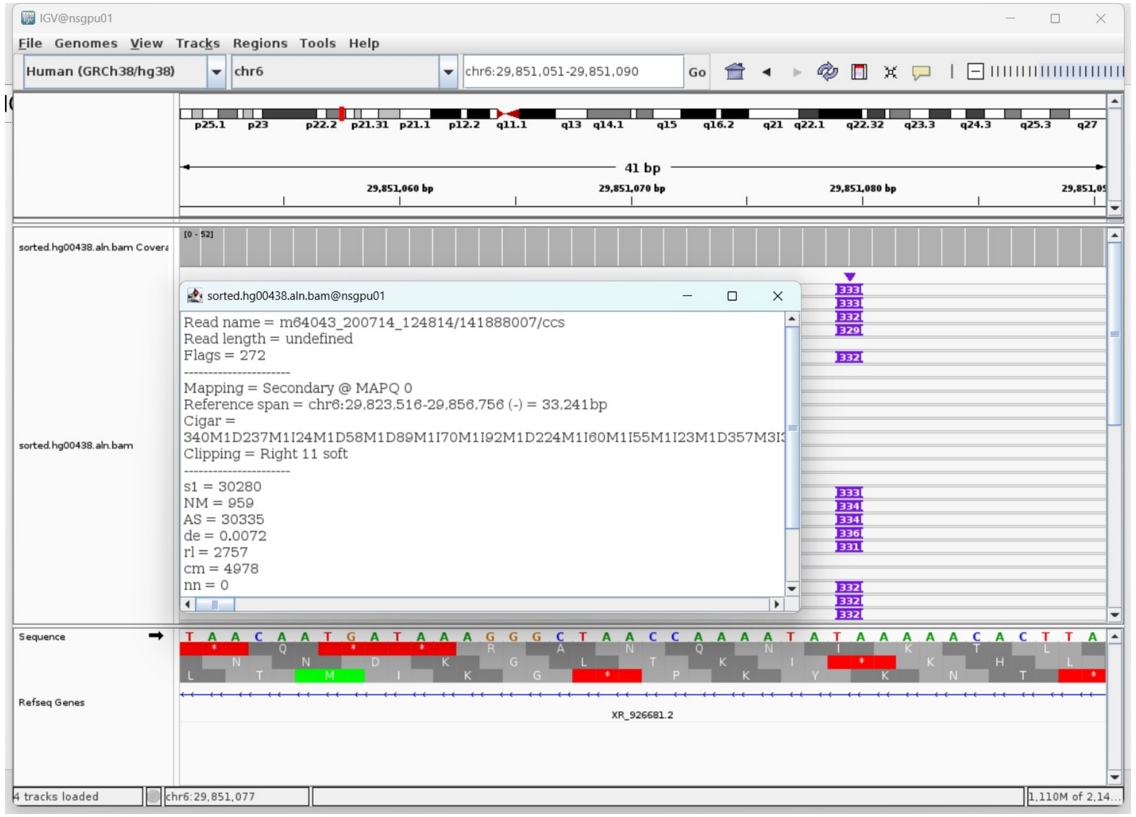


- Fetch one read (m64043_200710_174426/117966428/ccs, 30339 bp) and BLAT it to the hg38 genome.

The best alignment is to chr6_GL000252v2_alt (an alternative sequence of chr6) , which contains an *Alu*Yb8 element.

| ACTIONS | | | QUERY | SCORE | START | END | QSIZE | IDENTITY | CHROM | STRAND | START | END | SPAN |
|---|---|---|---|---|---|---|---|---|---|---|---|---|---|
| browser | new tab | details | YourSeq | 30284 | 1 | 30339 | 30339 | 100.0% | chr6_GL000252v2_alt | - | 1088707 | 1119056 | 30350 |
| browser | new tab | details | YourSeq | 30267 | 1 | 30339 | 30339 | 100.0% | chr6_GL000254v2_alt | - | 1088365 | 1118704 | 30340 |
| browser | new tab | details | YourSeq | 29883 | 1 | 30339 | 30339 | 99.8% | chr6_GL000253v2_alt | - | 1088381 | 1118386 | 30006 |
| browser | new tab | details | YourSeq | 29861 | 1 | 30339 | 30339 | 99.8% | chr6 | - | 29825894 | 29856264 | 30371 |

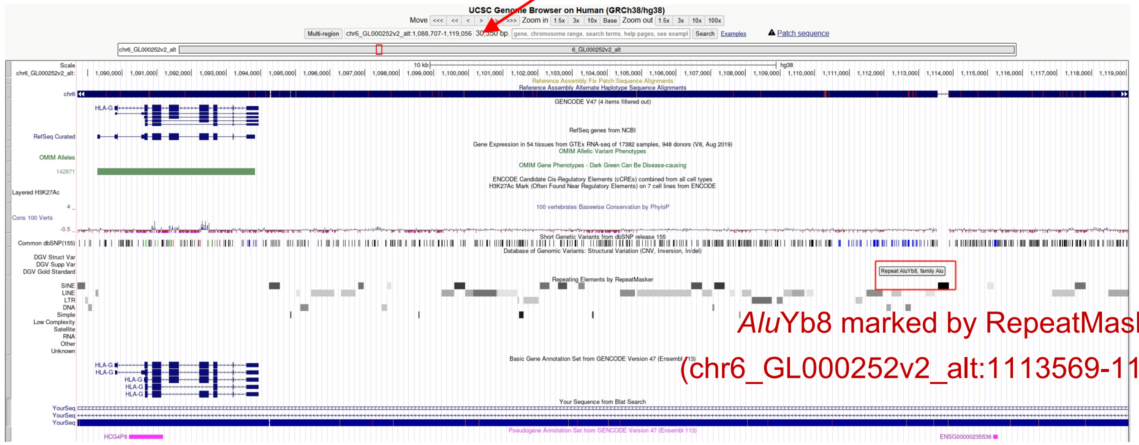


*Alu*Yb8 marked by RepeatMasker (chr6_GL000252v2_alt:1113569-1113875)

- Read:5223-5501 aligns to *Alu*Yb8:289-11, 99.64% identity

TGAGACGGAGTCTCGCTCTGTCGCCCAGGTCGGACTGCGGACTGCAGTGGCGCAATCTCGGCTCACTGCAAGCTCCG
CTTCCCGGGTTCACGCCATTCTCCTGCCTCAGCCTCCCGAGTAGCTGGGACTACAGGCGCCCGCCACCGCGCCCGGC
TAATTTTTTGTATTTTTAGTAGAGACGGGGTTTCACCTTGTTAGCCAGGATGGTCTCGATCTCCTGACCTCATGATC
CACCCGCCTCGGCCTCCCAAAGTGCTGGGATTACAGGCGTGAGCCAC

- It's an *Alu*Yb8 in the chromosome chr6_GL000252v2_alt.

- Illustration:

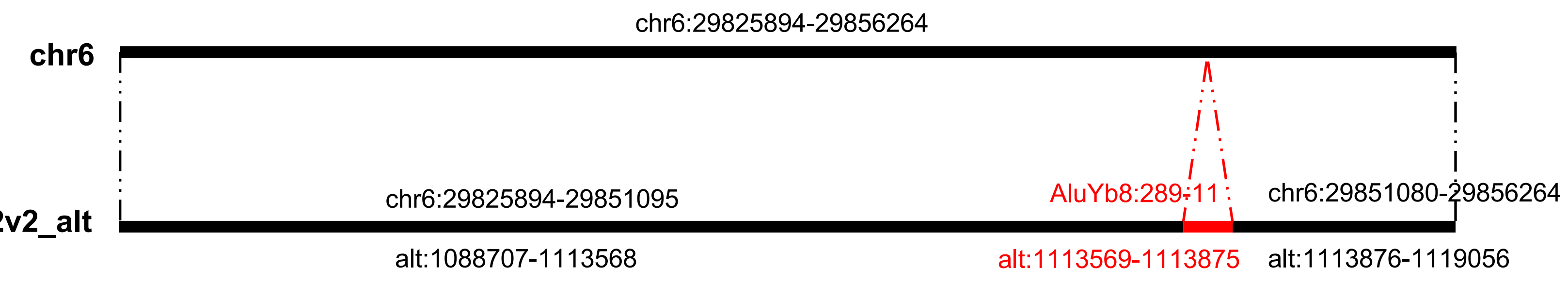

### HG00438 chr6:31471215

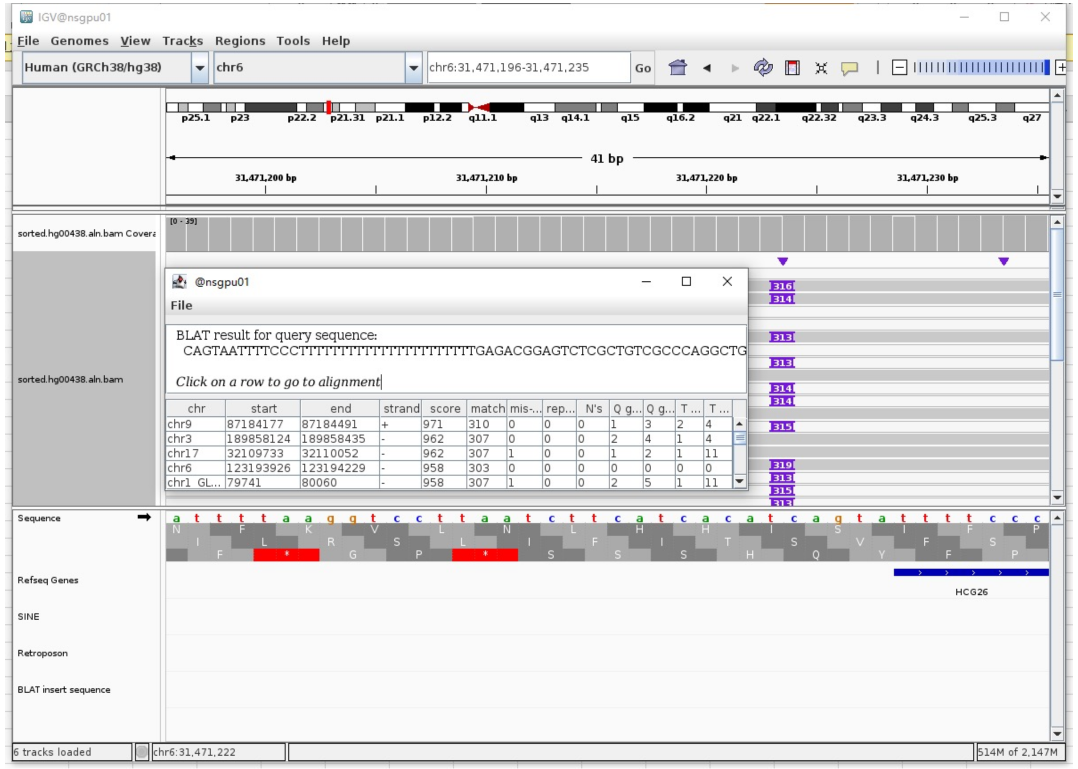


Insertion sequence (316bp)

- 5 bp TSD

CAGTA

- Insertion:6-33, polyT

ATTTTCCCTTTTTTTTTTTTTTTTTTTTTT

- Insertion:34-316 aligns to *Alu*Yc1:282-1, 97.85% identity

TGAGACGGAGTCTCGCTGTCGCCCAGGCTGGAGTGCAGTGGCGCAATCTCGGCTCACTGCAGG
CTCCGCCCCCTGGGGTTCACGCCATTCTCCTGCCTCAGCCTCCCGAGTAGCTGGGACTACAGG
CGCCCGCCACCTCGCCCGGCTAATTTTTTTTGTATTTTTAGTAGAGACGGGGTTTCACCGTGT
TAGCCAGGATGGTCTCGATCTCCTGACCTCGTGATCCGCCCGCCTCGGCCTCCCAAAGTGCTG
GGATTACAGGCGTGAGCCACCGCGCCCGGC

- Illustration:

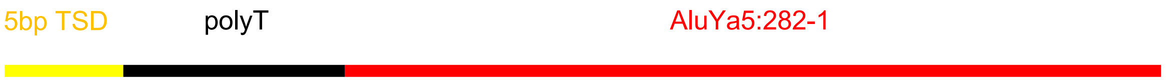

### HG00438 chr6:32805641

➢ Mapping quality is 0, indicating that this region may have alignment issues.

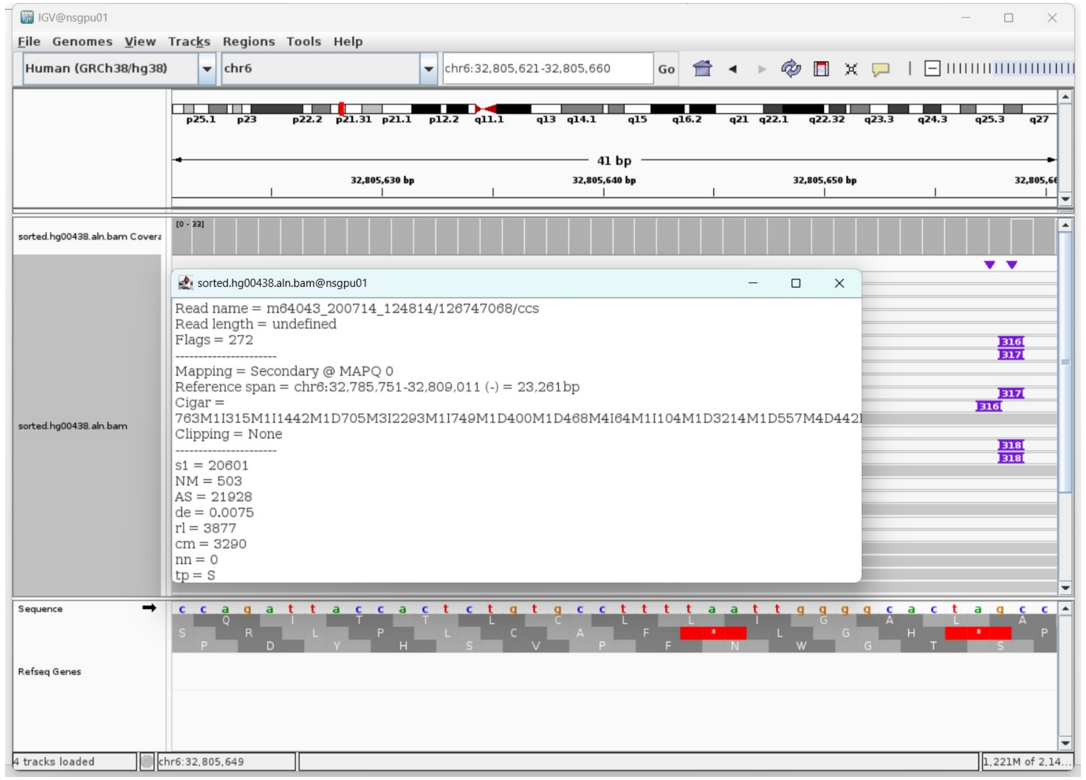


➢ Fetch one read (m64043_200714_124814/126747068/ccs, 23584 bp) and BLAT it to the hg38 genome.

The best alignment is to chr6_GL000252v2_alt (an alternative sequence of chr6) , which contains an *Alu*Ya5 element.

| ACTIONS | | | QUERY | SCORE | START | END | QSIZE | IDENTITY | CHROM | STRAND | START | END | SPAN |
|---|---|---|---|---|---|---|---|---|---|---|---|---|---|
| browser | new tab | details | YourSeq | 23520 | 1 | 23584 | 23584 | 100.0% | chr6_GL000252v2_alt | - | 4030203 | 4053781 | 23579 |
| browser | new tab | details | YourSeq | 23400 | 1 | 23584 | 23584 | 99.7% | chr6_GL000256v2_alt | - | 4185900 | 4209479 | 23580 |
| browser | new tab | details | YourSeq | 23347 | 1 | 23584 | 23584 | 99.6% | chr6_GL000250v2_alt | - | 4091508 | 4115091 | 23584 |
| browser | new tab | details | YourSeq | 23012 | 1 | 23584 | 23584 | 99.5% | chr6_GL000254v2_alt | - | 4084831 | 4108103 | 23273 |
| browser | new tab | details | YourSeq | 23005 | 1 | 23584 | 23584 | 99.5% | chr6_GL000251v2_alt | - | 4199023 | 4222271 | 23249 |
| browser | new tab | details | YourSeq | 22940 | 1 | 23584 | 23584 | 99.3% | chr6_GL000253v2_alt | - | 4205097 | 4228364 | 23268 |
| browser | new tab | details | YourSeq | 22922 | 1 | 23584 | 23584 | 99.3% | chr6 | - | 32785751 | 32809011 | 23261 |

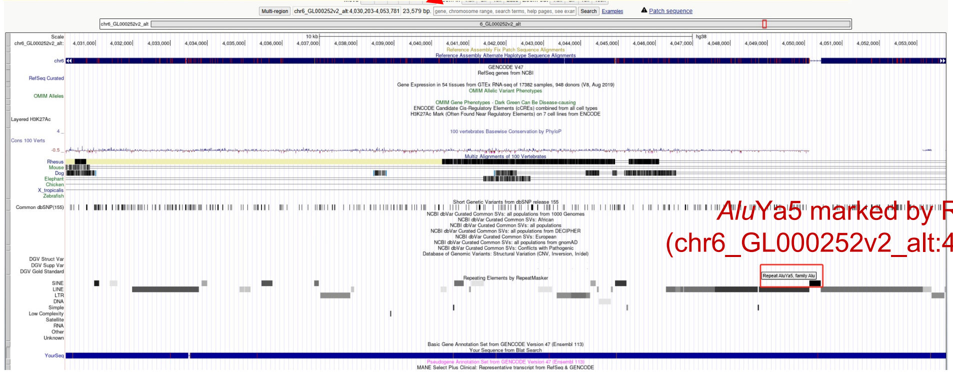

*Alu*Ya5 marked by RepeatMasker (chr6_GL000252v2_alt:4050120-4050430)

➢ Read:3352-3634 aligns to *Alu*Ya5:1-282, 98.94% identity

GGCCGGGCACGGTGGCTCACGCCTGTAATCCCAGCACTTTGGGAGGCCGAGGCGGGCGGATCACGAGGTCAGGAGAT
CGAGACGATCCCGGCTAAAACGGTGAAACCCCGTCTCTACTAAAAATACAAAAAATTAGCCGGGCGTAGTGGCAGGC
GCCTGTAGTCCCAGCTACTTGGGAGGCTGAGGCAGGAGAATGGCGTGAACCCGGGAGGCGGAGCTTGCAGTGAGCCG
AGATCCCGCCACTGCACTCCAGCCTGGGCGACAGAGCGAGACTCCGTCTCA

➢ It's an *Alu*Ya5 in the chromosome chr6_GL000252v2_alt.

- Illustration (*Alu* in chr6_GL000252v2_alt):

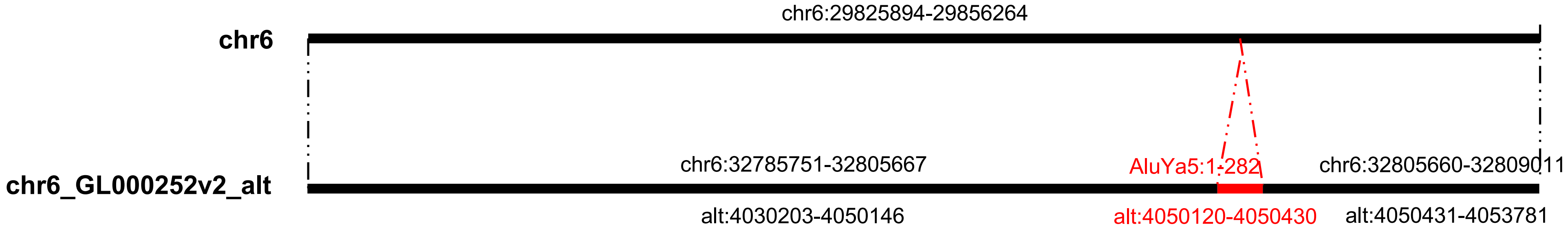

### HG00438 chr6:104041958

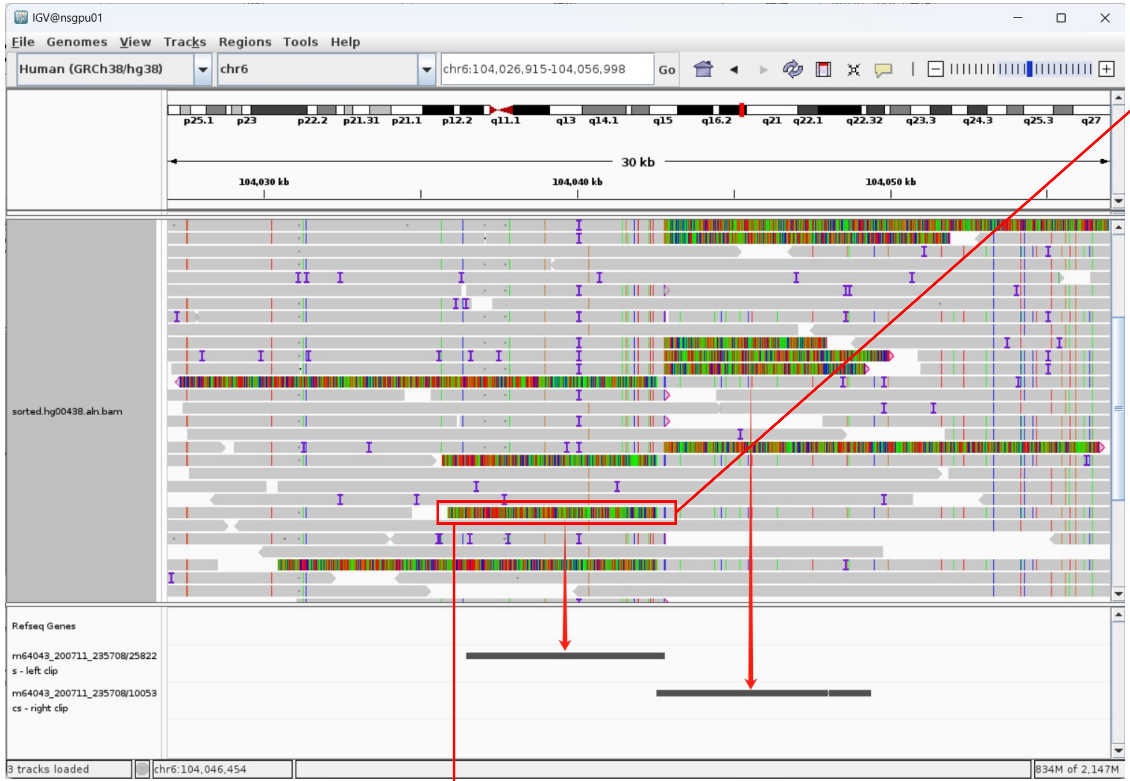


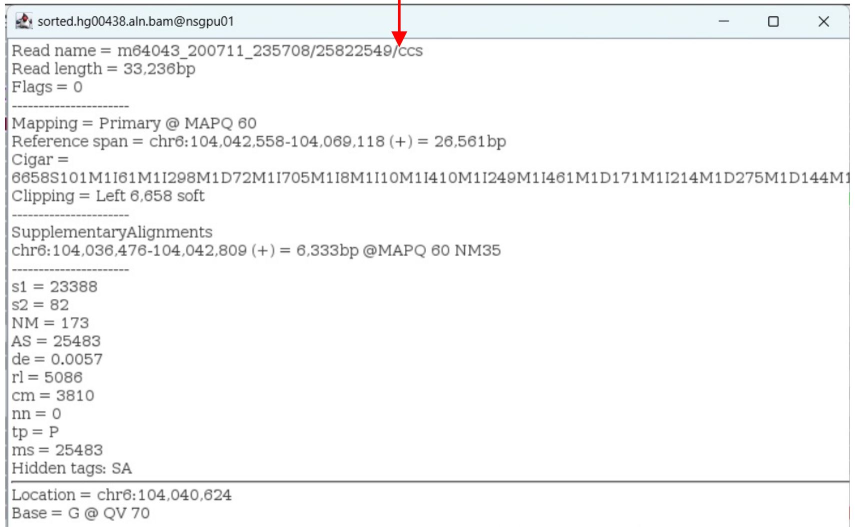
sorted.hg00438.aln.bam@nsgpu01

Read name = m64043_200711_235708/25822549/ccs
Read length = 33,236bp
Flags = 0
--------------------
Mapping = Primary @ MAPQ 60
Reference span = chr6:104,042,558-104,069,118 (+) = 26,561bp
Cigar =
6658S101M1I61M1I298M1D72M1I705M1I8M1I10M1I410M1I249M1I461M1D171M1I214M1D275M1D144M1
Clipping = Left 6,658 soft
--------------------
SupplementaryAlignments
chr6:104,036,476-104,042,809 (+) = 6,333bp @MAPQ 60 NM35
--------------------
s1 = 23388
s2 = 82
NM = 173
AS = 25483
de = 0.0057
rl = 5086
cm = 3810
nn = 0
tp = P
ms = 25483
Hidden tags: SA
--------------------
Location = chr6:104,040,624
Base = G @ QV 70

Left-clipped read (6658bp)

- 239bp TSD (same as chr6:104042558-104042797)

AAGAAACACCATTCTGGACATTGGCCTTGAGAAAGAATTTATCATTAAGTGCTCAAAAG
CAATTGCAACAAAAACAAAATTTGACAAGTCAGACCTAATTCAACAAAAGAGCTTGTGC
ACAACAATAGAAACAGAGTAGACAGATAACCTGTAGGATGGGAGAAAATATTAACAAAC
CATGCATTCAACAAAGGCCTCATATCCAGAGTATATAAAGAACTTAATCCAACAAGCAA
AAAA

| ACTIONS | | | QUERY | SCORE | START | END | QSIZE | IDENTITY | CHROM | STRAND | START | END | SPAN |
|---|---|---|---|---|---|---|---|---|---|---|---|---|---|
| browser | new tab | details | YourSeq | 6284 | 1 | 6343 | 6343 | 99.5% | chr6 | + | 104036476 | 104042808 | 6333 |
| browser | new tab | details | YourSeq | 664 | 4559 | 6262 | 6343 | 79.0% | chr14 | - | 28685291 | 28686613 | 1323 |

- Clip:6343-6626 aligns to *Alu*Ya5:1-282, 97.16% identity

CAACCTCTTTAGGCTGGGCGCAGTGGCTCACGCCTGTAATCCCAGCACTTTGGGAGGCC
GAGGCGGGCGGATCACGAGGTCAGGAGATCGAGACCATCCTGGCTAAAACGGTGAAACC
CCGTCTCTACTAAAAATACAAAAAATTAGCCGGGCGTAGTGGCGGGCGCCTGTAGTCCC
AGCTACTCGGGAGGCTGAGGCAGGAGAATGGCGTGAACCCGGGAGGCAGAGCTTGCAGT
GAGCCGAGATCAGGCCACTGCACTCTAGCCTGGGCGACAGAGCGAGACTCCGTCTCA

- Clip: 6627-6658, polyA AAAAAAAAAAAAAAAAAAAAAAAAAAAAAAAA

- Illustration:

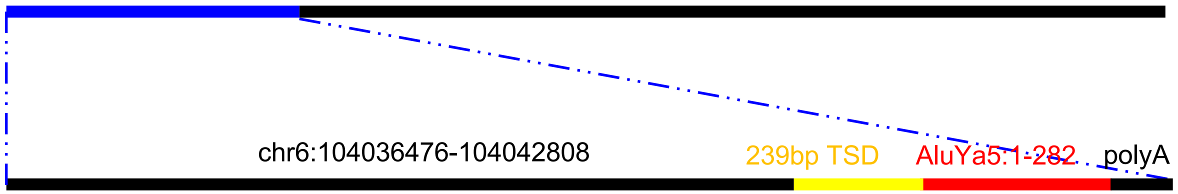

### HG00438 chr14:93364572

- Mapping quality is 0, indicating that this region may have alignment issues.

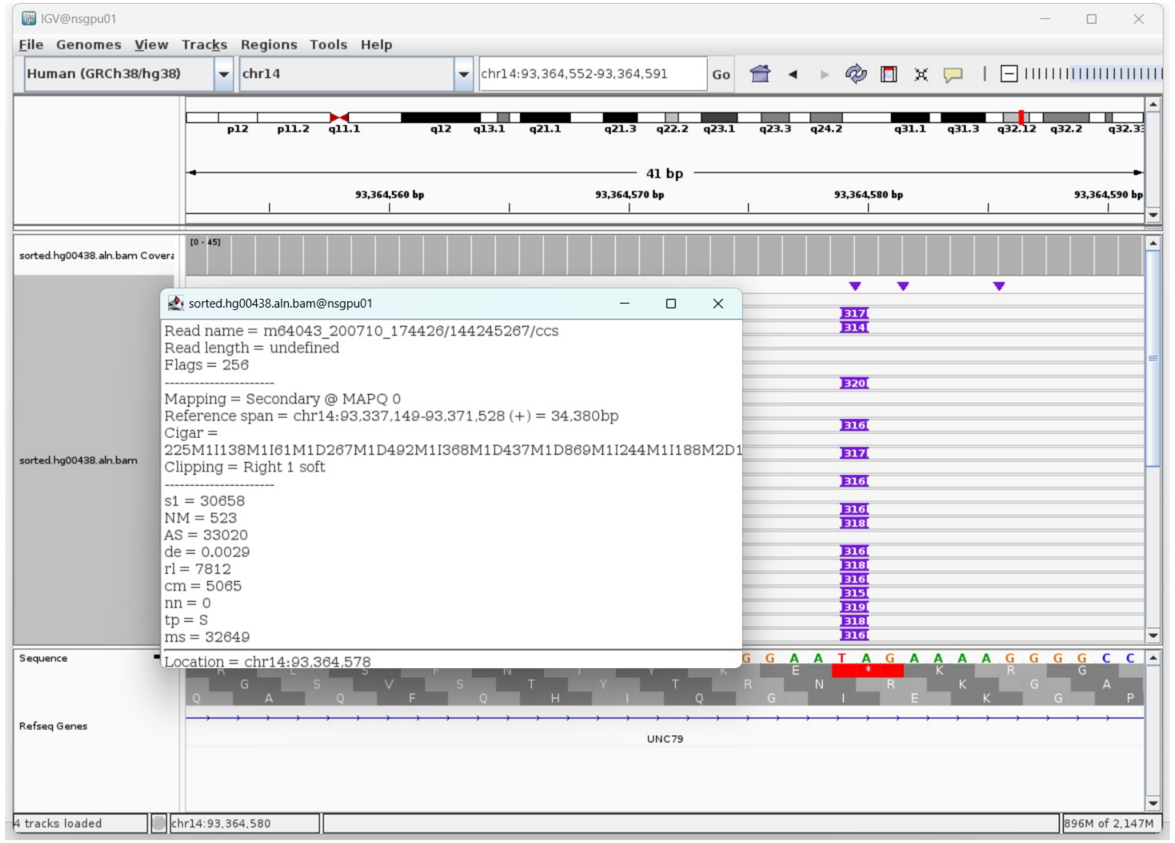


- Fetch one read (m64043_200714_124814/121833366/ccs, 20476 bp) and BLAT it to the hg38 genome.

  The best alignment is to chr14_KI270847v1_alt (an alternative sequence of chr14) and chr14.

| QUERY | SCORE | START | END | QSIZE | IDENTITY | CHROM | STRAND | START | END | SPAN |
|---|---|---|---|---|---|---|---|---|---|---|
| YourSeq | 20082 | 1 | 20476 | 20476 | 99.9% | chr14_KI270847v1_alt | - | 462950 | 483216 | 20267 |
| YourSeq | 20082 | 1 | 20476 | 20476 | 99.9% | chr14 | - | 93348388 | 93368654 | 20267 |

- Read:1-4071 aligns to chr14_KI270847v1_alt and chr14

| QUERY | SCORE | START | END | QSIZE | IDENTITY | CHROM | STRAND | START | END | SPAN |
|---|---|---|---|---|---|---|---|---|---|---|
| YourSeq | 4064 | 1 | 4071 | 4089 | 100.0% | chr14_KI270847v1_alt | - | 479142 | 483216 | 4075 |
| YourSeq | 4064 | 1 | 4071 | 4089 | 100.0% | chr14 | - | 93364580 | 93368654 | 4075 |

- Read:4072-4088, polyT

  TTTTTTTTTTTTTTTTTT

- Read:4089-4371 aligns to *Alu*Yc1:282-1, 98.94% identity

  TGAGACGGAGTCTCACTCTGTCGCCCAGGCTGGAGTGCAGTGGCACGATCTCGGCTCACTGCAAGCTCCGCCTCC
  CGGGTTCACGCCATTCTCCTGCCTCAGCCTCCCGAGTAGCTGGGACTACAGGCGCCCGCTACAACGCCCGGCTAA
  TTTTTTGTATTTTTAGTAGAGACGGGGTTTCACCGTGTTAGCCAGGATGGTCTCGATCTCCTGACCTCGTGATCC
  GCCCGCCTCGGCCTCCCAAAGTGCTGGGATTACAGGCGTGAGCCACCGCGCCCGGCC

- Read:4372- 20476 also aligns to chr14_KI270847v1_alt and chr14

| QUERY | SCORE | START | END | QSIZE | IDENTITY | CHROM | STRAND | START | END | SPAN |
|---|---|---|---|---|---|---|---|---|---|---|
| YourSeq | 16036 | 1 | 16105 | 16105 | 99.9% | chr14_KI270847v1_alt | - | 462950 | 479157 | 16208 |
| YourSeq | 16036 | 1 | 16105 | 16105 | 99.9% | chr14 | - | 93348388 | 93364595 | 16208 |

- It's an *Alu*Yc1

- Illustration:

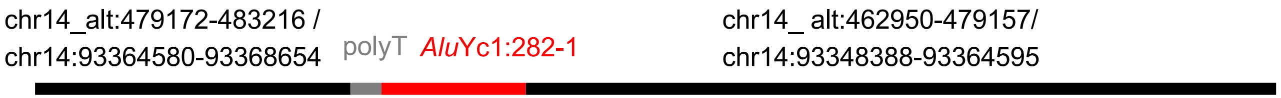

### HG00438 chr15:32821084

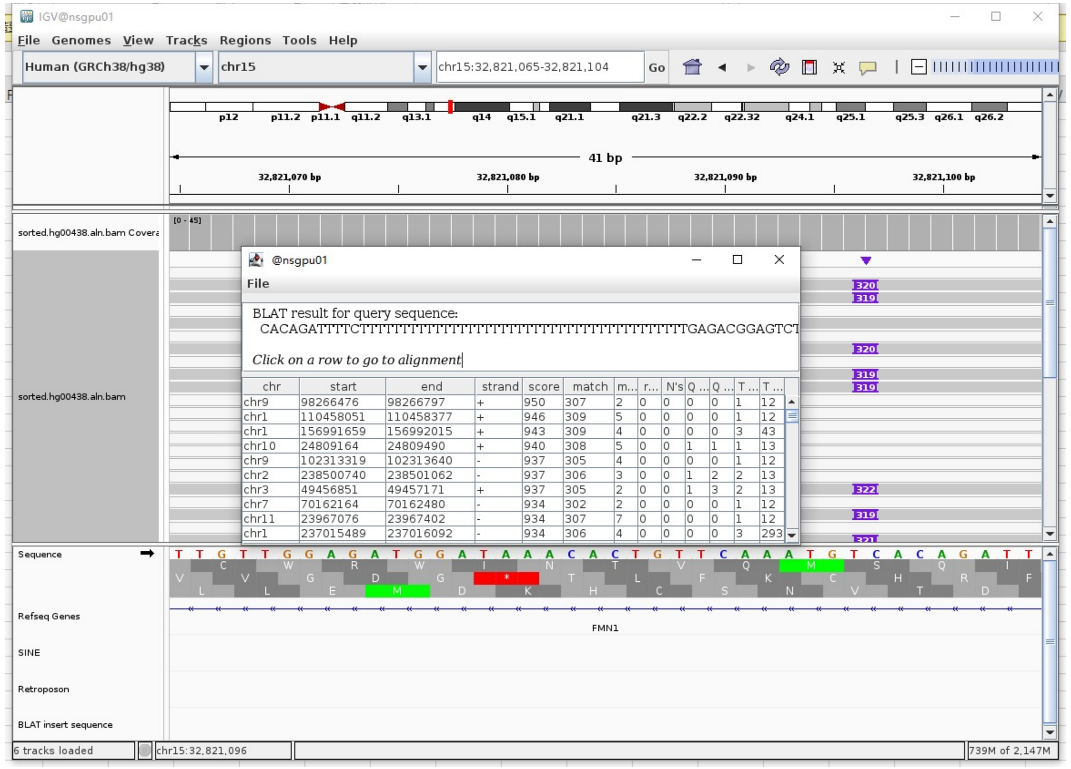


Insertion sequence (320)

- Insertion:1-12, 12bp **TSD**

CACAGATTTTCT

- Insertion:13-49, polyT

TTTTTTTTTTTTTTTTTTTTTTTTTTTTTTTTTTTTT

- Insertion:50-320 aligns to *Alu*Ya5:282-1, 98.89% identity

TGAGACGGAGTCTCGCTCTGTCGCCCAGGCTGGACTGCAGTGGCGGGATCTCGGCTCACT
GCAAGCTCCGCCTCCCGGGTTCATGCCATTCTCCTGCCTCAGCCTCCCAAGTAGCTGGGA
CTACAGGCGCCCGCCACTACGCCCGGCTAATTTTTTGCATTTTTAGTAGAGACGGGGTTT
CACCGTGGTCTCGATCTCCTGACCTCGTGATCCGCCCGCCTCGGCCTCCCAAAGTGCTGG
GATTACAGGCGTGAGCCACCGCGCCCGGCC

- Illustration:

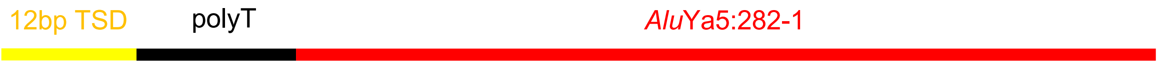

### HG00438 chr18:51774994

- Mapping quality is 0, indicating that this region may have alignment issues.

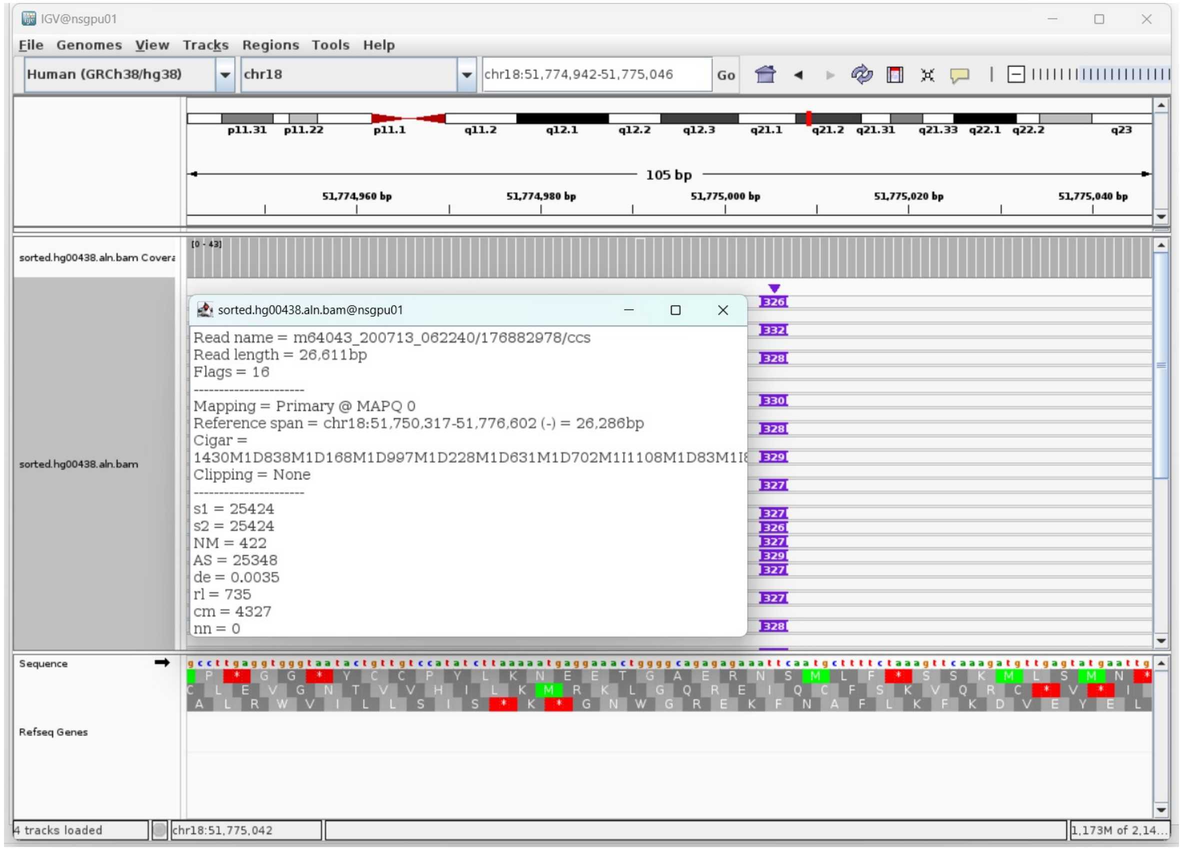


- Fetch one read (m64043_200713_062240/176882978/ccs, 26613 bp) and BLAT it to the hg38 genome.

The best alignment is to chr18_GL383570v1_alt (an alternative sequence of chr18) and chr18.

| ACTIONS | QUERY | SCORE | START | END | QSIZE | IDENTITY | CHROM | STRAND | START | END | SPAN |
|---|---|---|---|---|---|---|---|---|---|---|---|
| browser new tab details | YourSeq | 26150 | 1 | 26613 | 26613 | 99.7% | chr18_GL383570v1_alt | - | 45041 | 71326 | 26286 |
| browser new tab details | YourSeq | 26150 | 1 | 26613 | 26613 | 99.7% | chr18 | + | 51750317 | 51776602 | 26286 |

- Read:1-24734 aligns to chr18_GL383570v1_alt and chr18

| ACTIONS | QUERY | SCORE | START | END | QSIZE | IDENTITY | CHROM | STRAND | START | END | SPAN |
|---|---|---|---|---|---|---|---|---|---|---|---|
| browser new tab details | YourSeq | 24575 | 1 | 24703 | 24734 | 99.8% | chr18_GL383570v1_alt | - | 46625 | 71326 | 24702 |
| browser new tab details | YourSeq | 24575 | 1 | 24703 | 24734 | 99.8% | chr18 | + | 51750317 | 51775018 | 24702 |

- Read:24734-25017 aligns to *Alu*Ya5:282-1, 99.65% identity

TGAGACGGAGTCTCGCTCTGTCGCCCAGGCTGGAGTGCAGTGGCGGATCTCGGCTCACTGCAAGCTCCGCCTCC
CGGGTTCACGCCATTCTCCTGCCTCAGCCTCCCAAGTAGCTGGGACTACAGGCGCCCGCCACTACGCCCGGCTAA
TTTTTTTGTATTTTTAGTAGAGACGGGGTTTCACCATTTTAGCCGGGATGGTCTCGATCTCCTGACCTCGTGATC
CGCCCGCCTCGGCCTCCCAAAGTGCTGGGATTACAGGCGTGAGCCACCGCGCCCGG

13bp TSD

TCAATGCTTTTCT

- Read:25015-26613 also aligns to chr18_GL383570v1_alt and chr18

| ACTIONS | QUERY | SCORE | START | END | QSIZE | IDENTITY | CHROM | STRAND | START | END | SPAN |
|---|---|---|---|---|---|---|---|---|---|---|---|
| browser new tab details | YourSeq | 1589 | 3 | 1598 | 1598 | 99.9% | chr18_GL383570v1_alt | - | 45041 | 46637 | 1597 |
| browser new tab details | YourSeq | 1589 | 3 | 1598 | 1598 | 99.9% | chr18 | + | 51775006 | 51776602 | 1597 |

- It's an *Alu*Ya5 insertion.

- Illustration:

chr18_alt:71326-46625 / chr18:51750317-51775018 AluYa5:282-1 13bp TSD chr18_ alt:46636-45041/ chr18:51775006-51776602

### HG00621 chr4:21532942

- ➢ Using the built-in BLAT tool in IGV to check where the inserted sequence aligned.

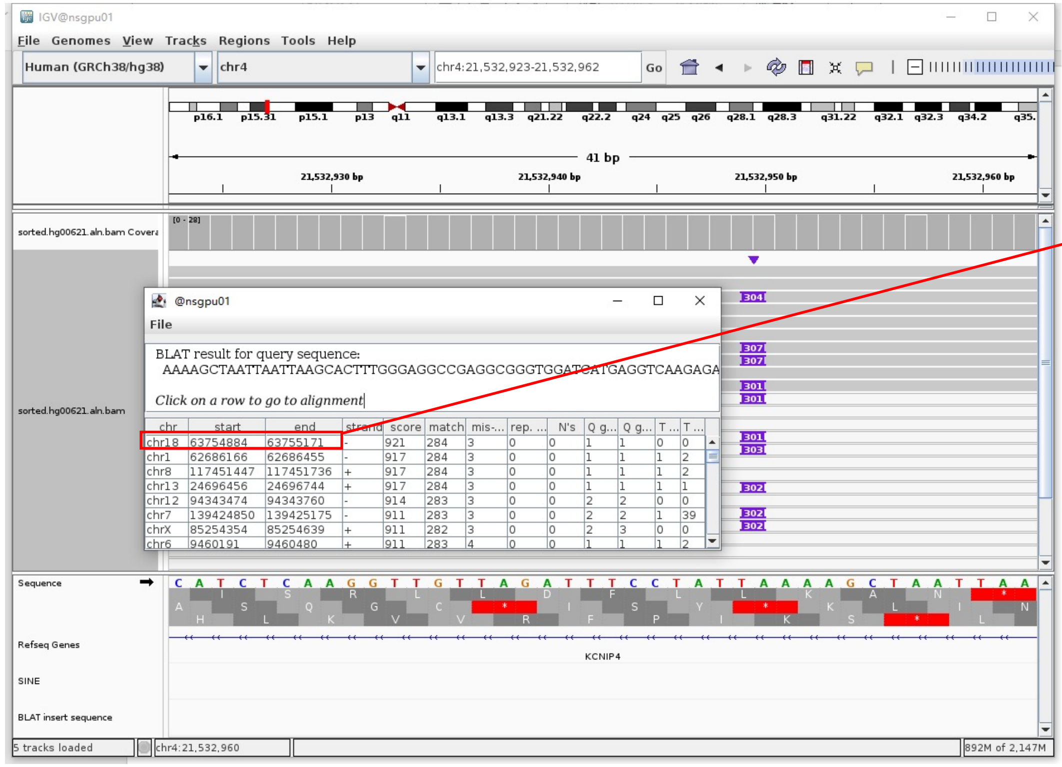


- ➢ The annotation at the aligned location indicated it as *Alu*Y.

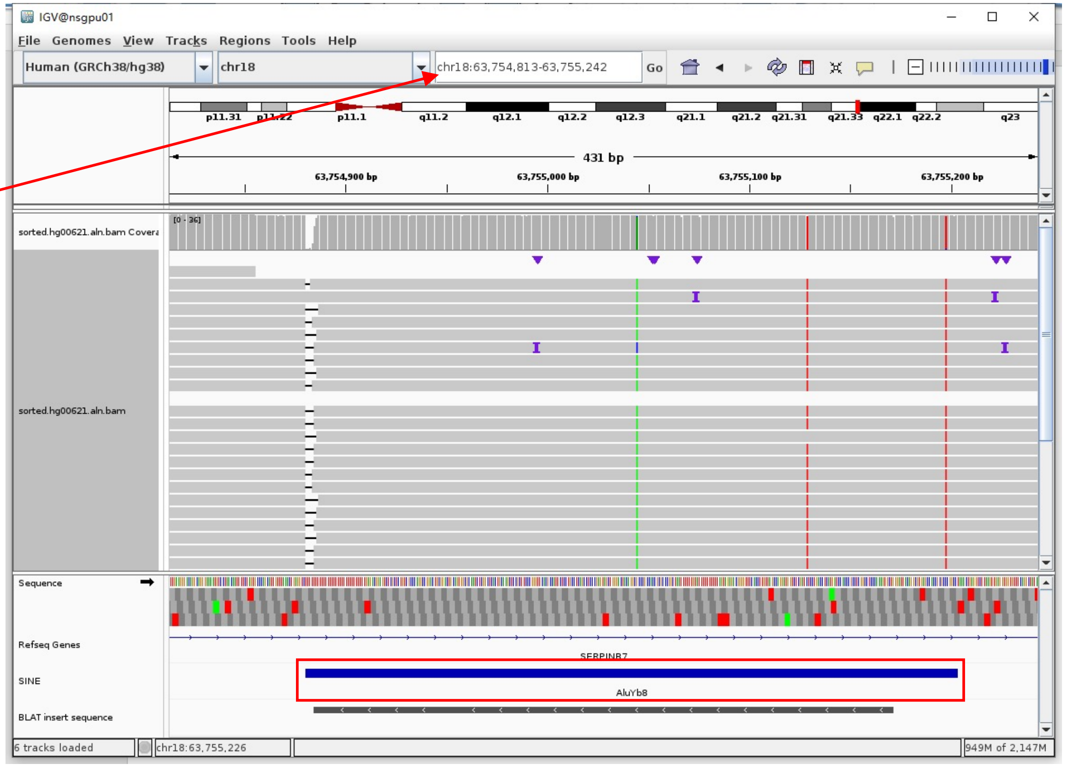


Insertion sequence (301bp)

- ➢ 17bp TSD

AAAAGCTAATTAATTAA

- ➢ Insertion:16-279 aligns to *Alu*Yb8:32-289, 98.83% identity

AGCACTTTGGGAGGCCGAGGCGGGTGGATCATGAGGTCAAGAGATCGAGACCATCCTGGCTAACAAGGTGAAACCCCGTCTCTACTAAAAAAAAAATACAAAAAATTAGCTGGGCGCGGTGGC
GGGCGCCTGTAGTCCCAGCTACTCGGGAGGCTGAGGCAGGAGAATGGCGTGAACCCGGGAAGCGGAGCTTGCAGTGAGCGGAGATTGCGCCACTGCAGTCCCGCAGTCCGGCCTGGGCGACAG
AGCGAGACTCCGTCTCAAAAAAAAAAAAAAAAAAAAAAAAAA

### HG00621 chr6:29851067

- The same genomic coordinate as in HG00438 (page 3)

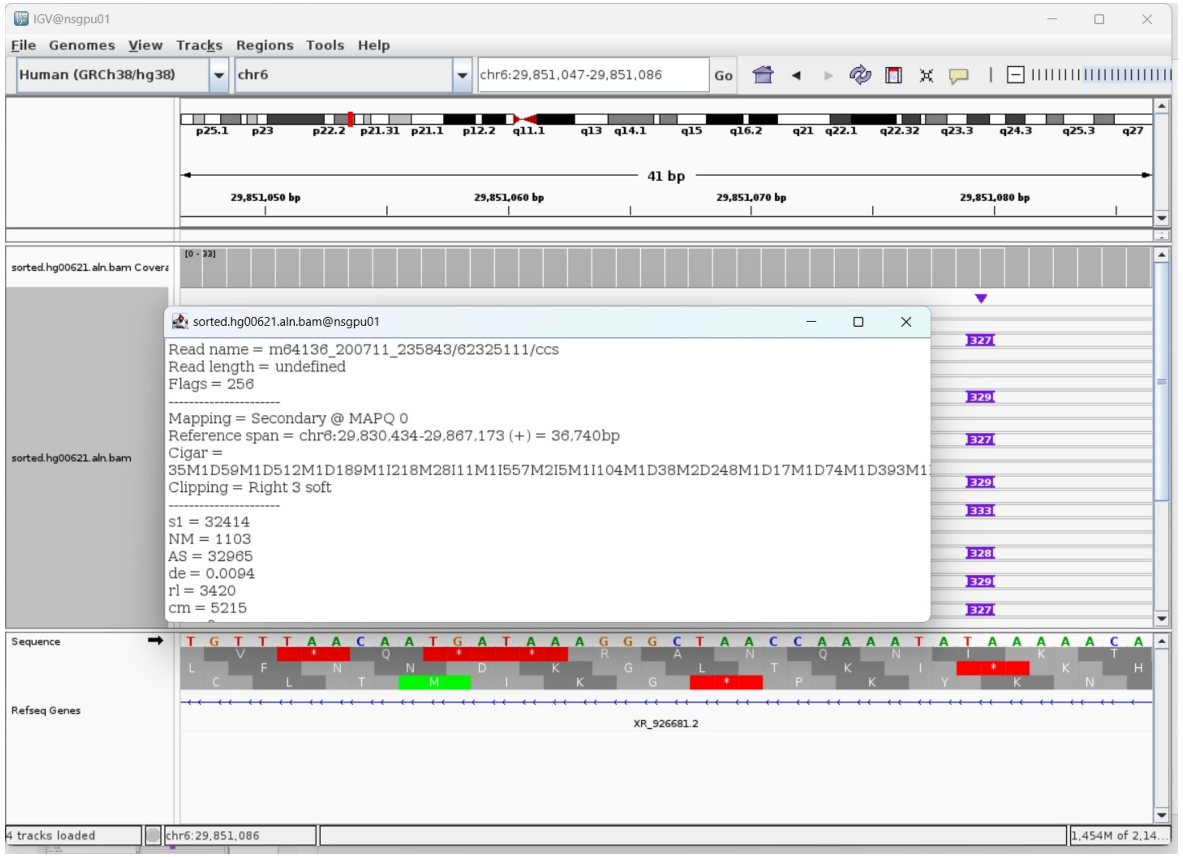

### HG00621 chr12:11026415

- Mapping quality is 0, indicating that this region may have alignment issues.

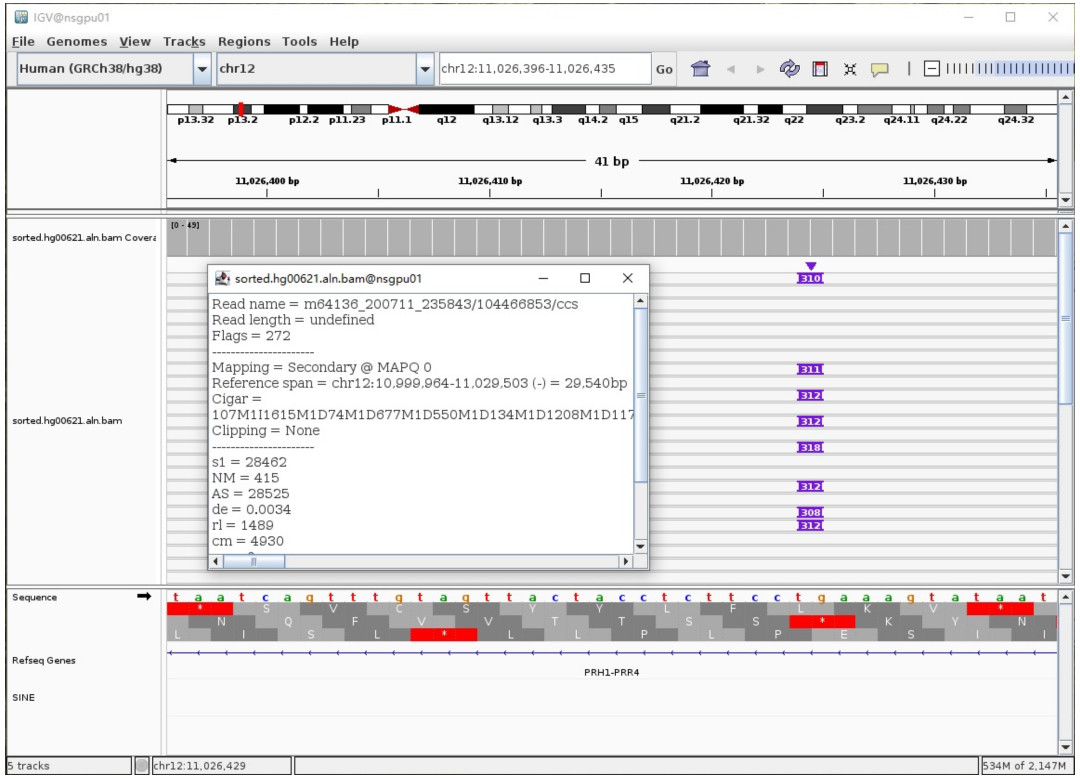


- Fetch one read (m64136_200711_235843/104466853/ccs, 23584 bp) and BLAT it to the hg38 genome.

  The best alignment is to chr12_KI270904v1_alt (an alternative sequence of chr12) , which contains an *Alu*Y element.

| ACTIONS | | | QUERY | SCORE | START | END | QSIZE | IDENTITY | CHROM | STRAND | START | END | SPAN |
|---|---|---|---|---|---|---|---|---|---|---|---|---|---|
| browser | new tab | details | YourSeq | 29810 | 1 | 29856 | 29856 | 100.0% | chr12_KI270904v1_alt | - | 198662 | 228508 | 29847 |
| browser | new tab | details | YourSeq | 29394 | 1 | 29856 | 29856 | 99.8% | chr12_GL877876v1_alt | - | 198673 | 228209 | 29537 |
| browser | new tab | details | YourSeq | 29394 | 1 | 29856 | 29856 | 99.8% | chr12 | - | 10999967 | 11029503 | 29537 |

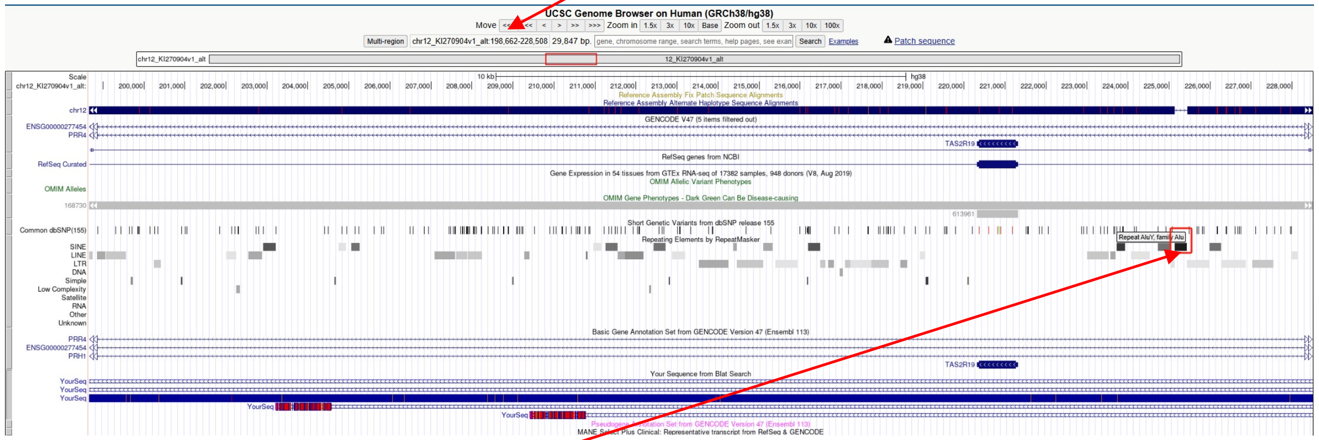


chr12_KI270904v1_alt:225135-225432 (*Alu*Y marked by RepeatMasker)

- Read:3106-3379 aligns to *Alu*Yc1:282-16, 98.12% identity

TGAGACAGAGTCTCGCTCTGTCGCCCAGGCTGGAGTGCAGTGGCGCGATCTCGGCTCACTGCAAGCTCCGCCTCCCGG
GTTCACGCCATTCTCCTGCCTCAGCCTCCGGAGTAGCTGGGACTACAGGCGCCCGCTACCACGCCCGGCTAATTTTTT
GTATTTTTTTTTTTTTAGTAGAGACGGGGTTTCACCATGTTAGCCAGGATGGTCTCGATCTCCTGACCTCGTGATCCGC
CCGCCTCGGCCTCCCAAAGTGCTGGGATTACAGGCTTCA

- It's an *Alu*Y in the chromosome chr12_KI270904v1_alt.

### HG00621 chr13:112339461

- Using the built-in BLAT tool in IGV to check where the inserted sequence aligned.
- The annotation at the aligned location indicated it as *Alu*Y.

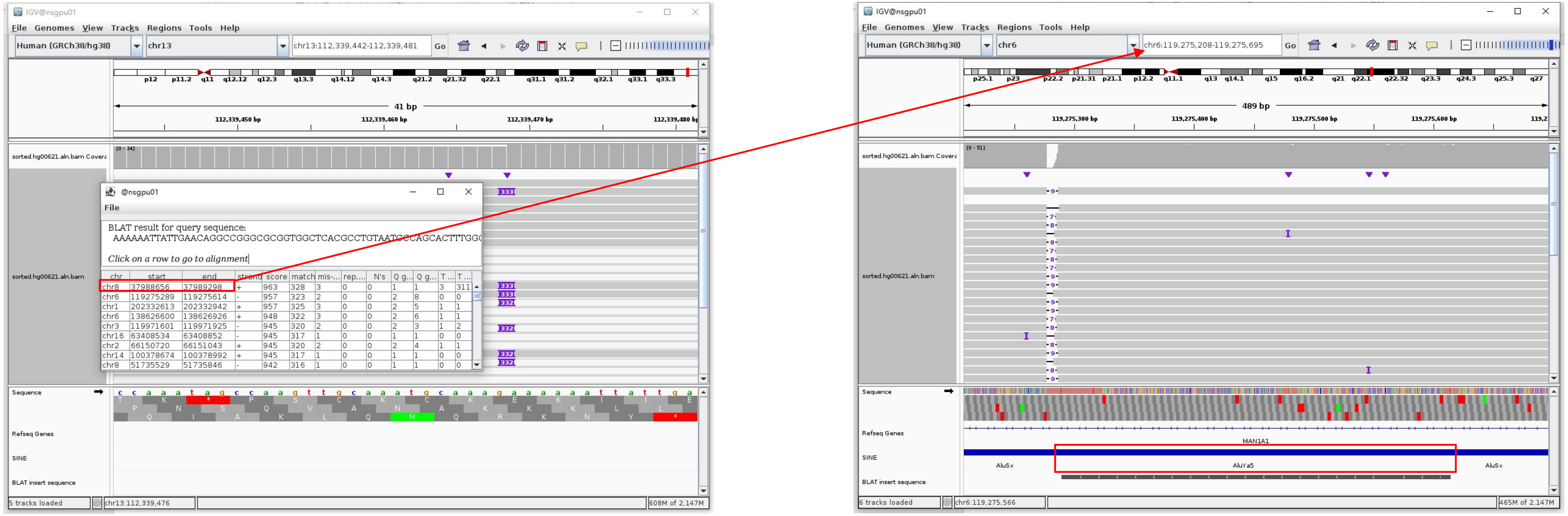


Insertion sequence (332bp)

- 17bp TSD

AAAAAATTATTGAACAG

- Insertion:16-299 aligns to *Alu*Ya5:2-282, 99.65% identity

GGCCGGGCGCGGTGGCTCACGCCTGTAATCCCAGCACTTTGGGAGGCCCGAGGCGGGCGGATCACGAGGTCAGGAGATCGAGACCATCCCGGCTAAAACGGTGAAACCCCGTCTCTACTAAAA
ATACAAAAAATTAGCCGGGCGTAGTGGCGGGCGCCTGTAGTCCCAGCTACTTGGGAGGCTGAGGCAGGAGAATGGCGTGAACCCGGGAGGCGGAGCTTGCAGTGAGCCGAGATCGCGCCACTG
CACTCCAGCCTGGGCGACAGAGCGAGACTCCGTCTCA………poly(A)

### HG00621 chr15:32662471

- ➢ Using the built-in BLAT tool in IGV to check where the inserted sequence aligned.

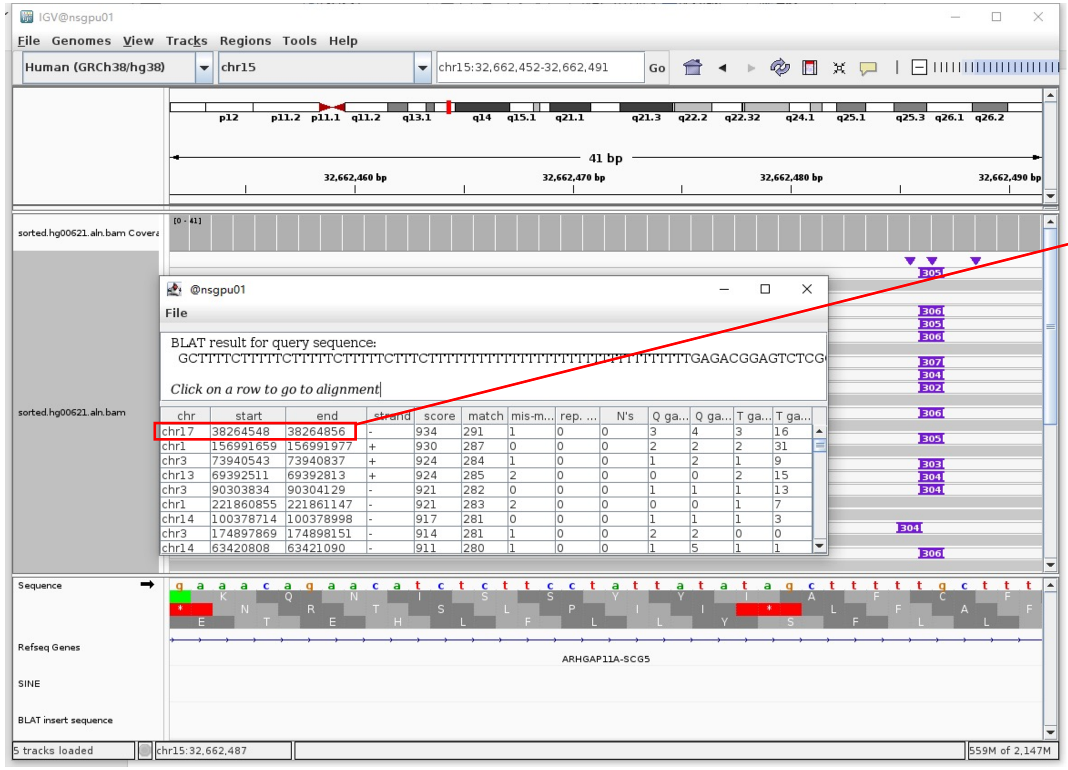


- ➢ The annotation at the aligned location indicated it as *AluY*.

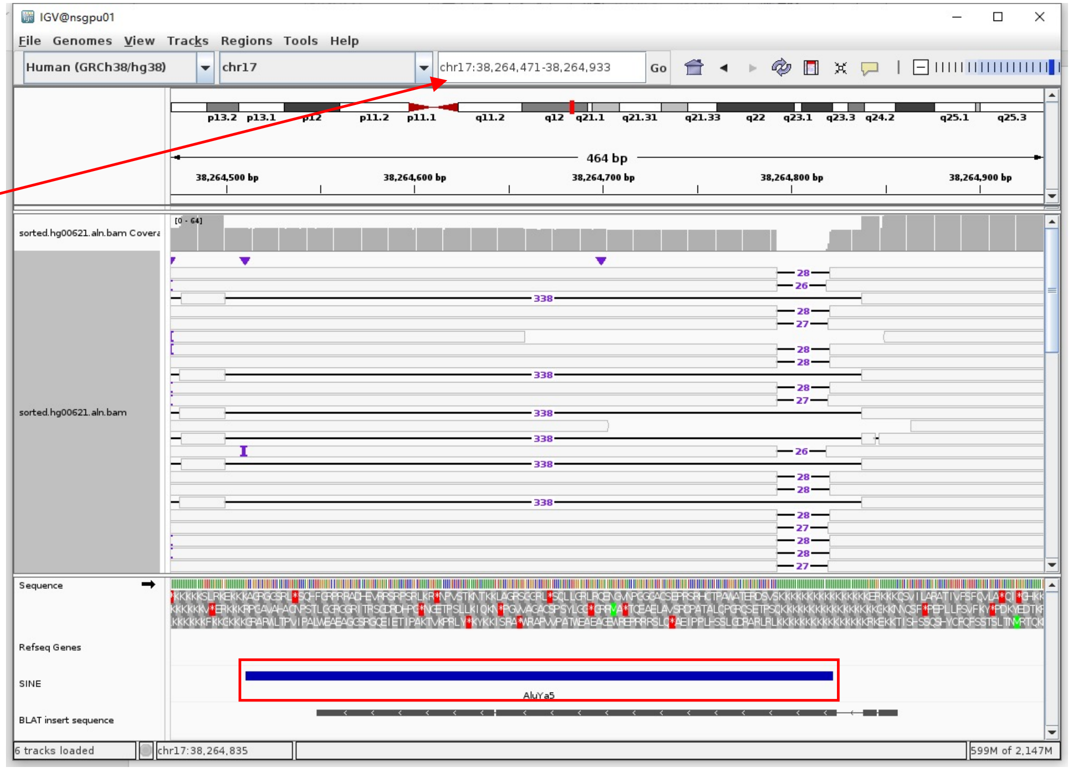


Insertion sequence (304bp)

- ➢ 9bp TSD

GCTTTTCTT

- ➢ Insertion:60-304 aligns to *AluYa5*:282-38, 100% identity

TGAGACGGAGTCTCGCTCTGTCGCCCAGGCTGGAGTGCAGTGGCGGGATCTCGGCTCACTGCAAGCTCCGCCTCCCGGGTTCACGCCATTCTCCTGCCTCAGCCTCCCAAGTAGCTGGGACTA
CAGGCGCCCGCCACTACGCCCGGCTAATTTTTTGTATTTTTAGTAGAGACGGGGTTTCACCGTTTTAGCCGGGATGGTCTCGATCTCCTGACCTCGTGATCCGCCCGCCTCGGCCTCCCAA

### HG00621 chr17:37561902

- ➢ Using the built-in BLAT tool in IGV to check where the inserted sequence aligned.
- ➢ The annotation at the aligned location indicated it as *Alu*Y.

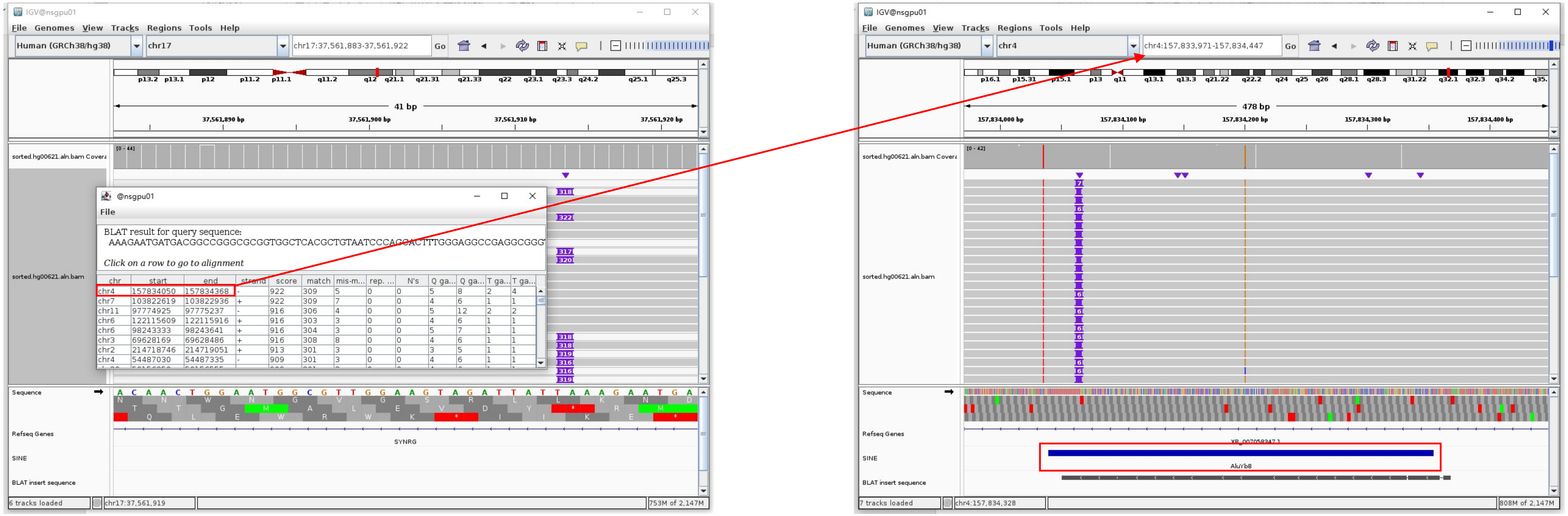


Insertion sequence (316bp)

- ➢ 9bp TSD

GCTTTTCTT

- ➢ Insertion:13-307 aligns to *Alu*Yb8:1-289, 98.96% identity

GGCCGGGCGCGGTGGCTCACGCTGTAATCCCAGCACTTTGGGAGGCCGAGGCGGGTGGATCATAAGGTCAGGAGATCGAGACCATCCTGGTTAACAATGTGAAACCCCGTCTCTACTAAAAAT
ACAAAAAAATTAGCCGGGCGCGGTGGCGGGCGCCTGTAGTCCCAGCTACTCGGGAGGCTGAGGCAGGGGGAGAATGGCGTGAACCCGGGAAGCGGAGCTTGCAGTGAGCCGAGATTGCGCCA
CTGCAGTCCGCAGTCCGGGCCTGGGCGACAGAGCGAGACTCCGTCTCA………poly(A)

### HG00621 chr18:51774994

- ➢ The same genomic coordinate as in HG00438 (page 9)

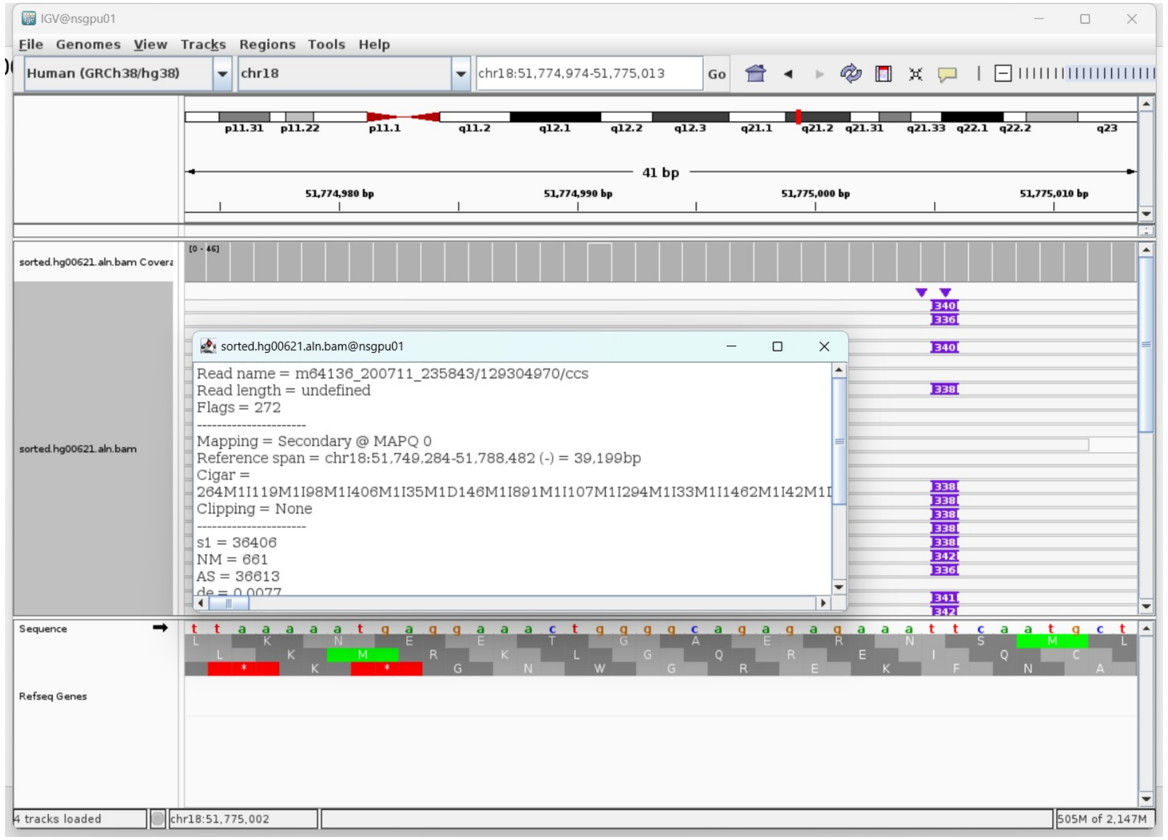

### HG00621 chr19:52384792

- ➢ Using the built-in BLAT tool in IGV to check where the inserted sequence aligned.

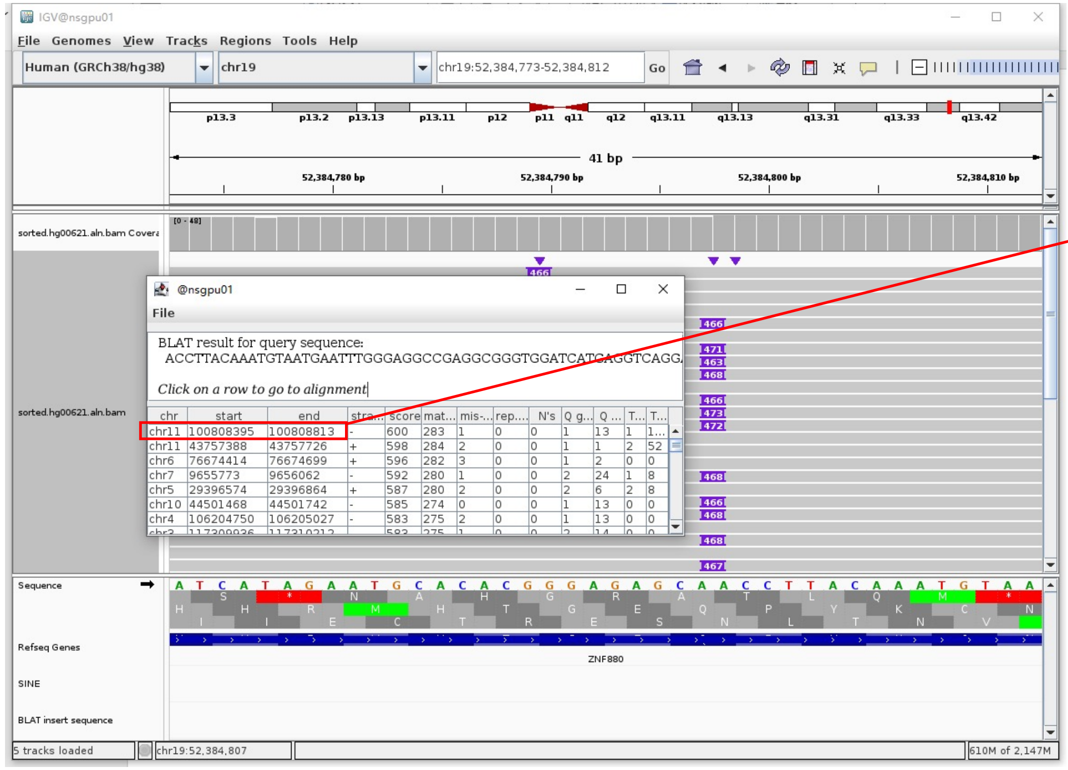


- ➢ The annotation at the aligned location indicated it as *Alu*Y.

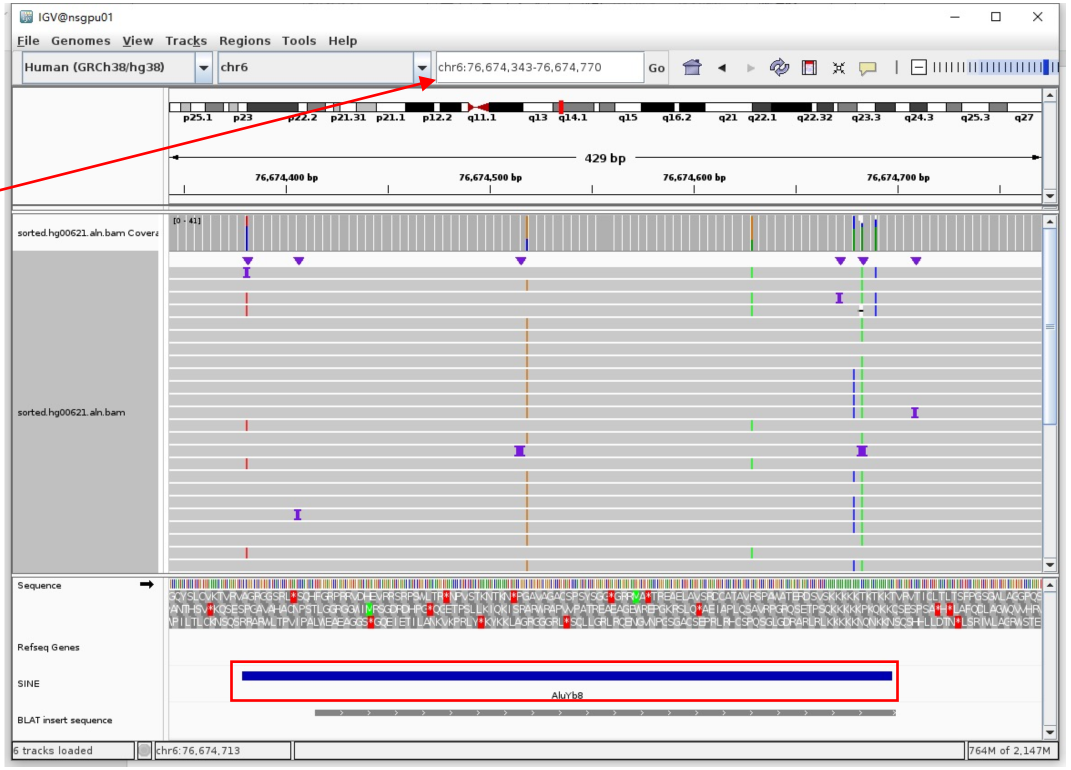


Insertion sequence (466bp)

- ➢ 19bp TSD

ACCTTACAAATGTAATGAAT

- ➢ Insertion:2-271 aligns to *Alu*Yb8:15-289, 97.77% identity

CTTACAAATGTAATGAATTTGGGAGGCCGAGGCGGGTGGATCATGAGGTCAGGAGATCGAGACCATCCTGGCTAACAAGGTGAAACCCCATCTCTACTAAAAATACAAAAAATTAGCCGGGCG
CGGTGGCGGGCGCCTGTAGTCCCAGCTACTCGGGAGGCTGAGGCAGGAGAATGGCGTGAACCCGGGAAGCGGAGCTTGCAGTGAGCCGAGATTGCGCCACTGCAGTCCGCAGTCCGGCCTGGG
CGACAGAGCGAGACTCCGTCTCA

### HG00621 chr21:22207227

➢ Mapping quality is 0, indicating that this region may have alignment issues.

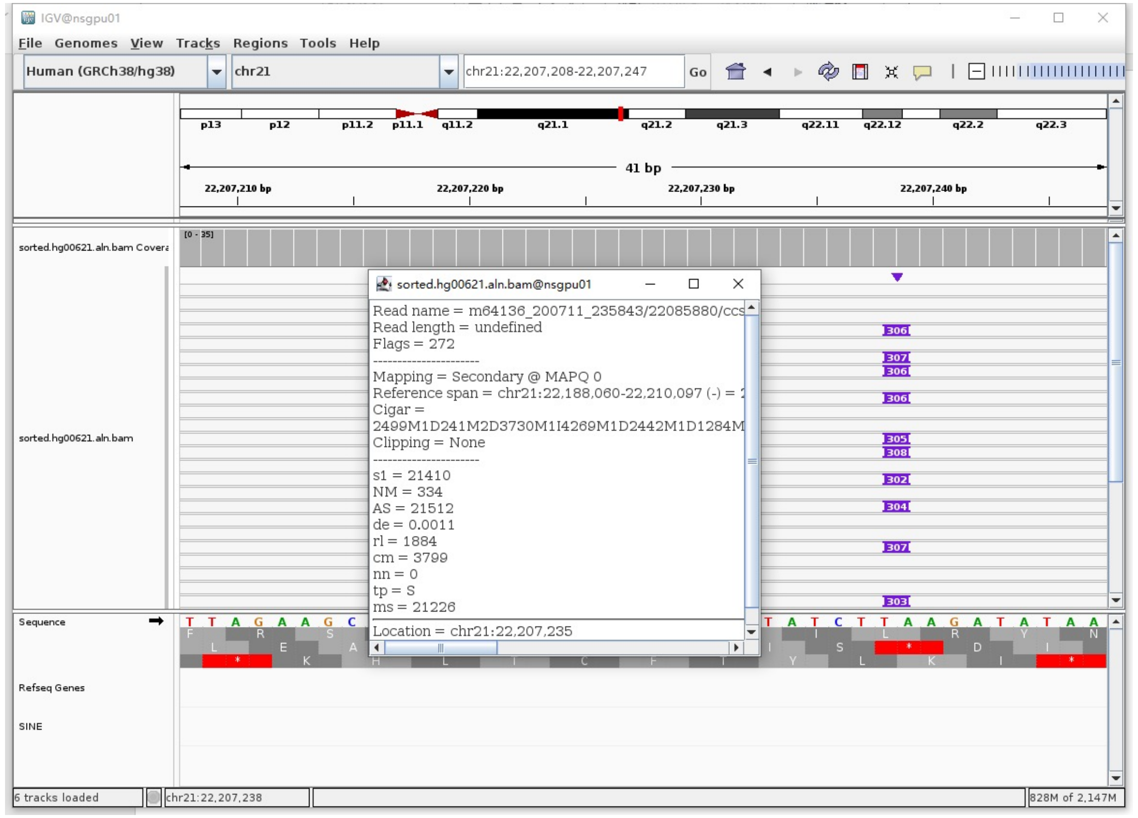


➢ Fetch one read (m64136_200714_125149/139985255/ccs, 18193 bp) and BLAT it to the hg38 genome.

The best alignment is to chr21_GL383579v2_alt (an alternative sequence of chr21) and chr21.

| ACTIONS | QUERY | SCORE | START | END | QSIZE | IDENTITY | CHROM | STRAND | START | END | SPAN |
|---|---|---|---|---|---|---|---|---|---|---|---|
| browser new tab details | YourSeq | 17874 | 1 | 18193 | 18193 | 99.9% | chr21_GL383579v2_alt | - | 100443 | 118331 | 17889 |
| browser new tab details | YourSeq | 17874 | 1 | 18193 | 18193 | 99.9% | chr21 | - | 22202916 | 22220804 | 17889 |

- Read:1-13607 aligns to chr21_GL383579v2_alt

| ACTIONS | QUERY | SCORE | START | END | QSIZE | IDENTITY | CHROM | STRAND | START | END | SPAN |
|---|---|---|---|---|---|---|---|---|---|---|---|
| browser new tab details | YourSeq | 13553 | 1 | 13565 | 13607 | 100.0% | chr21_GL383579v2_alt | - | 104766 | 118331 | 13566 |
| browser new tab details | YourSeq | 13553 | 1 | 13565 | 13607 | 100.0% | chr21 | - | 22207239 | 22220804 | 13566 |

- Read:13608-13856 aligns to *Alu*Ya5:282-34, 100% identity

TGAGACGGAGTCTCGCTCTGTCGCCCAGGCTGGAGTGCAGTGGCGGGATCTCGGCTCACTGCAAGCTCCGCCTCC
CGGGTTCACGCCATTCTCCTGCCTCAGCCTCCCAAGTAGCTGGGACTACAGGCGCCCGCCACTACGCCCGGCTAA
TTTTTTGTATTTTTAGTAGAGACGGGGTTTCACCGTTTTAGCCGGGATGGTCTCGATCTCCTGACCTCGTGATCC
GCCCGCCTCGGCCTCCCAAAGTG

12bp TSD

AGATATAATTTT

- Read:13857-18193 aligns to chr21_GL383579v2_alt

| ACTIONS | QUERY | SCORE | START | END | QSIZE | IDENTITY | CHROM | STRAND | START | END | SPAN |
|---|---|---|---|---|---|---|---|---|---|---|---|
| browser new tab details | YourSeq | 4334 | 1 | 4336 | 4336 | 100.0% | chr21_GL383579v2_alt | - | 100443 | 104777 | 4335 |
| browser new tab details | YourSeq | 4334 | 1 | 4336 | 4336 | 100.0% | chr21 | - | 22202916 | 22207250 | 4335 |

➢ It's an *Alu*Ya5 insertion.

### HG01106 chr1:82577503

- Using the built-in BLAT tool in IGV to check where the inserted sequence aligned.
- The annotation at the aligned location indicated it as *Alu*Y.

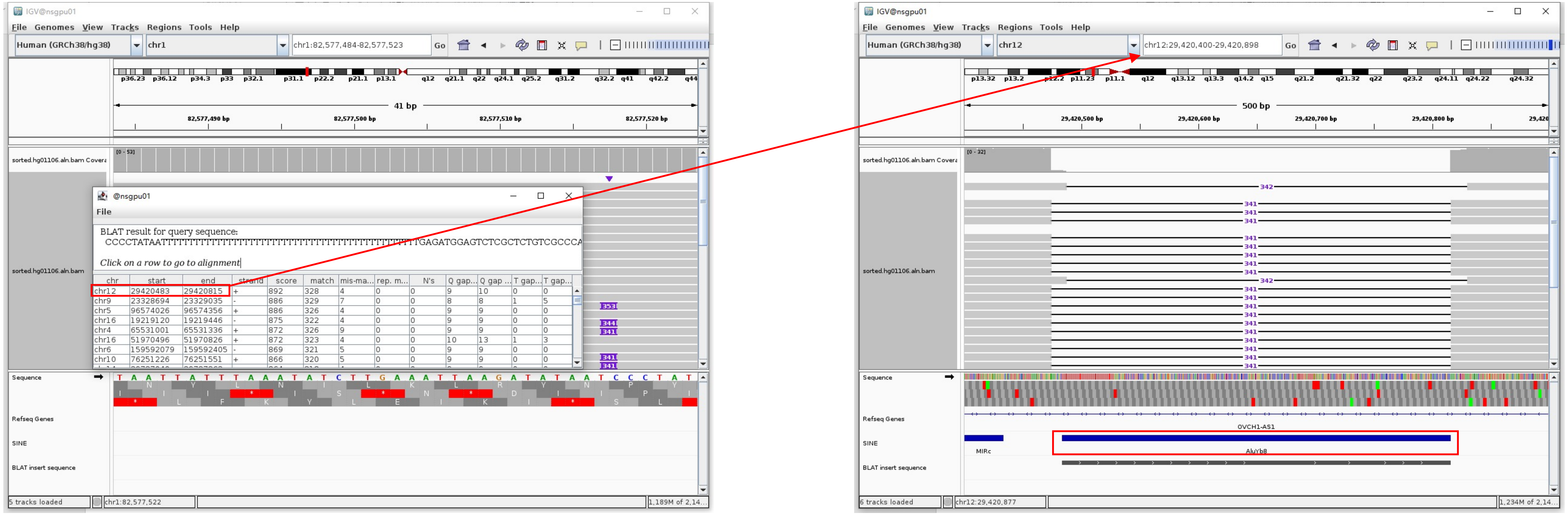


Insertion sequence (355bp)

- 10bp TSD

CCCTATAATT

- Insertion:55-355 aligns to *Alu*Yb8:289-1, 98.62% identity

TGAGATGGAGTCTCGCTCTGTCGCCCAGGCCGGACTGCGGACTGCAGTGGAGCAATCTCGGCTCACTGCAAGCTCCGCTTCCCGGGTTCACGCCATTCTCCTGCCTCAGCCTCCCCGAGTAGC
TGGGACTAACAGGCGCCTGCGCACCGGCGCCCGGCTAATTTTTTTGTATTTTTAGTAGAGACGTGGGTTTCCACCTTGTTAGCCAGGATGGTCTCGATCTCCCTGACCTCGTGATCCACCCGC
CTCGGCCTCCCAAAGTGCTGGGATTACAGGCGTGAGCCCACCGCGCCCGGCC

### HG01106 chr6:31282439

➢ Mapping quality is 0, indicating that this region may have alignment issues.

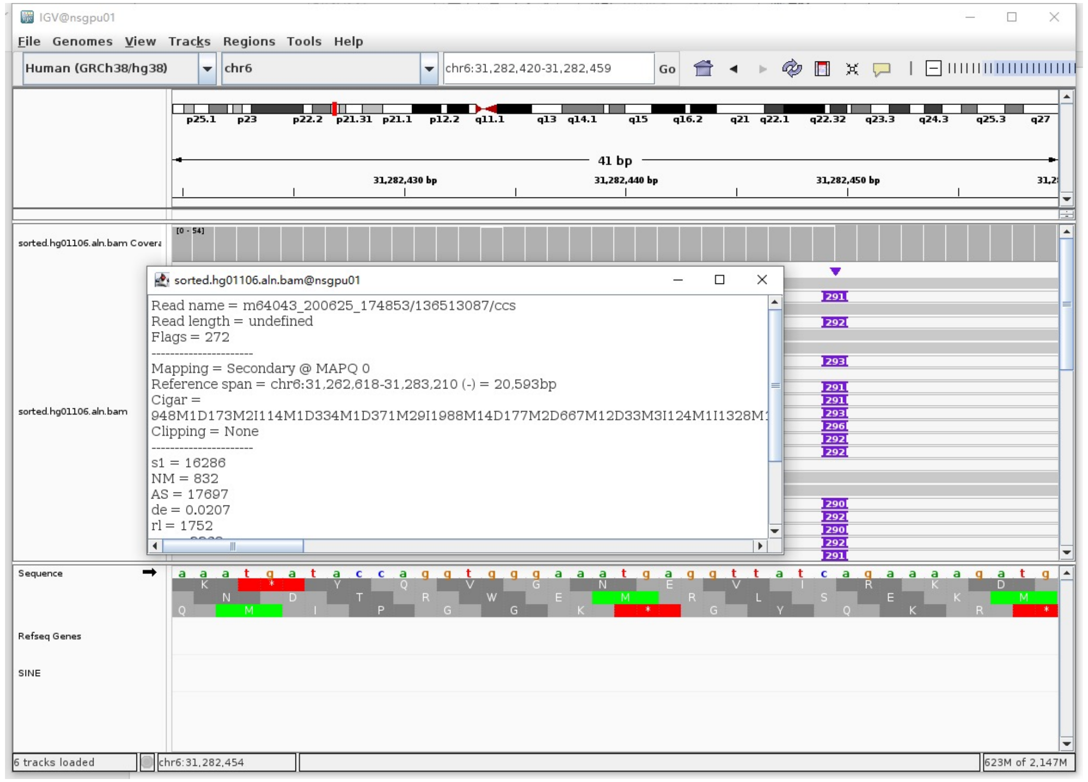


➢ Fetch one read (m64043_200625_174853/136513087/ccs, 20838 bp) and BLAT it to the hg38 genome.

The best alignment is to chr6_GL000256v2_alt (an alternative sequence of chr6), which contains an *Alu*Y element.

| ACTIONS | | | QUERY | SCORE | START | END | QSIZE | IDENTITY | CHROM | STRAND | START | END | SPAN |
|---|---|---|---|---|---|---|---|---|---|---|---|---|---|
| browser | new tab | details | YourSeq | 20813 | 1 | 20838 | 20838 | 100.0% | chr6_GL000256v2_alt | - | 2564572 | 2585407 | 20836 |
| browser | new tab | details | YourSeq | 20807 | 1 | 20838 | 20838 | 100.0% | chr6_GL000255v2_alt | - | 2518046 | 2538881 | 20836 |
| browser | new tab | details | YourSeq | 20228 | 1 | 20838 | 20838 | 99.2% | chr6_GL000253v2_alt | - | 2571656 | 2592210 | 20555 |
| browser | new tab | details | YourSeq | 20213 | 1 | 20838 | 20838 | 99.2% | chr6_GL000252v2_alt | - | 2520403 | 2540950 | 20548 |
| browser | new tab | details | YourSeq | 20181 | 1 | 20838 | 20838 | 99.1% | chr6_GL000254v2_alt | - | 2605337 | 2625896 | 20560 |
| browser | new tab | details | YourSeq | 19999 | 1 | 20838 | 20838 | 98.7% | chr6_GL000251v2_alt | - | 2743529 | 2764121 | 20593 |
| browser | new tab | details | YourSeq | 19691 | 1 | 20838 | 20838 | 98.1% | chr6 | - | 31262618 | 31283210 | 20593 |

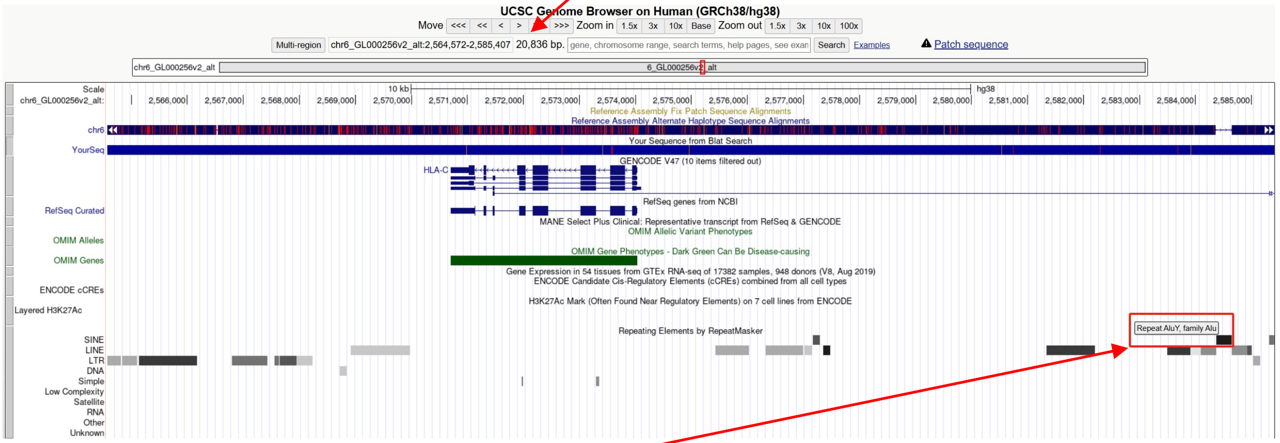


chr6_GL000256v2_alt:2584370-2584651 (*Alu*Y marked by RepeatMasker)

➢ Read:783-1038 aligns to *Alu*Yc1:282-32, 97.60% identity

TGAGACGGAGTCTCGCTCTGTCGCCCAGGCTGGAGTGCAGTGGCGCGATCTCGGCTCACTGCAAGCTCCGCCTCC
CGGGTTCACGCCATTCTCCTGCCTCAGCCTCCCGCGCAGCTGGGACTACAGGCGCCCGCCACCACGCCCGGCTAA
TTTTTTTTGTGTGTTTTTTAGTAGAGACGGGGTTTCACTGTGTTAGCCAGGATGGTCTCGATCTCCTGACCTCGT
GATCCGCCCGCCTCGGCCTCCCAAAGTGAT

➢ It's an *Alu*Y in the chromosome chr6_GL000256v2_alt.

### HG01106 chr10:52920858

- Using the built-in BLAT tool in IGV to check where the inserted sequence aligned.
- The annotation at the aligned location indicated it as *AluY*.

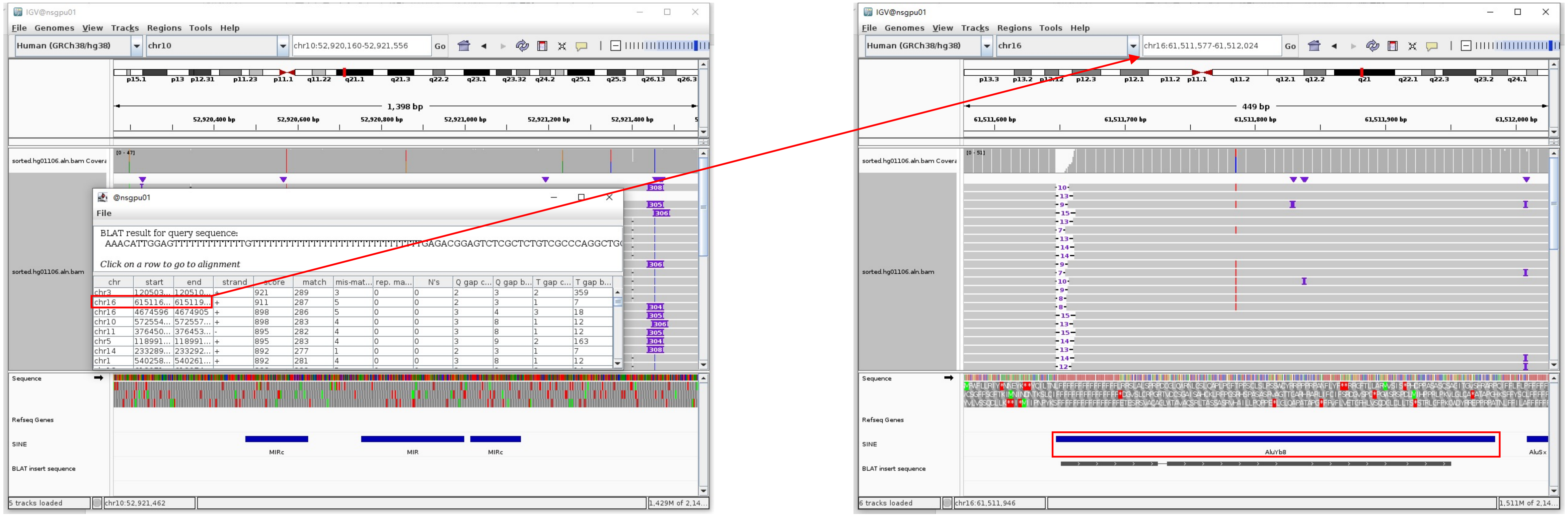


Insertion sequence (305bp)

- 15bp TSD

AAACATTGGAGTTTT

- Insertion:57-305 aligns to *AluY*c1:282-34, 97.18% identity

TGAGACGGAGTCTCGCTCTGTCGCCCAGGCTGGAGTGCAGTGGCGGGATCTCGGCTCACTGCAAGCTCCGCTTCCCGGGTTCACGCCATTCTCCTGCCTCAGCCTCCCGAGTAGCTGGGACTA
CAGGCGCCCGCCACCGCGCCCGGCTAATTTTTTGTATTTTTAGTAGAGACGGGGTTTCACCTTGTTAGCCAGGATGGTCTCGATCTCCTGACCTCATGATCCACCCGCCTCGGCCTCCCAAAG
TG

### HG01106 chr17:227341

- Using the built-in BLAT tool in IGV to check where the inserted sequence aligned.
- The annotation at the aligned location indicated it as *AluY*.

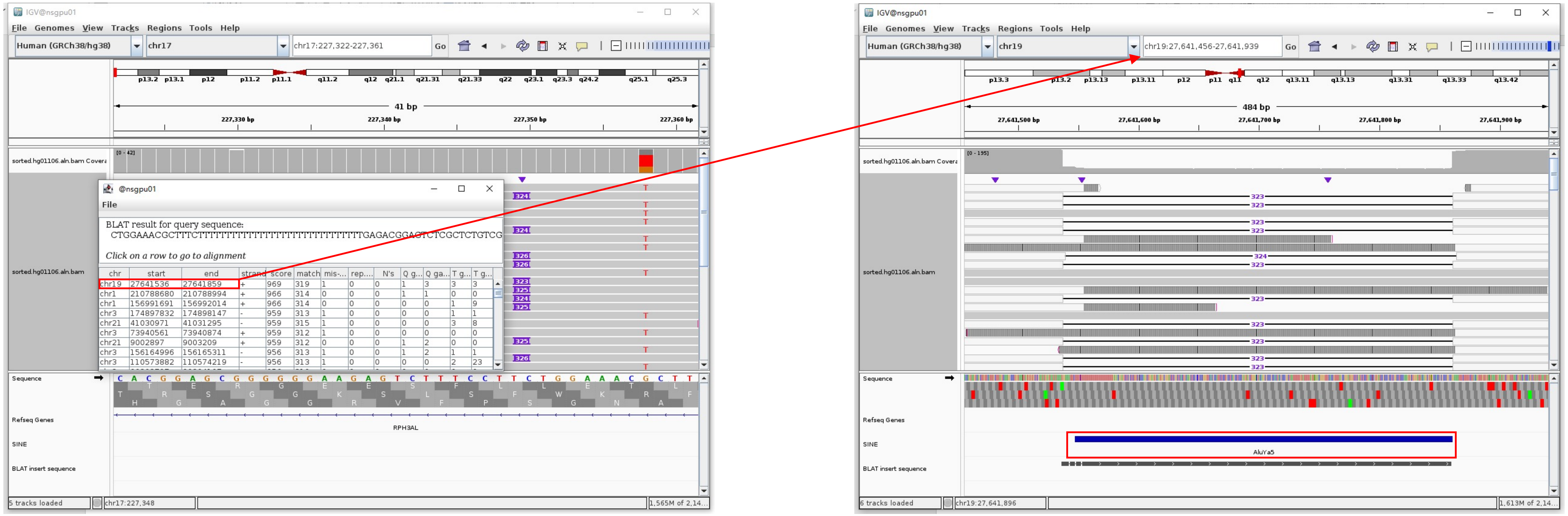


Insertion sequence (324bp)

- 16bp TSD

CTGGAAACGCTTTCTT

- Insertion:43-324 aligns to *Alu*Ya5:282-1, 100% identity

TGAGACGGAGTCTCGCTCTGTCGCCCAGGCTGGAGTGCAGTGGCGGGATCTCGGCTCACTGCAAGCTCCGCCTCCCGGGTTCACGCCATTCTCCTGCCTCAGCCTCCCAAGTAGCTGGGACTA
CAGGCGCCCGCCACTACGCCCGGCTAATTTTTTGTATTTTTAGTAGAGACGGGGTTTCACCGTTTTAGCCGGGATGGTCTCGATCTCCTGACCTCGTGATCCGCCCGCCTCGGCCTCCCAAAG
TGCTGGGATTACAGGCGTGAGCCACCGCGCCCGGC

### HG01106 chr19:52384793

- Using the built-in BLAT tool in IGV to check where the inserted sequence aligned.

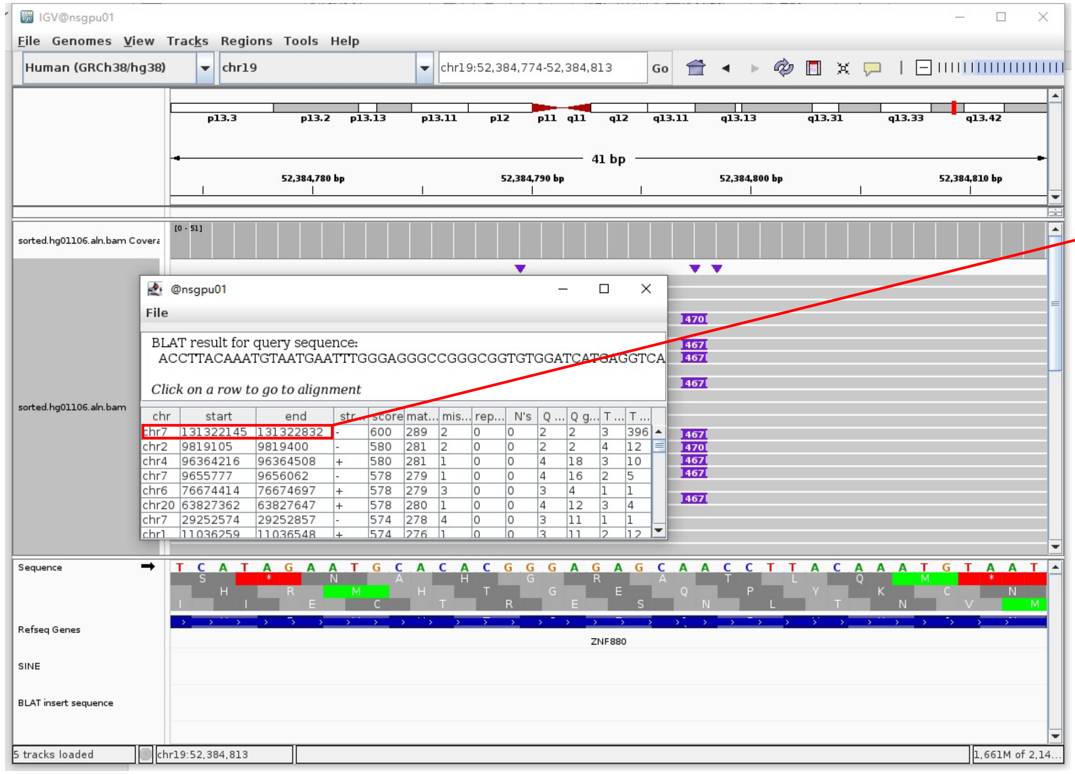


- The annotation at the aligned location indicated it as *Alu*Y.

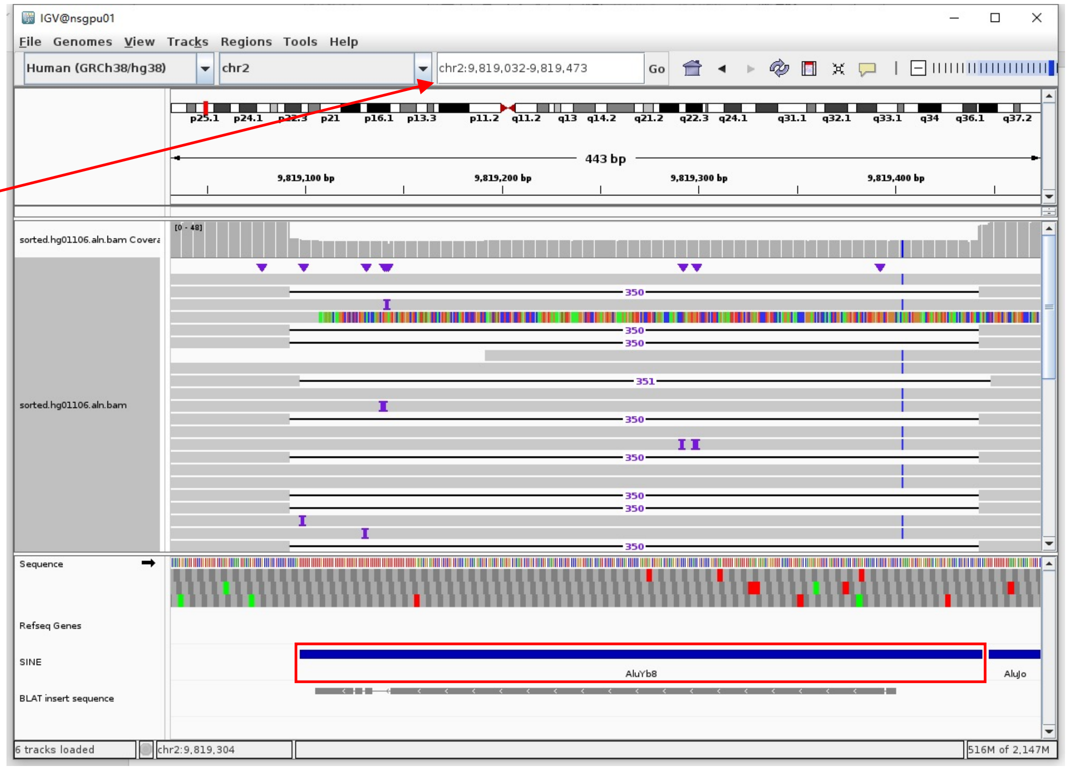


Insertion sequence (467bp)

- 20bp TSD

ACCTTACAAATGTAATGAAT

- Insertion:2-272 aligns to *Alu*Yb8:15-289, 96.65% identity

CTTACAAATGTAATGAATTTGGAGGGCCGGGCGGTGTGGATCATGAGGTCAGGAGATCGAGACCATCCTGGCTAACAAGGTGAAACCCCATCTCTACTAAAAATACAAAAAATTAGCCGGGC
GCGGTGGCGGGCGCCTGTAGTCCCAGCTACTCGGGAGGCTGAGGCAGGAGAATGGCGTGAACCCGGGAAGCGGAGCTTGCAGTGAGCCGAGATTGCGCCACTGCAGTCCGCAGTCCGGCCTGG
GCGACAGAGCGAGACTCCGTCTCA

### HG02630 chr1:82577503

- The same genomic coordinate as in HG01106 (page 19)

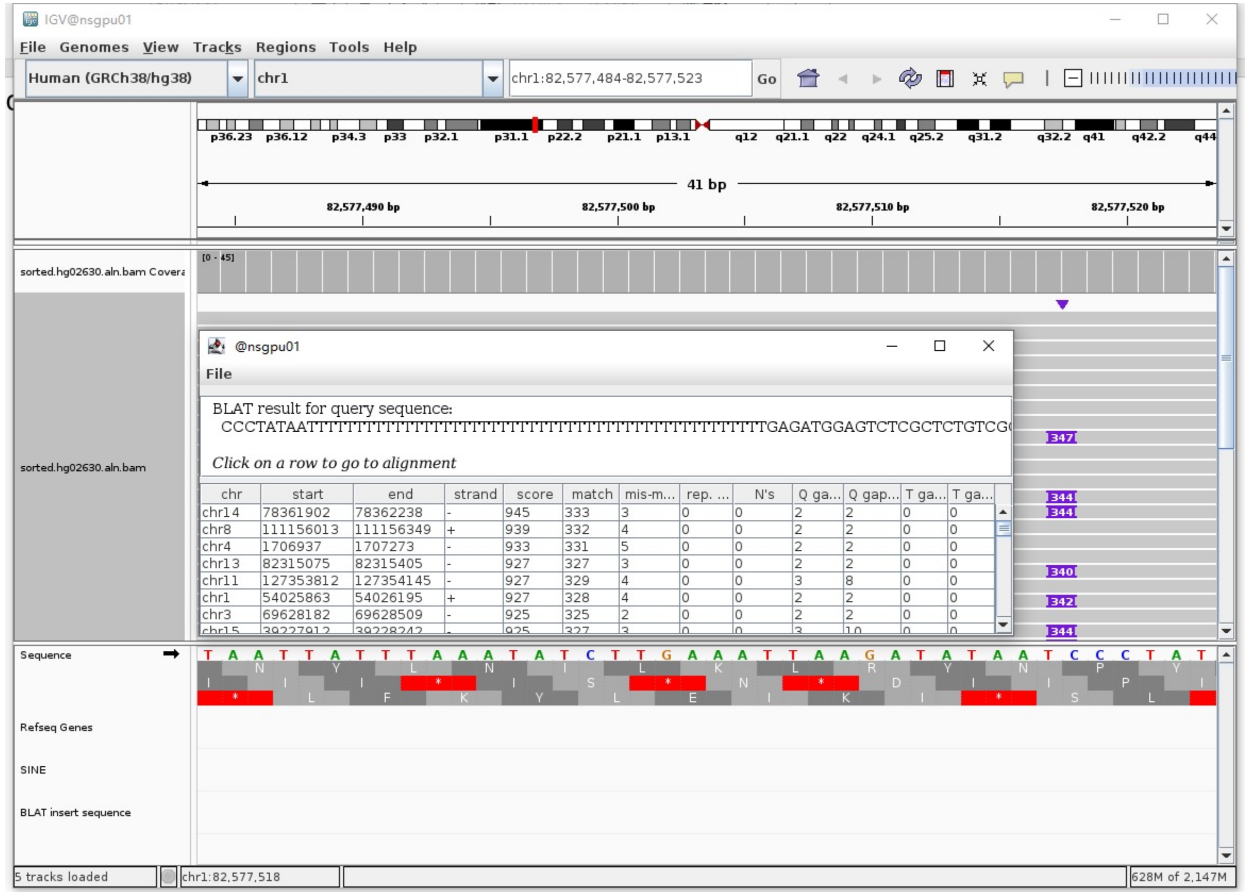

### HG02630 chr2:1467932

- ➢ Using the built-in BLAT tool in IGV to check where the inserted sequence aligned.
- ➢ The annotation at the aligned location indicated it as *Alu*Y.

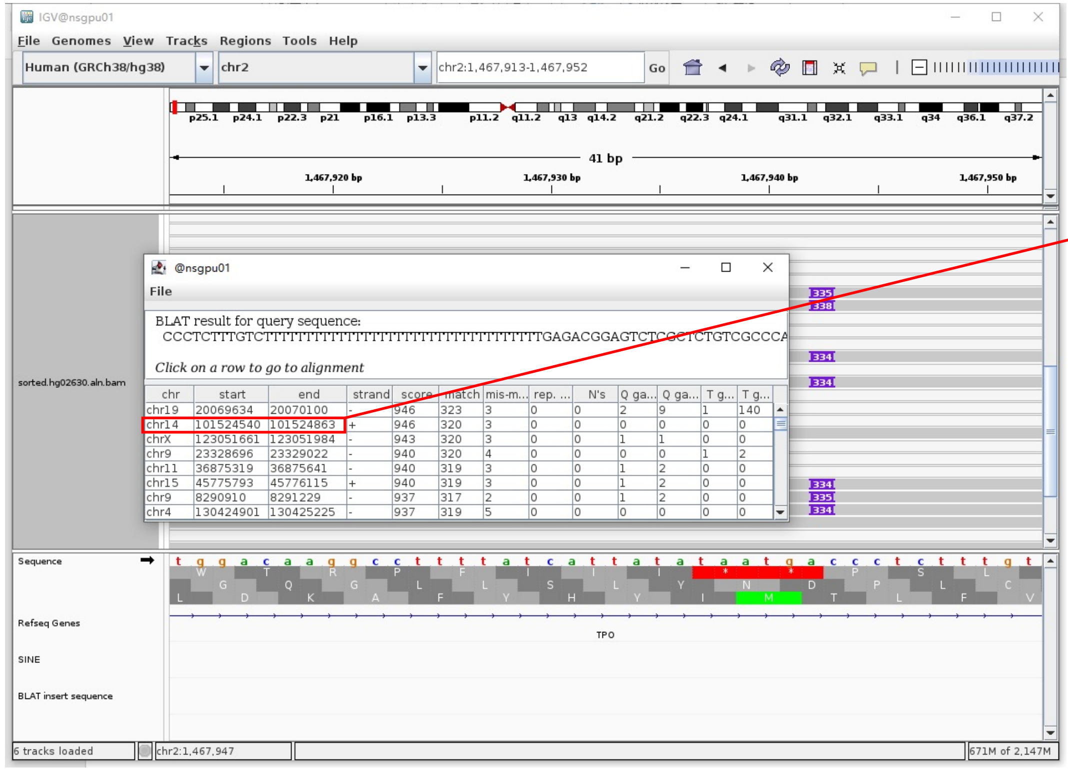


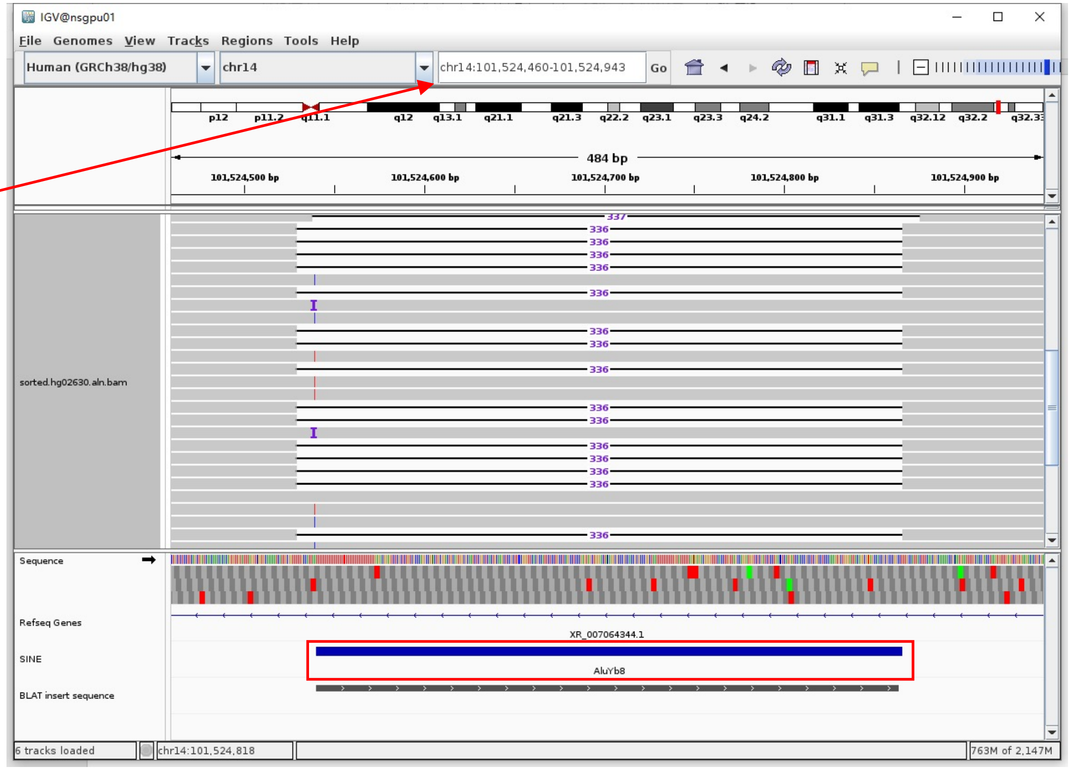


Insertion sequence (338bp)

- ➢ 18bp TSD

CCCTCTTTGTCTTTTTTT

- ➢ Insertion:45-338 aligns to *Alu*Yb8:289-1, 98.96% identity

TGAGACGGAGTCTCGCTCTGTCGCCCAGGCGGACTGCGGACTGCAGTGGCGCAATCTCGGCTCACTGCAAGCTCCGCCTCCCGGGTTCACGCCATTCTCCTGCCTCAGCCTCCCGAGTAGCT
GGGACTACAGGCGCCCGCCACCGCGCCCGGCTAATTTTTTTTTTTGTATTTTTAGTAGAGACGGGGTTTCACCTTGTTAGCCAGGATGGTCTCGATCTCCTGACCTCATGATCCACCTGCCTC
GGCCTCCCAAAGTGCTGGGATTACAGGCGTGAGCCACCGCGCCCGGC

### HG02630 chr2:36290645

- ➢ Mapping quality is 0, indicating that this region may have alignment issues.

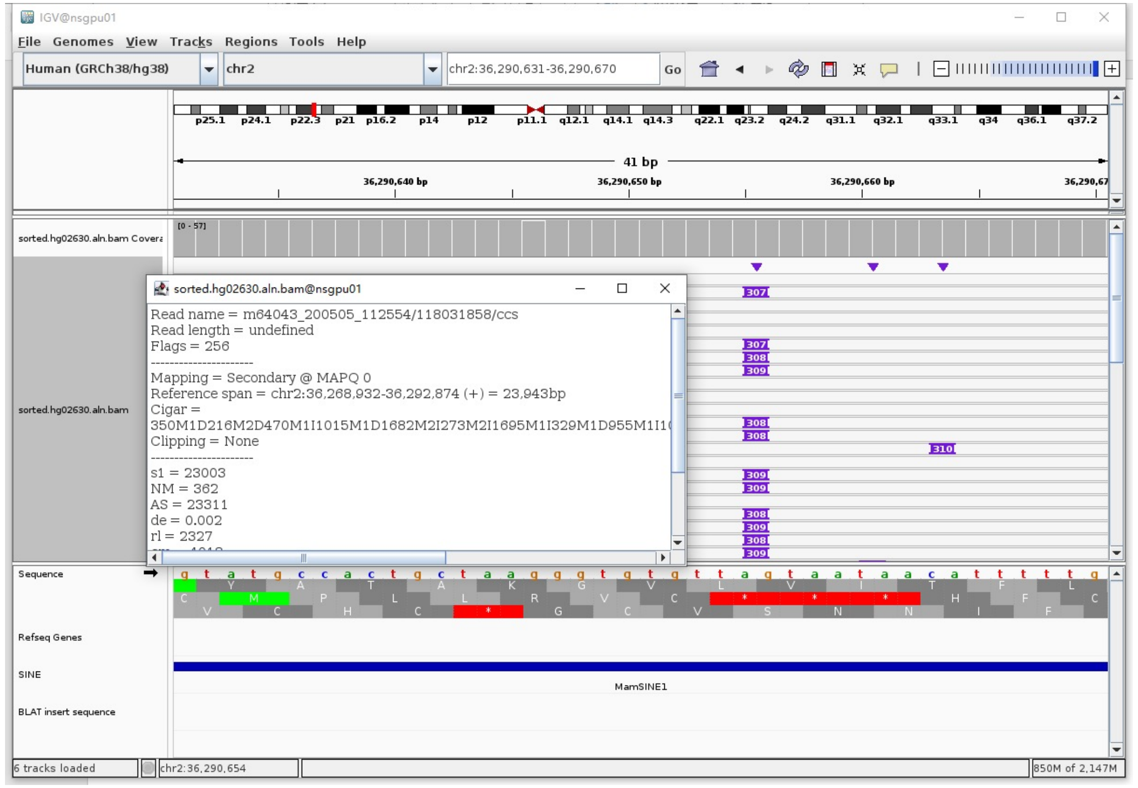


- ➢ Fetch one read (m64043_200502_223511/121243910/ccs, 23663 bp) and BLAT it to the hg38 genome.

  The best alignment is to chr2_GL383521v1_alt (an alternative sequence of chr2) and chr2.

| ACTIONS | QUERY | SCORE | START | END | QSIZE | IDENTITY | CHROM | STRAND | START | END | SPAN |
|---|---|---|---|---|---|---|---|---|---|---|---|
| browser new tab details | YourSeq | 23278 | 1 | 23663 | 23663 | 99.8% | chr2 | + | 36277037 | 36300385 | 23349 |
| browser new tab details | YourSeq | 23275 | 1 | 23663 | 23663 | 99.8% | chr2_GL383521v1_alt | - | 68964 | 92312 | 23349 |

- Read:1-13640 aligns to chr2_GL383521v1_alt

| ACTIONS | QUERY | SCORE | START | END | QSIZE | IDENTITY | CHROM | STRAND | START | END | SPAN |
|---|---|---|---|---|---|---|---|---|---|---|---|
| browser new tab details | YourSeq | 13586 | 1 | 13630 | 13640 | 99.9% | chr2_GL383521v1_alt | - | 78680 | 92312 | 13633 |
| browser new tab details | YourSeq | 13586 | 1 | 13630 | 13640 | 99.9% | chr2 | + | 36277037 | 36290669 | 13633 |

- Read:13641-13923 aligns to *Alu*Ya5:282-1, 99.29% identity

  TGAGACGGAGTCTCGCTCTGTCGCCCAGGCTGGAGTGCAGTGGCGCAATCTCGGCTCACTGCAAGCTCCGCCT
  CCCGGGTTCACGCCATTCTCCTGCCTCAGCCTCCCAAGTAGCTGGGACTACAGGCGCCCGCCACTACGCCCGG
  CTAATTTTTTGTATTTTTAGTAGAGACGGGGTTTCACCGTTTTAGCCGGGATGGTCTCGATCTCCTGACCTCG
  TGATCCGCCCGCCTCGGCCTCCCAAAGTGCTGGGATTACAGGCGTGAGCCACCGCGCCCGGCC

  8bp TSD

  GTAATAAC

- Read:13924-23663 also aligns to chr2_GL383521v1_alt

| ACTIONS | QUERY | SCORE | START | END | QSIZE | IDENTITY | CHROM | STRAND | START | END | SPAN |
|---|---|---|---|---|---|---|---|---|---|---|---|
| browser new tab details | YourSeq | 9705 | 1 | 9739 | 9739 | 99.9% | chr2_GL383521v1_alt | - | 68964 | 78693 | 9730 |
| browser new tab details | YourSeq | 9705 | 1 | 9739 | 9739 | 99.9% | chr2 | + | 36290656 | 36300385 | 9730 |

- ➢ It's an *Alu*Ya5

### HG02630 chr6:29416032

- Using the built-in BLAT tool in IGV to check where the inserted sequence aligned.
- The annotation at the aligned location indicated it as *Alu*Y.

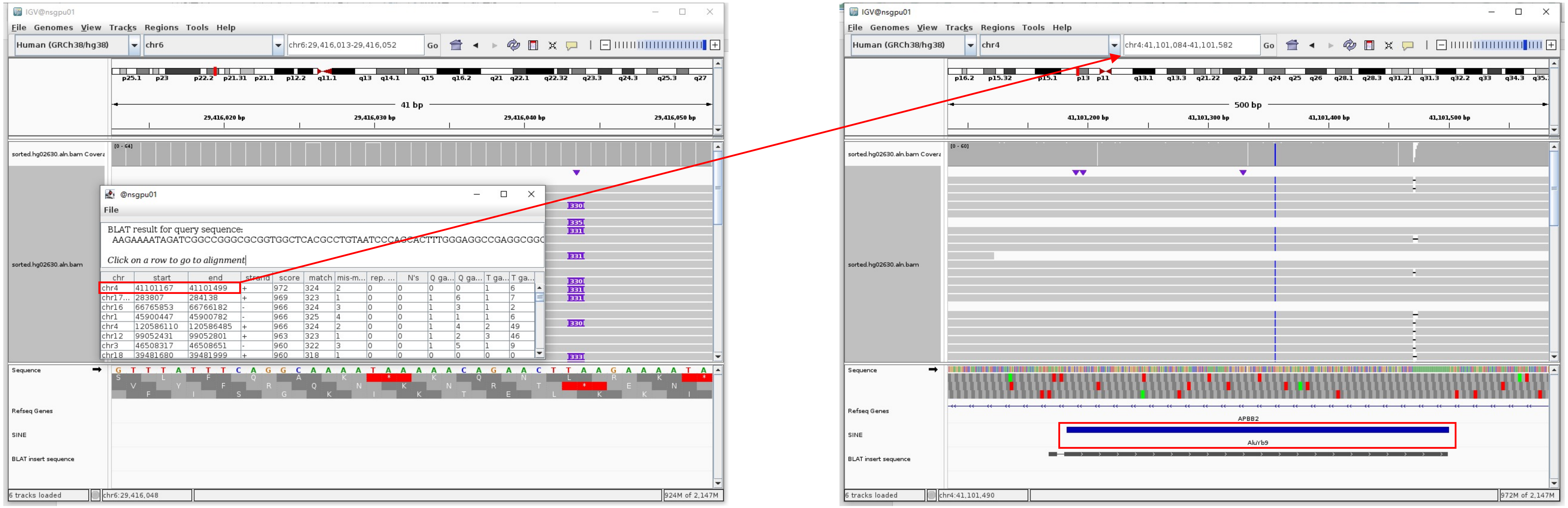


Insertion sequence (330bp)

- 13bp TSD

AAGAAAATAGATC

- Insertion:13-302 aligns to *Alu*Yb8:1-289, 99.31% identity

GGCCGGGCGCGGTGGCTCACGCCTGTAATCCCAGCACTTTGGGAGGCCGAGGCGGGTGGATCATGAGGTCAGGAGATCGAGACCATCCTGGCTAACAAGGTGAAACCCCGTCTCTACTAAAAA
TACAAAAAATTAGCCGGGCGCGGTGGCGGGCGCCTGTAGTCCCAGCTACTCGGGAGGCTGAGGCAGGAGAATGGCGTGAACCCGGGAAGCGGAGCTTGCAGTGAGCCGAGATTGCGCCACTGC
AGTCCGCAGTCCGACGTGGGCGACAGAGCGAGACTCCGTCTCA.........poly(A)

### HG02630 chr6:29851071

➢ The same genomic coordinate as in HG00438 (page 3)

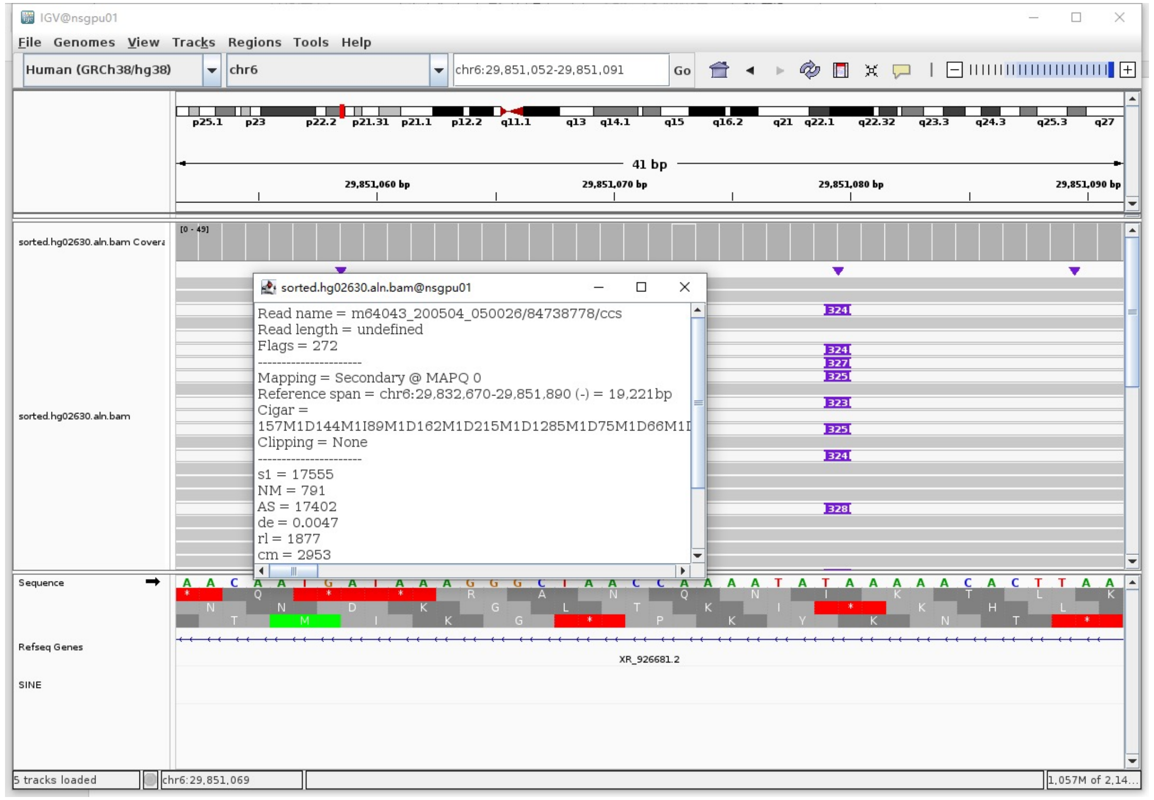

### HG02630 chr6:31471215

- The same genomic coordinate as in HG00438 (page 4)

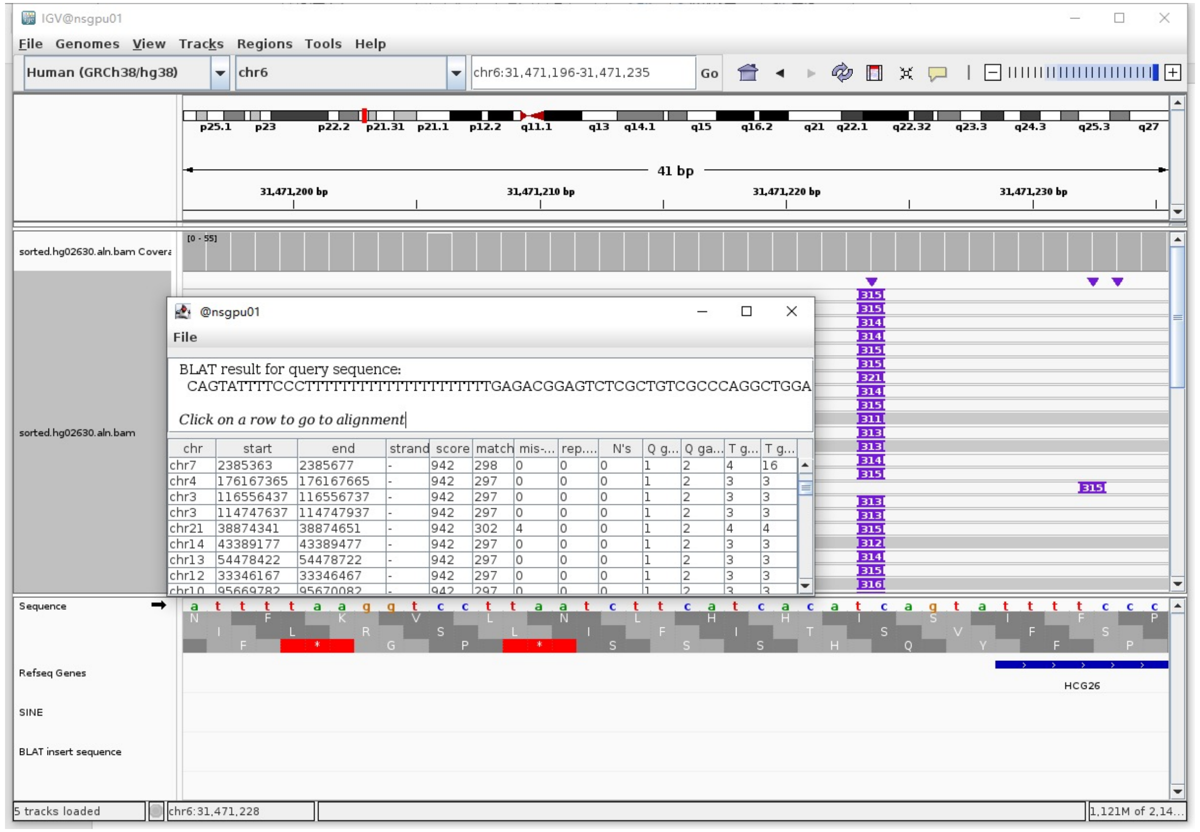

### HG02630 chr6:79370701

- ➢ Using the built-in BLAT tool in IGV to check where the inserted sequence aligned.
- ➢ The annotation at the aligned location indicated it as *Alu*Y.

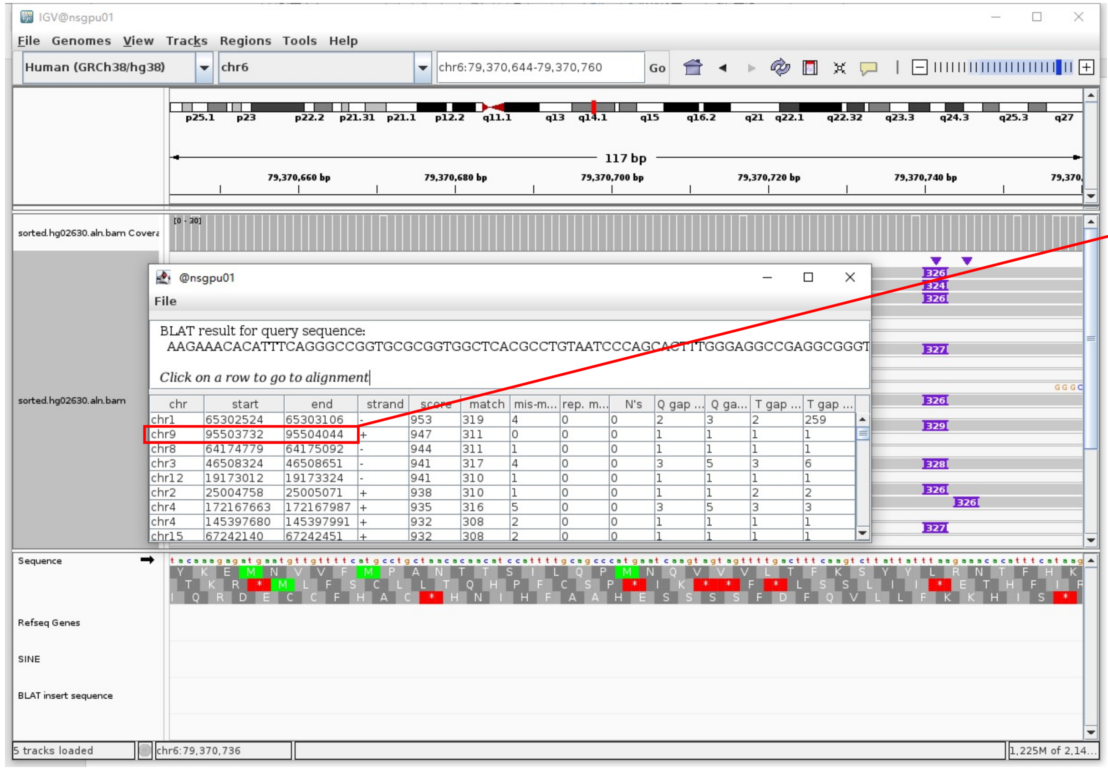


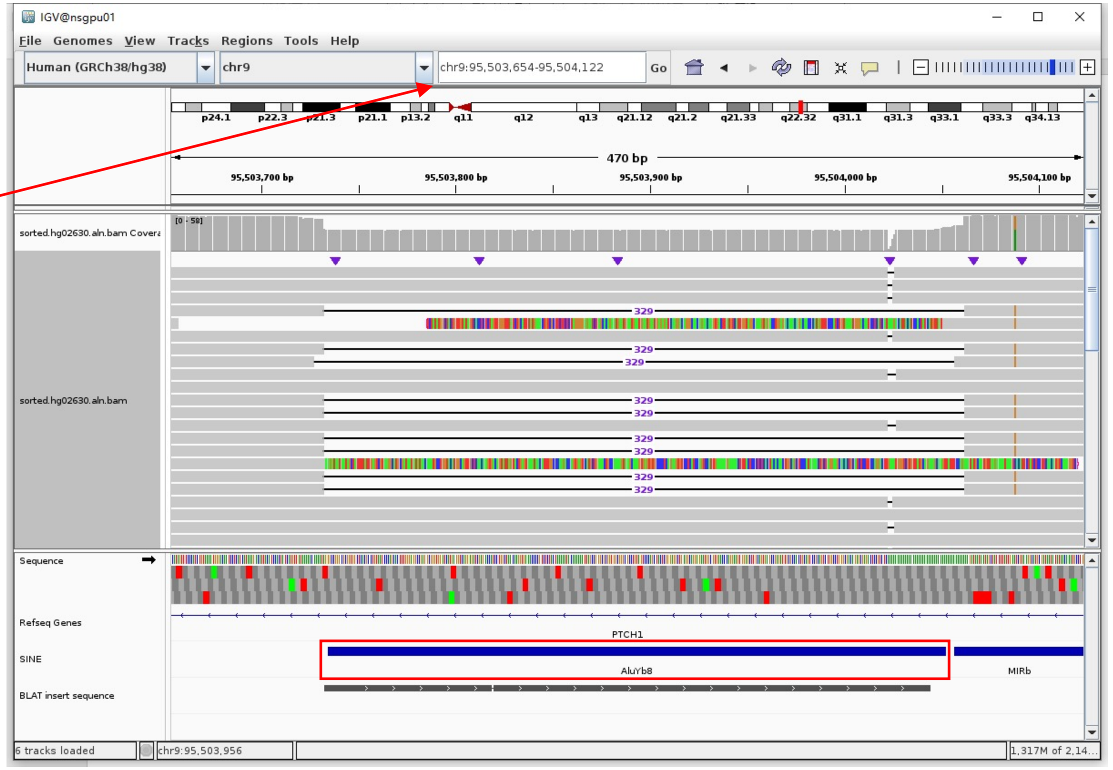


Insertion sequence (324bp)

- ➢ 15bp TSD

AAGAAACACATTTCA

- ➢ Insertion:16-305 aligns to *Alu*Yb8:1-289, 98.96% identity

GGCCGGTGCGCGGTGGCTCACGCCTGTAATCCCAGCACTTTGGGAGGCCGAGGCGGGTGGATCATGAGGTCAGGAGATCGAGACCTCCTGGCTAACAAGGTGAAACCCCGTCTCTACTAAAAA
TACAAAAAATTAGCCGGGCGCGGTGGCGGGCGCCTGTAGTCCCAGCTACTCGGGAGGCTGAGGCAGGAGAATGGCGTGAACCCAGGAAGTGGAGCTTGCAGTGAGCCGAGATTGCGCCACTGC
AGTCCGCAGTCTGGCCTGGGCGACAGAGCGAGACTCCGTCTCA………poly(A)

### HG02630 chr7:155318862

- Using the built-in BLAT tool in IGV to check where the inserted sequence aligned.
- The annotation at the aligned location indicated it as *AluY*.

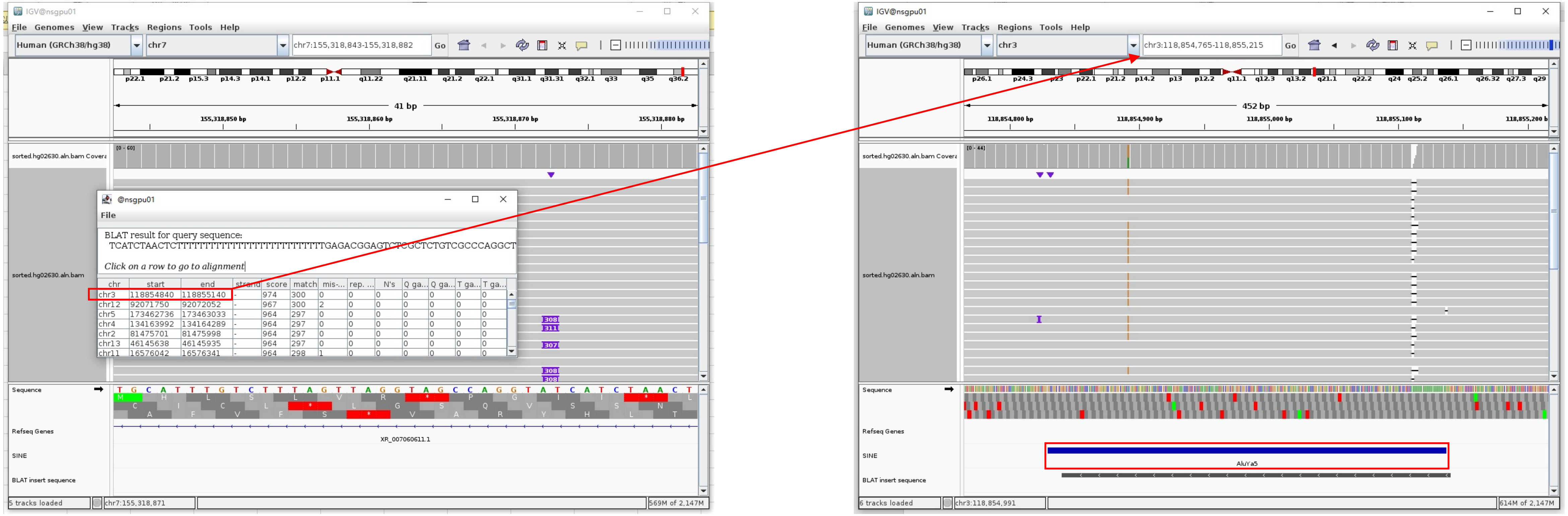


Insertion sequence (308bp)

- 14bp TSD

TCATCTAACTCTTT

- Insertion:37-308 aligns to *AluYa5*:282-11, 99.63% identity

TGAGACGGAGTCTCGCTCTGTCGCCCAGGCTGGAGTGCAGTGGCGGGATCTCGGCTCACTGCAAGCTCCGCCTCCCGGGTTCACGCCATTCTCCTGCCTCAGCCTCCCAAGTAGCTGGGACTA
CAGGCGCCCGCCACTACGCCCGGCTAATTTTTTGTATTTTTAGTAGAGACGGGGTTTCACCATTTTAGCCGGGATGGTCTCGATCTCCTGACCTCGTGATCCGCCCGCCTCGGCCTCCCAAAG
TGCTGGGATTACAGGCGTGAGCCAC

### HG02630 chr12:11026415

- The same genomic coordinate as in HG00621 (page 12)

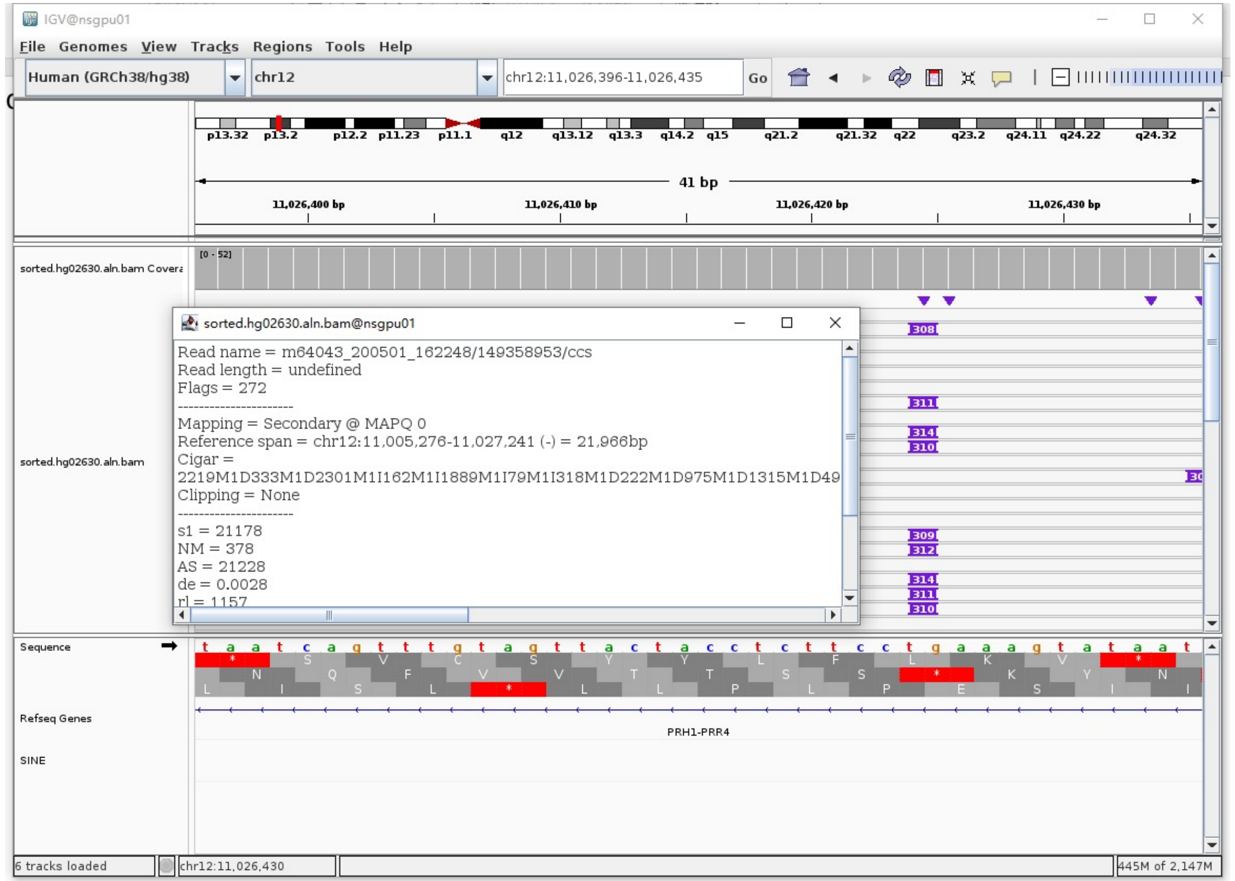

### HG02630 chr12:58064416

- Mapping quality is 0, indicating that this region may have alignment issues.

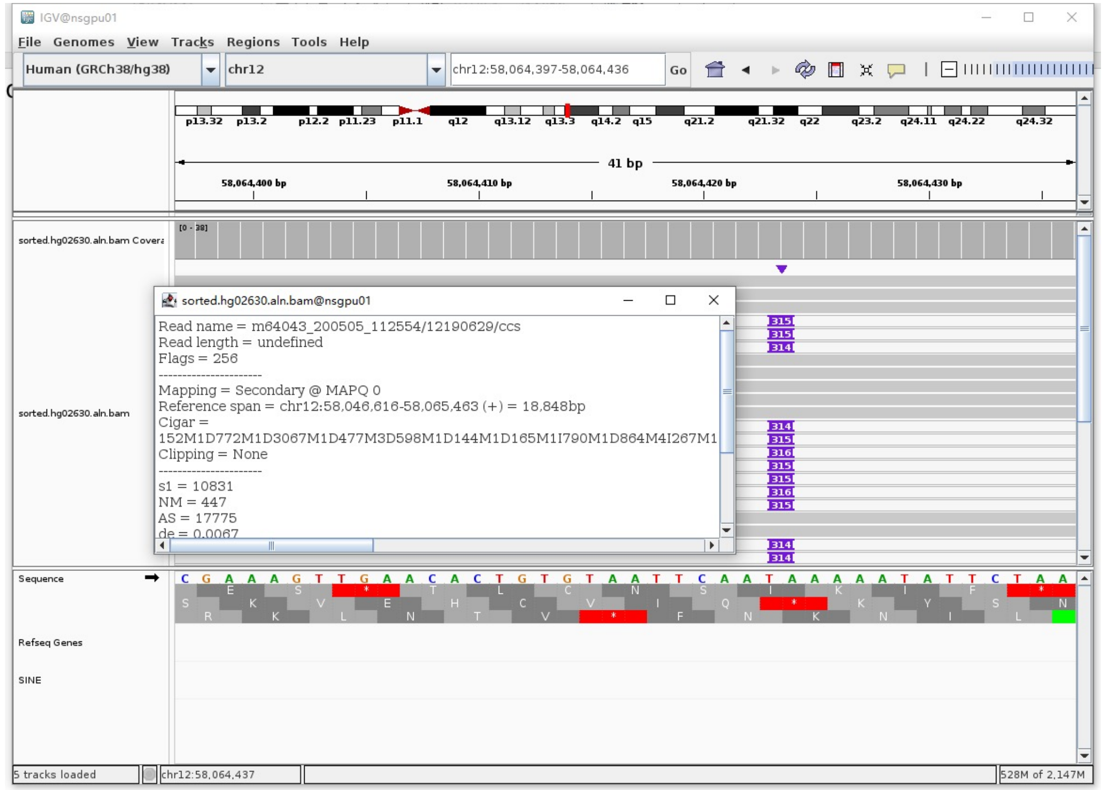


- Fetch one read (m64043_200505_112554/12190629/ccs, 19150 bp) and BLAT it to the hg38 genome.

  The best alignment is to chr12_GL383550v2_alt (an alternative sequence of chr12), which contains an *Alu*Ya5 element.

| ACTIONS | | | QUERY | SCORE | START | END | QSIZE | IDENTITY | CHROM | STRAND | START | END | SPAN |
|---|---|---|---|---|---|---|---|---|---|---|---|---|---|
| browser | new tab | details | YourSeq | 19124 | 1 | 19150 | 19150 | 100.0% | chr12_GL383550v2_alt | - | 36144 | 55299 | 19156 |
| browser | new tab | details | YourSeq | 17792 | 1 | 18321 | 19150 | 99.4% | chr12 | + | 58046616 | 58064623 | 18008 |

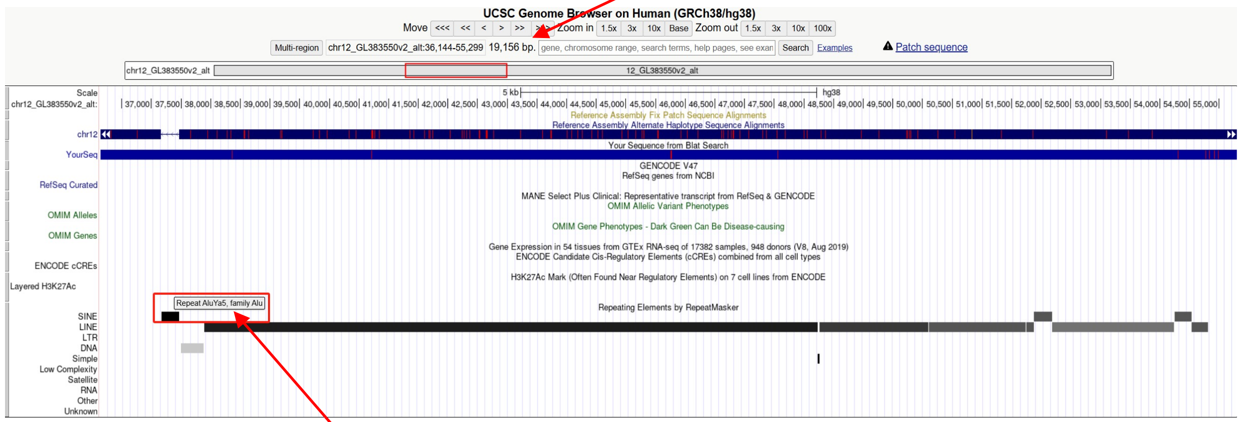


chr12_GL383550v2_alt:37169-37469 (*Alu*Ya5 marked by RepeatMasker)

- Read:17825-18107 aligns to *Alu*Ya5:1-282, 100% identity

  GGCCGGGCGCGGTGGCTCACGCCTGTAATCCCAGCACTTTGGGAGGCCGAGGCGGGCGGATCACGAGGTCAGGAG
  ATCGAGACCATCCCGGCTAAAACGGTGAAACCCCGTCTCTACTAAAAATACAAAAAATTAGCCGGGCGTAGTGGC
  GGGCGCCTGTAGTCCCAGCTACTTGGGAGGCTGAGGCAGGAGAATGGCGTGAACCCGGGAGGCGGAGCTTGCAGT
  GAGCCGAGATCCCGCCACTGCACTCCAGCCTGGGCGACAGAGCGAGACTCCGTCTCA

- It’s an *Alu*Ya5 in the chromosome chr12_GL383550v2_alt.

### HG02630 chr12:58961589

- ➢ Mapping quality is 0, indicating that this region may have alignment issues.

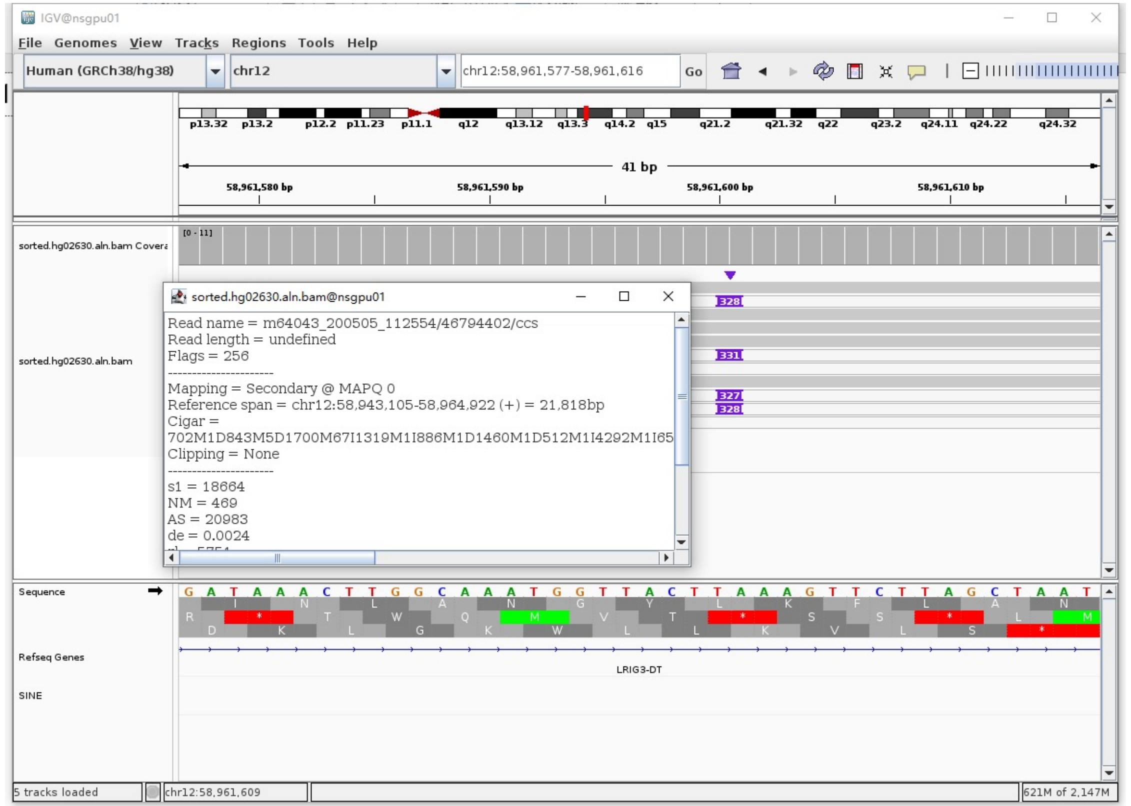


- ➢ Fetch one read (m64043_200505_112554/46794402/ccs, 22190 bp) and BLAT it to the hg38 genome.

  The best alignment is to chr12_GL383552v1_alt (an alternative sequence of chr12).

| ACTIONS | | | QUERY | SCORE | START | END | QSIZE | IDENTITY | CHROM | STRAND | START | END | SPAN |
|---|---|---|---|---|---|---|---|---|---|---|---|---|---|
| browser | new tab | details | YourSeq | 21722 | 1 | 22190 | 22190 | 99.8% | chr12_GL383552v1_alt | + | 13847 | 35657 | 21811 |
| browser | new tab | details | YourSeq | 21710 | 1 | 22190 | 22190 | 99.8% | chr12 | + | 58943105 | 58964922 | 21818 |

- Read:1-18560 aligns to chr12_GL383552v1_alt

| ACTIONS | | | QUERY | SCORE | START | END | QSIZE | IDENTITY | CHROM | STRAND | START | END | SPAN |
|---|---|---|---|---|---|---|---|---|---|---|---|---|---|
| browser | new tab | details | YourSeq | 18437 | 1 | 18559 | 18559 | 99.8% | chr12_GL383552v1_alt | + | 13848 | 32361 | 18514 |
| browser | new tab | details | YourSeq | 18426 | 1 | 18559 | 18559 | 99.8% | chr12 | + | 58943106 | 58961612 | 18507 |

- Read:18561-18843 aligns to *Alu*Ya5:1-282, 99.65% identity

GGCCGGGCGCGGTGGCTCACGCCTGTAATCCCAGCACTTTGGGAGGCCGAGGCGGGCGGATCACGAGGTCAGGAGA
TCGAGACCATCCCGGCTAAAACGGTGAAACCCCGTCTCTACTAAAAATACAAAAAATTAGCCGGGCGTAGTGGCGG
GCGCCTGTAGTCCCAGCTACTTGGGAGGCTGAGGCGGGAGAATGGCGTGAACCCGGGAGGCGGAGCTTGCAGTGAG
CCGAGATCCCGCCACTGCACTCCAGCCTGGGCGACAGAGCGAGACTCCGTCTCA

13bp TSD

AAAGTTCTTAGCT

- Read:18844-22190 also aligns to chr12_GL383552v1_alt

| ACTIONS | | | QUERY | SCORE | START | END | QSIZE | IDENTITY | CHROM | STRAND | START | END | SPAN |
|---|---|---|---|---|---|---|---|---|---|---|---|---|---|
| browser | new tab | details | YourSeq | 3297 | 33 | 3346 | 3346 | 99.7% | chr12_GL383552v1_alt | + | 32350 | 35657 | 3308 |
| browser | new tab | details | YourSeq | 3296 | 33 | 3346 | 3346 | 99.9% | chr12 | + | 58961601 | 58964922 | 3322 |

- ➢ It's an *Alu*Ya5 insertion.

### HG02630 chr14:91780893

- Mapping quality is 0, indicating that this region may have alignment issues.

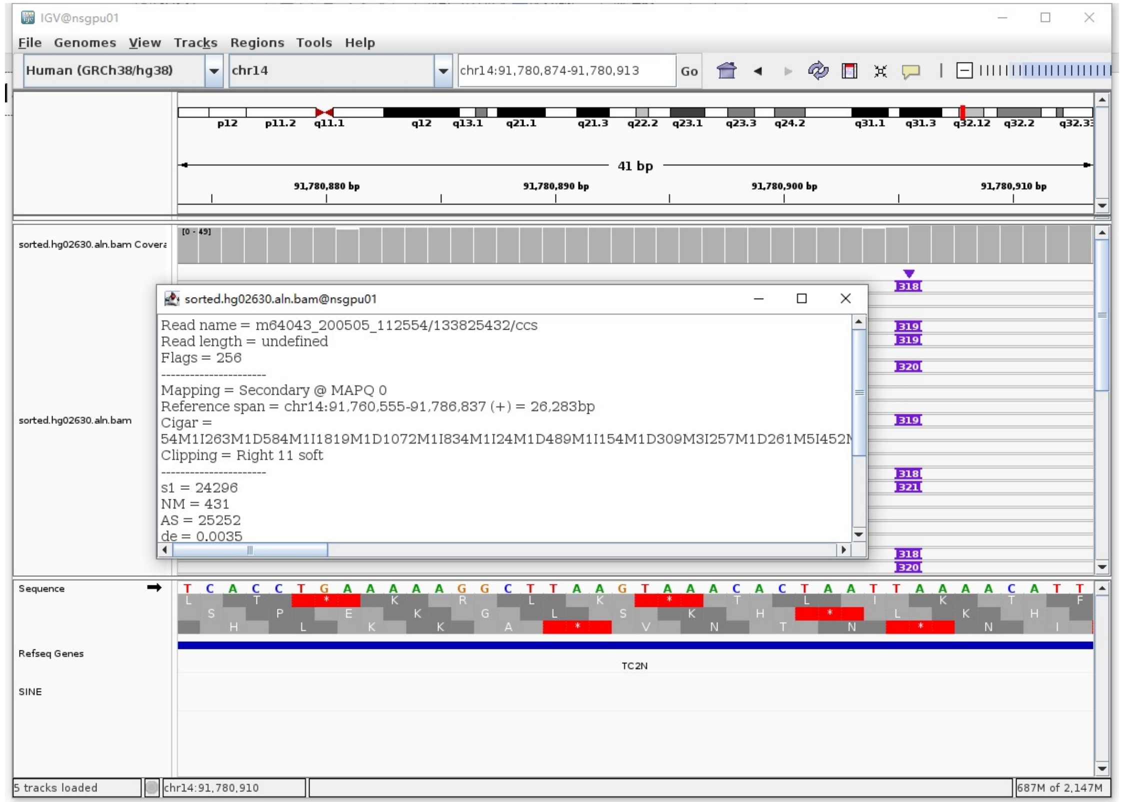


- Fetch one read (m64043_200502_223511/53084995/ccs, 19913 bp) and BLAT it to the hg38 genome.

  The best alignment is to chr14_KI270844v1_alt (an alternative sequence of chr14) and chr14.

| ACTIONS | QUERY | SCORE | START | END | QSIZE | IDENTITY | CHROM | STRAND | START | END | SPAN |
|---|---|---|---|---|---|---|---|---|---|---|---|
| browser new tab details | YourSeq | 19483 | 1 | 19913 | 19913 | 99.7% | chr14_KI270844v1_alt | - | 234122 | 253718 | 19597 |
| browser new tab details | YourSeq | 19483 | 1 | 19913 | 19913 | 99.7% | chr14 | - | 91766327 | 91785923 | 19597 |

- Read:1-5041 aligns to chr14_KI270844v1_alt

| ACTIONS | QUERY | SCORE | START | END | QSIZE | IDENTITY | CHROM | STRAND | START | END | SPAN |
|---|---|---|---|---|---|---|---|---|---|---|---|
| browser new tab details | YourSeq | 4997 | 1 | 5017 | 5040 | 99.9% | chr14_KI270844v1_alt | - | 248701 | 253717 | 5017 |
| browser new tab details | YourSeq | 4997 | 1 | 5017 | 5040 | 99.9% | chr14 | - | 91780906 | 91785922 | 5017 |

- Read:5042-5324 aligns to *Alu*Ya5:282-1, 99.65% identity

  TGAGACGGAGTCTCGCTCTGTCGCCCAGGCTGGAGTGCAGTGGCGGGATCTCGGCTCACTGCAAGCTCCGCCTC
  CCGGGTTCACGCCATTCTCCTGCCTCAGCCTCCCAAGTAGCTGGGACTACAGGCGCCTGCCACTACGCCCGGCT
  AATTTTTTGTATTTTTAGTAGAGACGGGGTTTCACCGTTTTAGCCGGGATGGTCTCGATCTCCTGACCTCGTGA
  TCCGCCCGCCTCGGCCTCCCAAAGTGCTGGGATTACAGGCGTGAGCCACCGCGCCCGGCC

  12bp TSD

  AAAACATTTAGG

- Read:5325-19913 also aligns to chr14_KI270844v1_alt

| ACTIONS | QUERY | SCORE | START | END | QSIZE | IDENTITY | CHROM | STRAND | START | END | SPAN |
|---|---|---|---|---|---|---|---|---|---|---|---|
| browser new tab details | YourSeq | 14496 | 1 | 14588 | 14588 | 99.8% | chr14_KI270844v1_alt | - | 234122 | 248710 | 14589 |
| browser new tab details | YourSeq | 14496 | 1 | 14588 | 14588 | 99.8% | chr14 | - | 91766327 | 91780915 | 14589 |

- It's an *Alu*Ya5 insertion.

### HG02630 chr15:30856126

- Using the built-in BLAT tool in IGV to check where the inserted sequence aligned.
- The annotation at the aligned location indicated it as *AluY*.

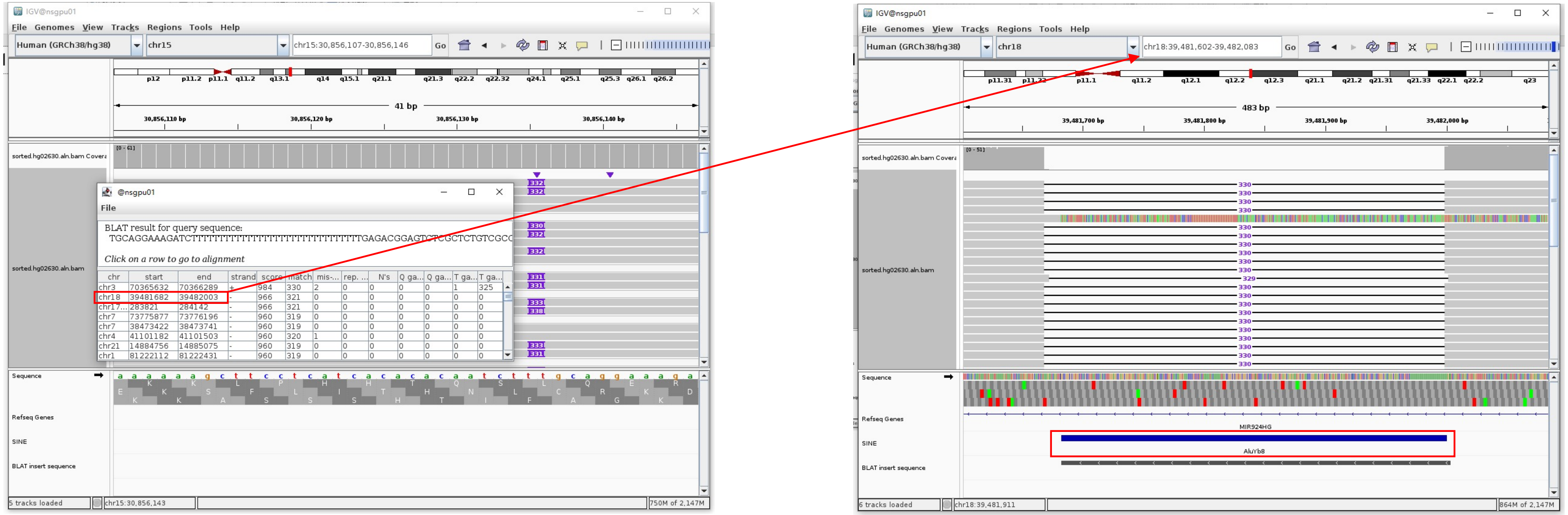


Insertion sequence (332bp)

- 15bp TSD

TGCAGGAAAGATCTT

- Insertion:43-332 aligns to *Alu*Yb8:289-1, 99.65% identity

TGAGACGGAGTCTCGCTCTGTCGCCCAGGTCGGACTGCGGACTGCAGTGGCGCAATCTCGGCTCACTGCAAGCTCCGCTTCCCGGGTTCACGCCATTCTCCTGCCTCAGCCTCCCGAGTAGCT
GGGACTACAGGCGCCCGCCACCGCGCCCGGCTAATTTTTTGTATTTTTAGTAGAGACGGGGTTTCACCTTGTTAGCCAGGATGGTCTCGATCTCCTGACCTCATGATCCACCCGCCTCGGCCT
CCCAAAGTGCTGGGATTACAGGCGTGAGCCACCGCGCCCGGCC

### HG02630 chr15:73690233

- ➢ Using the built-in BLAT tool in IGV to check where the inserted sequence aligned.

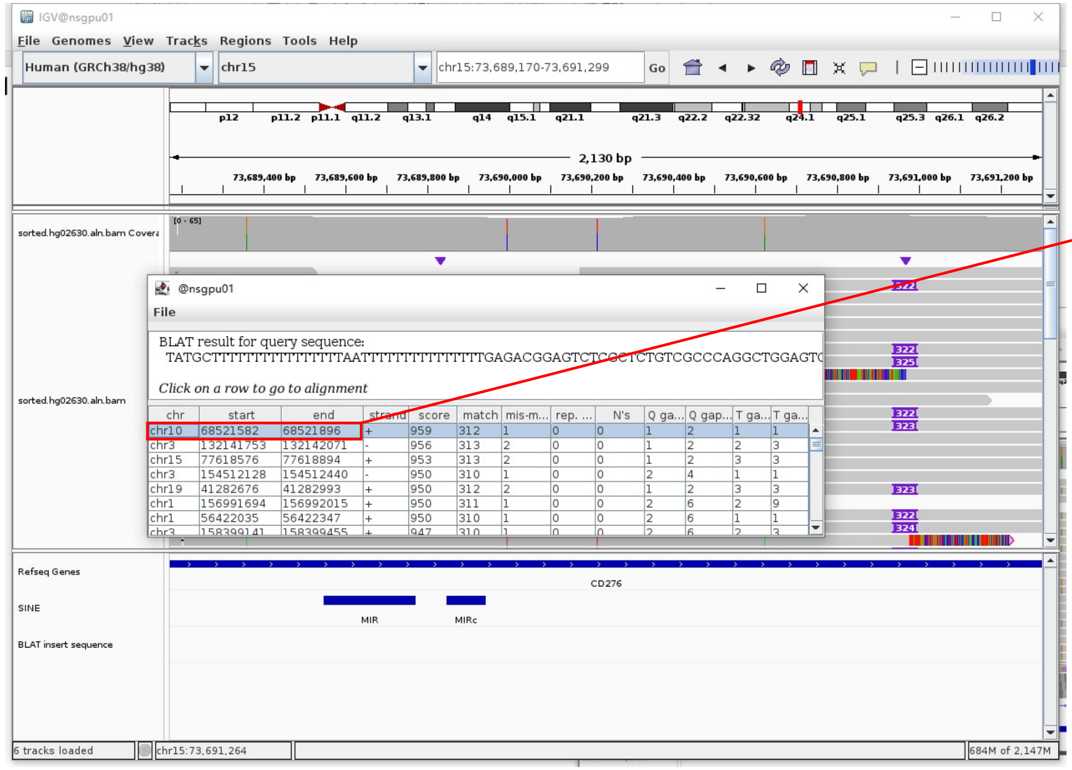

- ➢ The annotation at the aligned location indicated it as *AluY*.

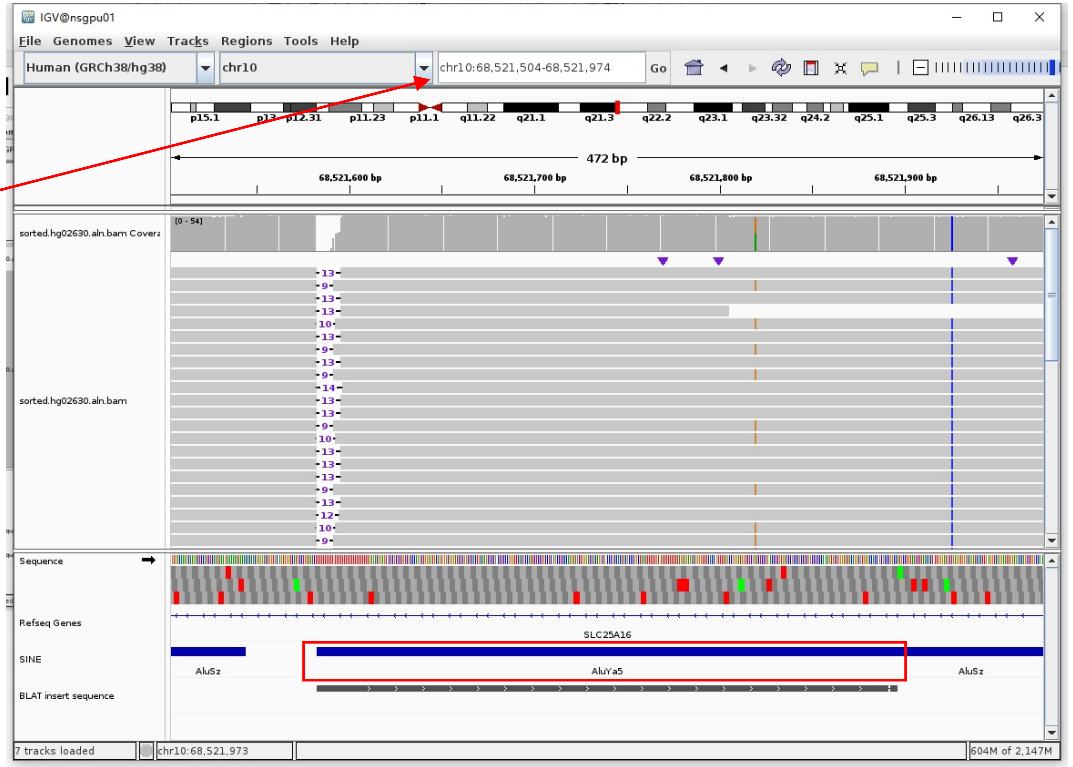

Insertion sequence (322bp)

- ➢ 9bp TSD

TATGCTTTT

- ➢ Insertion:37-322 aligns to *Alu*Ya5:282-1, 99.64% identity

TGAGACGGAGTCTCGCTCTGTCGCCCAGGCTGGAGTGCAGTGGCGGGATCTCGGCTCACTGCAAGCTCCGCCTCCCGGGTTCACGCCATTCTCCTGCCTCAGCCTCCCAAGTAGCTGGGACTA
CAGGCGCCCGCCACTACGCCCGGCTAATTTTTTGTATTTTTTTTTAGTAGAGACGGGGTTTCACCGTTTTAGCCAGGATGGTCTCGATCTCCTGACCTCGTGATCCGCCCGCCTCGGCCTCCC
AAAGTGCTGGGATTACAGGCGTGAGCCACCGCGCCGGCC

### HG02630 chr16:72351770

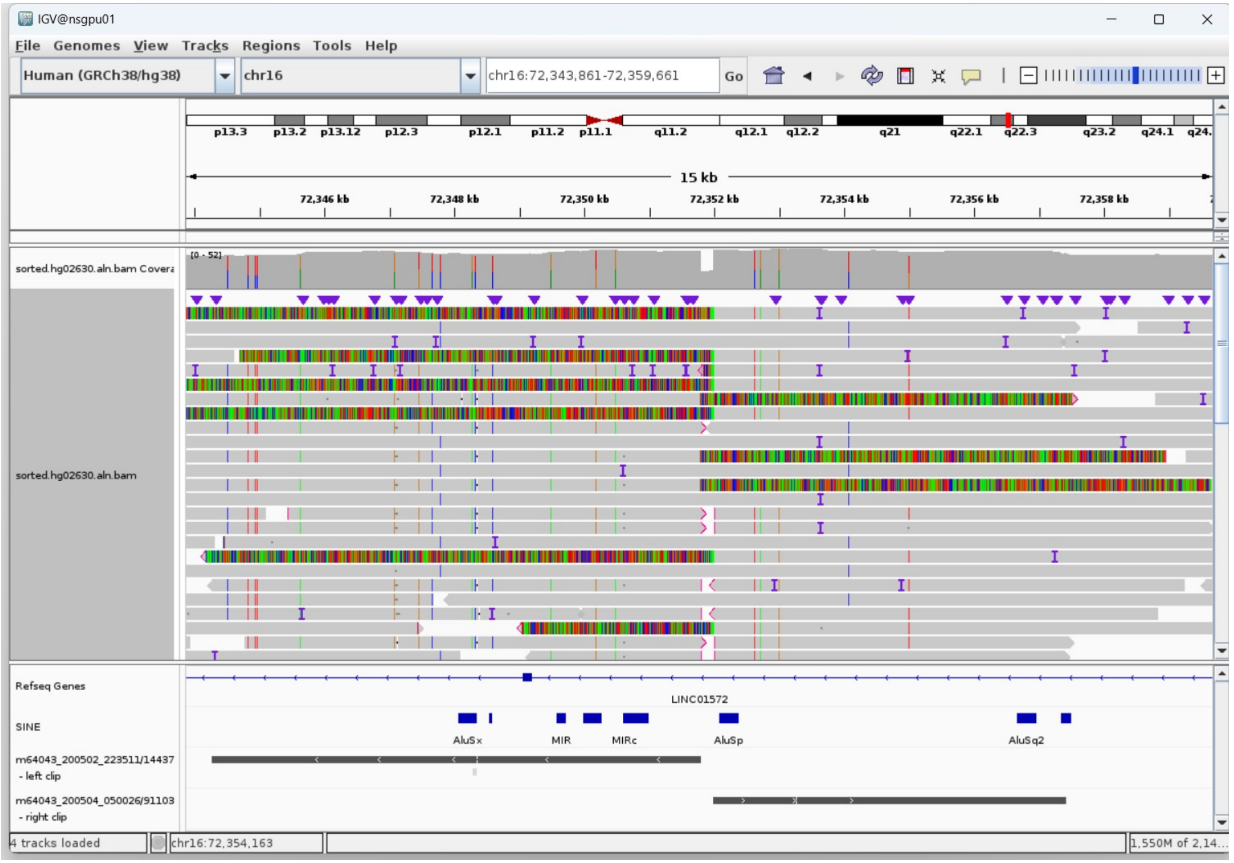


Right-clipped read (5722bp)

- Clip:1-284 aligns to *Alu*Yc1:1-282, 98.94% identity

TGGCCGGGCGCGGTGGCTCACGCCTGTAATCCCAGCACTTTGGGAGGCCGAGGCGGGT
GGATCACGAGGTCAGGAGATCGAGACCATCCTGGCTAACACGGTGAAACCCCGTCTC
TACTAAAAATACAAAAAATTAGCCGGGCGTGGTGGCGGGCGCCTGTAGTCCCAGCTAC
TCGGGAGGCTGAGGCAGGAGAATGGCGTGAACCCAGGAGGCGGAGCTTGCAGTGAGCC
GAGATCGCGCCACTGCACTCCAGCCTGGGCGACAGAGCGAGACTCCGTCTCA

- Clip:284-5722 aligns to chr16:72351994-72357416

### HG02630 chr17:37509872

- Mapping quality is 0, indicating that this region may have alignment issues.

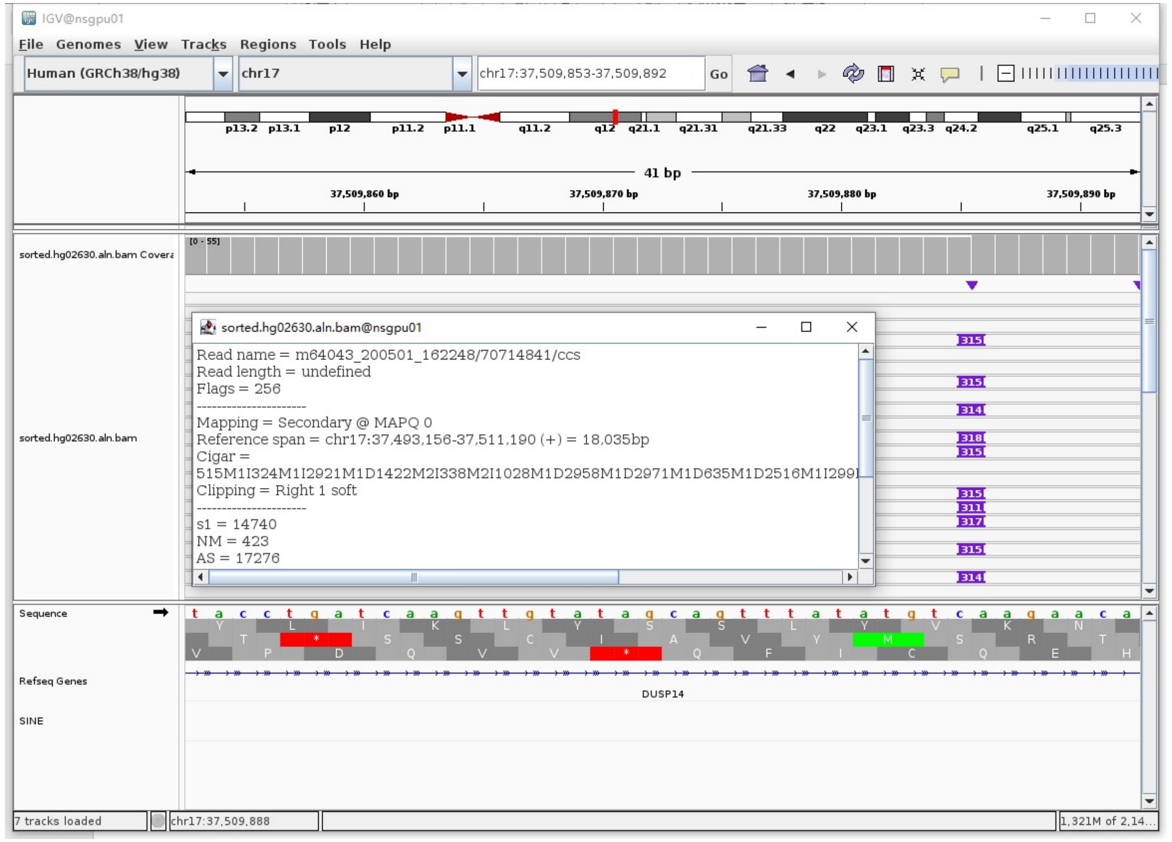


- Fetch one read (m64043_200501_162248/70714841/ccs, 18290 bp) and BLAT it to the hg38 genome.

  The best alignment is to chr17_KI270857v1_alt (an alternative sequence of chr17) and chr17.

| ACTIONS | QUERY | SCORE | START | END | QSIZE | IDENTITY | CHROM | STRAND | START | END | SPAN |
|---|---|---|---|---|---|---|---|---|---|---|---|
| browser new tab details | YourSeq | 6497 | 1 | 18290 | 18290 | 98.8% | chr17_KI270857v1_alt | + | 1732327 | 1750359 | 18033 |
| browser new tab details | YourSeq | 6493 | 1 | 18290 | 18290 | 98.8% | chr17 | + | 37493156 | 37511192 | 18037 |

- Read:1-16679 aligns to chr17_KI270857v1_alt

| ACTIONS | QUERY | SCORE | START | END | QSIZE | IDENTITY | CHROM | STRAND | START | END | SPAN |
|---|---|---|---|---|---|---|---|---|---|---|---|
| browser new tab details | YourSeq | 5947 | 1 | 16678 | 16678 | 99.1% | chr17_KI270857v1_alt | + | 1732328 | 1749063 | 16736 |
| browser new tab details | YourSeq | 5945 | 1 | 16678 | 16678 | 99.1% | chr17 | + | 37493157 | 37509896 | 16740 |

- Read:16679-16961 aligns to *Alu*Ya5:1-282, 99.65% identity

GGCCGGGCGCGGTGGCTCACGCCTGTAATCCCAGCACTTTGGGAGGCCGAGGCGGGCGGATCACGAGGTCAGGA
GATCGAGACCATCCCGGCTAAAACGGTGAAACCCCGTCTCTACTAAACATACAAAAAATTAGCCGGGCGTAGTG
GCGGGCGCCTGTAGTCCCAGCTACTTGGGAGGCTGAGGCAGGAGAATGGCGTGAACCCGGGAGGCGGAGCTTGC
AGTGAGCCGAGATCCCGCCACTGCACTCCAGCCTGGGCGACAGAGCGAGACTCCGTCTCA

11bp TSD

AAGAACACCTC

- Read:16962-18290 also aligns to chr17_KI270857v1_alt

| ACTIONS | QUERY | SCORE | START | END | QSIZE | IDENTITY | CHROM | STRAND | START | END | SPAN |
|---|---|---|---|---|---|---|---|---|---|---|---|
| browser new tab details | YourSeq | 1305 | 22 | 1328 | 1328 | 100.0% | chr17_KI270857v1_alt | + | 1749053 | 1750359 | 1307 |
| browser new tab details | YourSeq | 1303 | 22 | 1328 | 1328 | 99.9% | chr17 | + | 37509886 | 37511192 | 1307 |

- It's an *Alu*Ya5 insertion.

### HG02630 chr17:37561902

- The same genomic coordinate as in HG00621 (page 15)

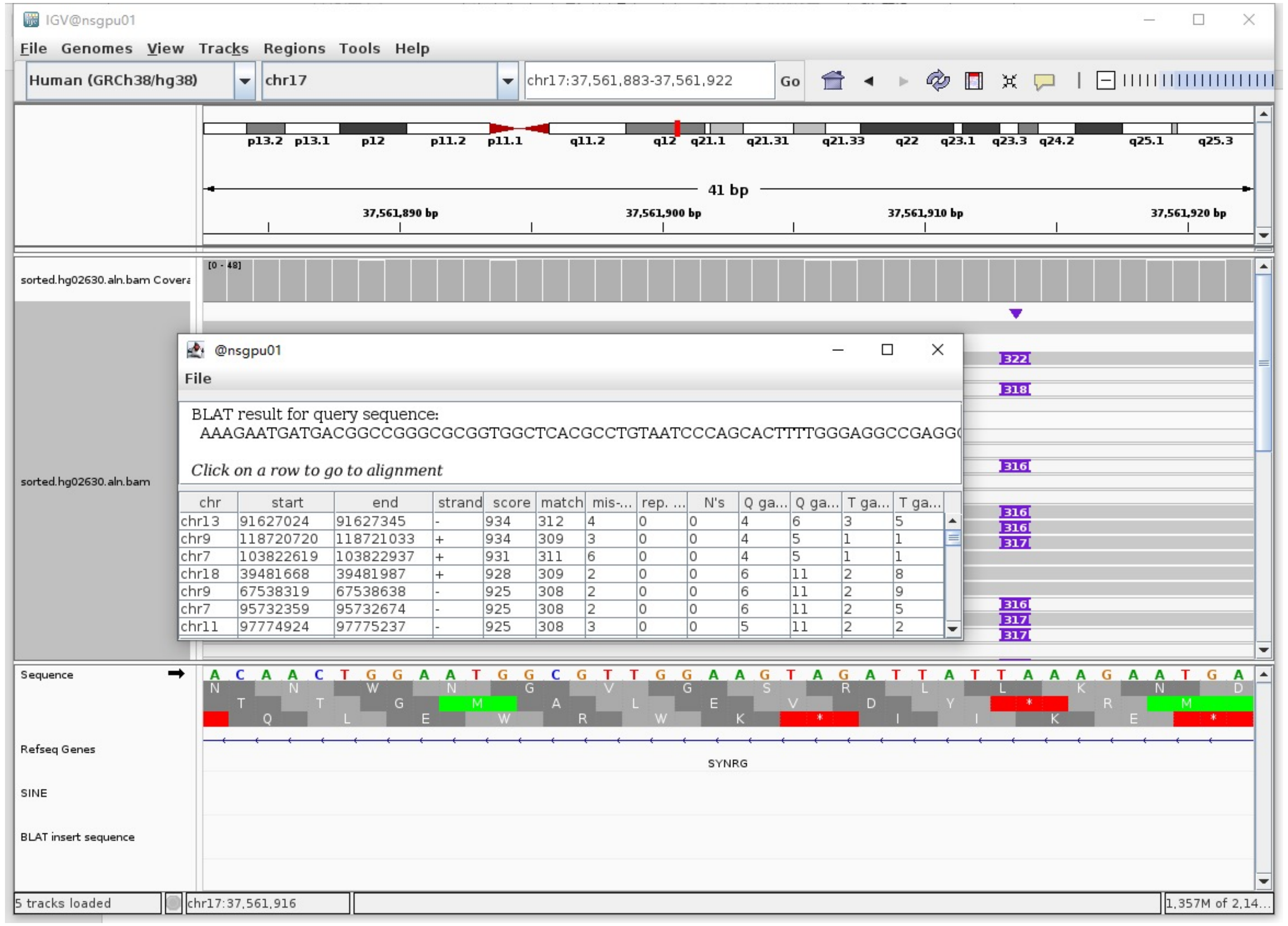

### HG02630 chr17:37660775

➢ Using the built-in BLAT tool in IGV to check where the inserted sequence aligned.

➢ The annotation at the aligned location indicated it as *Alu*Y.

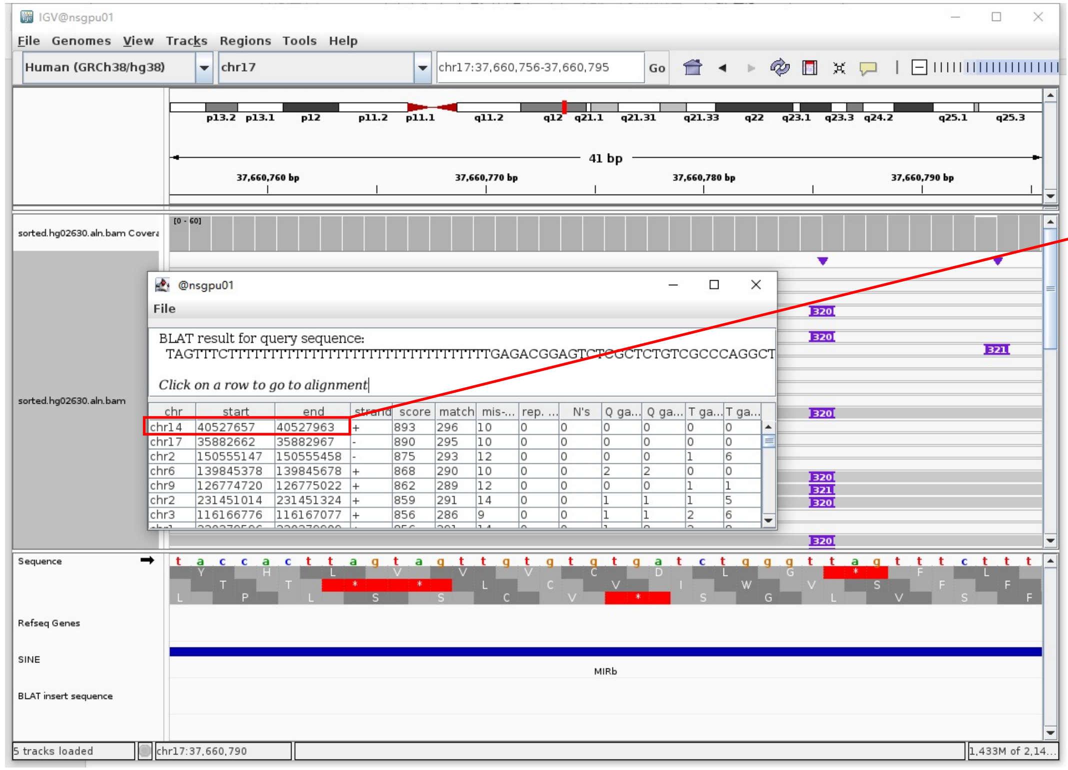


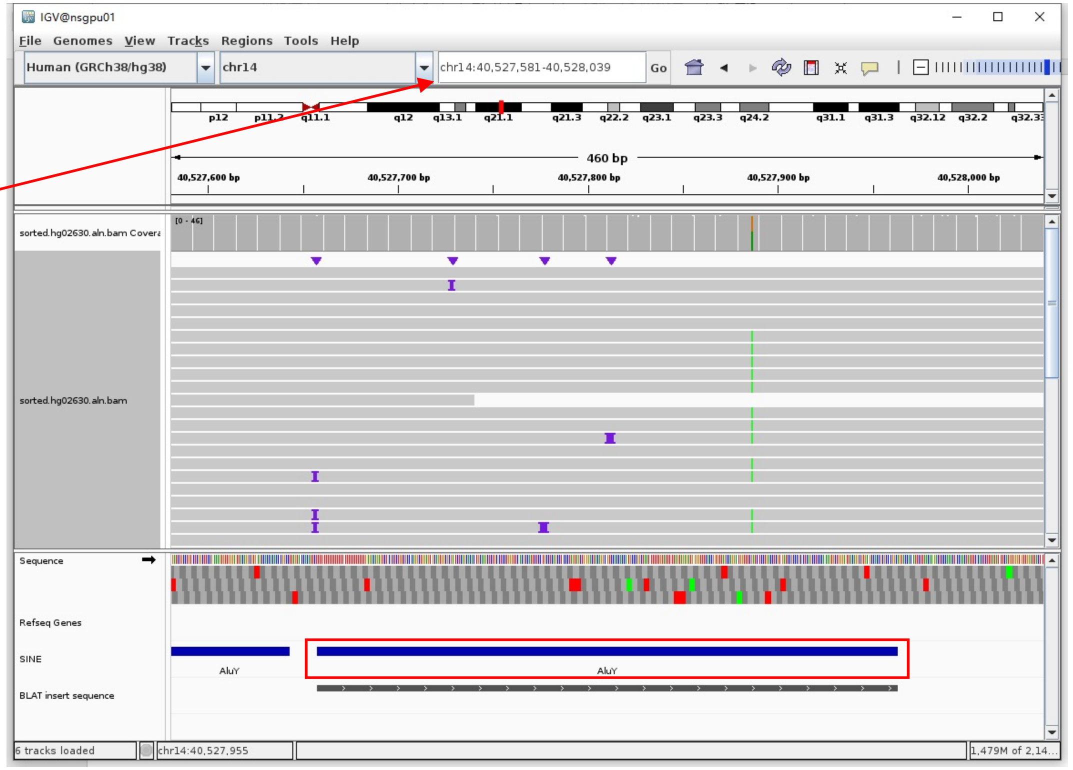


Insertion sequence (320bp)

➢ 14bp TSD

TAGTTTCTTTTTTT

➢ Insertion:36-318 aligns to *Alu*Yc1:282-1, 98.94% identity

TGAGACGGAGTCTCGCTCTGTCGCCCAGGCTGGAGTGCAGTGGCGCGATCTCAGCTCACTGCAAGCTCCGCCTCCTGGGTTCACGCCATTCTCCTGCCTCAGCCTCCCGAGTAGCTGGGACTA
CAGGCGCCCGCCACCACGCCCGGCTAATTTTTTGTATTTTTAGTAGAGACGGGGTTTCACCGTGTTAGCCAGGATGGTCTCGATCTCCTGACCTCGTGATCCGCCCGCCTCGGCCTCCCAAAG
TGCTGGGATTACAGGCGTGAGCCACCGCGCCCGGCC

### HG02630 chr17:37883970

- Using the built-in BLAT tool in IGV to check where the inserted sequence aligned.
- The annotation at the aligned location indicated it as *AluY*.

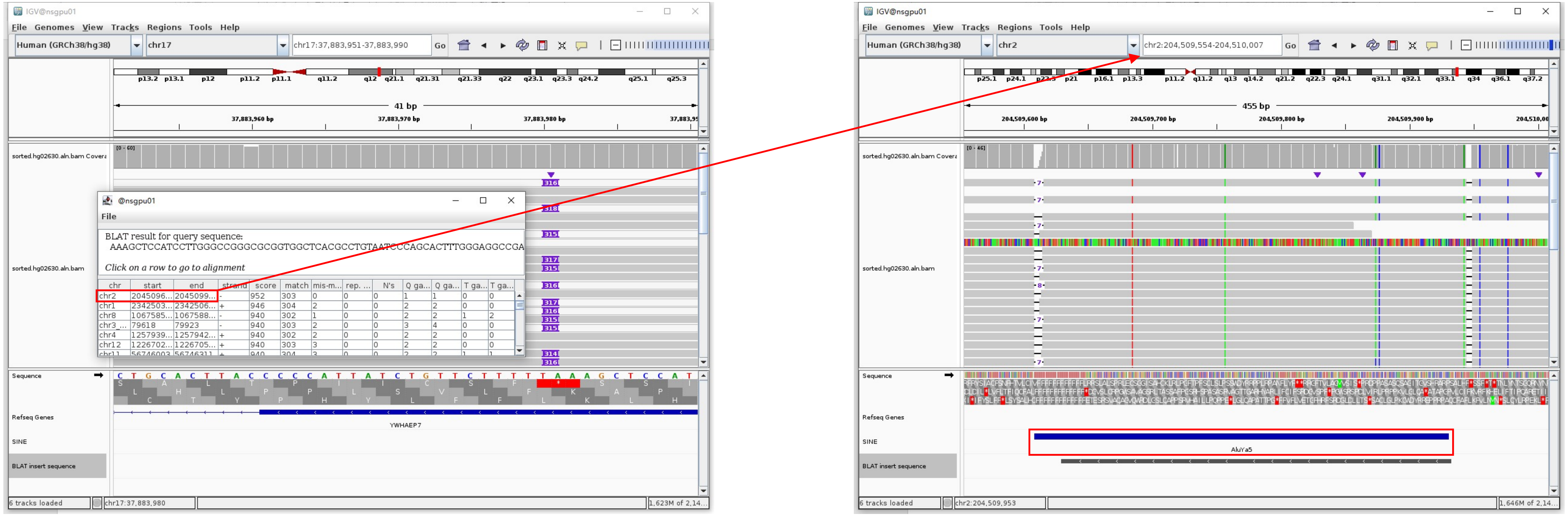


Insertion sequence (316bp)

- 14bp TSD

AAAGCTCCATCCTT

- Insertion:15-289 aligns to *Alu*Ya5:1-282, 99.29% identity

GGCCGGGCGCGGTGGCTCACGCCTGTAATCCCAGCACTTTGGGAGGCCGAGGCAGGCGGATCACGAGGTCAGGAGATCGAGACCATCCCGGCTAAAACGGTGAAACCCCGTCTCTACTAAAAA
TACAAAAAATTAGCCGGGCGTAGTGGCGGGCGCCTGTAGTCCCAGCTACTCGGGAGGCTGAGGCAGGAGAATGGCGTGAACCCGGGAAGGCGGAGCTTGCAGTGAGCCGAGATCCCGCCACTG
CACTCCAGCCTGGGCGACAGAGCGAGACTCCGTCTCA.........poly(A)

### HG02630 chr17:38528853

- ➢ Using the built-in BLAT tool in IGV to check where the inserted sequence aligned.
- ➢ The annotation at the aligned location indicated it as *AluY*.

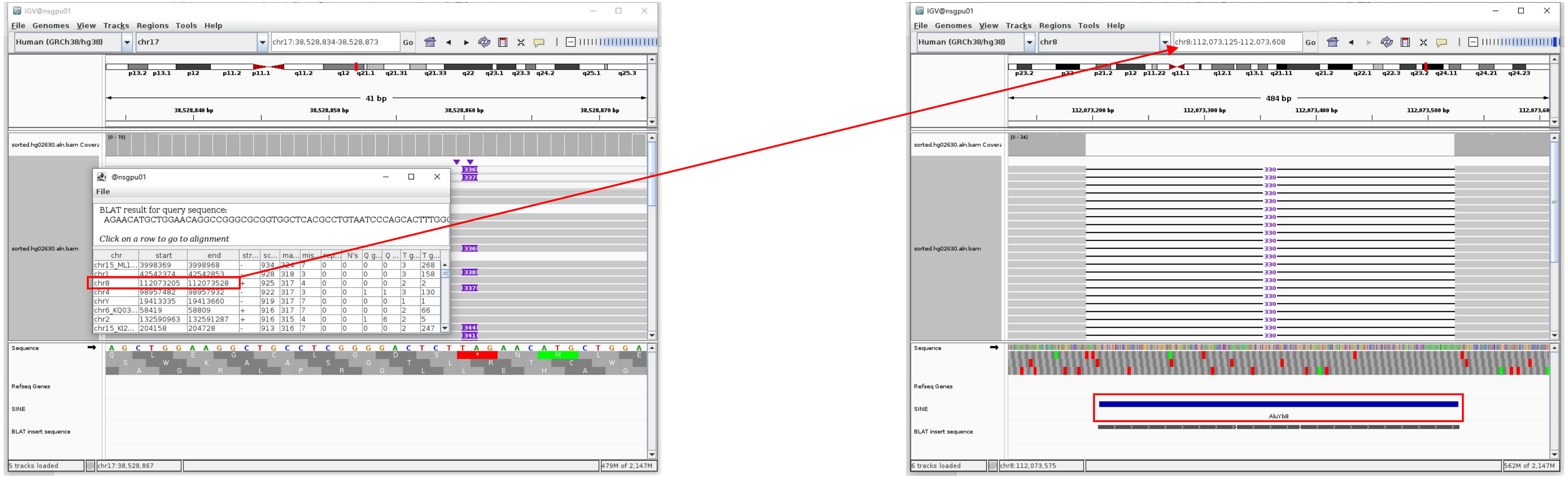


Insertion sequence (336bp)

- ➢ 16bp TSD

GAACATGCTGGAACAG

- ➢ Insertion:18-314 aligns to *Alu*Yb8:1-289, 98.96% identity

GGCCGGGCGCGGTGGCTCACGCCTGTAATCCCAGCACTTTGGGAGGGCCGAGGCGGGTGGATCATGAGGTCAGGAGATCGAGACCATCCTGGCTAACAAGGTGAAACCCCGTCTCTACTAAAA
ATACAAAAAAAAATTAGCCGGGCGCGGTGGCGGGCGCCTGTAGTCCCAGCTACTCGGGAGGCTGAGGCAGGAGAAATGGCGTGAACCCGGGAAGCGGAGCTTGCAGTGAGGCCGAGATTGCGC
CACTGCAGTCCGCAGTCCCGCGGGGGGCGACAGAGCGAGACTCCGTCTCA………poly(A)

### HG03492 chr1:198500196

➢ Mapping quality is 0, indicating that this region may have alignment issues.

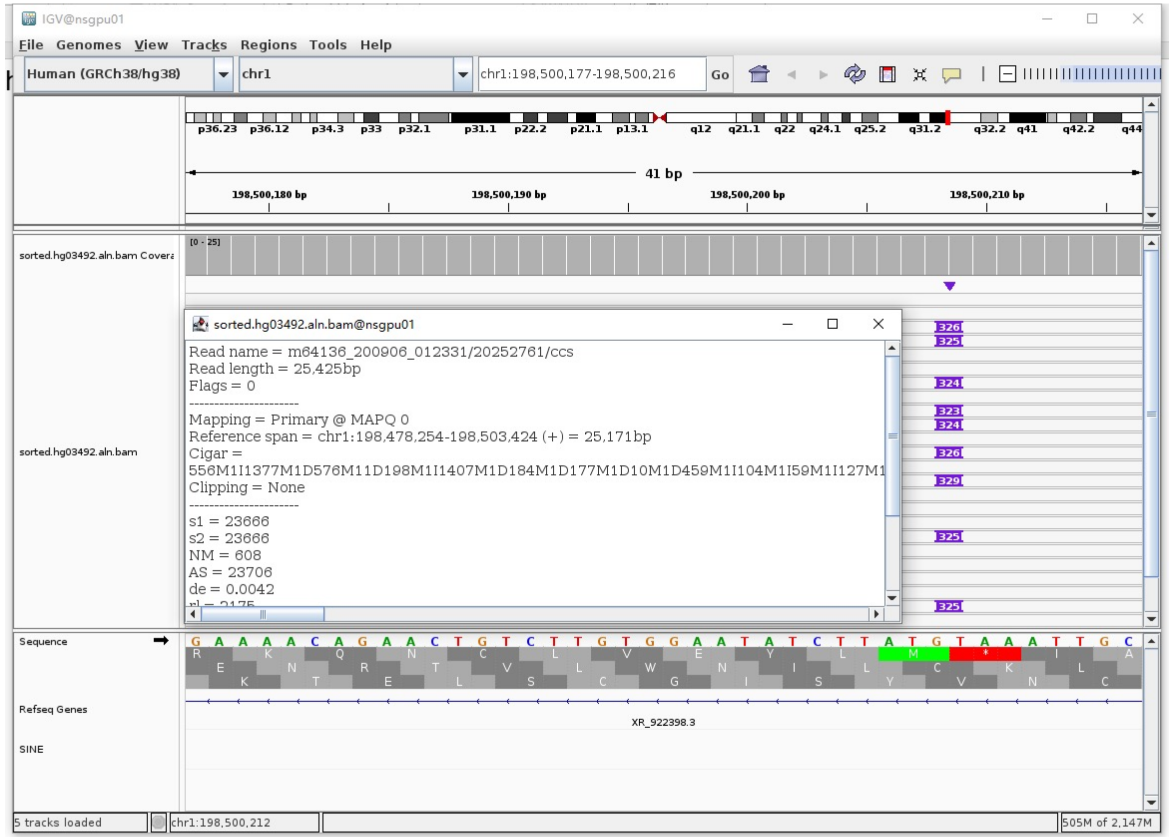


➢ Fetch one read (m64136_200906_012331/20252761/ccs, 25411 bp) and BLAT it to the hg38 genome.

The best alignment is to chr1_GL383520v2_alt (an alternative sequence of chr1) and chr1.

| ACTIONS | QUERY | SCORE | START | END | QSIZE | IDENTITY | CHROM | STRAND | START | END | SPAN |
|---|---|---|---|---|---|---|---|---|---|---|---|
| browser new tab details | YourSeq | 23628 | 1 | 25411 | 25411 | 99.8% | chr1_GL383520v2_alt | + | 108172 | 133342 | 25171 |
| browser new tab details | YourSeq | 23628 | 1 | 25411 | 25411 | 99.8% | chr1 | + | 198478254 | 198503424 | 25171 |

- Read:1-21877 aligns to chr1_GL383520v2_alt

| ACTIONS | QUERY | SCORE | START | END | QSIZE | IDENTITY | CHROM | STRAND | START | END | SPAN |
|---|---|---|---|---|---|---|---|---|---|---|---|
| browser new tab details | YourSeq | 20446 | 1 | 21848 | 21876 | 99.9% | chr1_GL383520v2_alt | + | 108173 | 130138 | 21966 |
| browser new tab details | YourSeq | 20446 | 1 | 21848 | 21876 | 99.9% | chr1 | + | 198478255 | 198500220 | 21966 |

- Read:21878-22160 aligns to *Alu*Ya5:282-1, 99.29% identity

TGAGACGGAGTCTTGCTCTGTCGCCCAGGCTGGAGTGCAGTGGCGGATCTCGGCTCACTGCAAGCTCCGCCTC
CCGGGTTCACGCCATTCTCCTGCCTCAGCCTCCCAAGTAGCTGGGACTACAGGCGCCCGCCACTACGCCCGGCT
AATTTTTTGTATTTTTAGTAGAGACGGGGTTTCACCGTTTTAGCCGGGATGGTCTCGATCTCCTGACCTCGTGA
TCCACCCGCCTCGGCCTCCCAAAGTGCTGGGATTACAGGCGTGAGCCACCGCGCCCGGCC

12bp TSD

TAAATTGCTCTT

- Read:16962-18290 also aligns to chr1_GL383520v2_alt

| ACTIONS | QUERY | SCORE | START | END | QSIZE | IDENTITY | CHROM | STRAND | START | END | SPAN |
|---|---|---|---|---|---|---|---|---|---|---|---|
| browser new tab details | YourSeq | 3193 | 1 | 3250 | 3250 | 99.3% | chr1_GL383520v2_alt | + | 130128 | 133342 | 3215 |
| browser new tab details | YourSeq | 3193 | 1 | 3250 | 3250 | 99.3% | chr1 | + | 198500210 | 198503424 | 3215 |

➢ It's an *Alu*Ya5 insertion.

### HG03492 chr3:40200107

- Using the built-in BLAT tool in IGV to check where the inserted sequence aligned.

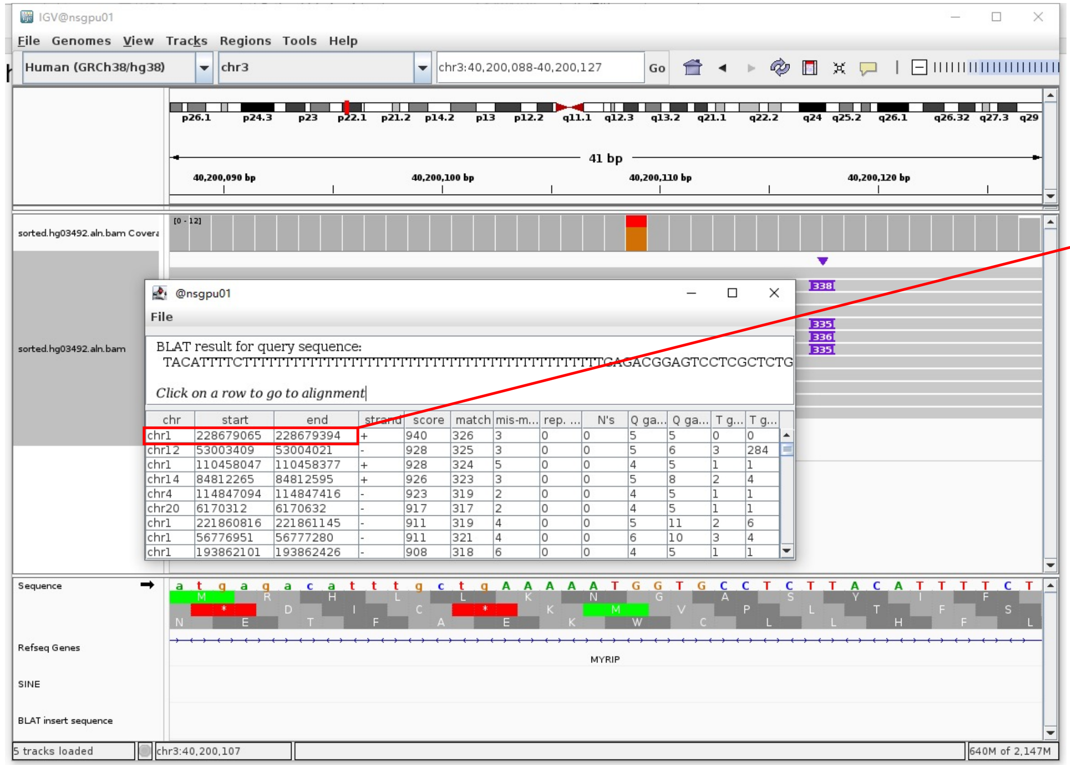


- The annotation at the aligned location indicated it as *Alu*Y.

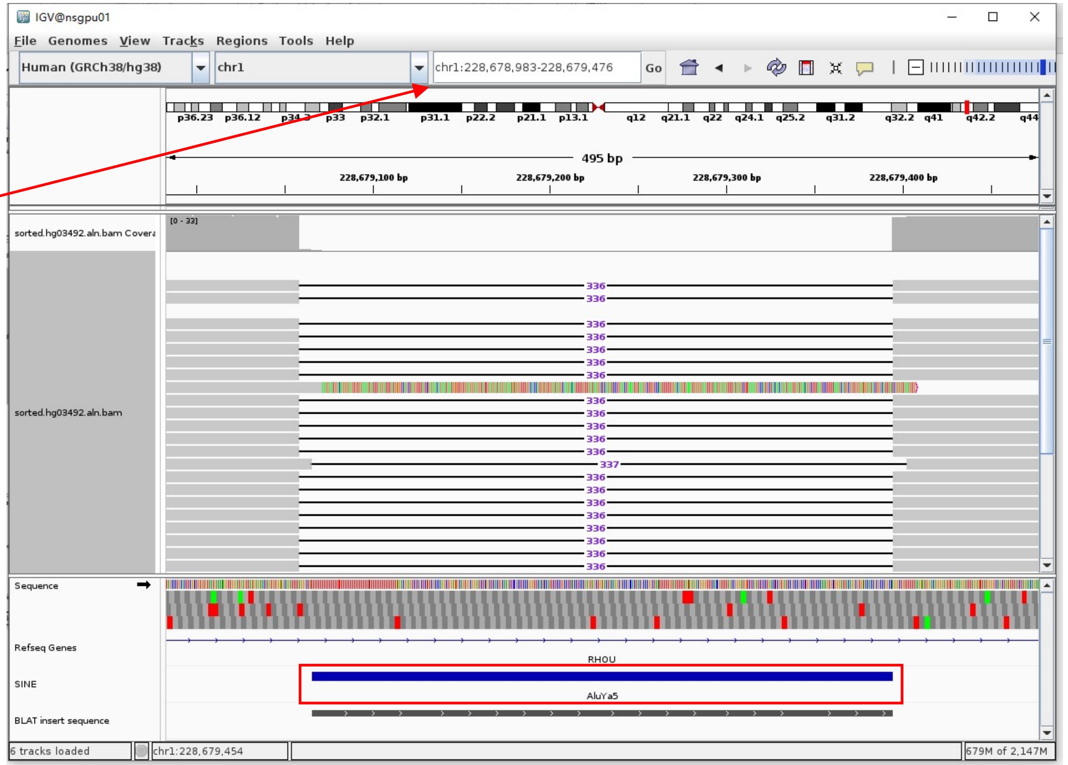


Insertion sequence (338bp)

- 14bp TSD

TACATTTTCTTTTT

- Insertion:52-338 aligns to *Alu*Ya5:282-1, 98.58% identity

TGAGACGGAGTCCTCGCTCTGTCGCCCAGGCTGGAGTGCAGTGGCGGGATCTCGGCTCACTGCAAGCTCCGCCTCCCGGGTTCACGCCATTCTCCTGCCTCAGCCTCCCCGAGTAGCTGGGAC
TACAGGCGCCCGCCACCACGCCCGGCTAATTTTTTGTATTTTCAGTAGAGACGGGGTTTCACCGTTTTAGCCGGGATGGTCTCGATCTCCTGACCTCGTGATCCGCCCGCTCTCGGCCTCCC
AAAGTGCTGGGATTACAGGCGTGAGCCACCACGCCCGGCC

### HG03492 chr4:21604842

- Using the built-in BLAT tool in IGV to check where the inserted sequence aligned.

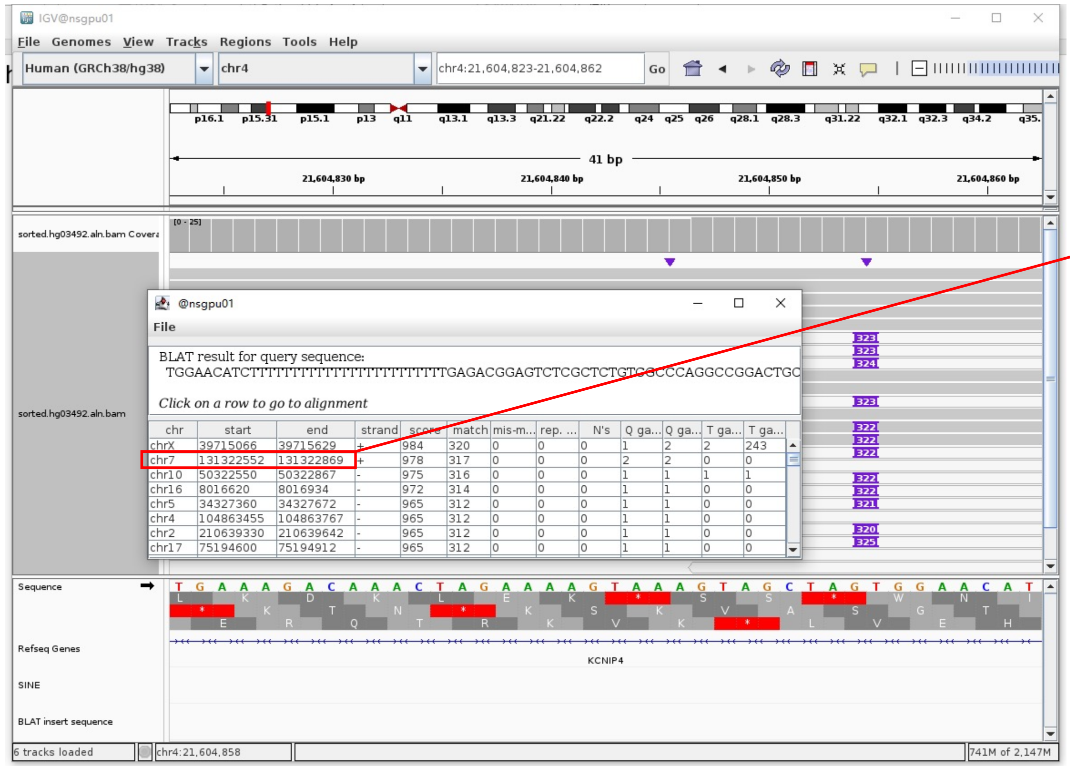


- The annotation at the aligned location indicated it as *AluY*.

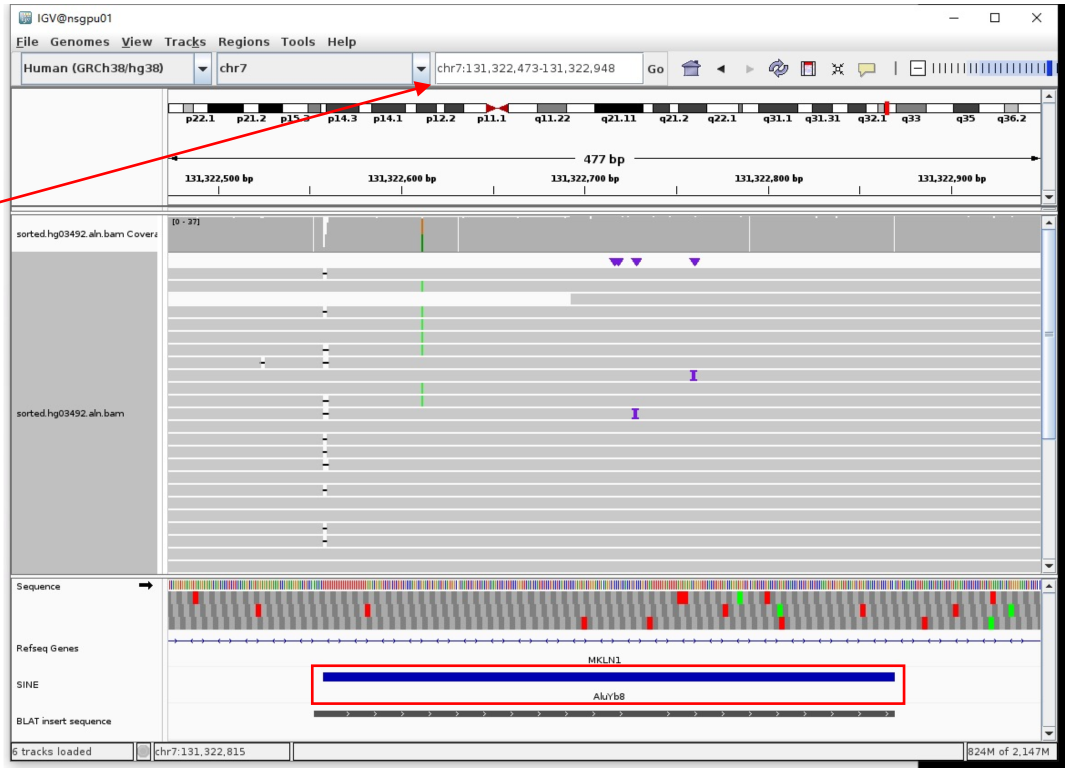


Insertion sequence (322bp)

- 12bp TSD

TGGAACATCTTT

- Insertion:32-322 aligns to *Alu*Yb8:289-0, 100% identity

TGAGACGGAGTCTCGCTCTGTCGCCCAGGCCGGACTGCGGACTGCAGTGGCGCAATCTCGGCTCACTGCAAGCTCCGCTTCCCGGGTTCACGCCATTCTCCTGCCTCAGCCTCCCGAGTAGCT
GGGACTACAGGCGCCCGCCACCGCGCCCGGCTAATTTTTTGTATTTTTAGTAGAGACGGGGTTTCACCTTGTTAGCCAGGATGGTCTCGATCTCCTGACCTCATGATCCACCCGCCTCGGCC
TCCCAAAGTGCTGGGATTACAGGCGTGAGCCACCGCGCCCGGCC

### HG03492 chr6:30030695

- Mapping quality is 0, indicating that this region may have alignment issues.

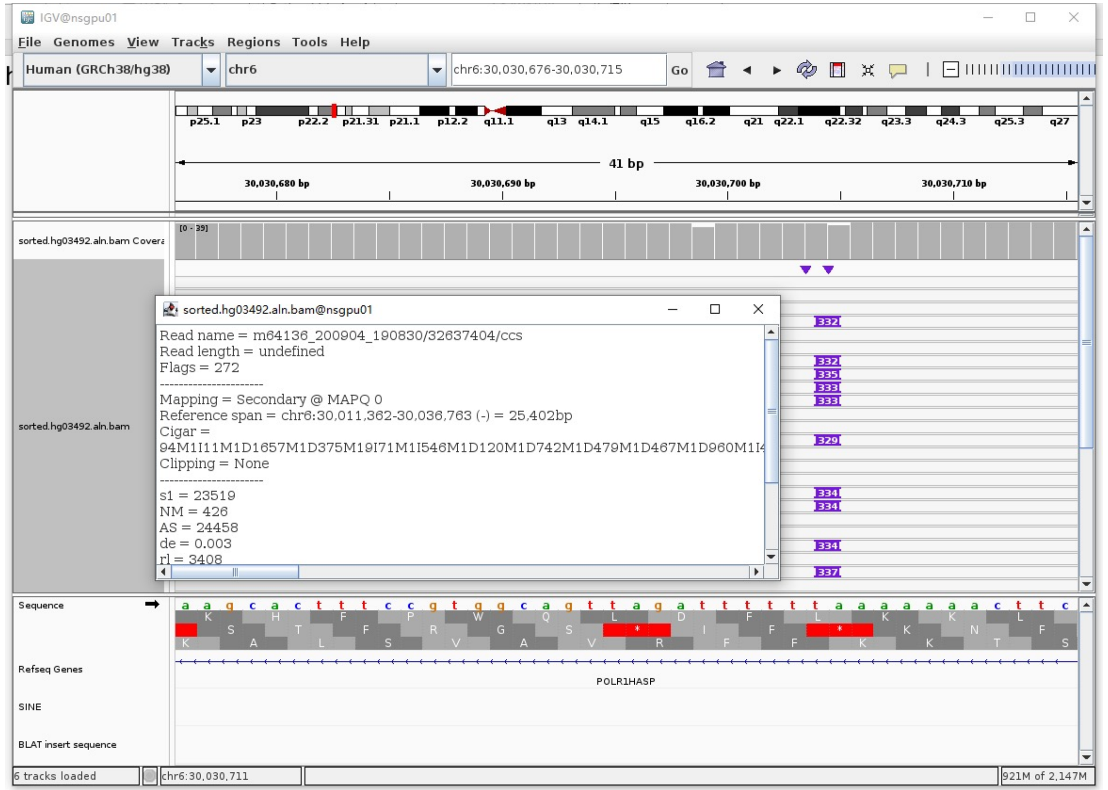


- Fetch one read (m64136_200904_190830/32637404/ccs, 25746 bp) and BLAT it to the hg38 genome.

  The best alignment is to chr6_GL000252v2_alt (an alternative sequence of chr6), which contains an *Alu*Yb8 element.

| ACTIONS | | | QUERY | SCORE | START | END | QSIZE | IDENTITY | CHROM | STRAND | START | END | SPAN |
|---|---|---|---|---|---|---|---|---|---|---|---|---|---|
| browser | new tab | details | YourSeq | 25727 | 1 | 25746 | 25746 | 100.0% | chr6_GL000252v2_alt | - | 1266897 | 1292649 | 25753 |
| browser | new tab | details | YourSeq | 25727 | 1 | 25746 | 25746 | 100.0% | chr6_GL000250v2_alt | - | 1269071 | 1294823 | 25753 |
| browser | new tab | details | YourSeq | 25723 | 1 | 25746 | 25746 | 100.0% | chr6_GL000251v2_alt | - | 1490747 | 1516498 | 25752 |
| browser | new tab | details | YourSeq | 25685 | 1 | 25746 | 25746 | 100.0% | chr6_GL000256v2_alt | - | 1309169 | 1335659 | 26491 |
| browser | new tab | details | YourSeq | 25344 | 1 | 25746 | 25746 | 99.9% | chr6_GL000254v2_alt | - | 1355418 | 1380837 | 25420 |
| browser | new tab | details | YourSeq | 25338 | 1 | 25746 | 25746 | 99.9% | chr6_GL000253v2_alt | - | 1272093 | 1297513 | 25421 |
| browser | new tab | details | YourSeq | 25316 | 1 | 25746 | 25746 | 99.8% | chr6 | - | 30011362 | 30036763 | 25402 |

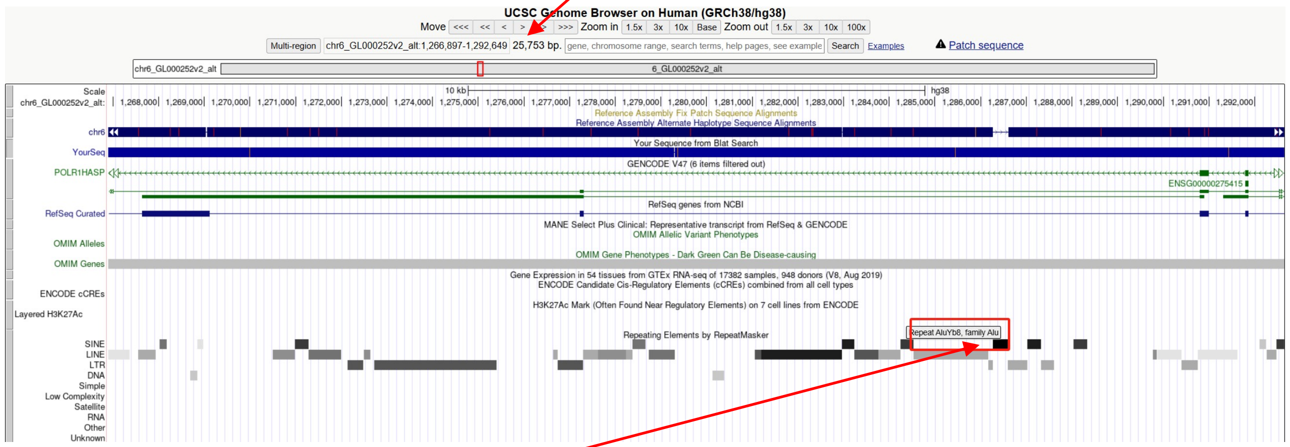


chr6_GL000252v2_alt:1286275-1286592 (*Alu*Yb8 marked by RepeatMasker)

- Read:6085-6375 aligns to *Alu*Yb8:289-1, 99.65% identity

  TTGAGACGGAGTCTCGCTCTGTCGCCCAGGCCGTACTGCGGACTGCAGTGGCGCAATCTCGGCTCACTGCAAGCTCC
  GCTTCCCGGGTTCACGCCATTCTCCTGCCTCAGCCTCCCGAGTAGCTGGGACTACAGGCGCCCGCCACCGCGCCCGG
  CTAATTTTTTGTATTTTTAGTAGAGACGGGGTTTCACCTTGTTAGCCAGGATGGTCTCGATCTCCTGACCTCATGAT
  CCACCCGCCTCGGCCTCCCAAAGTGCTGGGATTACAGGCGTGAGCCACCGCGCCCGGCC

- It's an *Alu*Yb8 in the chromosome chr6_GL000252v2_alt.

### HG03492 chr6:32720794 (False)

- Mapping quality is 0, indicating that this region may have alignment issues.

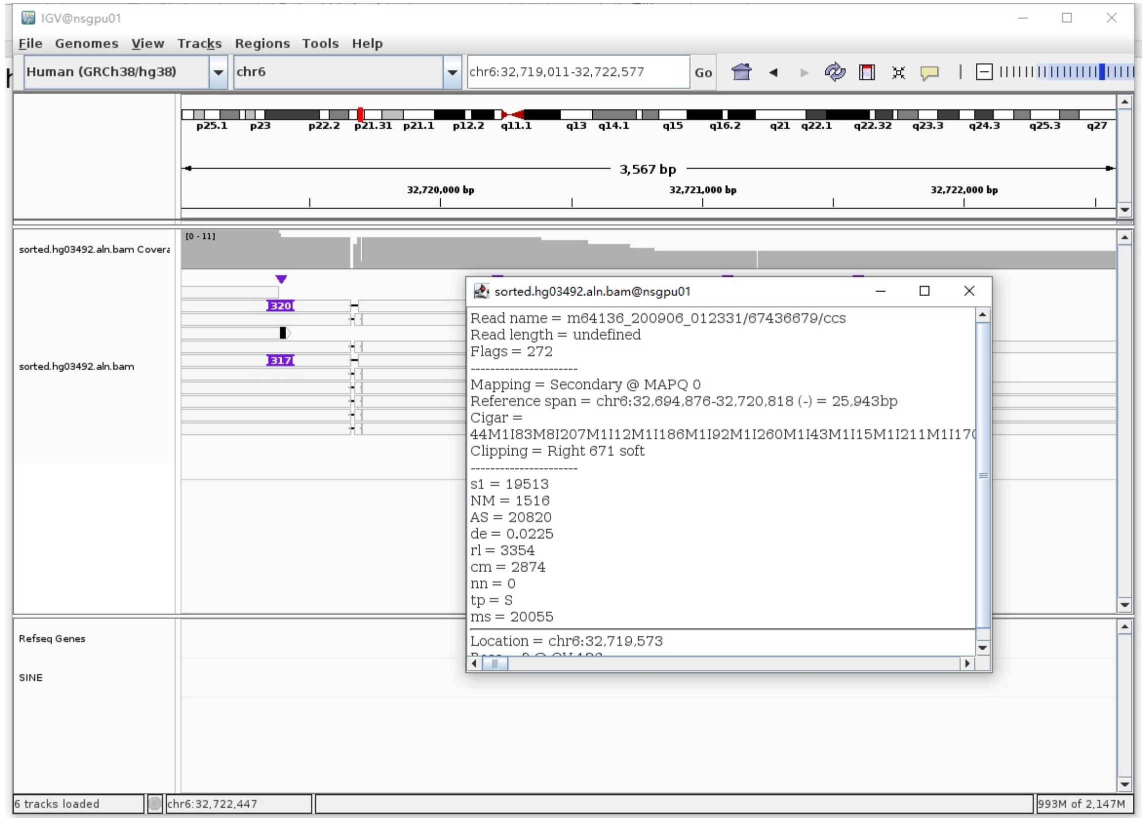


- Fetch one read (m64136_200906_012331/67436679/ccs, 26391 bp) and BLAT it to the hg38 genome.

  The best alignment is to chr6_GL000252v2_alt (an alternative sequence of chr6), which contains an *Alu*Y element.

| ACTIONS | | | QUERY | SCORE | START | END | QSIZE | IDENTITY | CHROM | STRAND | START | END | SPAN |
|---|---|---|---|---|---|---|---|---|---|---|---|---|---|
| browser | new tab | details | YourSeq | 26318 | 1 | 26391 | 26391 | 99.9% | chr6_GL000252v2_alt | - | 3939503 | 3965883 | 26381 |
| browser | new tab | details | YourSeq | 26189 | 1 | 26391 | 26391 | 99.7% | chr6_GL000254v2_alt | - | 3994138 | 4020505 | 26368 |
| browser | new tab | details | YourSeq | 25269 | 1 | 26391 | 26391 | 99.2% | chr6_GL000253v2_alt | - | 4114723 | 4140425 | 25703 |
| browser | new tab | details | YourSeq | 24826 | 1 | 26391 | 26391 | 98.4% | chr6_GL000251v2_alt | - | 4108202 | 4134396 | 26195 |
| browser | new tab | details | YourSeq | 24572 | 1 | 26391 | 26391 | 98.3% | chr6_GL000255v2_alt | - | 3889166 | 3915374 | 26209 |
| browser | new tab | details | YourSeq | 24546 | 1 | 26391 | 26391 | 98.3% | chr6 | - | 32694876 | 32721127 | 26252 |

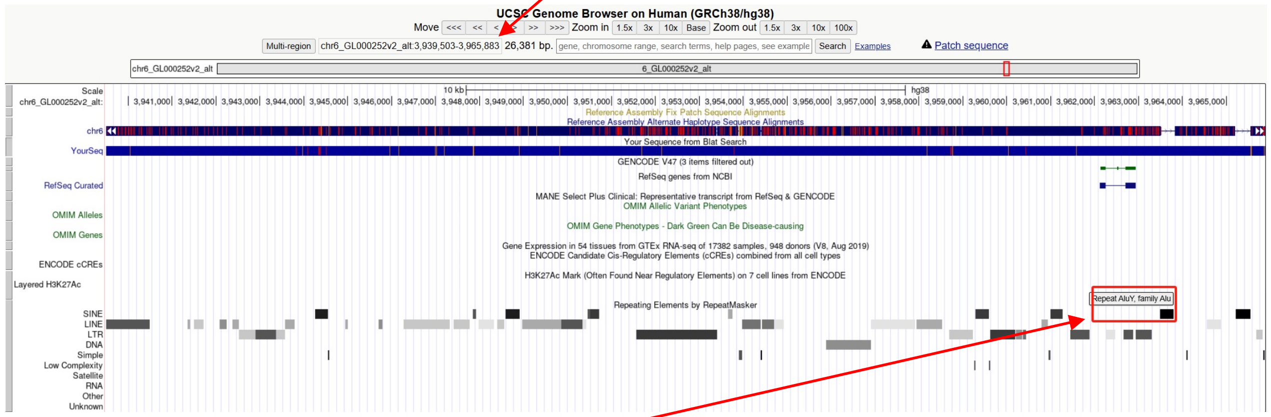


chr6_GL000252v2_alt:3963497-3963802 (*Alu*Y marked by RepeatMasker)

- Read:2085-2367 aligns to *Alu*Yb8:1-282, 98.58% identity

GGCCGGGCGCGGTGGCTCACGCCTGTAATCCCAGCACTTTGGGAGGCCGAGGCGGGCGGATCACGAGGTCAGGA
GATCGAGACCATCCTGGCTAACACGGTGAAACCCCGTCTCTACTAAAAATACAAAAAATTAGCCGGGCGTGGTA
GCGGGCACCTGTAGTTCCAGCTACTCGGGAGGCTGAGGCAGGAGAATGGTGTGAACCCGGGAGGTGGAGCTTGC
AGTGAGCCGAGATCGCGCCACTGCACTCCAGCCTGGGCGACAGAGCGAGACTCCGTCTCA

- It's an *Alu*Y in the chr6_GL000252v2_alt.

### HG03492 chr6:79377018

➢ Mapping quality is 0, indicating that this region may have alignment issues.

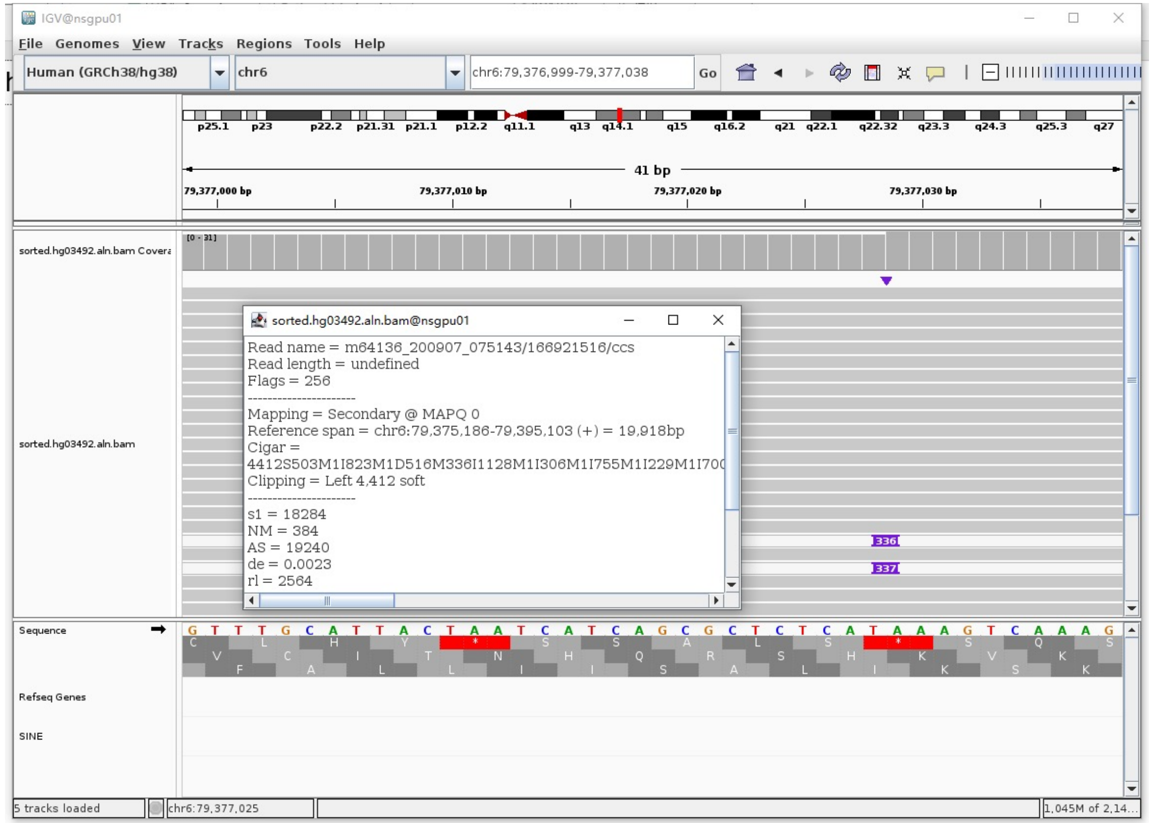


➢ Fetch one read (m64136_200907_075143/166921516/ccs, 24658 bp) and BLAT it to the hg38 genome.

The best alignment is to chr6_GL383533v1_alt (an alternative sequence of chr1).

| ACTIONS | | | QUERY | SCORE | START | END | QSIZE | IDENTITY | CHROM | STRAND | START | END | SPAN |
|---|---|---|---|---|---|---|---|---|---|---|---|---|---|
| browser | new tab | details | YourSeq | 24249 | 1 | 24658 | 24658 | 99.9% | chr6_GL383533v1_alt | - | 51809 | 76135 | 24327 |
| browser | new tab | details | YourSeq | 19867 | 4409 | 24658 | 24658 | 99.9% | chr6 | + | 79375186 | 79395103 | 19918 |

● Read:1-6297 aligns to chr6_GL383533v1_alt

| ACTIONS | | | QUERY | SCORE | START | END | QSIZE | IDENTITY | CHROM | STRAND | START | END | SPAN |
|---|---|---|---|---|---|---|---|---|---|---|---|---|---|
| browser | new tab | details | YourSeq | 6235 | 1 | 6264 | 6296 | 99.9% | chr6_GL383533v1_alt | - | 69865 | 76134 | 6270 |
| browser | new tab | details | YourSeq | 1853 | 4408 | 6264 | 6296 | 99.9% | chr6 | + | 79375186 | 79377042 | 1857 |

● Read:6298-6587 aligns to *Alu*Yb8:289-1, 99.65% identity

TGAGACGGAGTCTCGCTCTGTTGCCCAGGCCGGACTGCGGACTGCAGTGGCGCAATCTCGGCTCACTGCAAGCT
CCGCTTCCCGGGTTCACGCCATTCTCCTGCCTCAGCCTCCCGAGTAGCTGGGACTACAGGCGCCCGCCACCGCG
CCCGGCTAATTTTTTGTATTTTTAGTAGAGACGGGGTTTCACCTTGTTAGCCAGGATGGTCTCGATCTCCTGAC
CTCATGATCCACCCGCCTCGGCCTCCCAAAGTGCTGGGATTACAGGCGTGAGCCACCGCGCCCGGCC

13bp TSD

AAAGTCAAAGTTT

● Read:6588-24658 also aligns to chr6_GL383533v1_alt

| ACTIONS | | | QUERY | SCORE | START | END | QSIZE | IDENTITY | CHROM | STRAND | START | END | SPAN |
|---|---|---|---|---|---|---|---|---|---|---|---|---|---|
| browser | new tab | details | YourSeq | 18028 | 1 | 18070 | 18070 | 99.9% | chr6 | + | 79377030 | 79395103 | 18074 |
| browser | new tab | details | YourSeq | 18027 | 1 | 18070 | 18070 | 99.9% | chr6_GL383533v1_alt | - | 51809 | 69877 | 18069 |

➢ It's an *Alu*Yb8 insertion.

### HG03492 chr6:104041958

- The same genomic coordinate as in HG00438 (page 6)

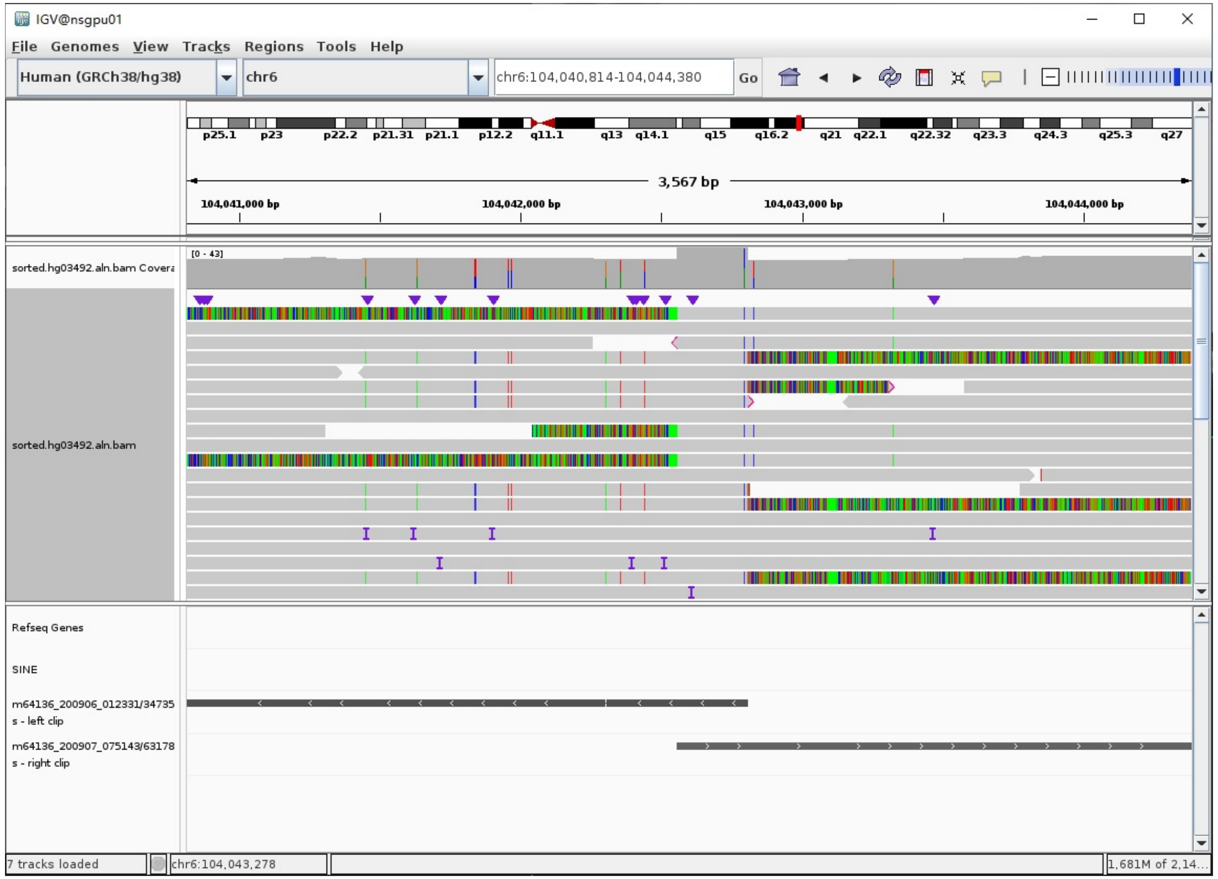

### HG03492 chr8:1362327

- Using the built-in BLAT tool in IGV to check where the inserted sequence aligned.

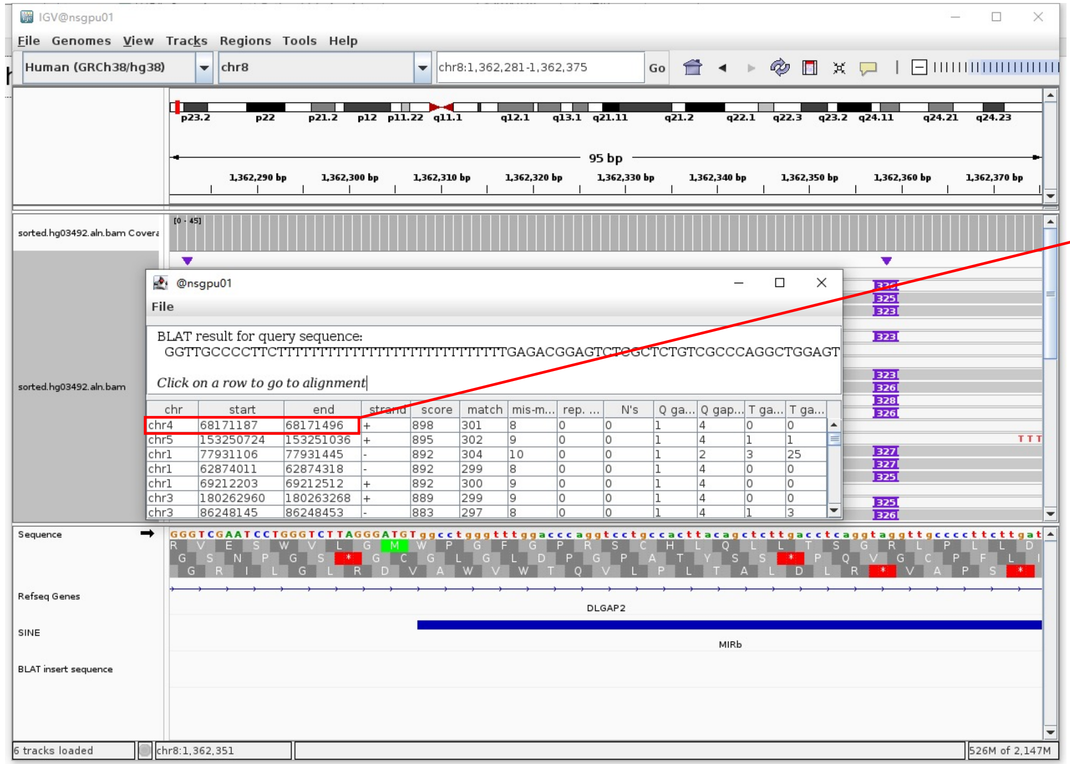


- The annotation at the aligned location indicated it as *AluY*.

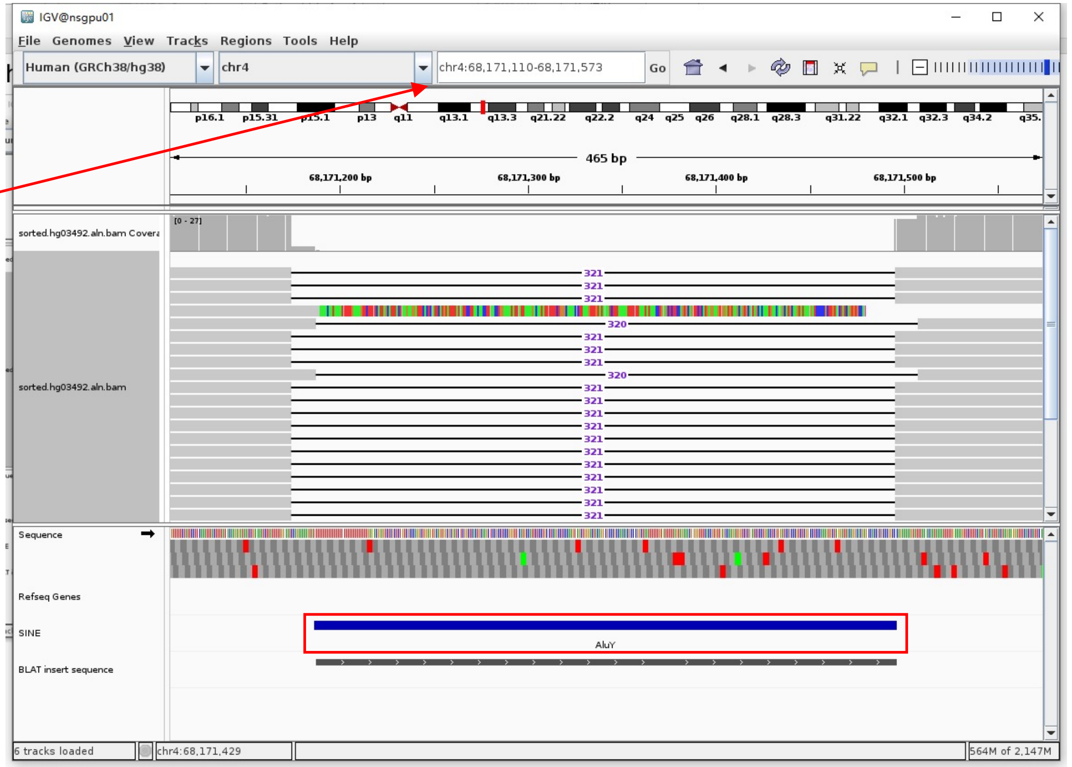


Insertion sequence (325bp)

- 14bp TSD

GGTTGCCCCTTCTT

- Insertion:39-325 aligns to *Alu*Yc1:282-1, 99.29% identity

TGAGACGGAGTCTCGCTCTGTCGCCCAGGCTGGAGTGCAGTGGCGCGATCTCGGCTCACTGCAAGCTCCGCCTCCCGGGTTCACGCCATTCTCCTGCCTCAGCCTCCCGAGTAGCTGGGACTA
CAGGCGCCCGCCAACACGCCCGGCTAATTTTTTTTTTGTATTTTTAGTAGAGACGGGGTTTCACCGTGTTAGCCAGGATGGTCTCGATCTCCTGACCTCGTGATCCGCCCGCCTCGGCCTCCC
AAAGTGCTGGGATTACAGGCGTGAGCCACCGCGCCCGGCC

### HG03492 chr8:39822492

- ➢ Using the built-in BLAT tool in IGV to check where the inserted sequence aligned.
- ➢ The annotation at the aligned location indicated it as *AluY*.

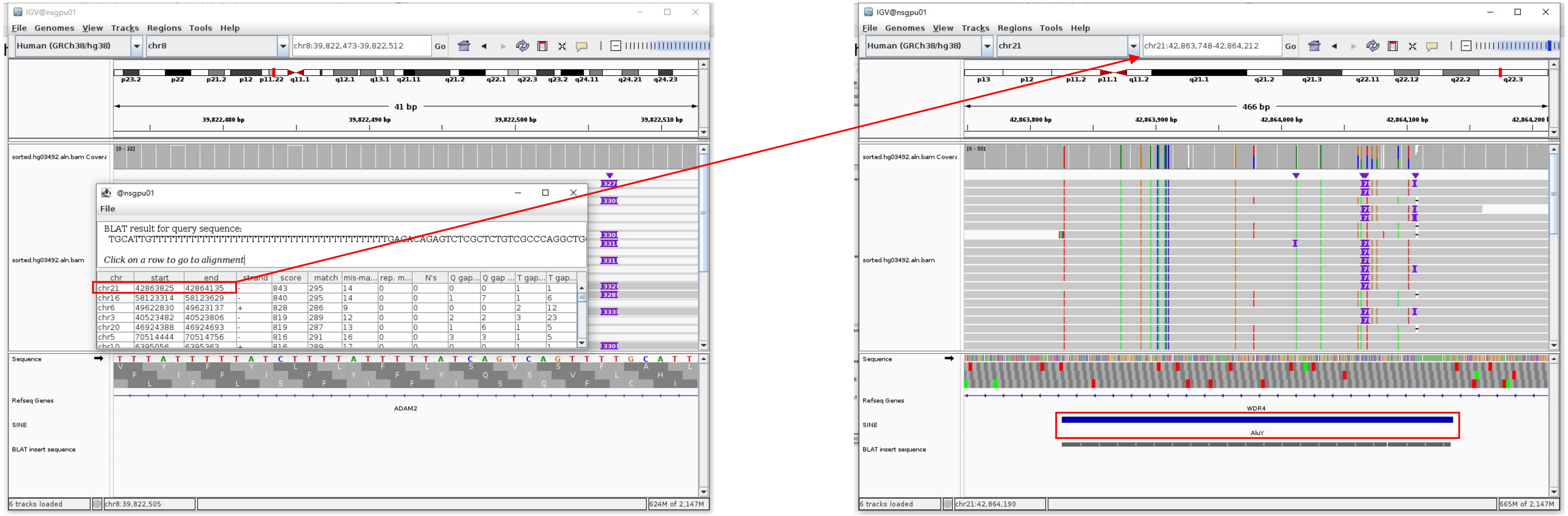


Insertion sequence (332bp)

- ➢ 10bp TSD

TGCATTGTTT

- ➢ Insertion:49-332 aligns to *Alu*Yc1:282-0, 98.94% identity

TGAGACAGAGTCTCGCTCTGTCGCCCAGGCTGGAGTGCAGTGGCGCGATCTCGGCTCACTGCAAGCTCCGCCTCCTGGGTTCACGCCATTCTCCTGCCTCAGCCTCCCGAGTAGCTGGGACTA
CAGGCGCCCGCCACCACGCCCGGCTAATTTTTTGTATTTTTAGTAGAGACGGGGTTTCACCGTGTTAGCCAGGATGGTCTCGATCTCCTGACCTCGTGATCCGCCCGCCTCGGCCTCCCAAA
GTGCTGGGATTACAGGCGTGAGCCACCGCGCCCGGCC

### HG03492 chr11:134789637

➢ Using the built-in BLAT tool in IGV to check where the inserted sequence aligned.

➢ The annotation at the aligned location indicated it as *AluY*.

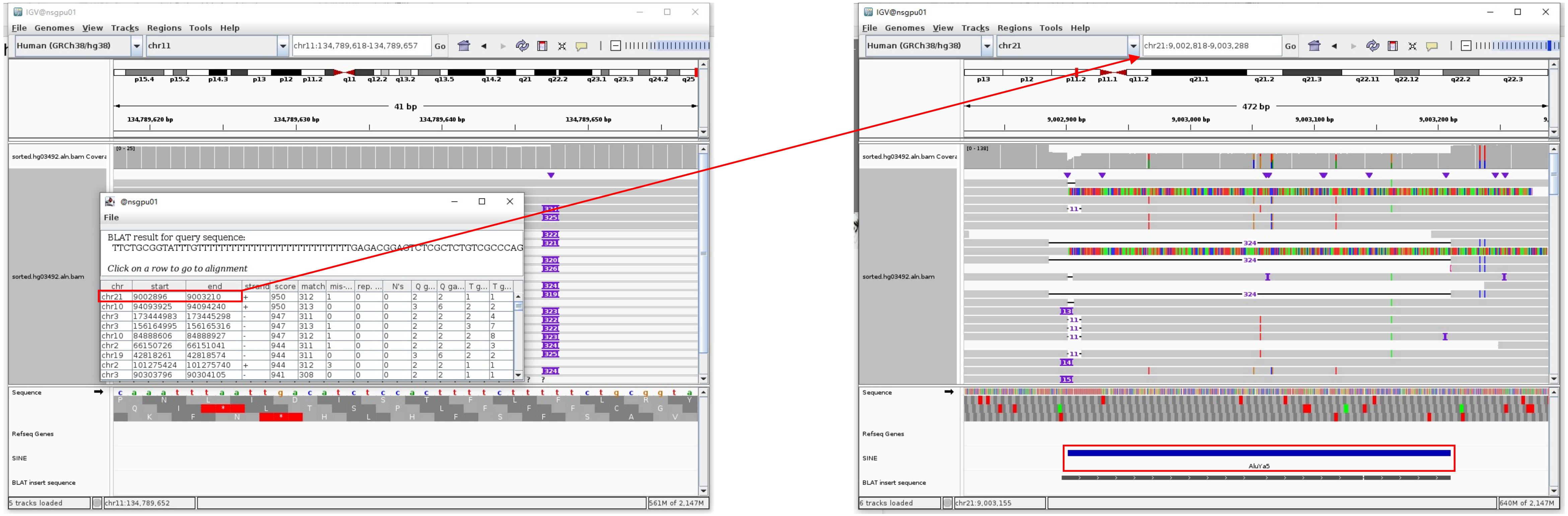


Insertion sequence (324bp)

➢ 14bp TSD

TTCTGCGGTATTTG

➢ Insertion:41-324 aligns to *Alu*Ya5:282-1, 100% identity

TGAGACGGAGTCTCGCTCTGTCGCCCAGGCTGGAGTGCAGTGGCGGGATCTCGGCTCACTGCAAGCTCCGCCTCCCGGGTTCACGCCATTCTCCTGCCTCAGCCTCCCAAGTAGCTGGGACT
ACAGGCGCCCGCCACTACGCCCGGCTAATTTTTTGTATTTTTAGTAGAGACGGGGTTTCACCGTTTTAGCCGGGATGGTCTCGATTCTCCGACCTCGTGATCCGCCCGCCTCGGCCTCCCAAA
GTGCTGGGATTACAGGCGTGAGCCACCGCGCCCGGCC

### HG03492 chr13:89749568

- Using the built-in BLAT tool in IGV to check where the inserted sequence aligned.

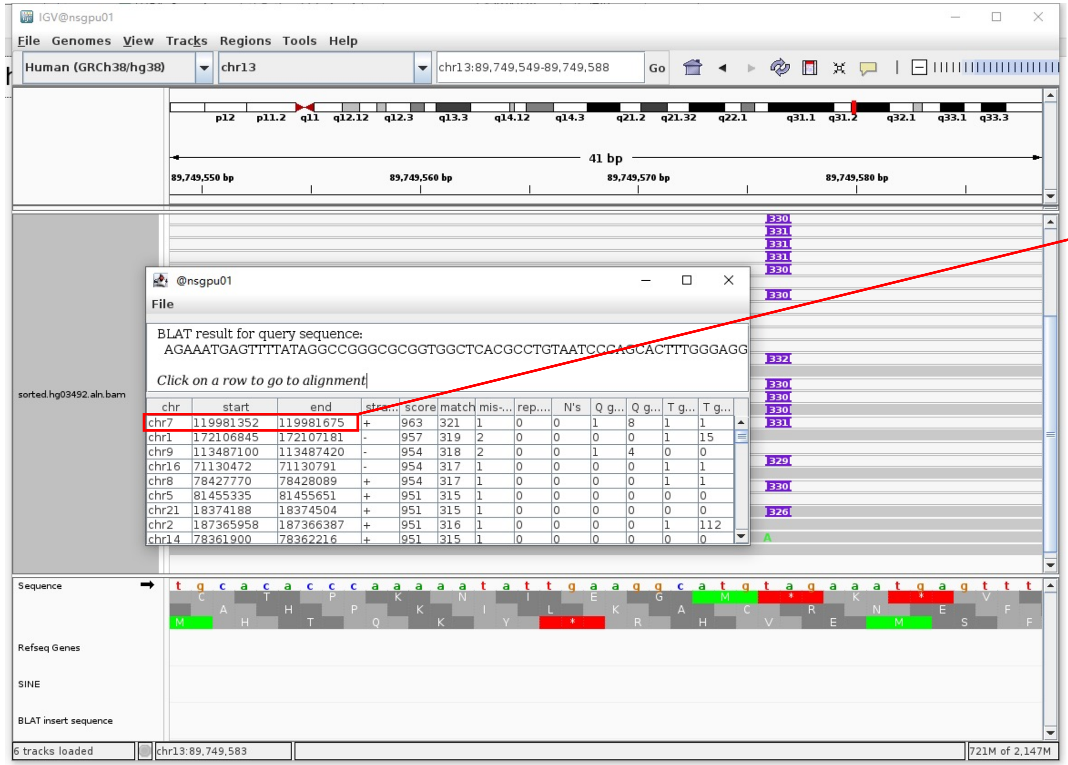


- The annotation at the aligned location indicated it as *Alu*Y.

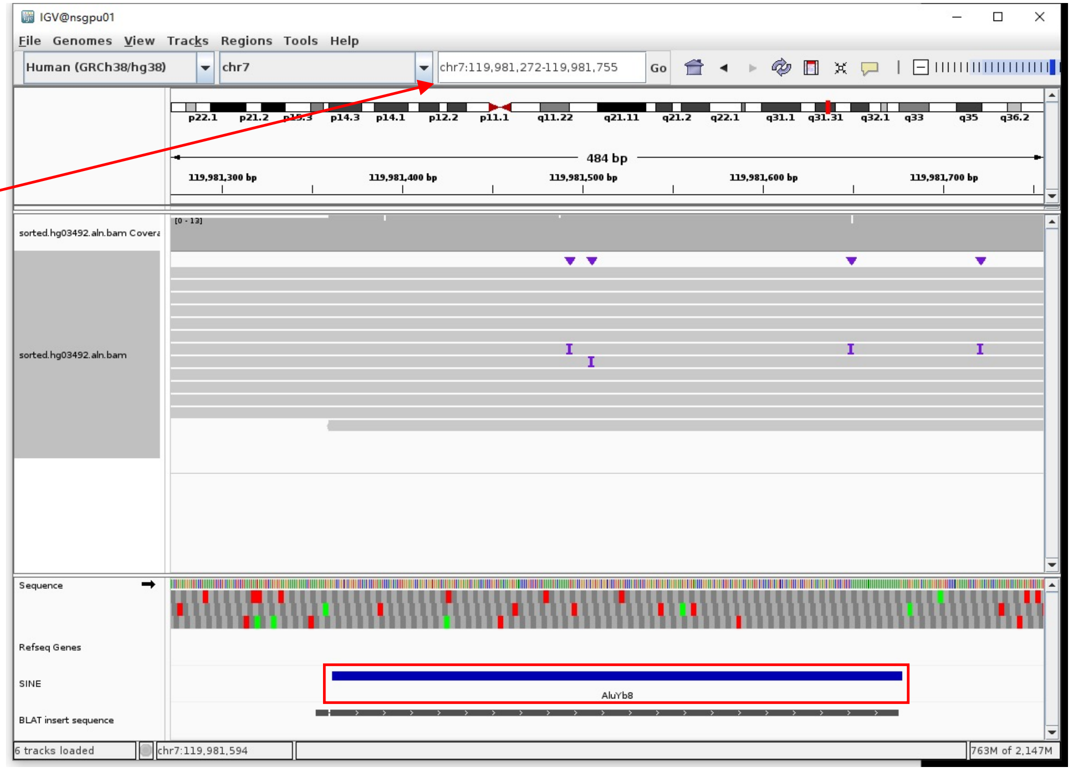


Insertion sequence (330bp)

- 16bp TSD

AGAAATGAGTTTTATA

- Insertion:16-304 aligns to *Alu*Yb8:1-289, 99.65% identity

GGCCGGGCGCGGTGGCTCACGCCTGTAATCCCAGCACTTTGGGAGGCCGAGGCGGGTGGATCATGAGGTCAGGAGATCGAGACCATCCTGGCTAACAAGGTGAAACCCCGTCTCTACTAAAAA
TACAAAAAATTAGCCGGGCGCGGTGGCGGGTGCTGTAGTCCCAGCTACTCGGGAGGCTGAGGCAGGAGAATGGCGTGAACCCGGGAAGCGGAGCTTGCAGTGAGCCGAGATTGCGCCACTGCA
GTCCGCAGTCCGGCCTGGGCGACAGAGCGAGACTCCGTCTCA………poly(A)

### HG03492 chr14:93364571

- The same genomic coordinate as in HG00438 (page 7)

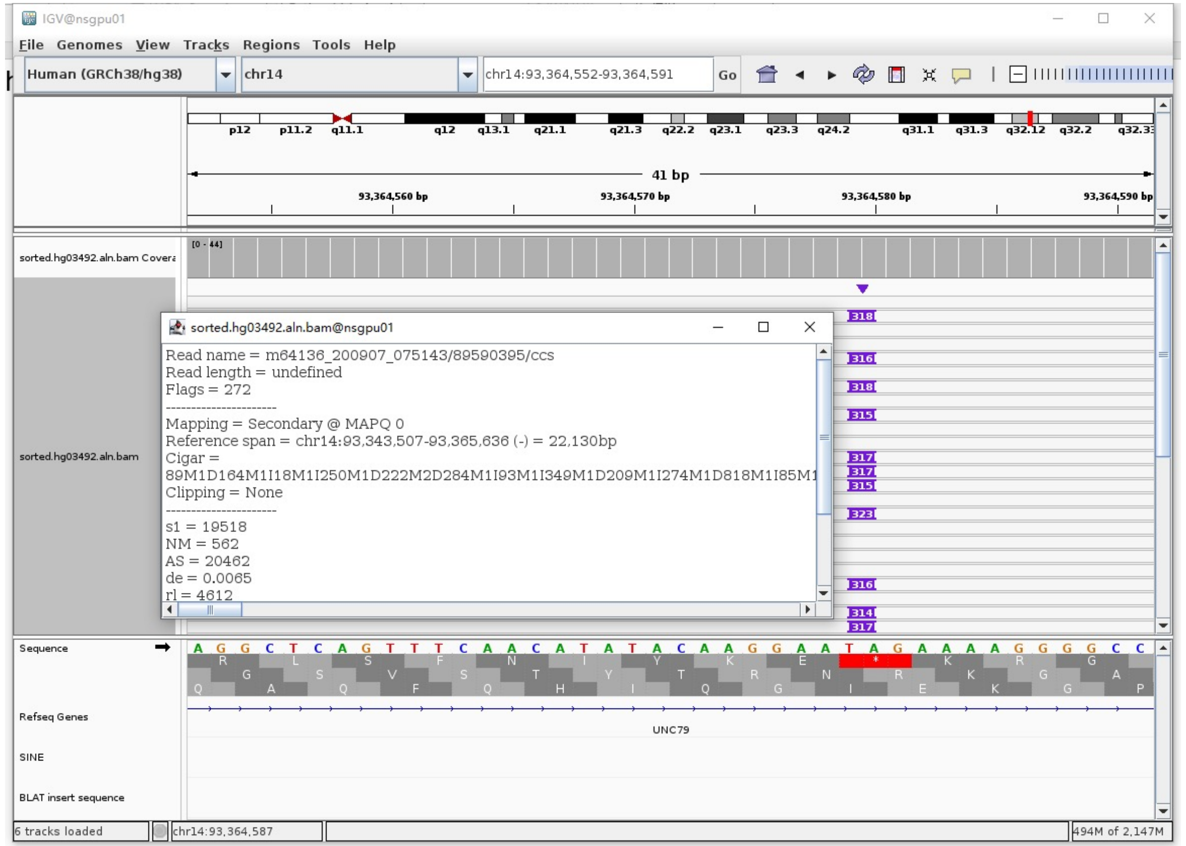

### HG03492 chr17:41020755

➢ Mapping quality is 0, indicating that this region may have alignment issues.

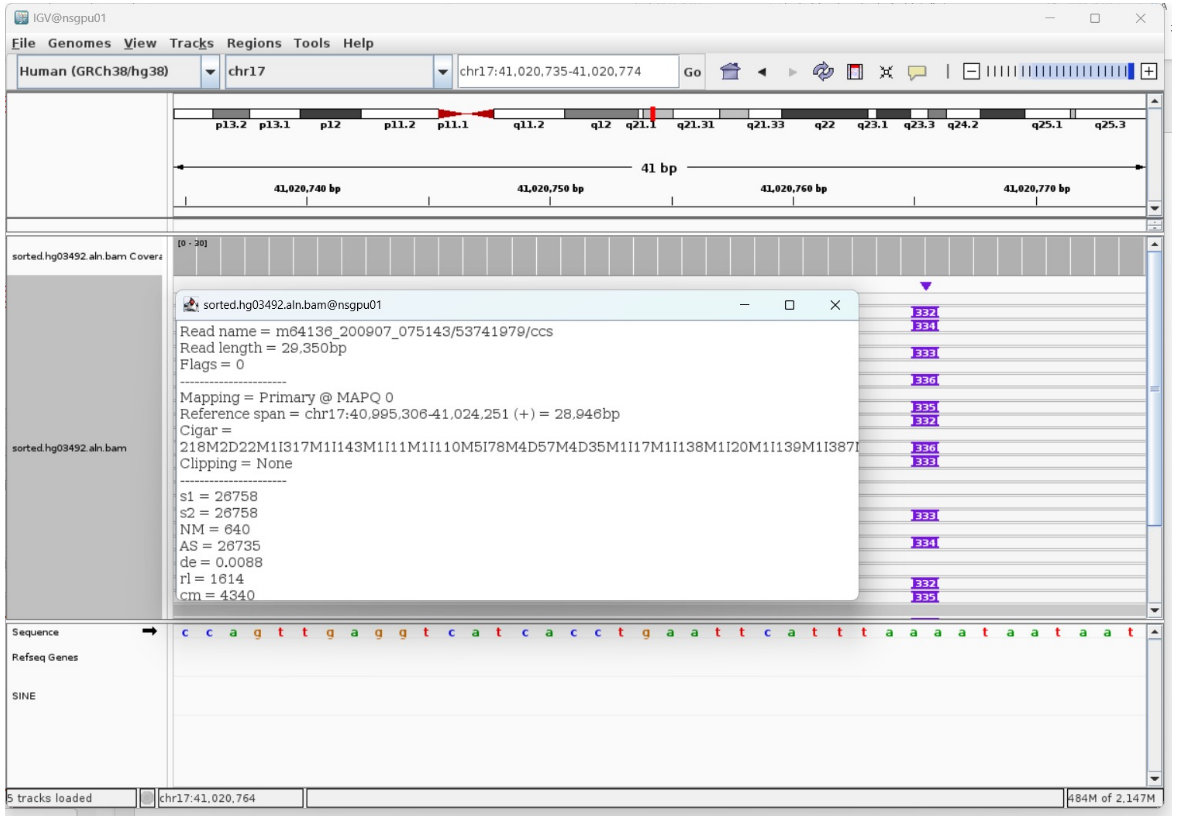


➢ Fetch one read (m64136_200907_075143/53741979/ccs, 29309 bp) and BLAT it to the hg38 genome.

The best alignment is to chr17_JH159146v1_alt (an alternative sequence of chr17) and chr17.

| ACTIONS | QUERY | SCORE | START | END | QSIZE | IDENTITY | CHROM | STRAND | START | END | SPAN |
|---|---|---|---|---|---|---|---|---|---|---|---|
| browser new tab details | YourSeq | 28618 | 1 | 29309 | 29309 | 99.5% | chr17_JH159146v1_alt | + | 144701 | 173646 | 28946 |
| browser new tab details | YourSeq | 28618 | 1 | 29309 | 29309 | 99.5% | chr17 | + | 40995306 | 41024251 | 28946 |

● Read:1-25501 aligns to chr17_JH159146v1_alt

| ACTIONS | QUERY | SCORE | START | END | QSIZE | IDENTITY | CHROM | STRAND | START | END | SPAN |
|---|---|---|---|---|---|---|---|---|---|---|---|
| browser new tab details | YourSeq | 25163 | 1 | 25500 | 25500 | 99.5% | chr17_JH159146v1_alt | + | 144702 | 170173 | 25472 |
| browser new tab details | YourSeq | 25163 | 1 | 25500 | 25500 | 99.5% | chr17 | + | 40995307 | 41020778 | 25472 |

● Read:25502-25785 aligns to *Alu*Ya5:1-282, 99.65% identity

GGCCGGGCGCGGTGGCTCACGCCTGTAATCCCAGCACTTTGGGAGGCCGAGGCGGGCGGATCACGAGGTCAAGA
GATCGAGACCATCCCGGCTAAAACGGTGAAACCCCGTCTCTACTAAAAATACAAAAAATTAGCCGGGCGTAGT
GGCGGGCGCCTGTAGTCCCAGCTACTTGGGAGGCTGAGGCAGGAGAATGGCGTGAACCCGGGAGGCGGAGCTTG
CAGTGAGCCGAGATCCCGCCACTGCACTCCAGCCTGGGCGACAGAGCGAGACTCCGTCTCA

14bp TSD

AATAATAATACATA

● Read:25786-29309 also aligns to chr17_JH159146v1_alt

| ACTIONS | QUERY | SCORE | START | END | QSIZE | IDENTITY | CHROM | STRAND | START | END | SPAN |
|---|---|---|---|---|---|---|---|---|---|---|---|
| browser new tab details | YourSeq | 3468 | 36 | 3523 | 3523 | 99.7% | chr17_JH159146v1_alt | + | 170161 | 173646 | 3486 |
| browser new tab details | YourSeq | 3468 | 36 | 3523 | 3523 | 99.7% | chr17 | + | 41020766 | 41024251 | 3486 |

➢ It's an *Alu*Ya5 insertion.

### HG03492 chr18:29602115

➢ Using the built-in BLAT tool in IGV to check where the inserted sequence aligned.

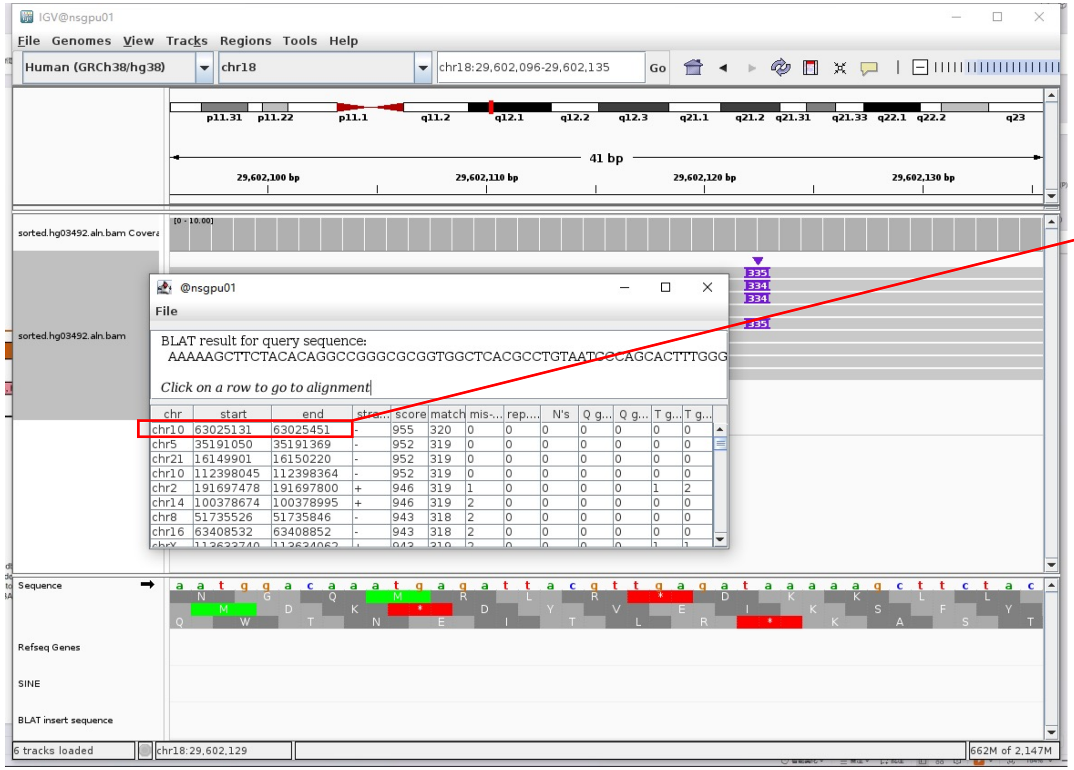


➢ The annotation at the aligned location indicated it as *AluY*.

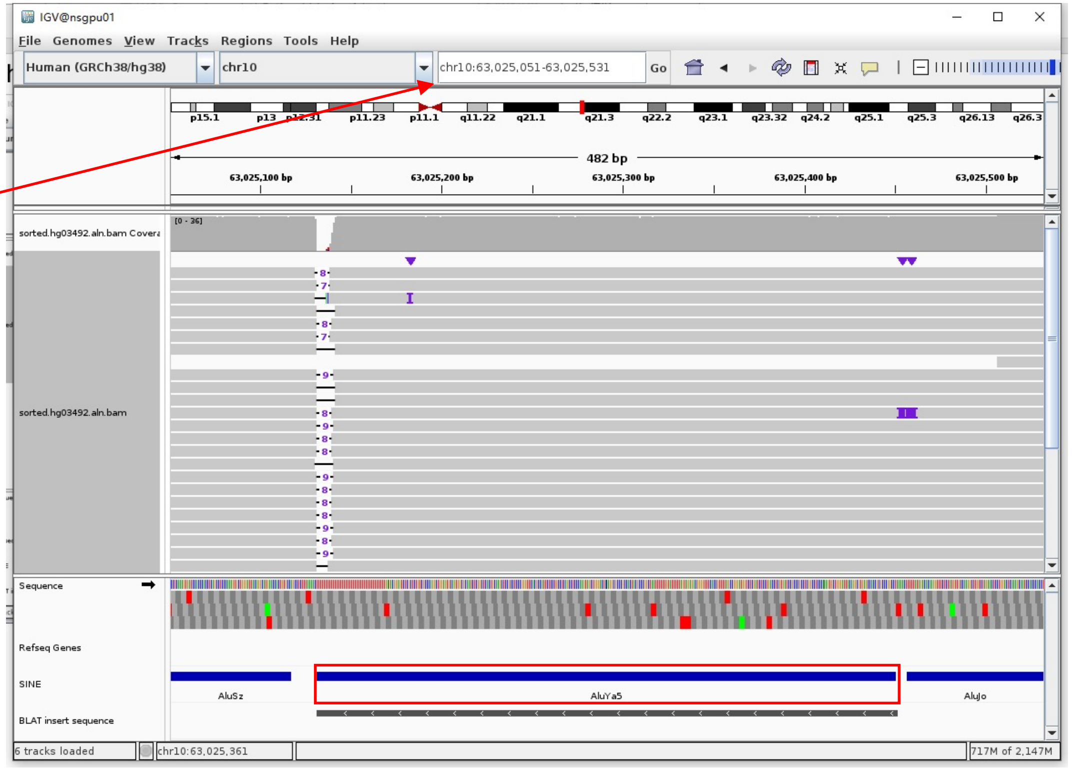


Insertion sequence (334bp)

➢ 17bp TSD

AAAAAGCTTCTACACAG

➢ Insertion:16-298 aligns to *Alu*Ya5:1-282, 99.29% identity

GGCCGGGCGCGGTGGCTCACGCCTGTAATCCCAGCACTTTGGGAGGCCGAGGCGGGCGGATCACGAGGTCAGGAGATCGAGACCATCCCGGCTAAAACGGTGAAACCCCGTCTCTACTAAAAA
TACAAAAAATTAGCCGGGCGTAGTGGCGGGCGCCTGTAGTCCCAGCTACTTGGGAGGCTGAGGCAGGAGAATGGCGTGAACCCGGGAGGCGGAGCTTGCAGTGAGCCGAGATTGCGCCACTGC
ACTCCAGCCTGGGCGACAGAGCGAGACTCCGTCTCA.........poly(A)

### HG03492 chr19:52384796

- Using the built-in BLAT tool in IGV to check where the inserted sequence aligned.
- The annotation at the aligned location indicated it as *AluY*.

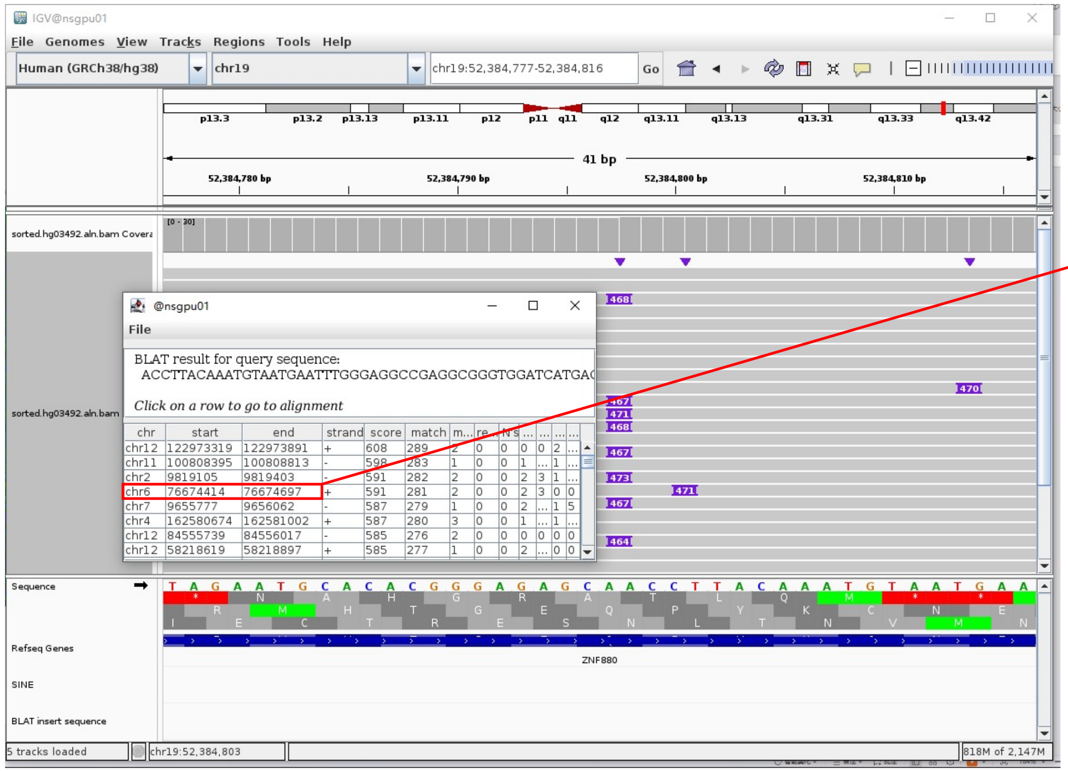


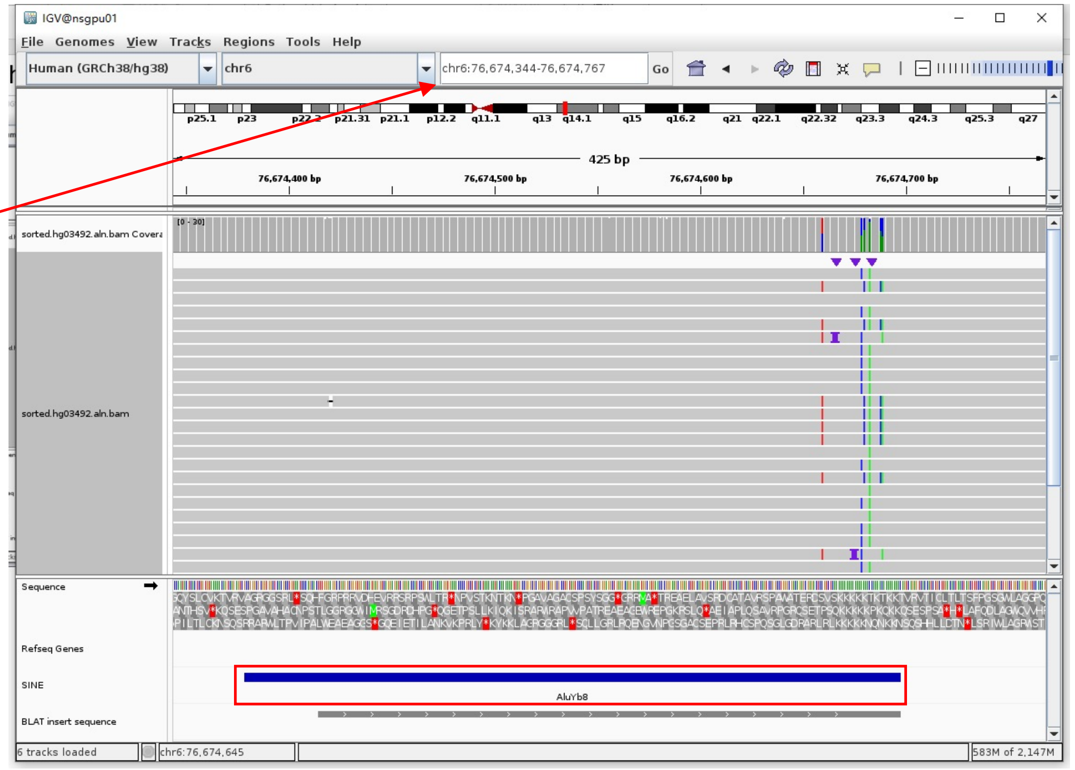


Insertion sequence (464bp)

- 20bp TSD

ACCTTACAAATGTAATGAAT

- Insertion:2-271 aligns to *Alu*Yb8:15-289, 97.77% identity

CTTACAAATGTAATGAATTTGGGAGGCCGAGGCGGGTGGATCATGAGGTCAGGAGATCGAGACCATCCTGGCTAACAAGGTGAAACCCCATCTCTACTAAAAATACAAAAAATTAGCCGGGCG
CGGTGGCGGGCGCCTGTAGTCCCAGCTACTCGGGAGGCTGAGGCAGGAGAATGGCGTGAACCCGGGAAGCGGAGCTTGCAGTGAGCCGAGATTGCGCCACTGCAGTCCGCAGTCCGGCCTGGG
CGACAGAGCGAGACTCCGTCTCA.........poly(A)

### HG03516 chr5:43541665

- ➢ Using the built-in BLAT tool in IGV to check where the inserted sequence aligned.

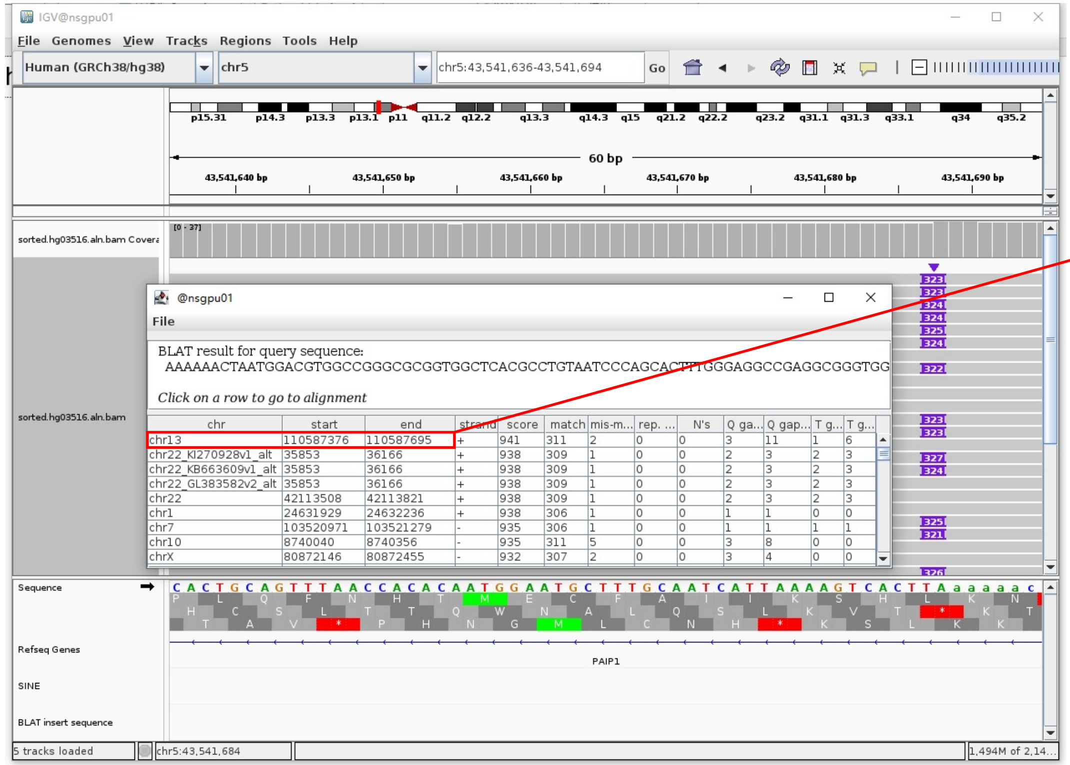


- ➢ The annotation at the aligned location indicated it as *AluY*.

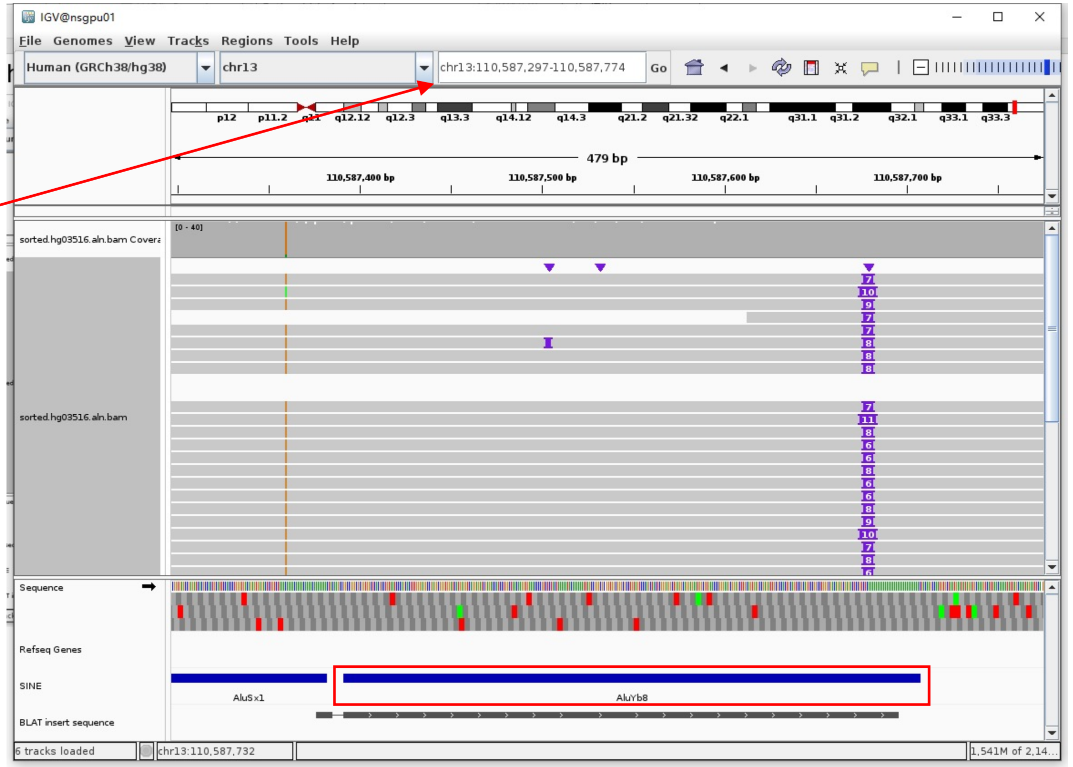


Insertion sequence (322bp)

- ➢ 16bp TSD

AAAAAACTAATGGACG

- ➢ Insertion:17-309 aligns to *Alu*Yb8:1-289, 99.31% identity

GGCCGGGCGCGGTGGCTCACGCCTGTAATCCCAGCACTTTGGGAGGCCGAGGCGGGTGGATCATGAGGTCAGGAGATCGAGACCATCCTGGCTAACAAGGTGAAACCCCGTCTCCACTAAAAATACAAAAAAAATTAGCCGGGCGCGGTGGCGGGCGCCTGTAGTCCCAGCTACTCGGGAGGCTGAGGCAGGAGAATGGCGTGAACCCGGGAAGCGGAGCTTGCAGTGAGCCGAGATTGCGCCACTGCAGTCCGCAGTCCCGCCTGGGCGACAGAGCGAGACTCCGTCTCA………poly(A)

### HG03516 chr5:69513668

- Mapping quality is 0, indicating that this region may have alignment issues.

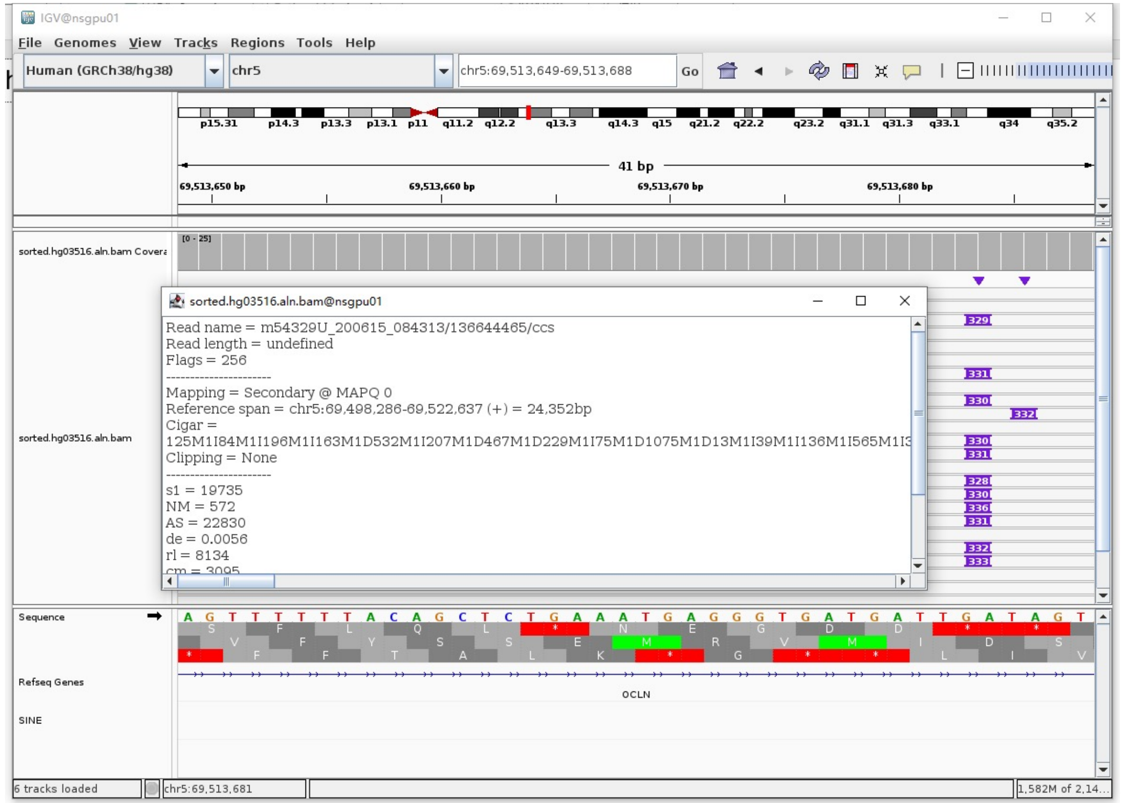


- Fetch one read (m54329U_200615_084313/87623274/ccs, 14628 bp) and BLAT it to the hg38 genome.

  The best alignment is to chr5_MU273354v1_fix, chr5_GL339449v2_alt (an alternative sequence of chr5) and chr5.

| ACTIONS | | | QUERY | SCORE | START | END | QSIZE | IDENTITY | CHROM | STRAND | START | END | SPAN |
|---|---|---|---|---|---|---|---|---|---|---|---|---|---|
| browser | new tab | details | YourSeq | 14281 | 1 | 14628 | 14628 | 99.9% | chr5_MU273354v1_fix | + | 170517 | 184816 | 14300 |
| browser | new tab | details | YourSeq | 14186 | 1 | 14628 | 14628 | 99.9% | chr5_GL339449v2_alt | + | 286852 | 301052 | 14201 |
| browser | new tab | details | YourSeq | 14168 | 1 | 14628 | 14628 | 99.8% | chr5 | + | 69503955 | 69518154 | 14200 |

- Read:1-9771 aligns to chr5_GL339449v2_alt

| ACTIONS | | | QUERY | SCORE | START | END | QSIZE | IDENTITY | CHROM | STRAND | START | END | SPAN |
|---|---|---|---|---|---|---|---|---|---|---|---|---|---|
| browser | new tab | details | YourSeq | 9737 | 1 | 9740 | 9771 | 100.0% | chr5_GL339449v2_alt | + | 286852 | 296592 | 9741 |
| browser | new tab | details | YourSeq | 9736 | 1 | 9740 | 9771 | 100.0% | chr5_MU273354v1_fix | + | 170517 | 180258 | 9742 |
| browser | new tab | details | YourSeq | 9722 | 1 | 9740 | 9771 | 99.9% | chr5 | + | 69503955 | 69513692 | 9738 |

- Read:9772-10061 aligns to *Alu*Yb8:289-1, 99.31% identity

  TGAGACGGAGTCTCGCTCTGTCGCCCAGGCCGGACTGCGGACCGCAGTGGCGCAATCTCGGCTCACTGCAAGCT
  CCGCCTCCCGGGTTCACGCCATTCTCCTGCCTCAGCCTCCCGAGTAGCTGGGACTACAGGCGCCCGCCACCGCG
  CCCGGCTAATTTTTTGTATTTTTAGTAGAGACGGGGTTTCACCTTGTTAGCCAGGATGGTCTCGATCTCCTGAC
  CTCATGATCCACCCGCCTCGGCCTCCCAAAGTGCTGGGATTACAGGCGTGAGCCACCGCGCCCGGCC

  9bp TSD

  ATAGTTCCT

- Read:10062-14628 aligns to chr5_MU273354v1_fix, chr5_GL339449v2_alt

| ACTIONS | | | QUERY | SCORE | START | END | QSIZE | IDENTITY | CHROM | STRAND | START | END | SPAN |
|---|---|---|---|---|---|---|---|---|---|---|---|---|---|
| browser | new tab | details | YourSeq | 4555 | 1 | 4566 | 4566 | 100.0% | chr5_MU273354v1_fix | + | 180250 | 184816 | 4567 |
| browser | new tab | details | YourSeq | 4459 | 1 | 4566 | 4566 | 99.7% | chr5_GL339449v2_alt | + | 296584 | 301052 | 4469 |
| browser | new tab | details | YourSeq | 4456 | 1 | 4566 | 4566 | 99.6% | chr5 | + | 69513684 | 69518154 | 4471 |

- It's an *Alu*Yb8 insertion.

### HG03516 chr6:30030695

- The same genomic coordinate as in HG03492 (page 47)

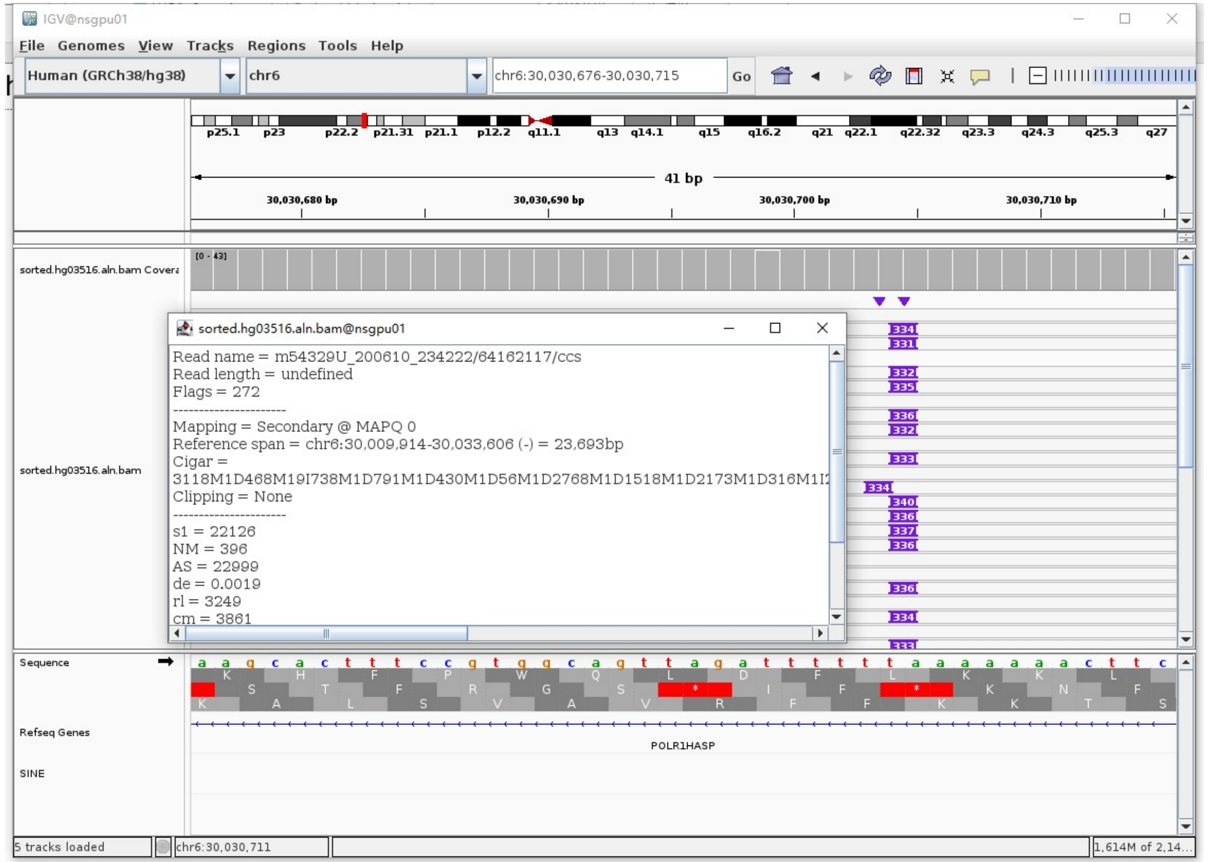

### HG03516 chr6:32740195

➢ Mapping quality is 0, indicating that this region may have alignment issues.

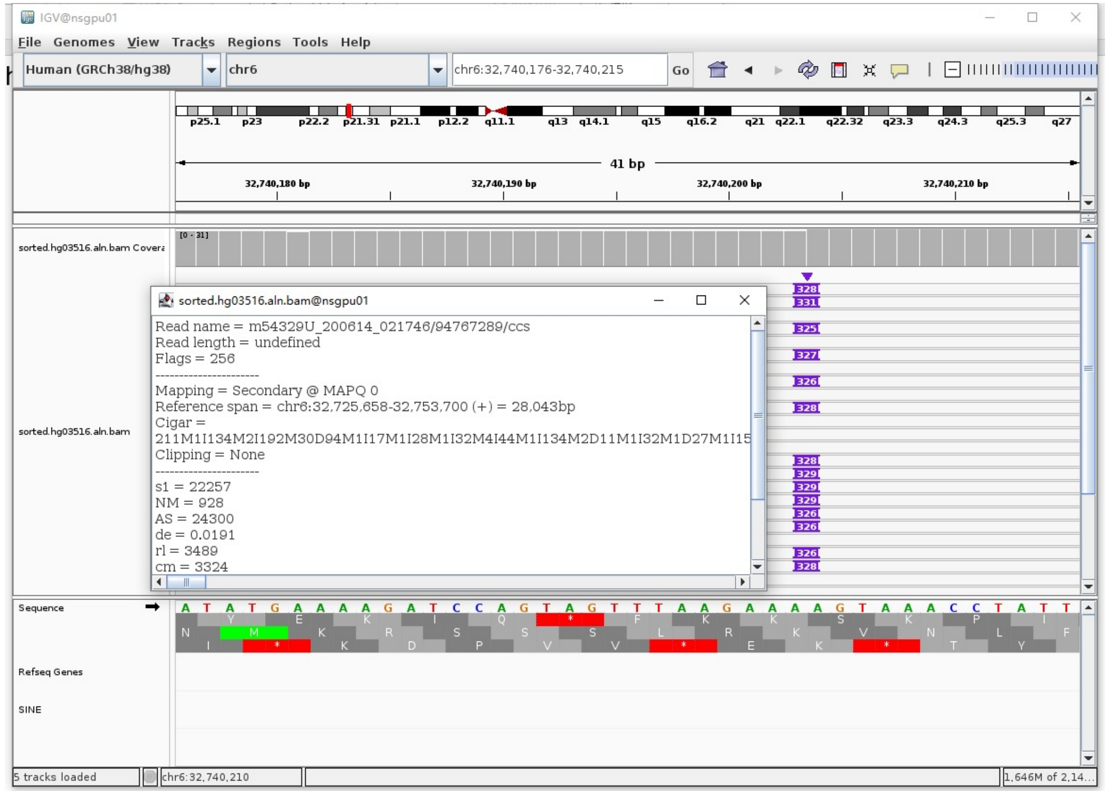


➢ Fetch one read (m54329U_200612_200443/124324370/ccs, 18723 bp) and BLAT it to the hg38 genome.

The best alignment is to chr6_GL000251v2_alt (an alternative sequence of chr6), which contains an *Alu*Ya5 element.

| ACTIONS | | | QUERY | SCORE | START | END | QSIZE | IDENTITY | CHROM | STRAND | START | END | SPAN |
|---|---|---|---|---|---|---|---|---|---|---|---|---|---|
| browser | new tab | details | YourSeq | 18664 | 1 | 18723 | 18723 | 99.9% | chr6_GL000251v2_alt | - | 4145388 | 4164106 | 18719 |
| browser | new tab | details | YourSeq | 18662 | 1 | 18723 | 18723 | 99.9% | chr6_GL000250v2_alt | - | 4037954 | 4056672 | 18719 |
| browser | new tab | details | YourSeq | 18078 | 1 | 18723 | 18723 | 99.1% | chr6_GL000256v2_alt | - | 4132280 | 4150678 | 18399 |
| browser | new tab | details | YourSeq | 18076 | 1 | 18723 | 18723 | 99.1% | chr6 | - | 32732133 | 32750532 | 18400 |

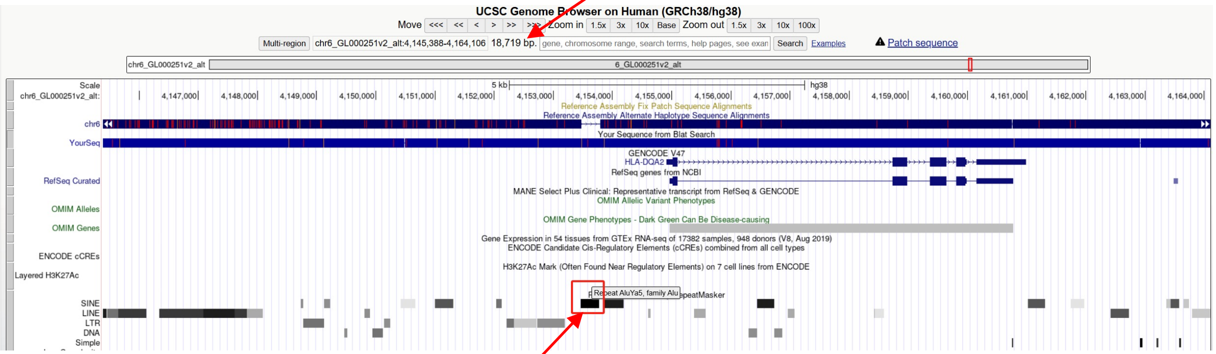


chr6_GL000251v2_alt:4153465-4153777 (*Alu*Ya5 marked by RepeatMasker)

➢ Read:10326-10608 aligns to *Alu*Ya5:1-282, 98.94% identity

GGCCGGGCGCGGTGGCTCACGCCTGTAATCCCAGCACTTTGGGAGGCCGAGGCGGGCGGATCACGAGGTCAGGAG
ATCGAGACCATCCTGGCTAACAAGGTGAAACCCCGTCTCTACTAAAAATACAAAAAATTAGCCGGGCGTAGTGGC
GGGCGCCTGTAGTCCCAGCTACTTGGGAGGCTGAGGCAGGAGAATGGCGTGAACCCGGGAGGCGGAGCTTGCAGT
GAGCCGAGATCCCGCCACTGCACTCCAGCCTGGGCGACAGAGCGAGACTCCGTCTCA

➢ It's an *Alu*Ya5 in the chr6_GL000251v2_alt.

### HG03516 chr7:142976545

- Mapping quality is 0, indicating that this region may have alignment issues.

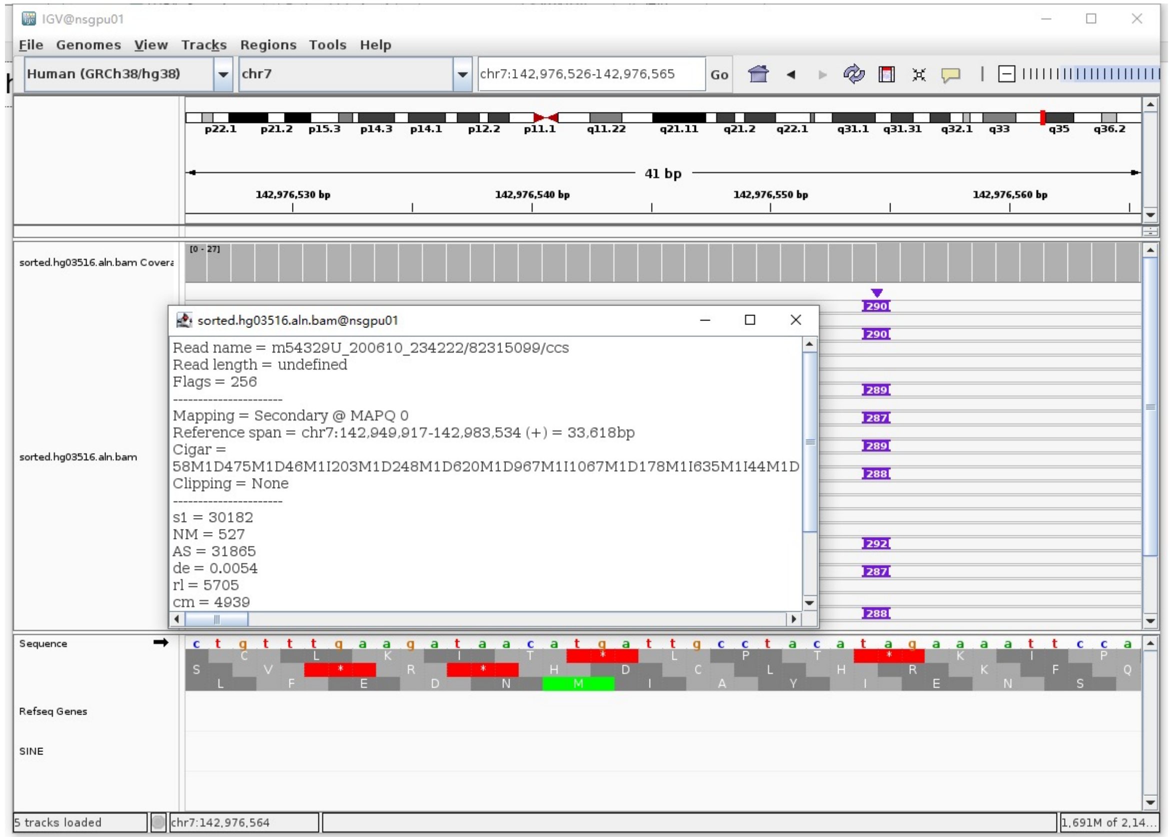


- Fetch one read (m54329U_200612_200443/54199051/ccs, 16162 bp) and BLAT it to the hg38 genome.

  The best alignment is to chr7_KI270803v1_alt (an alternative sequence of chr7) and chr7.

| ACTIONS | QUERY | SCORE | START | END | QSIZE | IDENTITY | CHROM | STRAND | START | END | SPAN |
|---|---|---|---|---|---|---|---|---|---|---|---|
| browser new tab details | YourSeq | 15727 | 1 | 16162 | 16162 | 99.5% | chr7_KI270803v1_alt | + | 986872 | 1002792 | 15921 |
| browser new tab details | YourSeq | 15727 | 1 | 16162 | 16162 | 99.5% | chr7 | + | 142963805 | 142979725 | 15921 |

- Read:1-12714 aligns to chr7_KI270803v1_alt

| ACTIONS | QUERY | SCORE | START | END | QSIZE | IDENTITY | CHROM | STRAND | START | END | SPAN |
|---|---|---|---|---|---|---|---|---|---|---|---|
| browser new tab details | YourSeq | 12587 | 1 | 12714 | 12714 | 99.6% | chr7_KI270803v1_alt | + | 986872 | 999634 | 12763 |
| browser new tab details | YourSeq | 12587 | 1 | 12714 | 12714 | 99.6% | chr7 | + | 142963805 | 142976567 | 12763 |

- Read:12715-12970 aligns to *Alu*Yc1:27-282, 99.61% identity

  ATCCCAGCACTTTGGGAGGCCGAGGCGGGCGGATCACGAGGTCAGGAGATCGAGACCATCCTGGCTAACACGGT
  GAAACCCCGTCTCTACTAAAAATACAAAAAATTAGCCGGGCGTGGTAGCGGGCGCCTGTAGTCCCAGCTACTCG
  GGAGGCTGAGGCAGGAGAATGGCGTGAACCTGGGAGGCGGAGCTTGCAGTGAGCCGAGATCGCGCCACTGCACT
  CCAGCCTGGGCGACAGAGCGAGACTCCGTCTCA

  15bp TSD

  AGAAAATTCCAAAGA

- Read:12971-16162 also aligns to chr7_KI270803v1_alt

| ACTIONS | QUERY | SCORE | START | END | QSIZE | IDENTITY | CHROM | STRAND | START | END | SPAN |
|---|---|---|---|---|---|---|---|---|---|---|---|
| browser new tab details | YourSeq | 3171 | 1 | 3191 | 3191 | 99.8% | chr7_KI270803v1_alt | + | 999381 | 1002792 | 3412 |
| browser new tab details | YourSeq | 3171 | 1 | 3191 | 3191 | 99.8% | chr7 | + | 142976314 | 142979725 | 3412 |

- It's an *Alu*Yc1 insertion.

### HG03516 chr8:39842932

➢ Mapping quality is 0, indicating that this region may have alignment issues.

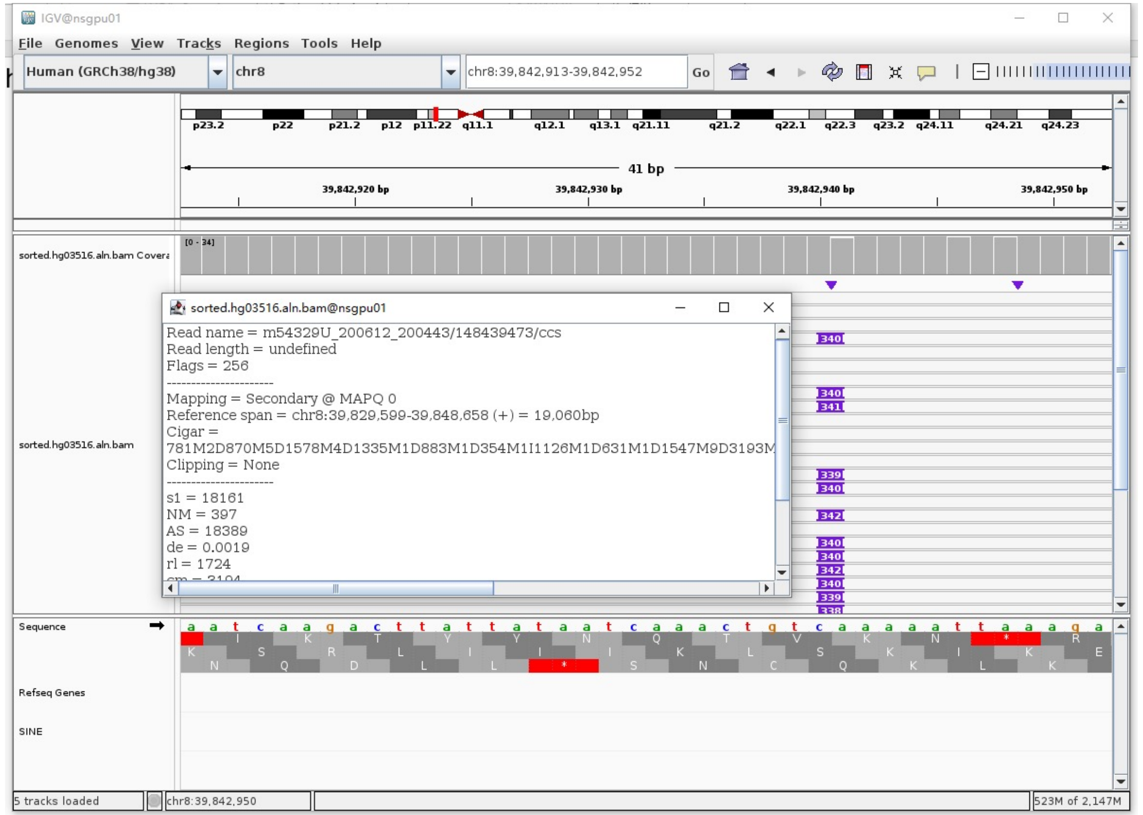


➢ Fetch one read (m54329U_200614_021746/13304069/ccs, 20067 bp) and BLAT it to the hg38 genome.

The best alignment is to chr8_KI270822v1_alt (an alternative sequence of chr8) and chr8.

| ACTIONS | | | QUERY | SCORE | START | END | QSIZE | IDENTITY | CHROM | STRAND | START | END | SPAN |
|---|---|---|---|---|---|---|---|---|---|---|---|---|---|
| browser | new tab | details | YourSeq | 19670 | 1 | 20067 | 20067 | 99.8% | chr8_KI270822v1_alt | - | 578125 | 597873 | 19749 |
| browser | new tab | details | YourSeq | 19670 | 1 | 20067 | 20067 | 99.8% | chr8 | - | 39827489 | 39847237 | 19749 |

- Read:1-4321 aligns to chr8_KI270822v1_alt

| ACTIONS | | | QUERY | SCORE | START | END | QSIZE | IDENTITY | CHROM | STRAND | START | END | SPAN |
|---|---|---|---|---|---|---|---|---|---|---|---|---|---|
| browser | new tab | details | YourSeq | 4284 | 1 | 4292 | 4320 | 100.0% | chr8_KI270822v1_alt | - | 593577 | 597872 | 4296 |
| browser | new tab | details | YourSeq | 4284 | 1 | 4292 | 4320 | 100.0% | chr8 | - | 39842941 | 39847236 | 4296 |

- Read:4322-4617 aligns to *Alu*Yc1:282-1, 93.55% identity

TTGAGACGGAGTCTCGCTGTCGCCCAGGCTGGAGTGCAGTGGCGCAATCTCGGCTCACTGCAGGCTCCGCCCCC
TGGGGTTCGCGCCATTCTCCTGCCTCAGCCTCCGGAGTAGCTGGGACTACAGACGCCCGCCACCTCGCCCGGCT
AATTTTTTTTTTTTTTTTTTTTTGTATTTTTAGTAGAGACGGGGTTTCACCGTGTTAGCCAGGATGGTCTCGAT
CTCCTGACCTCGTGATCCGCCCGCCTCGGCCTCCCAAAGTGCTGGGATTACAGGCGTGAGCCACCGCGCCCGG

16bp TSD

CTTCTCTTTAATTTTT

- Read:4618-20067 also aligns to chr8_KI270822v1_alt

| ACTIONS | | | QUERY | SCORE | START | END | QSIZE | IDENTITY | CHROM | STRAND | START | END | SPAN |
|---|---|---|---|---|---|---|---|---|---|---|---|---|---|
| browser | new tab | details | YourSeq | 15401 | 1 | 15449 | 15449 | 99.9% | chr8_KI270822v1_alt | - | 578125 | 593591 | 15467 |
| browser | new tab | details | YourSeq | 15401 | 1 | 15449 | 15449 | 99.9% | chr8 | - | 39827489 | 39842955 | 15467 |

➢ It's an *Alu*Yc1 insertion.

### HG03516 chr10:517814

- ➢ Using the built-in BLAT tool in IGV to check where the inserted sequence aligned.

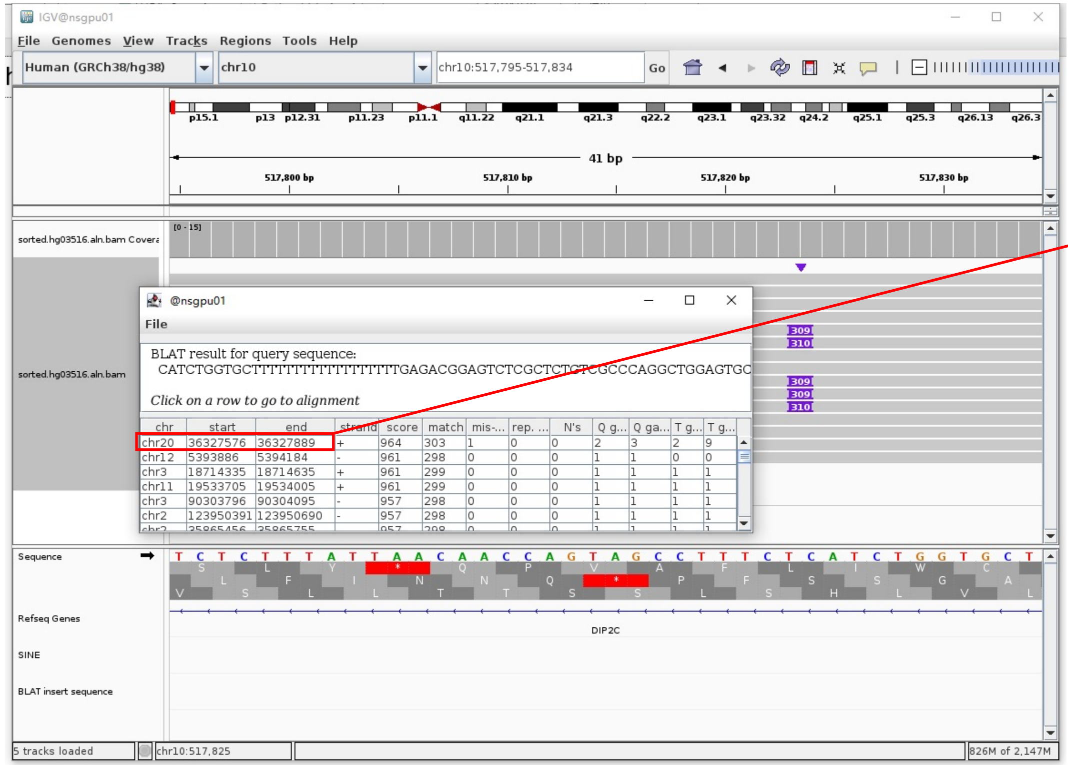


- ➢ The annotation at the aligned location indicated it as *AluY*.

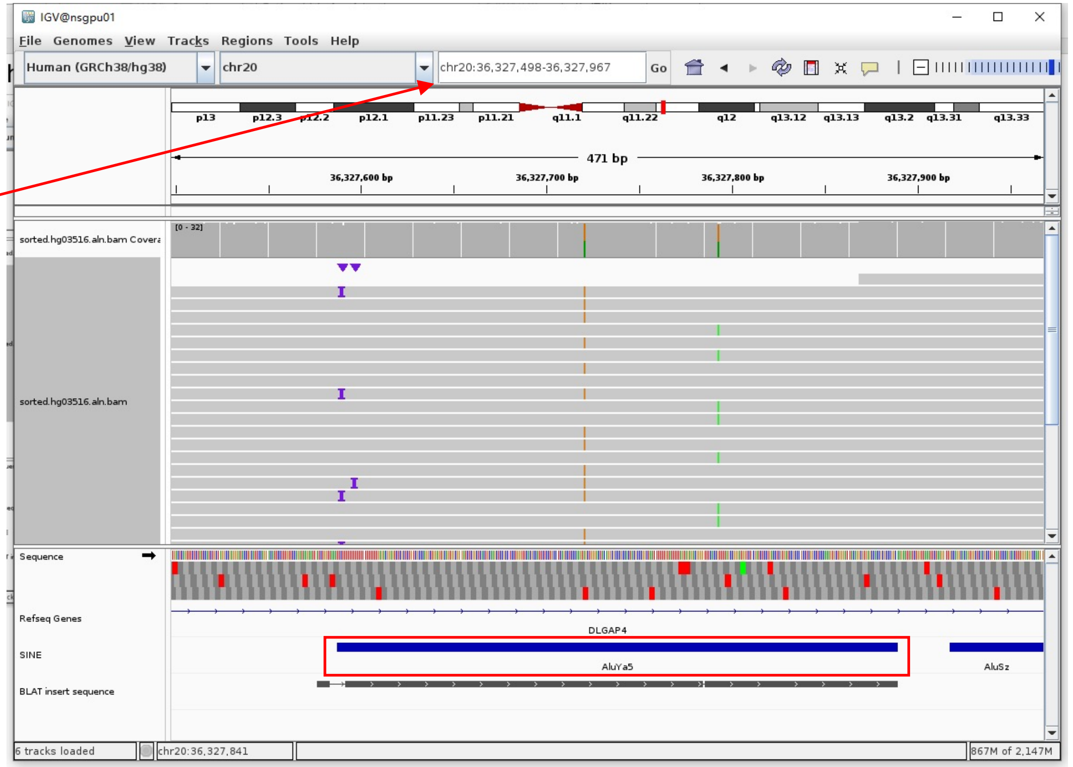


Insertion sequence (309bp)

- ➢ 15bp TSD

CATCTGGTGCTTTTT

- ➢ Insertion:27-309 aligns to *Alu*Ya5:282-1, 100% identity

TGAGACGGAGTCTCGCTCTGTCGCCCAGGCTGGAGTGCAGTGGCGGGATCTCGGCTCACTGCAAGCTCCGCCTCCCGGGTTCACGCCATTCTCCTGCCTCAGCCTCCCAAGTAGCTGGGACTA
CAGGCGCCCGCCACTACGCCCGGCTAATTTTTTGTATTTTTAGTAGAGACGGGTTTCACCGTTTTAGCCGGGATGGTCTCGATCTCCTTGACCTCGTGATCCGCCCGCCTCGGCCTCCCAAAG
TGCTGGGATTACAGGCGTGAGCCACCGCGCCCGGCC

### HG03516 chr12:58064416

- The same genomic coordinate as in HG02630 (page 33)

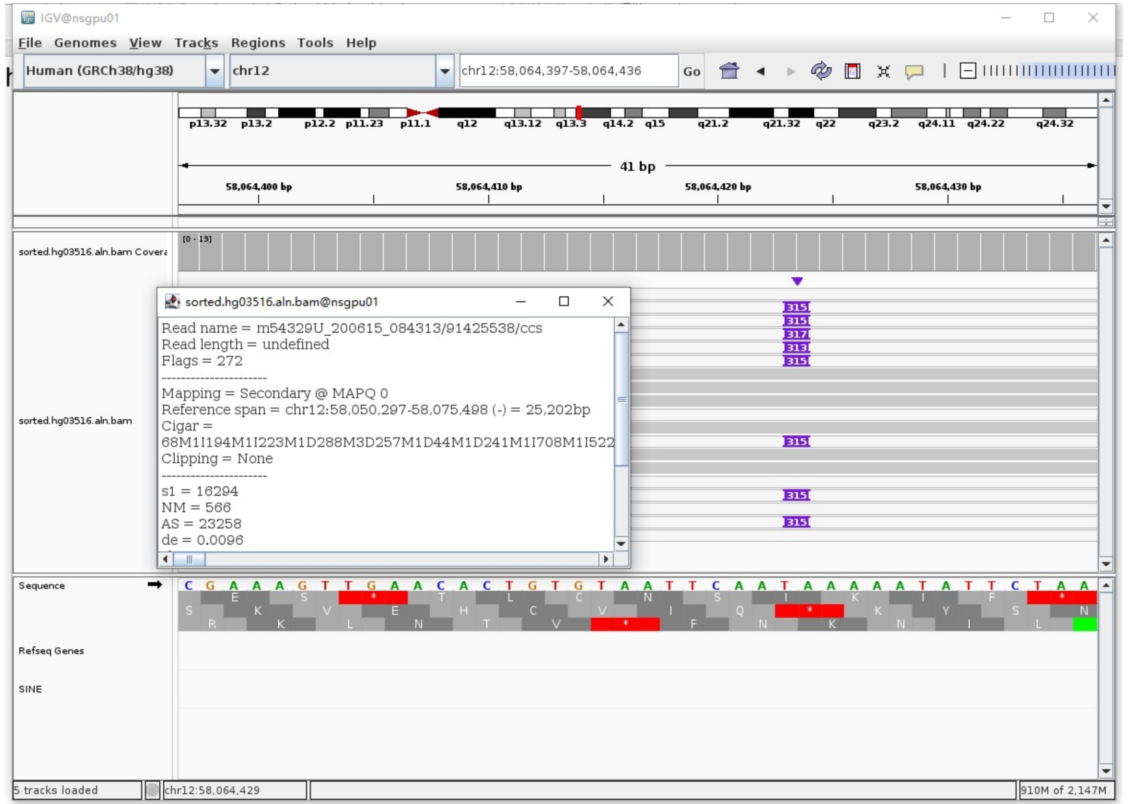

### HG03516 chr12:58961589

- The same genomic coordinate as in HG2630 (page 34)

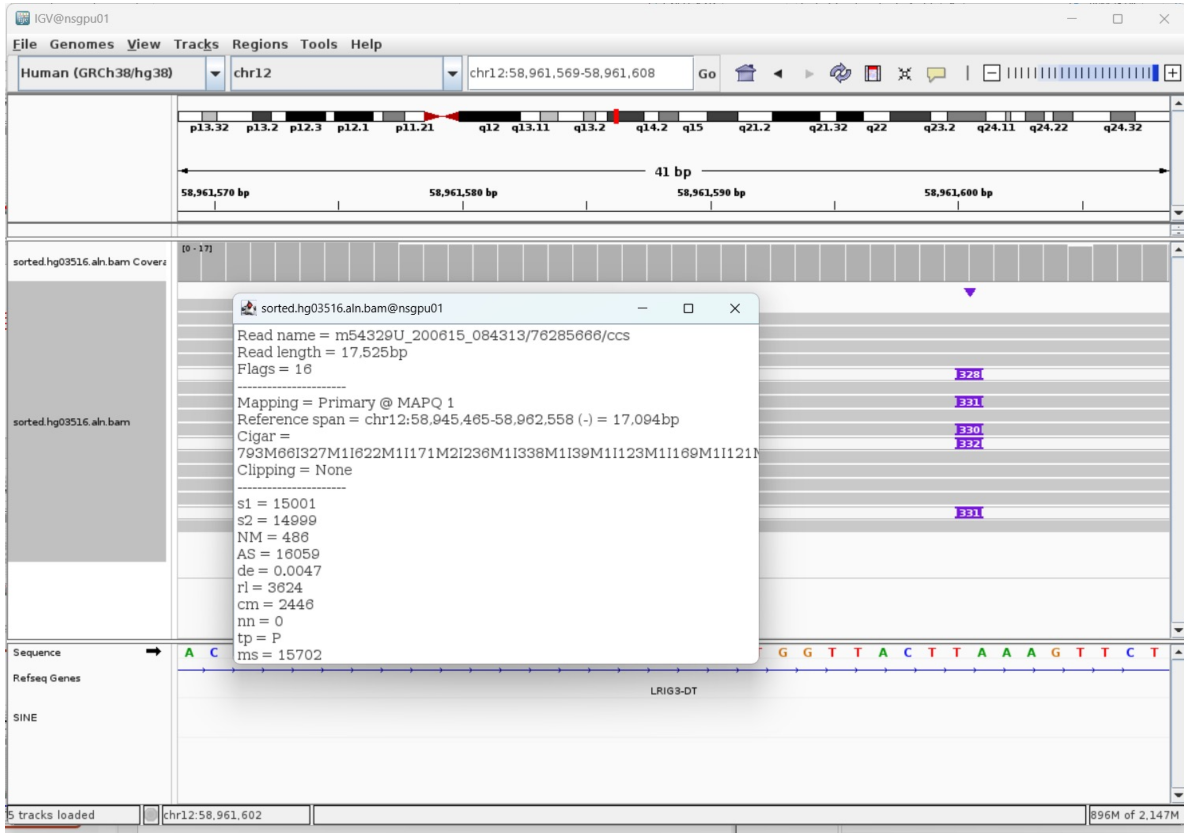

### HG03516 chr14:39716321

- Using the built-in BLAT tool in IGV to check where the inserted sequence aligned.
- The annotation at the aligned location indicated it as *AluY*.

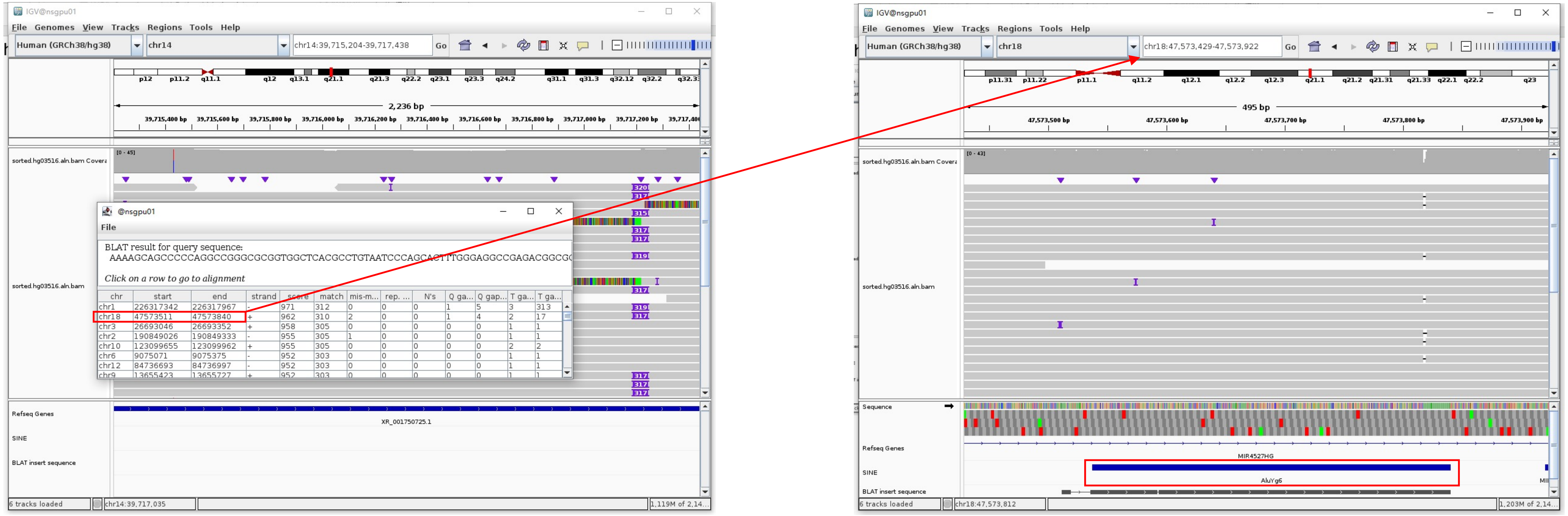


Insertion sequence (315bp)

- 15bp TSD

AAAAGCAGCCCCCAG

- Insertion:14-298 aligns to *AluY*c1:1-282, 96.80% identity

GGCCGGGGCGCGGTGGCTCACGCCTGTAATCCCAGCACTTTGGGAGGCCGAGACGGGGCGGGATCACGAGGTCAGGAGATCGAGACCATCCTGGCTAACACGGTGAAACCCCGTCTCTACTAA
AAATACAAAAATTAGCCGGGCATGGTGGCGCGTGCCTGTAGTCCCAGCTACACAGGAGGCTGAGGCAGGAGAATGGCGTGAACCCGGGAGGCGGAGCTTGCAGTGAGTCGAGATCGCGCCACT
GCACTCCAGCCTGGGCGACAGAGCGAAACTCCGTCTCA.........poly(A)

### HG03516 chr14:91658700

➢ Mapping quality is 0, indicating that this region may have alignment issues.

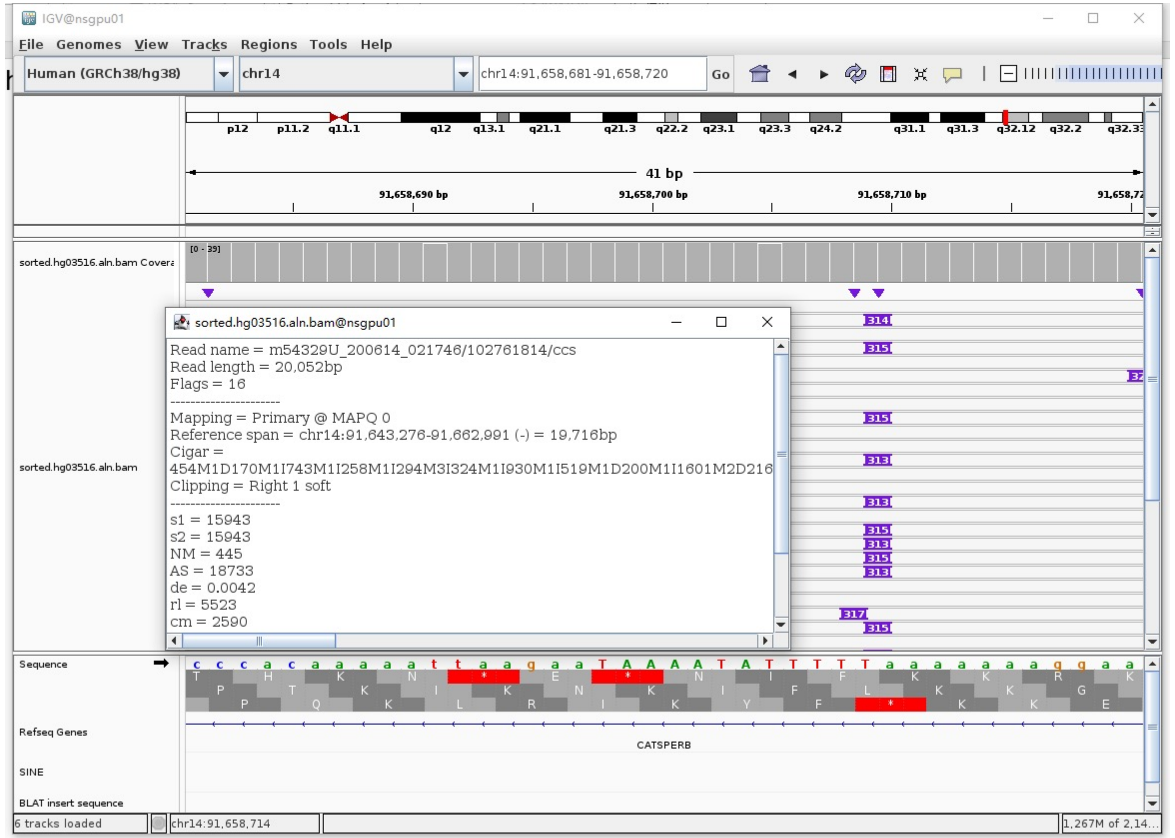


➢ Fetch one read (m54329U_200615_084313/118949166/ccs, 18769 bp) and BLAT it to the hg38 genome.

The best alignment is to chr14_KI270844v1_alt (an alternative sequence of chr14) and chr14.

| ACTIONS | QUERY | SCORE | START | END | QSIZE | IDENTITY | CHROM | STRAND | START | END | SPAN |
|---|---|---|---|---|---|---|---|---|---|---|---|
| browser new tab details | YourSeq | 18308 | 1 | 18769 | 18769 | 99.7% | chr14_KI270844v1_alt | + | 112587 | 131024 | 18438 |
| browser new tab details | YourSeq | 18308 | 1 | 18769 | 18769 | 99.7% | chr14 | + | 91644792 | 91663229 | 18438 |

- Read:1-13929 aligns to chr14_KI270844v1_alt

| ACTIONS | QUERY | SCORE | START | END | QSIZE | IDENTITY | CHROM | STRAND | START | END | SPAN |
|---|---|---|---|---|---|---|---|---|---|---|---|
| browser new tab details | YourSeq | 13843 | 1 | 13929 | 13929 | 99.8% | chr14_KI270844v1_alt | + | 112587 | 126518 | 13932 |
| browser new tab details | YourSeq | 13843 | 1 | 13929 | 13929 | 99.8% | chr14 | + | 91644792 | 91658723 | 13932 |

- Read:13930-14212 aligns to *Alu*Ya5:1-282, 99.65% identity

GGCCGGGCGCGGTGGCTCACGCCTGTAATCCCAGCACTTTGGGAGGCCGAGGCGGGCGGATCACGAGGTCAGGA
GATCGAGACCATCCCGGCTAAAACGGTGAAACCCCGTCTCTACTAAAAATACAAAAAATTAGCCGGGCGTAGTG
GCGGGCGCCTGTAGTCCCAGCTACTTGGGAGGCTGAGGCAGGAGAATGGTGTGAACCCGGGAGGCGGAGCTTGC
AGTGAGCCGAGATCCCGCCACTGCACTCCAGCCTGGGCGACAGAGCGAGACTCCGTCTCA

15bp TSD

AAAAAAAGGAAGGGA

- Read:14213-18769 also aligns to chr14_KI270844v1_alt

| ACTIONS | QUERY | SCORE | START | END | QSIZE | IDENTITY | CHROM | STRAND | START | END | SPAN |
|---|---|---|---|---|---|---|---|---|---|---|---|
| browser new tab details | YourSeq | 4480 | 16 | 4556 | 4556 | 99.5% | chr14_KI270844v1_alt | + | 126505 | 131024 | 4520 |
| browser new tab details | YourSeq | 4480 | 16 | 4556 | 4556 | 99.5% | chr14 | + | 91658710 | 91663229 | 4520 |

➢ It's an *Alu*Ya5 insertion.

### HG03516 chr14:106651070

- ➢ Mapping quality is 0, indicating that this region may have alignment issues.

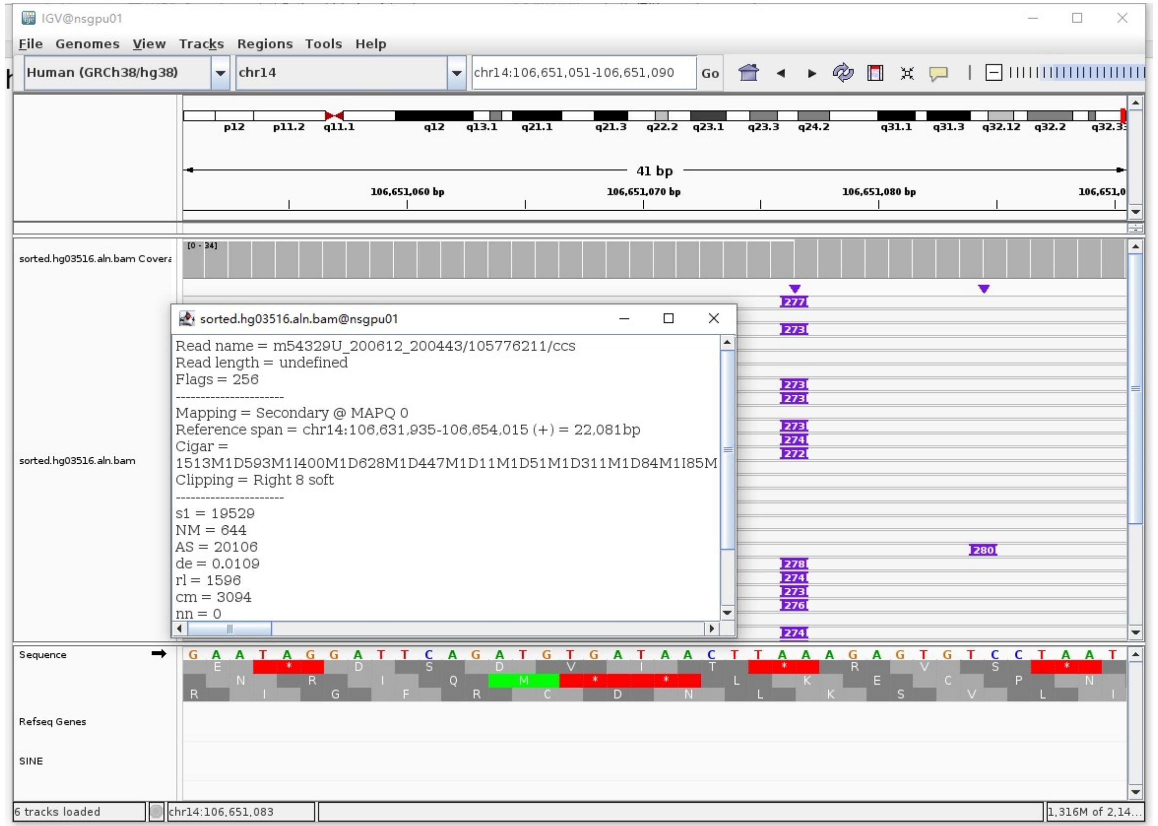


- ➢ Fetch one read (m54329U_200612_200443/105776211/ccs, 22290 bp) and BLAT it to the hg38 genome.

  The best alignment is to chr14_KI270846v1_alt (an alternative sequence of chr14) and chr14.

| ACTIONS | QUERY | SCORE | START | END | QSIZE | IDENTITY | CHROM | STRAND | START | END | SPAN |
|---|---|---|---|---|---|---|---|---|---|---|---|
| browser new tab details | YourSeq | 21600 | 1 | 22290 | 22290 | 99.1% | chr14_KI270846v1_alt | + | 1123614 | 1145703 | 22090 |
| browser new tab details | YourSeq | 21600 | 1 | 22290 | 22290 | 99.1% | chr14 | + | 106631935 | 106654024 | 22090 |

- Read:1-19109 aligns to chr14_KI270846v1_alt

| ACTIONS | QUERY | SCORE | START | END | QSIZE | IDENTITY | CHROM | STRAND | START | END | SPAN |
|---|---|---|---|---|---|---|---|---|---|---|---|
| browser new tab details | YourSeq | 18685 | 1 | 19076 | 19109 | 99.1% | chr14_KI270846v1_alt | + | 1123614 | 1142773 | 19160 |
| browser new tab details | YourSeq | 18685 | 1 | 19076 | 19109 | 99.1% | chr14 | + | 106631935 | 106651094 | 19160 |

- Read:19110-19390 aligns to *Alu*Yb8:289-21, 93.66% identity

  TGAGACGGAGTCTCGCTCTGTCGCCCAGGCCGGACTGCGGACTGCAGTGGCGCAATCTCGGCTCACTGCAAGCT
  CCGCTTCCCGGGTTCACGCCATTCTCCTGCCTCAGCCTCCCGAGTAGCTGGGACTACAGGCGCCCGCCACCGCG
  CCCGGCTAATTTTTTGTATTTTTAGTAGAGACGGGGTTTCACCTTGTTAGCCAGGATGGTCTCGATCTCCTGAA
  GAGTGTCCTAATTCTTAACCCACAATTAGGCCTGAGAAGCAGTCACAGGCACTGGAGG

- Read:19391-22290 also aligns to chr14_KI270846v1_alt

| ACTIONS | QUERY | SCORE | START | END | QSIZE | IDENTITY | CHROM | STRAND | START | END | SPAN |
|---|---|---|---|---|---|---|---|---|---|---|---|
| browser new tab details | YourSeq | 2873 | 1 | 2899 | 2899 | 99.5% | chr14_KI270846v1_alt | + | 1142817 | 1145703 | 2887 |
| browser new tab details | YourSeq | 2873 | 1 | 2899 | 2899 | 99.5% | chr14 | + | 106651138 | 106654024 | 2887 |

- ➢ It's an *Alu*Yb8 insertion.

### HG03516 chr17:37561902

➢ The same genomic coordinate as in HG00621 (page 15)

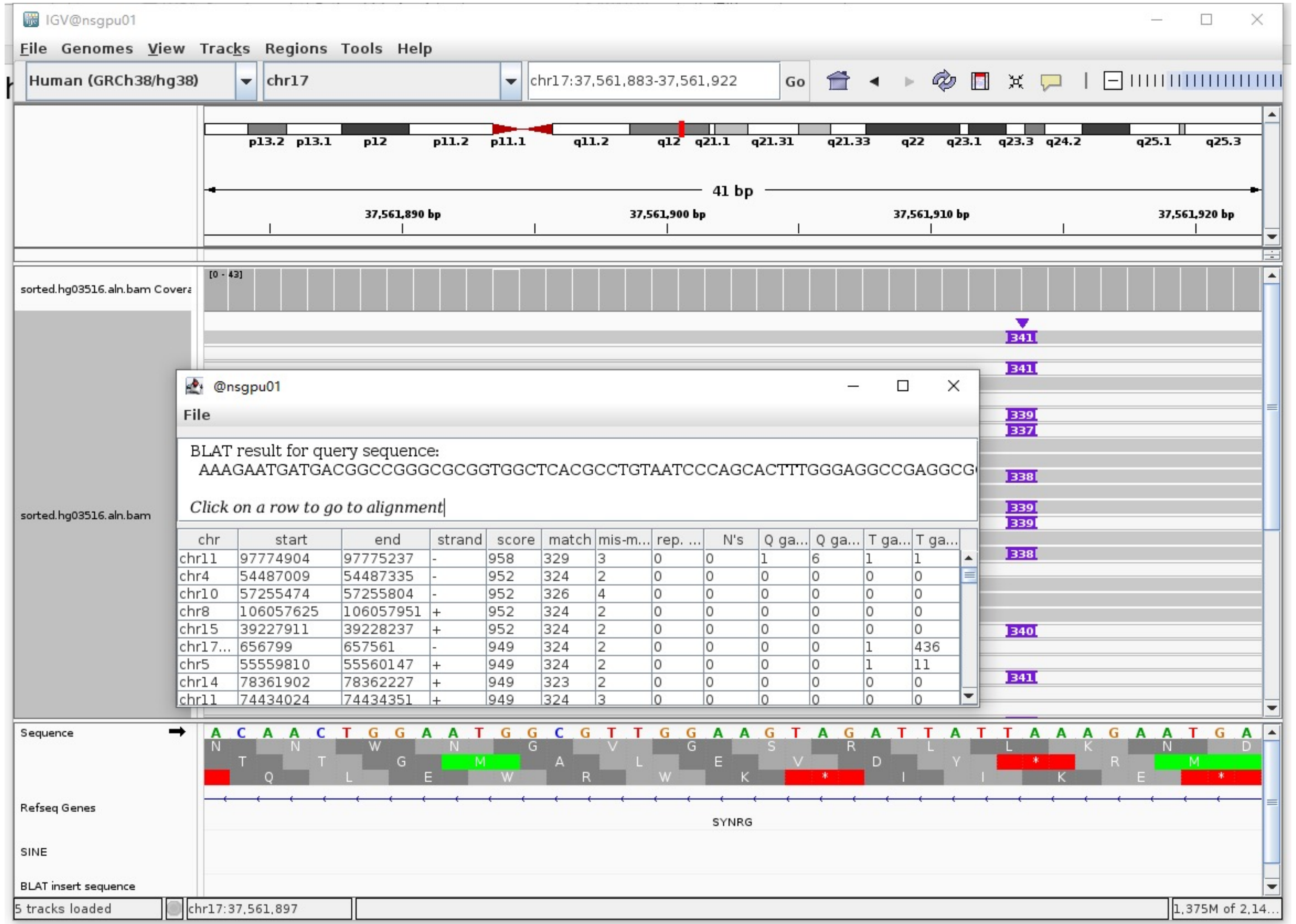

### HG03540 chr1:82577503

- The same genomic coordinate as in HG01106 (page 19)

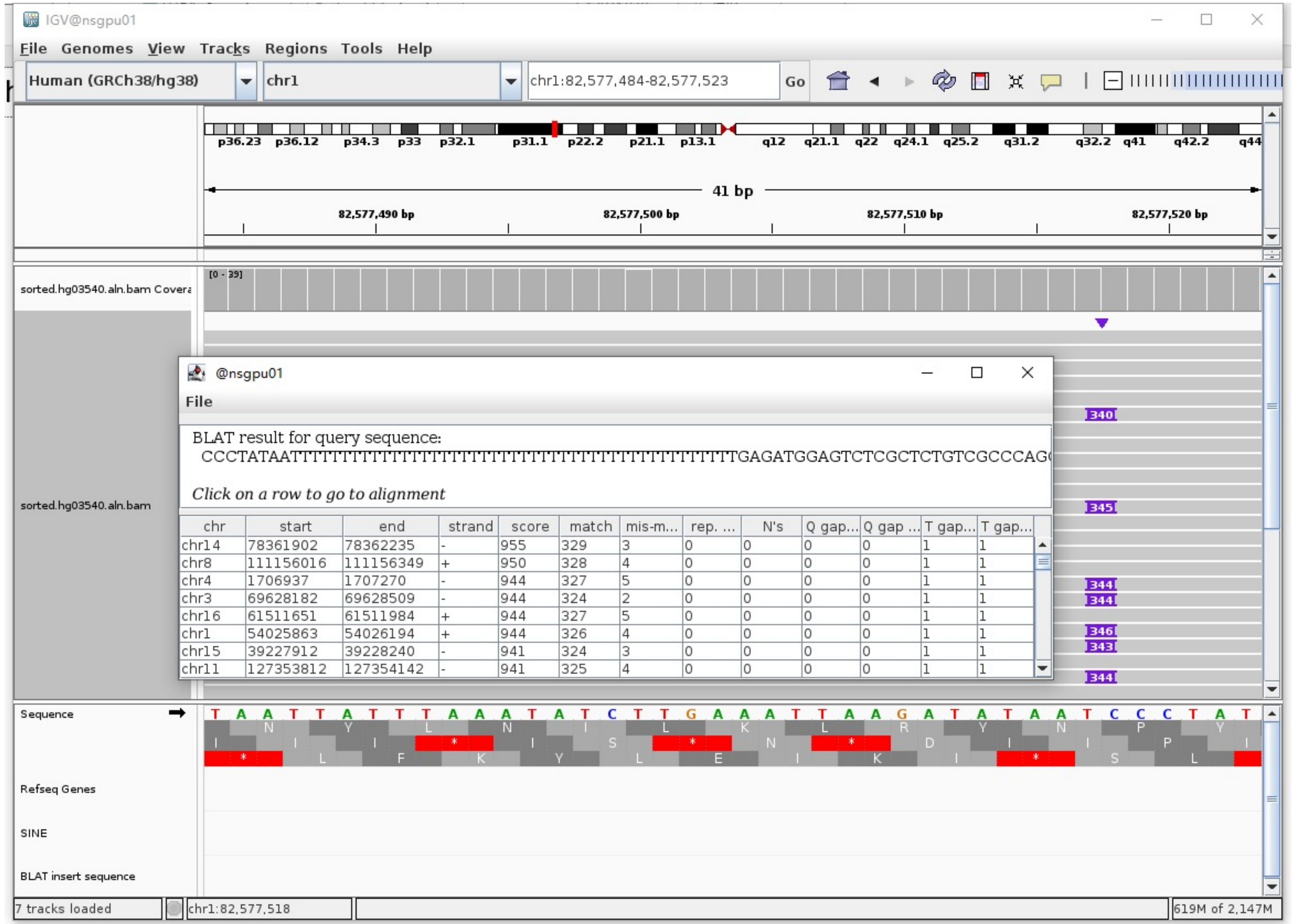

### HG03540 chr1:170913134

- Using the built-in BLAT tool in IGV to check where the inserted sequence aligned.
- The annotation at the aligned location indicated it as *Alu*Y.

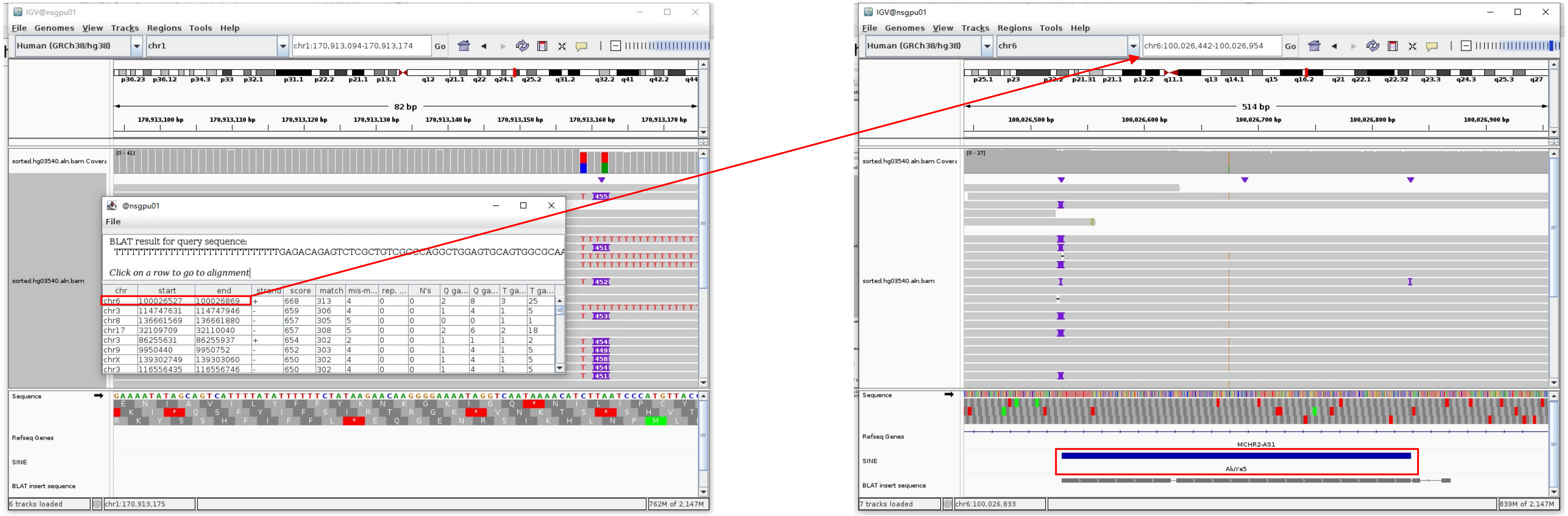


Insertion sequence (452bp)

- Insertion:26-306 aligns to *Alu*Yc1:282-1, 97.86% identity

TGAGACAGAGTCTCGCTGTCGCCCAGGCTGGAGTGCAGTGGCGCAATCTCGGCTCACTGCAAGCTCCGCCTCCCGGGTTCACGCCATTCTCCTGCCTCAGCCTCCCAAGTAGCTGGGACTACAGGCGCCCACCAC
CTCGCCCGGCTAATTTTTTGTATTTTTAGTAGAGACGGGGTTTCACCGTGTTAGCCAGGATGGTCTCGATCTCCTGACCTCGTGATCCGCCCGCCTCGGCCTCCCAAAGTGCTGGGATTACAGGCGTGAGCCACC
GCGCCCGGCC

- Insertion:307-452 aligns to chr1:170913147-170913292, 100% identity

TCAATAAAACATCTTAATCCCATGTTACCATTTTGTATGTTATGGATTTTTTTTATTAGAGAAAAATACACATTTACAGTAGAAAATTTGGGAAATATTCAAATAAAAACTGATATAAGAAAAAGAAATCATAAA
ATATGATAAAA

### HG03540 chr1:170913303

- ➢ Using the built-in BLAT tool in IGV to check where the inserted sequence aligned.
- ➢ The annotation at the aligned location indicated it as *AluY*.

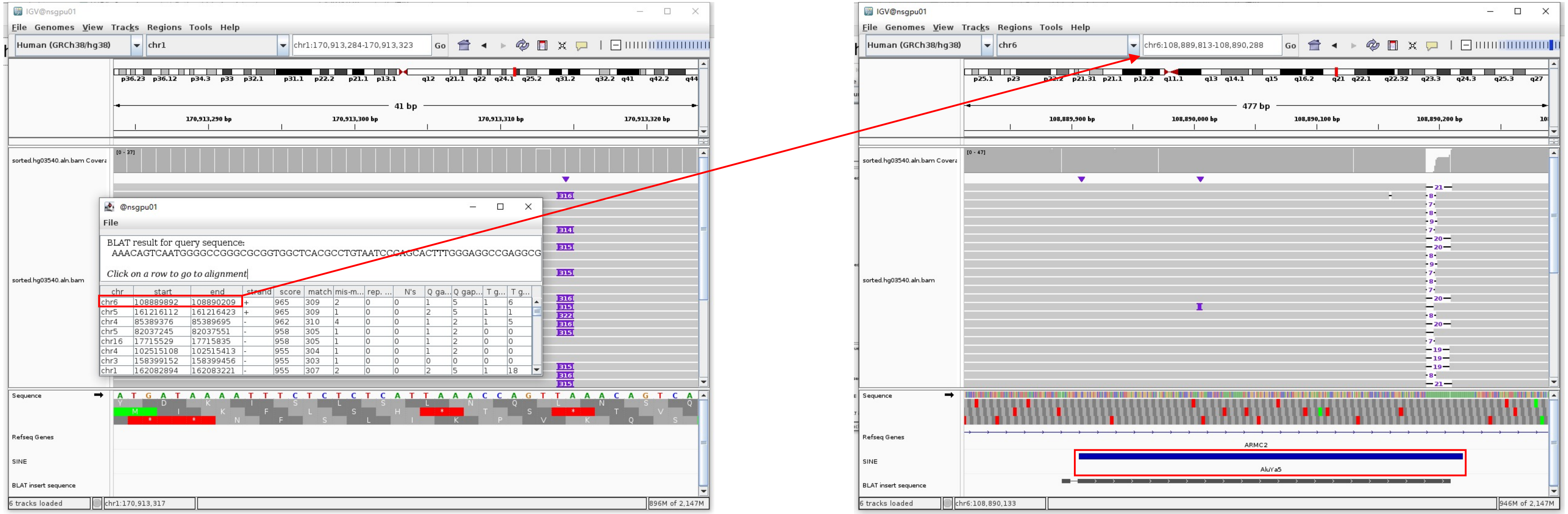


Insertion sequence (314bp)

- ➢ 13bp TSD

AAAAGCAGCCCCCAG

- ➢ Insertion:13-297 aligns to *Alu*Ya5:1-282, 99.65% identity

GGCCGGGCGCGGTGGCTCACGCCTGTAATCCCAGCACTTTGGGAGGCCGAGGCGGGTGGATCACGAGGTCAGGAGATCGAGACCATCCCGGCTAAAACGGTGAAACCCCGTCTCTACTAAAAA
TACAAAAAAAATTAGCCGGGCGTAGTGGCGGGCGCCTGTAGTCCCAGCTACTTGGGAGGCTGAGGCAGGAGAATGGCGTGAACCCGGGAGGCGGAGCTTGCAGTGAGCCGAGATCCCGCCACT
GCACTCCAGCCTGGGCGACAGAGCGAGACTCCGTCTCA.........poly(A)

### HG03540 chr1:238293249

- Using the built-in BLAT tool in IGV to check where the inserted sequence aligned.
- The annotation at the aligned location indicated it as *AluY*.

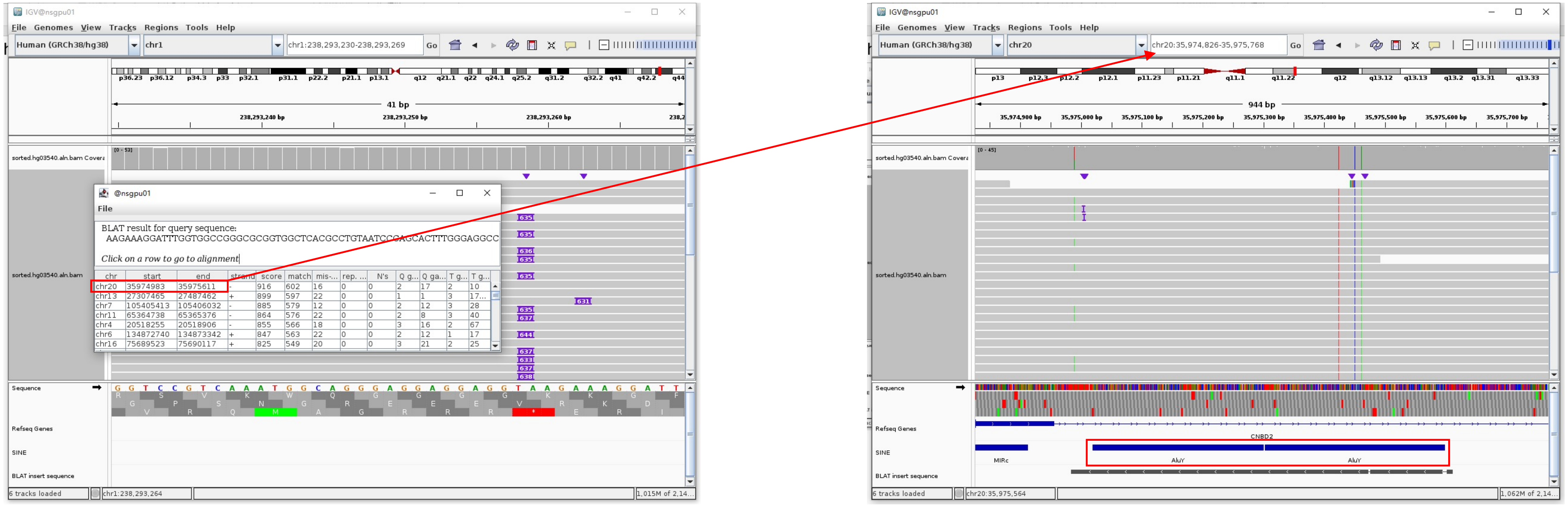


Insertion sequence (635bp)

- 15bp TSD

AAGAAAGGATTTGGT

- Insertion:15-301 aligns to *Alu*Yc1:0-282, 98.58% identity

GGCCGGGCGCGGTGGCTCACGCCTGTAATCCCAGCACTTTGGGAGGCCGAGGCGGGCGGATCACGAGGTCAGGAGATCGAGACCATCCTGGCTAACACGGTGAAACCCCGTATCTACTAAAAACACACAAAAAAAT
TAGCCGGGCGTGGTGGCGGGCGCCTGTAGTCCCAGCTACGCGGGAGGCTGAGGCAGGAGAATGGCGTGAACCCGGGAGGCGGAGCTTGCAGTGAGCCGAGATCGCGCCACTGCACTCCAGCCTGGGCGACAGAGCGA
GACTCCGTCTCA………poly(A)

- Insertion:318-600 aligns to *Alu*Yc1:0-282, 99.65% identity

GGCCGGGCGCGGTGGCTCACGCCTGTAATCCCAGCACTTTGGGAGGCCGAGGCGGGCGGATCACGAGGTCAGGAGATCGAGACCATCGTGGCTAACACGGTGAAACCCCGTCTCTACTAAAAATACAAAAAATTAGC
CGGGCGTGGTAGCGGGCGCCTGTAGTCCCAGCTACTCGGGAGGCTGAGGCAGGAGAATGGCGTGAACCCGGGAGGCGGAGCTTGCAGTGAGCCGAGATCGCGCCACTGCACTCCAGCCTGGGCGACAGAGCGAGACT
CCGTCTCA………poly(A)

### HG03540 chr4:63331066

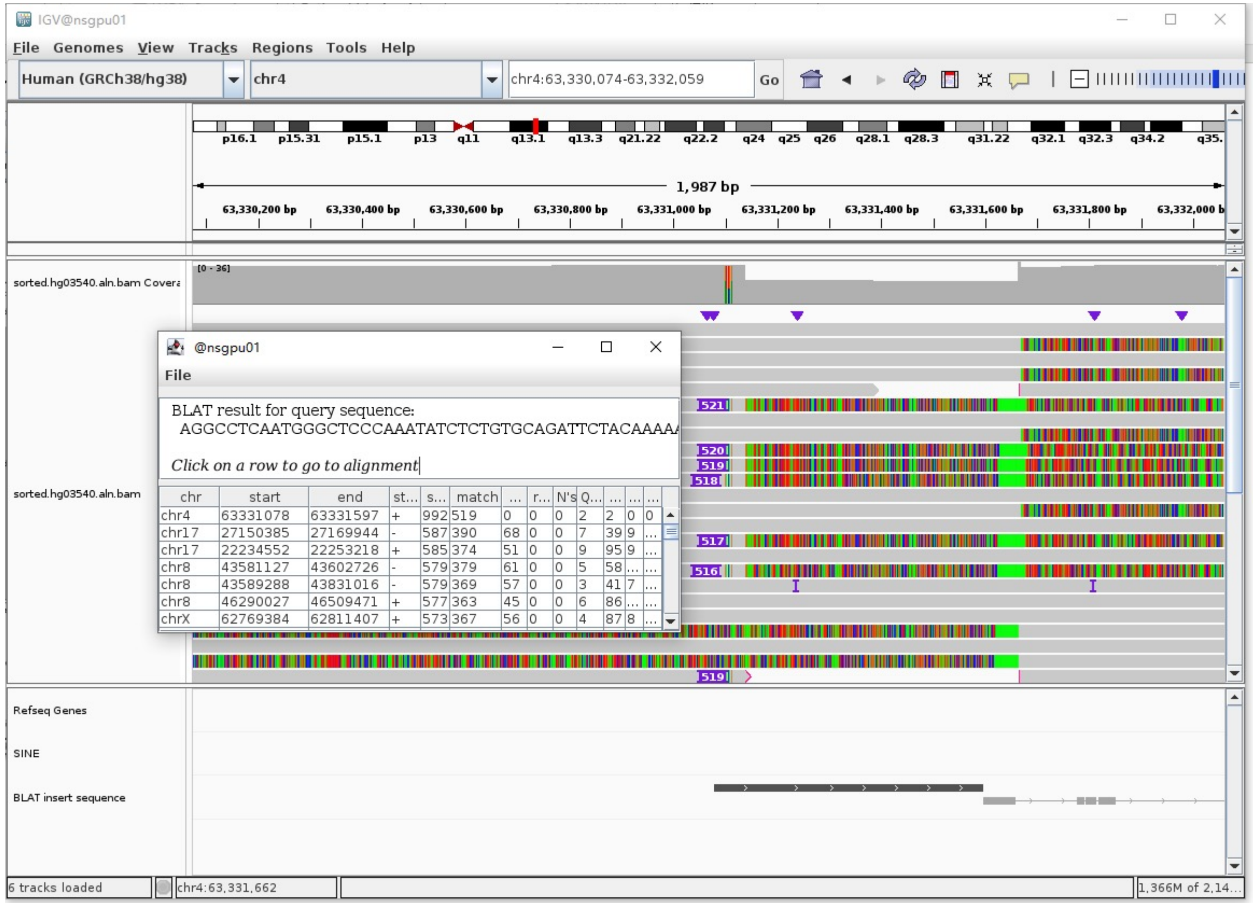


Insertion sequence (516bp)

- Insertion:1-516 aligns to chr4:63331068-63331585

AGGCCTCAATGGGCCTCCCAAATATCTCTGTGCAGATTCTACAAAAAGAGTGTTTCCA
ACCTGCAGAATCAAACAAAAAAGTTAAACTCTGTGTGATTTATCCACACATTGGAAAG
CATTTTCACACATACCTTTTTTCTAGTTTTTGTCACAGAATATTCGGTTTTTCACTAT
AGACATAAAAGGGCTCCAAAATGTCCTTTCGCAGATTCAACAAAAAGACTGTTTCCAA
CATGCTGAATCAAAAGAAAGGTTTAACTCTGTGAGTGAATCCACACATTGCAAAGCAT
TTTCAGAGCTAGCTTCTTTCCAGTTTTTGTTGAGGGATATTCAGTTTTTTCACTCTAT
GCTTCAATCGGCTCCCAAATGTCCCTTTGTAGATTCTACAAAAAAAAGTGATCCCAAC
CTGCTGAATGAAAATAAAGGTTTATCTCTGTGAGATGATTCCAACATCACAAAGCATT
TTCAAGGAAAGCTTCTTTCTAGTTTTTATCATGTGATATGAGTTTTTCACTAC

Right-clipped read (5134bp)

- Clip:1-197 aligns to chr4:63331659-63332029, 100% identity
- Clip:198-487 aligns to *Alu*Yb8:1-289, 98.96% identity

GGCCGGGCGCGGTGGCTCACGCCTGTAGTCCCAGCACTTTGGGAGGCCGAGGCGGGTG
GATCATGAGGTCAGGAGATCGAGACCATCCTGGCTAACACGGTGAAACCCCGTCTCTA
CTAAAAATACAAAAAATTAGCCGGGCGCGGTGGCGGGCGCCTGTAGTCCCAGCTACTG
GGGAGGCTGAGGCAGGAGAATGGCGTGAACCCGGGAAGCGGAGCTTGCAGTGAGCCGA
GATTGCGCCACTGCAGTCCGCAGTCCGGCCTGGGCGACAGAGCGAGACTCCGTCTCA

- Clip:534-5134 aligns to chr4:63331666-63336280, 99.7% identity

### HG03540 chr4:68078452

➢ Using the built-in BLAT tool in IGV to check where the inserted sequence aligned.

➢ The annotation at the aligned location indicated it as *Alu*Y.

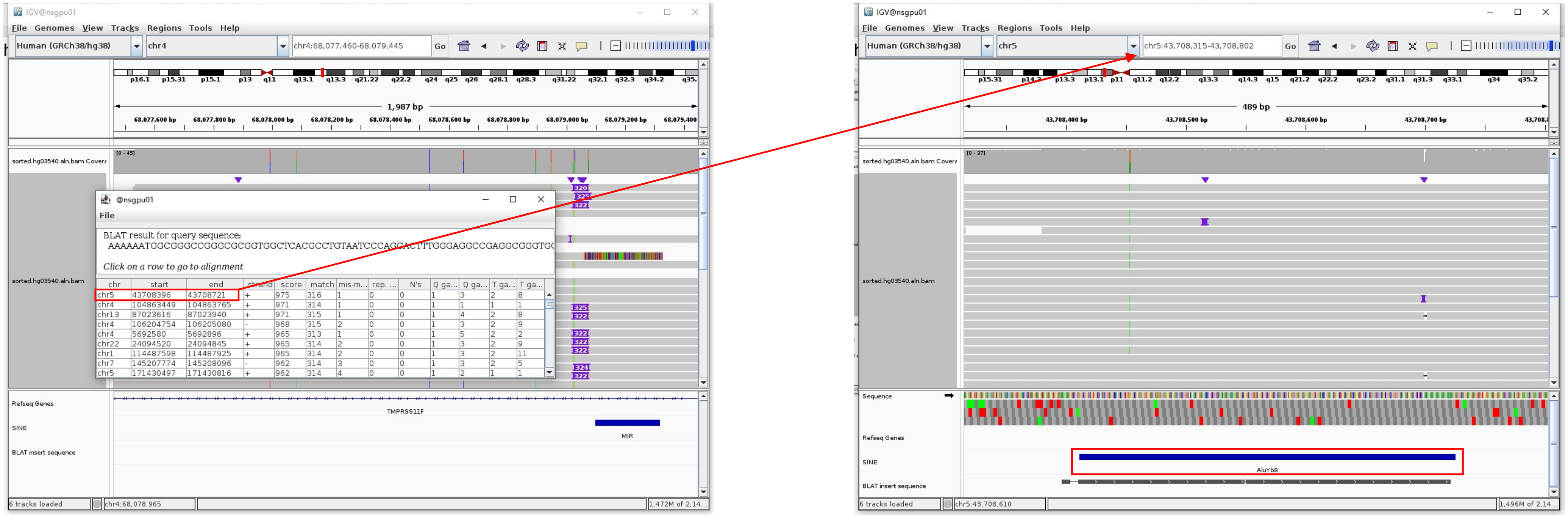


Insertion sequence (320bp)

➢ 10bp TSD

AAAAAATGGC

➢ Insertion:11-299 aligns to *Alu*Yb8:1-289, 99.65% identity

GGCCGGGCGCGGTGGCTCACGCCTGTAATCCCAGCACTTTGGGAGGCCGAGGCGGGTGGATCATGAGGTCAGGAGATCGAGACCATCCTGGCTAACAAGGTGAAACCCCGTCTCTACTAAGAA
TACAAAAAATTAGCCGGGCGCGGTGGCGGGCGCCTGTAGTCCCAGCTACTCGGGAGGCTGAGGCAGGAGAATGGCGTGAACCCGGGAAGCGGAGCTTGCAGTGAGCCGAGATTGCGCCACTGCA
GTCCGCAGTCCGGCCTGGGCGACAGAGCGAGACTCCGTCTCA.........poly(A)

### HG03540 chr4:68329636

➢ Mapping quality is 0, indicating that this region may have alignment issues.

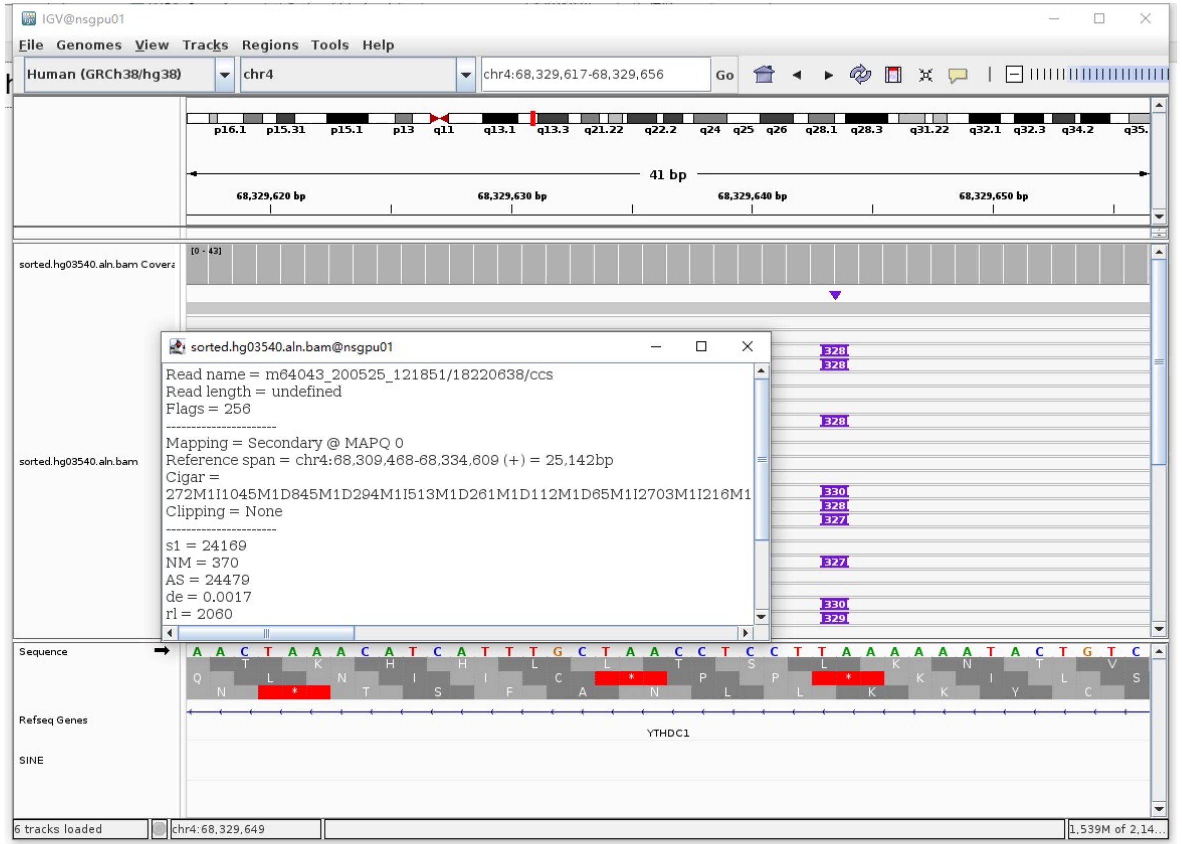


➢ Fetch one read (m64043_200525_121851/74779323/ccs, 21119 bp) and BLAT it to the hg38 genome.

The best alignment is to chr4_GL000257v2_alt (an alternative sequence of chr4) and chr4.

| ACTIONS | QUERY | SCORE | START | END | QSIZE | IDENTITY | CHROM | STRAND | START | END | SPAN |
|---|---|---|---|---|---|---|---|---|---|---|---|
| browser new tab details | YourSeq | 20752 | 1 | 21119 | 21119 | 99.9% | chr4_GL000257v2_alt | + | 8764 | 29549 | 20786 |
| browser new tab details | YourSeq | 20752 | 1 | 21119 | 21119 | 99.9% | chr4 | + | 68315122 | 68335907 | 20786 |

● Read:1-14542 aligns to chr4_GL000257v2_alt

| ACTIONS | QUERY | SCORE | START | END | QSIZE | IDENTITY | CHROM | STRAND | START | END | SPAN |
|---|---|---|---|---|---|---|---|---|---|---|---|
| browser new tab details | YourSeq | 14510 | 1 | 14541 | 14541 | 99.9% | chr4_GL000257v2_alt | + | 8765 | 23300 | 14536 |
| browser new tab details | YourSeq | 14510 | 1 | 14541 | 14541 | 99.9% | chr4 | + | 68315123 | 68329658 | 14536 |

● Read:14543-14825 aligns to *Alu*Ya5:1-282, 98.58% identity

GGCCGGGCGCGGTGGCTCACGCCTGTAATCCCAGCACTTTGGGAGGCCGAGGCGGGCGGATCACGAGGTCAGGA
GATCGAGACCATCCTGGCTAACAAGGTGAAACCCCGTCTCTACTAAAAATACAAAAAATTAGCCGGGCGTAGTG
GCGGGCGCCTGTAGTCCCAGCTACTTGGGAGGCTGAGGCAGGAGAATGGCGTGAACCCGGGAGGCGGAGTTTGC
AGTGAGCCGAGATCCCGCCACTGCACTCCAGCCTGGGCGACAGAGCGAGACTCCGTCTCA

16bp TSD

AAAAATACTGTCCACG

● Read:14826-21119 also aligns to chr4_GL000257v2_alt

| ACTIONS | QUERY | SCORE | START | END | QSIZE | IDENTITY | CHROM | STRAND | START | END | SPAN |
|---|---|---|---|---|---|---|---|---|---|---|---|
| browser new tab details | YourSeq | 6257 | 31 | 6293 | 6293 | 100.0% | chr4_GL000257v2_alt | + | 23286 | 29549 | 6264 |
| browser new tab details | YourSeq | 6257 | 31 | 6293 | 6293 | 100.0% | chr4 | + | 68329644 | 68335907 | 6264 |

➢ It's an *Alu*Ya5 insertion

### HG03540 chr6:29925071

- ➢ Mapping quality is 0, indicating that this region may have alignment issues.

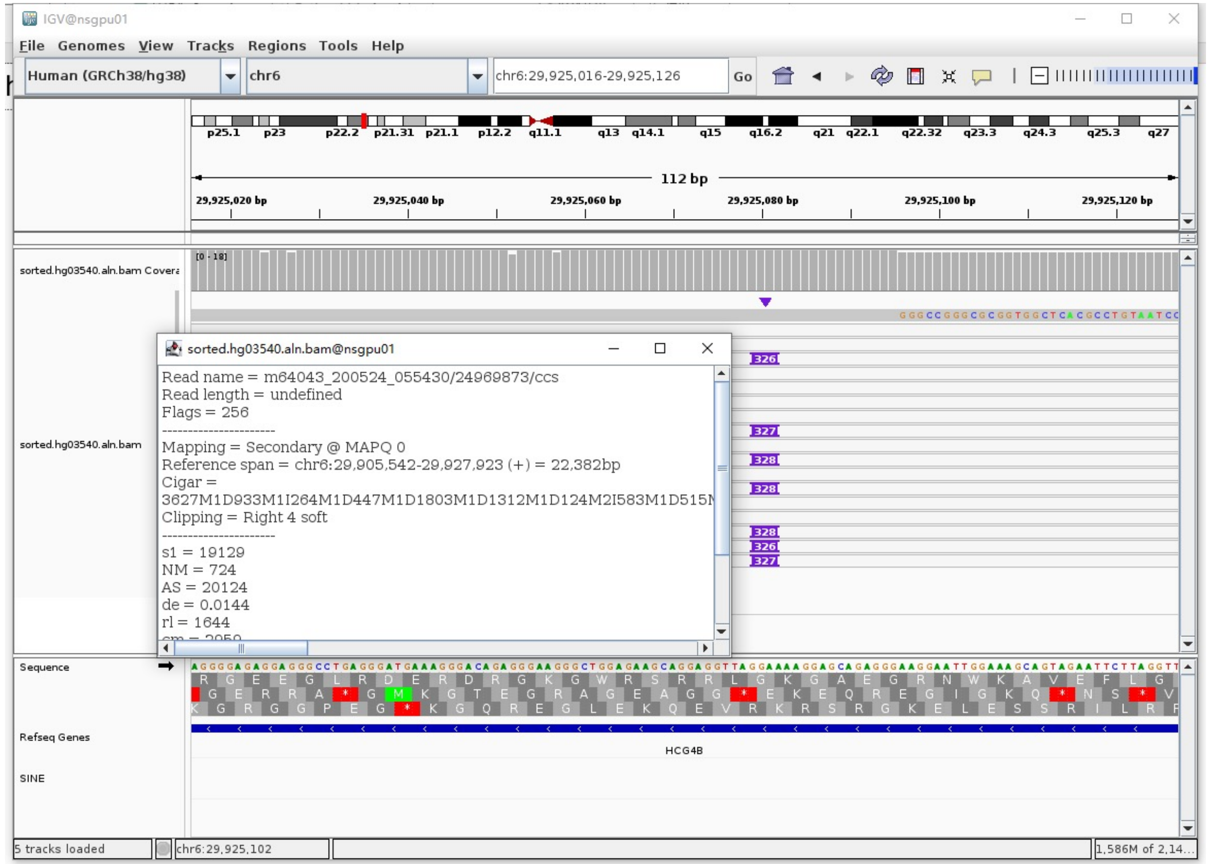


- ➢ Fetch one read (m64043_200524_055430/24969873/ccs, 22693 bp) and BLAT it to the hg38 genome.

  The best alignment is to chr6_GL000253v2_alt (an alternative sequence of chr6).

| ACTIONS | | | QUERY | SCORE | START | END | QSIZE | IDENTITY | CHROM | STRAND | START | END | SPAN |
|---|---|---|---|---|---|---|---|---|---|---|---|---|---|
| browser | new tab | details | YourSeq | 21987 | 1 | 22693 | 22693 | 99.2% | chr6_GL000253v2_alt | + | 1161876 | 1184289 | 22414 |
| browser | new tab | details | YourSeq | 21782 | 1 | 22693 | 22693 | 98.8% | chr6_GL000250v2_alt | + | 1165160 | 1187602 | 22443 |
| browser | new tab | details | YourSeq | 21778 | 1 | 22693 | 22693 | 98.8% | chr6_GL000251v2_alt | + | 1386834 | 1409276 | 22443 |
| browser | new tab | details | YourSeq | 21730 | 1 | 22693 | 22693 | 98.7% | chr6_GL000256v2_alt | + | 1205269 | 1227615 | 22347 |
| browser | new tab | details | YourSeq | 21715 | 1 | 22693 | 22693 | 98.6% | chr6_GL000255v2_alt | + | 1162290 | 1184715 | 22426 |
| browser | new tab | details | YourSeq | 21696 | 1 | 22693 | 22693 | 98.6% | chr6 | + | 29905542 | 29927927 | 22386 |

- ● Read:1-19580 aligns to chr6_GL000253v2_alt

| ACTIONS | | | QUERY | SCORE | START | END | QSIZE | IDENTITY | CHROM | STRAND | START | END | SPAN |
|---|---|---|---|---|---|---|---|---|---|---|---|---|---|
| browser | new tab | details | YourSeq | 19226 | 1 | 19579 | 19579 | 99.2% | chr6_GL000253v2_alt | + | 1161877 | 1181509 | 19633 |
| browser | new tab | details | YourSeq | 19041 | 1 | 19579 | 19579 | 98.8% | chr6_GL000256v2_alt | + | 1205270 | 1224832 | 19563 |
| browser | new tab | details | YourSeq | 19032 | 1 | 19579 | 19579 | 98.8% | chr6_GL000250v2_alt | + | 1165161 | 1184814 | 19654 |
| browser | new tab | details | YourSeq | 19028 | 1 | 19579 | 19579 | 98.8% | chr6_GL000251v2_alt | + | 1386835 | 1406488 | 19654 |
| browser | new tab | details | YourSeq | 18993 | 1 | 19579 | 19579 | 98.7% | chr6_GL000255v2_alt | + | 1162291 | 1181923 | 19633 |
| browser | new tab | details | YourSeq | 18977 | 1 | 19579 | 19579 | 98.6% | chr6_GL000252v2_alt | + | 1162646 | 1182293 | 19648 |
| browser | new tab | details | YourSeq | 18976 | 1 | 19579 | 19579 | 98.6% | chr6 | + | 29905543 | 29925095 | 19553 |

- ● Read:19581-19870 aligns to *Alu*Yb8:1-289, 99.65% identity

GGCCGGGCGCGGTGGCTCACGCCTGTAATCCCAGCACTTTGGGAGGCCGAGGCGGGTGGATCATGAGGTCAGGA
GATCGAGACCATCCTGGCTAACAAGGTGAAACCCCGTCTCTACTAAAAATACAAAAAATTAGCCGGGCGCGGTG
GCGGGCGCCTGTAGTCCCAGCTACTCGGGAGGCTGAGGCAGGAGAATGGCGTGAACCCGGGAAGCGGAGCTTGC
AGTGAGCCGAGATTGCGCCACTGCAGTCCGCAGTCCGGCCTGGGCGACAGAGCGAGACTCAGTCTCA

15bp TSD

AAAAGGAGCAGAGGG

- ● Read:19871-22693 also aligns to chr6_GL000253v2_alt

| ACTIONS | | | QUERY | SCORE | START | END | QSIZE | IDENTITY | CHROM | STRAND | START | END | SPAN |
|---|---|---|---|---|---|---|---|---|---|---|---|---|---|
| browser | new tab | details | YourSeq | 2776 | 21 | 2822 | 2822 | 99.5% | chr6_GL000253v2_alt | + | 1181495 | 1184289 | 2795 |
| browser | new tab | details | YourSeq | 2765 | 21 | 2822 | 2822 | 99.4% | chr6_GL000251v2_alt | + | 1406474 | 1409276 | 2803 |
| browser | new tab | details | YourSeq | 2765 | 21 | 2822 | 2822 | 99.4% | chr6_GL000250v2_alt | + | 1184800 | 1187602 | 2803 |
| browser | new tab | details | YourSeq | 2737 | 21 | 2822 | 2822 | 98.9% | chr6_GL000255v2_alt | + | 1181909 | 1184715 | 2807 |
| browser | new tab | details | YourSeq | 2735 | 21 | 2822 | 2822 | 98.9% | chr6 | + | 29925081 | 29927927 | 2847 |

- ➢ It's an *Alu*Yb8 insertion.

### HG03540 chr6:30254196

- ➢ Using the built-in BLAT tool in IGV to check where the inserted sequence aligned.

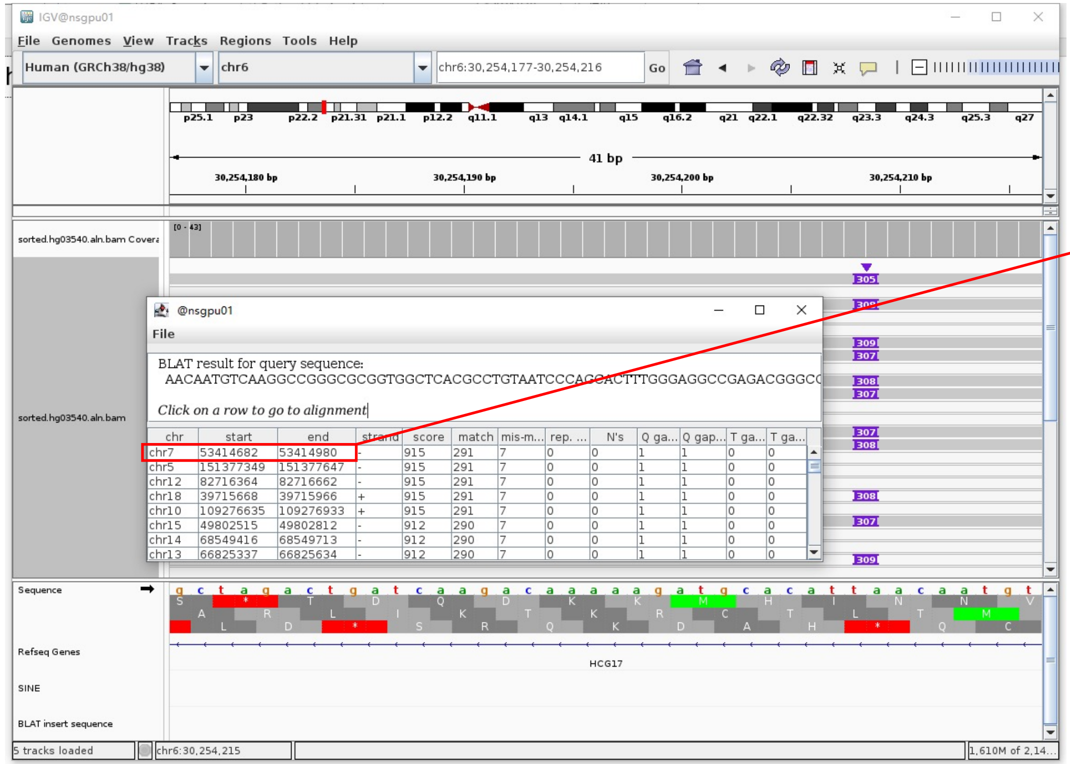


- ➢ The annotation at the aligned location indicated it as *AluY*.

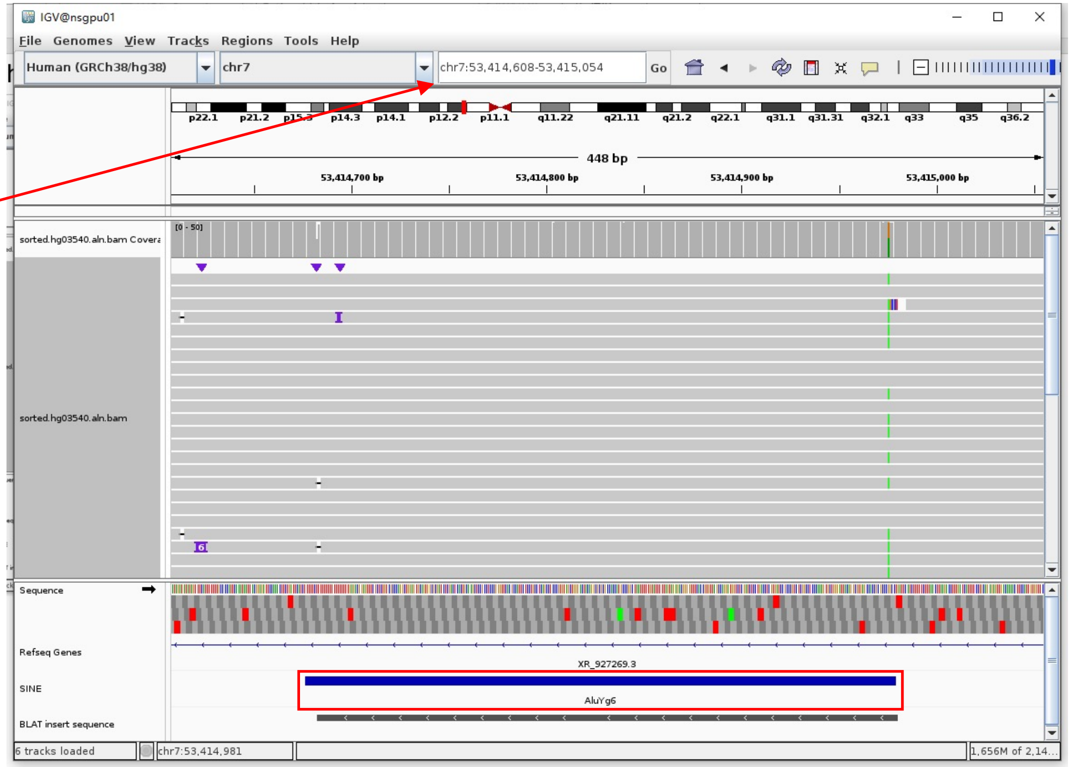


Insertion sequence (305bp)

- ➢ 12bp TSD

AACAATGTCAAG

- ➢ Insertion:11-289 aligns to *Alu*Yc1:1-282, 99.65% identity

GGCCGGGCGCGGTGGCTCACGCCTGTAATCCCAGCACTTTGGGAGGCCGAGACGGGCGGATCACGAGGTCAGGAGATCGAGACATCCTGGCTAACACGGTGAAACCCGTCTCTACTAAAAATA
CAAAATTAGCCGGGCATGGTGGCGGGCGCCTGTAGTCCCAGCTACACGGGAGGCTGAGGCAGGAGAATGGCGTGAACCCGGGAGGCGGAGCTTGCAGTGAGCGAGATTGCGCCACTGCCACTC
CAGCCTGGGCGACAAAGCGAGACTCCGTCTCA.........poly(A)

### HG03540 chr6:131694946

➢ Using the built-in BLAT tool in IGV to check where the inserted sequence aligned.

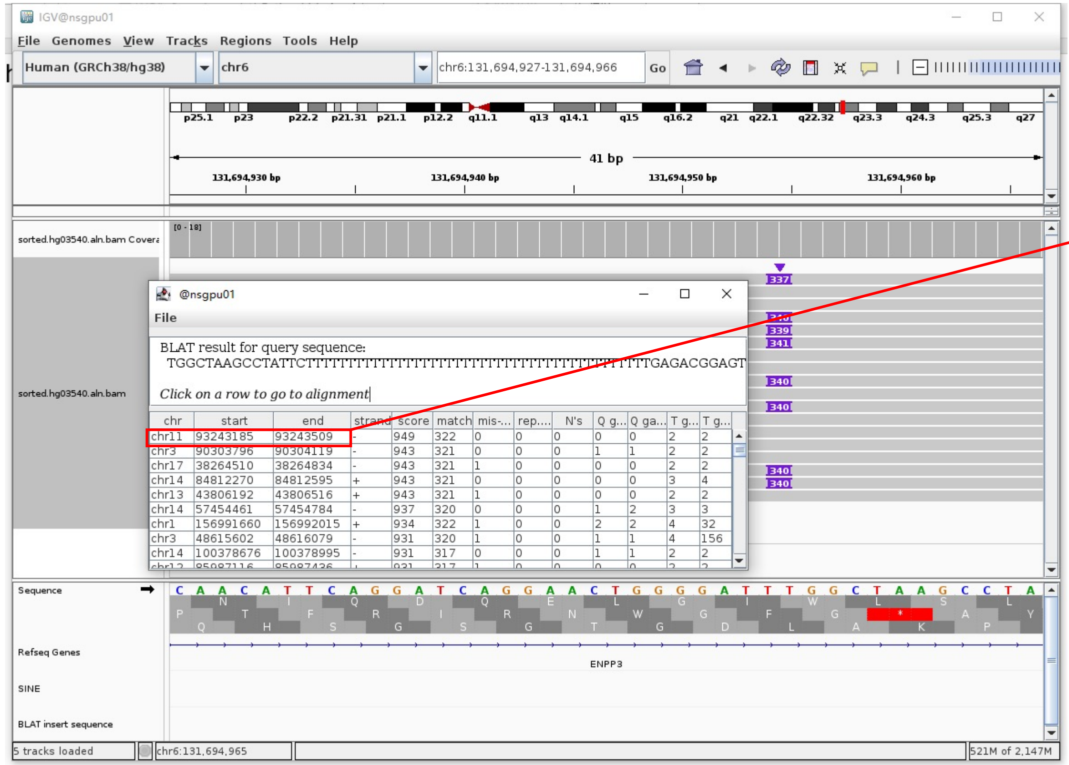


➢ The annotation at the aligned location indicated it as *Alu*Y.

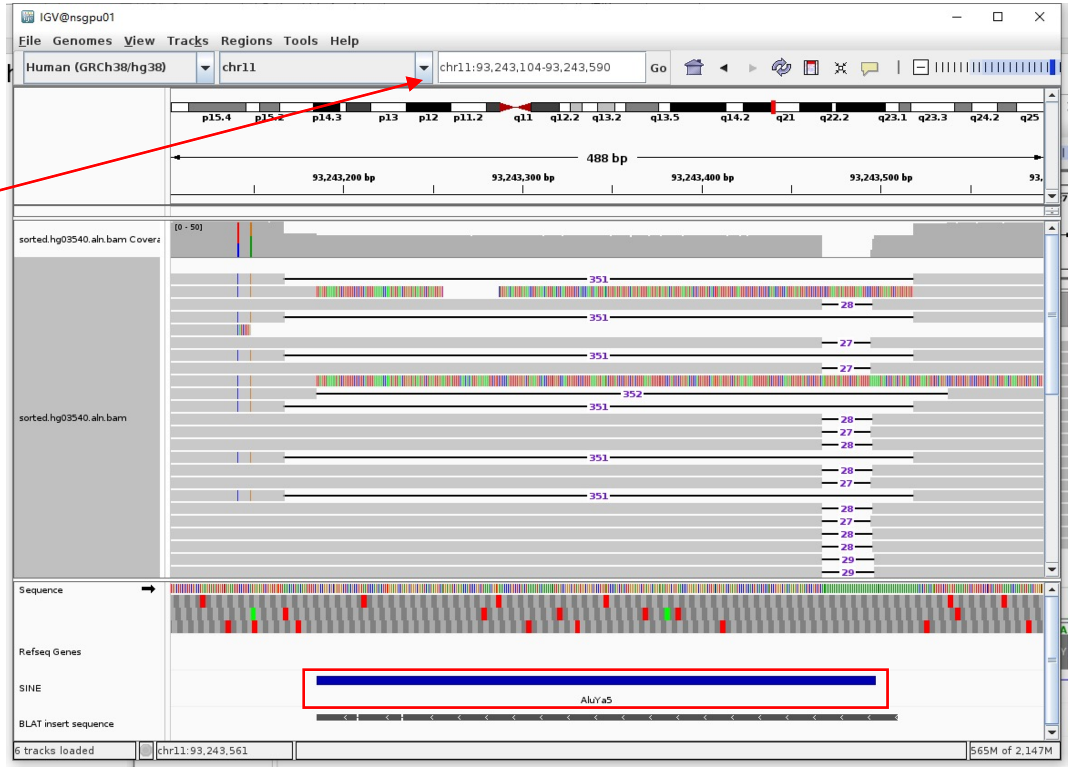


Insertion sequence (337bp)

➢ 16bp TSD

TGGCTAAGCCTATTCT

➢ Insertion:56-337 aligns to *Alu*Ya5:282-1, 100% identity

TGAGACGGAGTCTCGCTCTGTCGCCCAGGCTGGAGTGCAGTGGCGGGATCTCGGCTCACTGCAAGCTCCGCCTCCCGGGTTCACGCCATTCTCCTGCCTCAGCCTCCCAAGTAGCTGGGACTA
CAGGCGCCCGCCACTACGCCCGGCTAATTTTTTGTATTTTTAGTAGAGACGGGGTTTCACCGTTTTAGCCGGGATGGTCTCGATCTCCTGACCTCGTGATCCGCCCGCCTCGCCTCCCAAAG
TGCTGGGATTACAGCGTGAGCCACCGCGCCCGGCC

### HG03540 chr8:39842808

- The same genomic coordinate as in HG03516 (page 64)

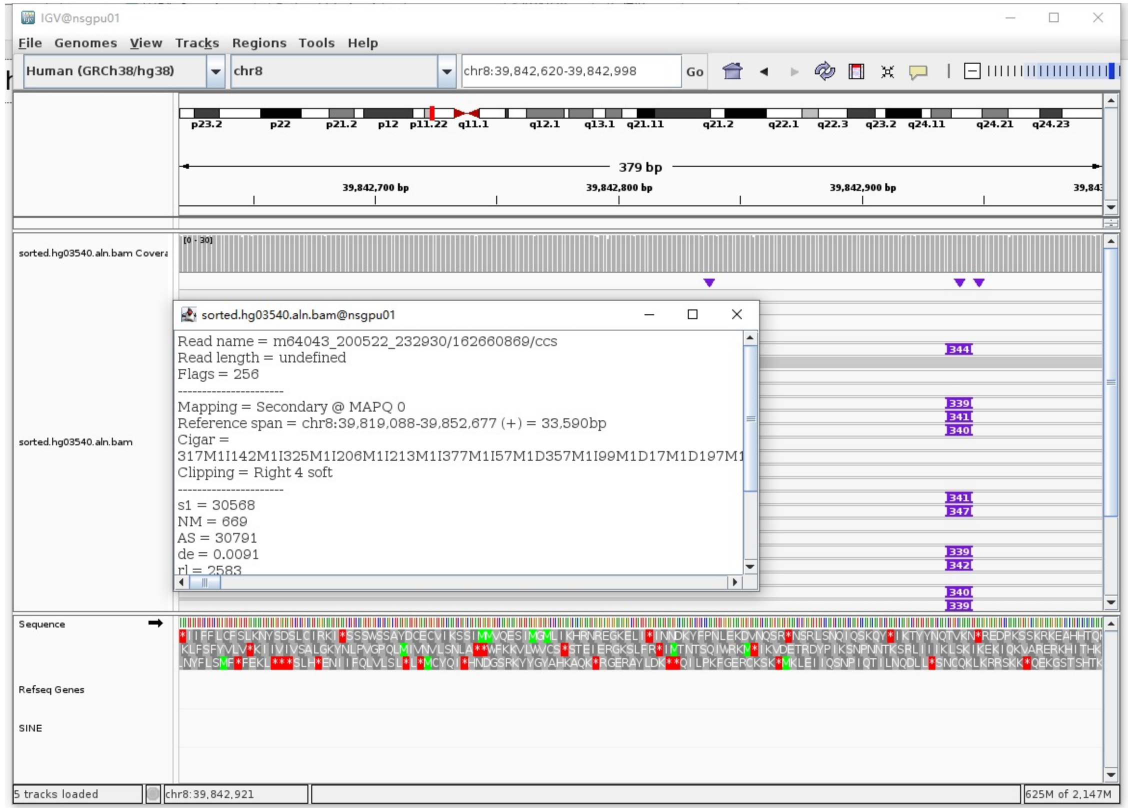

### HG03540 chr9:101285278

➢ Mapping quality is 0, indicating that this region may have alignment issues.

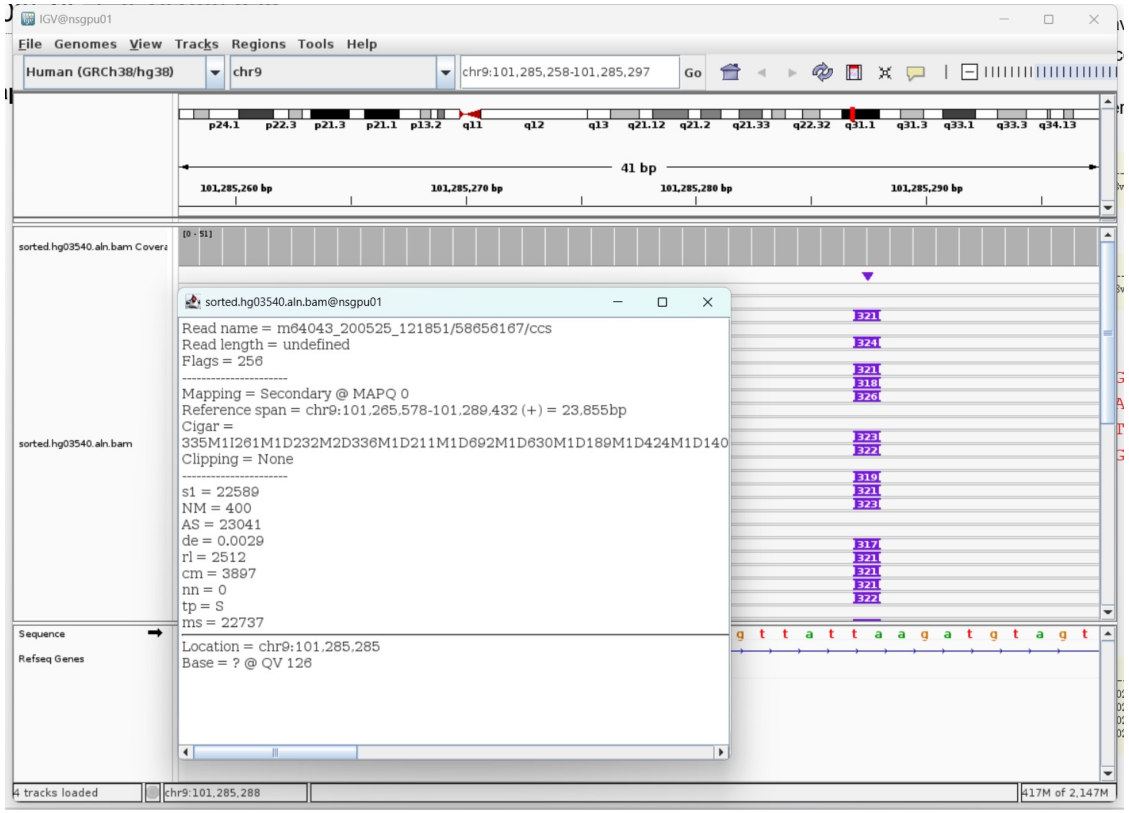


➢ Fetch one read (m64043_200521_171703/63374804/ccs, 18180 bp) and BLAT it to the hg38 genome.

The best alignment is to chr9_KI270823v1_alt (an alternative sequence of chr9) and chr9.

| ACTIONS | QUERY | SCORE | START | END | QSIZE | IDENTITY | CHROM | STRAND | START | END | SPAN |
|---|---|---|---|---|---|---|---|---|---|---|---|
| browser new tab details | YourSeq | 17767 | 1 | 18180 | 18180 | 99.7% | chr9_KI270823v1_alt | - | 323387 | 341247 | 17861 |
| browser new tab details | YourSeq | 17767 | 1 | 18180 | 18180 | 99.7% | chr9 | - | 101272184 | 101290044 | 17861 |

- Read:1-4809 aligns to chr9_KI270823v1_alt

| ACTIONS | QUERY | SCORE | START | END | QSIZE | IDENTITY | CHROM | STRAND | START | END | SPAN |
|---|---|---|---|---|---|---|---|---|---|---|---|
| browser new tab details | YourSeq | 4739 | 1 | 4759 | 4808 | 99.8% | chr9_KI270823v1_alt | - | 336491 | 341246 | 4756 |
| browser new tab details | YourSeq | 4739 | 1 | 4759 | 4808 | 99.8% | chr9 | - | 101285288 | 101290043 | 4756 |

- Read:4810-5071 aligns to *Alu*Ya5:282-21, 100% identity

TGAGACGGAGTCTCGCTCTGTCGCCCAGGCTGGAGTGCAGTGGCGGGATCTCGGCTCACTGCAAGCTCC
GCCTCCCGGGTTCACGCCATTCTCCTGCCTCAGCCTCCCAAGTAGCTGGGACTACAGGCGCCCGCCACT
ACGCCCGGCTAATTTTTTGTATTTTTAGTAGAGACGGGGTTTCACCGTTTTAGCCGGGATGGTCTCGAT
CTCCTGACCTCGTGATCCGCCCGCCTCGGCCTCCCAAAGTGCTGGGATTACAGG

15bp TSD

AAGATGTAGTCCCTG

- Read:5072-18180 also aligns to chr9_KI270823v1_alt

| ACTIONS | QUERY | SCORE | START | END | QSIZE | IDENTITY | CHROM | STRAND | START | END | SPAN |
|---|---|---|---|---|---|---|---|---|---|---|---|
| browser new tab details | YourSeq | 13038 | 1 | 13108 | 13108 | 99.8% | chr9_KI270823v1_alt | - | 323387 | 336500 | 13114 |
| browser new tab details | YourSeq | 13038 | 1 | 13108 | 13108 | 99.8% | chr9 | - | 101272184 | 101285297 | 13114 |

➢ It's an *Alu*Ya5 insertion.

### HG03540 chr13:89749560

- Using the built-in BLAT tool in IGV to check where the inserted sequence aligned.

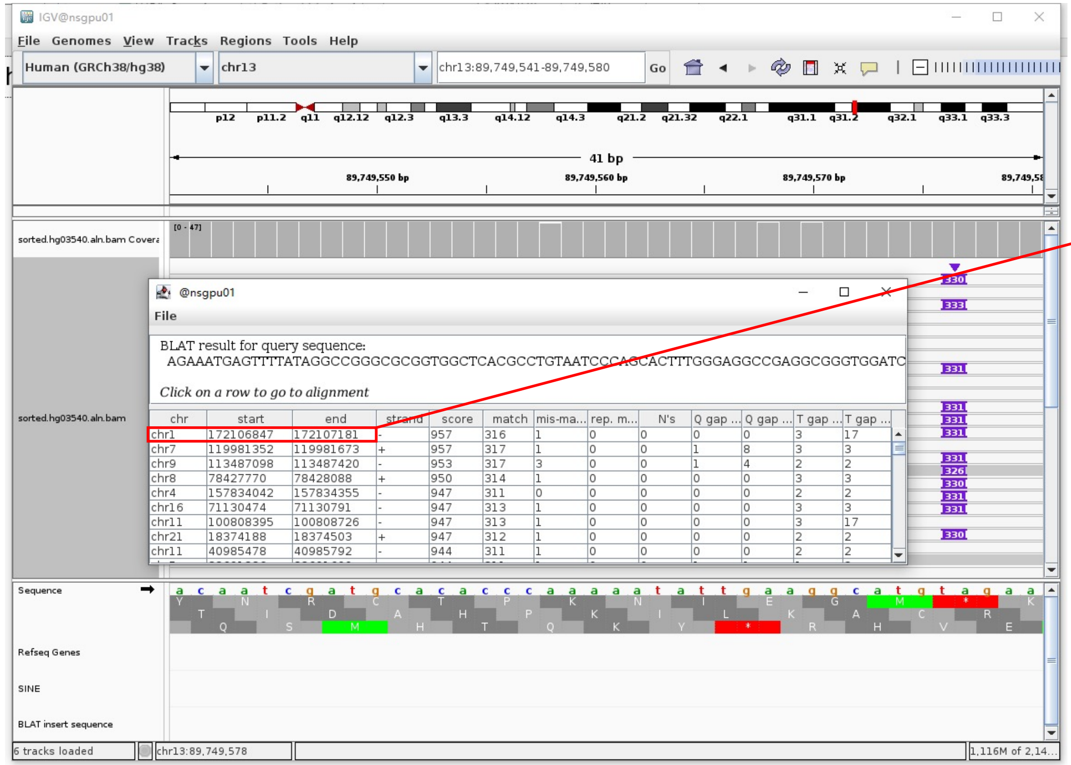


- The annotation at the aligned location indicated it as *AluY*.

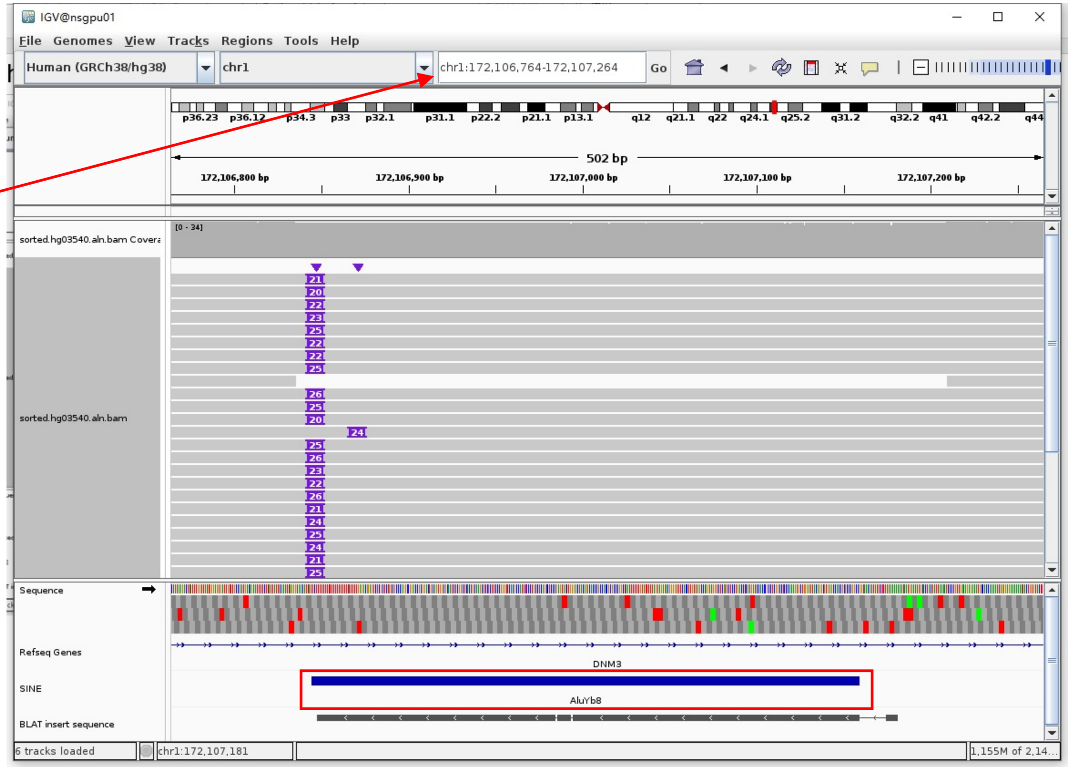


Insertion sequence (330bp)

- 16bp TSD

AGAAATGAGTTTTATA

- Insertion:16-303 aligns to *Alu*Yb8:1-289, 99.65% identity

GGCCGGGCGCGGTGGCTCACGCCTGTAATCCCAGCACTTTGGGAGGCCGAGGCGGGTGGATCATGAGGTCAGGAGATCGAGACCATCCTGGCTAACAAGGTGAAACCCCGTCTCTACTAAAAA
TACAAAAAATTAGCCGGGCGCGGTGGCGGGTGCCTGTAGTCCAGCTACTCGGAGGCTGAGGCAGGAGAATGGCGTGAACCCGGGAAGCGGAGCTTGCAGTGAGCCGAGATTGCGCCACTGCAG
TCCGCAGTCCGGCCTGGGCGACAGAGCGAGACTCCGTCTCA.........poly(A)

### HG03540 chr14:91780893

➢ The same genomic coordinate as in HG02630 (page 35)

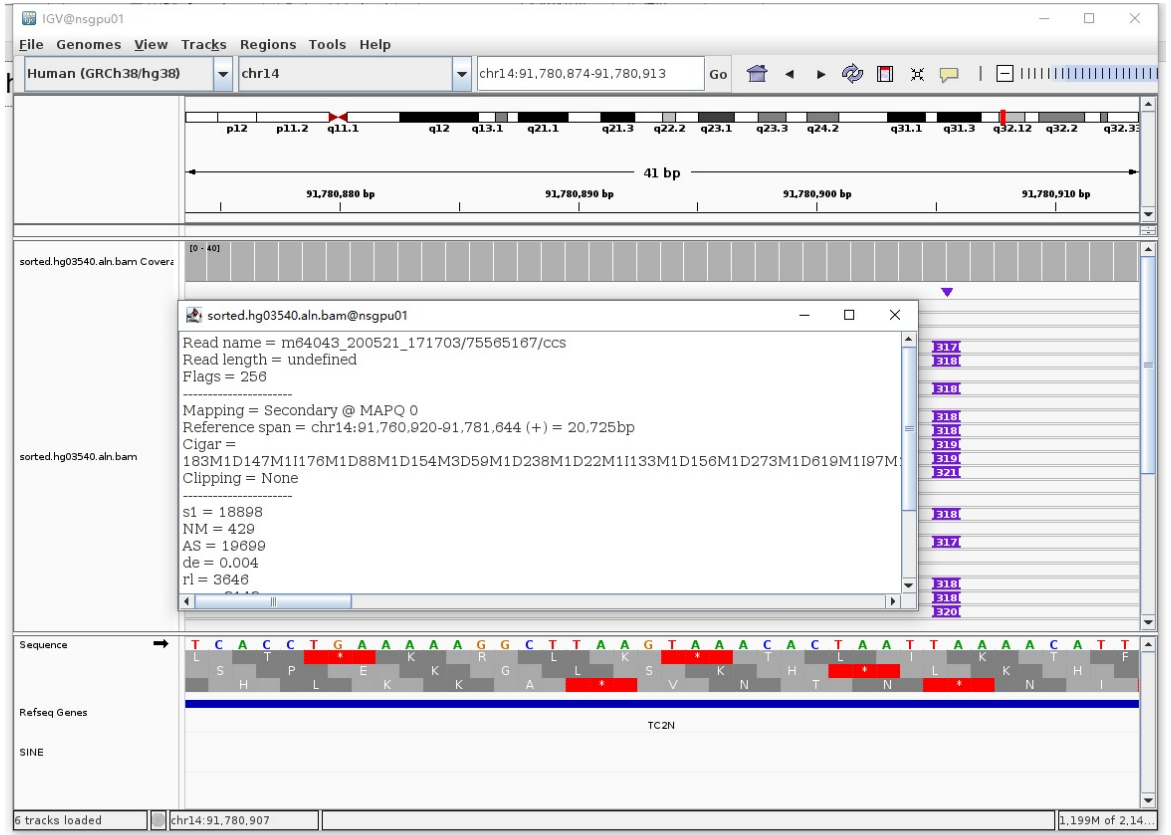

### HG03540 chr14:93364571

- ➢ The same genomic coordinate as in HG00438 (page 7)

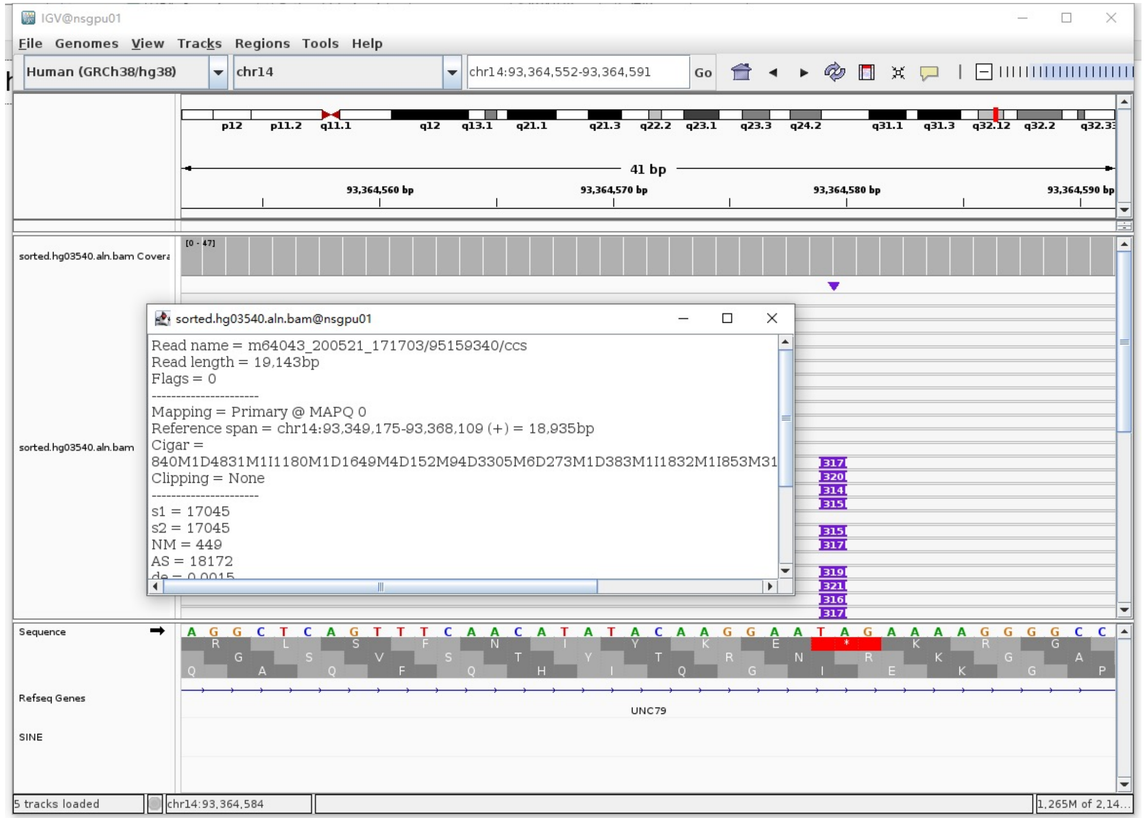

### HG03540 chr14:93400345

- ➢ Mapping quality is 0, indicating that this region may have alignment issues.

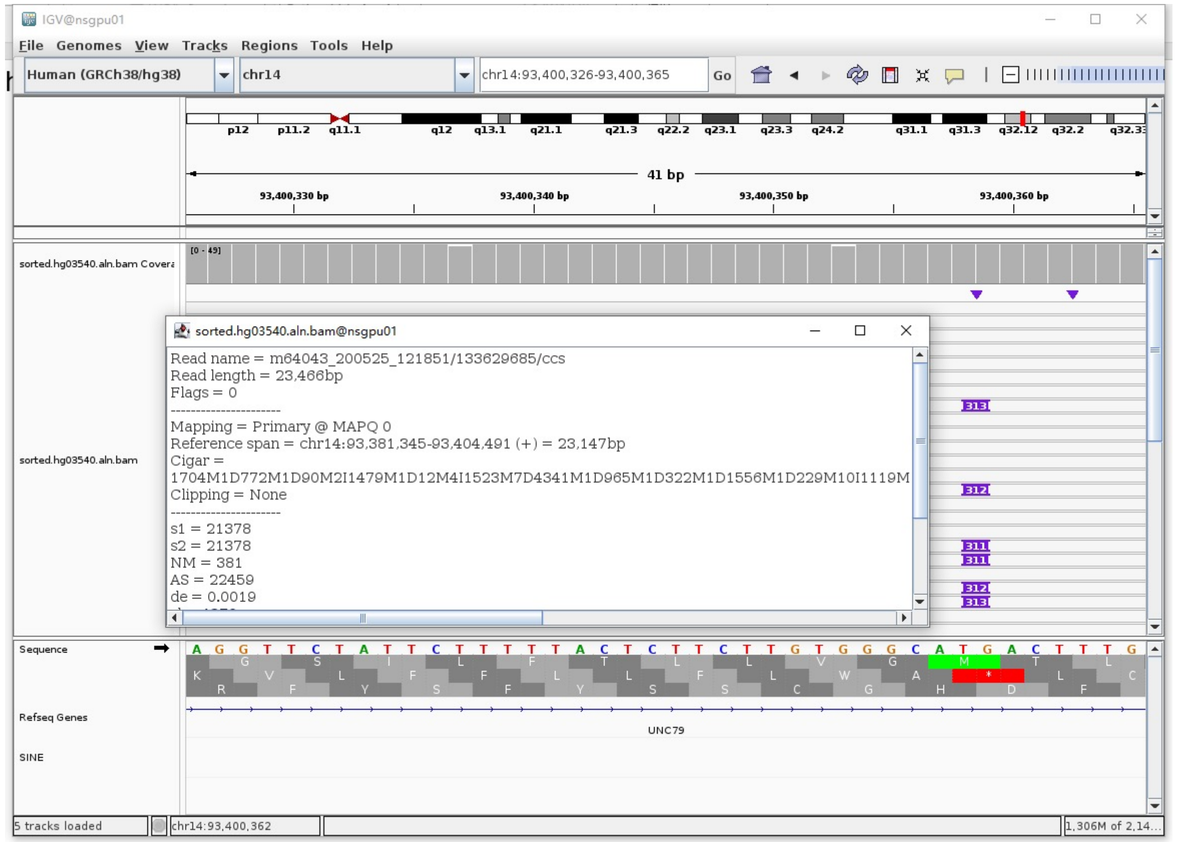


- ➢ Fetch one read (m64043_200525_121851/133629685/ccs, 23477 bp) and BLAT it to the hg38 genome.

  The best alignment is to chr14_KI270847v1_alt (an alternative sequence of chr14) and chr14.

| ACTIONS | | | QUERY | SCORE | START | END | QSIZE | IDENTITY | CHROM | STRAND | START | END | SPAN |
|---|---|---|---|---|---|---|---|---|---|---|---|---|---|
| browser | new tab | details | YourSeq | 23080 | 1 | 23477 | 23477 | 99.9% | chr14_KI270847v1_alt | + | 495907 | 519053 | 23147 |
| browser | new tab | details | YourSeq | 23079 | 1 | 23477 | 23477 | 99.9% | chr14 | + | 93381345 | 93404491 | 23147 |

- ● Read:1-19060 aligns to chr14_KI270847v1_alt

| ACTIONS | | | QUERY | SCORE | START | END | QSIZE | IDENTITY | CHROM | STRAND | START | END | SPAN |
|---|---|---|---|---|---|---|---|---|---|---|---|---|---|
| browser | new tab | details | YourSeq | 18966 | 1 | 19039 | 19059 | 99.9% | chr14_KI270847v1_alt | + | 495908 | 514931 | 19024 |
| browser | new tab | details | YourSeq | 18965 | 1 | 19039 | 19059 | 99.9% | chr14 | + | 93381346 | 93400369 | 19024 |

- ● Read:19061-19342 aligns to *Alu*Ya5:282-1, 98.93% identity

TGAGACGGAGTCTCACTCTGTCGCCCAGGCTGGAGTGCAGTGGCGGGATCTCGGCTCACTGCAAGCTCCGCCTC
CCGGGTTCACGCCATTCTCCTGCCTCAGCCTCCCGAGTAGCTGGGACTACAGGCGCCCGCCACTACGCCCGGCT
AATTTTTTGTATTTTCAGTAGAGACGGGGTTTCACCGTTTTAGCCGGGATGGTCTCGATCTCCTGACCTCGTGA
TCCGCCCGCCTCGGCCTCCCAAAGTGCTGGGATTACAGGCGTGAGCCACCGCGCCCGGC

11bp TSD

GACTTTGTTTT

- ● Read:19342-23477 also aligns to chr14_KI270847v1_alt

| ACTIONS | | | QUERY | SCORE | START | END | QSIZE | IDENTITY | CHROM | STRAND | START | END | SPAN |
|---|---|---|---|---|---|---|---|---|---|---|---|---|---|
| browser | new tab | details | YourSeq | 4125 | 1 | 4135 | 4135 | 99.9% | chr14_KI270847v1_alt | + | 514921 | 519053 | 4133 |
| browser | new tab | details | YourSeq | 4125 | 1 | 4135 | 4135 | 99.9% | chr14 | + | 93400359 | 93404491 | 4133 |

- ➢ It's an *Alu*Ya5 insertion.

### HG03540 chr17:37561902

- ➤ The same genomic coordinate as in HG00621 (page 15)

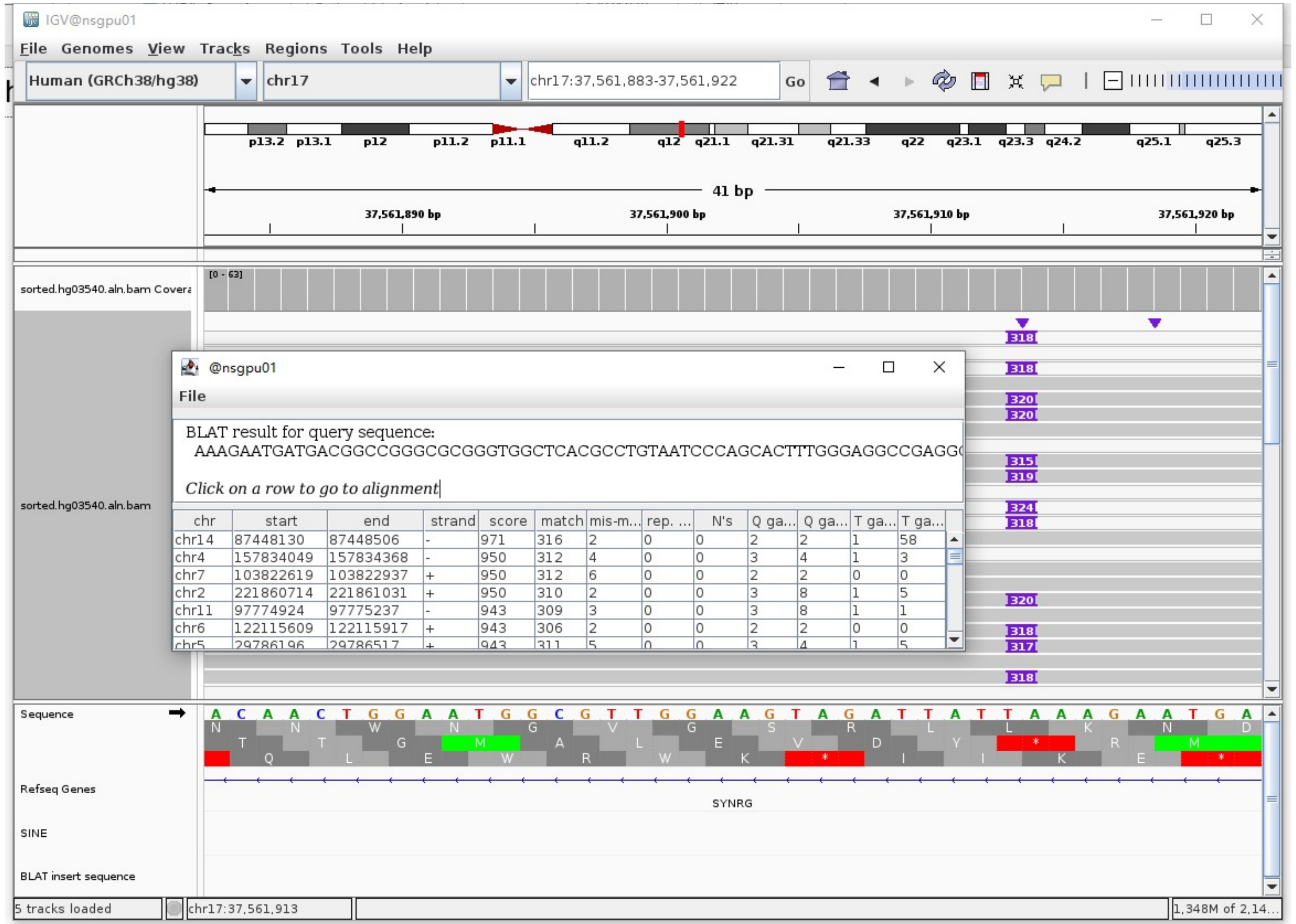

### HG03540 chr17:70457036

➢ Mapping quality is 0, indicating that this region may have alignment issues.

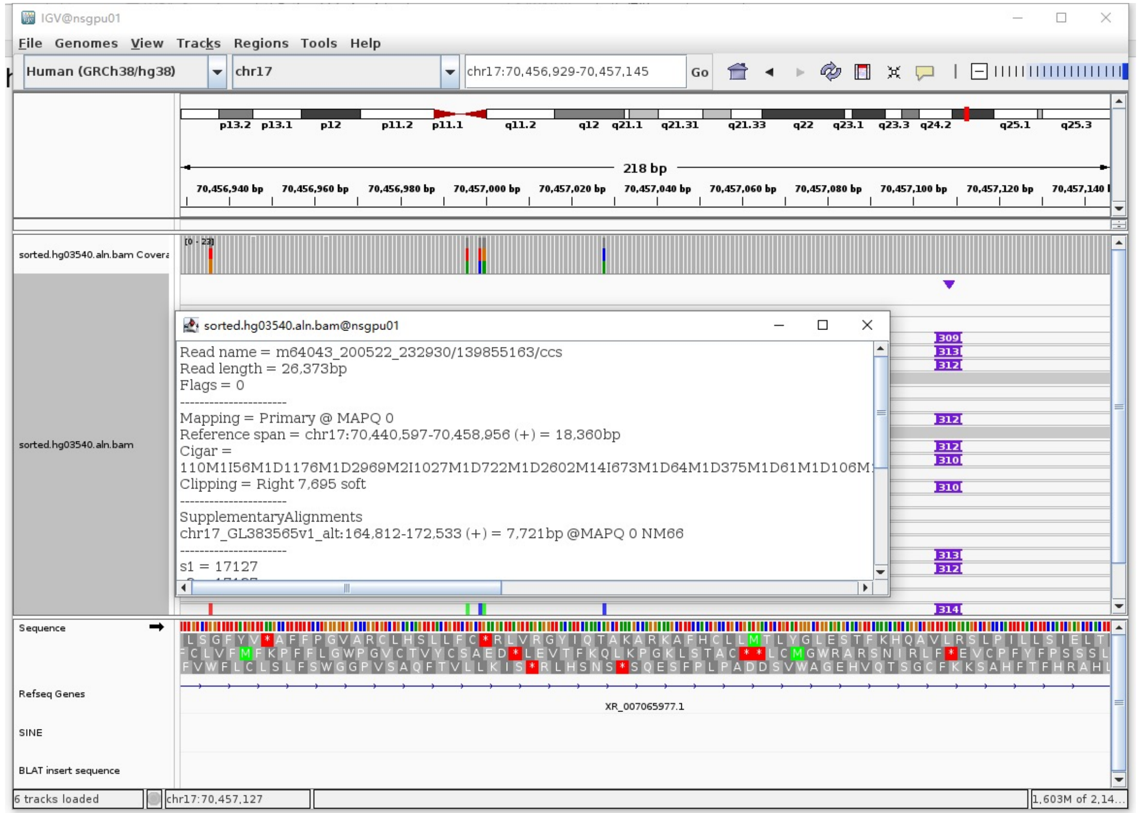


➢ Fetch one read (m64043_200521_171703/145294648/ccs, 19408 bp) and BLAT it to the hg38 genome.

The best alignment is to chr17_GL383565v1_alt (an alternative sequence of chr17) and chr17.

| ACTIONS | QUERY | SCORE | START | END | QSIZE | IDENTITY | CHROM | STRAND | START | END | SPAN |
|---|---|---|---|---|---|---|---|---|---|---|---|
| browser new tab details | YourSeq | 18914 | 1 | 19408 | 19408 | 99.6% | chr17_GL383565v1_alt | + | 138999 | 158089 | 19091 |
| browser new tab details | YourSeq | 18914 | 1 | 19408 | 19408 | 99.6% | chr17 | + | 70439223 | 70458313 | 19091 |

- Read:1-17905 aligns to chr17_GL383565v1_alt

| ACTIONS | QUERY | SCORE | START | END | QSIZE | IDENTITY | CHROM | STRAND | START | END | SPAN |
|---|---|---|---|---|---|---|---|---|---|---|---|
| browser new tab details | YourSeq | 17741 | 1 | 17904 | 17904 | 99.7% | chr17_GL383565v1_alt | + | 139000 | 156896 | 17897 |
| browser new tab details | YourSeq | 17741 | 1 | 17904 | 17904 | 99.7% | chr17 | + | 70439224 | 70457120 | 17897 |

- Read:17906-18188 aligns to *Alu*Ya5:1-282, 99.65% identity

GGCCGGGCGCGGTGGCTCACGCCTGTAATCCCAGCACTTTGGGAGGCCGAGGCGGGCGGATCACGAGGTCAGGA
GATCGAGACCATCCCGGCTAAAACGGTGAAACCCCGTCTCTACTAAAAATACAAAAAATTAGCCGGGCGTAGTG
GCGGGCGCCTGTAGTCCCAGCTACTTGGGAGGCTGAGGCAGGAGAATGGCGTGAACCCGGGAGGCGCAGCTTGC
AGTGAGCCGAGATCCCGCCACTGCACTCCAGCCTGGGCGACAGAGCGAGACTCCGTCTCA

13bp TSD

AAGAAGTCTGCCC

- Read:18189-19408 also aligns to chr17_GL383565v1_alt

| ACTIONS | QUERY | SCORE | START | END | QSIZE | IDENTITY | CHROM | STRAND | START | END | SPAN |
|---|---|---|---|---|---|---|---|---|---|---|---|
| browser new tab details | YourSeq | 1185 | 16 | 1219 | 1219 | 99.3% | chr17_GL383565v1_alt | + | 156885 | 158089 | 1205 |
| browser new tab details | YourSeq | 1185 | 16 | 1219 | 1219 | 99.3% | chr17 | + | 70457109 | 70458313 | 1205 |

➢ It's an *Alu*Ya5 insertion.

### HG03540 chr18:76926745

➢ Using the built-in BLAT tool in IGV to check where the inserted sequence aligned.

➢ The annotation at the aligned location indicated it as *AluY*.

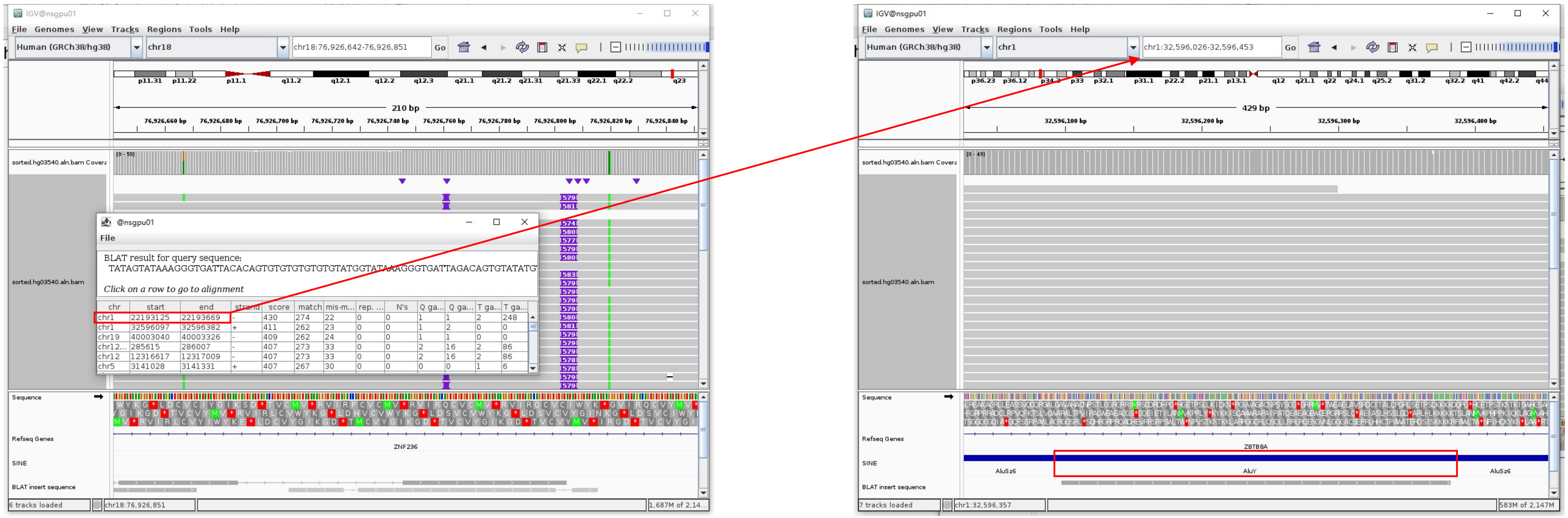


Insertion sequence (574bp)

➢ 16bp TSD

AGAAATGAGTTTTATA

➢ Insertion:1-224 aligns to chr18:76926553-76926791, 95.7% identity

➢ Insertion:225-505 aligns to *Alu*Yc1:1-282, 89.93% identity

GGCCGGGTGTGGTGGCTCATGCCTGTAATCCCAGCACTTTGGGAGGCCGAGATCGACGGATCACGAGGTCAGGAGATCGAGATCATCCTGGCTAACATGGTGAAACCCCGTCTCTACTAAAAA
TACAAAAAAAATTACCAGTGTGGTGGCTGAGCCTGTAGTCCAGCTACTCAGGAGGGTGAGGGAGGAGAATGGTGTGAACCTGGGAGGCGGAGCTTGCAGTGAGCTGAGATGGCACCACTGCAC
TCTAGTCTGGGCGACTGAGCGAGAGACTGTCTCAAAAAAAAAAAAAAGGGTGATTAGACAGTGTGTGTGTATGGTATAAAGGGTGATTAGACAGTATGTGTGTG

### HG03540 chr19:52384793

- ➢ The same genomic coordinate as in HG01106 (page 23)

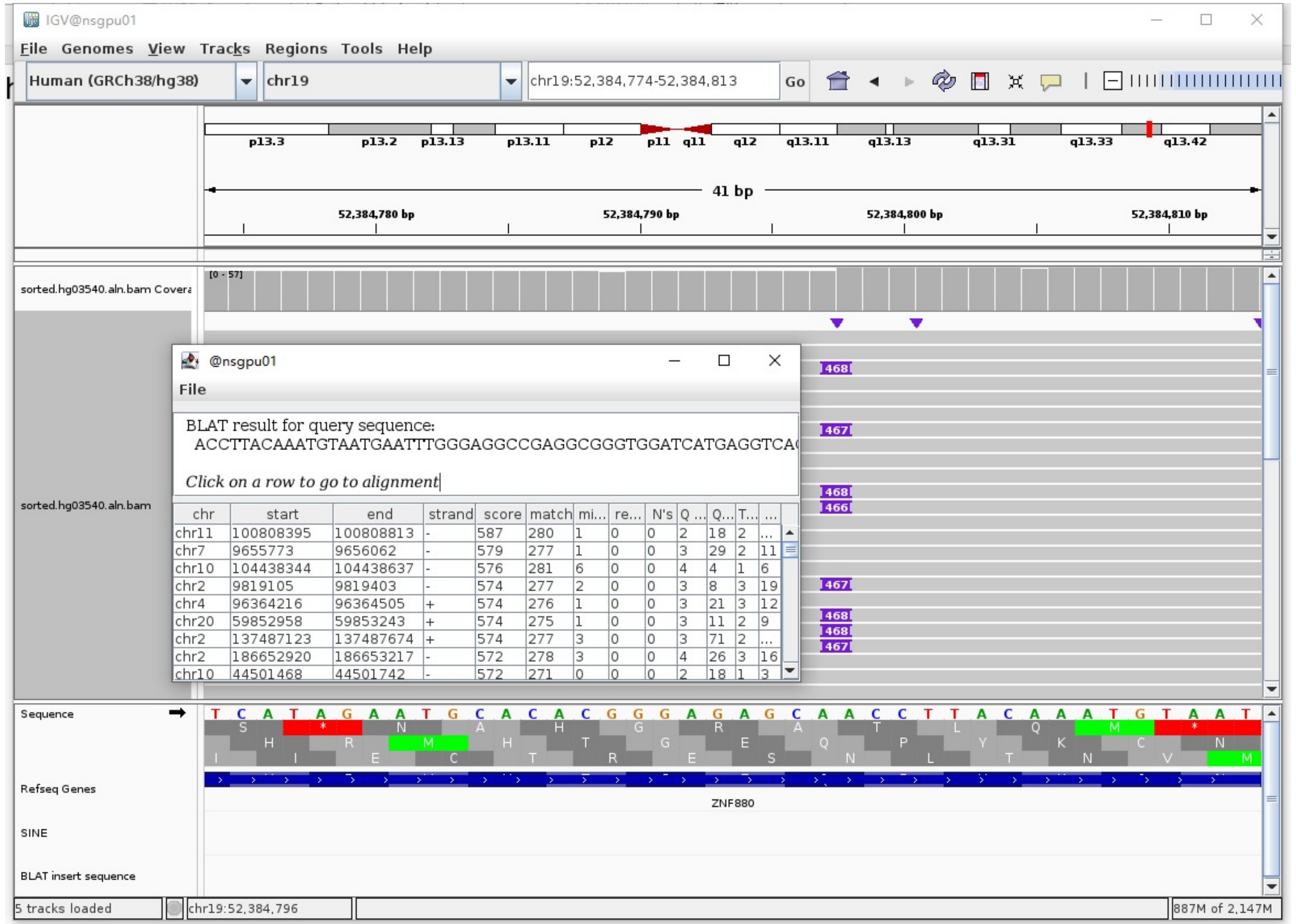

### HG03540 chr22:23928251

- Mapping quality is 0, indicating that this region may have alignment issues.

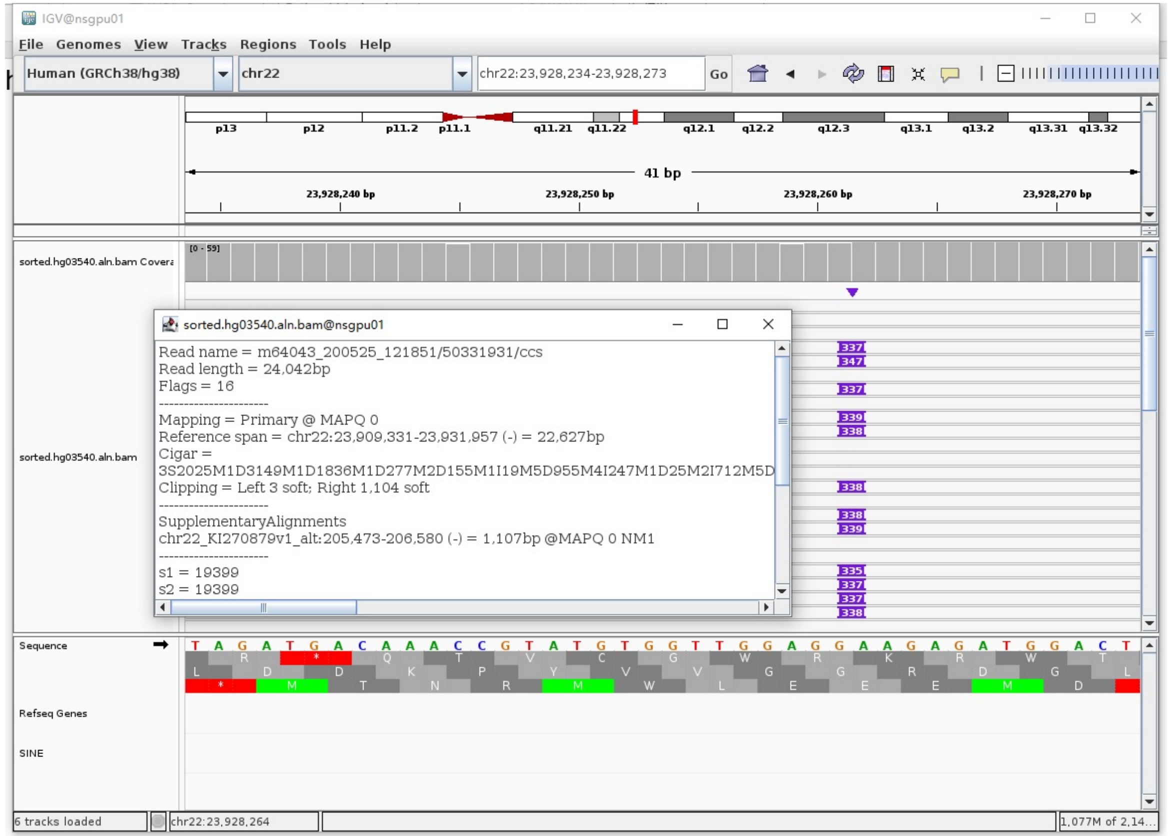


- Fetch one read (m64043_200524_055430/165545135/ccs, 20520 bp) and BLAT it to the hg38 genome.

  The best alignment is to chr22_KI270879v1_alt (an alternative sequence of chr22) and chr22.

| ACTIONS | | | QUERY | SCORE | START | END | QSIZE | IDENTITY | CHROM | STRAND | START | END | SPAN |
|---|---|---|---|---|---|---|---|---|---|---|---|---|---|
| browser | new tab | details | YourSeq | 20005 | 1 | 20520 | 20520 | 99.6% | chr22 | + | 23909944 | 23930140 | 20197 |
| browser | new tab | details | YourSeq | 19690 | 1 | 20520 | 20520 | 99.6% | chr22_KI270879v1_alt | + | 146306 | 166502 | 20197 |

- Read:1-18312 aligns to chr22_KI270879v1_alt and chr22

| ACTIONS | | | QUERY | SCORE | START | END | QSIZE | IDENTITY | CHROM | STRAND | START | END | SPAN |
|---|---|---|---|---|---|---|---|---|---|---|---|---|---|
| browser | new tab | details | YourSeq | 18159 | 1 | 18311 | 18311 | 99.7% | chr22 | + | 23909945 | 23928274 | 18330 |
| browser | new tab | details | YourSeq | 17844 | 1 | 18311 | 18311 | 99.7% | chr22_KI270879v1_alt | + | 146307 | 164636 | 18330 |

- Read:18313-18601 aligns to *Alu*Yb8:1-289, 100% identity

  GGCCGGGCGCGGTGGCTCACGCCTGTAATCCCAGCACTTTGGGAGGCCGAGGCGGGTGGATCATGAGGTCAGGA
  GATCGAGACCATCCTGGCTAACAAGGTGAAACCCCGTCTCTACTAAAAATACAAAAATTAGCCGGGCGCGGTGG
  CGGGCGCCTGTAGTCCCAGCTACTCGGGAGGCTGAGGCAGGAGAATGGCGTGAACCCGGGAAGCGGAGCTTGCA
  GTGAGCCGAGATTGCGCCACTGCAGTCCGCAGTCCGGCCTGGGCGACAGAGCGAGACTCCGTCTCA

  14bp TSD

  AAGAGATGGACTGA

- Read:18602-20520 also aligns to chr22_KI270879v1_alt and chr22

| ACTIONS | | | QUERY | SCORE | START | END | QSIZE | IDENTITY | CHROM | STRAND | START | END | SPAN |
|---|---|---|---|---|---|---|---|---|---|---|---|---|---|
| browser | new tab | details | YourSeq | 1859 | 37 | 1918 | 1918 | 99.3% | chr22_KI270879v1_alt | + | 164624 | 166502 | 1879 |
| browser | new tab | details | YourSeq | 1859 | 37 | 1918 | 1918 | 99.3% | chr22 | + | 23928262 | 23930140 | 1879 |

- It’s an *Alu*Yb8 insertion.

### NA18906 chr1:246243393

➢ Using the built-in BLAT tool in IGV to check where the inserted sequence aligned.

➢ The annotation at the aligned location indicated it as *AluY*.

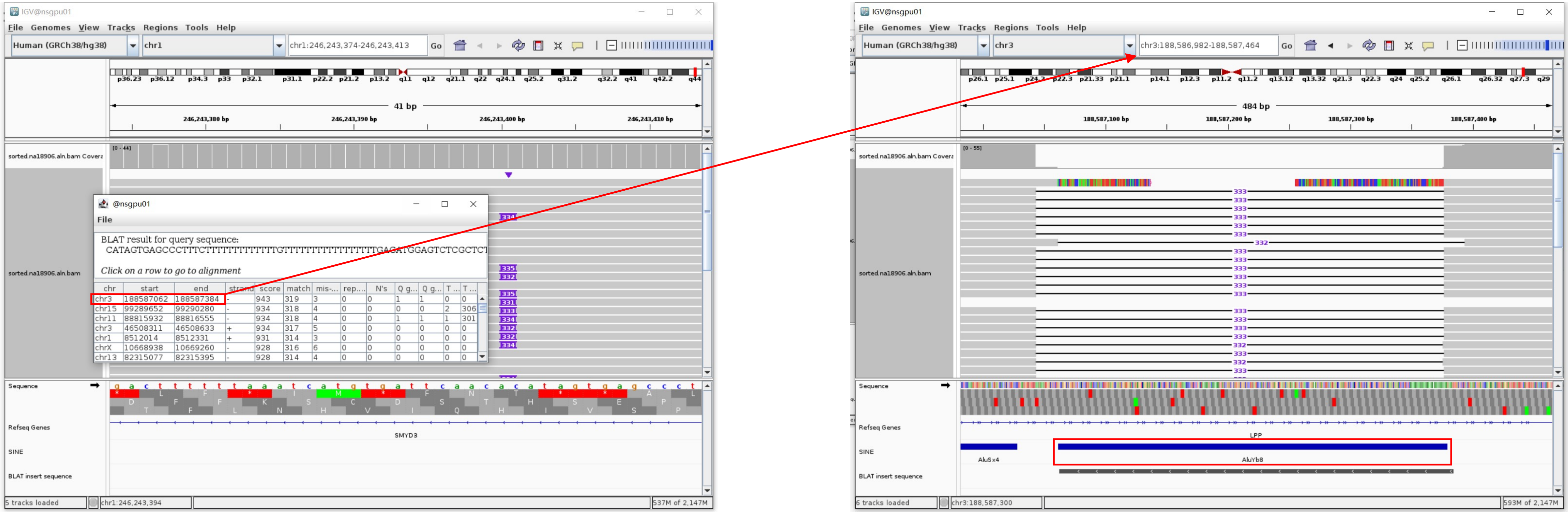


Insertion sequence (331bp)

➢ 17bp TSD

CATAGTGAGCCCTTTCT

➢ Insertion:43-330 aligns to *Alu*Yb8:289-1, 98.95% identity

TGAGATGGAGTCTCGCTCTGTCGCCCAGCCGGACTGCAGACTGCAGTGGTGCAATCTCGGCTCACTGCAAGCTCCGCTTCCCGGGTTCACGCCATTCTCCTGCCTCAGCCTCCCGAGTAGCTG
GGACTACAGGCGCCCGCCACCGCGCCCGGCTAATTTTTTGTATTTTTAGTAGAGACGGGGTTTCCACCTTGTTAGCCAGGATGGTCTCGATCTCCTGACCTCATGATCCACCCGCCTCGGCCT
CCCAAAGTGCTGGATTACAGGCGTGAGCCACCGCGCCCGGC

### NA18906 chr2:36249528

- ➢ Using the built-in BLAT tool in IGV to check where the inserted sequence aligned.

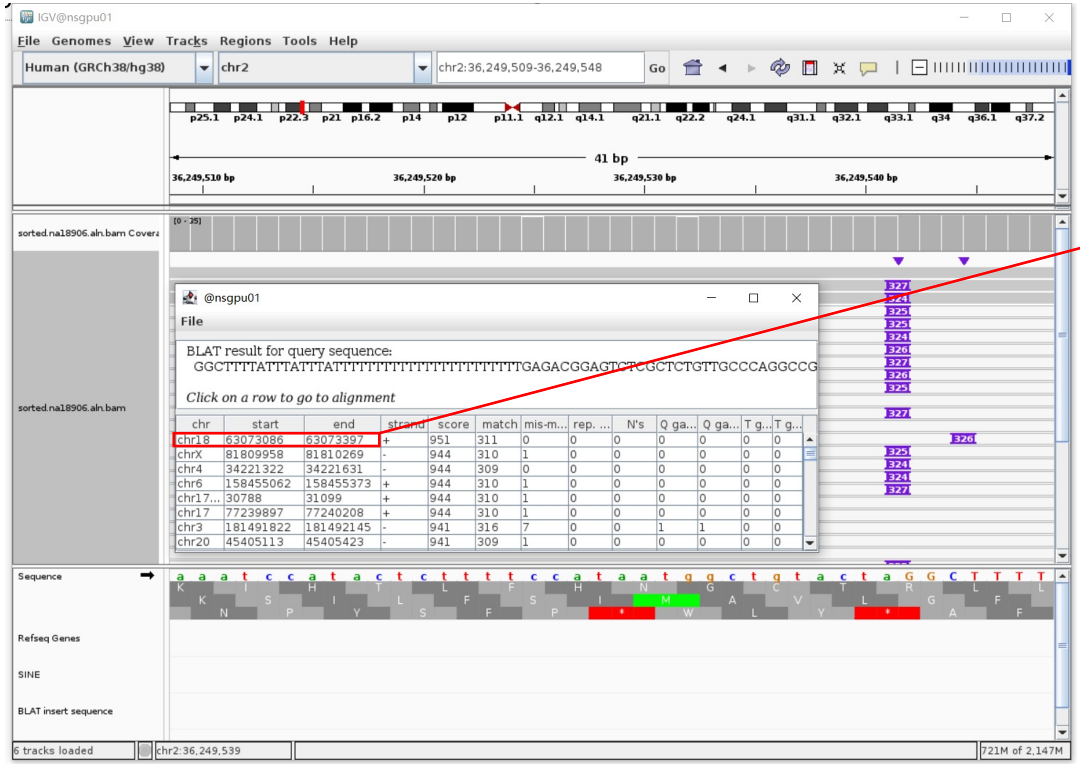


- ➢ The annotation at the aligned location indicated it as *AluY*.

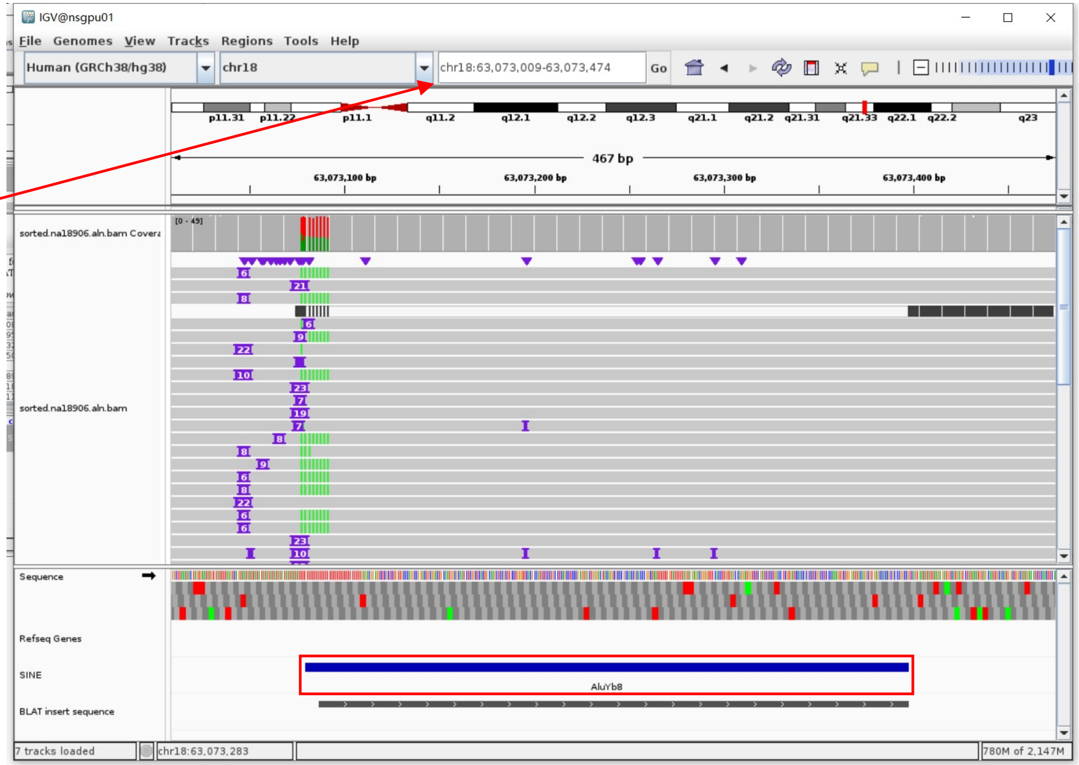


Insertion sequence (327bp)

- ➢ 11bp TSD

GGCTTTTATTT

- ➢ Insert:38-327 aligns to *AluYb8*:289-1, 98.27% identity

TGAGACGGAGTCTCGCTCTGTTGCCCAGGCCGGACTGCGGACTGCAATGGCGCAATCTCGGCTCACTGCAAGCTCCGCTTCCTGGGTTCACGCCATTCTCCTGCCTCAGCCTCCCGAGTAGCT
GGGACTACAGGCGCCCGCCACCGCACCCGGCTAATTTTTTGTATTTTTAGTAGAGACGGGGTTTCACCTTGTTAGCCAGGATGGTCTCGATCTCCTGACCTCGTGATCCACCCGCCTCGGCCT
CCCAAAGTGCTGGGATTACAGGCGTGAGCCACCGCGCCCGGCC

### NA18906 chr6:30030695

- The same genomic coordinate as in HG03492 (page 47)

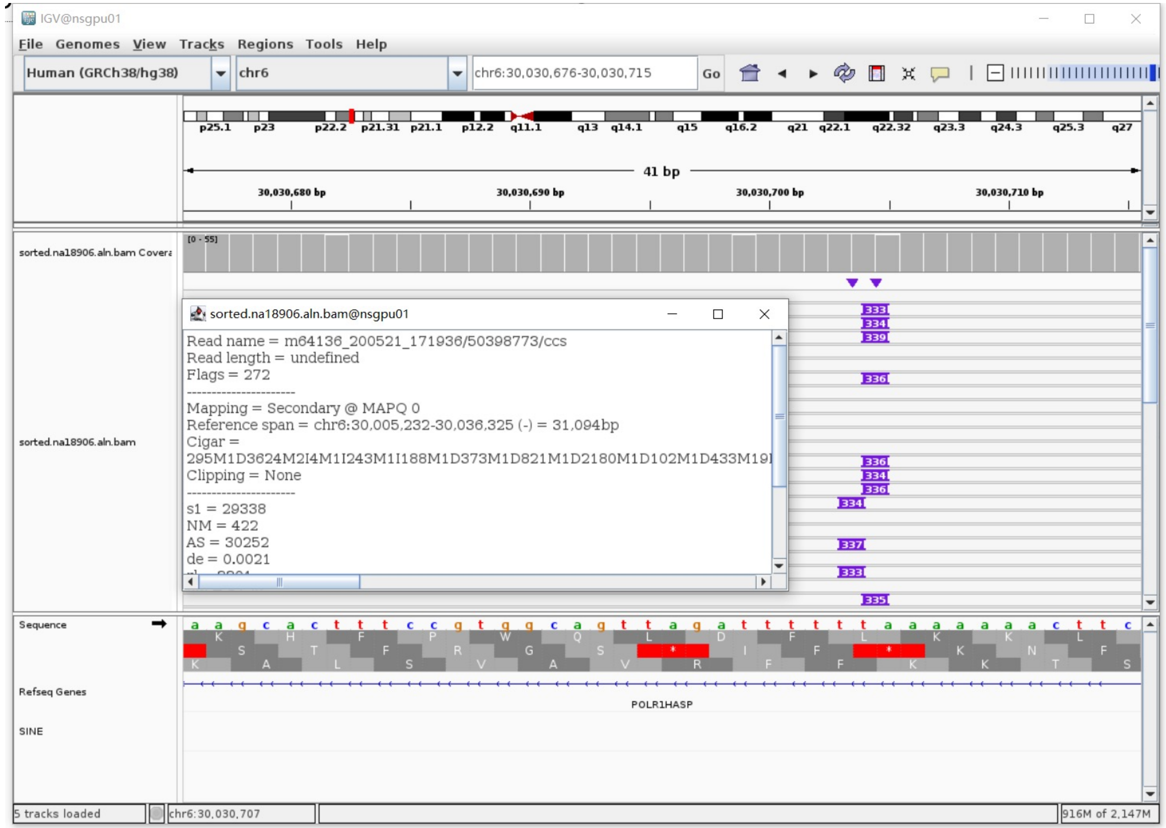

### NA18906 chr6:31471215

➢ The same genomic coordinate as in HG00438 (page 4)

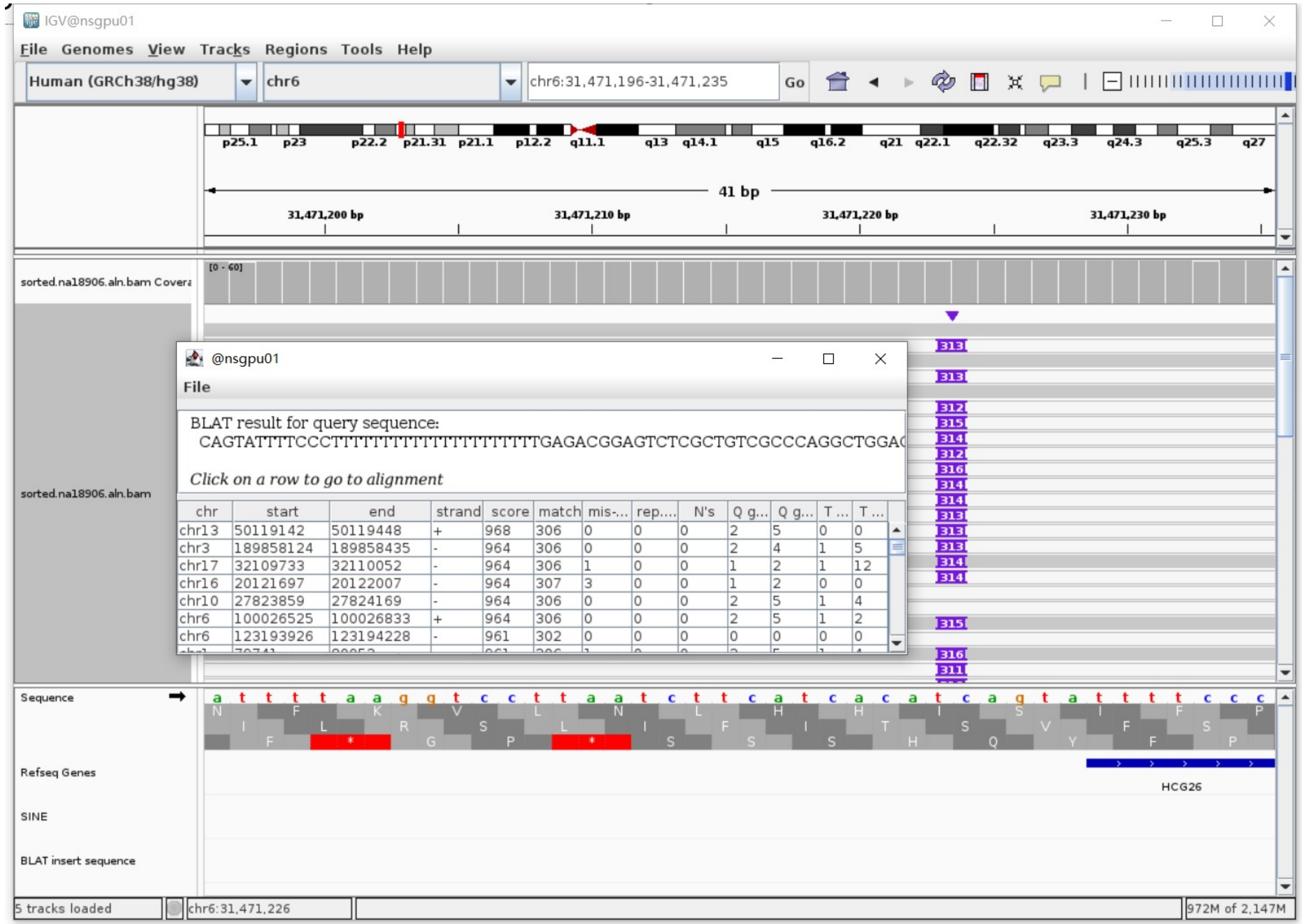

### NA18906 chr6:32719367

➢ Mapping quality is 0, indicating that this region may have alignment issues.

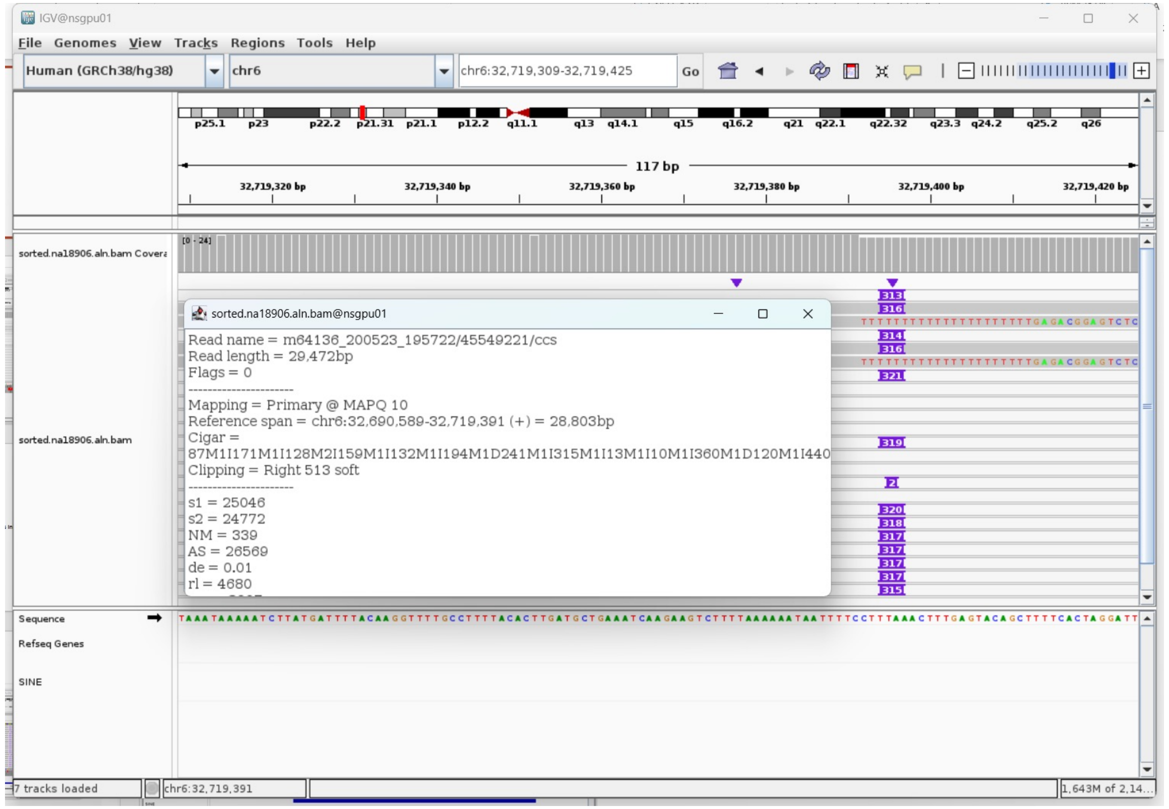


➢ Fetch one read (m64136_200523_195722/168559050/ccs, 24495 bp) and BLAT it to the hg38 genome.

The best alignment is to chr6_GL000254v2_alt (an alternative sequence of chr6).

| ACTIONS | | | QUERY | SCORE | START | END | QSIZE | IDENTITY | CHROM | STRAND | START | END | SPAN |
|---|---|---|---|---|---|---|---|---|---|---|---|---|---|
| browser | new tab | details | YourSeq | 23487 | 1 | 24495 | 24495 | 99.0% | chr6_GL000254v2_alt | + | 4000447 | 4024438 | 23992 |
| browser | new tab | details | YourSeq | 23446 | 1 | 24495 | 24495 | 99.2% | chr6_GL000251v2_alt | + | 4114515 | 4138341 | 23827 |
| browser | new tab | details | YourSeq | 23439 | 1 | 24495 | 24495 | 99.2% | chr6_GL000255v2_alt | + | 3895479 | 3919310 | 23832 |
| browser | new tab | details | YourSeq | 23429 | 1 | 24495 | 24495 | 98.9% | chr6_GL000252v2_alt | + | 3945812 | 3969819 | 24008 |
| browser | new tab | details | YourSeq | 23170 | 1 | 24495 | 24495 | 98.8% | chr6 | + | 32701196 | 32725051 | 23856 |

- Read:1-18203

| ACTIONS | | | QUERY | SCORE | START | END | QSIZE | IDENTITY | CHROM | STRAND | START | END | SPAN |
|---|---|---|---|---|---|---|---|---|---|---|---|---|---|
| browser | new tab | details | YourSeq | 18008 | 1 | 18185 | 18202 | 99.6% | chr6 | + | 32701197 | 32719391 | 18195 |
| browser | new tab | details | YourSeq | 17968 | 1 | 18189 | 18202 | 99.5% | chr6_GL000251v2_alt | + | 4114516 | 4132675 | 18160 |
| browser | new tab | details | YourSeq | 17926 | 1 | 18185 | 18202 | 99.4% | chr6_GL000255v2_alt | + | 3895480 | 3913643 | 18164 |
| browser | new tab | details | YourSeq | 17203 | 1 | 18202 | 18202 | 98.7% | chr6_GL000254v2_alt | + | 4000448 | 4018146 | 17699 |
| browser | new tab | details | YourSeq | 17142 | 1 | 18202 | 18202 | 98.5% | chr6_GL000252v2_alt | + | 3945813 | 3963518 | 17706 |
| browser | new tab | details | YourSeq | 16929 | 1 | 18185 | 18202 | 98.1% | chr6_GL000256v2_alt | + | 4101817 | 4119534 | 17718 |
| browser | new tab | details | YourSeq | 16477 | 1 | 18185 | 18202 | 98.2% | chr6_GL000253v2_alt | + | 4121024 | 4138694 | 17671 |

- Read:18204-18486 aligns to *Alu*Yc1:282-1, 98.58% identity

TGAGACGGAGTCTCGCTCTGTCGCCCAGGCTGGAGTGCAGTGGCGCGATCTCGGCTCACTGCAAGCTCCACCTC
CCGGGTTCACACCATTCTCCTGCCTCAGCCTCCCGAGTAGCTGGAACTACAGGTGCCCGCTACCACGCCCGGCT
AATTTTTTGTATTTTTAGTAGAGACGGGGTTTCACCGTGTTAGCCAGGATGGTCTCGATCTCCTGACCTCGTGA
TCCGCCCGCCTCGGCCTCCCAAAGTGCTGGGATTACAGGCGTGAGCCACCGCGCCCGGCC

14bp TSD

AAAAAATAATTTTCTTTT

- Read:18487-24495

| ACTIONS | | | QUERY | SCORE | START | END | QSIZE | IDENTITY | CHROM | STRAND | START | END | SPAN |
|---|---|---|---|---|---|---|---|---|---|---|---|---|---|
| browser | new tab | details | YourSeq | 6003 | 1 | 6008 | 6008 | 100.0% | chr6_GL000252v2_alt | + | 3963804 | 3969819 | 6016 |
| browser | new tab | details | YourSeq | 5999 | 1 | 6008 | 6008 | 100.0% | chr6_GL000254v2_alt | + | 4018431 | 4024438 | 6008 |
| browser | new tab | details | YourSeq | 5527 | 1 | 6008 | 6008 | 98.6% | chr6_GL000255v2_alt | + | 3913630 | 3919310 | 5681 |
| browser | new tab | details | YourSeq | 5527 | 1 | 6008 | 6008 | 98.6% | chr6_GL000253v2_alt | + | 4138681 | 4144361 | 5681 |
| browser | new tab | details | YourSeq | 5496 | 1 | 6008 | 6008 | 98.3% | chr6_GL000250v2_alt | + | 4025226 | 4030907 | 5682 |
| browser | new tab | details | YourSeq | 5495 | 1 | 6008 | 6008 | 98.3% | chr6_GL000251v2_alt | + | 4132658 | 4138341 | 5684 |
| browser | new tab | details | YourSeq | 5175 | 1 | 6008 | 6008 | 96.0% | chr6_GL000256v2_alt | + | 4119521 | 4125198 | 5678 |
| browser | new tab | details | YourSeq | 5175 | 1 | 6008 | 6008 | 96.0% | chr6 | + | 32719378 | 32725051 | 5674 |

➢ It's an *Alu*Yc1 insertion.

### NA18906 chr6:32805641

➢ The same genomic coordinate as in HG00438 (page 5)

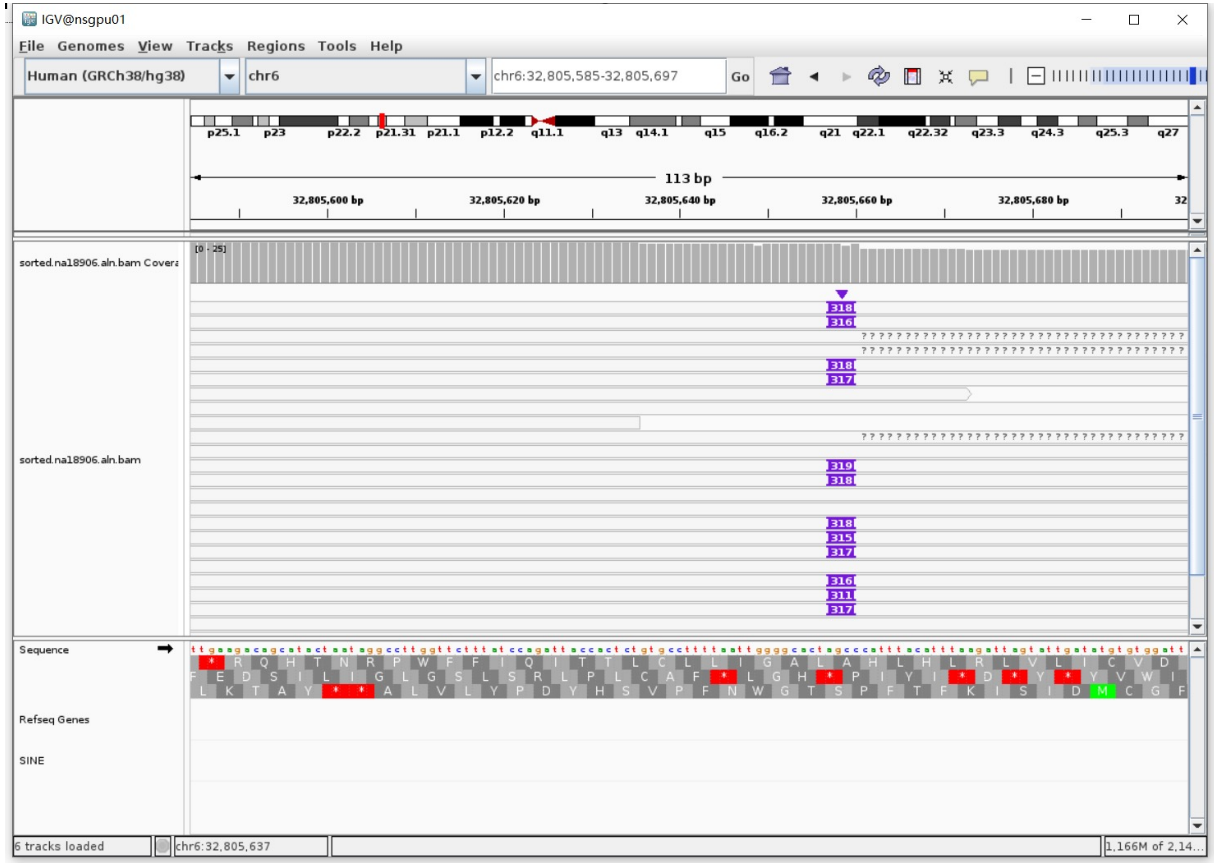

### NA18906 chr7:155318862

- ➢ The same genomic coordinate as in HG02630 (page 31)

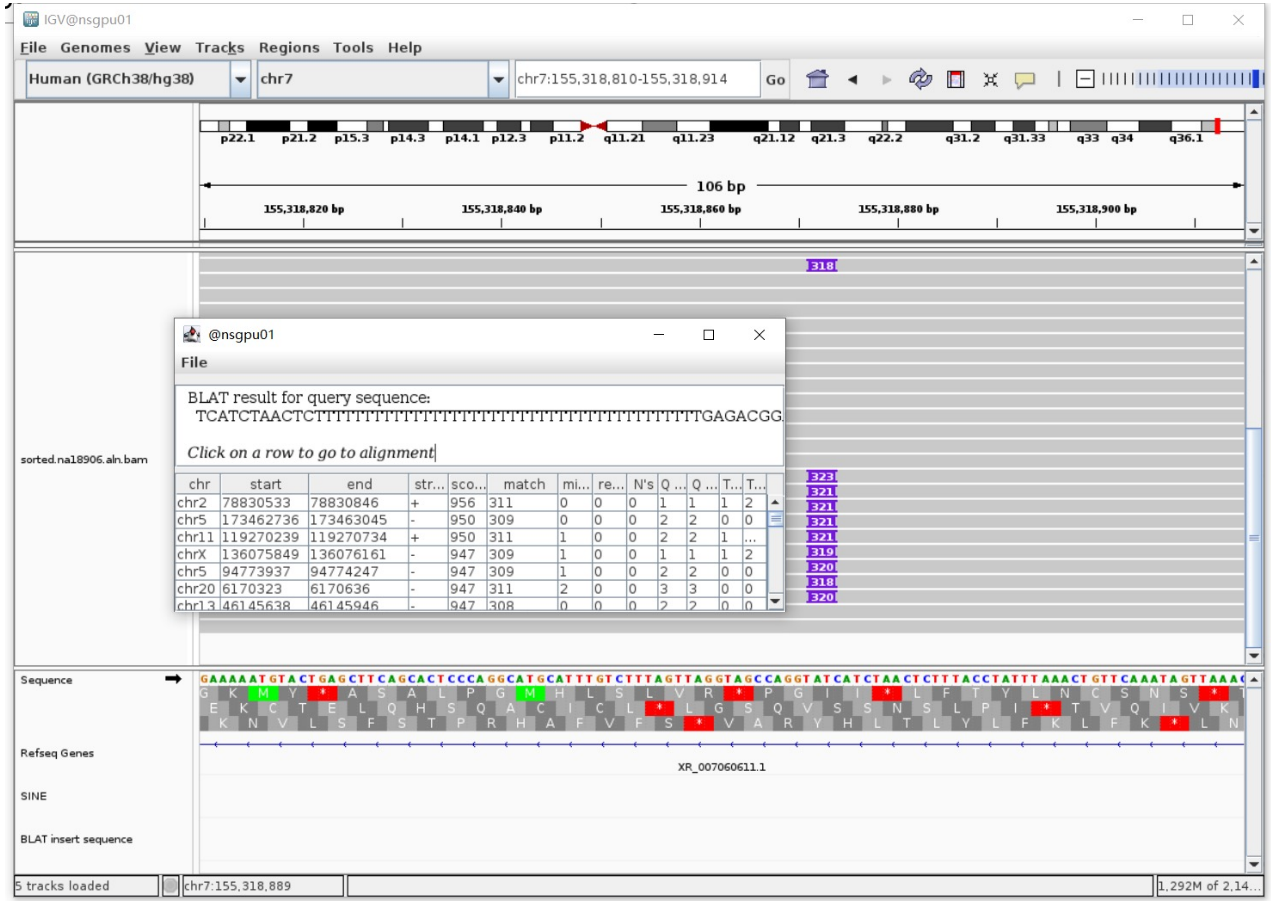

### NA18906 chr8:1362348

- Using the built-in BLAT tool in IGV to check where the inserted sequence aligned.

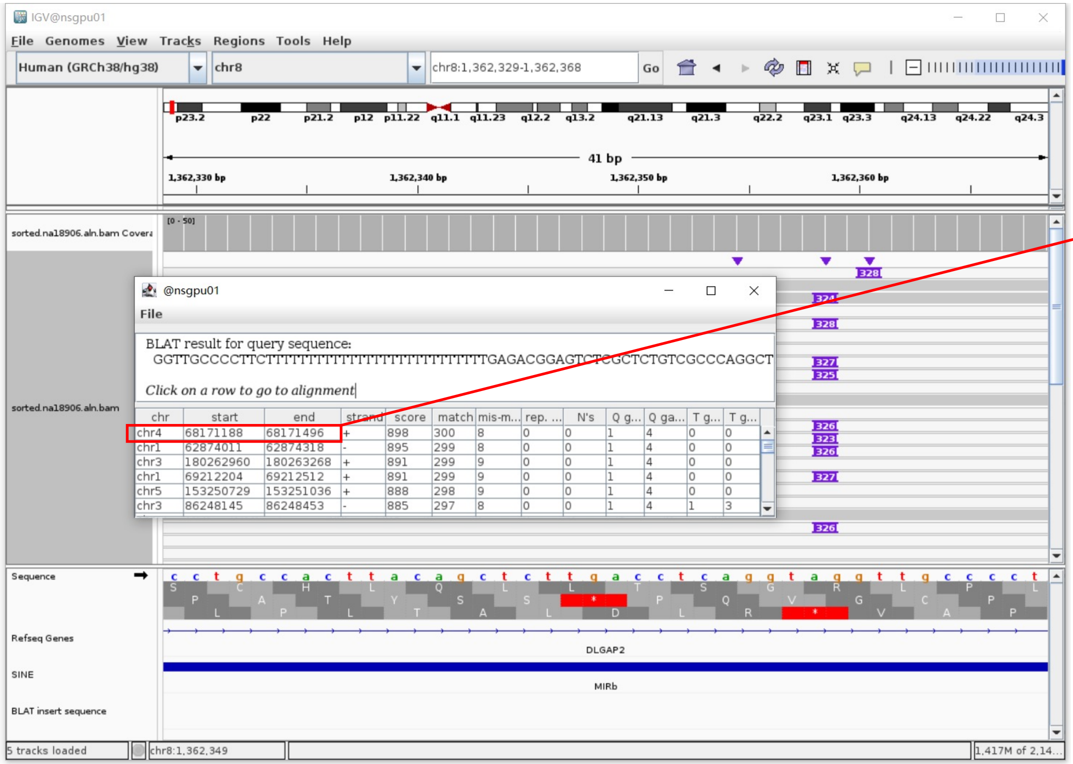


- The annotation at the aligned location indicated it as *AluY*.

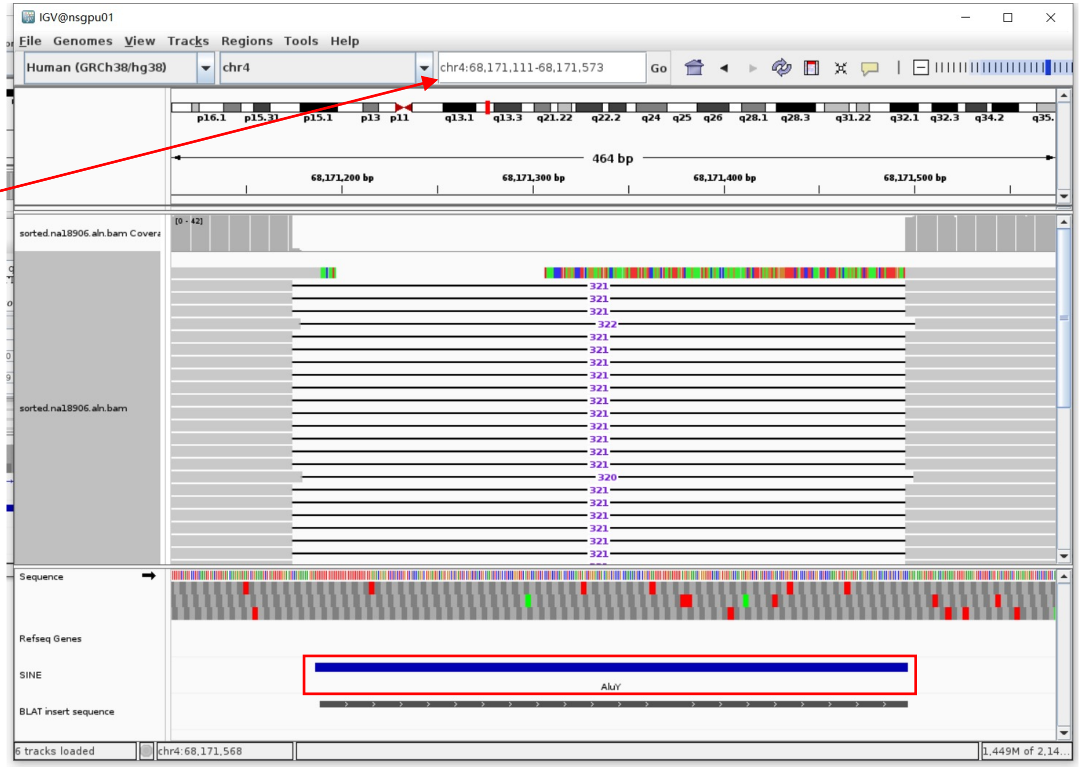


Insertion sequence (326bp)

- 14bp TSD

GGTTGCCCCTTCTT

- Insertion:40-326 aligns to *Alu*Yc1:282-1, 99.29% identity

TGAGACGGAGTCTCGCTCTGTCGCCCAGGCTGGAGTGCAGTGGCGCGATCTCGGCTCACTGCAAGCTCCGCCTCCCGGGTTCACGCCATTCTCCTGCCTCAGCCTCCGAGTAGCTGGGACTAC
AGGCGCCCGCCAACACGCCCGGCTAATTTTTTTTTGTATTTTTAGTAGAGACGGGGTTTCACCGTGTTAGCCAGGATGGTCTCGATCTCCTGACCTCGTGATCCGCCCGCCTCGGCCTCCCA
AAGTGCTGGGATTACAGGCGTGAGCCACCGCGCCCCGGCC

### NA18906 chr12:57965263

- Using the built-in BLAT tool in IGV to check where the inserted sequence aligned.

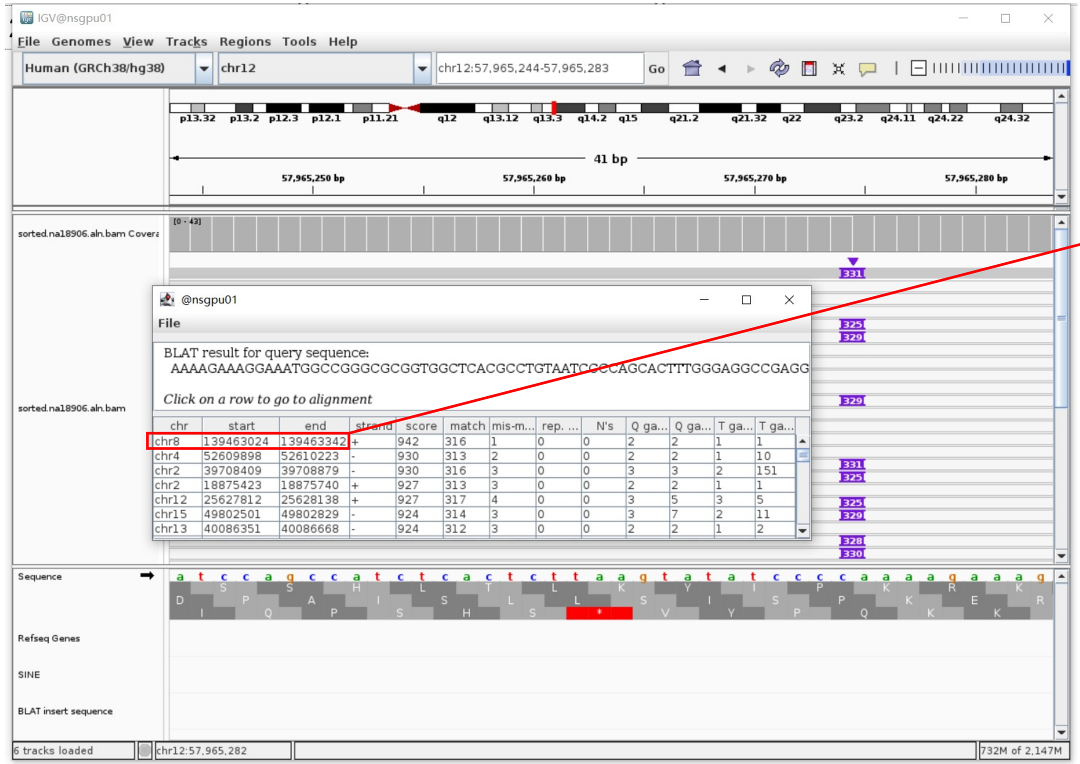


- The annotation at the aligned location indicated it as *AluY*.

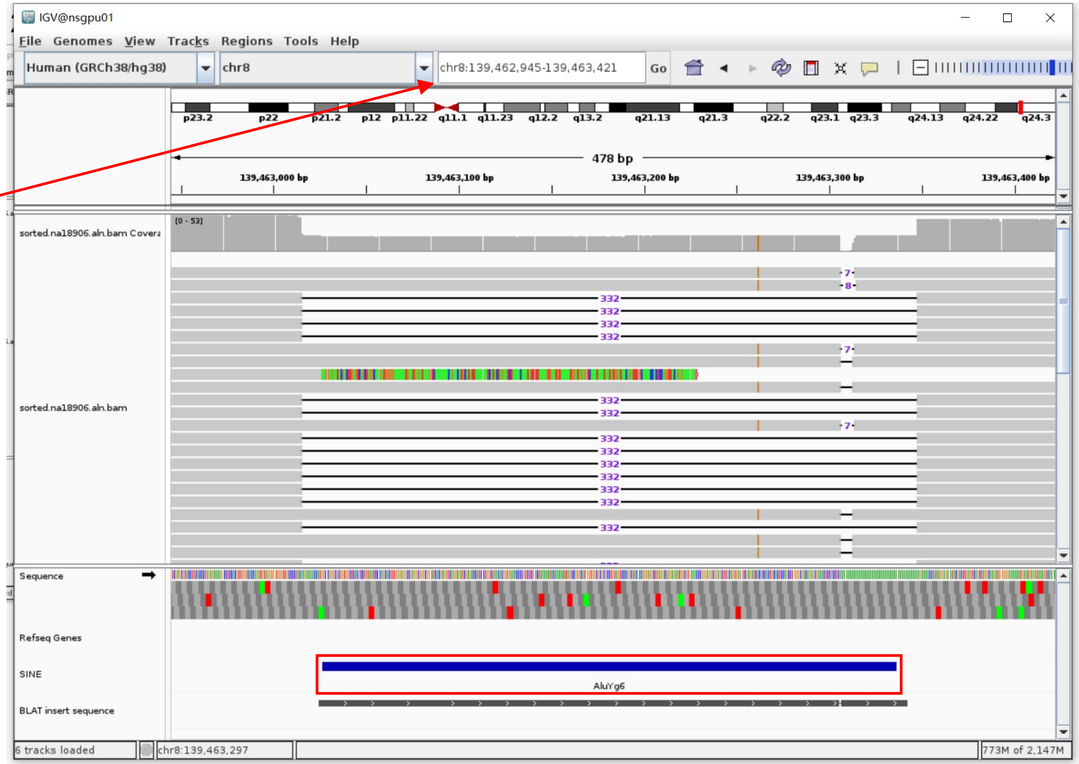


Insertion sequence (325bp)

- 14bp TSD

AAAAGAAAGGAAAT

- Insertion:14-297 aligns to *Alu*Yc1:1-282, 96.8% identity

GGCCGGGCGCGGTGGCTCACGCCTGTAATCCCAGCACTTTGGGAGGCCGAGGCGGGCGGATCACGAGGTCAGGAGATCGAGACCATCCTGGCTAACACGGTGAAACCCCGTCTCTACTAAA
AATACAAAAATTAGCCGGGCATGGTGGCGCGCGCCTGTAGTCCCAGCTACACGGGAGGCTGAGGCAGGAGAATGGCGTAAACCCGGGAGGCGGAGCTTGCAGTGAGTCGAGATCGCGCCACTG
CACTCCAGCCTGGGCGACAGAGCGAAACTCCGCCTA………poly(A)

### NA18906 chr12:58064417

- The same genomic coordinate as in HG02630 (page 33)

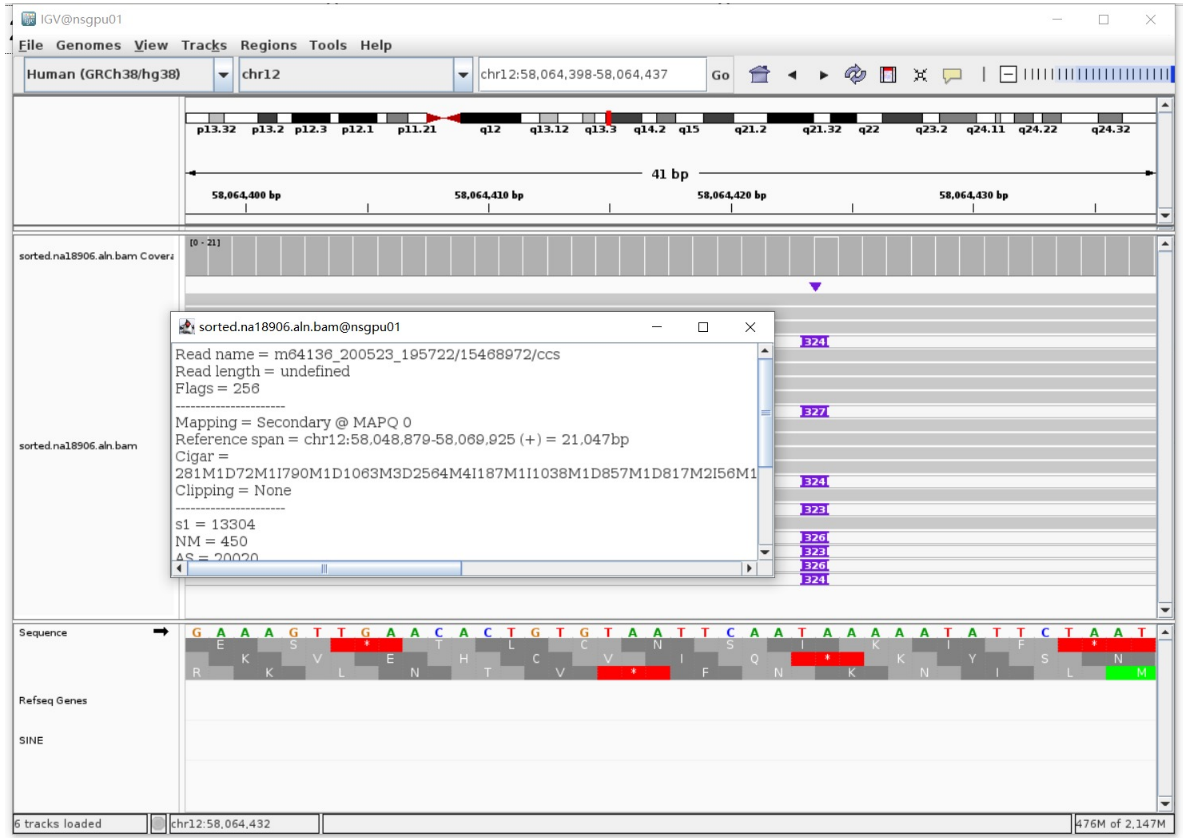

### NA18906 chr12:58987593

➢ Mapping quality is 0, indicating that this region may have alignment issues.

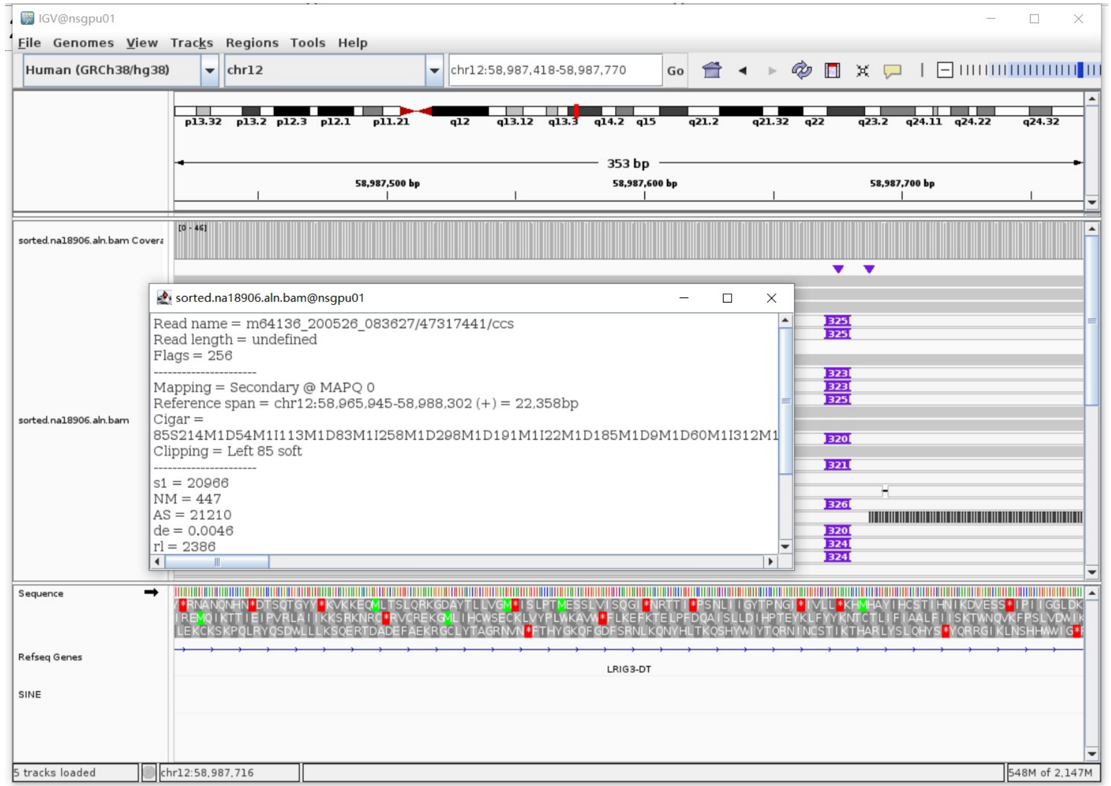


➢ Fetch one read (m64136_200526_083627/47317441/ccs, 22763 bp) and BLAT it to the hg38 genome.

The best alignment is to chr12_GL383552v1_alt (an alternative sequence of chr12).

| ACTIONS | QUERY | SCORE | START | END | QSIZE | IDENTITY | CHROM | STRAND | START | END | SPAN |
|---|---|---|---|---|---|---|---|---|---|---|---|
| browser new tab details | YourSeq | 22331 | 1 | 22763 | 22763 | 99.8% | chr12_GL383552v1_alt | + | 43645 | 66086 | 22442 |
| browser new tab details | YourSeq | 22235 | 86 | 22763 | 22763 | 99.8% | chr12 | + | 58965945 | 58988301 | 22357 |

- Read:1-21824 aligns to chr12_GL383552v1_alt

| ACTIONS | QUERY | SCORE | START | END | QSIZE | IDENTITY | CHROM | STRAND | START | END | SPAN |
|---|---|---|---|---|---|---|---|---|---|---|---|
| browser new tab details | YourSeq | 21717 | 1 | 21823 | 21823 | 99.9% | chr12_GL383552v1_alt | + | 43646 | 65471 | 21826 |
| browser new tab details | YourSeq | 21622 | 85 | 21823 | 21823 | 99.8% | chr12 | + | 58965945 | 58987686 | 21742 |

- Read:21825-22114 aligns to *Alu*Yb8:1-289, 99.65% identity

GGCCGGGCGCGGTGGCTCACGCCTGTAATCCCAGCACTTTGGGAGGCCGAGGCGGGTGGATCATGAGGTCAGGA
GATCGAGACCATCCTGGCTAACAAGGTGAAACCCCGTCTCTACTAAAAATACAAAAAATTAGCCGGGCGCGGTG
GCGGGCGCCTGTAGTCCCAGCTACTCGGGAGGCTGAGGCAGGAGAATGGCGTGAACCCGGGAAGCGGAGCTTGC
AGTGAGCCGAGATTGCGCCACTGCAGTCCGCAGTCCGACCTGGGCGACAGAGCGAGACTCCGTCTCA

12bp TSD

AAAAACACATGC

- Read:22115-22763 also aligns to chr12_GL383552v1_alt

| ACTIONS | QUERY | SCORE | START | END | QSIZE | IDENTITY | CHROM | STRAND | START | END | SPAN |
|---|---|---|---|---|---|---|---|---|---|---|---|
| browser new tab details | YourSeq | 625 | 22 | 648 | 648 | 99.6% | chr12_GL383552v1_alt | + | 65461 | 66086 | 626 |
| browser new tab details | YourSeq | 625 | 22 | 648 | 648 | 99.6% | chr12 | + | 58987676 | 58988301 | 626 |

➢ It's an *Alu*Yb8 insertion.

### NA18906 chr13:62014795

- ➢ Using the built-in BLAT tool in IGV to check where the inserted sequence aligned.
- ➢ The annotation at the aligned location indicated it as *AluY*.

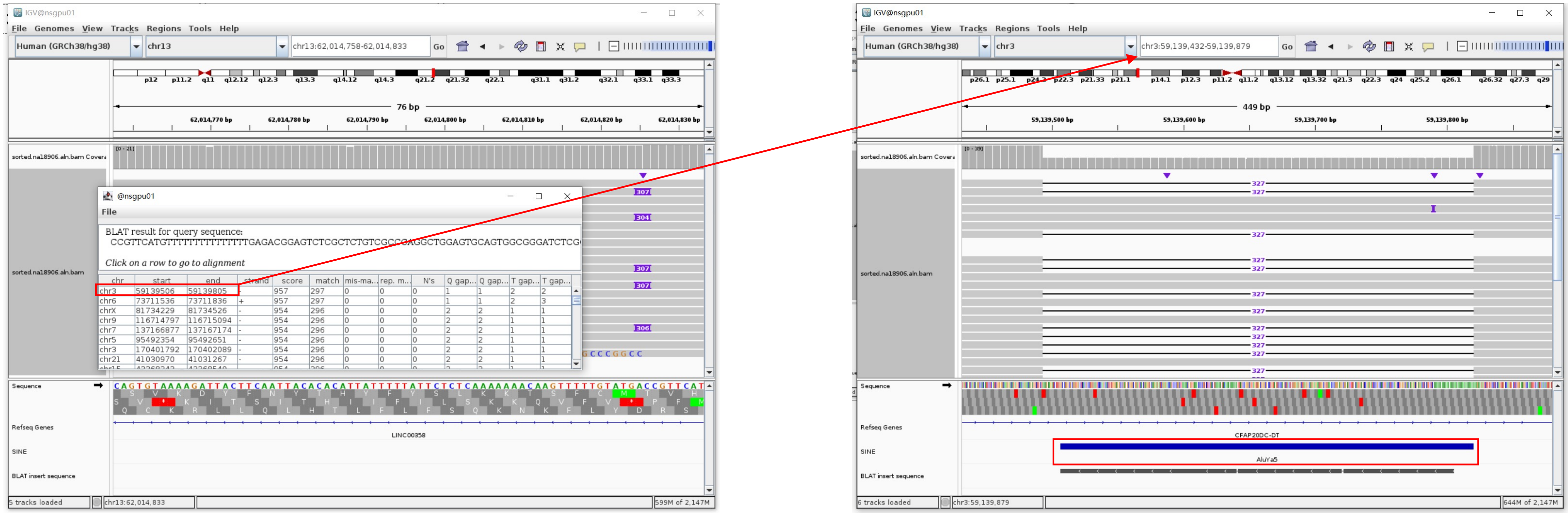


Insertion sequence (307bp)

- ➢ 15bp TSD

CCGTTCATGTTTTTT

- ➢ Insertion:23-307 aligns to *AluYa5*:282-1, 99.64% identity

TGAGACGGAGTCTCGCTCTGTCGCCCAGGCTGGAGTGCAGTGGCGGGATCTCGGCTCACTGCAAGCTCCGCTCCCGGGTTCACGCCATTCTCCTGCCTCAGCCTCCCAAGTAGCTGGGACTAC
AGGCGCCCGCCACTACGCCCGGCTAATTTTTTTGTATTTTTAGTAGAGACGGGGTTTCACCGTTTTTAGCCGGGATGGTCTCGATCTCTTGACCTCGTGATCCGCCCGCCTCGGCCTCCCAA
AGTGCTGGGATTACAGGCGTGAGCCACCGCGCCCGGCC

### NA18906 chr14:91780873

➢ The same genomic coordinate as in HG02630 (page 35)

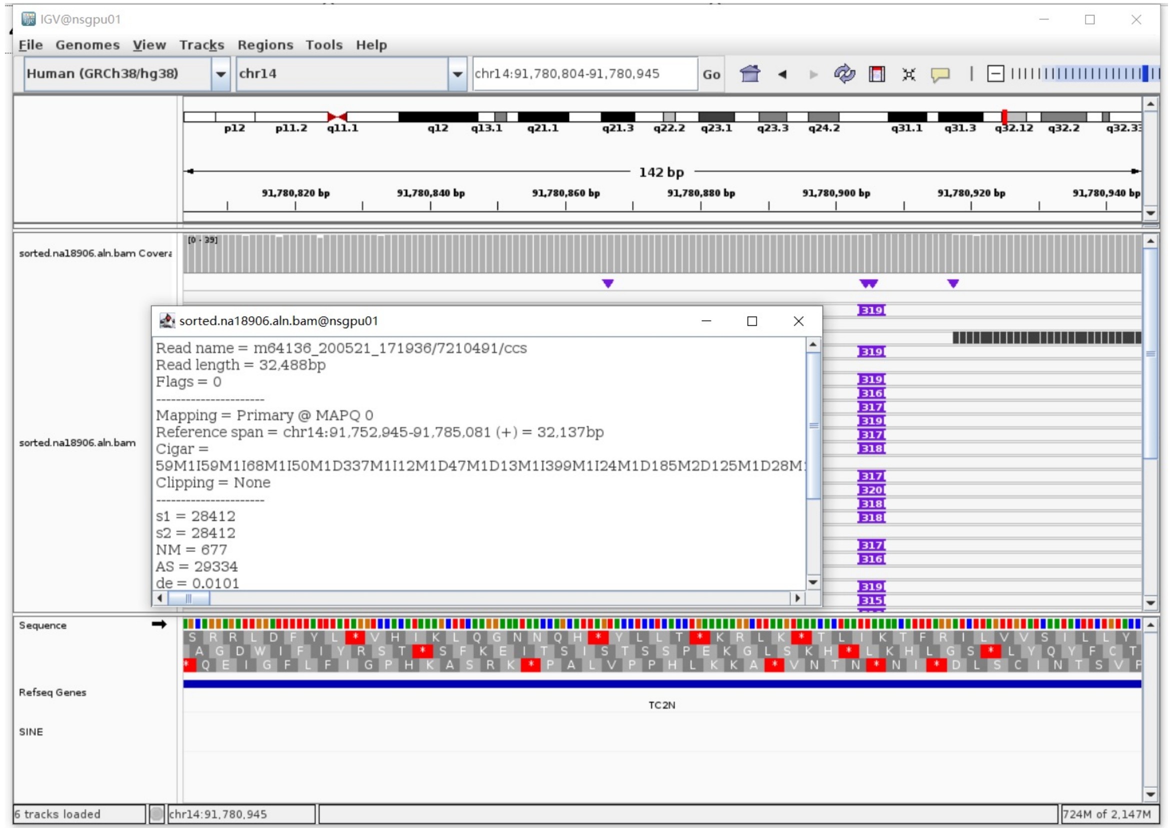

### NA18906 chr14:106651070

➢ The same genomic coordinate as in HG03516 (page 70)

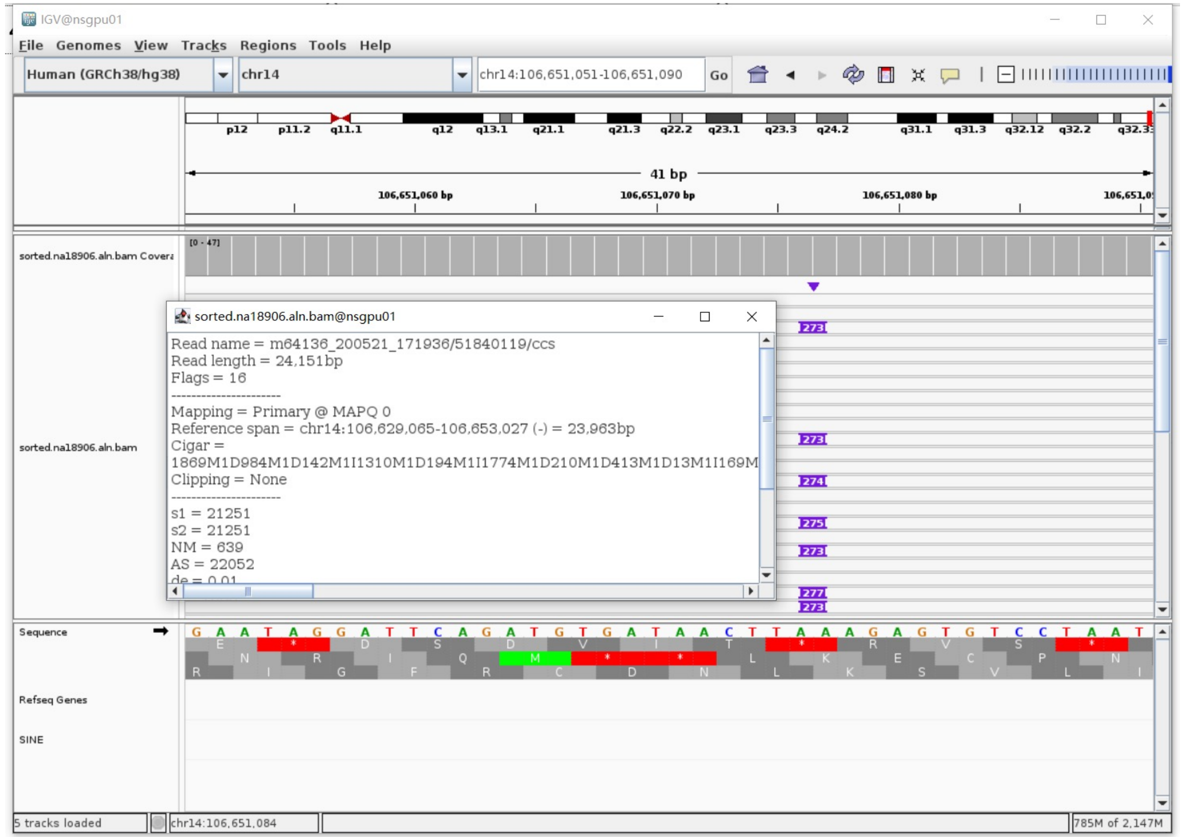

### NA18906 chr17:17831055

- Using the built-in BLAT tool in IGV to check where the inserted sequence aligned.

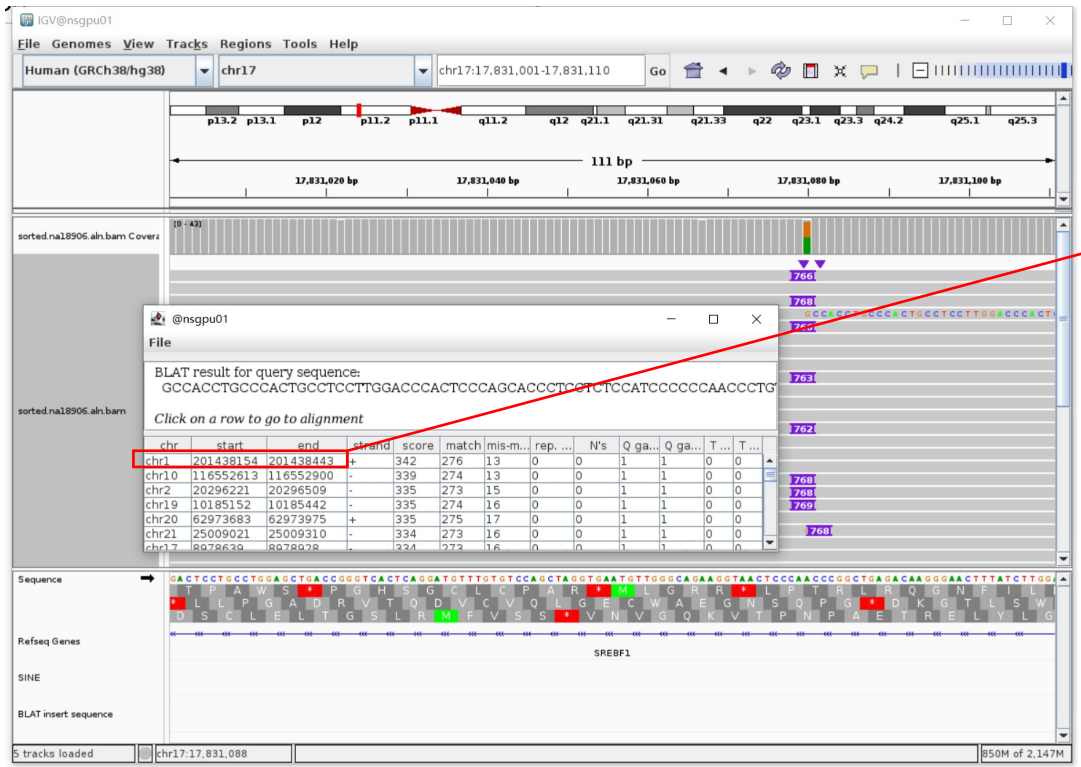


- The annotation at the aligned location indicated it as *AluY*.

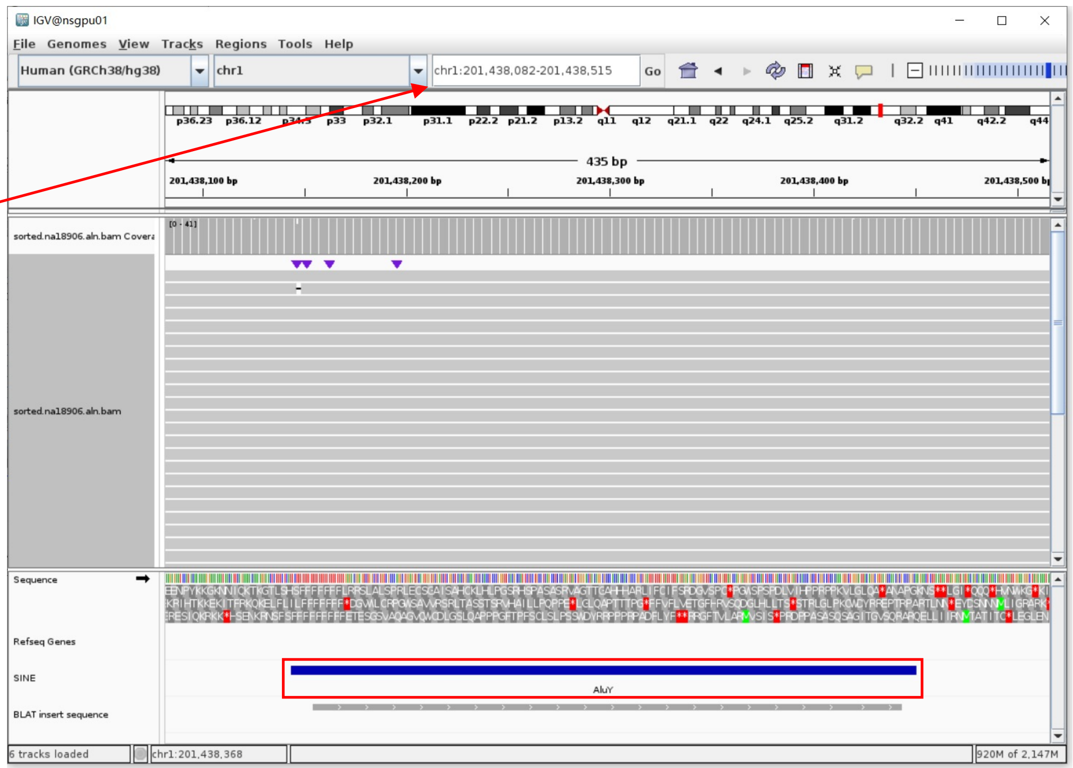


Insertion sequence (766bp)

- Insertion:491-766 aligns to *Alu*Yc1:282-8, 94.53% identity

TGAGACGGAGTCTCGCTCTGTCGCCCAGGCTGGAGTGCAGTGGCGGGATCTCGGCTCACTGCAAGCTCCGCTCCCGGGTTCACGCCATTCTCCTGCCTCAGCCTCCCAAGTAGCTGGGACTAC
AGGCGCCCGCCACTACGCCCGGCTAATTTTTTTGTATTTTTAGTAGAGACGGGGTTTCACCGTTTTAGCCGGGATGGTCTCGATCTCTTGACCTCGTGATCCGCCCGCCTCGGCCTCCCAA
AGTGCTGGGATTACAGGCGTGAGCCACCGCGCCCGGCC

### NA18906 chr17:37561902

- The same genomic coordinate as in HG00621 (page 15)

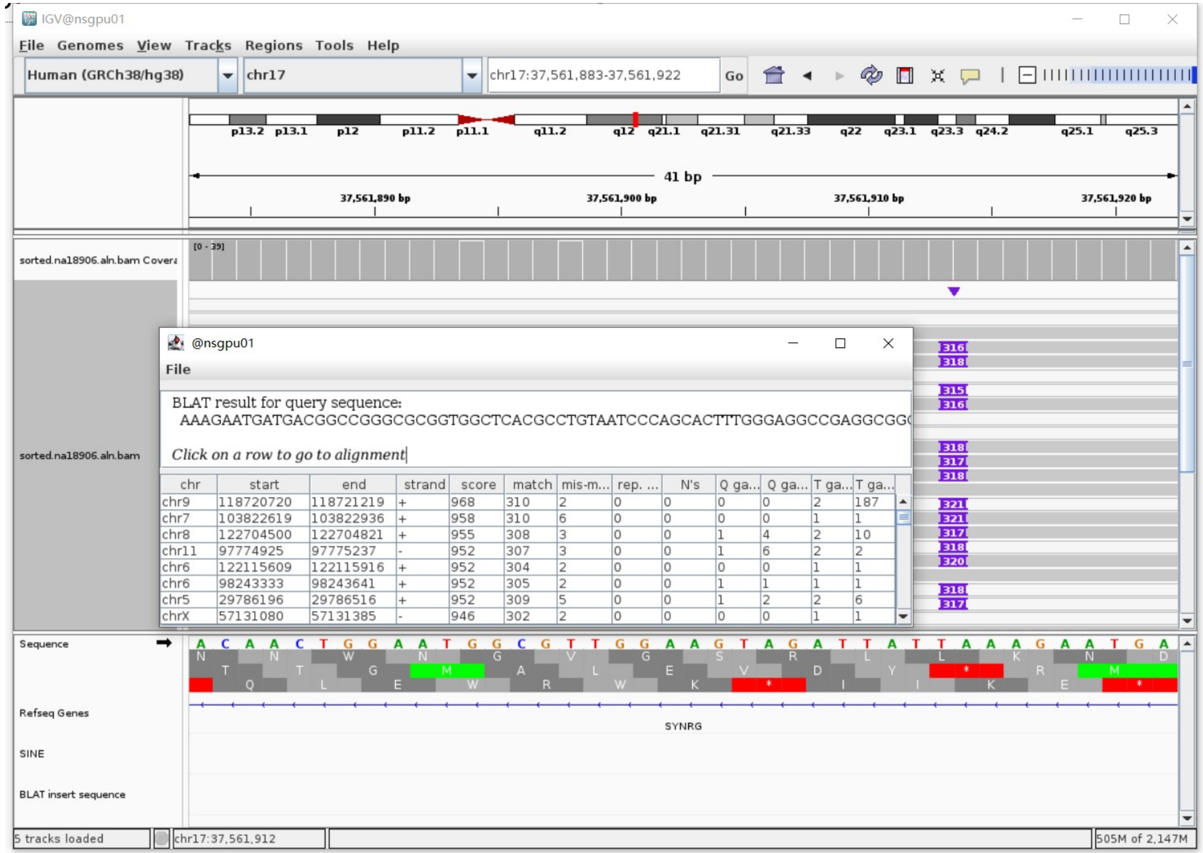

### NA18906 chr18:29602115

- The same genomic coordinate as in HG03492 (page 57)

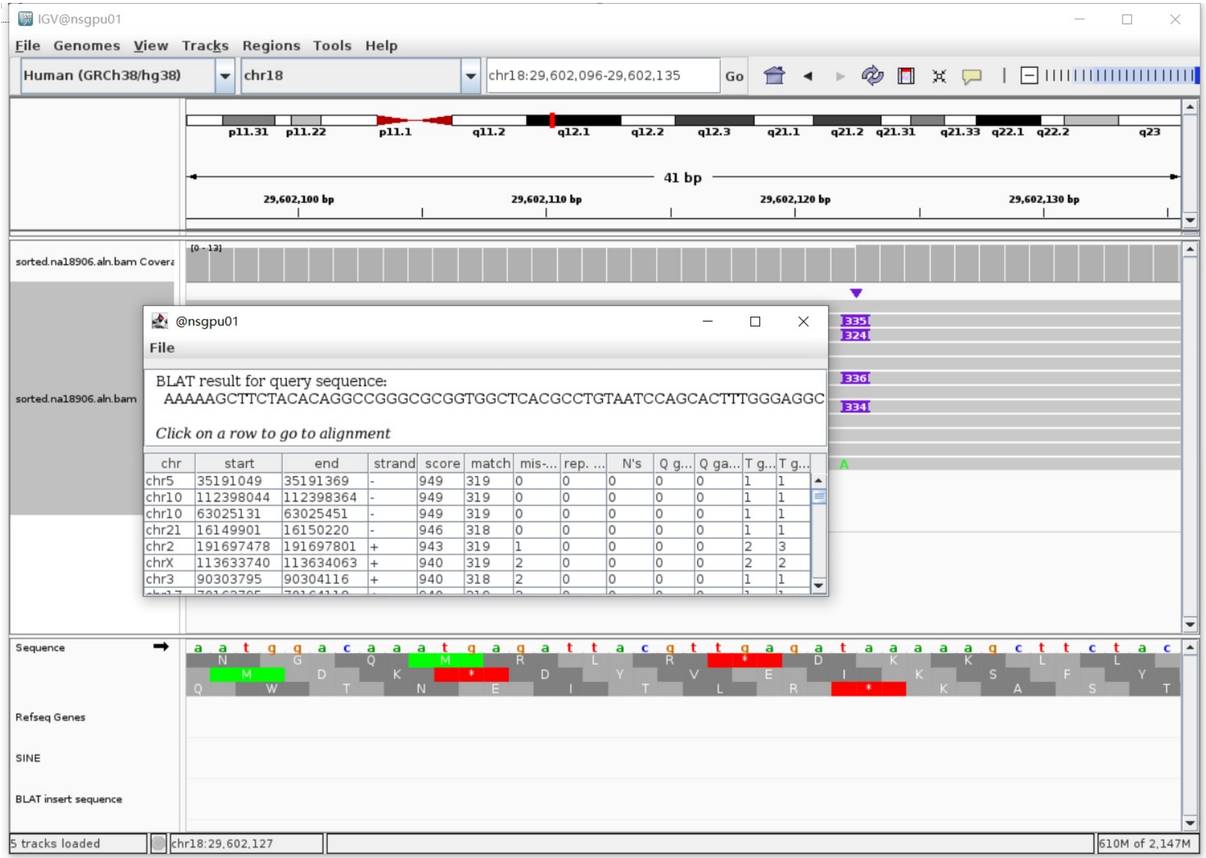

### NA18906 chr22:23928251

- The same genomic coordinate as in HG03540 (page 92)

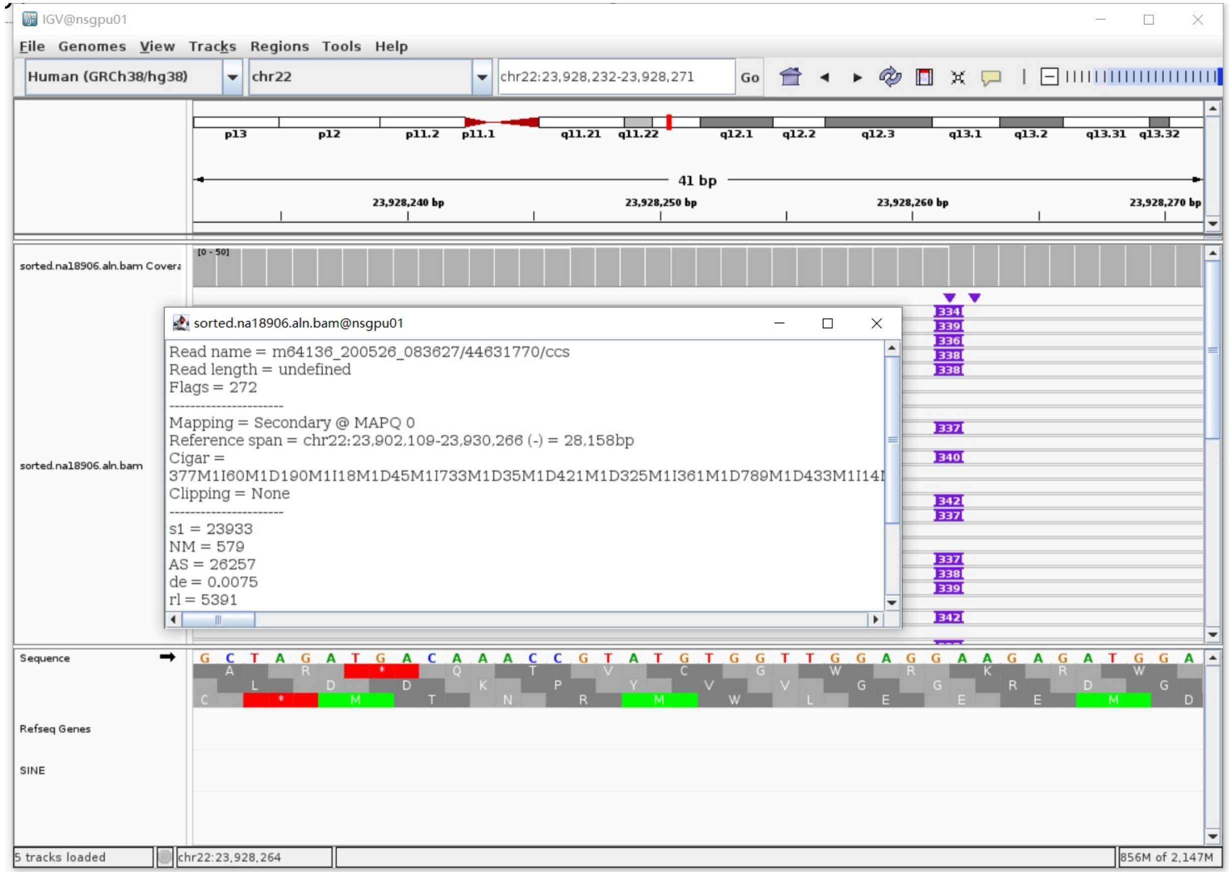

### NA20129 chr1:82577498

- Using the built-in BLAT tool in IGV to check where the inserted sequence aligned.

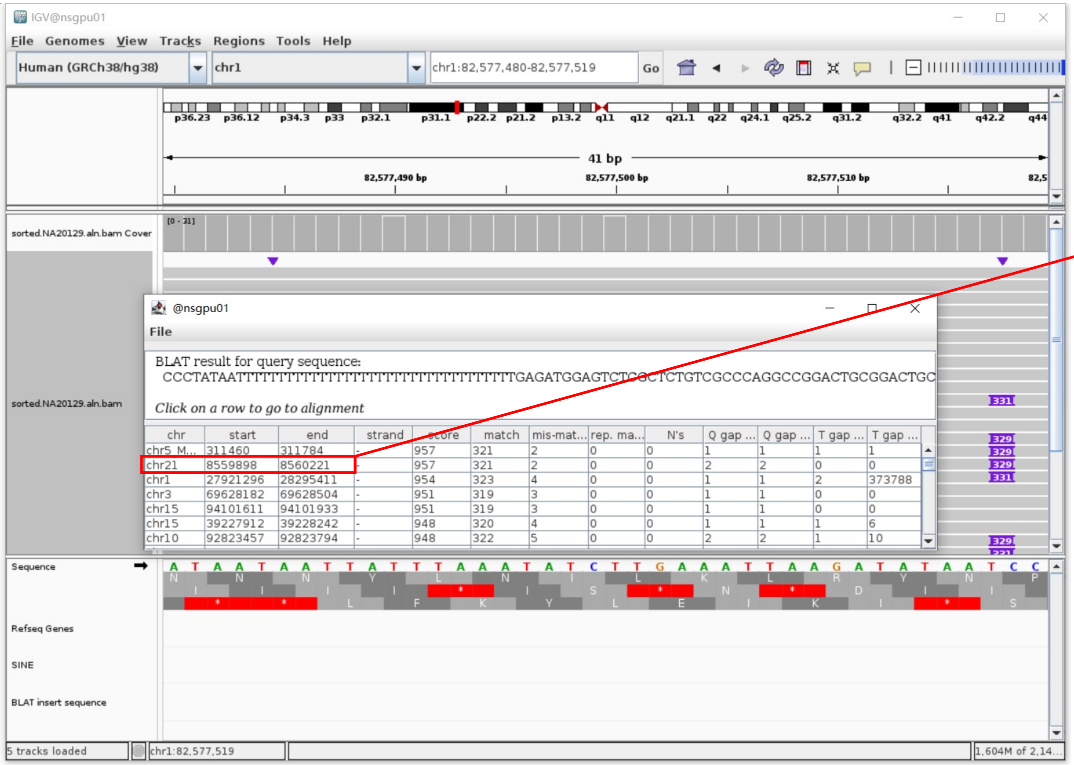


- The annotation at the aligned location indicated it as *AluY*.

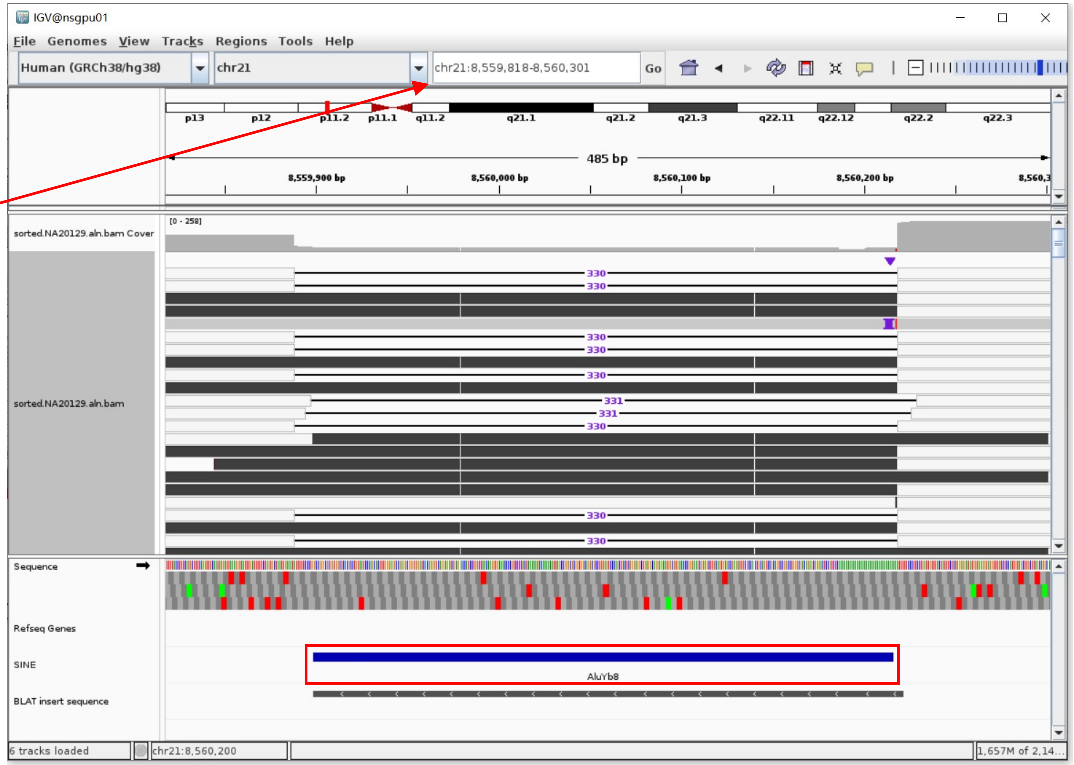


Insertion sequence (331bp)

- 10bp TSD

CCCTATAATT

- Insertion:41-331 aligns to *Alu*Yb8:289-1, 98.62% identity

TGAGATGGAGTCTCGCTCTGTCGCCCAGGCCGGACTGCGGACTGCAGTGGAGCAATCTCGGCTCACTGCAAGCTCCGCTTCCCGGGTTCACGCCATTCTCCTGCCTCAGCCTCCCGAGTAGCT
GGGGACTACAGGCGCCTGCCACCGCGCCCGGCTAATTTTTTGTATTTTTAGTAGAGACGGGGTTTCACCTTGTTAGCCAGGATGGTCTCGATCTCCTGACCTCGTGATCCACCCGCCTCGGCC
TCCCAAAGTGCTGGGATTACAGGCGTGAGCCACCGCGCCCGGCC

### NA20129 chr4:63331066

- The same genomic coordinate as in HG03540 (page 76)

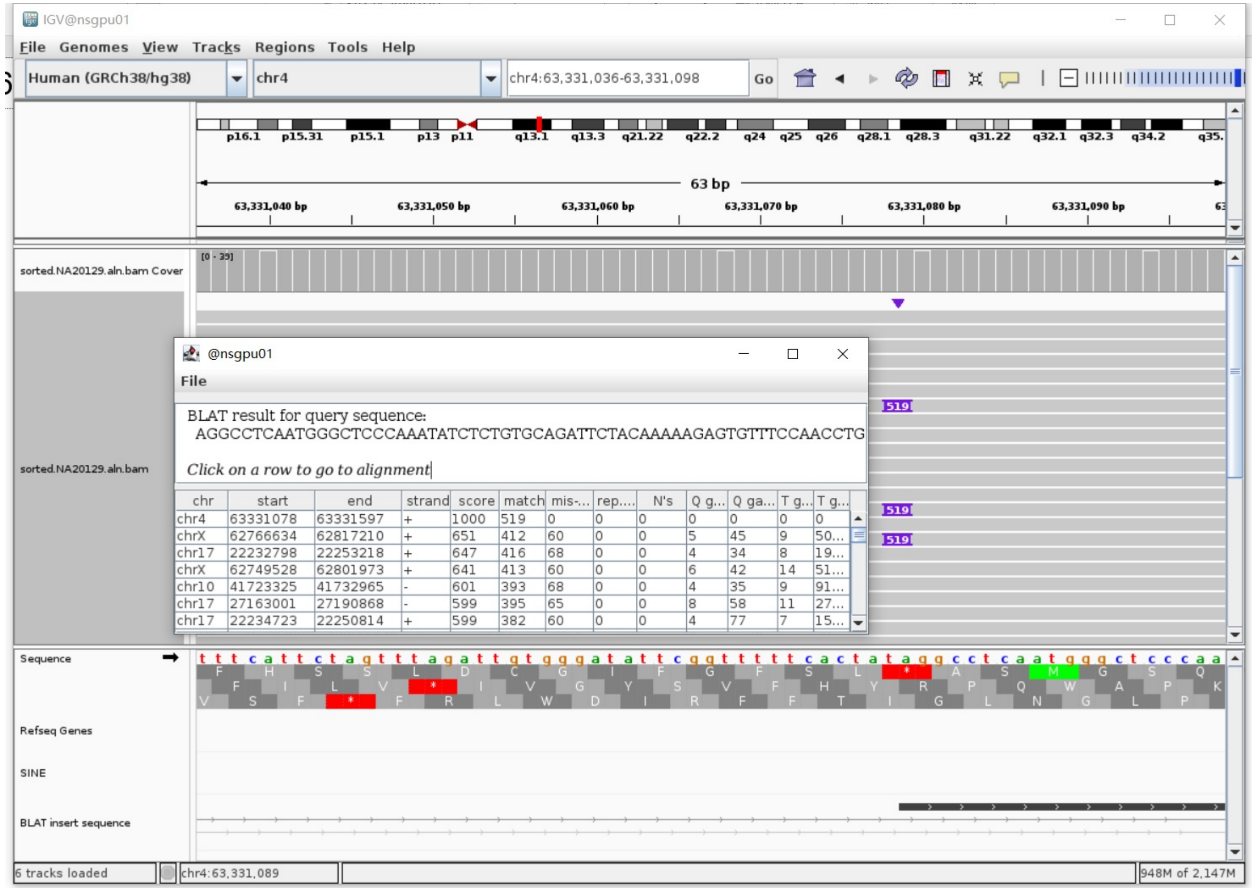


| chr | start | end | strand | score | match | mis-... | rep.... | N's | Q g... | Q ga... | T g... | T g... |
|---|---|---|---|---|---|---|---|---|---|---|---|---|
| chr4 | 63331078 | 63331597 | + | 1000 | 519 | 0 | 0 | 0 | 0 | 0 | 0 | 0 |
| chrX | 62766634 | 62817210 | + | 651 | 412 | 60 | 0 | 0 | 5 | 45 | 9 | 50... |
| chr17 | 22232798 | 22253218 | + | 647 | 416 | 68 | 0 | 0 | 4 | 34 | 8 | 19... |
| chrX | 62749528 | 62801973 | + | 641 | 413 | 60 | 0 | 0 | 6 | 42 | 14 | 51... |
| chr10 | 41723325 | 41732965 | - | 601 | 393 | 68 | 0 | 0 | 4 | 35 | 9 | 91... |
| chr17 | 27163001 | 27190868 | - | 599 | 395 | 65 | 0 | 0 | 8 | 58 | 11 | 27... |
| chr17 | 22234723 | 22250814 | + | 599 | 382 | 60 | 0 | 0 | 4 | 77 | 7 | 15... |

### NA20129 chr4:189727059

- ➢ Mapping quality is 0, indicating that this region may have alignment issues.

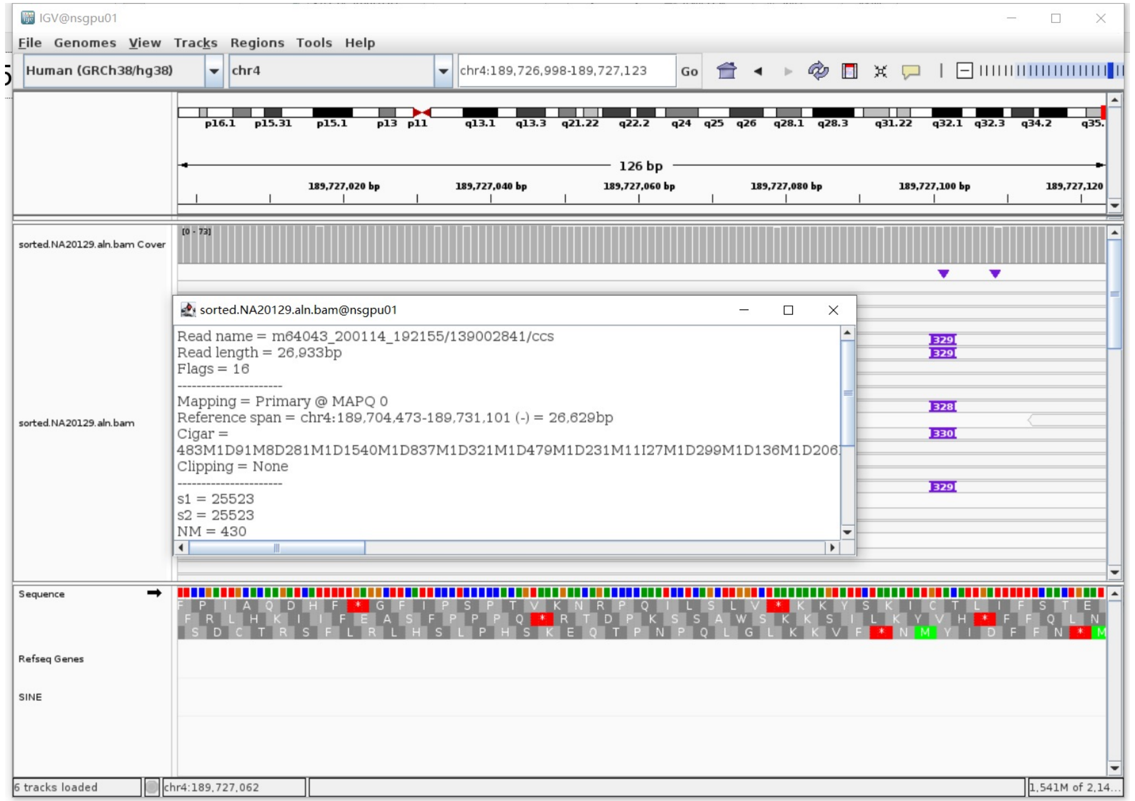


- ➢ Fetch one read (m64043_200114_192155/139002841/ccs, 26963 bp) and BLAT it to the hg38 genome.

  The best alignment is to chr4_KI270925v1_alt (an alternative sequence of chr4) and chr4.

| ACTIONS | | | QUERY | SCORE | START | END | QSIZE | IDENTITY | CHROM | STRAND | START | END | SPAN |
|---|---|---|---|---|---|---|---|---|---|---|---|---|---|
| browser | new tab | details | YourSeq | 26520 | 1 | 26963 | 26963 | 99.8% | chr4_KI270925v1_alt | - | 353202 | 379830 | 26629 |
| browser | new tab | details | YourSeq | 26520 | 1 | 26963 | 26963 | 99.8% | chr4_KI270896v1_alt | - | 38026 | 64654 | 26629 |
| browser | new tab | details | YourSeq | 26520 | 1 | 26963 | 26963 | 99.8% | chr4_KI270786v1_alt | - | 38026 | 64654 | 26629 |
| browser | new tab | details | YourSeq | 26520 | 1 | 26963 | 26963 | 99.8% | chr4 | - | 189704473 | 189731101 | 26629 |

- ● Read:1-4000 aligns to chr4_KI270925v1_alt

| ACTIONS | | | QUERY | SCORE | START | END | QSIZE | IDENTITY | CHROM | STRAND | START | END | SPAN |
|---|---|---|---|---|---|---|---|---|---|---|---|---|---|
| browser | new tab | details | YourSeq | 3994 | 1 | 3999 | 3999 | 100.0% | chr4_KI270925v1_alt | - | 375832 | 379829 | 3998 |
| browser | new tab | details | YourSeq | 3994 | 1 | 3999 | 3999 | 100.0% | chr4_KI270896v1_alt | - | 60656 | 64653 | 3998 |
| browser | new tab | details | YourSeq | 3994 | 1 | 3999 | 3999 | 100.0% | chr4_KI270786v1_alt | - | 60656 | 64653 | 3998 |
| browser | new tab | details | YourSeq | 3994 | 1 | 3999 | 3999 | 100.0% | chr4 | - | 189727103 | 189731100 | 3998 |

- ● Read:4001-4282 aligns to *Alu*Yc1:1-282, 97.86% identity

  GGCCGGGCGCGGTGGCTCACGCCTGTAATCCCAGCACTTTGGGAGGCCGAGGCGGGCGGATCACGAGGTCAGGA
  GATCGAGACCATCCTGGCTAACACGGTGAAACCCCGTCTCTACTAAAAATACAAAAAATTAGCCGGGCGCGGTG
  GCGGGCGCCTGTAGTCCCAGCTACTCGGGAGGCTGAGGCAGGAGAATGGCGTGAACCCCAGGGGGCGGAGCCTG
  CAGTGAGCCGAGATTGCGCCACTGCACTCCAGCCTGGGCGACAGCGAGACTCCGTCTCA

  12bp TSD

  AAAAATCAATGT

- ● Read:4283-26963 also aligns to chr4_KI270925v1_alt

| ACTIONS | | | QUERY | SCORE | START | END | QSIZE | IDENTITY | CHROM | STRAND | START | END | SPAN |
|---|---|---|---|---|---|---|---|---|---|---|---|---|---|
| browser | new tab | details | YourSeq | 22538 | 36 | 22680 | 22680 | 99.8% | chr4_KI270925v1_alt | - | 353202 | 375843 | 22642 |
| browser | new tab | details | YourSeq | 22538 | 36 | 22680 | 22680 | 99.8% | chr4_KI270896v1_alt | - | 38026 | 60667 | 22642 |
| browser | new tab | details | YourSeq | 22538 | 36 | 22680 | 22680 | 99.8% | chr4_KI270786v1_alt | - | 38026 | 60667 | 22642 |
| browser | new tab | details | YourSeq | 22538 | 36 | 22680 | 22680 | 99.8% | chr4 | - | 189704473 | 189727114 | 22642 |

- ➢ It's an *Alu*Yc1 insertion.

### NA20129 chr5:34717915

- ➢ Using the built-in BLAT tool in IGV to check where the inserted sequence aligned.

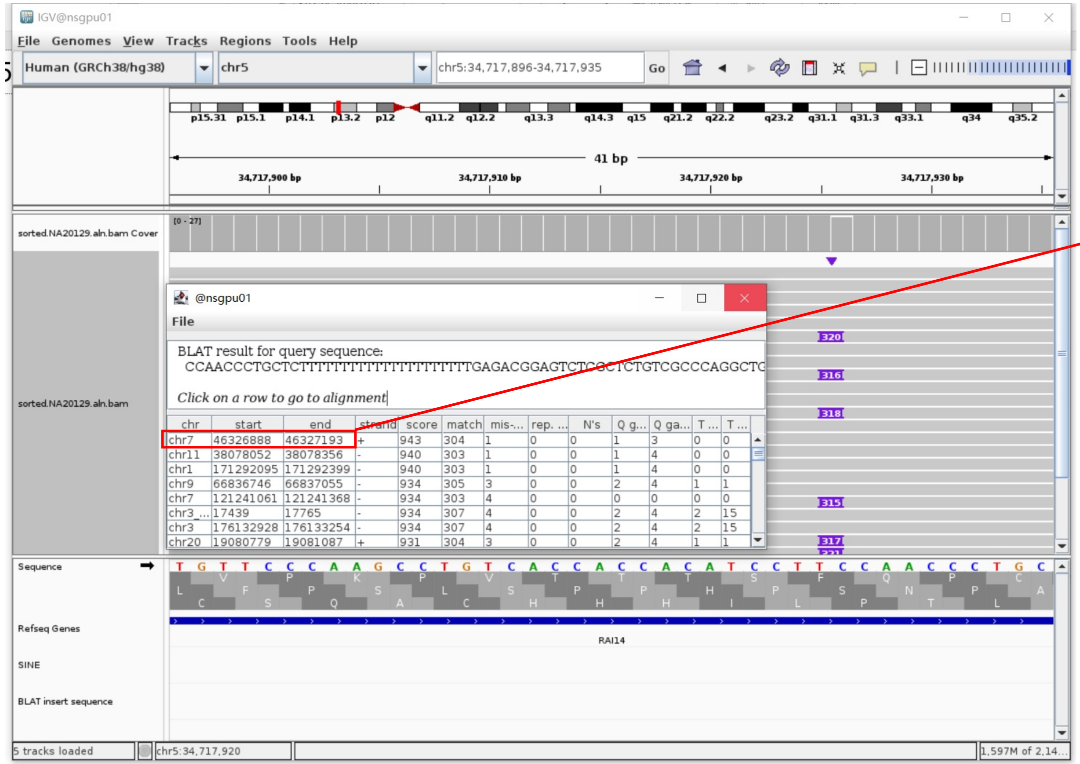


- ➢ The annotation at the aligned location indicated it as *AluY*.

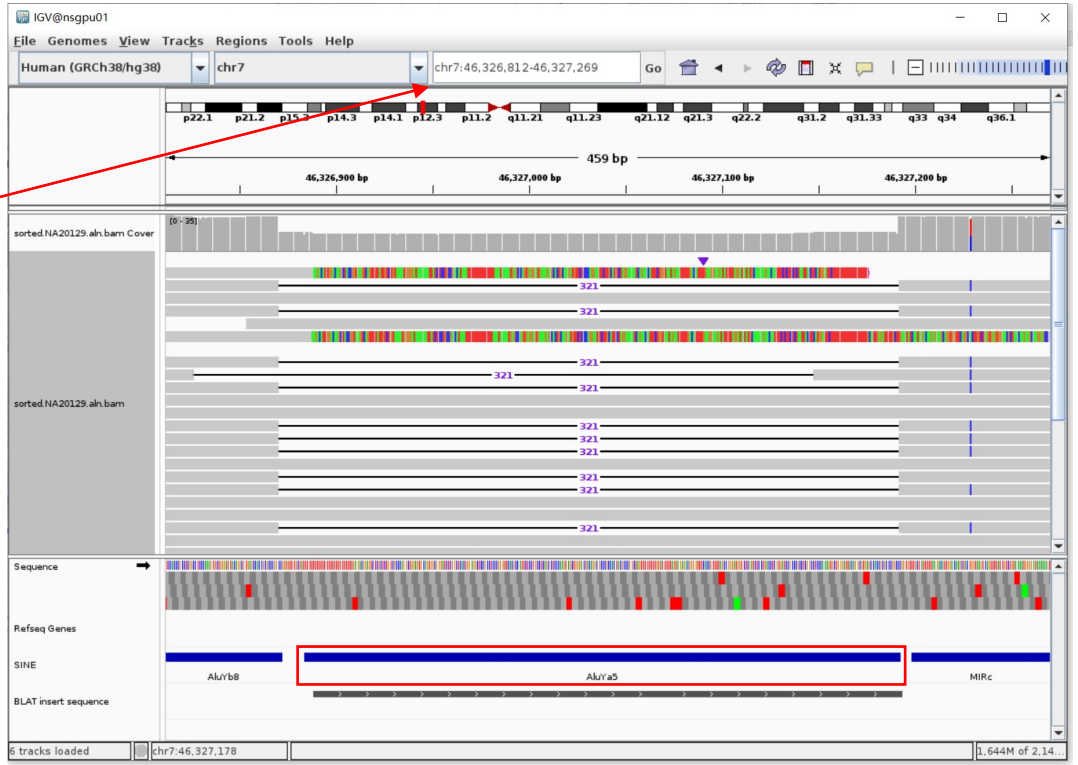


Insertion sequence (319bp)

- ➢ 14bp TSD

CCAACCCTGCTCTT

- ➢ Insertion:32-319 aligns to *AluYa5*:282-1, 98.58% identity

TGAGACGGAGTCTCGCTCTGTCGCCCAGGCTGGAGTGCAGTGGCGGGATCTCGGCTCACTGGAGGCTCCGCCTCCCGGGTTCACGCCATTCTCCTGCCTCAGCCTCCCAAGTAGCTGGGACTACCGGCGCCCGCCACTACGCCCGGCTAGTTTTTTTTTTGTATTTTTAGTAGAGACGGGGTTTCACCGTTTTAGCCGGGATGGTCTCGATCTCCTGACCTCGTGATCCGCCCGCCTCGGCCTCCCAAAGTGCTGGGATTACAGGCGTGAGCCACCGCGCCCGGCCC

### NA20129 chr6:28546544

- Mapping quality is 0, indicating that this region may have alignment issues.

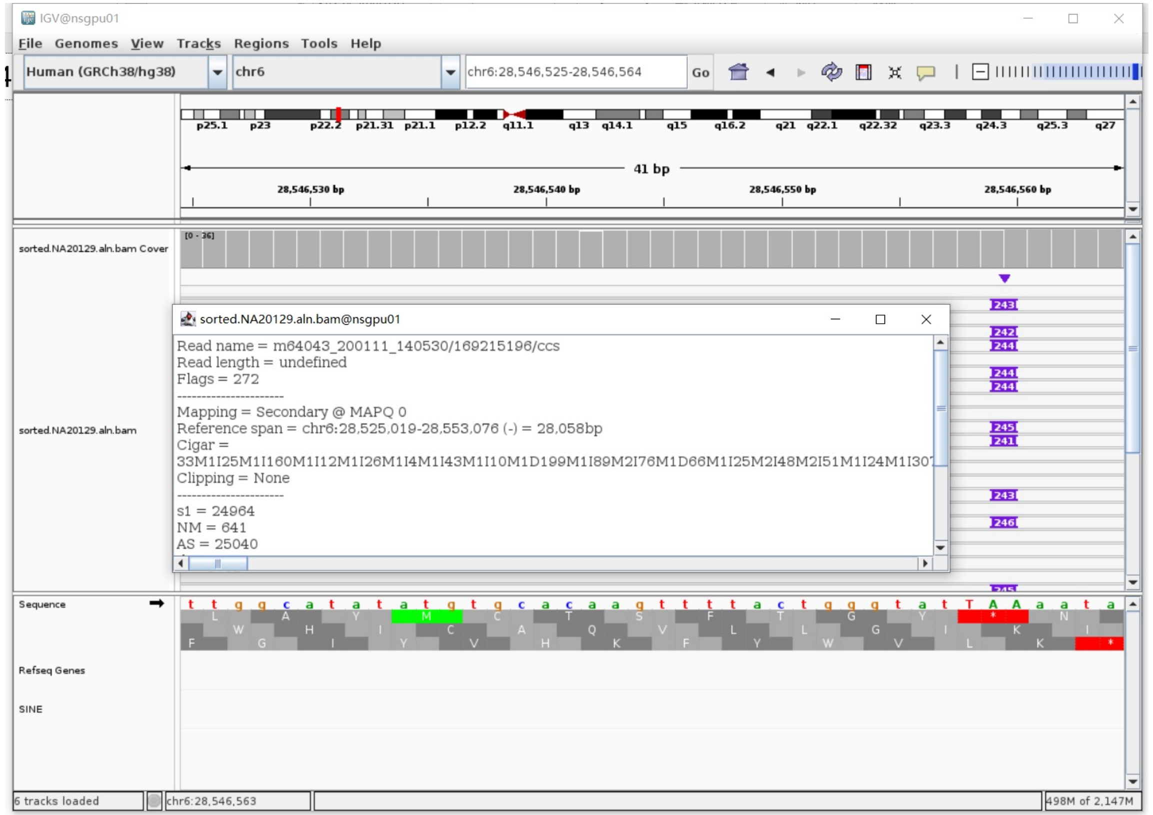


- Fetch one read (m64043_200111_140530/169215196/ccs, 28218 bp) and BLAT it to the hg38 genome.

  The best alignment is to chr6_GL000251v2_alt (an alternative sequence of chr6) and chr6.

| ACTIONS | QUERY | SCORE | START | END | QSIZE | IDENTITY | CHROM | STRAND | START | END | SPAN |
|---|---|---|---|---|---|---|---|---|---|---|---|
| browser new tab details | YourSeq | 27814 | 1 | 28218 | 28218 | 99.8% | chr6 | + | 28525019 | 28553072 | 28054 |
| browser new tab details | YourSeq | 27813 | 1 | 28218 | 28218 | 99.8% | chr6_GL000251v2_alt | + | 14900 | 42953 | 28054 |

- Read:1-21485 aligns to chr6_GL000251v2_alt and chr6

| ACTIONS | QUERY | SCORE | START | END | QSIZE | IDENTITY | CHROM | STRAND | START | END | SPAN |
|---|---|---|---|---|---|---|---|---|---|---|---|
| browser new tab details | YourSeq | 21354 | 1 | 21484 | 21484 | 99.8% | chr6 | + | 28525020 | 28546568 | 21549 |
| browser new tab details | YourSeq | 21353 | 1 | 21484 | 21484 | 99.8% | chr6_GL000251v2_alt | + | 14901 | 36449 | 21549 |

- Read:21486-21681 aligns to *Alu*Yb8:93-289, 98.46% identity

  AATAAGGTGAAACCCCGTCTCTACTAAAAATACAAAAAATTAGCCGGGCGTGGTGGCGGGCGCCTGTAGTCCC
  AGCTACTCGGGAGGCTGAGGCAGGAGAATGGCGTGAACCCGTGAAGCGGAGCTTGCAGTGAGCGAGATTGCGC
  CACTGCAGTCCGCAGTCCGGCCTGGGCGACAGAGCGAGACTCCGTCTCA

  15bp TSD

  AAATAAAGAGAATAA

- Read:21682-28218 also aligns to chr6_GL000251v2_alt and chr6

| ACTIONS | QUERY | SCORE | START | END | QSIZE | IDENTITY | CHROM | STRAND | START | END | SPAN |
|---|---|---|---|---|---|---|---|---|---|---|---|
| browser new tab details | YourSeq | 6469 | 38 | 6536 | 6536 | 100.0% | chr6_GL000251v2_alt | + | 36441 | 42953 | 6513 |
| browser new tab details | YourSeq | 6469 | 38 | 6536 | 6536 | 100.0% | chr6 | + | 28546560 | 28553072 | 6513 |

- It's an *Alu*Yb8 insertion.

### NA20129 chr6:29851072

- The same genomic coordinate as in HG00438 (page 3)

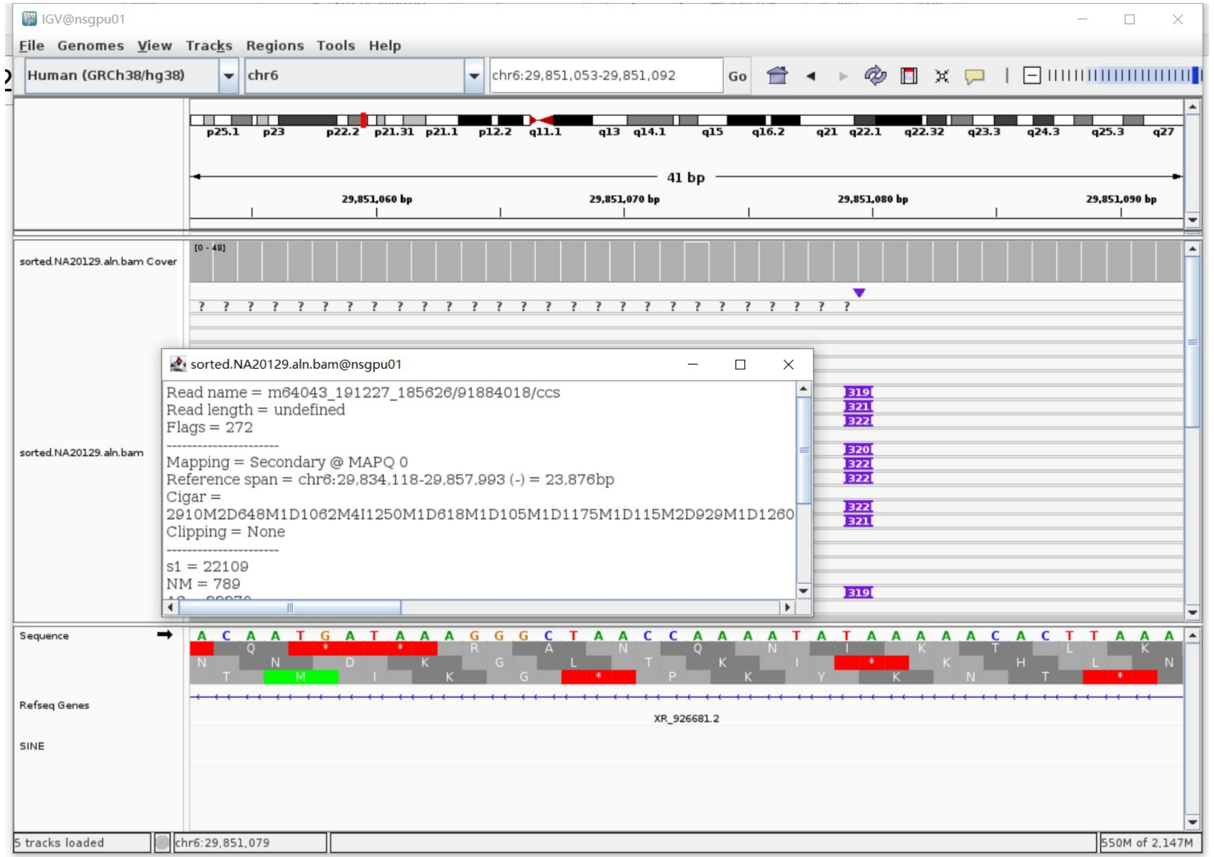

### NA20129 chr6:29925071

- The same genomic coordinate as in HG03540 (page 79)

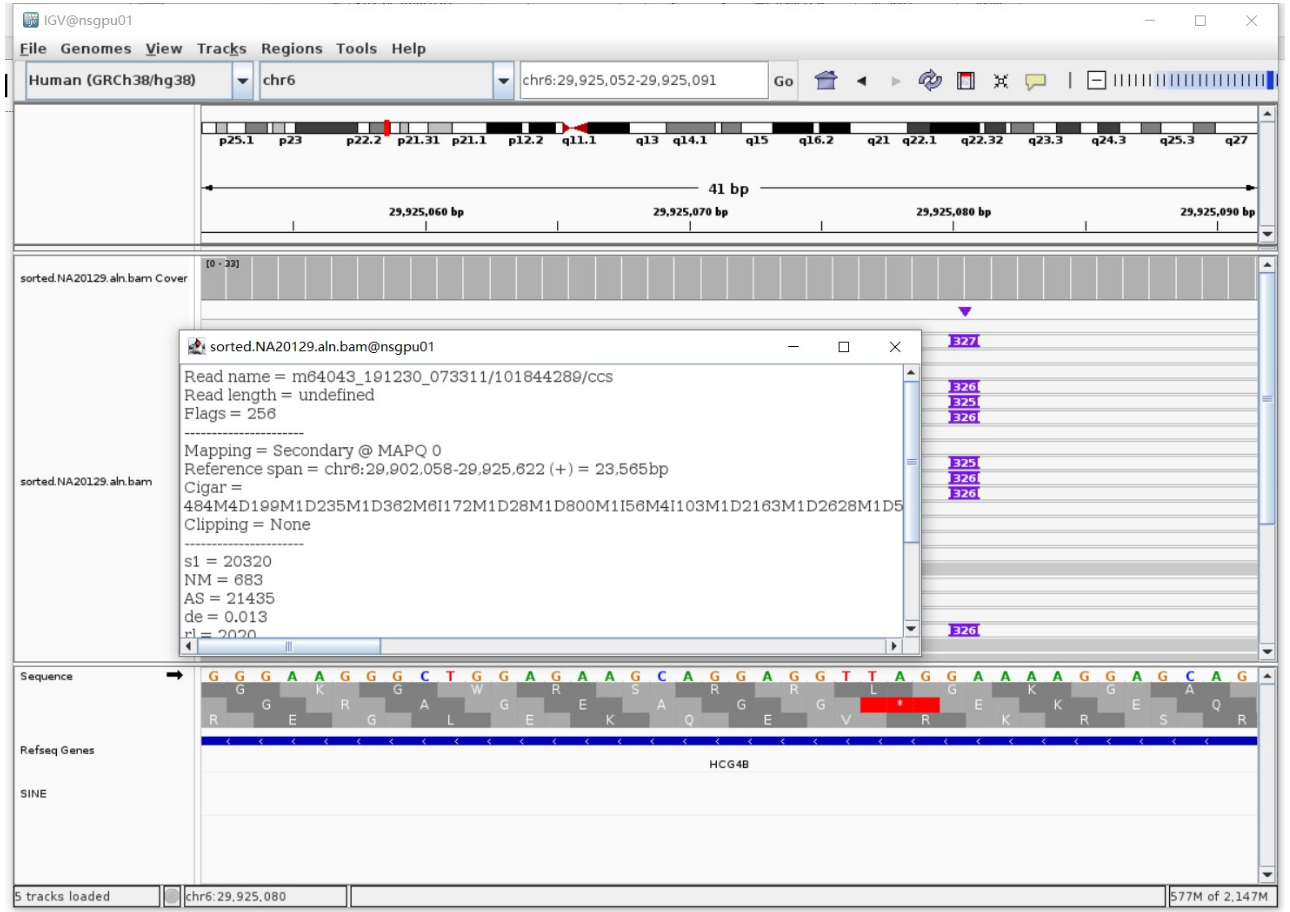

### NA20129 chr6:31282422

- The same genomic coordinate as in HG01106 (page 20)

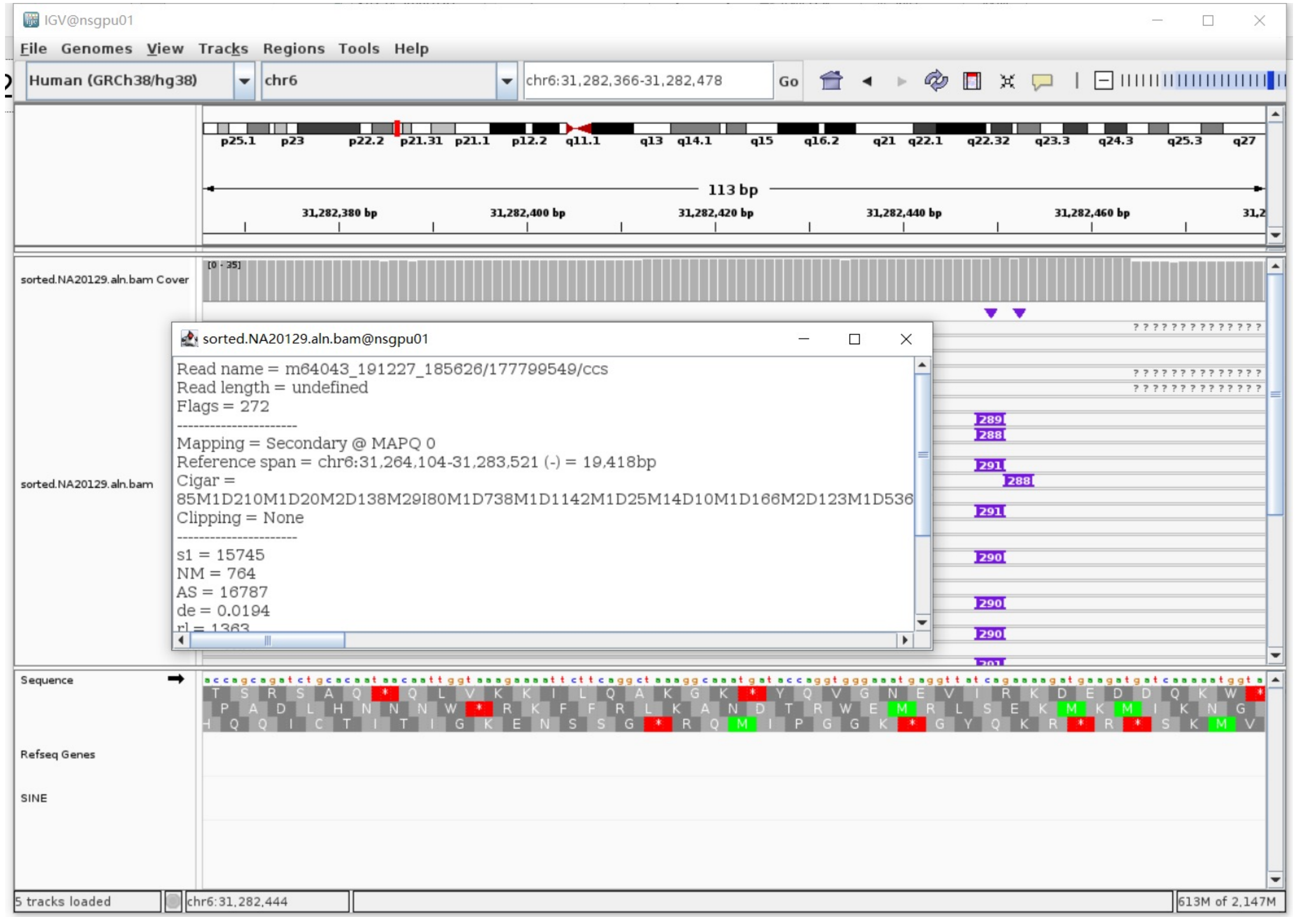

### NA20129 chr6:31336214

➢ Mapping quality is 0, indicating that this region may have alignment issues.

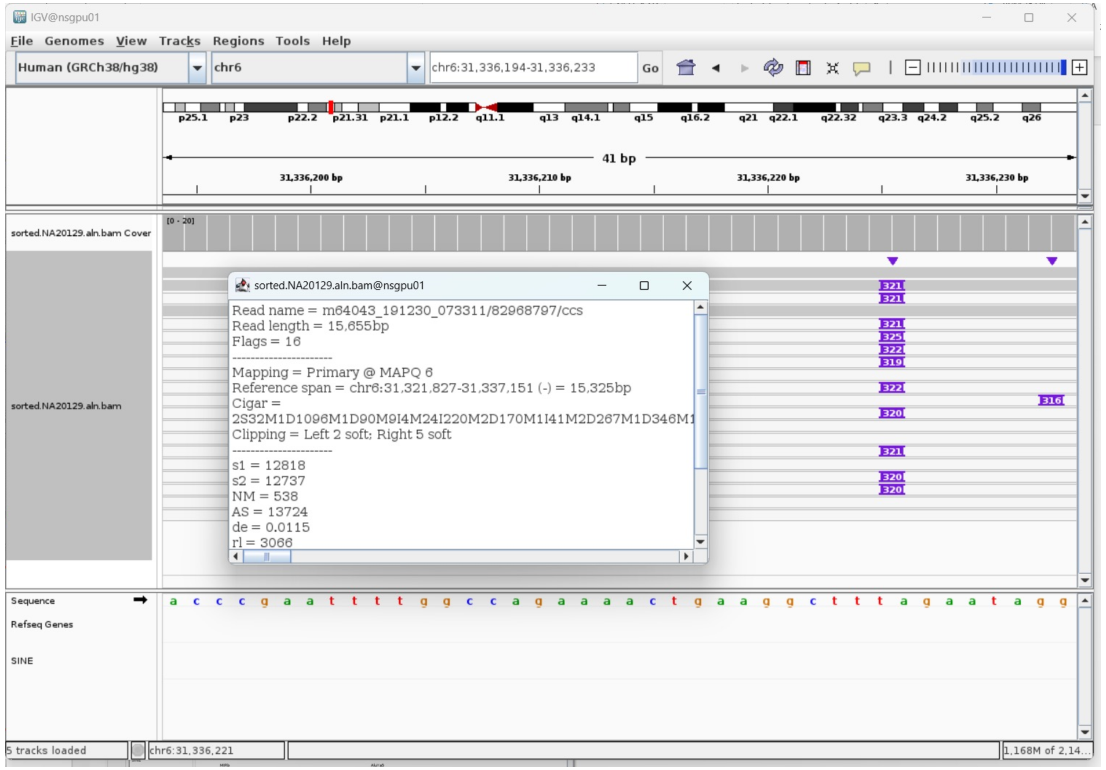


➢ Fetch one read (m64043_191230_073311/82968797/ccs, 15680 bp) and BLAT it to the hg38 genome.

The best alignment is to chr6_GL000256v2_alt (an alternative sequence of chr6) and chr6.

| ACTIONS | | | QUERY | SCORE | START | END | QSIZE | IDENTITY | CHROM | STRAND | START | END | SPAN |
|---|---|---|---|---|---|---|---|---|---|---|---|---|---|
| browser | new tab | details | YourSeq | 15105 | 1 | 15680 | 15680 | 99.3% | chr6 | - | 31321829 | 31337157 | 15329 |
| browser | new tab | details | YourSeq | 15077 | 1 | 15680 | 15680 | 99.2% | chr6_GL000256v2_alt | - | 2624030 | 2639355 | 15326 |
| browser | new tab | details | YourSeq | 15075 | 1 | 15680 | 15680 | 99.2% | chr6_GL000255v2_alt | - | 2577489 | 2592814 | 15326 |
| browser | new tab | details | YourSeq | 12430 | 1 | 15680 | 15680 | 96.0% | chr6_GL000251v2_alt | - | 2802756 | 2817495 | 14740 |
| browser | new tab | details | YourSeq | 12280 | 1 | 15680 | 15680 | 95.5% | chr6_GL000252v2_alt | - | 2569895 | 2584644 | 14750 |
| browser | new tab | details | YourSeq | 12275 | 1 | 15680 | 15680 | 95.5% | chr6_GL000253v2_alt | - | 2630924 | 2645724 | 14801 |
| browser | new tab | details | YourSeq | 12185 | 1 | 15680 | 15680 | 95.2% | chr6_GL000254v2_alt | - | 2664541 | 2679277 | 14737 |

- Read:1-956

| ACTIONS | | | QUERY | SCORE | START | END | QSIZE | IDENTITY | CHROM | STRAND | START | END | SPAN |
|---|---|---|---|---|---|---|---|---|---|---|---|---|---|
| browser | new tab | details | YourSeq | 927 | 1 | 930 | 955 | 99.9% | chr6 | - | 31336226 | 31337156 | 931 |
| browser | new tab | details | YourSeq | 923 | 1 | 930 | 955 | 99.7% | chr6_GL000251v2_alt | - | 2816564 | 2817494 | 931 |
| browser | new tab | details | YourSeq | 908 | 1 | 930 | 955 | 99.0% | chr6_GL000256v2_alt | - | 2638423 | 2639354 | 932 |
| browser | new tab | details | YourSeq | 908 | 1 | 930 | 955 | 99.0% | chr6_GL000255v2_alt | - | 2591882 | 2592813 | 932 |
| browser | new tab | details | YourSeq | 825 | 1 | 930 | 955 | 94.4% | chr6_GL000252v2_alt | - | 2583719 | 2584643 | 925 |
| browser | new tab | details | YourSeq | 821 | 1 | 930 | 955 | 94.2% | chr6_GL000254v2_alt | - | 2678352 | 2679276 | 925 |
| browser | new tab | details | YourSeq | 820 | 1 | 930 | 955 | 94.4% | chr6_GL000253v2_alt | - | 2644799 | 2645723 | 925 |

- Read:957-1239 aligns to *AluYa5*:282-1, 98.23% identity

TGAGACGGAGTCTCGCTCTGTCTCCCAGGCTGGAGTGCAGTGGCGGGATCTTGGCTCACTGCAAGCTCCGCCTC
CTGGGTTCACGCCATTCTCCTGCCTCAGCCTCCCAAGTAGCTGGGACTACAGGCGCCCGCCACTATGCCCGGCT
AATTTTTTGTATTTTTAGTAGAGACGGGGTTTCACCGTTTTAGCCGGGATGGTCTCGATCTCCTGACCTCGTGA
TCTGCCCGCCTCGGCCTCCCAAAGTGCTGGGATTACAGGCGTGAGCCACCGCGCCCGGCC

13bp TSD

AGAATAGGTTCCT

- Read:1240-15680

| ACTIONS | | | QUERY | SCORE | START | END | QSIZE | IDENTITY | CHROM | STRAND | START | END | SPAN |
|---|---|---|---|---|---|---|---|---|---|---|---|---|---|
| browser | new tab | details | YourSeq | 14190 | 1 | 14440 | 14440 | 99.3% | chr6 | - | 31321829 | 31336237 | 14409 |
| browser | new tab | details | YourSeq | 14181 | 1 | 14440 | 14440 | 99.3% | chr6_GL000256v2_alt | - | 2624030 | 2638434 | 14405 |
| browser | new tab | details | YourSeq | 14179 | 1 | 14440 | 14440 | 99.3% | chr6_GL000255v2_alt | - | 2577489 | 2591893 | 14405 |

➢ It's an *AluYa5* insertion.

### NA20129 chr6:32719367

- The same genomic coordinate as in NA18906 (page 97)

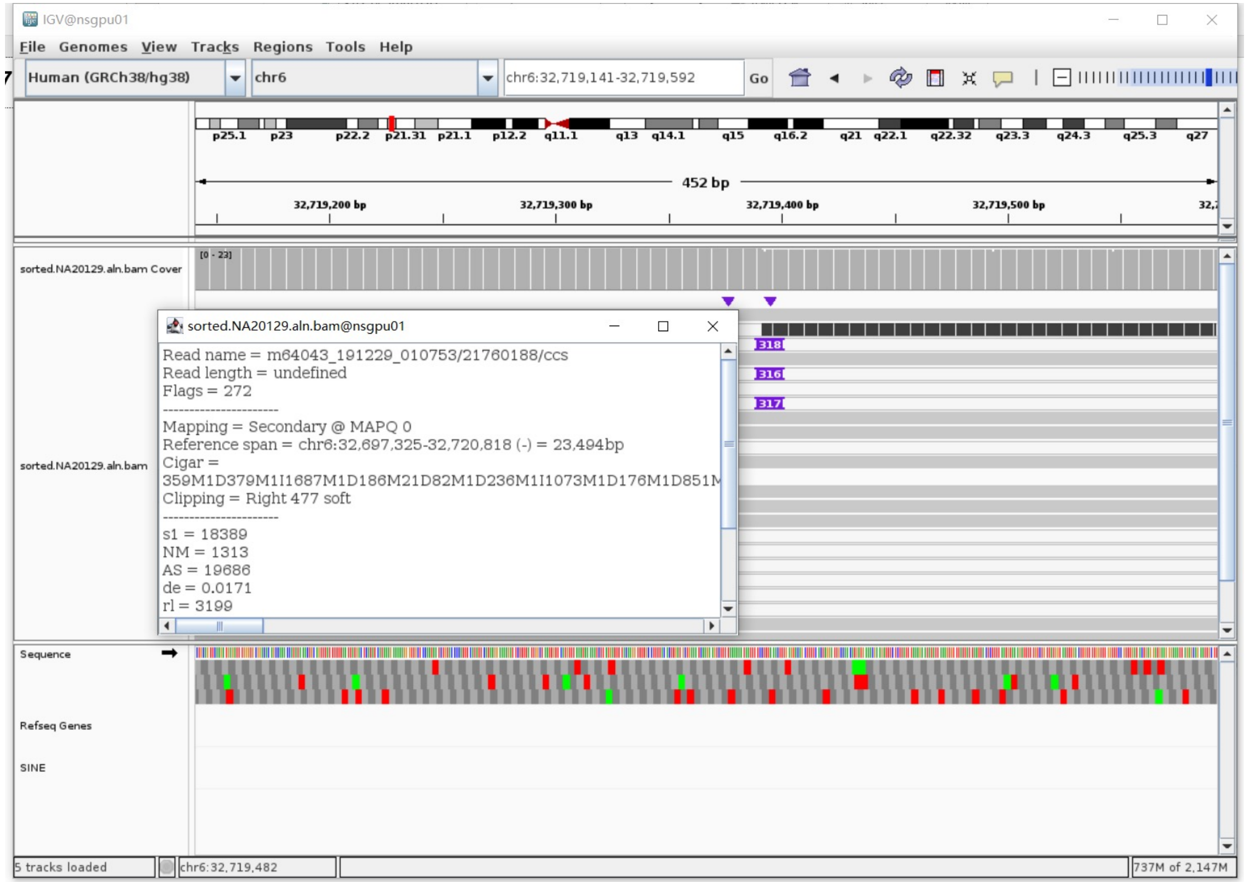

### NA20129 chr6:32720794

- The same genomic coordinate as in HG03492 (page 48)

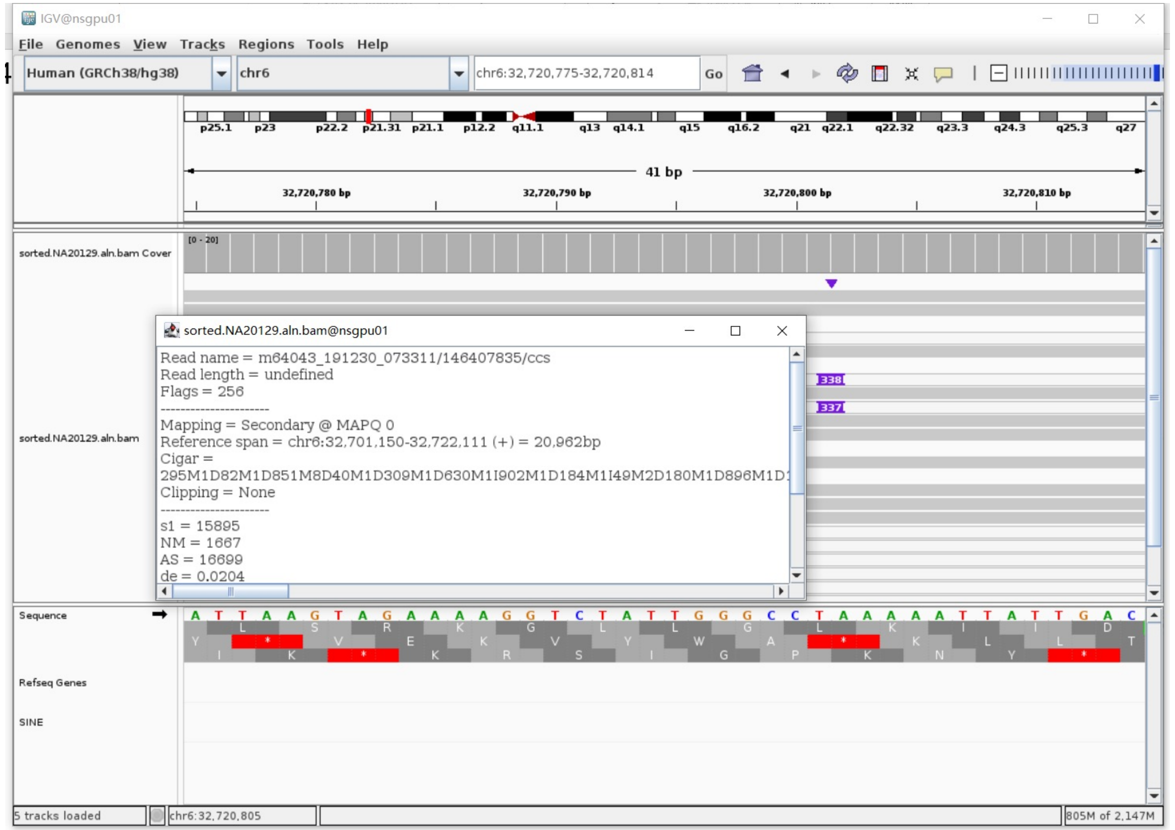

### NA20129 chr6:32805636

- The same genomic coordinate as in HG00438 (page 5)

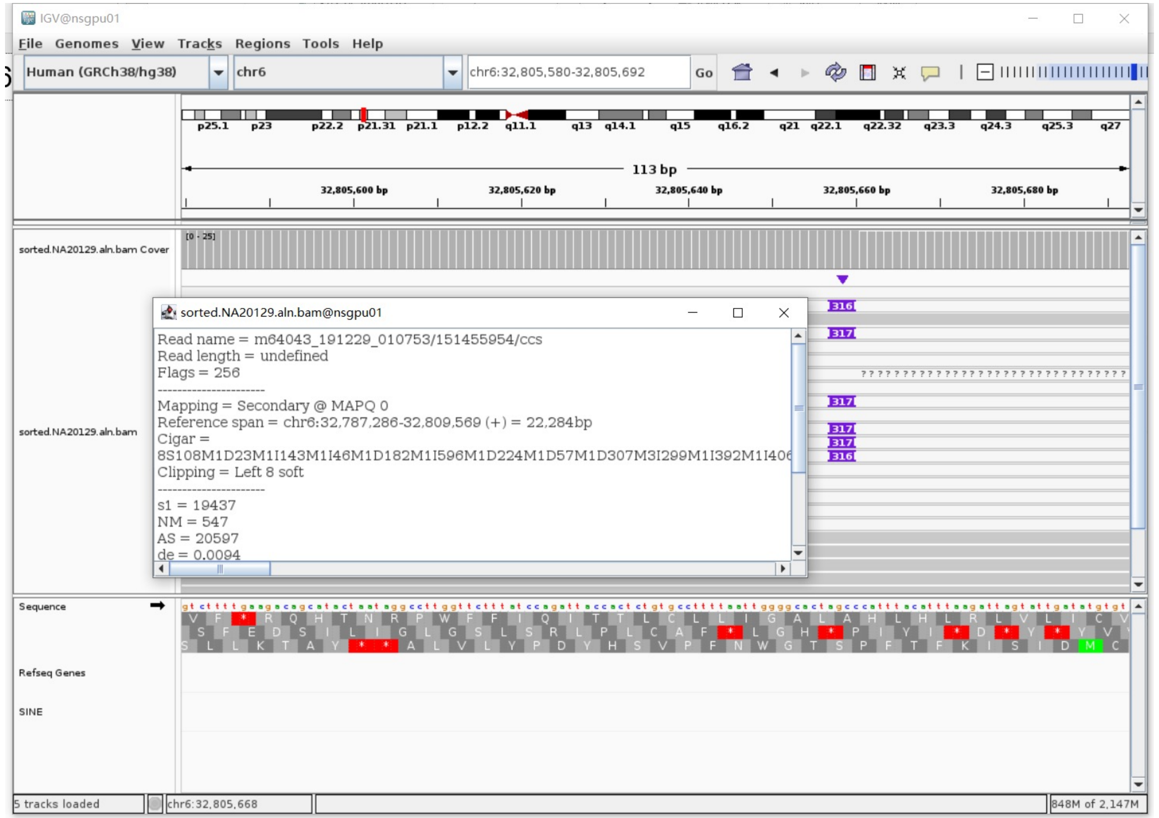

### NA20129 chr8:1362269

➢ Using the built-in BLAT tool in IGV to check where the inserted sequence aligned.

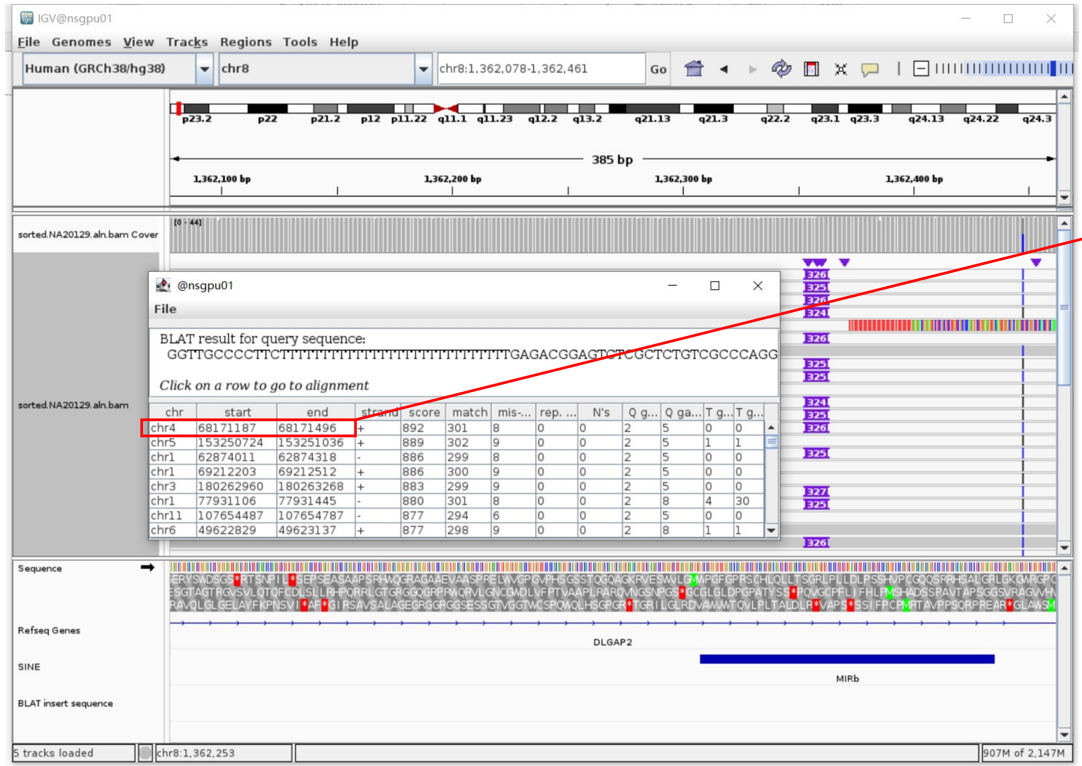


➢ The annotation at the aligned location indicated it as *AluY*.

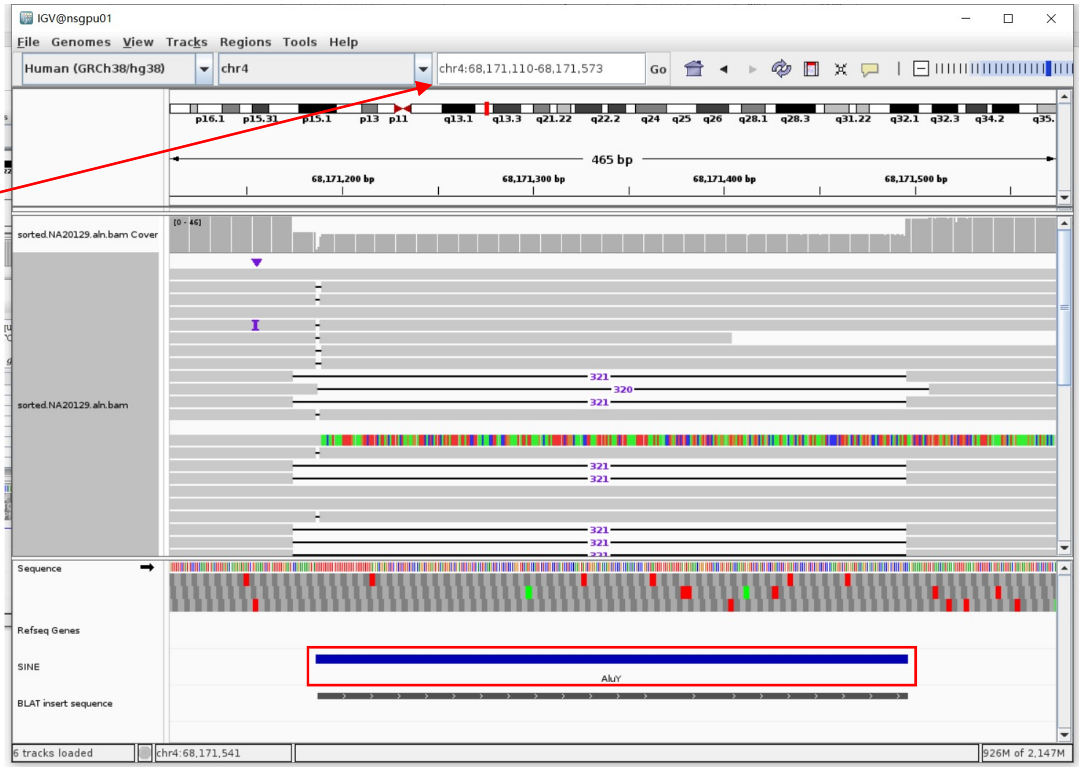


Insertion sequence (326bp)

➢ 14bp TSD

GGTTGCCCCTTCTT

➢ Insertion:39-326 aligns to *Alu*Yc1:282-1, 99.29% identity

TGAGACGGAGTCTCGCTCTGTCGCCCAGGCTGGAGTGCAGTGGCGCGATCTCGGCTCACTGCAAGCTCCGCCTCCCGGGTTCACGCCATTCTCCTGCCTCAGCCTCCCGAGTAGCTGGGACTACAGGCGCCCGCCAACACGCCCGGCTAATTTTTTTTTTGTATTTTTAGTAGAGACGGGGTTTCACCGTGTTAGCCAGGATGGTCTCGATCTCCTGACCTCGTGATCCGCCCGCCTCGGCCTCCCAAAGTGCTGGGATTACAGGCGTGAGCCACCGCGCCCGGCC

### NA20129 chr8:39842932

➢ The same genomic coordinate as in HG03516 (page 64)

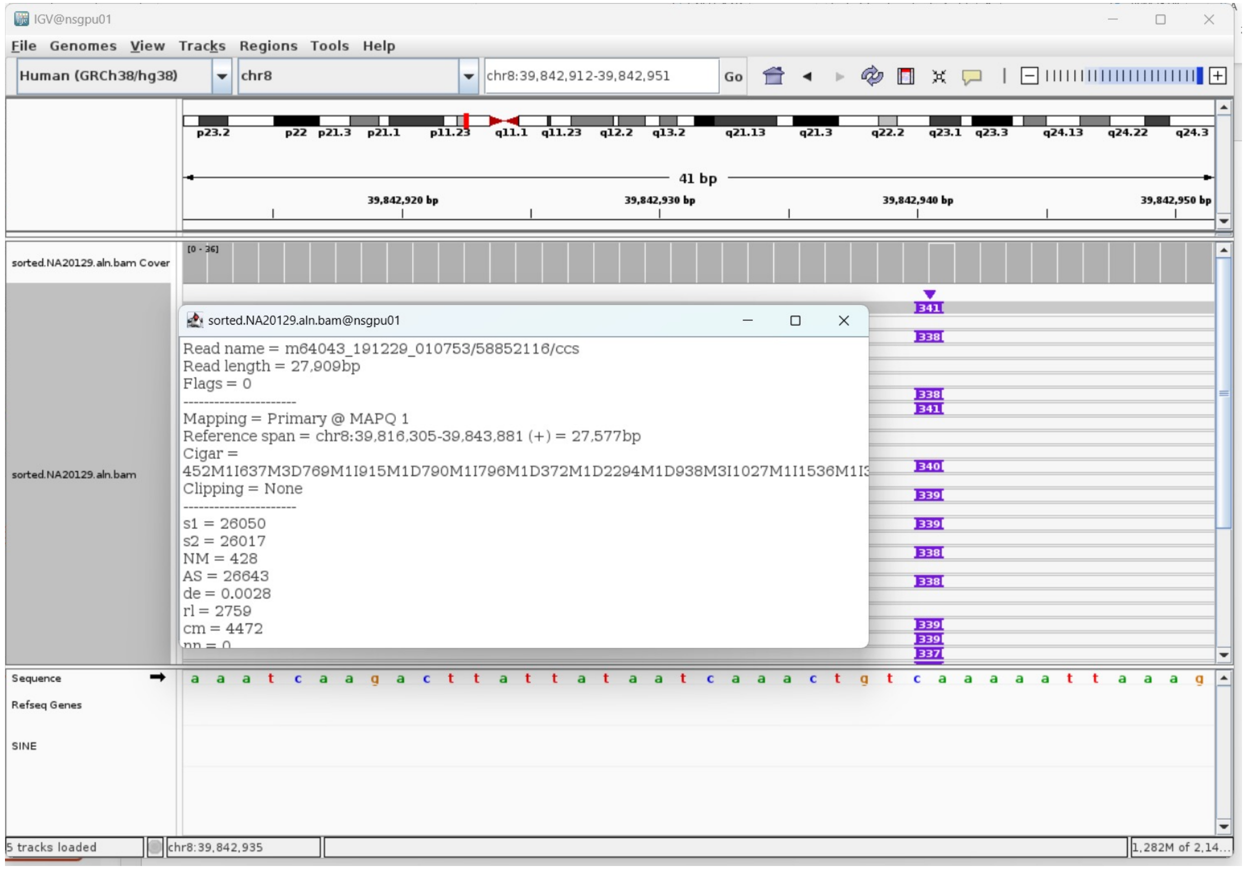

### NA20129 chr12:31253691

➢ Mapping quality is 0, indicating that this region may have alignment issues.

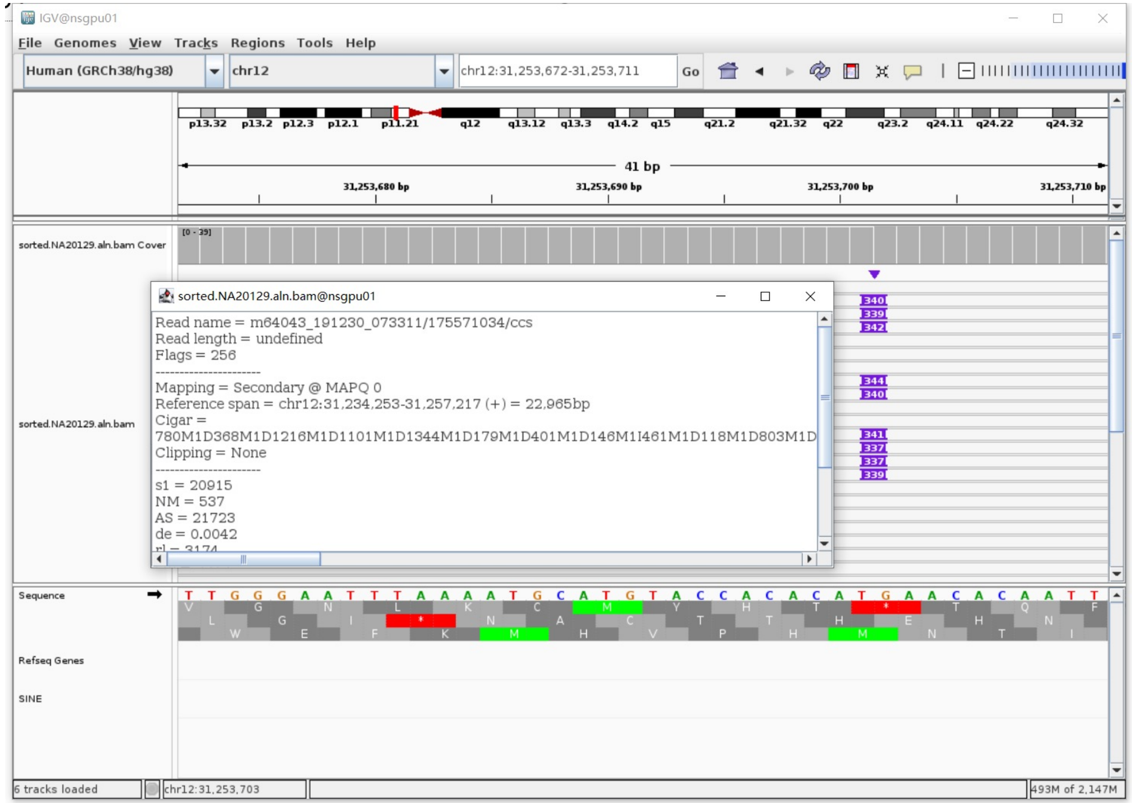


➢ Fetch one read (m64043_191230_073311/175571034/ccs, 23266 bp) and BLAT it to the hg38 genome.

The best alignment is to chr12_KI270835v1_alt (an alternative sequence of chr12) and chr12.

| ACTIONS | | | QUERY | SCORE | START | END | QSIZE | IDENTITY | CHROM | STRAND | START | END | SPAN |
|---|---|---|---|---|---|---|---|---|---|---|---|---|---|
| browser | new tab | details | YourSeq | 22769 | 1 | 23266 | 23266 | 99.7% | chr12 | + | 31234253 | 31257217 | 22965 |
| browser | new tab | details | YourSeq | 22716 | 1 | 23266 | 23266 | 99.8% | chr12_KI270835v1_alt | + | 125047 | 148011 | 22965 |

- Read:1-19519 aligns to chr12_KI270835v1_alt

| ACTIONS | | | QUERY | SCORE | START | END | QSIZE | IDENTITY | CHROM | STRAND | START | END | SPAN |
|---|---|---|---|---|---|---|---|---|---|---|---|---|---|
| browser | new tab | details | YourSeq | 19402 | 1 | 19473 | 19518 | 99.9% | chr12_KI270835v1_alt | + | 125048 | 144509 | 19462 |
| browser | new tab | details | YourSeq | 19402 | 1 | 19473 | 19518 | 99.9% | chr12 | + | 31234254 | 31253715 | 19462 |

- Read:19519-19800 aligns to *Alu*Ya5:282-1, 99.29% identity

TGAGACGGAGTCTCGCTCTGTCGCCCAGGCTGGAGTGCAGTGGCGCAATCTCGGCTCACTGCAAGCTCCGCCTC
CCGGGTTCACGCCATTCTCCTGCCTCAGCCTCCCAAGTAGCTGGGACTACAGGCGCCCGCCACTACGCCCGGCT
AATTTTTGTATTTTTAGTAGAGACGGGGTTTCACCGTTTTAGCCGGGATGGTCTCGATCTCCTGACCTCGTGAT
CCGCCCGCCTCGGCCTCCCAAAGTGCTGGGATTACAGGCGTGAGCCACCGCGCCCGGCC

14bp TSD

GAACACAATTCTTT

- Read:19801-23266 also aligns to chr12_KI270835v1_alt

| ACTIONS | | | QUERY | SCORE | START | END | QSIZE | IDENTITY | CHROM | STRAND | START | END | SPAN |
|---|---|---|---|---|---|---|---|---|---|---|---|---|---|
| browser | new tab | details | YourSeq | 3379 | 1 | 3465 | 3465 | 99.4% | chr12 | + | 31253703 | 31257217 | 3515 |
| browser | new tab | details | YourSeq | 3368 | 1 | 3465 | 3465 | 99.4% | chr12_KI270835v1_alt | + | 144497 | 148011 | 3515 |

➢ It's an *Alu*Ya5 insertion.

### NA20129 chr14:91780857

- The same genomic coordinate as in HG02630 (page 35)

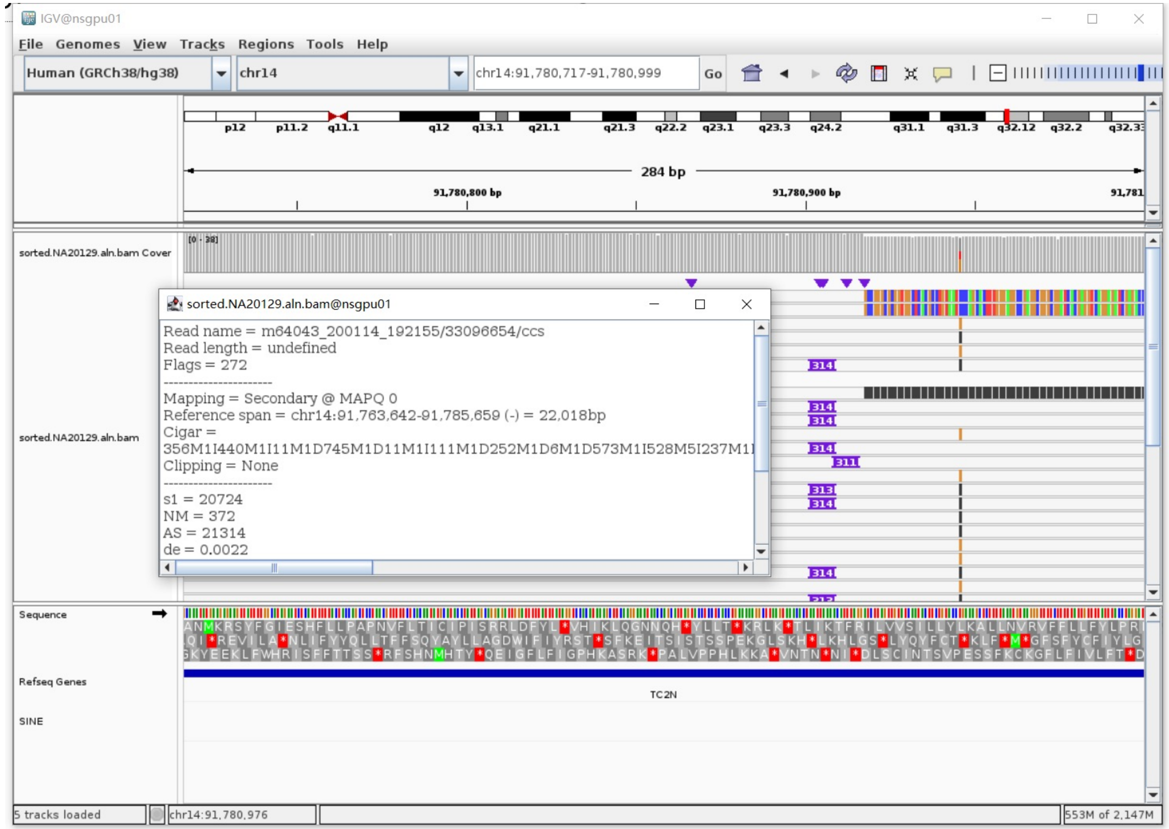

### NA20129 chr14:106651070

- ➢ The same genomic coordinate as in HG03516 (page 70)

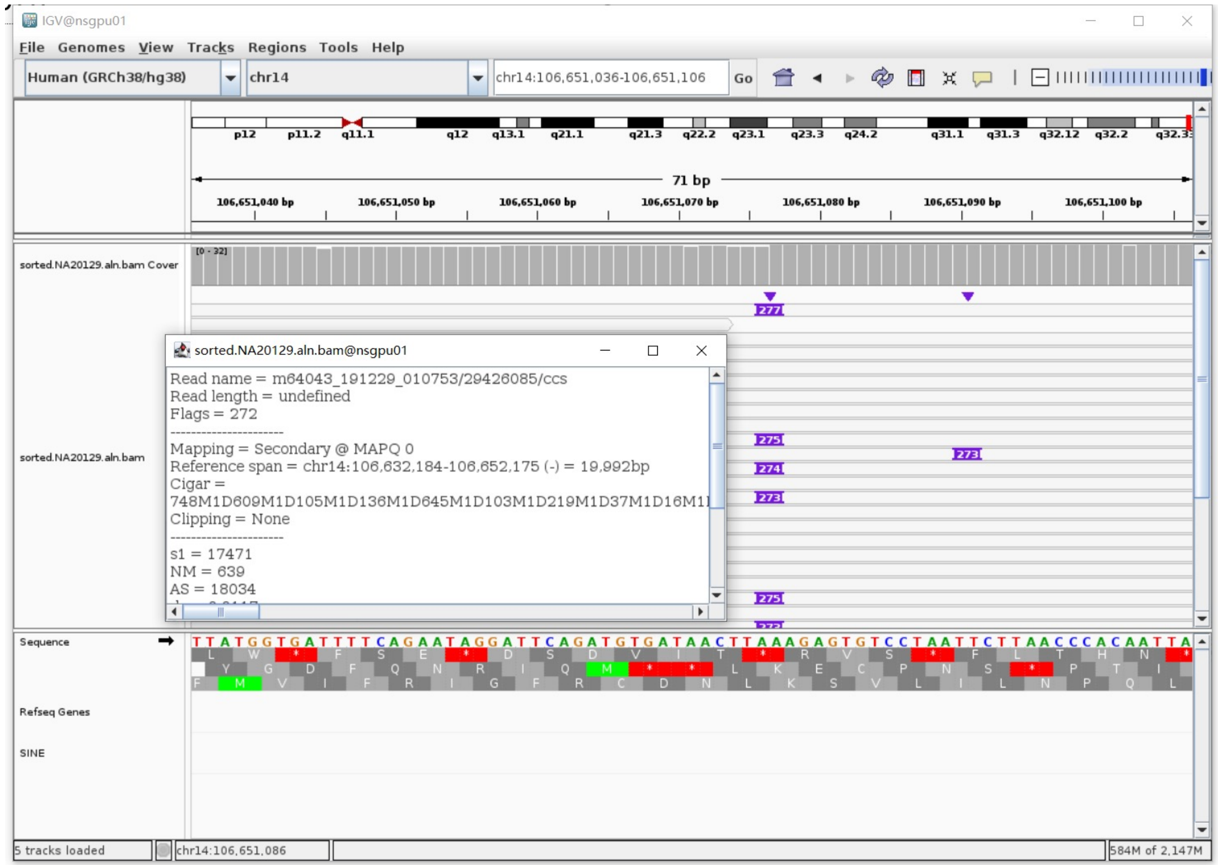

### NA20129 chr17:38594618

- ➢ Using the built-in BLAT tool in IGV to check where the inserted sequence aligned.

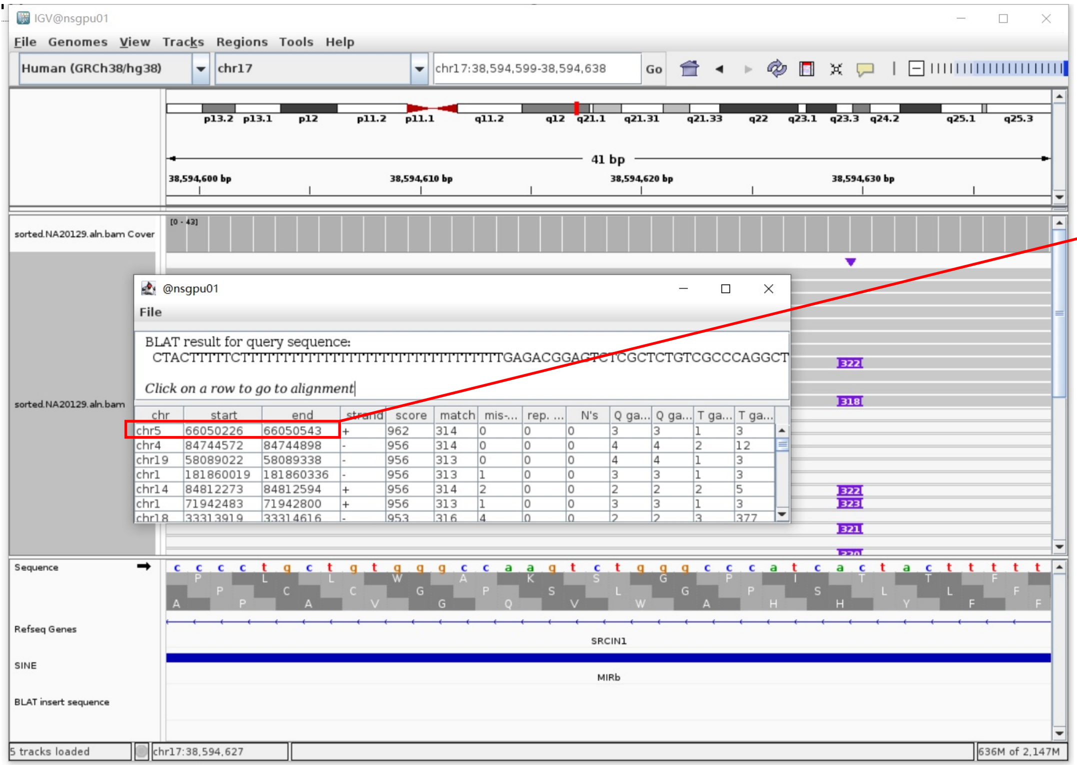


- ➢ The annotation at the aligned location indicated it as *AluY*.

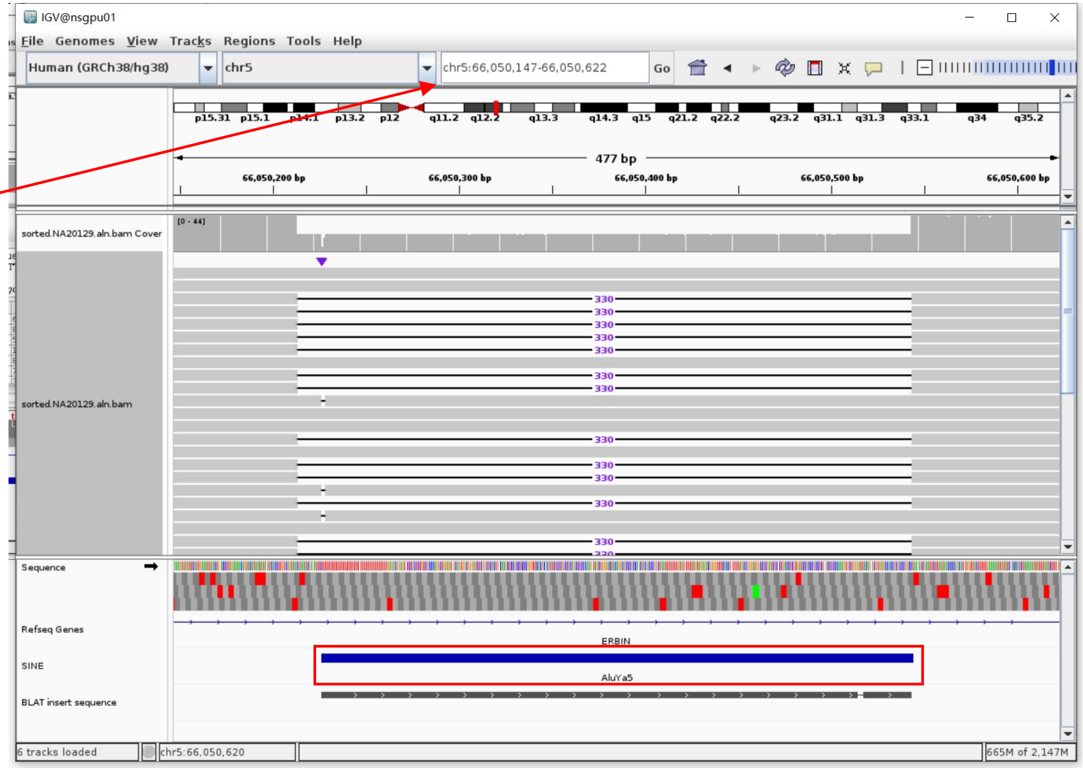


Insertion sequence (322bp)

- ➢ 13bp TSD

CTACTTTTTCTTT

- ➢ Insertion:41-322 aligns to *Alu*Ya5:282-1, 99.64% identity

TGAGACGGAGTCTCGCTCTGTCGCCCAGGCTGGAGTGCAGTGGCGGATCTCGGCTCACTGCAAGCTCCGCCTCCCGGGTTCACGCCATTCTCCTGCCTCAGCCTCCCAAGTAGCTGGGACTA
CAGGCGCCCCGCCACTACGCCCGGCTAATTTTTTTGTATTTTTAGTAGAGACGGGGTTTCACCGTTTTAGCCGGGATGGCCTCGATCTCCTGACCTCGTGATCCGCCCGCCTCGGCCTCCCAA
AGTGCTGGATACAGGCGTGAGCCACCGCGCCCGGC

### NA20129 chr18:51774994

- The same genomic coordinate as in HG00438 (page 9)

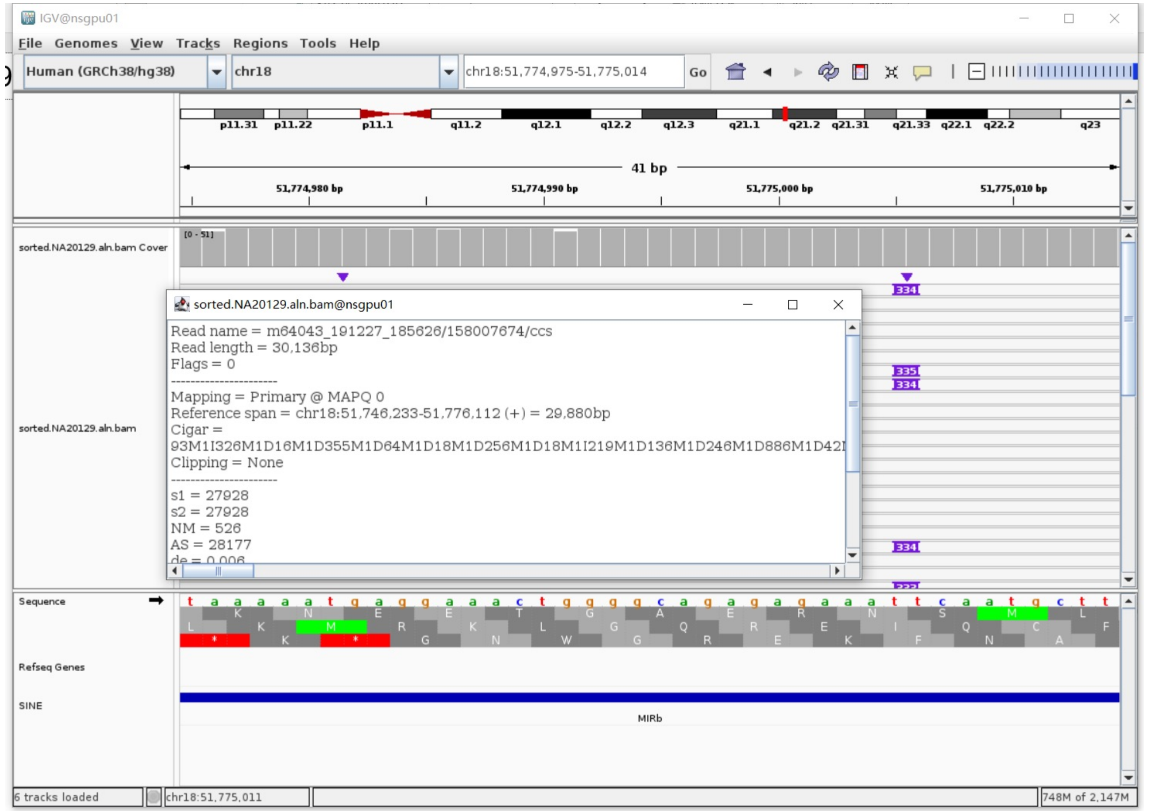

### ERX2355888 chr1:82577503

- The same genomic coordinate as in HG01106 (page 19)

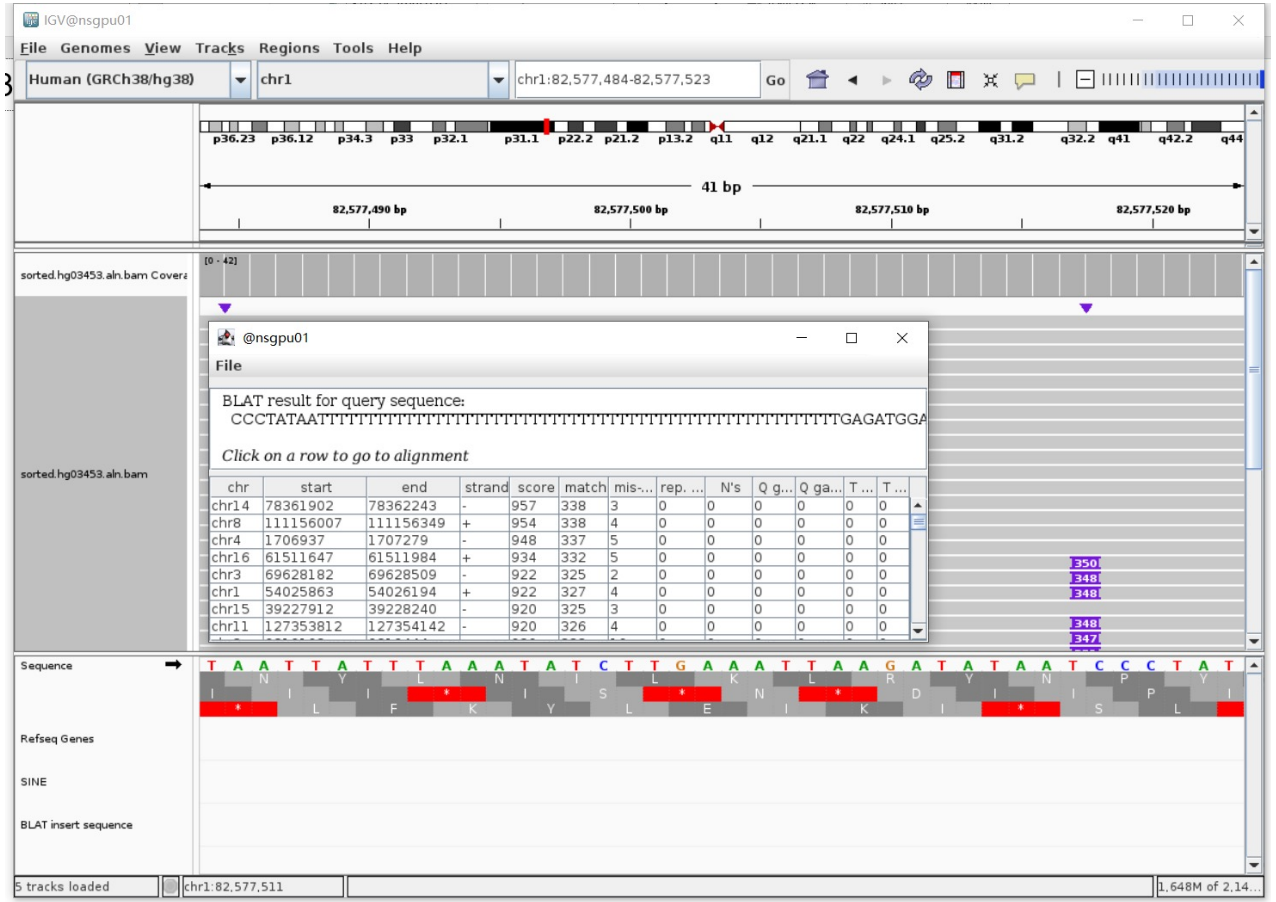


| chr | start | end | strand | score | match | mis-... | rep. ... | N's | Q g... | Q ga... | T ... | T ... |
|---|---|---|---|---|---|---|---|---|---|---|---|---|
| chr14 | 78361902 | 78362243 | - | 957 | 338 | 3 | 0 | 0 | 0 | 0 | 0 | 0 |
| chr8 | 111156007 | 111156349 | + | 954 | 338 | 4 | 0 | 0 | 0 | 0 | 0 | 0 |
| chr4 | 1706937 | 1707279 | - | 948 | 337 | 5 | 0 | 0 | 0 | 0 | 0 | 0 |
| chr16 | 61511647 | 61511984 | + | 934 | 332 | 5 | 0 | 0 | 0 | 0 | 0 | 0 |
| chr3 | 69628182 | 69628509 | - | 922 | 325 | 2 | 0 | 0 | 0 | 0 | 0 | 0 |
| chr1 | 54025863 | 54026194 | + | 922 | 327 | 4 | 0 | 0 | 0 | 0 | 0 | 0 |
| chr15 | 39227912 | 39228240 | - | 920 | 325 | 3 | 0 | 0 | 0 | 0 | 0 | 0 |
| chr11 | 127353812 | 127354142 | - | 920 | 326 | 4 | 0 | 0 | 0 | 0 | 0 | 0 |

### ERX2355888  chr2:36249528

➢ The same genomic coordinate as in NA18906 (page 94)

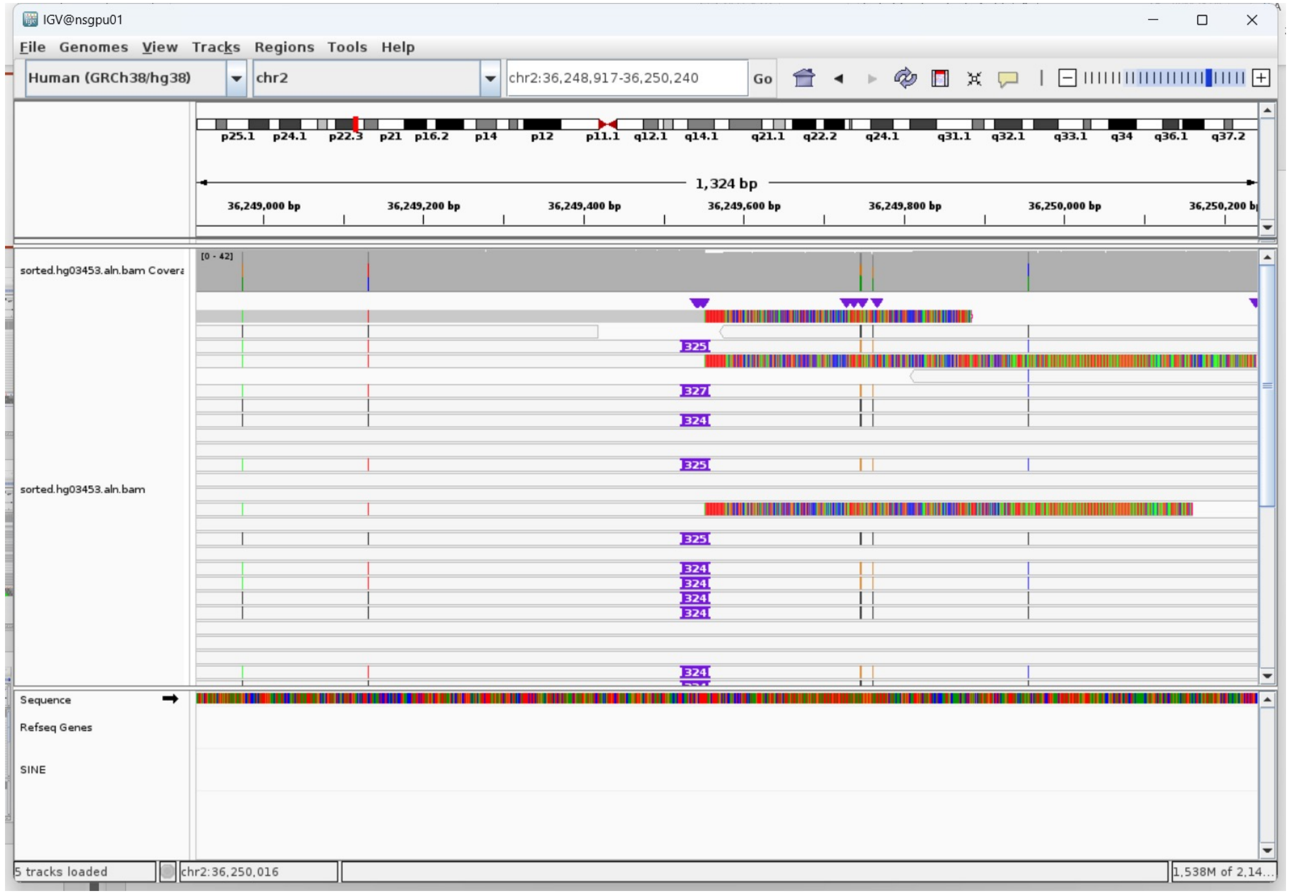

### ERX2355888 chr6:30030695

- The same genomic coordinate as in HG03492 (page 47)

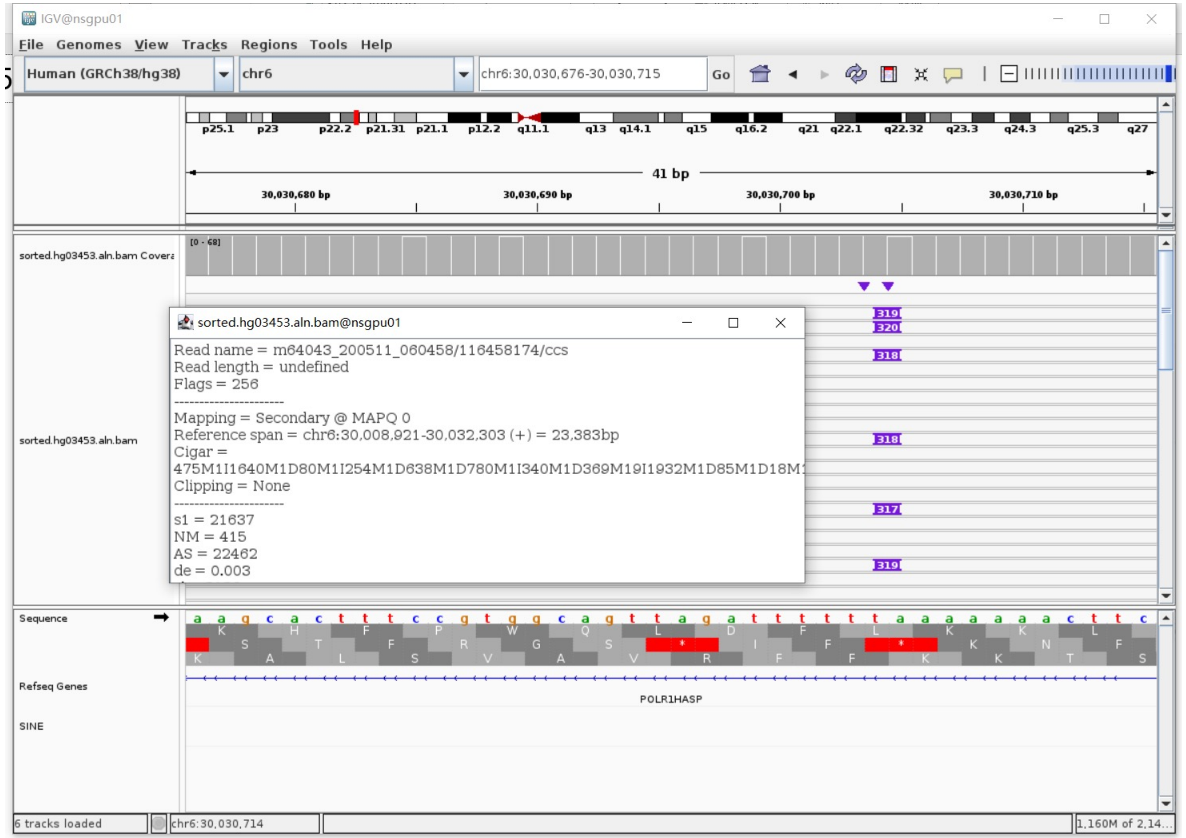

### ERX2355888 chr6:32805636

➢ The same genomic coordinate as in HG00438 (page 5)

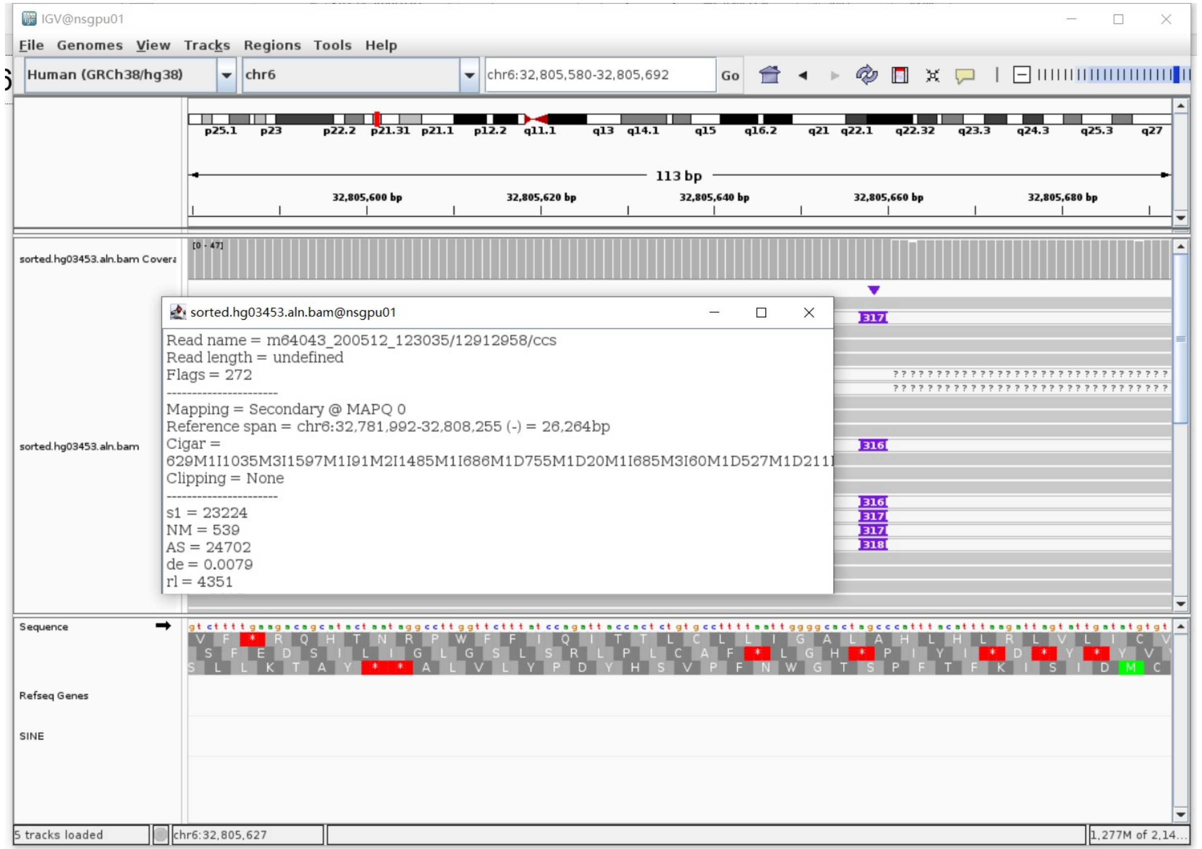

### ERX2355888 chr6:33235423

- Mapping quality is 0, indicating that this region may have alignment issues.

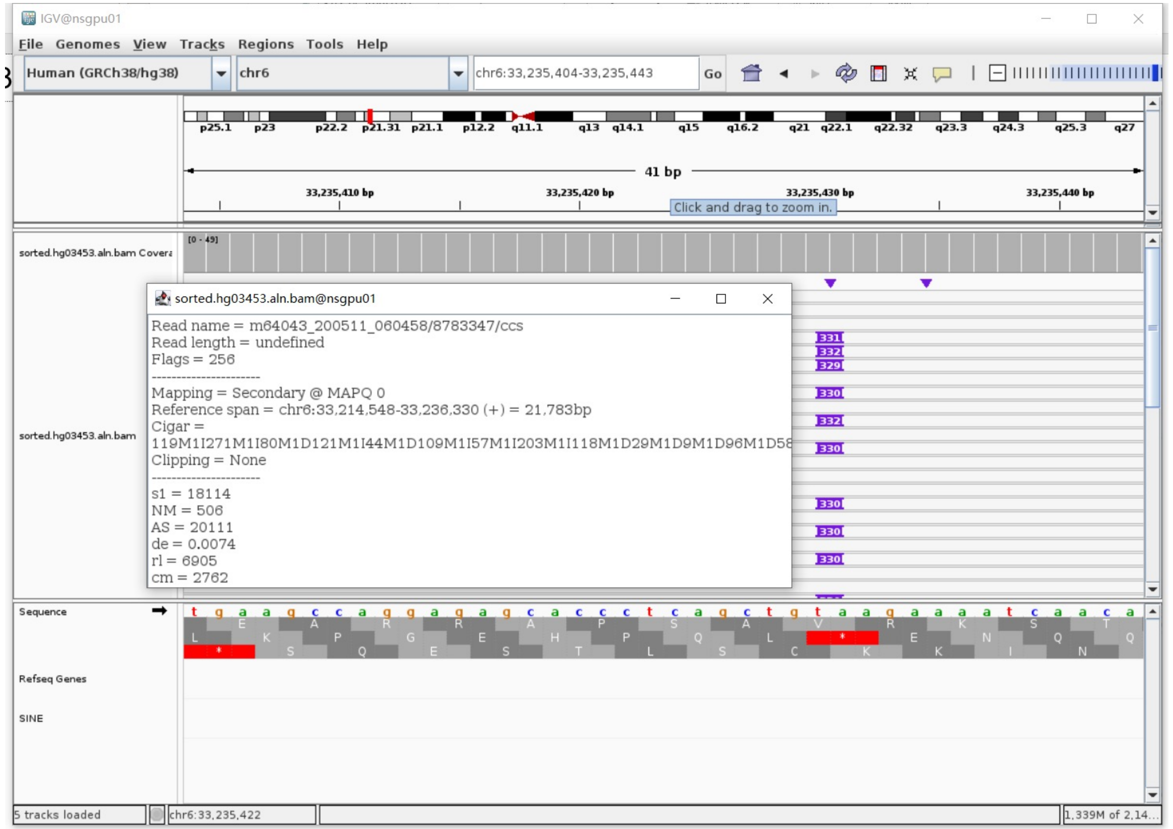


- Fetch one read (m64043_200509_233929/166857291/ccs, 23584 bp) and BLAT it to the hg38 genome.

  The best alignment is to chr6_GL000252v2_alt (an alternative sequence of chr6), which contains an *Alu*Yb8 element.

| ACTIONS | | | QUERY | SCORE | START | END | QSIZE | IDENTITY | CHROM | STRAND | START | END | SPAN |
|---|---|---|---|---|---|---|---|---|---|---|---|---|---|
| browser | new tab | details | YourSeq | 21375 | 1 | 21446 | 21446 | 99.9% | chr6_GL000252v2_alt | - | 1271338 | 1292801 | 21464 |
| browser | new tab | details | YourSeq | 21375 | 1 | 21446 | 21446 | 99.9% | chr6_GL000250v2_alt | - | 1273512 | 1294975 | 21464 |
| browser | new tab | details | YourSeq | 21373 | 1 | 21446 | 21446 | 99.9% | chr6_GL000251v2_alt | - | 1495188 | 1516650 | 21463 |
| browser | new tab | details | YourSeq | 21364 | 1 | 21446 | 21446 | 99.9% | chr6_GL000256v2_alt | - | 1313610 | 1335811 | 22202 |
| browser | new tab | details | YourSeq | 21044 | 1 | 21446 | 21446 | 99.8% | chr6_GL000254v2_alt | - | 1359859 | 1380989 | 21131 |
| browser | new tab | details | YourSeq | 21038 | 1 | 21446 | 21446 | 99.8% | chr6_GL000253v2_alt | - | 1276534 | 1297665 | 21132 |
| browser | new tab | details | YourSeq | 21038 | 1 | 21446 | 21446 | 99.8% | chr6 | - | 30015785 | 30036916 | 21132 |

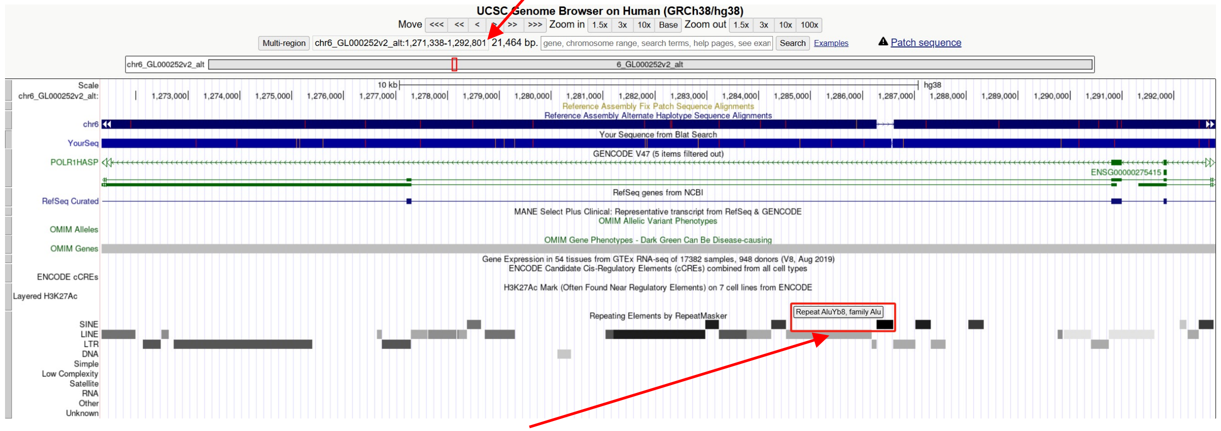


chr6_GL000252v2_alt:1286275-1286592 (*Alu*Y marked by RepeatMasker)

- Read:6223-6512 aligns to *Alu*Yb8:289-1, 99.65% identity

TGAGACGGAGTCTCGCTCTGTCGCCCAGGCCGTACTGCGGACTGCAGTGGCGCAATCTCGGCTCACTGCAAGCT
CCGCTTCCCGGGTTCACGCCATTCTCCTGCCTCAGCCTCCCGAGTAGCTGGGACTACAGGCGCCCGCCACCGCG
CCCGGCTAATTTTTTGTATTTTTAGTAGAGACGGGGTTTCACCTTGTTAGCCAGGATGGTCTCGATCTCCTGAC
CTCATGATCCACCCGCCTCGGCCTCCCAAAGTGCTGGGATTACAGGCGTGAGCCACCGCGCCCGGCC

- It's an *Alu*Yb1 in the chromosome chr6_GL000252v2_alt.

### ERX2355888 chr6:79370732

- Mapping quality is 0, indicating that this region may have alignment issues.

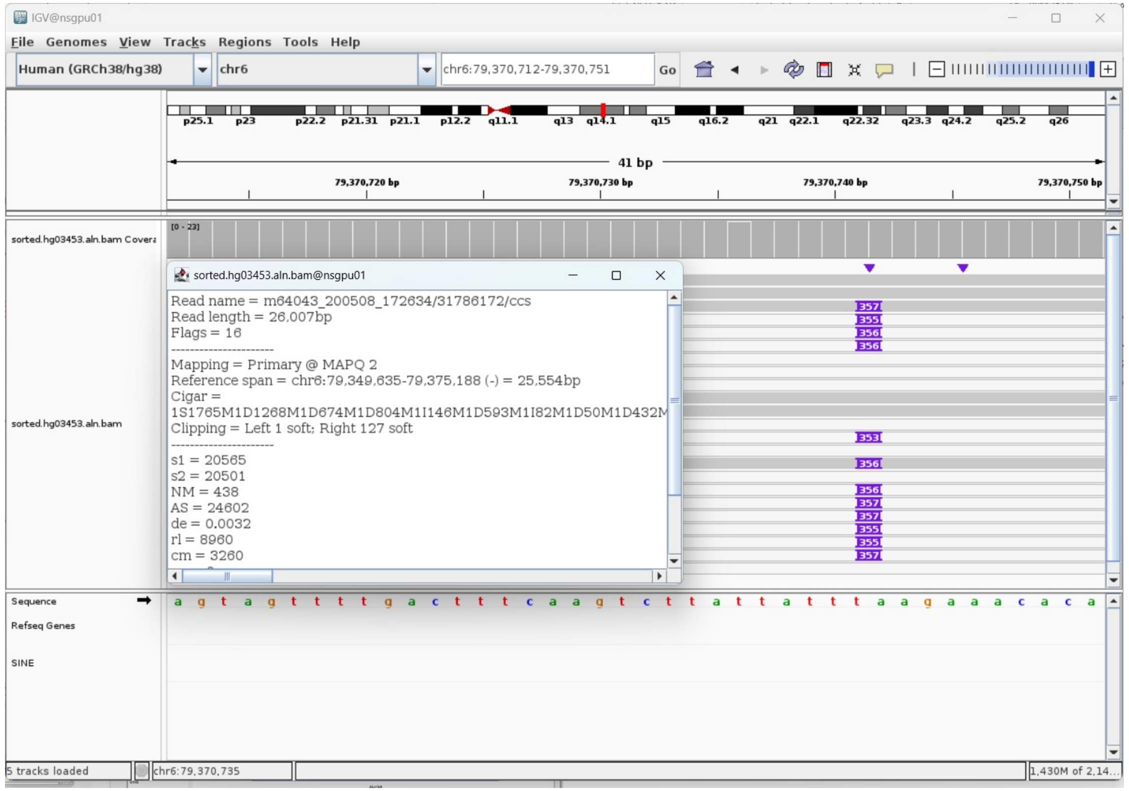


- Fetch one read (m64043_200508_172634/31786172/ccs, 26034 bp) and BLAT it to the hg38 genome.

  The best alignment is to chr6_GL383533v1_alt (an alternative sequence of chr6) and chr6.

| ACTIONS | QUERY | SCORE | START | END | QSIZE | IDENTITY | CHROM | STRAND | START | END | SPAN |
|---|---|---|---|---|---|---|---|---|---|---|---|
| browser new tab details | YourSeq | 25450 | 129 | 26034 | 26034 | 99.8% | chr6 | - | 79349635 | 79375188 | 25554 |
| browser new tab details | YourSeq | 25205 | 1 | 25661 | 26034 | 99.8% | chr6_GL383533v1_alt | + | 99428 | 124736 | 25309 |

- Read:1-4623 aligns to chr6_GL383533v1_alt and chr6

| ACTIONS | QUERY | SCORE | START | END | QSIZE | IDENTITY | CHROM | STRAND | START | END | SPAN |
|---|---|---|---|---|---|---|---|---|---|---|---|
| browser new tab details | YourSeq | 4552 | 1 | 4569 | 4622 | 99.9% | chr6_GL383533v1_alt | + | 99429 | 104002 | 4574 |
| browser new tab details | YourSeq | 4425 | 128 | 4569 | 4622 | 99.9% | chr6 | - | 79370742 | 79375188 | 4447 |

- Read:4624-4913 aligns to *Alu*Yb8:289-1, 98.96% identity

TGAGACGGAGTCTCGCTCTGTCGCCCAGGCCAGACTGCGGACTGCAGTGGCGCAATCTCGGCTCACTGCAAGC
TCCACTTCCTGGGTTCACGCCATTCTCCTGCCTCAGCCTCCCGAGTAGCTGGGACTACAGGCGCCCGCCACCG
CGCCCGGCTAATTTTTTGTATTTTTAGTAGAGACGGGGTTTCACCTTGTTAGCCAGGATGGTCTCGATCTCCT
GACCTCATGATCCACCCGCCTCGGCCTCCCAAAGTGCTGGGATTACAGGCGTGAGCCACCGCGCCCGGCC

13bp TSD

TGAAATGTGTTTC

- Read:4914-26034 also aligns to chr6_GL383533v1_alt and chr6

| ACTIONS | QUERY | SCORE | START | END | QSIZE | IDENTITY | CHROM | STRAND | START | END | SPAN |
|---|---|---|---|---|---|---|---|---|---|---|---|
| browser new tab details | YourSeq | 21041 | 1 | 21120 | 21120 | 99.9% | chr6 | - | 79349635 | 79370756 | 21122 |
| browser new tab details | YourSeq | 20668 | 1 | 20747 | 21120 | 99.9% | chr6_GL383533v1_alt | + | 103988 | 124736 | 20749 |

- It's an *Alu*Yb8 insertion.

### ERX2355888 chr6:104041958

➢ The same genomic coordinate as in HG00438 (page 6)

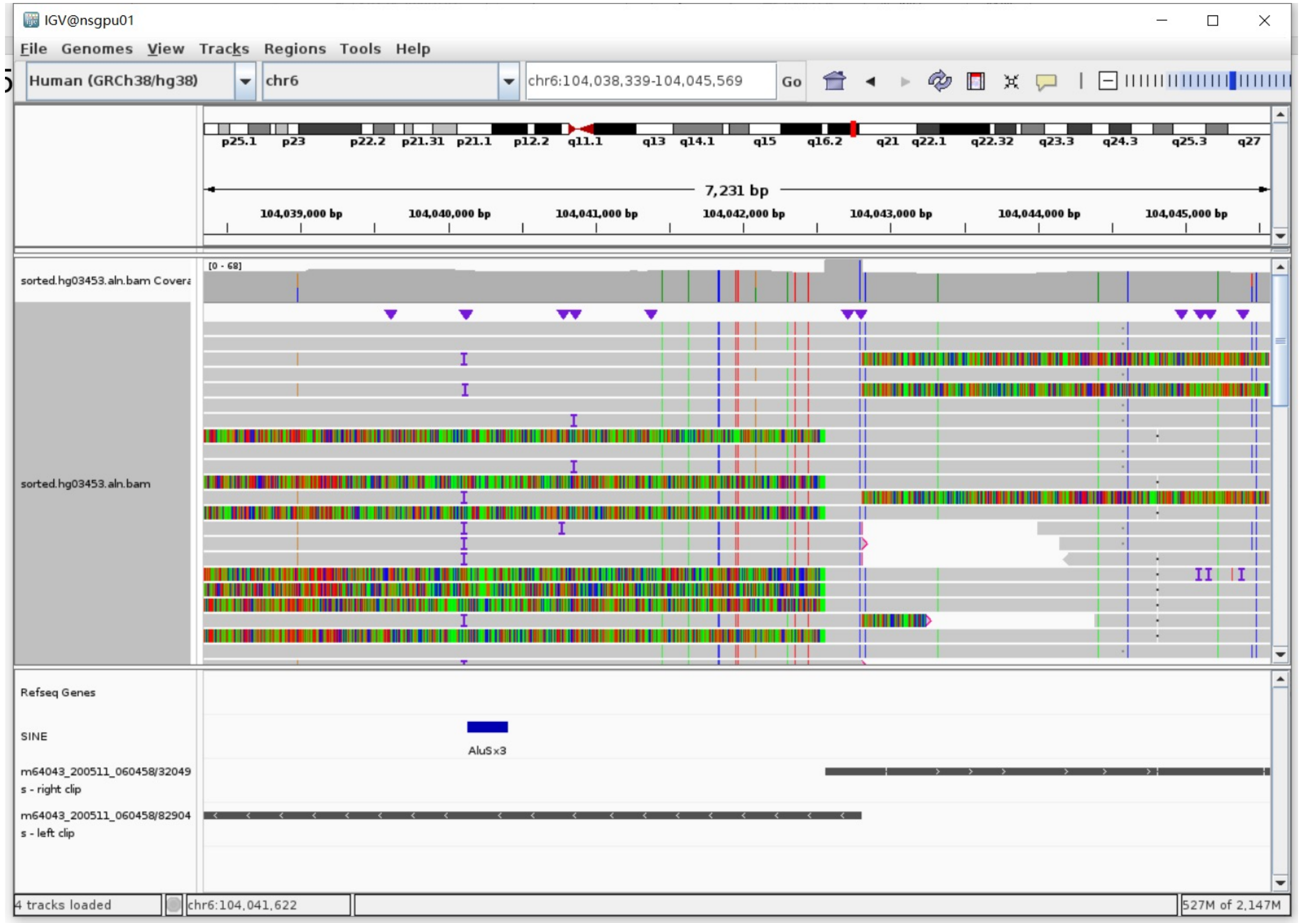

### ERX2355888  chr7:142906943

- ➢ Using the built-in BLAT tool in IGV to check where the inserted sequence aligned.
- ➢ The annotation at the aligned location indicated it as *AluY*.

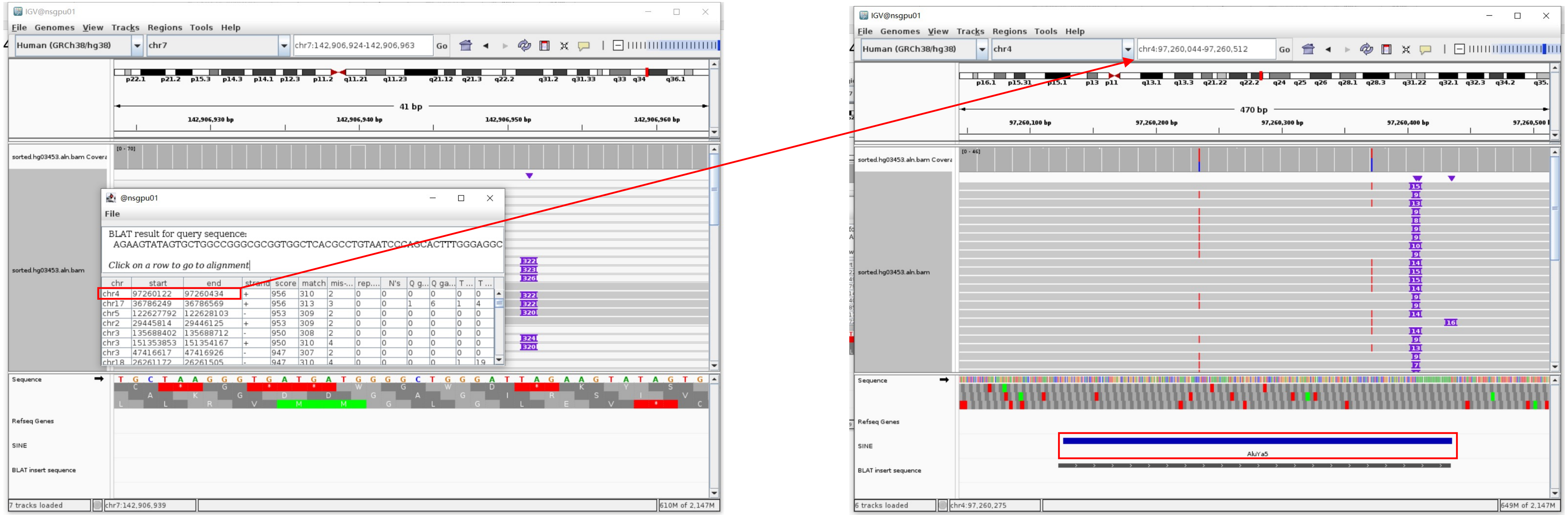


Insertion sequence (320bp)

- ➢ 16bp TSD

AGAAGTATAGTGCTGG

- ➢ Insertion:14-296 aligns to *AluYa5*:1-282, 99.65% identity

GGCCGGGCGCGGTGGCTCACGCCTGTAATCCCAGCACTTTGGGAGGCCGAGGCGGGCGGATCACGAGGTCAGGAGATCGAGACCATCCCGGCTAAAACGGTGAAACCCCGTCTCTACTAAAAA
TACAAAAAATTAGCCGGGCGTAGTGGCGGGCACCTGTAGTCCCAGCTACTTGGGAGGCTGAGGCAGGAGAATGGCGTGAACCCGGGAGGCGGAGCTTGCAGTGAGCCGAGATCCCGCCACTGC
ACTCCAGCCTGGGCGACAGAGCGAGACTCCGTCTCA………poly(A)

### ERX2355888 chr9:111517769

- Using the built-in BLAT tool in IGV to check where the inserted sequence aligned.

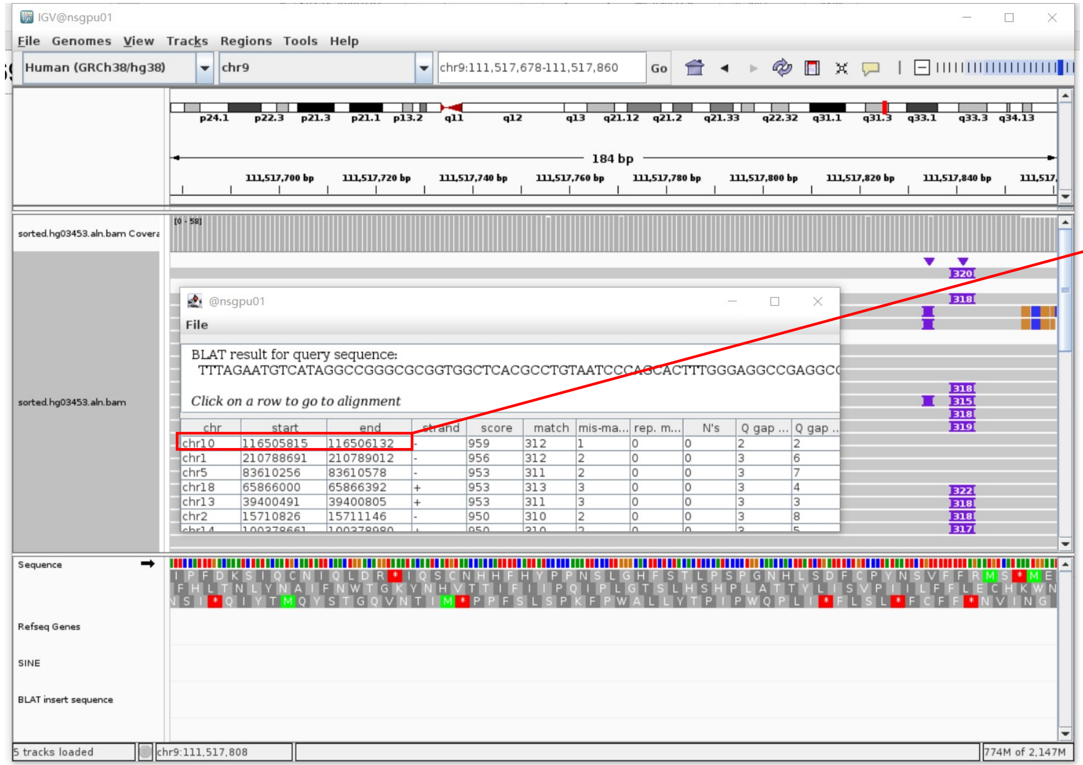


- The annotation at the aligned location indicated it as *AluY*.

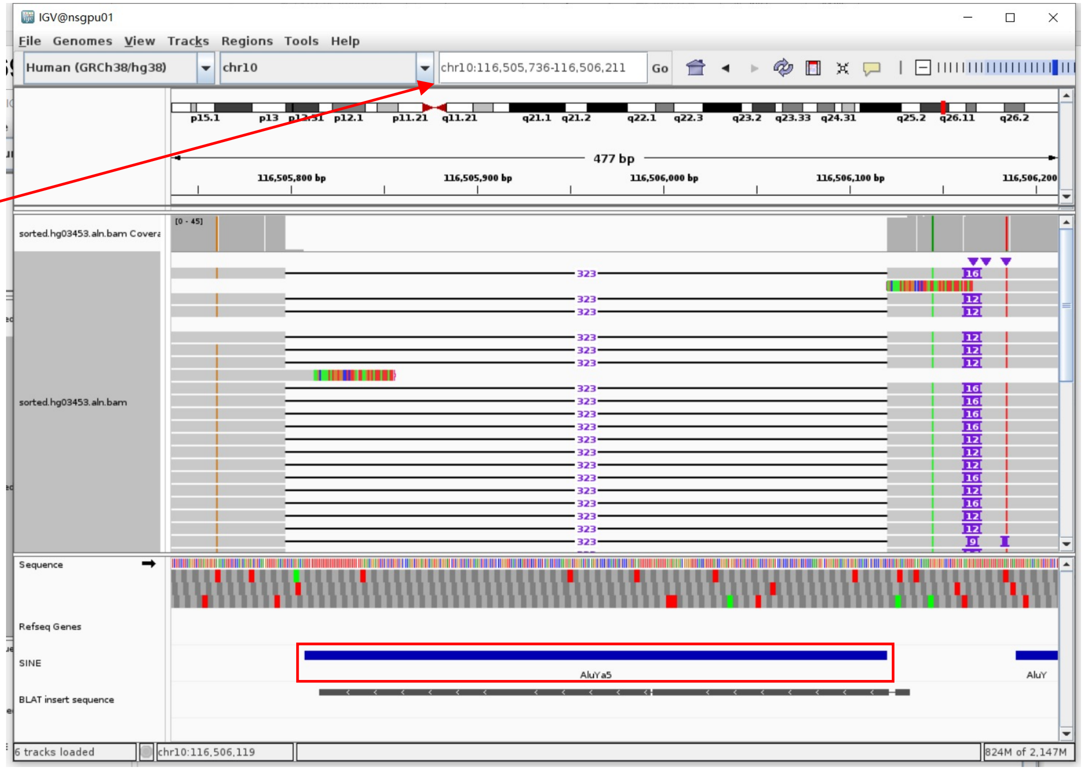


Insertion sequence (315bp)

- 14bp TSD

TTTAGAATGTCATA

- Insertion:14-298 aligns to *Alu*Ya5:1-282, 99.65% identity

GGCCGGGCGCGGTGGCTCACGCCTGTAATCCCAGCACTTTGGGAGGCCGAGGCGGGCGGATCACGAGGTCAAGAGATCGAGACCATCCCGGCTAAAACGGTGAAACCCCGTCTTCTACTAAAA
ATACAAAAAATTAGCCGGGCGTAGTGGCGGGCGCCTGTAGTCCCAGCTACTTGGGAGGCTGAGGCAGGAGAATGGCGTGAACCCCGGGAGGCGGAGCTTGCAGTGAGCCGAGATCCCGCCACT
GCACTCCAGCCTGGGCGACAGAGCGAGACTCCGTCTCA.........poly(A)

### ERX2355888 chr9:111518090

- Using the built-in BLAT tool in IGV to check where the inserted sequence aligned.
- The annotation at the aligned location indicated it as *AluY*.

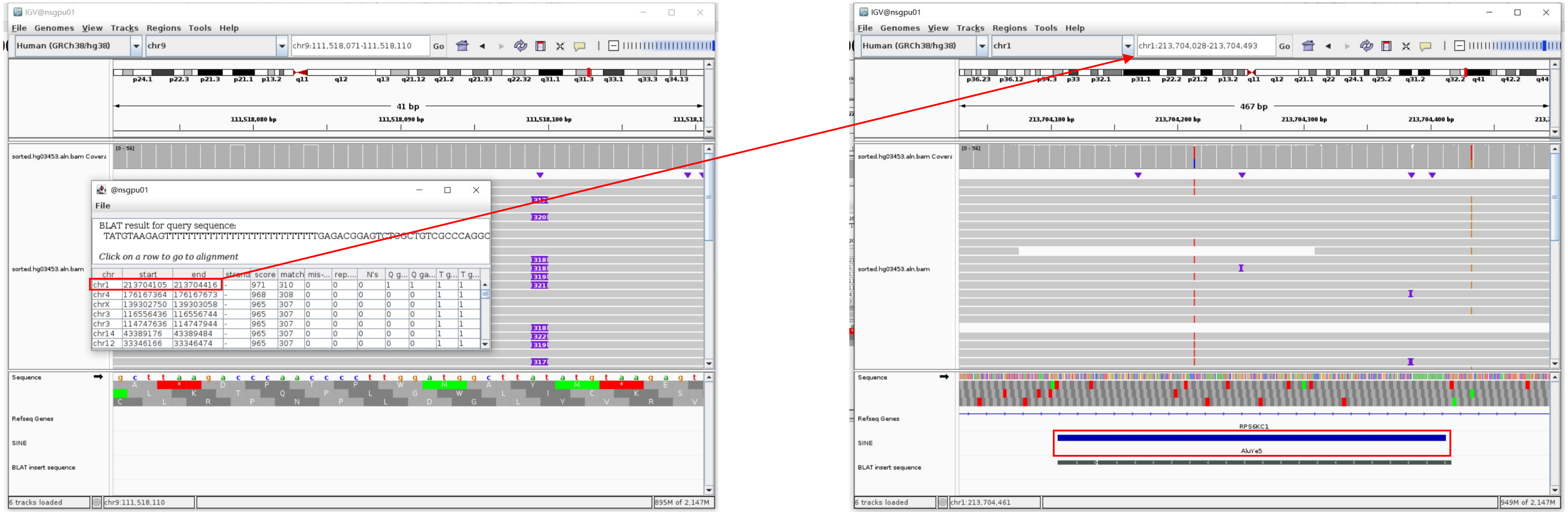


Insertion sequence (317bp)

- 15bp TSD

TATGTAAGAGTTTTT

- Insertion:37-317 aligns to *AluY*c1:282-1, 97.85% identity

TGAGACGGAGTCTCGCTGTCGCCCAGGCTGGAGTGCAGTGGCGCAATCTCGGCTCACTGCAGGCTCCGCCCCCTGGGGTTCACGCCATTCTCCTGCCTCAGCCTCCCGAGTAGCTGGGACTAC
AGGCGCCCGCCACCTCGCCCGGCTAATTTTTTGTATTTTTAGTAGAGACGGGGTTTCACCGTGTTAGCCAGGATGGTCTCGATCTCCTGACCTCGTGATCCGCCCGCCTCGGCCTCCCAAAGT
GCTGGATTACAGGCGTGAGCCACCGCGCCCGGCC

### ERX2355888 chr12:11026415

➤ The same genomic coordinate as in HG00621 (page 12)

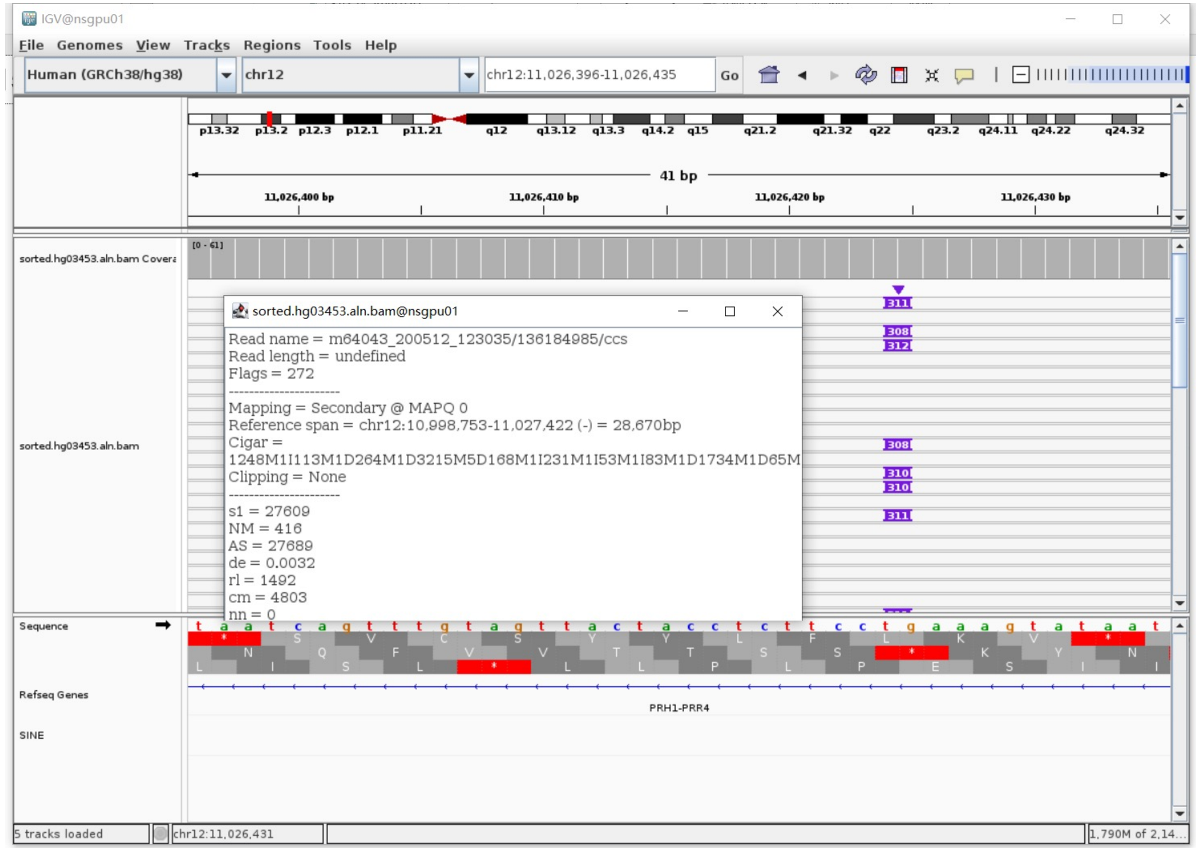

### ERX2355888 chr12:31253677

➢ The same genomic coordinate as in NA20129 (page 12)

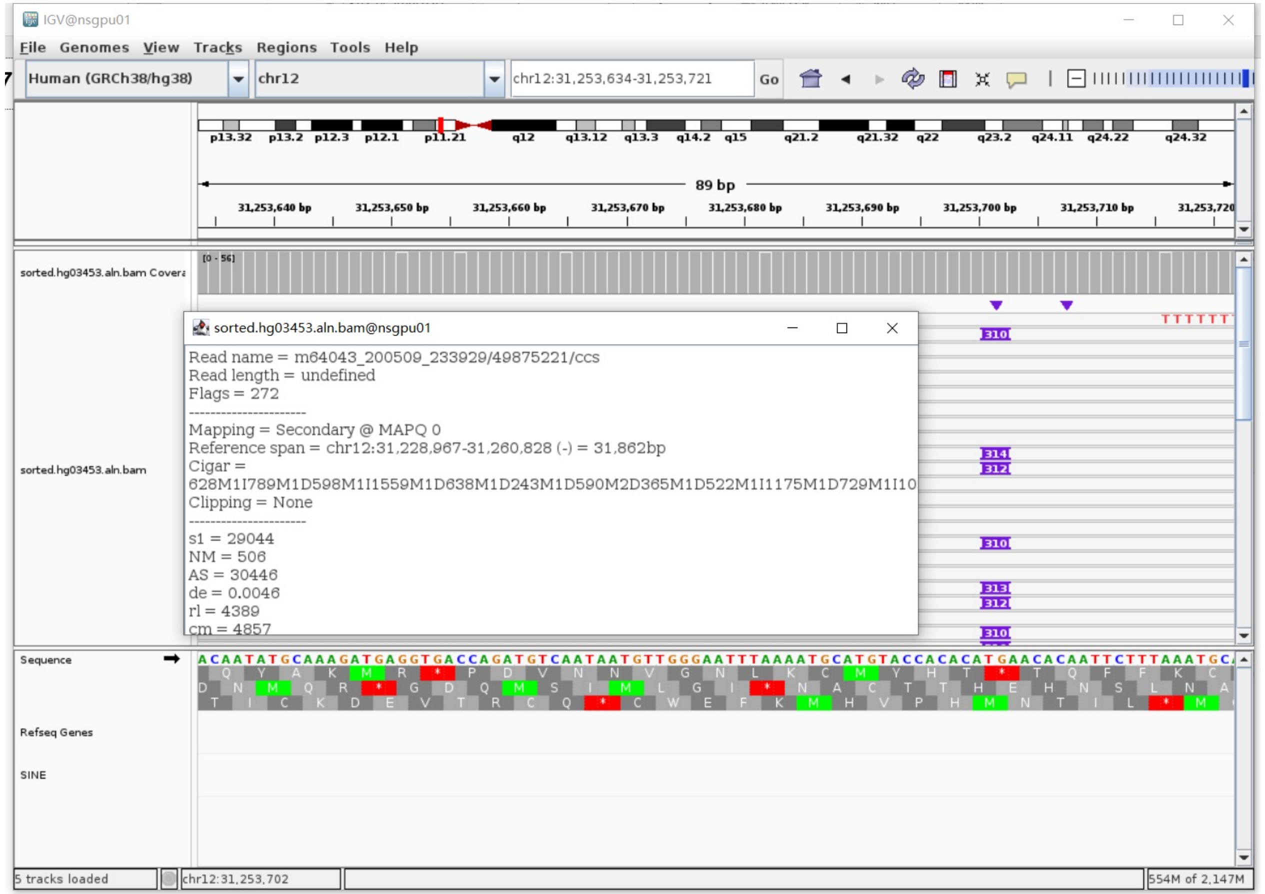

### ERX2355888 chr12:57965264

- The same genomic coordinate as in NA18906 (page 101)

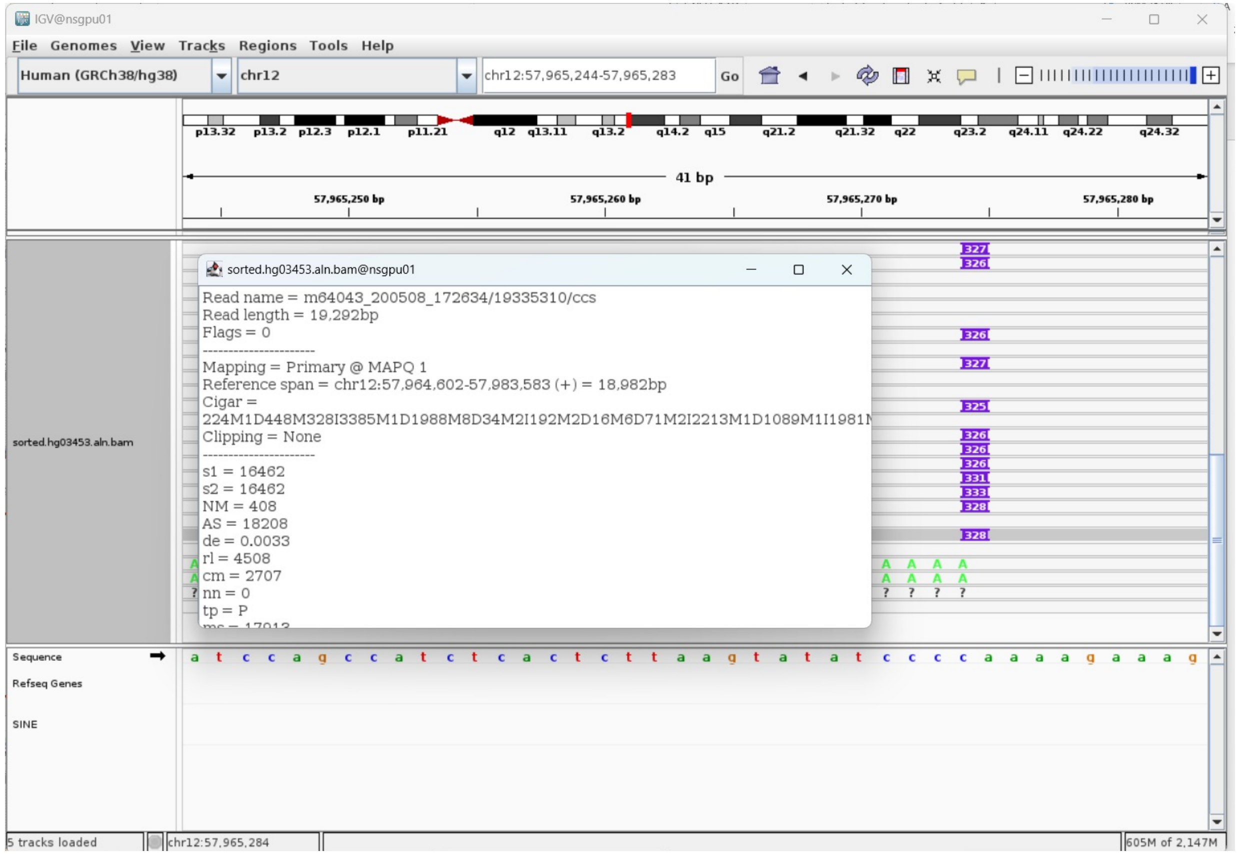

### ERX2355888 chr14:93364571

- The same genomic coordinate as in HG00438 (page 7)

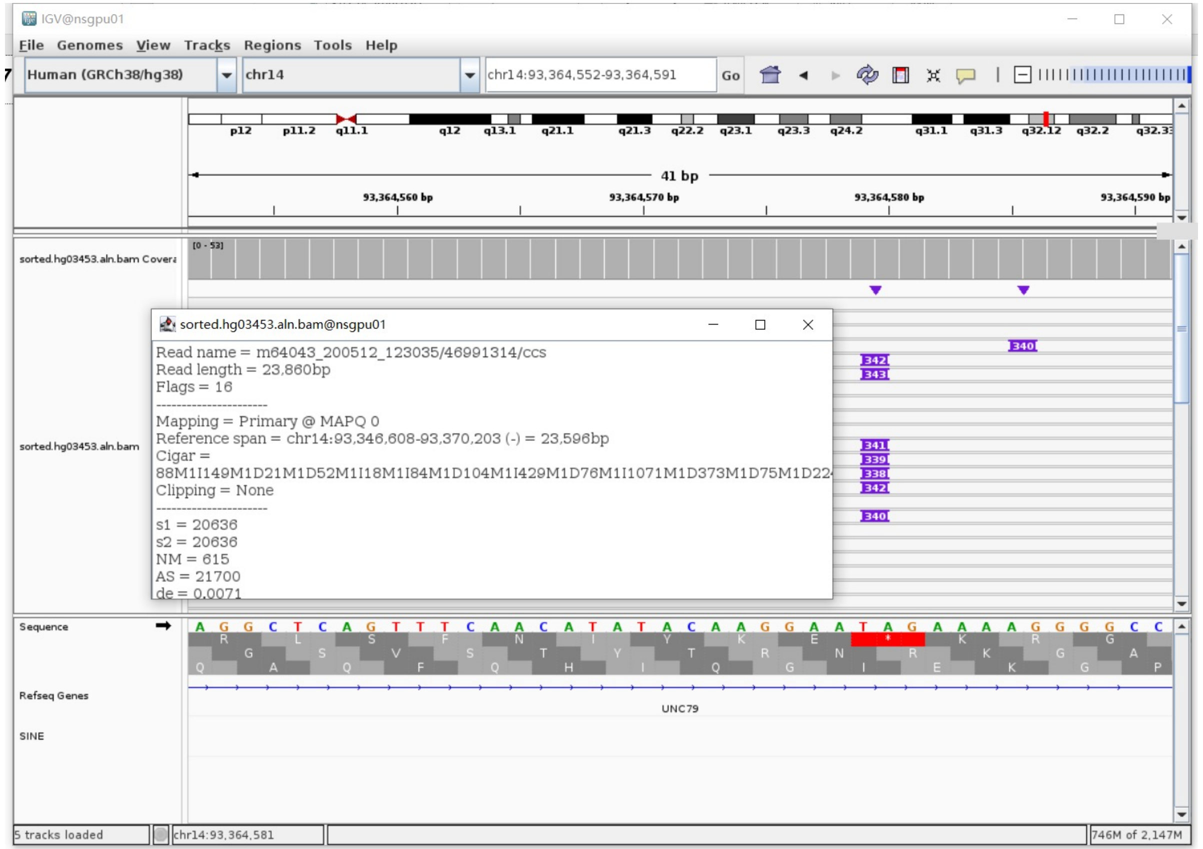

### ERX2355888 chr17:37561901

- The same genomic coordinate as in HG00621 (page 15)

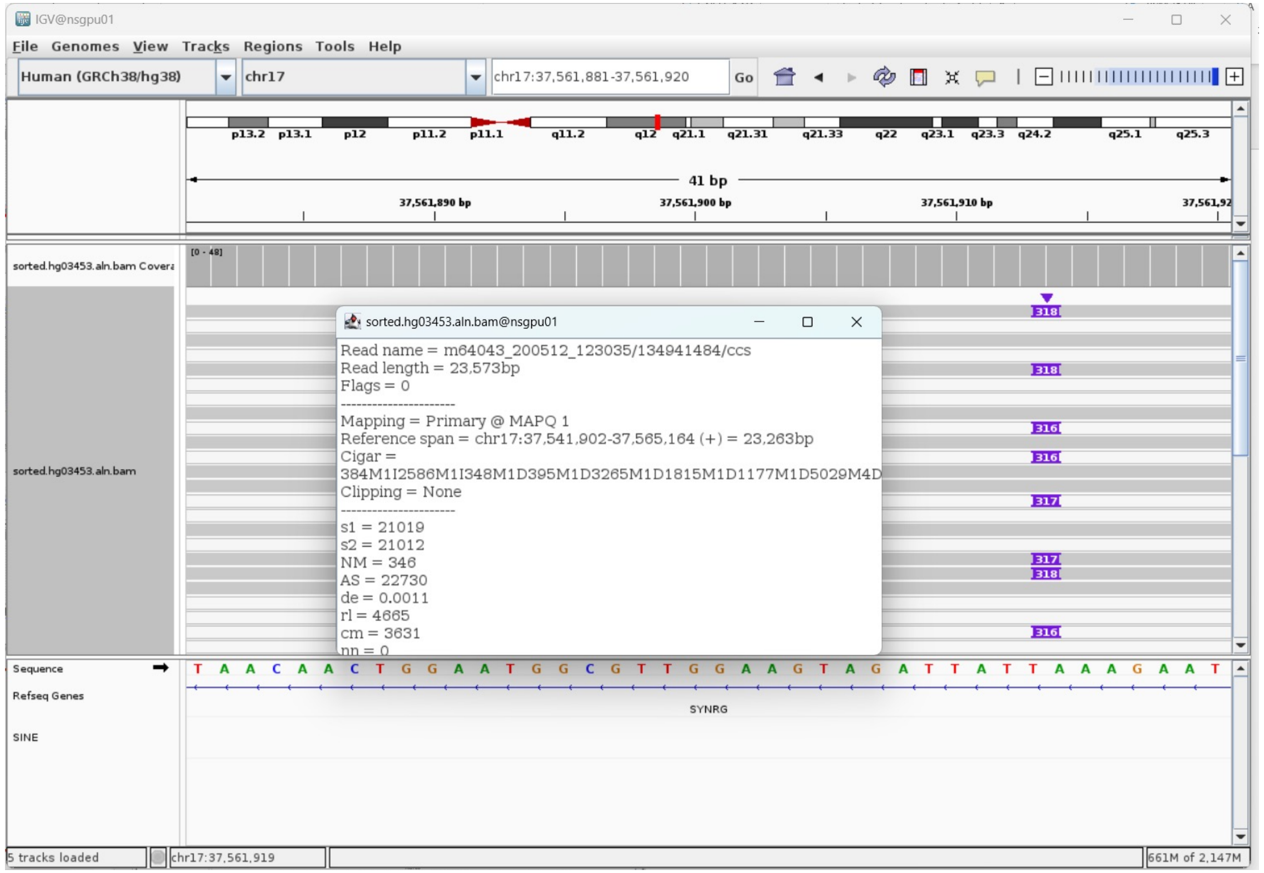

### ERX2355888 chr18:67471609

- ➢ Using the built-in BLAT tool in IGV to check where the inserted sequence aligned.
- ➢ The annotation at the aligned location indicated it as *AluY*.

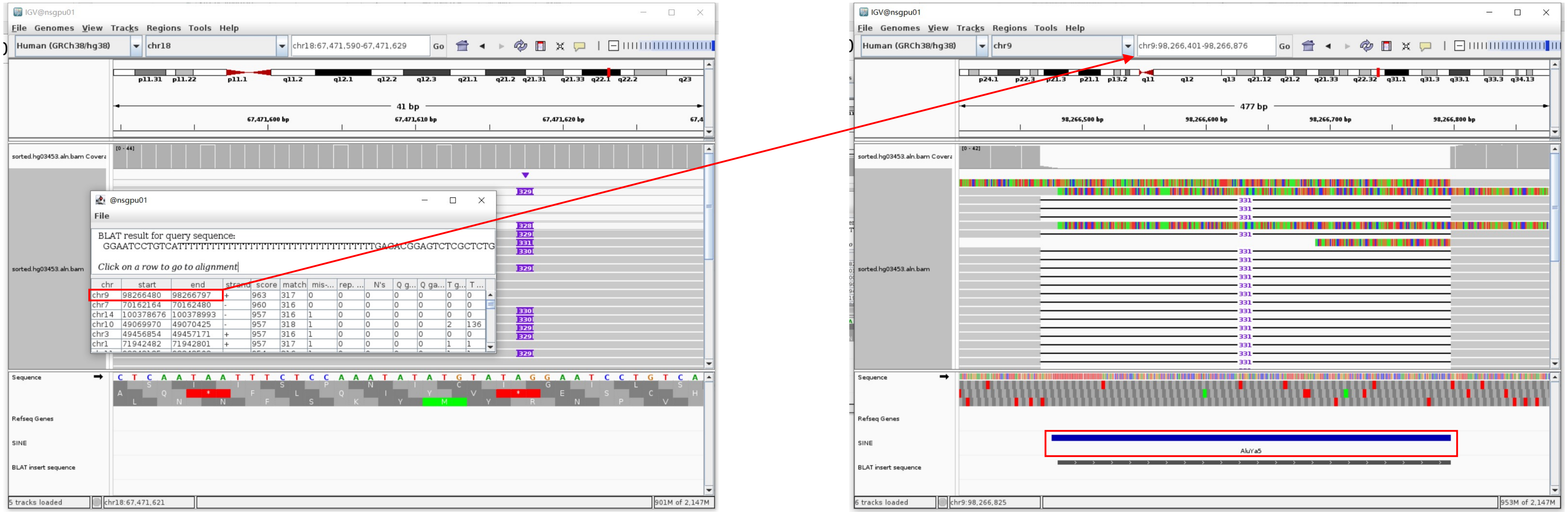


Insertion sequence (330bp)

- ➢ 16bp TSD

GGAATCCTGTCATTTT

- ➢ Insert:47-330 aligns to *AluYa5*:282-1, 99.65% identity

TGAGACGGAGTCTCGCTCTGTCTGCCCAGGCTGGAGTGCAGTGGCGGGATCTCGGCTCACTGCAAGCTCCGCCTCCCGGGTTCATGCCATTCTCCTGCCTCAGCCTCCCAAGTAGCTGGGACT
ACAGGCGCCCGCCACTACGCCCGGCTAATTTTTTGTATTTTTAGTAGAGACGGGGTTTCACCGTTTTAGCCGGGATGGTCTCGATCTCCTGACCTCGTGATCCGCCCGCCTCGGCCTCCCAAA
GTGCTGGGATTACAGGCGTGAGCCACCGCGCCCGGCC

### ERX2355888 chr22:42237075

- ➢ Using the built-in BLAT tool in IGV to check where the inserted sequence aligned.
- ➢ The annotation at the aligned location indicated it as *AluY*.

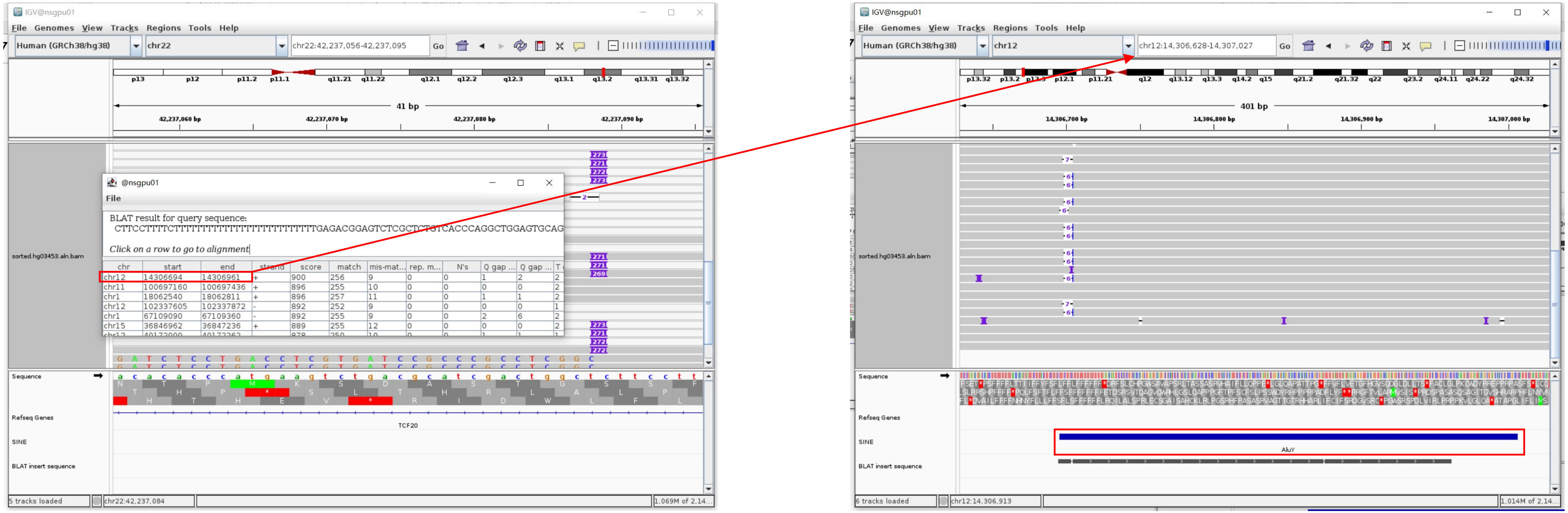


Insertion sequence (271bp)

- ➢ 11bp TSD

CTTCCTTTTCT

- ➢ Insertion:35-271 aligns to *Alu*Yc1:282-45, 98.73% identity

TGAGACGGAGTCTCGCTCTGTCACCCAGGCTGGAGTGCAGTGGCGCCATCTCGGCTCACTGCAAGCTCCGCCTCCCGGGTTCACGCCATTCTCCTGCCTCAGCCTCCCGAGTAGCTGGGACTA
CAGGCGCCCGCCACCACGCCCGGCTAATTTTTGTATTTTTAGTAGAGACGGGGTTTCACCGTGTTAGCCAGGATGGTCTCGATCTCCTGACCTCGTGATCCGCCCGCCTCGGC

### ERX2355869 chr1:82577503

- ➢ The same genomic coordinate as in HG01106 (page 19)

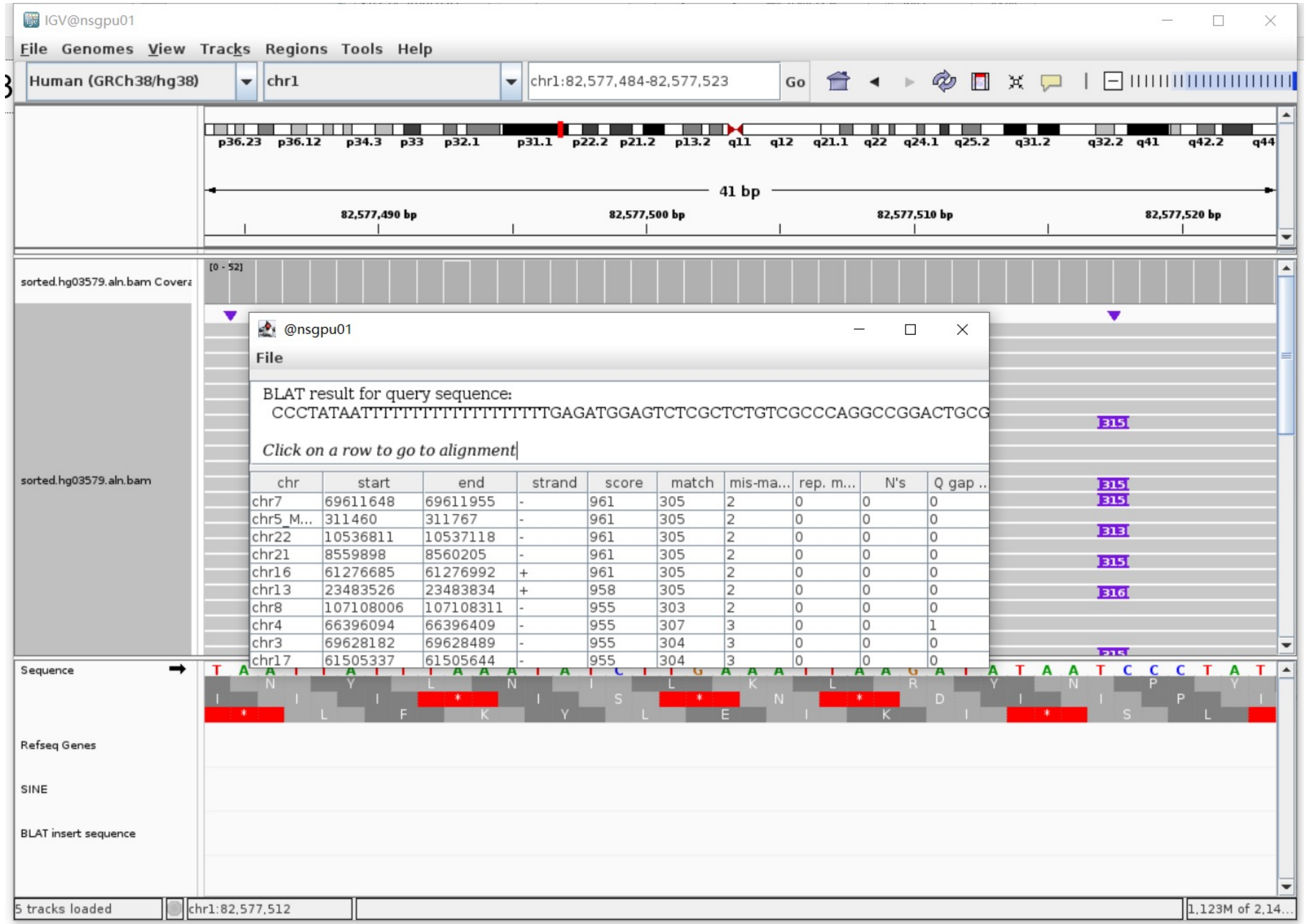

### ERX2355869 chr2:1467935

- The same genomic coordinate as in HG02630 (page 25)

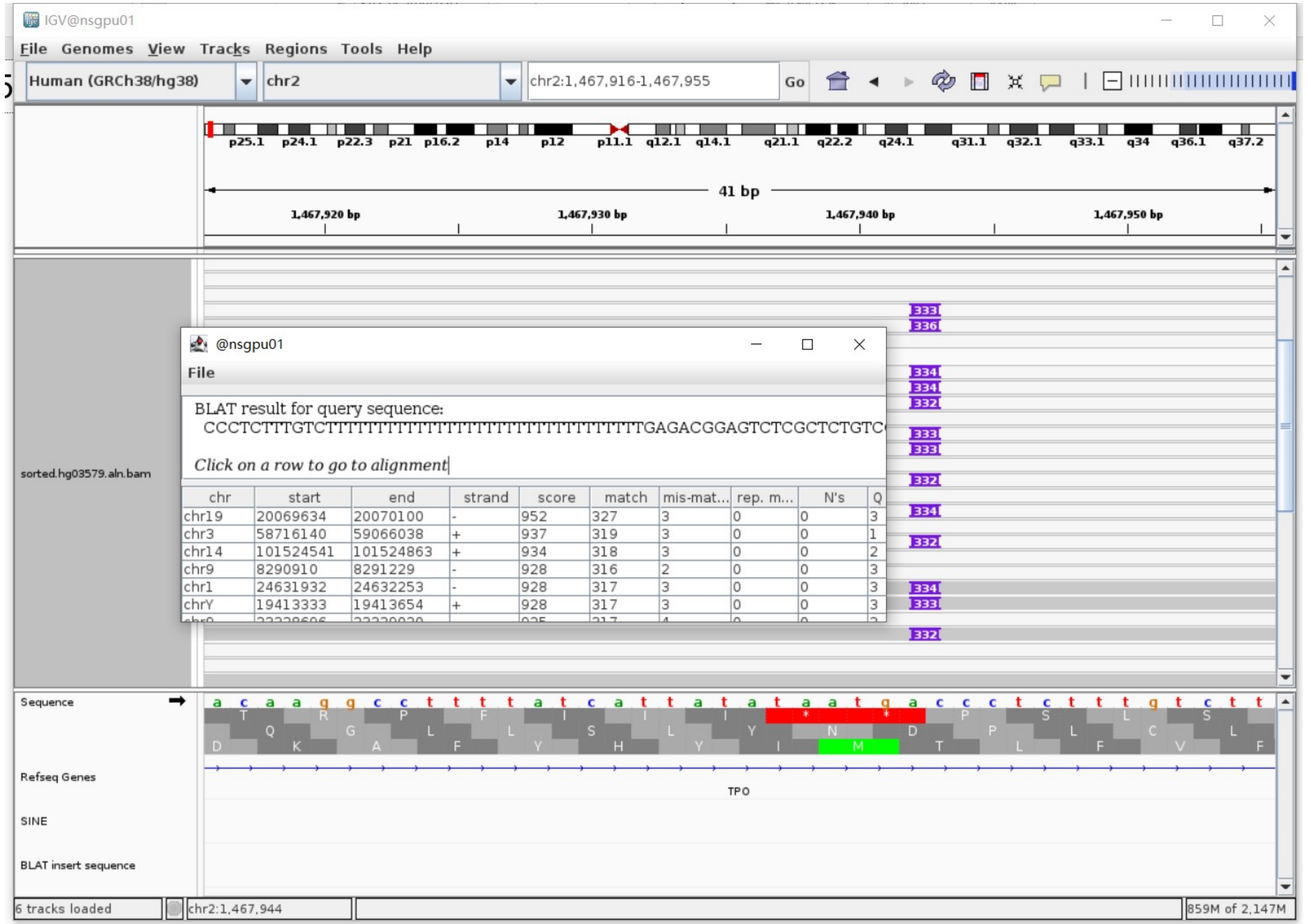

### ERX2355869 chr6:31282440

➢ The same genomic coordinate as in HG01106 (page 20)

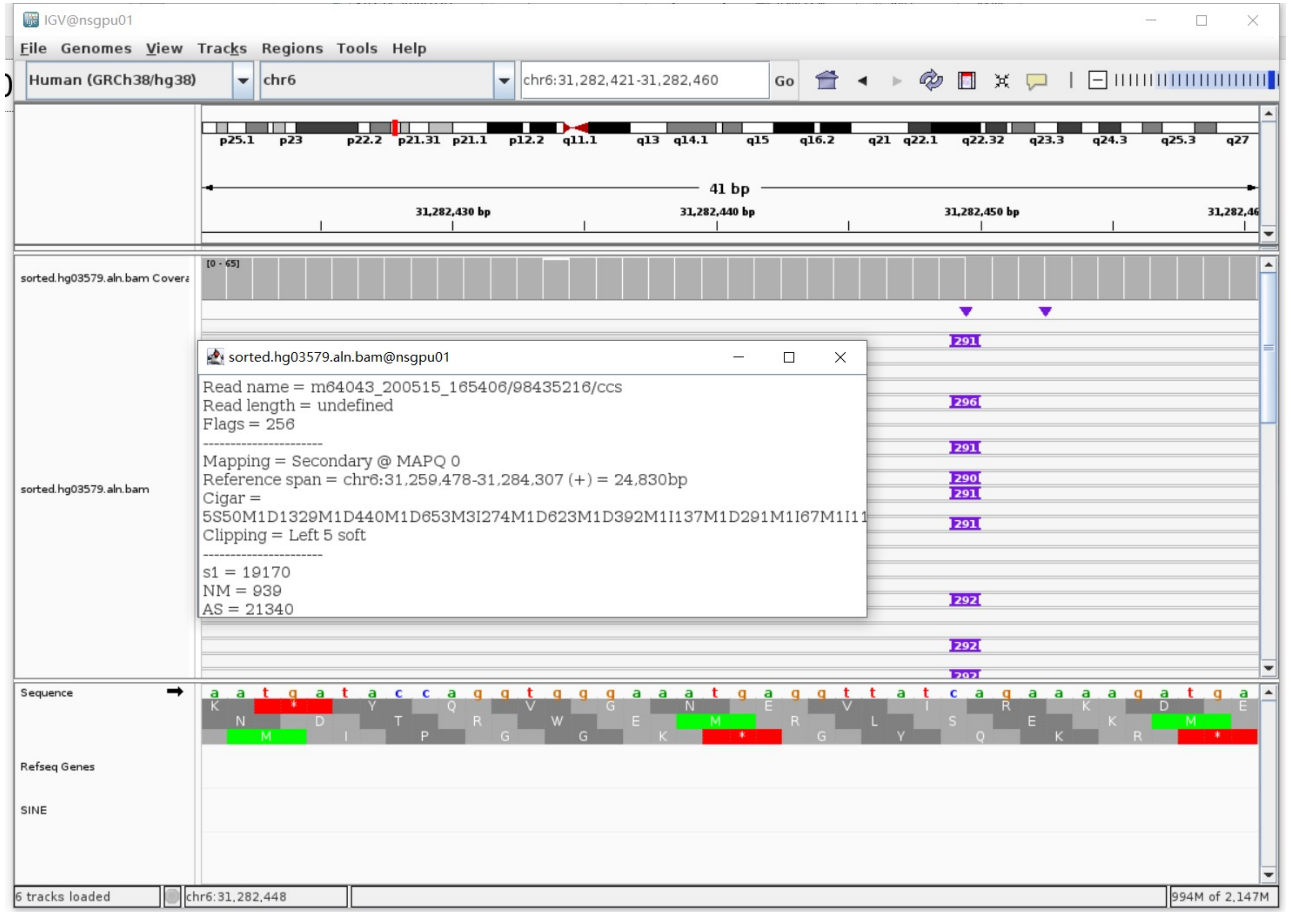

### ERX2355869 chr6:99022883

- ➢ Using the built-in BLAT tool in IGV to check where the inserted sequence aligned.

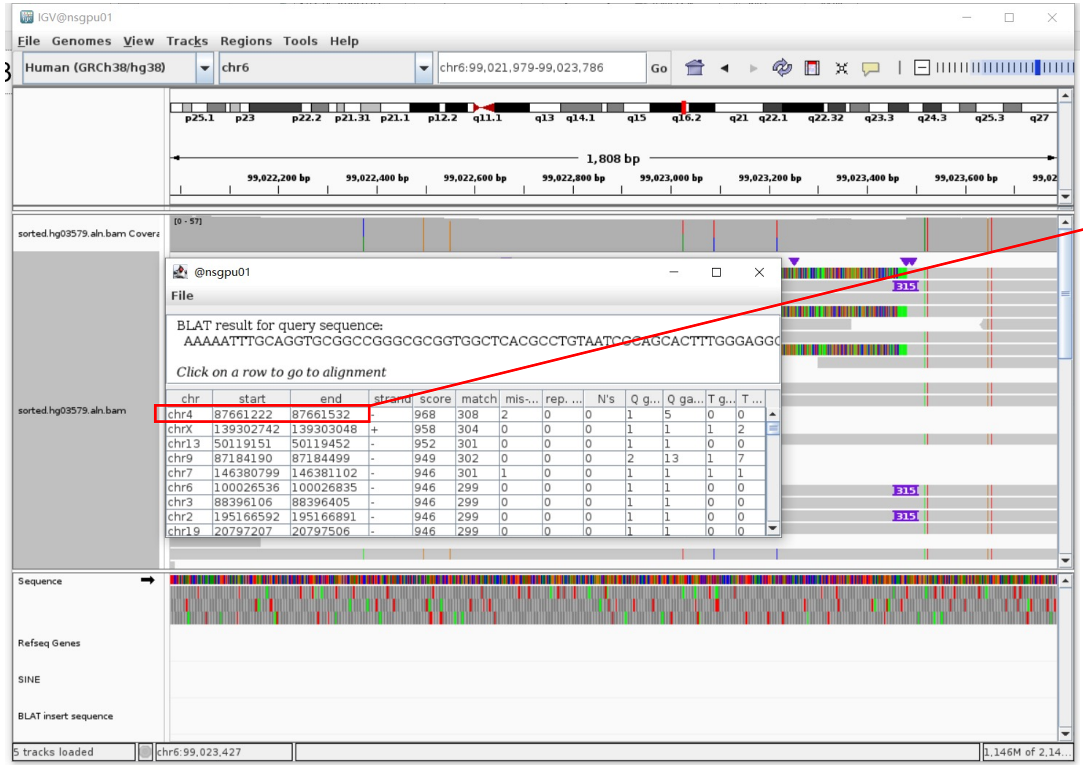


- ➢ The annotation at the aligned location indicated it as *AluY*.

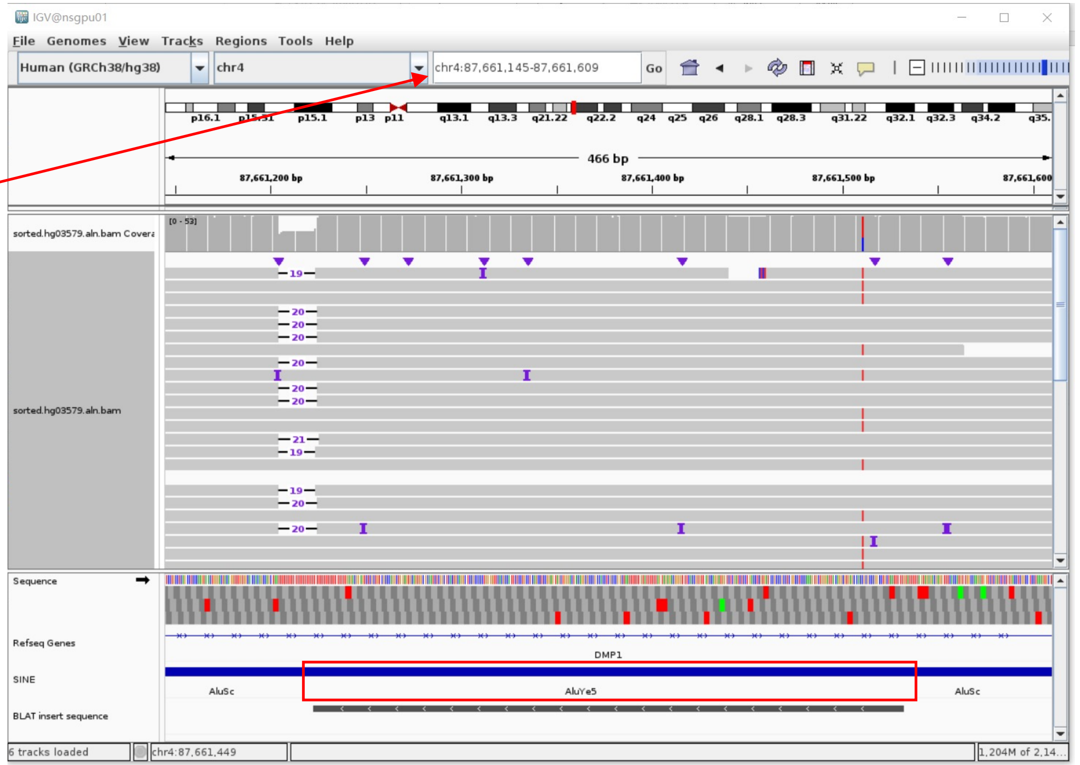


Insertion sequence (315bp)

- ➢ 16bp TSD

AAAAATTTGCAGGTGC

- ➢ Insertion:16-298 aligns to *AluY*c1:1-282, 97.86% identity

GGCCGGGCGCGGTGGCTCACGCCTGTAATCCCAGCACTTTGGGAGGCCGAGGCGGGCGGATCACGAGGTCAGGAGATCGAGACCATCCTGGCTAACACGGTGAAACCCCGTCTCTACTAAAAA
ATACAAAAAATTAGCCGGGCGAGGTGGCGGGCGCCTGTAGTCCCAGCTACTCGGGAGGCTGAGGCAGGAGAATGGCGTGAACCCAGGGGGCGGAGCCTGCAGTGAGCCGAGATTGCGCCACT
GCACTCCAGCCTGGGCGACAGCGAGACTCCGTCTCA………poly(A)

### ERX2355869 chr6:104042021

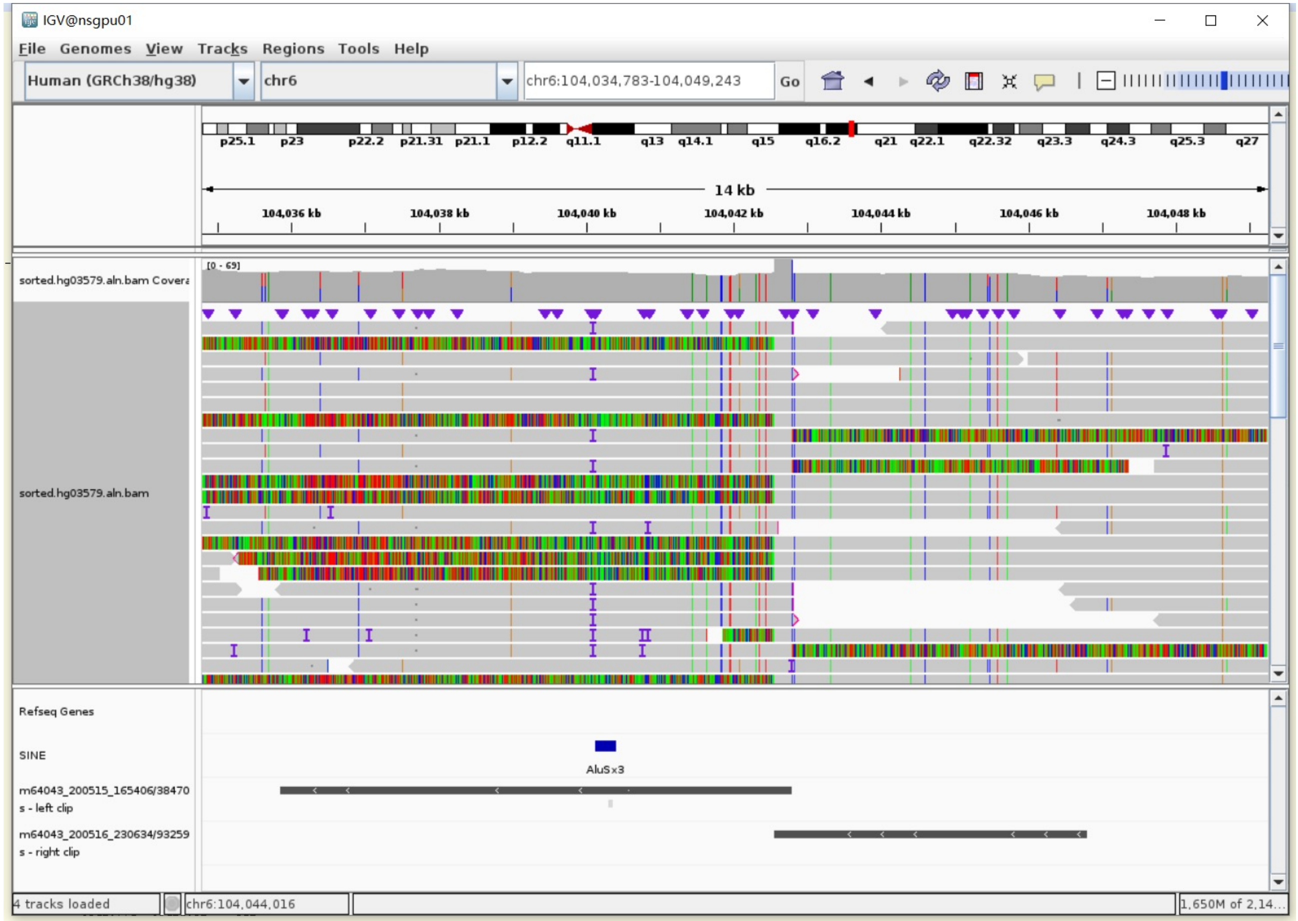


Right-clipped sequence

- 249bp TSD

AAGAAACACCATTCTGGACATTGGCCTTGAGAAAGAATTTATCATTAAGTGCTCAAAA
GCAATTGCAACAAAAACAAAATTTGACAAGTCAGACCTAATTCAACAAAAGAGCTTGT
GCACAACAATAGAAACAGAGTAGACAGATAACCTGTAGGATGGGAGAAAATATTAACA
AACCATGCATTCAACAAAGGCCTCATATCCAGAGTATATAAAGAACTTAATCCAACAA
GCAAAAAAAAACCTCTTT

- Clip:1-285 aligns to *Alu*Yb8:1-289, 95.04% identity

GGCCGGGCGCAGTGGCTCACGCCTGTAATCCCAGCACTTTGGGAGGCCGAGGCGGGCG
GGATCACGAGGTCAGGAGATCGAGACCATCCTGGCTAAAACGGTGAAACCCCGTCTCT
ACTAAAAATACAAAAAATTAGCCGGGCGTAGTGGCGGGCGCCTGTAGTCCCAGCTAC
TCGGGAGGCTGAGGCAGGAGAATGGCGTGAACCCGGGAGGCAGAGCTTGCAGTGAGCC
GAGATCAGGCCACTGCACTCTAGCCTGGGCGGACAGAGCGAGACTCCGTCTCA

### ERX2355869 chr8:1362348

- ➢ The same genomic coordinate as in NA18906 (page 100)

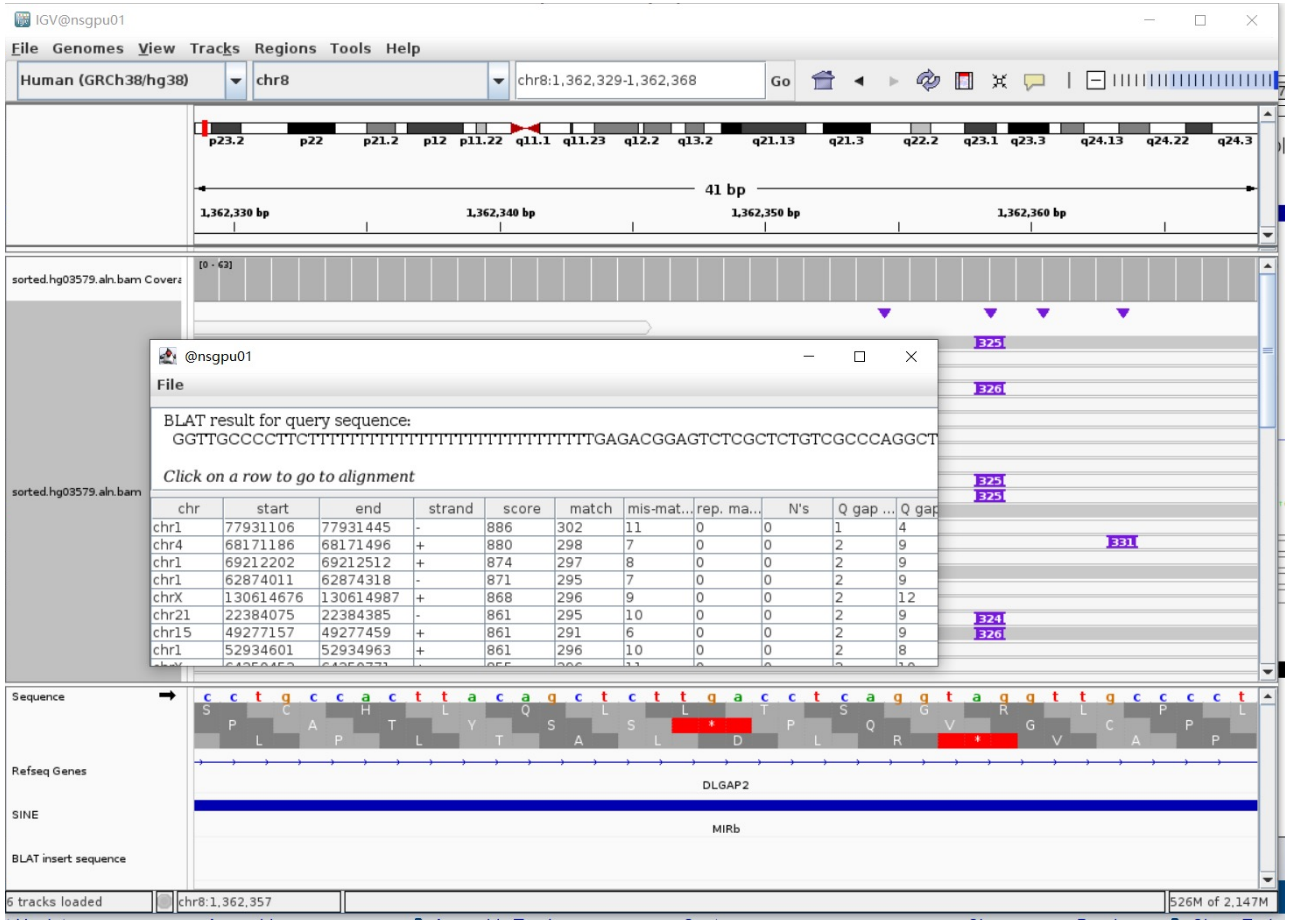

### ERX2355869 chr8:85019904

- Using the built-in BLAT tool in IGV to check where the inserted sequence aligned.

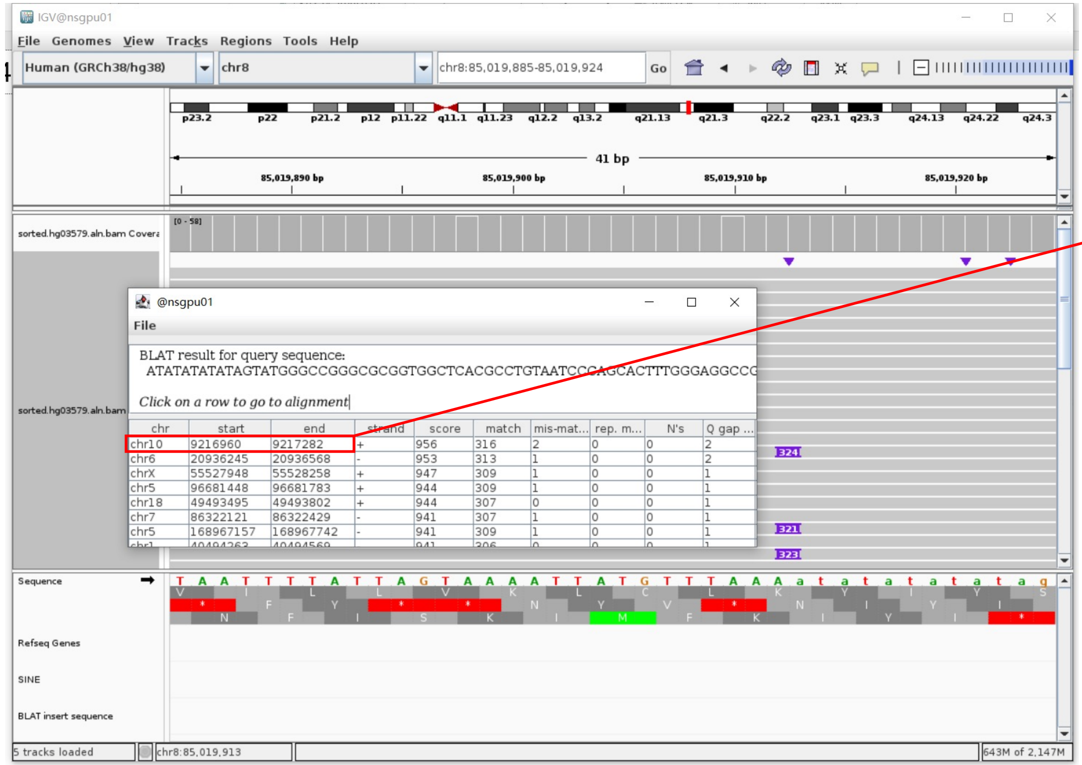


- The annotation at the aligned location indicated it as *AluY*.

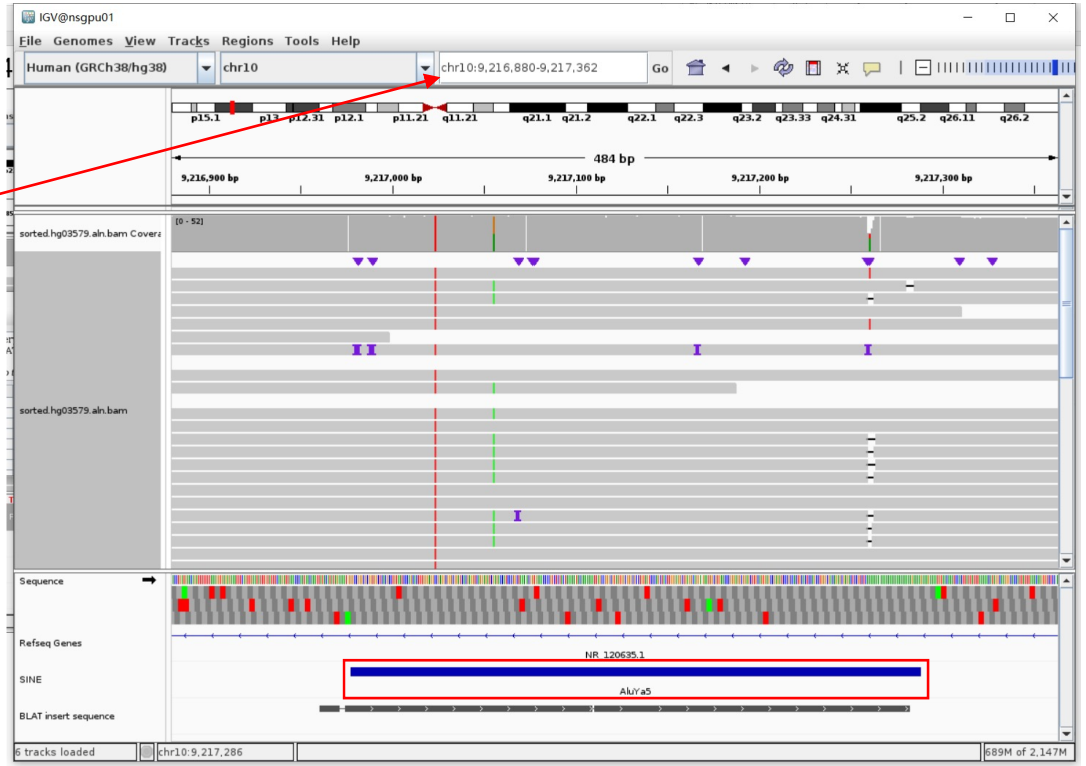


Insertion sequence (321bp)

- 16bp TSD

ATATATATATAGTATG

- Insertion:166-299 aligns to *Alu*Yb8:1-289, 96.10% identity

GGCCGGGCGCGGTGGCTCACGCCTGTAATCCCAGCACTTTGGGAGGCCGAGGCGGGCGGATCACGAGGTCAGGAGATCGAGACCATCCCGGCTAAAACGGTGAAACCCCGTCTCTACTAAAAA
TACAAAAAATTAGCCGGGCGTAGTGGCGGGCGCCTGTAGTCCCAGCTACTCGGGAGGCTGAGGCAGGAGAATGGCGTGAACCCGGGAGGCGGAGCTTGCAGTGAGCCGAGATCCCGCCACTG
CACTCCAGCCTGGGCGACAGAGCGAGACTCCGTCTCA.........poly(A)

### ERX2355869 chr1:246243395

- The same genomic coordinate as in NA18906 (page 93)

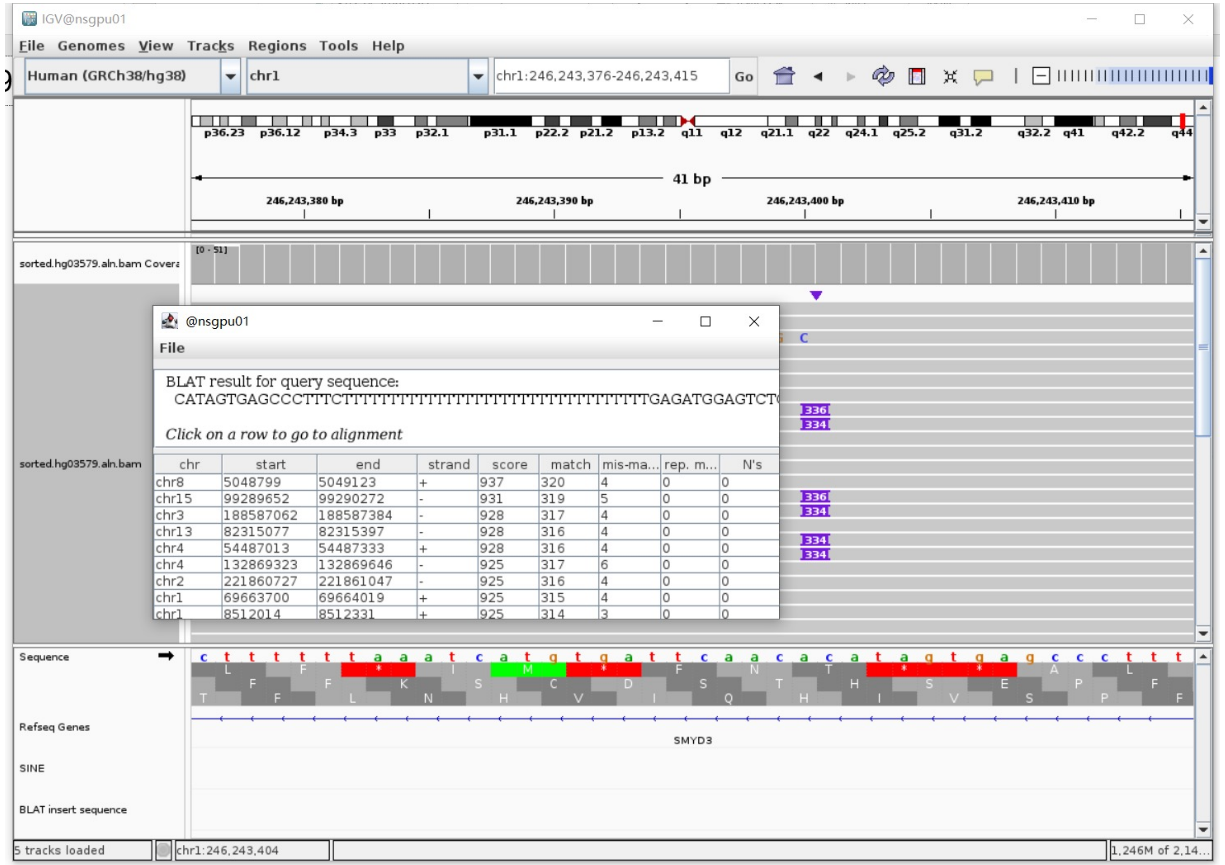

### ERX2355869 chr12:58064363

- ➢ The same genomic coordinate as in HG02630 (page 33)

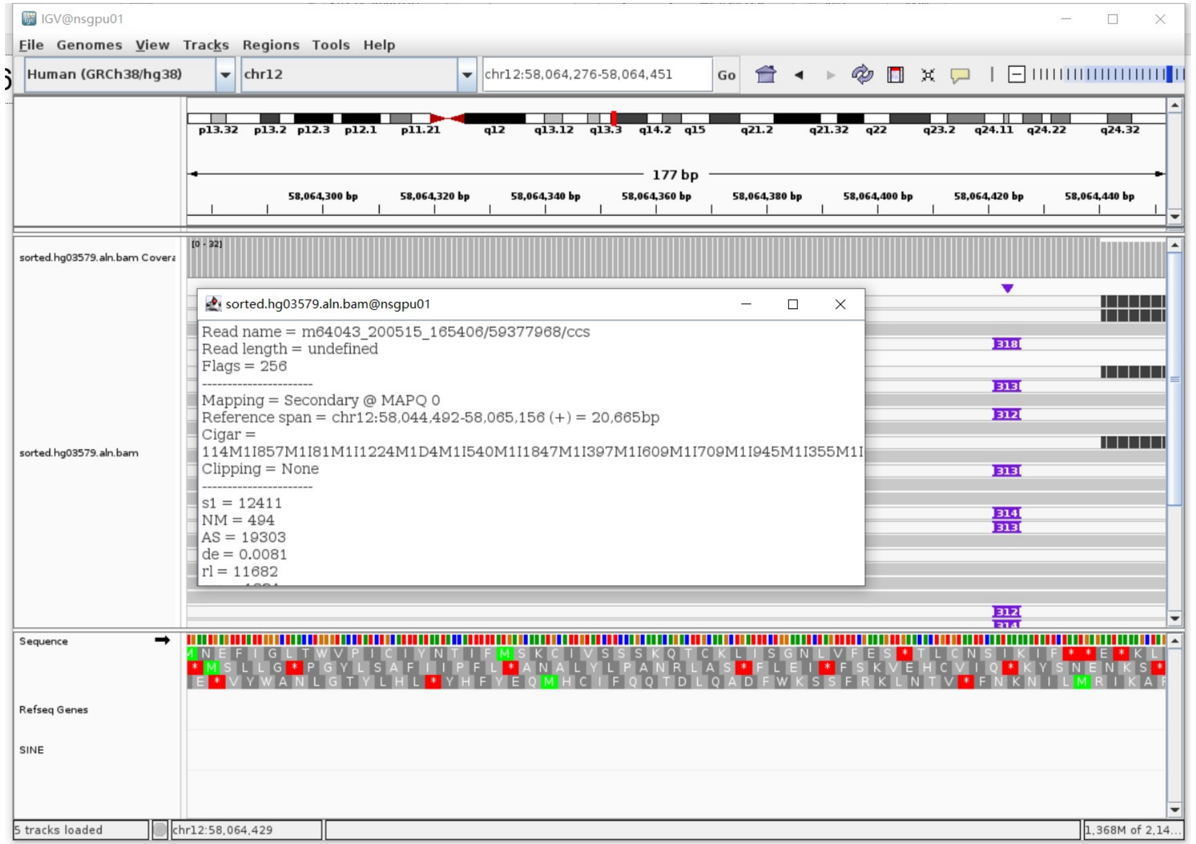

### ERX2355869 chr14:78516322

- Using the built-in BLAT tool in IGV to check where the inserted sequence aligned.
- The annotation at the aligned location indicated it as *Alu*Y.

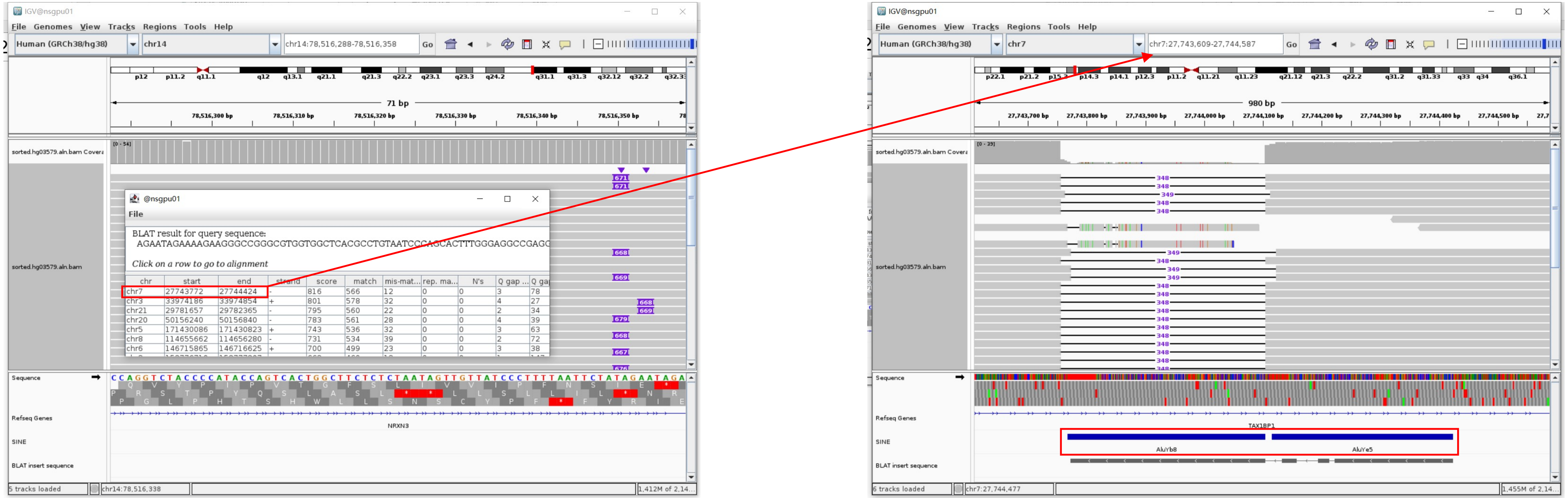


Insertion sequence (667bp)

- 16bp TSD

AGAATAGAAAAGAAGG

- Insertion:19-296 aligns to *Alu*Yc1:1-282, 95.73% identity

GGCCGGGCGTGGTGGCTCACGCCTGTAATCCCAGCACTTTGGGAGGCCGAGGCGGGCGGATCATGAGGTCAGGAGATCGAGACCATCCTGGCTAACACGGTGAAACCCCGTCTCTACTAAAAATACAAAAATTA
GCCGGGCGTGGTGGTGGGCGCCTGTAATCCCAGCTACTCGGGAGGCTGAGGCAGGAGAATGGCGTGAGCCCGGGAGGCGGAGCTTGCAGTGAGCCGGGATAGCGCCACTGCAGTCCAGCTTGGGCGAAAGAGTG
AGACTCCGTCTCA

- Insertion:339-628 aligns to *Alu*Yb8:1-289, 99.65% identity

GGCCGGGCGCGGTGGCTCACGCCTGTAATCCCAGCACTTTGGGAGGCCGAGGCGGGTGGATCATGAGGTCAGGAGATCGAGACCATCCTGGCTAACAAGGTGAAACCCCGTCTCTACTAAAAATACAAAAAATT
AGCCGGGCGCGGTGGCGGGCGCCTGTAGTCCCAGCTACTCGGGAGGCTGAGGCAGGAGAATGGCGTGAACCCGGGAAGCGGAGCTTGCAGTGAGCCGAGATTGCGCCACTGCAGTCCGCAGTCCGACCTGGGCG
ACAGAGCGAGACTCCGTCTCA.........poly(A)

### ERX2355869 chr15:30774821

➢ Mapping quality is 0, indicating that this region may have alignment issues.

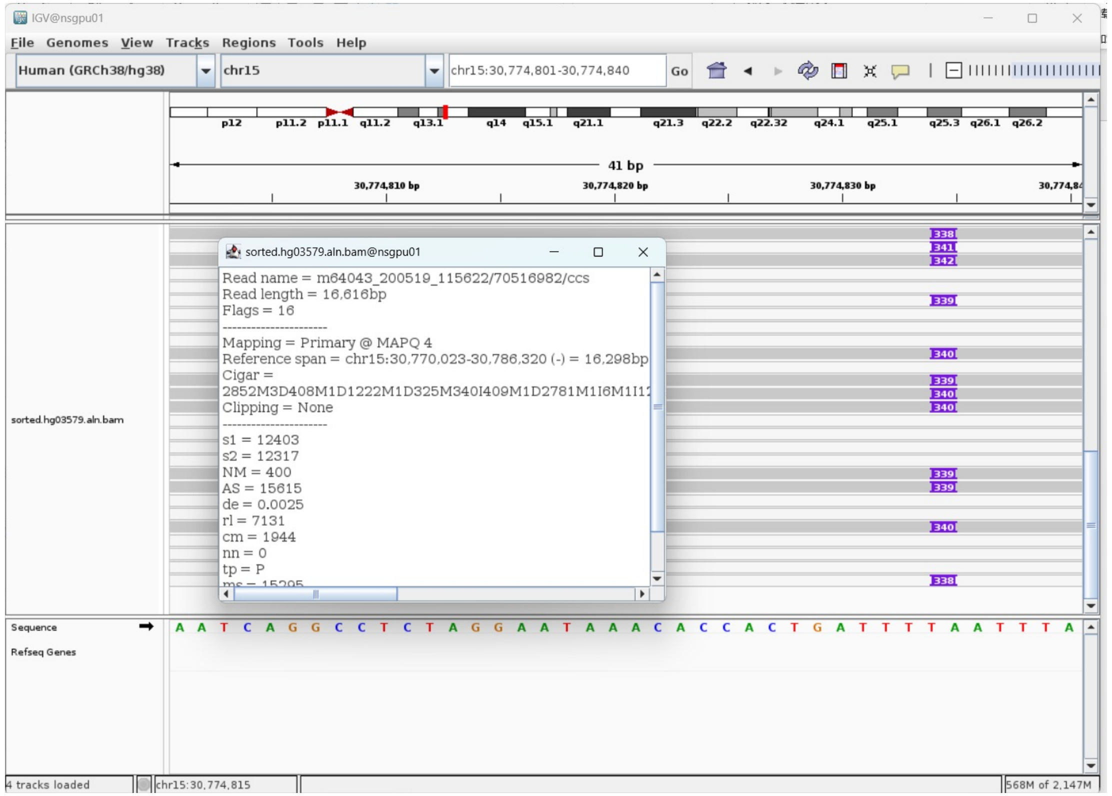


➢ Fetch one read (m64043_200518_053124/108265741/ccs, 21675 bp) and BLAT it to the hg38 genome.

The best alignment is to chr15_KI270905v1_alt (an alternative sequence of chr15) and chr15.

| ACTIONS | | | QUERY | SCORE | START | END | QSIZE | IDENTITY | CHROM | STRAND | START | END | SPAN |
|---|---|---|---|---|---|---|---|---|---|---|---|---|---|
| browser | new tab | details | YourSeq | 21259 | 1 | 21675 | 21675 | 99.8% | chr15_KN538374v1_fix | - | 2927730 | 2949069 | 21340 |
| browser | new tab | details | YourSeq | 21259 | 1 | 21675 | 21675 | 99.8% | chr15_KI270905v1_alt | - | 3040182 | 3061521 | 21340 |
| browser | new tab | details | YourSeq | 21233 | 1 | 21675 | 21675 | 99.7% | chr15 | - | 30754455 | 30775792 | 21338 |

- Read:1-958

| ACTIONS | | | QUERY | SCORE | START | END | QSIZE | IDENTITY | CHROM | STRAND | START | END | SPAN |
|---|---|---|---|---|---|---|---|---|---|---|---|---|---|
| browser | new tab | details | YourSeq | 955 | 1 | 957 | 957 | 99.9% | chr15_KN538374v1_fix | - | 2948112 | 2949068 | 957 |
| browser | new tab | details | YourSeq | 955 | 1 | 957 | 957 | 99.9% | chr15_KI270905v1_alt | - | 3060564 | 3061520 | 957 |
| browser | new tab | details | YourSeq | 953 | 1 | 957 | 957 | 99.8% | chr15 | - | 30774835 | 30775791 | 957 |

- Read:958-1248 aligns to *Alu*Yb8:1-289, 100% identity

GGGCCGGGCGCGGTGGCTCACGCCTGTAATCCCAGCACTTTGGGAGGCCGAGGCGGGTGGATCATGAGGTCAGG
AGATCGAGACCATCCTGGCTAACAAGGTGAAACCCCGTCTCTACTAAAAATACAAAAAATTAGCCGGGCGCGGT
GGCGGGCGCCTGTAGTCCCAGCTACTCGGGAGGCTGAGGCAGGAGAATGGCGTGAACCCGGGAAGCGGAGCTTG
CAGTGAGCCGAGATTGCGCCACTGCAGTCCGCAGTCCGGCCTGGGCGACAGAGCGAGACTCCGTCTCA

11bp TSD

AACATTAAATT

- Read:1249-21675

| ACTIONS | | | QUERY | SCORE | START | END | QSIZE | IDENTITY | CHROM | STRAND | START | END | SPAN |
|---|---|---|---|---|---|---|---|---|---|---|---|---|---|
| browser | new tab | details | YourSeq | 20315 | 38 | 20426 | 20426 | 99.9% | chr15_KN538374v1_fix | - | 2927730 | 2948122 | 20393 |
| browser | new tab | details | YourSeq | 20315 | 38 | 20426 | 20426 | 99.9% | chr15_KI270905v1_alt | - | 3040182 | 3060574 | 20393 |
| browser | new tab | details | YourSeq | 20291 | 38 | 20426 | 20426 | 99.8% | chr15 | - | 30754455 | 30774845 | 20391 |

➢ It's an *Alu*Yb8 insertion.

### ERX2355869 chr17:37883970

- The same genomic coordinate as in HG02630 (page 42)

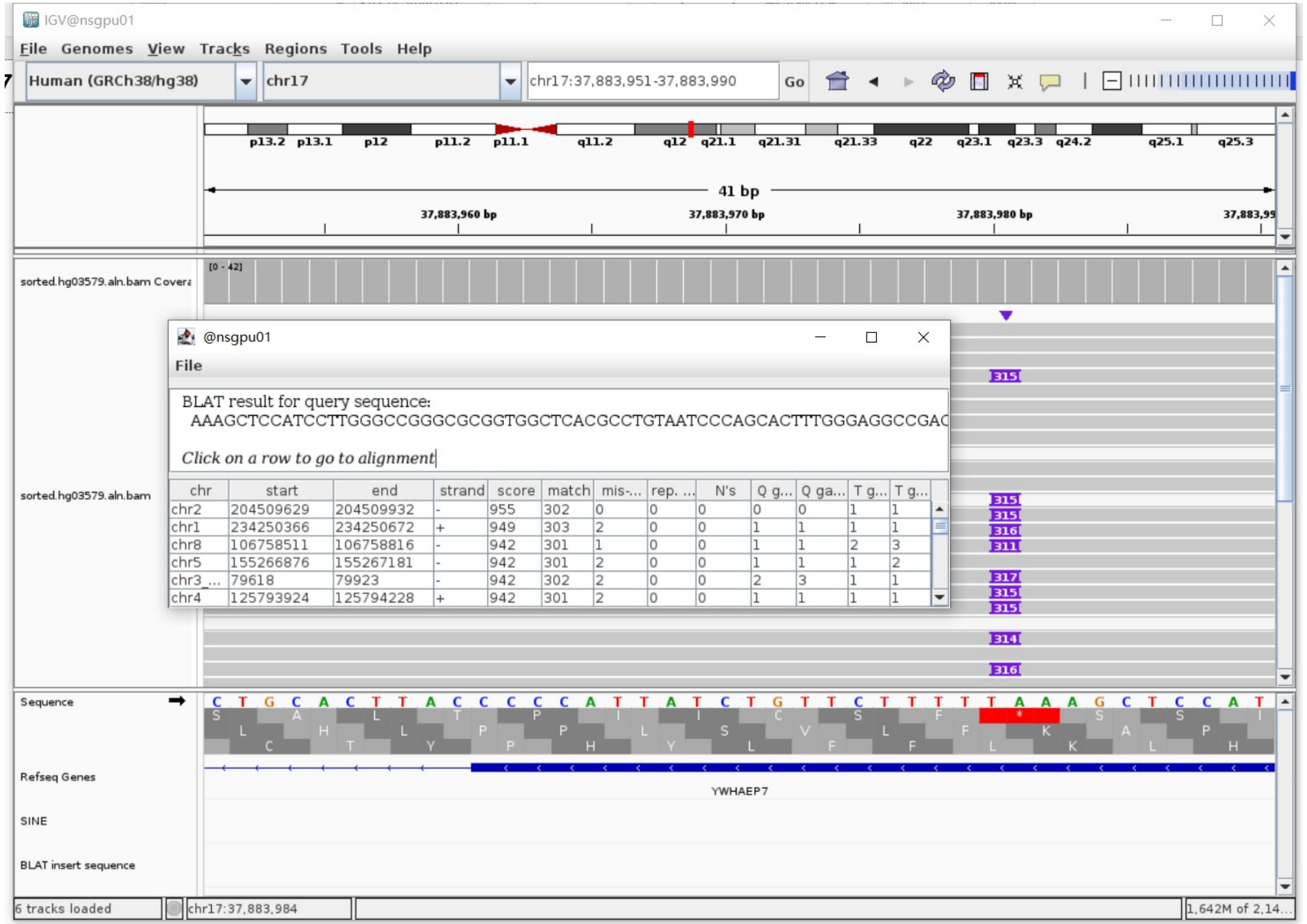

### ERX2355869 chr17:54885771

- ➢ Using the built-in BLAT tool in IGV to check where the inserted sequence aligned.
- ➢ The annotation at the aligned location indicated it as *AluY*.

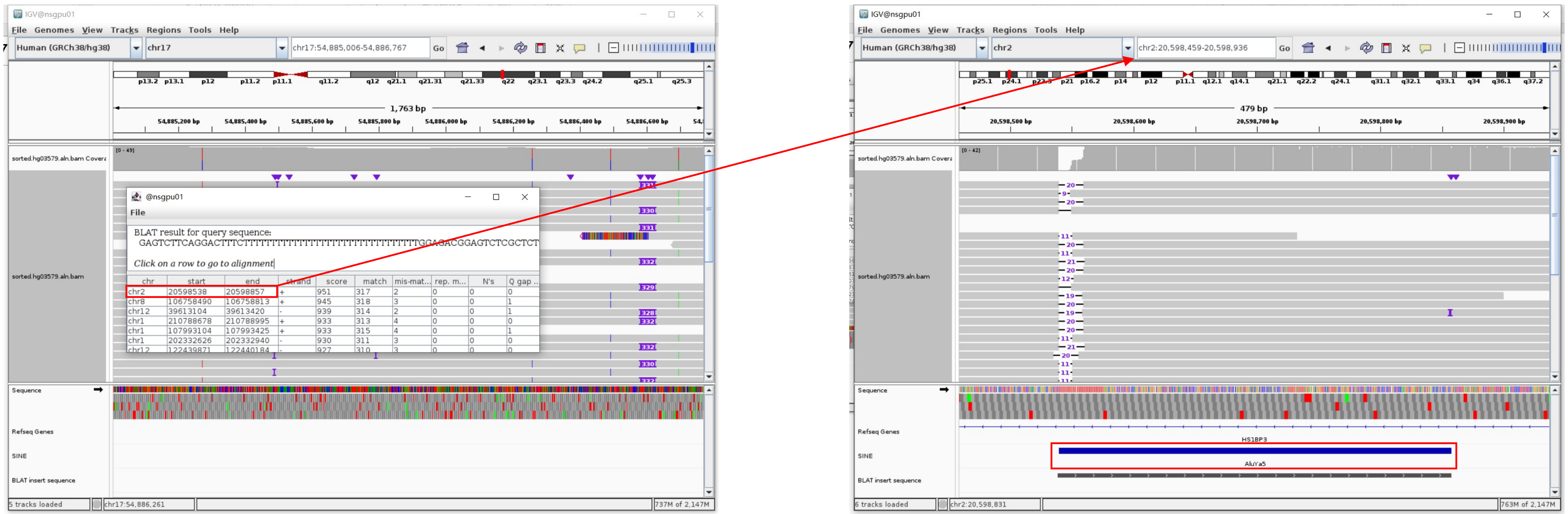


Insertion sequence (331bp)

- ➢ 18bp TSD

GAGTCTTCAGGACTTTCT

- ➢ Insertion:50-331 aligns to *AluYa5*:281-1, 99.29% identity

GAGACGGAGTCTCGCTCTGTCGCCCAGGCTGGAGTGCAGTGGCGGGATCTCGGCTCACTGCAAGCTCCGCCTCCCGGGTTCACGCCATTCTCCTGCCTCAGCCTCCCGAGTAGCTGGGACTAC
AGGCGCCCGCCACTACGCCCGGCTAATTTTTTGTATTTTTAGTAGAGACGGGGTTTCACTGTTTTAGCCGGGATGGTCTCGATCTCCTGACCTCGTGATCCGCCCGCCTCGGCCTCCCAAAGT
GCTGGGATTACAGGCGTGAGCCACCGCGCCCGGCC
