## Extended Files for "Image-based DNA Sequencing Encoding for Detecting Low-Mosaicism Somatic Mobile Element Insertions": Extended Files 3.pdf

### Manual check of SVA label that were not identified by Pangenome xTea_long PacBio calls or the Polymorphic SVA dataset

(Additional explanation for Supplementary Table 8)

| ID | Chr | Coordinate | Manual Inspection |
|---|---|---|---|
| HG00438 | chr4 | 9703198 | TRUE |
| HG00621 | chr5 | 721226 | TRUE |
| | chr7 | 128575317 | TRUE |
| HG02630 | chr4 | 87384744 | TRUE |
| HG03540 | chr7 | 21349667 | TRUE |
| NA20129 | chr19 | 50004904 | TRUE |
| | chr7 | 128575317 | TRUE |
| ERX2355888 | chr13 | 24314860 | TRUE |

### HG00438 chr4:9703198

- Using IGV to check the insert in PacBio

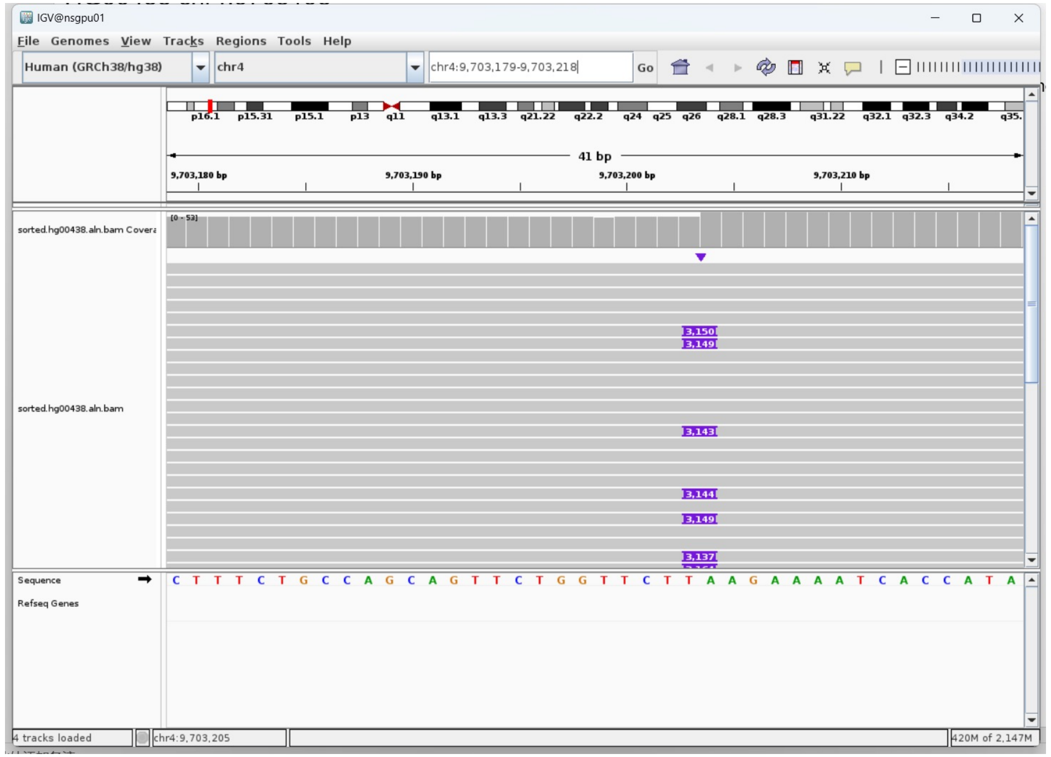


- 19 bp TSD

AAGAAAATCACCATAAATC

- Blat insert sequence to hg38 genome. Annotations from RepearMasker indicated it as SVA_D. Annotations from the polymorphic SVA dataset defined by Chu et al.(2023) indicated it as SVA_E.

| ACTIONS | | | QUERY | SCORE | START | END | QSIZE | IDENTITY | CHROM | STRAND | START | END | SPAN |
|---|---|---|---|---|---|---|---|---|---|---|---|---|---|
| browser | new tab | details | YourSeq | 2544 | 21 | 3063 | 3150 | 96.6% | chr1 | + | 7952202 | 8503136 | 550935 |
| browser | new tab | details | YourSeq | 2416 | 21 | 3135 | 3150 | 96.2% | chr3 | - | 106072784 | 106075681 | 2898 |
| browser | new tab | details | YourSeq | 2349 | 21 | 3126 | 3150 | 96.2% | chr8 | + | 116678115 | 116680856 | 2742 |

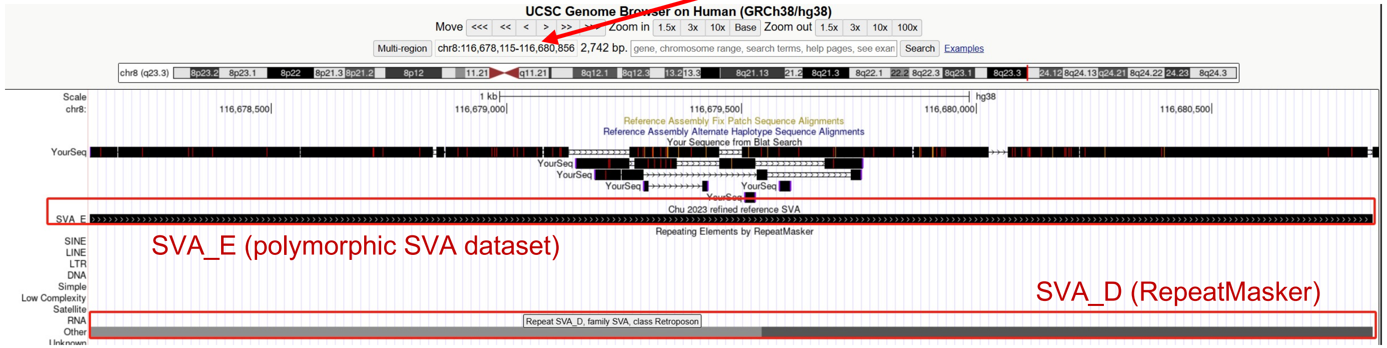


- Blast insertion sequence to the polymorphic SVA dataset defined by Chu et al. Insertion: 360-3150 aligns to **SVA_E:**1-2714 (at chr4:9703208 of HG02818, 97% identity).

```
>HG02818_chr4_9703208
Length=2726

 Score = 4578 bits (5076),  Expect = 0.0
 Identities = 2708/2794 (97%), Gaps = 83/2794 (3%)
 Strand=Plus/Plus

Query  360   CGTCTGGGATATGAGGAGCCTCTCTGCCTGGCTGCACAGTCTGGAAAGTGAGGAGCGTCT  419
             ||||||||||||||||||||||||||||||||||||||||||||||||||||||||||||
Sbjct  1     CGTCTGGGATATGAGGAGCCTCTCTGCCTGGCTGCACAGTCTGGAAAGTGAGGAGCGTCT  60

Query  420   CTGCCCGGCCGCCATCCCATCTAGGAAGCGAGGAGCGCCTCTTCCCCCGCCGCCATCCCA  479
             |||||||||||||||||||||||||||||||||||||||||||||| |||||||||||||
Sbjct  61    CTGCCCGGCCGCCATCCCATCTAGGAAGCGAGGAGCGCCTCTTCCCC-GCCGCCATCCCA  119
```

.........

```
Query  3058  aaaGAAAGAAAATAACATTCGTAACTTTGAGG-AACAGAATAAATAGGAGATTTATTAAA  3116
             |||||||||||||||||||||||||||||||| |||||||||||||||||||||||||||
Sbjct  2621  AAAGAAAGAAAATAACATTCGTAACTTTGAGGAAACAGAATAAATAGGAGATTTATTAAA  2680

Query  3117  Taaaaaaaaaaaaaaaaaaaaaaaaaaaaaaa  3150
             ||||||||||||||||||||||||||||||||
Sbjct  2681  TAAAAAAAAAAAAAAAAAAAAAAAAAAAAAAA  2714
```

### HG00621 chr5:721226

- Using IGV to check the insert in PacBio

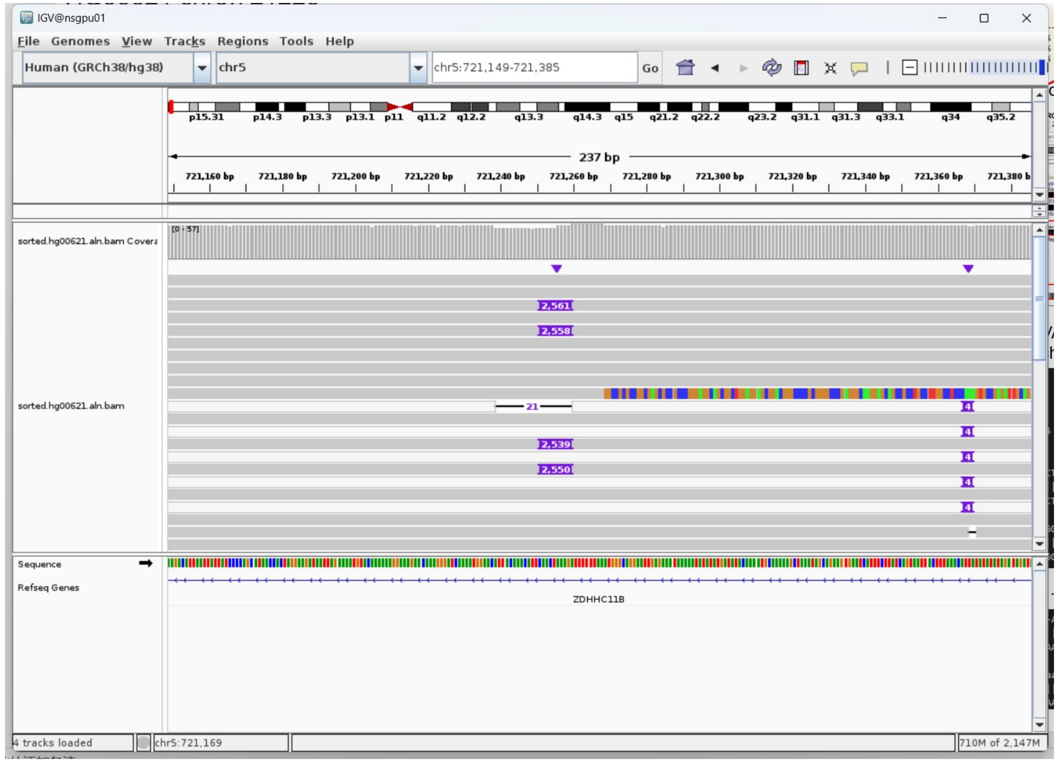

Insertion sequence (2558bp):

- Insertion:1-13，13bp TSD
  AAAGATTTTTTAA
- Insertion:15-2546 aligns to chr1:46240429-46242954. Annotations from RepearMasker and the polymorphic SVA dataset defined by Chu et al. indicated it as SVA_F.

| ACTIONS | QUERY | SCORE | START | END | QSIZE | IDENTITY | CHROM | STRAND | START | END | SPAN |
|---|---|---|---|---|---|---|---|---|---|---|---|
| browser new tab details | YourSeq | 2254 | 15 | 2546 | 2561 | 97.2% | chr1 | + | 46240429 | 46242954 | 2526 |
| browser new tab details | YourSeq | 2246 | 47 | 2561 | 2561 | 97.7% | chr10 | - | 92673703 | 92676023 | 2321 |
| browser new tab details | YourSeq | 2227 | 18 | 2548 | 2561 | [illegible] | chr1 | + | 113068744 | 113071160 | 2417 |

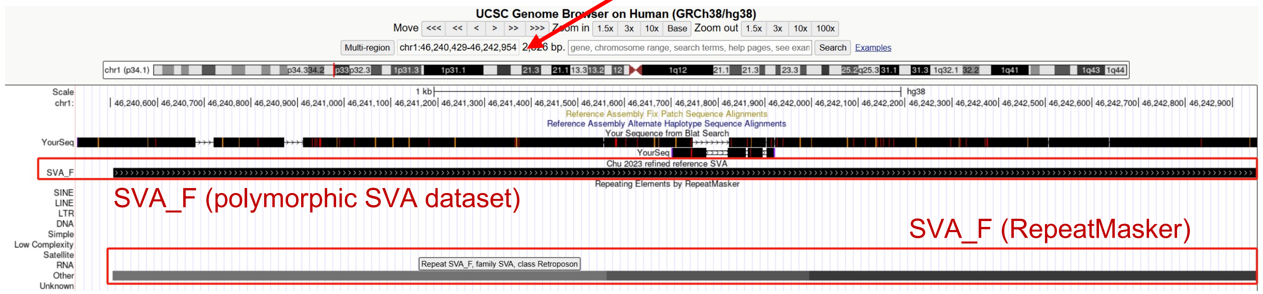


- Blast insertion sequence to the polymorphic SVA dataset defined by Chu et al. Insertion:47-2558 aligns to **SVA_F:**1-2476 (at chr5:721262 of HG005, 98% identity).

```
>HG005_chr5_721262
Length=2478

 Score = 4205 bits (4663),  Expect = 0.0
 Identities = 2470/2514 (98%), Gaps = 40/2514 (2%)
 Strand=Plus/Plus

Query  47    GGCTGGAGGAGCGGGACGGGCCCCGCGGGGCCCGAGGGCAAGGAGCAGCCGCCTGCCTTG  106
             ||||||||||||| ||||||||||||| |||||||||||||||||||||||||||||||
Sbjct  1     GGCTGGAGGAGCGG-ACGGGCCCCGCGGG-CCCGAGGGCAAGGAGCAGCCGCCTGCCTTG  58
```

.........

```
Query  2445  TGACCTTCCCTCCACTATTGTCCCATGACCCTGCCAAATCCCCCTCTGTGAGAAACACCC  2504
             ||||||||||||||||||||||||||||||||||||||||||||||||||||||||||||
Sbjct  2363  TGACCTTCCCTCCACTATTGTCCCATGACCCTGCCAAATCCCCCTCTGTGAGAAACACCC  2422

Query  2505  AAGAATTATCAATaaaaaaataaattaaaaaaaaaaaaaaaaaaaaaaaaaaa  2558
             ||||||||||||||||||||||||||||||||||||||||||||||||||||||
Sbjct  2423  AAGAATTATCAATAAAAAAATAAATTAAAAAAAAAAAAAAAAAAAAAAAAAAA  2476
```

### HG00621 chr7:128575317

- The same genomic coordinate as in NA20129 (page 8)

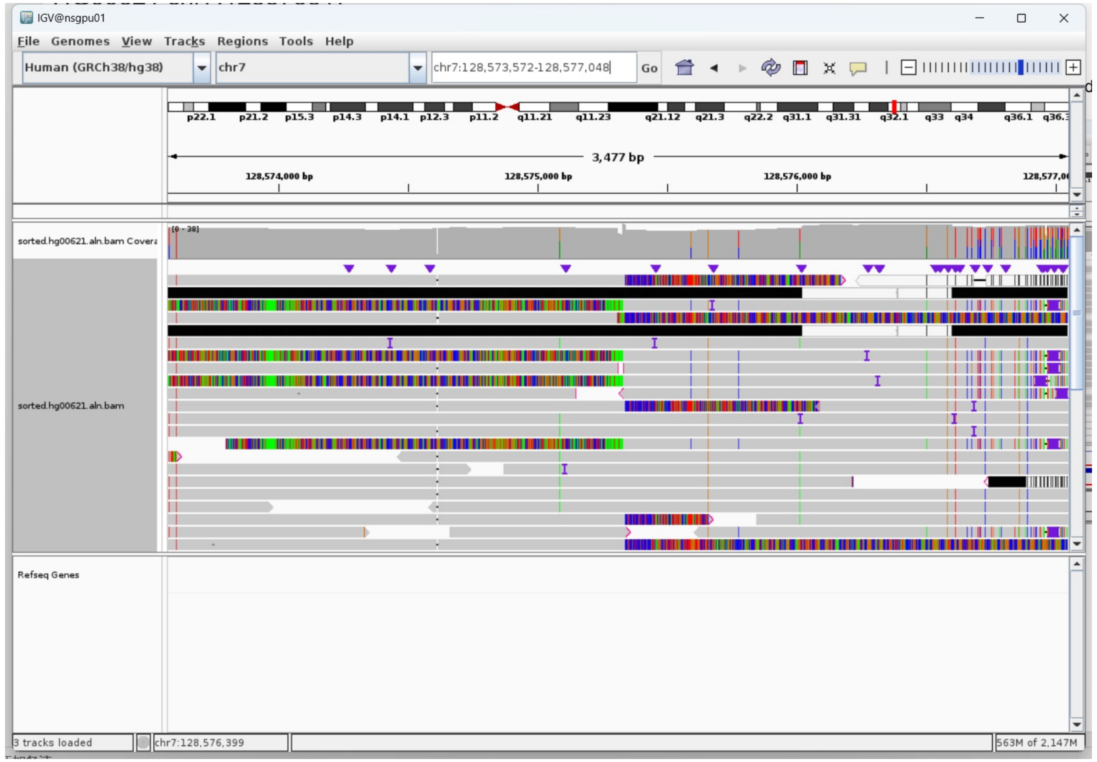


- 8 bp TSD

AAAATTGA

### HG02630 chr4:87384744

➢ Using IGV to check the insert in PacBio

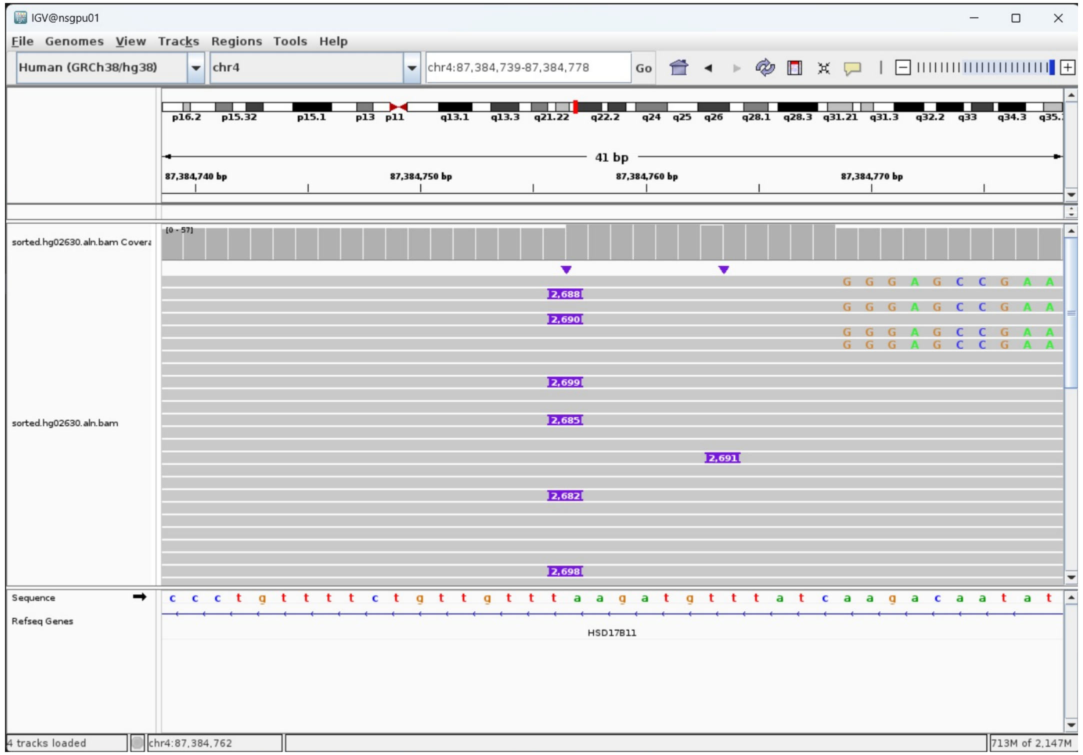


➢ 12 bp TSD

AAGATGTTTATC

➢ Blat insert sequence to hg38 genome. Annotations from RepearMasker and the polymorphic SVA dataset defined by Chu et al.(2023) indicated it as SVA_F.

| ACTIONS | | | QUERY | SCORE | START | END | QSIZE | IDENTITY | CHROM | STRAND | START | END | SPAN |
|---|---|---|---|---|---|---|---|---|---|---|---|---|---|
| browser | new tab | details | YourSeq | 2458 | 14 | 2687 | 2688 | 98.2% | chr7 | + | 1037302 | 1040204 | 2903 |
| browser | new tab | details | YourSeq | 2448 | 14 | 2681 | 2688 | 98.6% | chr6 | - | 122847777 | 122850260 | 2484 |
| browser | new tab | details | YourSeq | 2406 | 14 | 2688 | 2688 | 98.2% | chr2 | - | 127118884 | 127789661 | 670778 |

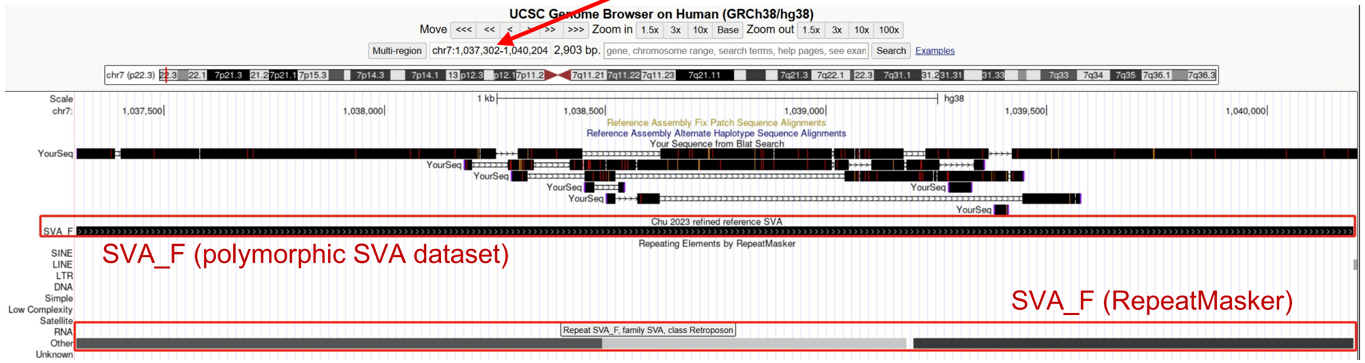


➢ Blast insert sequence to polymorphic SVA defined by Chu et al. Insertion:14-2684 aligns to **SVA_F:**37-2701 (at chr21:39908212 of HG002, 99% identity).

```
>HG002_ccs_chr21_39908212
Length=2712

 Score = 4690 bits (5201),  Expect = 0.0
 Identities = 2650/2673 (99%), Gaps = 10/2673 (0%)
 Strand=Plus/Plus

Query  14    GGAGCCGAAGCTGGACTGTACTGCTGCCATCTCGGCTCACTGCAACCTCCCTGCCTGATT  73
             ||||||||||||||||||||||||||||||||||||||||||||||||||||||||||||
Sbjct  37    GGAGCCGAAGCTGGACTGTACTGCTGCCATCTCGGCTCACTGCAACCTCCCTGCCTGATT  96

Query  74    CTCCTGCCTCAGCCTGCCCAGTGCCTGCAATGGCGCCGCCACGCCTGACTGGTTTTGGTG  133
             ||||||||||||||||||||||||||||||||||||||||||||||||||||||||||||
Sbjct  97    CTCCTGCCTCAGCCTGCCCAGTGCCTGCAATGGCGCCGCCACGCCTGACTGGTTTTGGTG  156
.........
Query  2592  CACTATTGTCCCATGACCCTGCCAAATCCCCCTCTGTGAGAAACACCCAAGAATTATCAA  2651
             ||||||||||||||||||||||||||||||||||||||||||||||||||||||||||||
Sbjct  2609  CACTATTGTCCCATGACCCTGCCAAATCCCCCTCTGTGAGAAACACCCAAGAATTATCAA  2668

Query  2652  Taaaaaaataaattaaaaaaaaaaaaaaaaaa  2684
             ||||||||||||| |||||||||||||| ||
Sbjct  2669  TAAAAAAATAAATTTAAAAAAAAAAAAAAAGAA  2701
```

### HG03540 chr7:21349667

- Using IGV to check the insert in PacBio

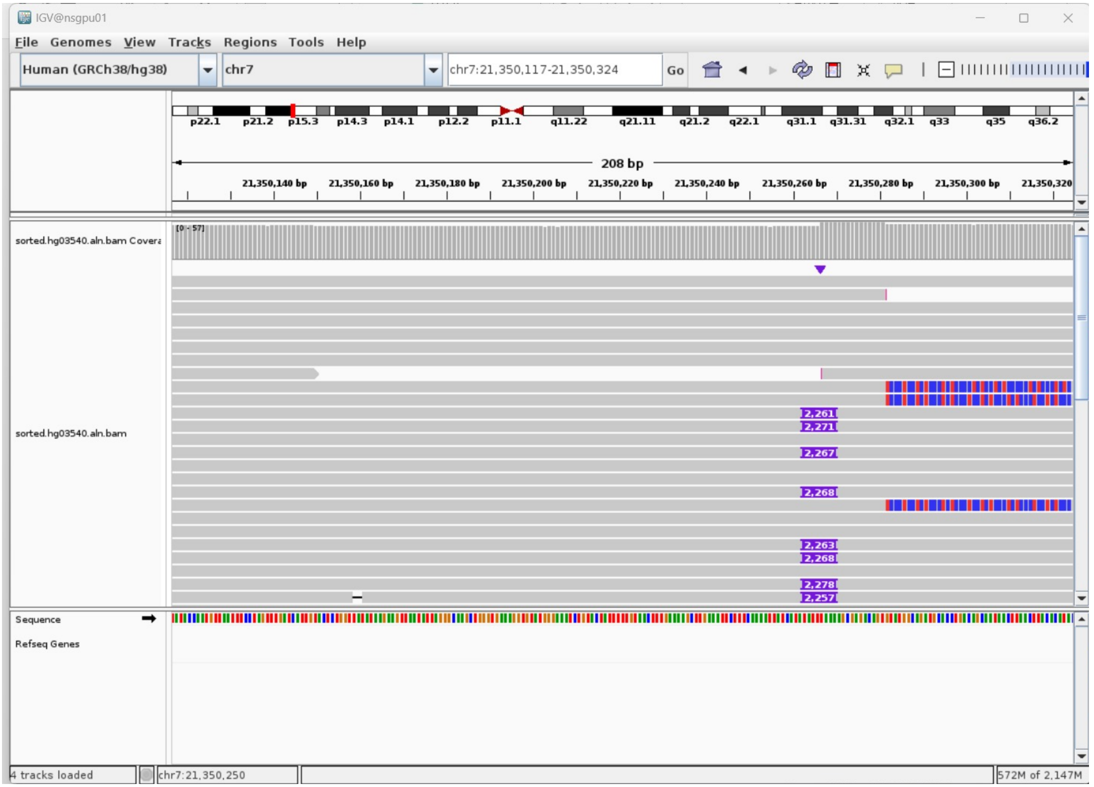


- 15 bp TSD

TAAAAGCGAGACAGT

- Blat insert sequence to hg38 genome. Annotations from RepearMasker and the polymorphic SVA dataset defined by Chu et al.(2023) indicated it as SVA_D.

| ACTIONS | | | QUERY | SCORE | START | END | QSIZE | IDENTITY | CHROM | STRAND | START | END | SPAN |
|---|---|---|---|---|---|---|---|---|---|---|---|---|---|
| browser | new tab | details | YourSeq | 2196 | 16 | 2241 | 2261 | 99.5% | chr10 | + | 80533316 | 80535592 | 2277 |
| browser | new tab | details | YourSeq | 2067 | 40 | 2260 | 2261 | 98.3% | chr20 | + | 36999420 | 37001792 | 2373 |
| browser | new tab | details | YourSeq | 1984 | 33 | 2248 | 2261 | 97.4% | chr1 | + | 143702020 | 143704154 | 2135 |

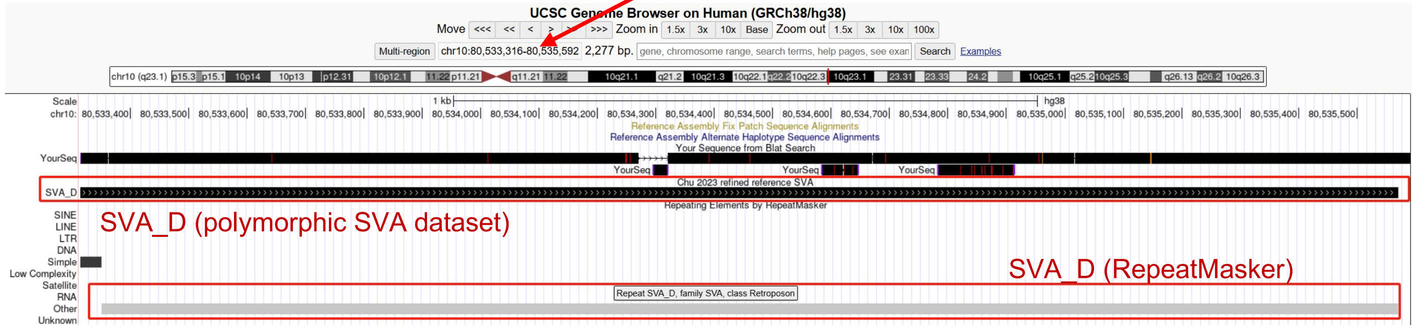


- Blast insert sequence to polymorphic SVA defined by Chu et al. Insertion: 34-2241 aligns to **SVA_D:**1-2258 (at chr16:75469417 of HG002, 97% identity).

```
>HG002_ccs_chr16_75469417
Length=2258

 Score = 3832 bits (4249),  Expect = 0.0
 Identities = 2201/2260 (97%), Gaps = 54/2260 (2%)
 Strand=Plus/Plus

Query  34    ctcctctccctctccctcctctccctctcc-tcctctccctctccctctccctctcccca  92
             |||||||||||||||||||||||||||||| ||||||||||||||||||||||||||||| |
Sbjct  1     CTCCTCTCCCTCTCCCTCCTCTCCCTCTCCCTCCTCTCCCTCTCCCTCTCCCTCTCCCTA  60

Query  93    cggtctccctctccctctctttccacggtctcccccTGATGCCGAGCCAAAGCTGGACTG  152
             ||||||||||||||||||||||||||||||||||||||||||||||||||||||||||||
Sbjct  61    CGGTCTCCCTCTCCCTCTCTTTCCACGGTCTCCCCCTGATGCCGAGCCAAAGCTGGACTG  120
.........
Query  2142  CTCCACTATTGTCCTGTGACCCTGCCAAATCCCCCTCTGCGAGAAACACCCAAGAATGAT  2201
             ||||||||||||||||||||||||||||||||||||||||||||||||||||||||||||
Sbjct  2159  CTCCACTATTGTCCTGTGACCCTGCCAAATCCCCCTCTGCGAGAAACACCCAAGAATGAT  2218

Query  2202  CAATaaaaaagaaaaaaaTGGATTACCCTGTGCACAGTa  2241
             |||||||||||||||||||||||||||||||||||||||
Sbjct  2219  CAATAAAAAAGAAAAAAAATGGATTACCCTGTGCACAGTA  2258
```

### NA20129 chr19:50004904

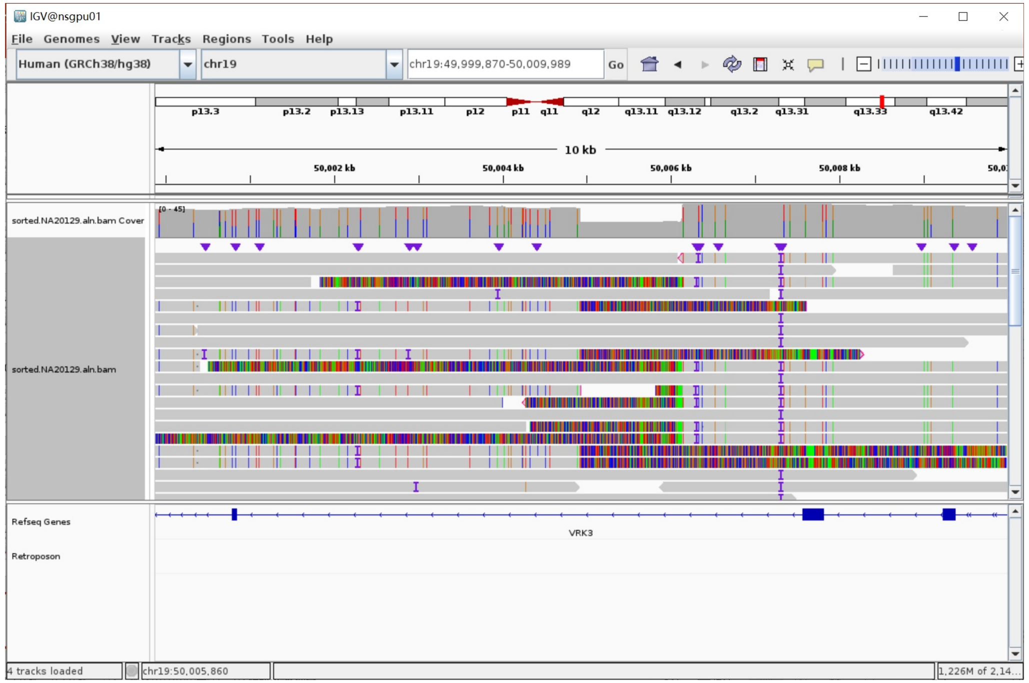


- Fetch one read (m64043_191230_073311/47121434/ccs, 21791 bp) and BLAT it to the hg38 genome.
- Read:1-6791 aligns to chr19:49998130-50004928

| ACTIONS | | | QUERY | SCORE | START | END | QSIZE | IDENTITY | CHROM | STRAND | START | END | SPAN |
|---|---|---|---|---|---|---|---|---|---|---|---|---|---|
| browser | new tab | details | YourSeq | 6638 | 1 | 6777 | 6791 | 99.2% | chr19 | + | 49998130 | 50004928 | 6799 |
| browser | new tab | details | YourSeq | 530 | 78 | 6737 | 6791 | 88.2% | chr19 | - | 41970672 | 42261035 | 290364 |
| browser | new tab | details | YourSeq | 266 | 798 | 6601 | 6791 | 86.0% | chr16 | - | 22285867 | 22428227 | 142361 |

- Read:6792-9504 aligns to chr7:23635867-23638724. Annotations from RepearMasker and the polymorphic SVA dataset defined by Chu et al. indicated it as SVA_E.

| ACTIONS | | | QUERY | SCORE | START | END | QSIZE | IDENTITY | CHROM | STRAND | START | END | SPAN |
|---|---|---|---|---|---|---|---|---|---|---|---|---|---|
| browser | new tab | details | YourSeq | 2401 | 3 | 2712 | 2712 | 96.0% | chr7 | + | 23635867 | 23638724 | 2858 |
| browser | new tab | details | YourSeq | 2356 | 1 | 2712 | 2712 | 97.0% | chr7 | + | 64855505 | 65260248 | 404744 |
| browser | new tab | details | YourSeq | 2293 | 16 | 2712 | 2712 | 95.3% | chr4 | - | 183734274 | 183737817 | 3544 |

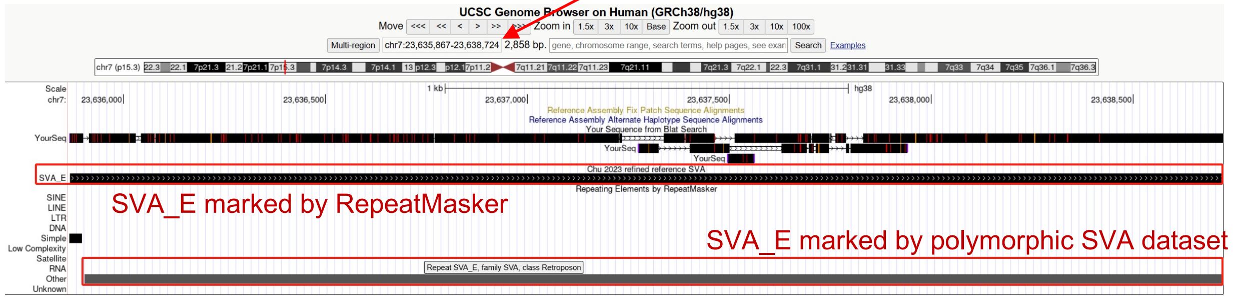


- Illustration:

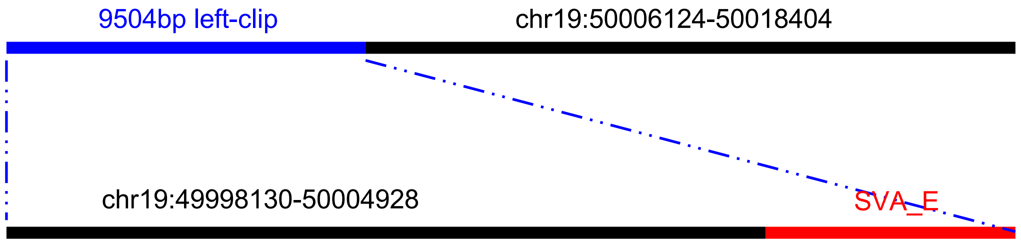


- Read:9505-21791 blat to chr19:50006124-50018404

| ACTIONS | | | QUERY | SCORE | START | END | QSIZE | IDENTITY | CHROM | STRAND | START | END | SPAN |
|---|---|---|---|---|---|---|---|---|---|---|---|---|---|
| browser | new tab | details | YourSeq | 11999 | 27 | 12284 | 12284 | 99.3% | chr19 | + | 50006124 | 50018404 | 12281 |
| browser | new tab | details | YourSeq | 435 | 6469 | 7029 | 12284 | 90.2% | chr18 | + | 50244790 | 50245409 | 620 |

- SVA-mediated deletion (chr19:50004929-50006123)

### NA20129 chr7:128575317

- Using IGV to check the insert in PacBio

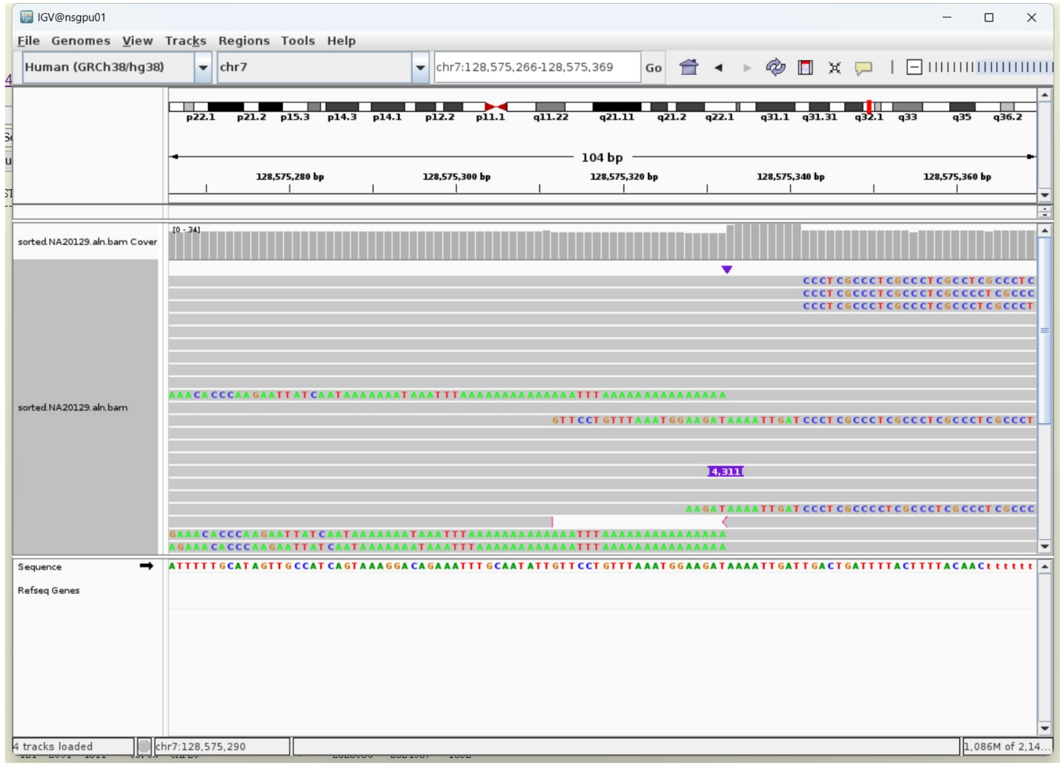


- 8 bp TSD

AAAATTGA

- Blat insert sequence to hg38 genome. Annotations from RepearMasker and the polymorphic SVA dataset defined by Chu et al.(2023) indicated it as SVA_E/F.

| ACTIONS | | | QUERY | SCORE | START | END | QSIZE | IDENTITY | CHROM | STRAND | START | END | SPAN |
|---|---|---|---|---|---|---|---|---|---|---|---|---|---|
| browser | new tab | details | YourSeq | 3542 | 120 | 4298 | 4311 | 99.1% | chr14 | + | 22636430 | 22640013 | 3584 |
| browser | new tab | details | YourSeq | 3216 | 100 | 4296 | 4311 | 95.8% | chr19 | - | 52592756 | 53188600 | 595845 |
| browser | new tab | details | YourSeq | 2981 | 121 | 4311 | 4311 | 97.1% | chr22 | - | 40500369 | 40746385 | 246017 |

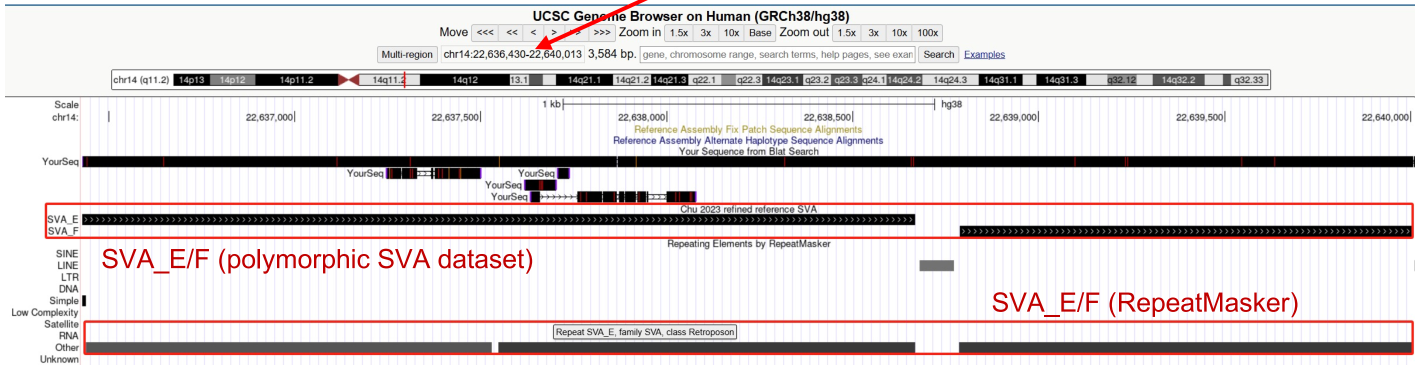


- Blast insert sequence to polymorphic SVA defined by Chu et al. Insertion: 91-2975 aligns to **SVA_E:**1-2881 (at chr17:20092293 of HG002, 96% identity).

```
>HG002_ccs_chr17_20092293
Length=2933

 Score = 4740 bits (5256),  Expect = 0.0
 Identities = 2801/2918 (96%), Gaps = 70/2918 (2%)
 Strand=Plus/Plus

Query  91    tcgccctcgccctcgccctcgccctcgccc-tctccctccacggtctccctcTGATGCCG  149
              || ||||| ||||| ||||| |||||  |  ||||| ||||||||||||||||||||||
Sbjct  1     TCTCCCTCTCCCTCTCCCTCTCCCTCCACAGTCTCCTTCCACGGTCTCCCTCTGATGCCG  60

Query  150   AGCCAAGGCTGGACGGTGCTGCTGCCATCTCGGCTCACTGCAGCCTCCCTGCCTGATTCT  209
             ||||||||||||||||||||||||||||||||||||||||||||||||||||||||||||
Sbjct  61    AGCCAAGGCTGGACGGTGCTGCTGCCATCTCGGCTCACTGCAGCCTCCCTGCCTGATTCT  120
.........
Query  2878  CACTATTGTCCTATGACCCTGCCAAATCCCCCTCTGCGAGAAACACCCAAGAATGATCAA  2937
             |||||||||||||||||||||||||||||||||||| |||||||||||||||||||||||
Sbjct  2784  CACTATTGTCCTATGACCCTGCCAAATCCCCCTCTGTGAGAAACACCCAAGAATGATCAA  2843

Query  2938  TaaaaaaaaaaaaaaaaaaaaaaaaaaaaaaaTTAA  2975
             |||||||||||| |||||| |||||| ||||||| ||
Sbjct  2844  TAAAAAAAAAAAATAAAAAATAAAAAATAAAAAATAAA  2881
```

### ERX2355888 chr13:24314860

- Using IGV to check the insert in PacBio

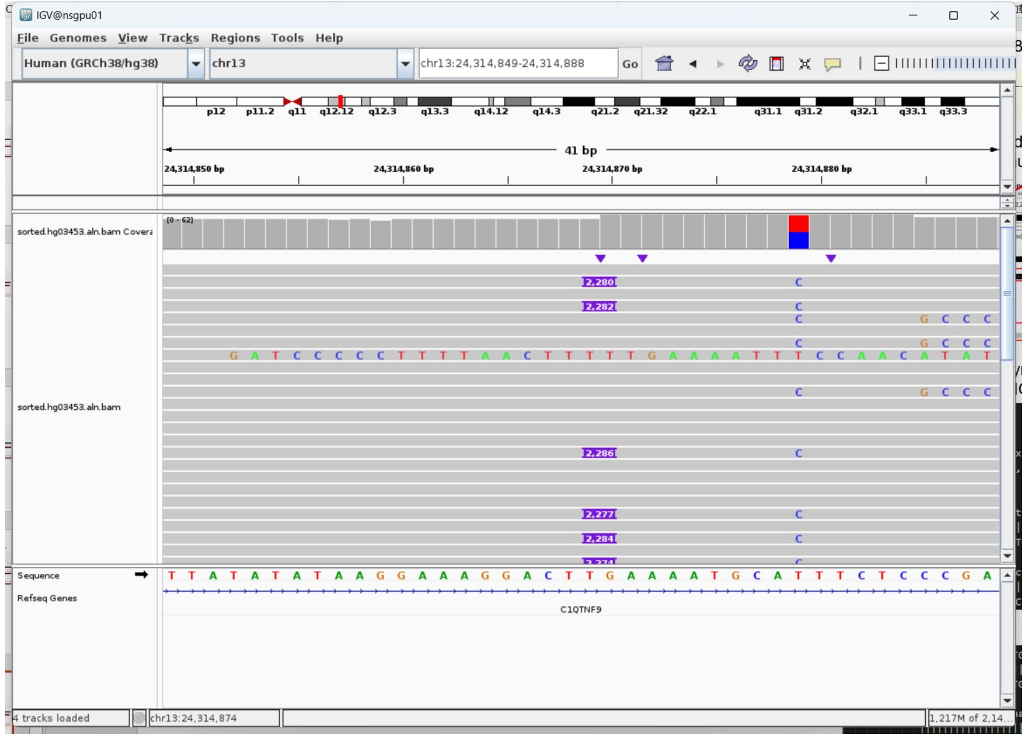


- 9 bp TSD

GAAAATGCA

- Blat insert sequence to hg38 genome. Annotations from RepearMasker and the polymorphic SVA dataset defined by Chu et al.(2023) indicated it as SVA_E.

| ACTIONS | QUERY | SCORE | START | END | QSIZE | IDENTITY | CHROM | STRAND | START | END | SPAN |
|---|---|---|---|---|---|---|---|---|---|---|---|
| browser new tab details | YourSeq | 2206 | 17 | 2261 | 2280 | 99.1% | chr1 | + | 151599499 | 151601739 | 2241 |
| browser new tab details | YourSeq | 2138 | 17 | 2275 | 2280 | 98.5% | chr10 | - | 96940304 | 96942475 | 2172 |

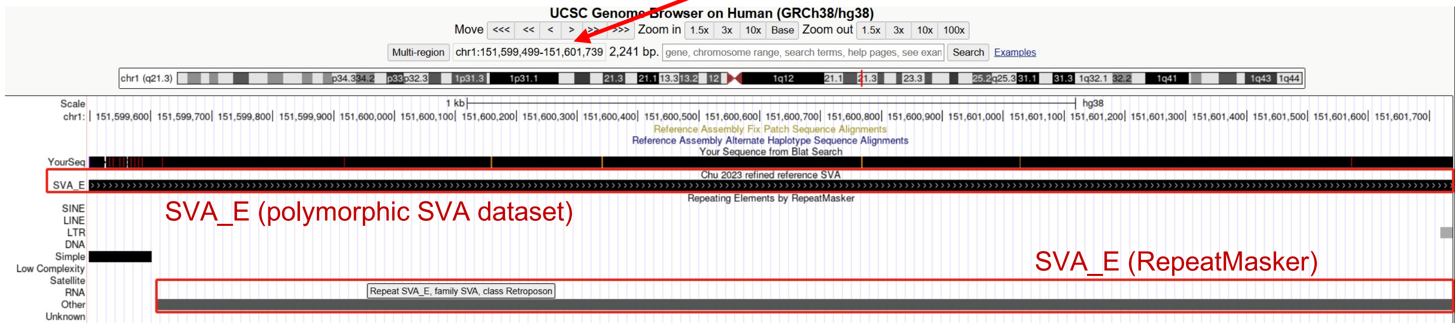


- Blast insert sequence to polymorphic SVA defined by Chu et al. Insertion:13-2257 aligns to **SVA_E:**215-2454 (at chr5:134619330 of HG002, 99% identity).

```
>HG002_ccs_chr5_134619330_2
Length=2454

 Score = 3922 bits (4349),  Expect = 0.0
 Identities = 2223/2246 (99%), Gaps = 7/2246 (0%)
 Strand=Plus/Plus

Query  13    ctcgccctctccctctccctctccctctccctcccccctcccccctcccccctcccccctccc  72
             ||| ||||||||||||||||||||||||||||| ||||| ||||| ||| || ||||| |
Sbjct  215   CTCTCCCTCTCCCTCTCCCTCTCCCTCTCCCTCTCCCTCTCCCTCTCCC-TCTCCCTCTC  273

Query  73    cctccccc-ctccccctccccctccccctccccctccctctccctctctccacggtctcc  131
             |||| ||| ||||| ||||| ||||| ||||| |||||||||||||||||||||||||
Sbjct  274   CCTCTCCCTCTCCCTCTCCCTCTCCCTCTCCCTCTCCCTCTCCCTCTCTCCACGGTCTCC  333
```

………

```
Query  2172  CCTTCCCTCCACTGTTGTCCTATGACCCTGCCAAATCCCCCTCTGCGAGAAACACCCAAG  2231
             ||||||||||||||||||||||||||||||||||||||||||||||||||||||||||||
Sbjct  2369  CCTTCCCTCCACTGTTGTCCTATGACCCTGCCAAATCCCCCTCTGCGAGAAACACCCAAG  2428

Query  2232  AATGATCAATaaaaaaaaaaaaaaaa  2257
             ||||||||||||||||||||||||||
Sbjct  2429  AATGATCAATAAAAAAAAAAAAAAAA  2454
```
