## Extended Files for "Image-based DNA Sequencing Encoding for Detecting Low-Mosaicism Somatic Mobile Element Insertions": Extended Files 4.pdf

### Detailed visualization of benchmarked tumor somatic L1 insertions in HG008-T based on PacBio data

(Additional explanation for Supplementary Data 7)

| Chr | Start | End | L1Hs:Start-End | Supporting Reads | Depth | Allele Frequency | Estimated Mosaicism in PacBio | Manual inspect |
|---|---|---|---|---|---|---|---|---|
| chr2 | 95103766 | 95103767 | L1HS:6064-812 | 2 | 158 | 0.013 | 2.53% | near centromere; 3'transduction; 13bp TSD; ACA TAG |
| chr3 | 137297551 | 137297552 | L1HS:5342-5131:L1HS:6002-6064 | 2 | 238 | 0.008 | 1.68% | inversion; internal deletion; 17bp TSD |
| chr3 | 175249151 | 175249152 | L1HS:5590-6053 | 2 | 224 | 0.009 | 1.79% | 15bp TSD; ACA TAG |
| chr4 | 14331582 | 14331584 | L1HS:6064-5516 | 125 | 190 | 0.658 | 100.00% | short 3'transduction; no TSD; ACA TAG; insert in L1PB1(divergence: 10.0%) |
| chr4 | 112897566 | 112897582 | L1HS:4697-5227 | 89 | 129 | 0.690 | 100.00% | inversion; internal deletion; 3'transduction; 18bp TSD |
| chr4 | 134389657 | 134389677 | L1HS:5867-6064 | 105 | 108 | 0.972 | 100.00% | 19bp TSD; ACA TAG |
| chr5 | 4450306 | 4450341 | L1HS:6058-5827 | 34 | 168 | 0.202 | 40.48% | 3'transduction; 15bp TSD; ACA TAG |
| chr8 | 62600957 | 62600958 | L1HS:4908-6064 | 11 | 223 | 0.049 | 9.87% | 6bp TSD; ACA TAG |
| chr8 | 82189195 | 82189200 | L1HS:5935-6064 | 8 | 200 | 0.040 | 8.00% | 3'transduction; 4bp TSD; TAG |
| chr8 | 104941340 | 104941341 | L1HS:6060-5786;L1HS:5662-5778 | 107 | 229 | 0.467 | 93.45% | inversion; 14bp TSD; ACA TAG |
| chr8 | 107947957 | 107947991 | L1HS:6064-5723;L1HS:5121-5327 | 67 | 184 | 0.364 | 72.83% | Inversion; 16bp TSD; ACA TAG |
| chr9 | 104414095 | 104414109 | L1HS:6064-5441;L1HS:5249-5408 | 8 | 131 | 0.061 | 12.21% | inversion; 14bp TSD; ACA TAG |
| chr10 | 25515229 | 25515242 | L1HS:5893:6064 | 35 | 142 | 0.246 | 49.30% | inversion; 3'transduction; 12bp TSD; ACA TAG |
| chr10 | 84002439 | 84002440 | L1HS:4089-6064 | 3 | 132 | 0.023 | 4.55% | 4bp TSD; ACA TAG |
| chr12 | 7380776 | 7380777 | L1HS:5860-6064 | 134 | 256 | 0.523 | 100% | no TSD; ACA TAG |
| chr20 | 54619796 | 54619809 | L1HS:6064-5238;L1HS:4001-5227 | 10 | 320 | 0.031 | 6.25% | inversion; 18bp TSD; ACA TAG |
| chrX | 69807091 | 69807093 | L1HS:5602-6064 | 101 | 227 | 0.445 | 88.99% | 5bp TSD; ACA TAG |
| chrX | 95987540 | 95987541 | L1HS:4929-6047 | 2 | 185 | 0.011 | 2.16% | 3'transduction; no TSD; ACA TAG |
| chrX | 143137164 | 143137165 | L1HS:5989-6054 | 2 | 185 | 0.011 | 2.16% | 3'transduction; no TSD; TAG |

**Note:**

Insertion sequences of L1 were extracted using the IGV visualization tool and aligned (as query) against the L1Hs reference sequence (as subject) via NCBI BLAST using default parameters. Retrotransposition hallmarks were systematically evaluated, including target site duplication (TSD) and L1Hs hallmark alleles (ACA[5927-5929], TAG[6010-6012]). Structural features characteristic of L1-mediated insertions, including 5' inversions and 3' transduction events, were assessed through established tools such as NCBI BLAST, UCSC BLAT and UCSC genome browser.

In the insertion sequence, red text represents the fragment aligns to L1Hs, while purple text represents the fragment inversely aligns to L1Hs. The yellow background signifies target site duplication (TSD), the blue background signifies L1 hallmark alleles ACA[5927-5929] and TAG[6010-6012], the green background signifies 3' transduction.

Bam files of the tumor and normal datasets were accessed through the following links:
HG008-T (PB-Hifi-1): https://42basepairs.com/download/s3/giab/data_somatic/HG008/Liss_lab/BCM_Revio_20240313/HG008-T_PacBio-HiFi-Revio_20240313_106x_GRCh38-GIABv3.bam

HG008-T (PB-Hifi-2): https://42basepairs.com/download/s3/giab/data_somatic/HG008/Liss_lab/PacBio_Revio_20240125/HG008-T_PacBio-HiFi-Revio_20240125_116x_GRCh38-GIABv3.bam

### chr2:95103767

near centromere; 3'transduction; 13bp TSD; ACA TAG

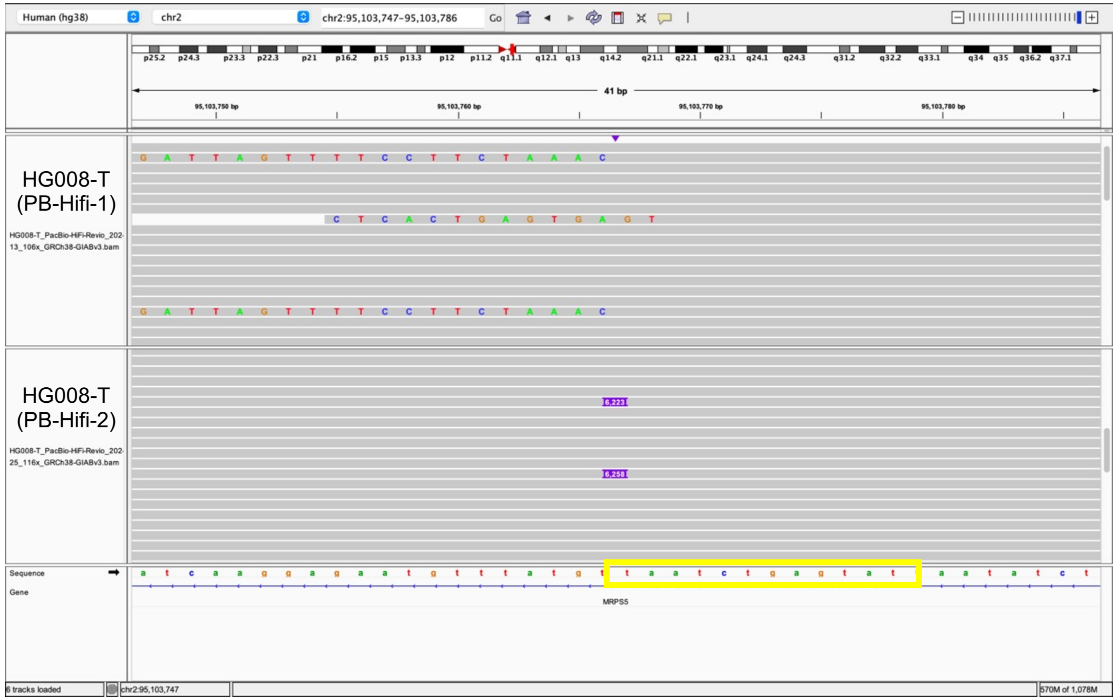


```
Insertion sequence (6223 bp):
3'-TAATCTGAGTATTTTTTTTTTTTTTTTTTTTTTTTTTTTTTTTTTTTTTTTTTTTTTTTTTTTTTTTTTT
TTTTTTTTTCATTCAACAAATATTTATTAAACACCTTCTTTCTTTAAATTCCAAATTCCTCATTTGAGGCAGTCCAAATG
CTCAGCTGTCCCAAGGCTGTTCTACTCTCCATACAAAGCTTTAGTTTCTCCAGCATATCCCCATTACCACATCTCATTTT
CCATCCAGGATTCTTCCTATTTCCATATTTACTCATTTTGCCTGAAATGATCTTCCCTTTTTCCTCCACCAATTCAAAT
CTGAATCATTCTTCAAAGCCTAGCTGAAGTCACCCCTCGCCCCCGGACCCTGGCCTAATTGCCTTCATGCCATTCTGTGT
GACATTATAATATTGAGCTTAAGTTGCATTCCACACAAACACTTGATATTTCTTGATTTGAATAGTTCTATTTTTATTAT
TGTTATTAGAGACAGGACCTTCCTCTGTTTGGAGTGCAGTGACACAACCATAGCTCACTGCAGCCTTAAACTCCTGGGCT
CAAGGGGTCCTCCTTCTTCAGTCTCTCAAGTATCTAGTACTACAAATATATGCCACCACACCCAGATAATTTTTATTGTT
TGTAGAGATAGGGTCTCACGGTATTGTTCAGGCTGGTCTCGAACTTCTGGCTTCAAGCAATCCTCTCATCTCAGCCTCCC
AAAGCACTGAGACTATAGGCATCAGCCATCATGCCTGGCCTGTTCTCTTATTTTTGTTTTTTTTTTTTTTTTTTTTTTTT
TTTTTTTTTTTTTTATTATACTCTAAGTTTTAGGGTACATGTGCACATTGTGCAGGTTAGTTACATATGTATACATGTGC
CATGCTGGTGCGCTGCACCCACTAATGTGTCATCTAGC-5000bp-CTGTTTTTTCCCATCTTTGTGGCTACTTTTGGTC
TTTGATGATGGTGATGTACAGATGGGTTTTCGGTGTAGATGTCCTTTCTGGTTGTTAGTTTTCCTTCTATATATATATAG
TATGTGGCCAAAGGAAACCAGGCCTAACAGCTACTTTTGTAATTTTCCTAAATTTTATGTTCATATTATTATCATACATC
TTTAAAAAATTTACATAAAGGATCACATTATAGCCATAATTTTGCACCTTGATTTTTCTGACATAGTATTATATTCAGA
GATATTAATACTCAGATTAGTTTAGAAGGAAAAC-5'
```

- **12bp TSD**
- Insertion:768-6018 aligns to L1Hs:6064-813, **99% identity**

**Range 1: 813 to 6064** Graphics ▼ Next Match ▲

| Score | Expect | Identities | Gaps | Strand |
|---|---|---|---|---|
| 9520 bits(5155) | 0.0 | 5227/5259(99%) | 15/5259(0%) | Plus/Minus |

- L1 hallmark alleles **ACA** (TGT reserve complement), **TAG** (CTA reserve complement)
- **3'transduction:** Insertion:85-767 aligns to chr14:58753673-58754355, adjacent to a full-length L1 insertion in this subject

| SCORE | START | END | QSIZE | IDENTITY | CHROM | STRAND | START | END | SPAN |
|---|---|---|---|---|---|---|---|---|---|
| 680 | 71 | 753 | 753 | 99.9% | chr14 | - | 58753673 | 58754355 | 683 |
| 152 | 486 | 743 | 753 | 81.9% | chr1 | - | 39579033 | 39579305 | 273 |
| 151 | 493 | 753 | 753 | 80.0% | chr4 | + | 164802283 | 164802553 | 271 |
| 138 | 435 | 707 | 753 | 80.9% | chr17 | + | 42176888 | 42177462 | 575 |
| 136 | 66 | 213 | 753 | 96.0% | chrX | - | 30955387 | 30955534 | 148 |

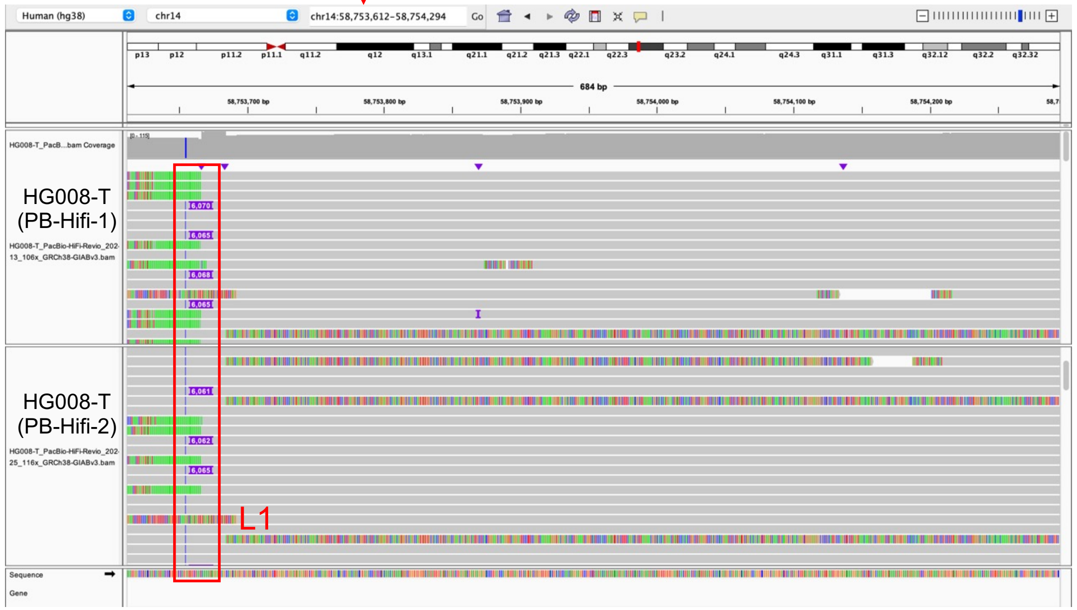


- Illustration of this L1

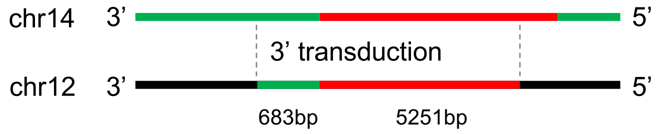

### chr3:137297551

inversion; internal deletion; 17bp TSD

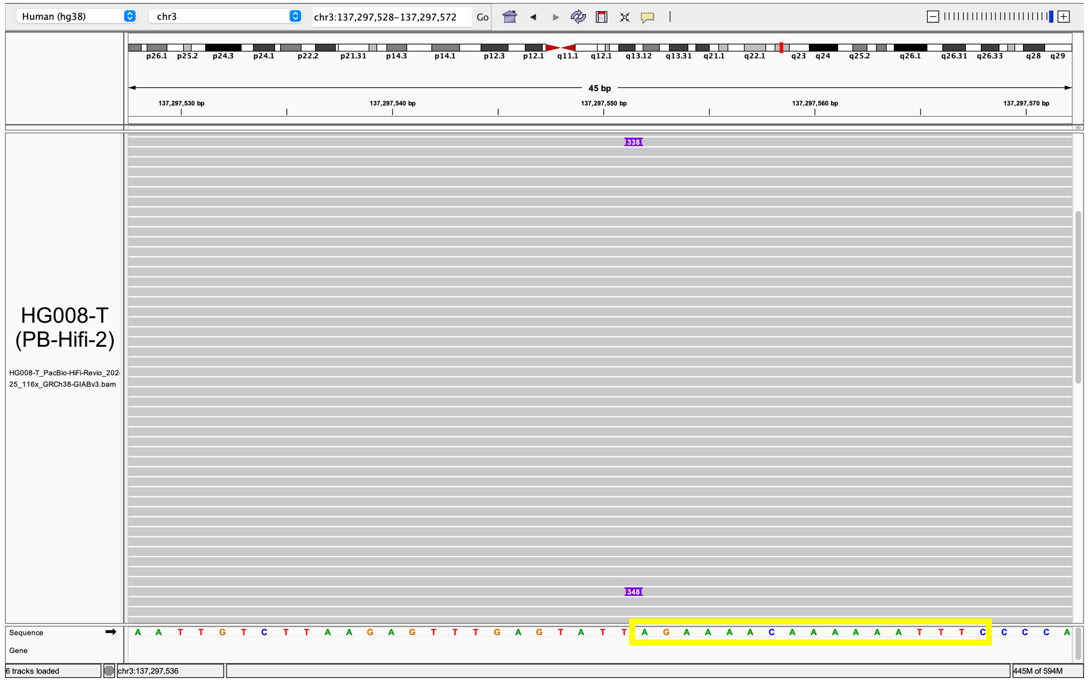


Insertion sequence (338 bp):
5’-AGAAAACAAAAAATTTCTACCTGGTGTGAGATGATATCTCATAGTGGTTTTGATTTGCATTTCTC
TGATGGCCAGTGATGATGAGCATTTCTTCATGGTTTTTTGGCTGCATAAATGTCTTCTTTTGAGAAGT
GTCTGTTCATGTCCTTCGCCCACTTTTTGATGGGGTTGTTTGTTTTTTTCTTGTAAATTTGTTTGAGT
TCATTGTAGATTCTGGATATTAGAGAGTATAATAAAAAAAATAAAATAAAATAAAATAAAAATAAATG
AAAAAAAAAAAAAAAAAAAAAAAAAAAAAAAAAAAAAAAAAAAAAAAAAAAAAAAAAAAAAAAAAAAA
A-3’

- **17bp TSD**
- Insertion:21-224 aligns to L1Hs:5336-5132, **99% identity**

**Range 1: 5132 to 5336** Graphics ▼ Next Match ▲ Previous Match

| Score | Expect | Identities | Gaps | Strand |
|---|---|---|---|---|
| 372 bits(201) | 2e-106 | 204/205(99%) | 1/205(0%) | Plus/Minus |

```
Query  4     CTGGTGTGAGATGATATCTCATAGTGGTTTTGATTTGCATTTCTCTGATGGCCAGTGATG  63
             ||||||||||||||||||||||||||||||||||||||||||||||||||||||||||||
Sbjct  5336  CTGGTGTGAGATGATATCTCATAGTGGTTTTGATTTGCATTTCTCTGATGGCCAGTGATG  5277

Query  64    ATGAGCATTTCTTCATG-GTTTTTTGGCTGCATAAATGTCTTCTTTTGAGAAGTGTCTGT  122
             ||||||||||||||||| ||||||||||||||||||||||||||||||||||||||||||
Sbjct  5276  ATGAGCATTTCTTCATGTGTTTTTTGGCTGCATAAATGTCTTCTTTTGAGAAGTGTCTGT  5217

Query  123   TCATGTCCTTCGCCCACTTTTTGATGGGGTTGTTTGtttttttCTTGTAAATTTGTTTGA  182
             ||||||||||||||||||||||||||||||||||||||||||||||||||||||||||||
Sbjct  5216  TCATGTCCTTCGCCCACTTTTTGATGGGGTTGTTTGTTTTTTTCTTGTAAATTTGTTTGA  5157

Query  183   GTTCATTGTAGATTCTGGATATTAG  207
             |||||||||||||||||||||||||
Sbjct  5156  GTTCATTGTAGATTCTGGATATTAG  5132
```

- The remain sequence aligns to the poly(A) tail of L1, suggest there’s a 5’ inversion event and internal deletion

poly(A): AGAGTATAATAAAAAAAAAAAAAAAAAAAAAAAAAAAAAAAAAAAAAAAAAAATAAATAAAAA

- Illustration of this L1

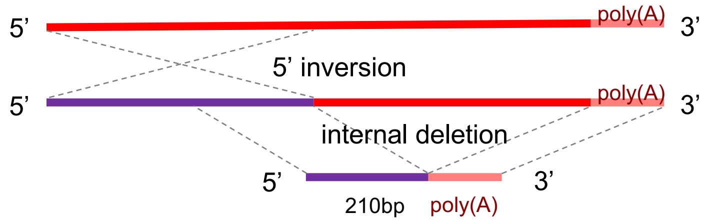

### chr3:175249152

#### 15bp TSD; ACA TAG

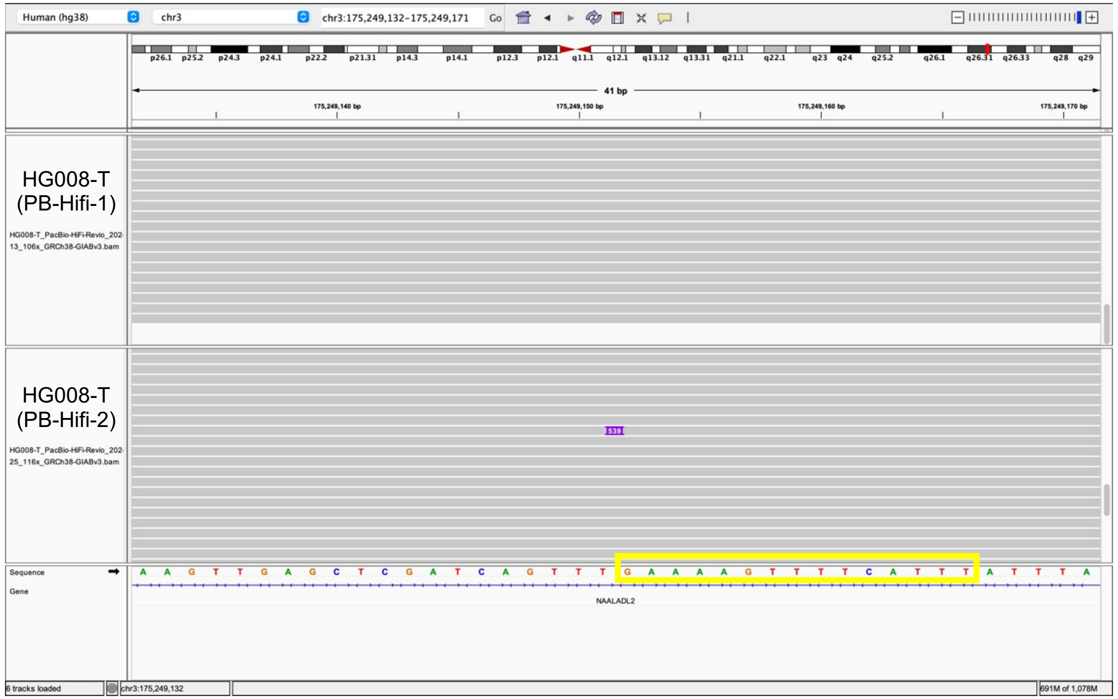


Insertion sequence:

```
5'-GAAAAGTTTTCATTTCACAATAGCAAAGACTTGGAACCAACCCAAATGTCCAACAATGATAGACTGGATT
AAGAAAATGTGGCACATATACACCATGGAATACTATGCAGCCATAAAAAATGATGAGTTCATATCCTTTGTAG
GGACATGGATGAAATTGGAAACCATCATTCTCAGTAAACTATCGCAAGAACAAAAAACCAAACACCGCATATT
CTCACTCATAGGTGGGAATTGAACAATGAGATCACATGGACACAGGAAGGGGAATATCACACTCTGGGGACTG
TGGTGGGGTCGGGGGAGGGGGGAGGGATAGCATTGGGAGATATACCTAATGCTAGATGACACATTAGTGGGTG
CAGCGCACCAGCATGGCACATGTATACATATGTAACTAACCTGCACAATGTGCACATGTACCCTAAAACTTAG
AGTATAATAAAAAAAAAAAAAAAAAAAAAAAAAAAAAAAAAAGAACTGTGCAAAAAAAAAAAAAAAAAAAAAA
AAAAAAAAAAAAAAAAAAAAAAAAAAAAAA-3'
```

- **15bp TSD**
- Insertion:16-477 aligns to L1Hs:5593-6054, **99% identity**

**Range 1: 5593 to 6054** Graphics ▼ Next Match ▲ Previous Match

| Score | Expect | Identities | Gaps | Strand |
|---|---|---|---|---|
| 848 bits(459) | 0.0 | 461/462(99%) | 0/462(0%) | Plus/Plus |

- L1 hallmark alleles **ACA, TAG**

```
Query  301   ATAGCATTGGGAGATATACCTAATGCTAGATGACACATTAGTGGGTGCAGCGCACCAGCA  360
             |||||||||||||||||||||||||||||||||||| |||||||||||||||||||||||
Sbjct  5893  ATAGCATTGGGAGATATACCTAATGCTAGATGACACGTTAGTGGGTGCAGCGCACCAGCA  5952

Query  361   TGGCACATGTATACATATGTAACTAACCTGCACAATGTGCACATGTACCCTAAAACTTAG  420
             ||||||||||||||||||||||||||||||||||||||||||||||||||||||||||||
Sbjct  5953  TGGCACATGTATACATATGTAACTAACCTGCACAATGTGCACATGTACCCTAAAACTTAG  6012

Query  421   AGTATAATaaaaaaaaaaaaaaaaaaaaaaaaaaaaaaaaaa  462
             ||||||||||||||||||||||||||||||||||||||||||
Sbjct  6013  AGTATAATAAAAAAAAAAAAAAAAAAAAAAAAAAAAAAAAAA  6054
```

### chr4:14331602

short 3'transduction; no TSD; ACA TAG; insert in L1PB1(divergence: 10.0%)

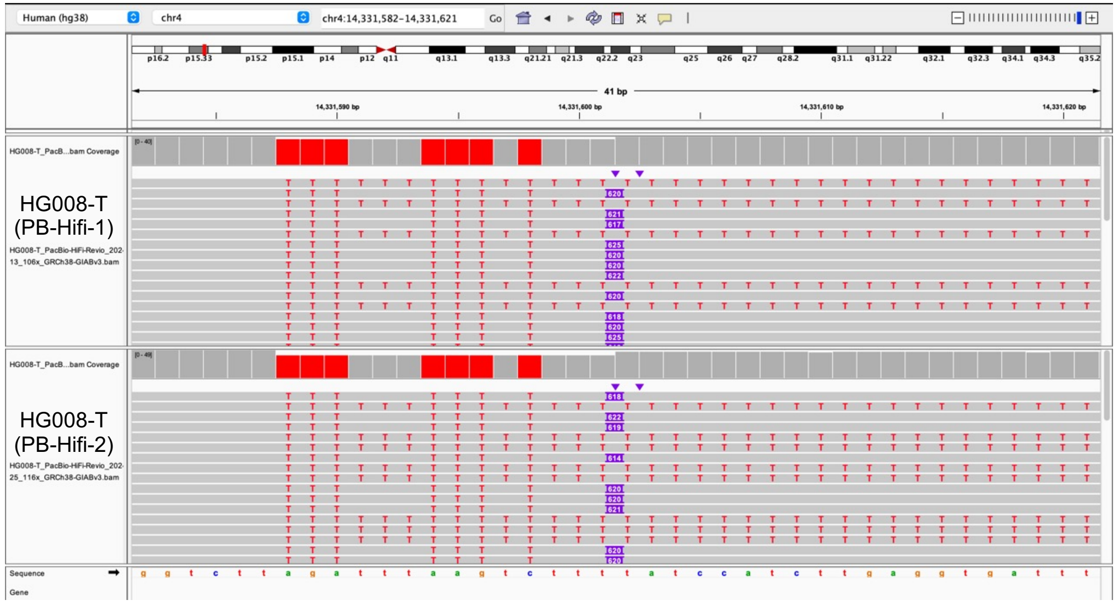


```
Insertion sequence (614 bp):
3'-TTTTTTTTTTTTTTTTTTTTTTTTTTTTTTCCTCTCACATATTCTTCTTTGTTTTATTTATAATTTTTTTTTTTTACAATTCT
TTTTTTTTTTTTTTTTTTTTTTTTTTTATTATACTCTAAGTTTTAGGGTACATGTGCACATTGTGCAGGTTAGTTACATATGTATA
CATGTGCCATGCTGGTGCACTGCACCCACTAATGTGTCATCTAGCATTAGGTATATCTCCCAATGCTATCCCTCCCCCCTCCCCCG
ACCCCACCACAGTCCCCAGAGTGTGATATTCCCCTTCCTGTGTCCATGTGATCTCATTGTTCAATTCCCACCTATGAGTGAGAATA
TGCGGTGTTTGGTTTTTTGTTCTTGCGATAGTTTACTGAGAATGATGGTTTCCAATTTCATCCATGTCCCTACAAAGGATATGAAC
TCATCATTTTTTATGGCTGCATAGTATTCCATGGTGTATATGTGCCACATTTTCTTAATCCAGTCTATCATTGTTGGGCATTTGGG
TTGGTTCCAAGTCTTTGCTATTGTGAATAGTGCCGCAATAAACATATGTGTGCATGTGTCTTTATAGCAGCATGATTTATACTCAT
TTGGGTATATACCCA-5'
```

- This L1 is inserted in L1PB1(divergence: 10.0%)

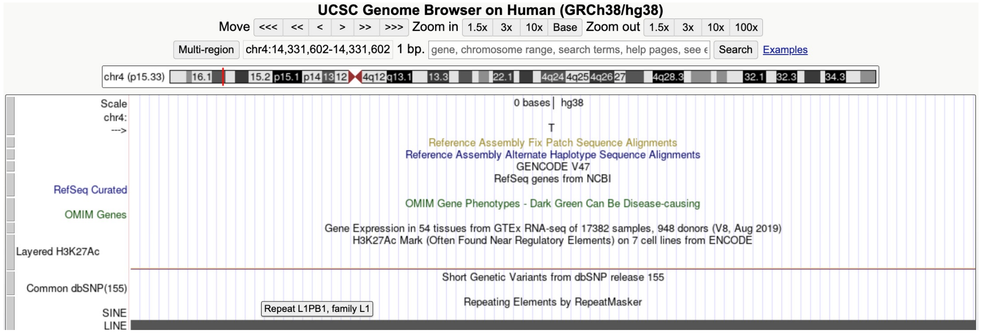


- Insertion:67-614 aligns to L1Hs:6064-5517, **98% identity**

**Range 1: 5517 to 6064** Graphics ▼ Next Match

| Score | Expect | Identities | Gaps | Strand |
|---|---|---|---|---|
| 990 bits(515) | 0.0 | 537/548(98%) | 0/548(0%) | Plus/Minus |

- L1 hallmark alleles **ACA** (TGT reserve complement), **TAG** (CTA reserve complement)
- **3'transduction:** Insertion:1-66 aligns to chr1:82661836-82661928, adjacent to a L1Hs annotated by RepeatMasker

| QUERY | SCORE | START | END | QSIZE | IDENTITY | CHROM | STRAND | START | END | SPAN |
|---|---|---|---|---|---|---|---|---|---|---|
| YourSeq | 65 | 1 | 66 | 66 | 100.0% | chr1 | - | 82661836 | 82661923 | 88 |
| YourSeq | 65 | 1 | 66 | 66 | 100.0% | chr1 | + | 84052256 | 84052345 | 90 |
| YourSeq | 52 | 3 | 66 | 66 | 83.7% | chr9 | + | 12556845 | 12556899 | 55 |
| YourSeq | 42 | 15 | 66 | 66 | 81.4% | chr7 | + | 96846590 | 96846632 | 43 |

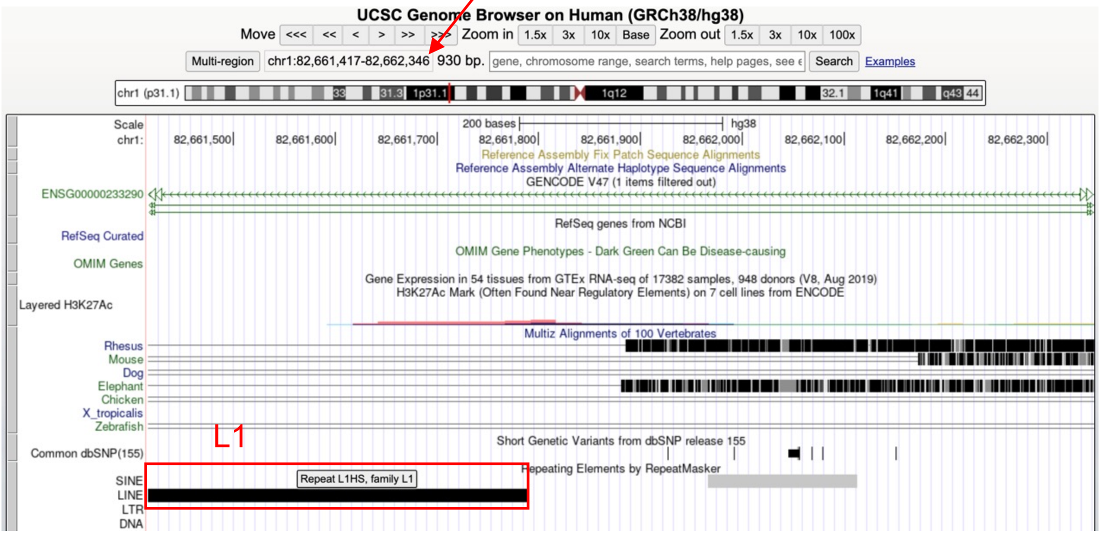


- Illustration of this L1

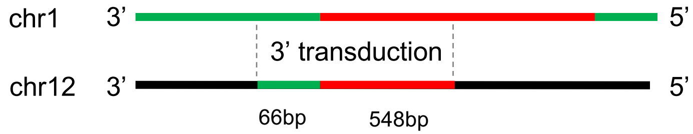

### chr4:112897566

inversion; internal deletion; 3'transduction; 18bp TSD

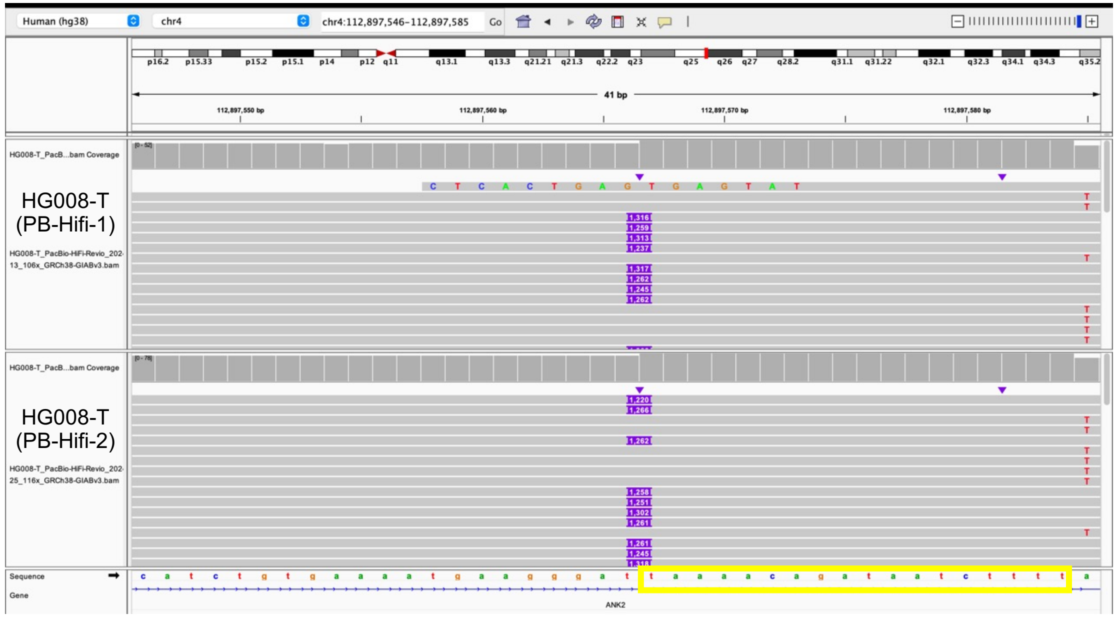


Insertion sequence (1245 bp):

3'-TAAAACAGATAATCTTTTTTTTTTTTTTTTTTTTTTTTTTTTTTTTTTTTTTTTCATTCAACAAATATTT
ATTAAACACCTTCTTTCTTTAAATTCCAAATTCCTCATTTGAAGCAGTCCAAATGCTCAGCTGTCCCAAGGCT
GTTCTACTCTCCATACAAAGCTTTAGTTTCTCCAGCATATCCCCATTACCACATCTCATTTTCCATCCAGGAT
TCTTCCTATTTCCATATTTTTACTCATTTTGCCTGAAATGATCTTCCCTTTTTCCTCCACCAATTCAAATCTG
AATCATTCTTCAAAGCCTAGCTGAAGTCACCCTCGCCCCCGGACCCTGGCCTAATTGCCTTCATGCCATTCTG
TGTGACATTATAATATTGAGCTTAAGTTGCATTCCACACAAACACTTGATATTTCTTGATTTGAATAGTTCTA
TTTTTATTATTGTTAGTAGAGACAGGACCTTCCTCTGTTTGGAGTGCAGTGACACAACCATAGCTCACTGCAG
CCTTAAACTCCTGGGCTCAAGGGGTCCTCCTTCTTCAGTCTCTCAAGTATCTAGTACTACAAATATATGCCAC
CACACCCAGATAATTTTTATTGTTTGTAGAGATAGGGTCTCACGGTATTGTTCAGGCTGGTCTCGAACTTCTG
GCTTCAAGCAATCCTCTCATCTCAGCCTCCCAAAGCACTGAGACTATAGGCATCAGCCATCCCTCAGAAATAA
TGCCGCATATCTACAACTATCTGATCTTTGACAAACCTGAGAAAAACAAGCAATGGGGAAAGGATTCCCTATT
TAATAAATGGTGCTGGGAAAACTGGCTAGCCATATGTAGAAAGCTGAAACTGGATCCCTTCCTTACACCTTAT
ACAAAAATCAATTCAAGATGGATTAAAGATTTAAACGTTAAACCTAAAACCATAAAAACCCTAGAAGAAAACC
TAGGCATTACCATTCAGGACATAGGCGTGGGCAAGGACTTCATGTCCAAAACACCAAAAGCAATGGCAACAAA
AGACAAAATTGACAAATGGGATCTAATTAAACTAAAGAGCTTCTGCACAGCAAAAGAAACTACCATCAGAGTG
AACAGGCAACCTACAACATGGGAGAAAATTTTTGCAACCTACTCATCTGACAAAGGGCTAATATCCAGAATCT
ACAATGAACTCAAACAATTTACAAGAAAAAAACAAACAACCCCATCAAAAAGTGGGCGAAGGACATGAACAG
ACACTTC-5'

- **18bp TSD**

- Insertion:715-1245 aligns to L1Hs:4697-5227, **98% identity**

**Range 1: 4697 to 5227** Graphics ▼ Next Match

| Score | Expect | Identities | Gaps | Strand |
|---|---|---|---|---|
| 1004 bits(522) | 0.0 | 528/531(99%) | 0/531(0%) | Plus/Plus |

- **3'transduction:** Insertion:53-714 of the sequence aligns to chr14:58753694-58754355, adjacent to a full-length L1 insertion in this subject

| SCORE | START | END | QSIZE | IDENTITY | CHROM | STRAND | START | END | SPAN |
|---|---|---|---|---|---|---|---|---|---|
| 658 | 53 | 714 | 714 | 99.7% | chr14 | - | 58753694 | 58754355 | 662 |
| 145 | 468 | 704 | 714 | 81.6% | chr1 | - | 39579053 | 39579305 | 253 |
| 141 | 475 | 713 | 714 | 80.0% | chr3 | - | 49735085 | 50104613 | 369529 |

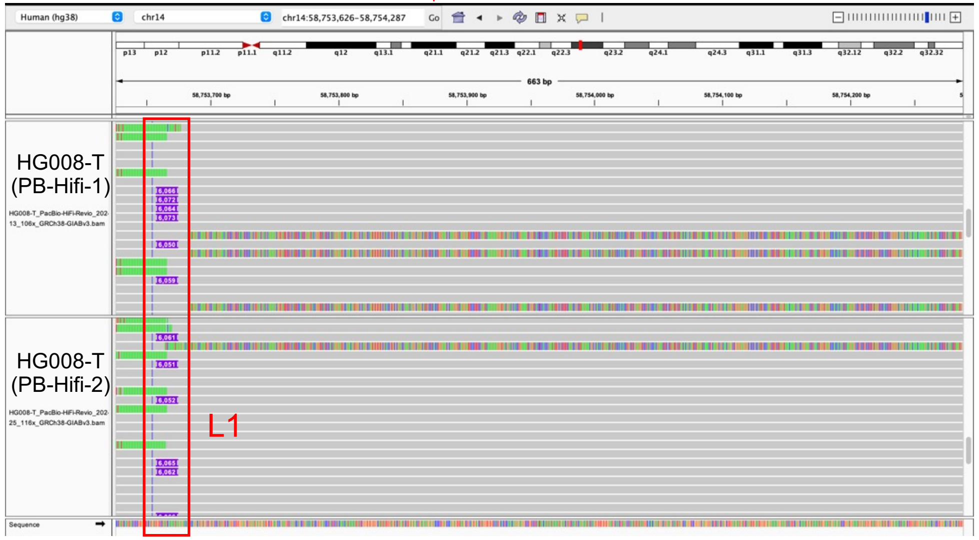


- Illustration of this L1

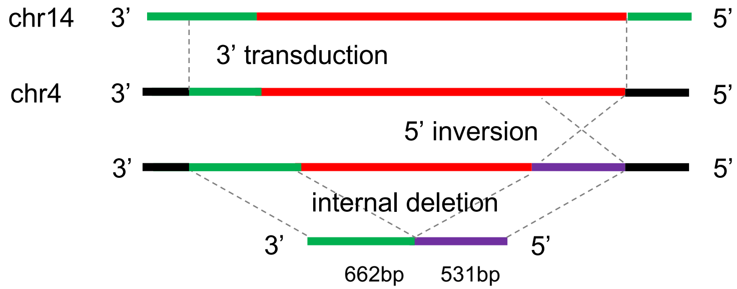

### chr4:134389658

#### 19bp TSD; ACA TAG

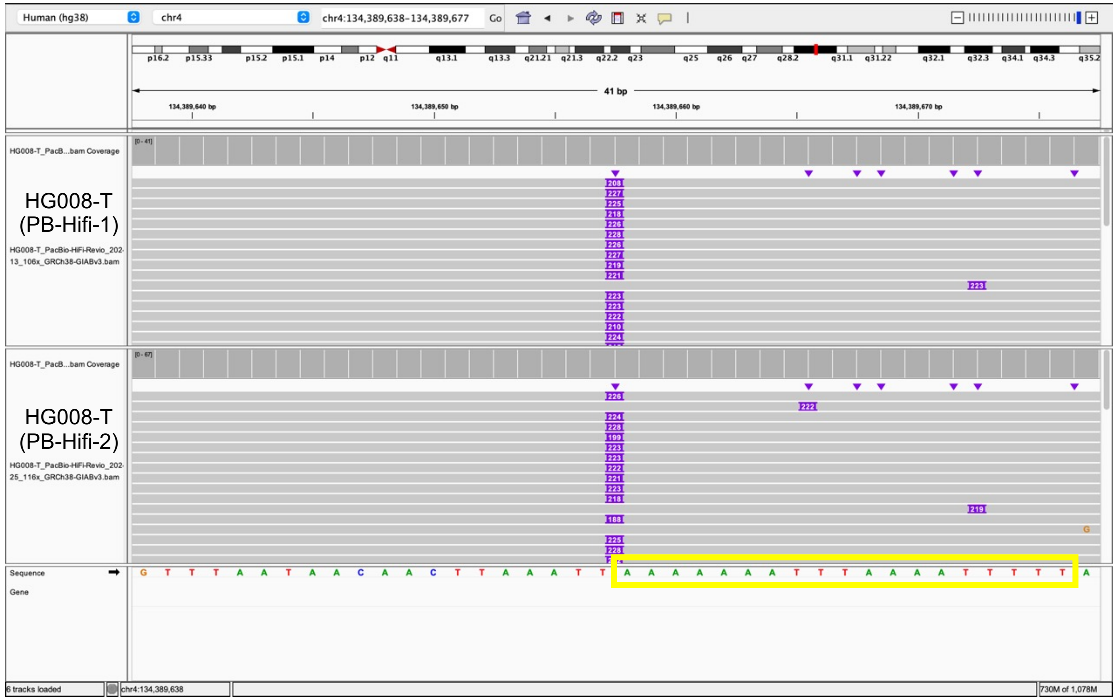


Insertion sequence:
5'-AAAAAAATTTAAAATTTTTGGTGGGGTCGGGGGAGGGGGGAGGGATAGCATTGGGAGATATACCTAATGC
TAGATGACACAATTAGTGGGTGCAGTGCACCAGCATGGCACATGTATACATATGTAACTAACCTGCACAATGTG
CACATGTACCCTAAAACTTAGAGTATAATAAAAAAAAAAAAAAAAAAAAAAAAAAAAAAAAAAAAAAAAAAAAA
AAAAAAAAAAA-3'

- **19bp TSD**
- Insertion:20-216 aligns to L1Hs:5868-6064, **98% identity**

Range 1: 5868 to 6064 Graphics ▼ Next Match ▲ Previous Match

| Score | Expect | Identities | Gaps | Strand |
|---|---|---|---|---|
| 342 bits(185) | 8e-98 | 193/197(98%) | 0/197(0%) | Plus/Plus |

- L1 hallmark alleles **ACA, TAG**

```
Query  1     GGTggggtcgggggaggggggagggATAGCATTGGGAGATATACCTAATGCTAGATGACA  60
Sbjct  5868  GGTGGGGTCGGGGGAGGGGGGAGGGATAGCATTGGGAGATATACCTAATGCTAGATGACA  5927

Query  61    CATTAGTGGGTGCAGTGCACCAGCATGGCACATGTATACATATGTAACTAACCTGCACAA  120
Sbjct  5928  CGTTAGTGGGTGCAGCGCACCAGCATGGCACATGTATACATATGTAACTAACCTGCACAA  5987

Query  121   TGTGCACATGTACCCTAAAACTTAGAGTATAATaaaaaaaaaaaaaaaaaaaaaaaaaaa  180
Sbjct  5988  TGTGCACATGTACCCTAAAACTTAGAGTATAATAAAAAAAAAAAAAAAAAAAAAAAAAAA  6047

Query  181   aaaaaaaaaaaaaaaaa  197
Sbjct  6048  AAAAAAATAAATAAAAA  6064
```

### chr5:4450306

3'transduction; 15bp TSD; ACA TAG

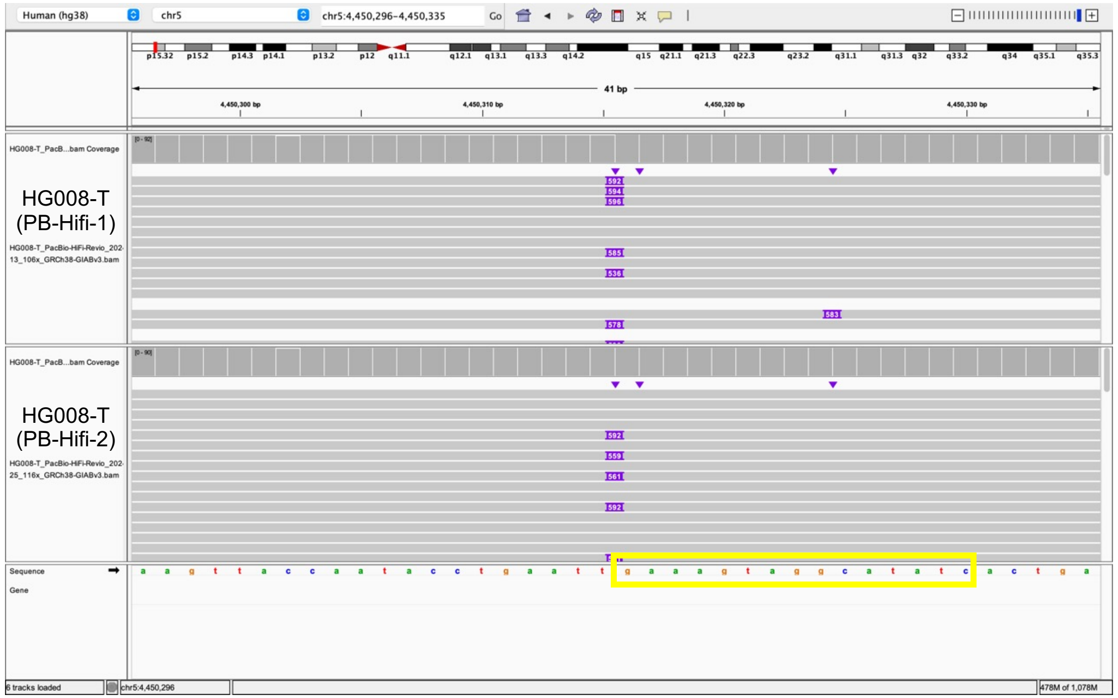


Insertion sequence (592 bp):

3’-GAAAGTAGGCATATCGTACACAAATTTGATGAGTTTTGACACATGCAAACAGCCATGAAACCATCATAAC
AATTAAAATAACAAACATTTCCATGTTTGCTTTGTTTTGTTTGTTTTTTTTTTTTTTTTCATGTGTTTTTTTT
TTTTTTTTTTTTTTTTTTTTTTTTTTATTATACTCTAAGTTTTAGGGTACATGTGCACATTGTGCAGGTTAGT
TACATATGTATACATGTGCCATGCTTGTGCGCTGCACCCACTAATGTGTCATCTAGCATTAGGTATATCTCCC
AATGCTATCCCTCCCCCCTCCCCCGACCCCACCACAGTCCCCAGAGTGTGATATTCCCCTTCCTGTGTCCATG
TAAATTTGTGTACGTTAAATATGTGAAACTTATTGTATGCTGGTTACACCTCAATAAAGCTGTTAAATTTTTT
TAAAAAAAAAAAAAAAAAAAAAAAAAAAGATGTAAGTAGAAATAGCAAAAAGTTAAAAAGCAGGACAAATTAA
AATAGAGTTTTTATTAAAAAAAAAAAAAAAAAAAAAAAAAAAAAAAAAAAAAAAAAAAAAAAAAAAAAAAAAA
AAAAAAAAAAA-5’

- **15bp TSD**
- Insertion:125-363 aligns to L1Hs:6064-5827, **98% identity**

**Range 1: 5827 to 6064** Graphics ▼ Next Match ▲

| Score | Expect | Identities | Gaps | Strand |
|---|---|---|---|---|
| 429 bits(223) | 2e-123 | 233/238(98%) | 0/238(0%) | Plus/Minus |

- L1 hallmark alleles **ACA** (TGT reserve complement), **TAG** (CTA reserve complement)
- **3’transduction:** Insertion:15-124 aligns to chr6:70016374-70016502, with a L1Hs at that location

| SCORE | START | END | QSIZE | IDENTITY | CHROM | STRAND | START | END | SPAN |
|---|---|---|---|---|---|---|---|---|---|
| 109 | 15 | 124 | 124 | 100.0% | chr6 | - | 70016374 | 70016502 | 129 |
| 109 | 15 | 124 | 124 | 100.0% | chrX | + | 11935152 | 11935280 | 129 |
| 109 | 15 | 124 | 124 | 100.0% | chr5 | + | 24370557 | 24370667 | 111 |
| 44 | 23 | 93 | 124 | 79.7% | chr1 | + | 6710935 | 6711003 | 69 |
| 34 | 23 | 66 | 124 | 88.7% | chr1 | + | 210576330 | 210576373 | 44 |

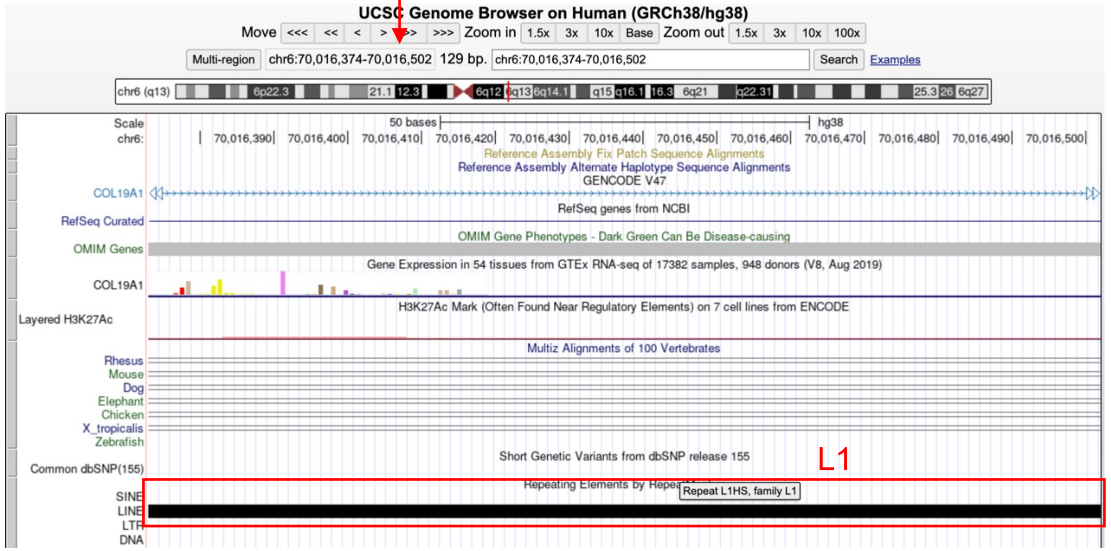


- Illustration of this L1

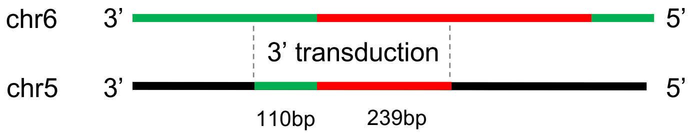

### chr8:62600958

#### 6bp TSD; ACA TAG

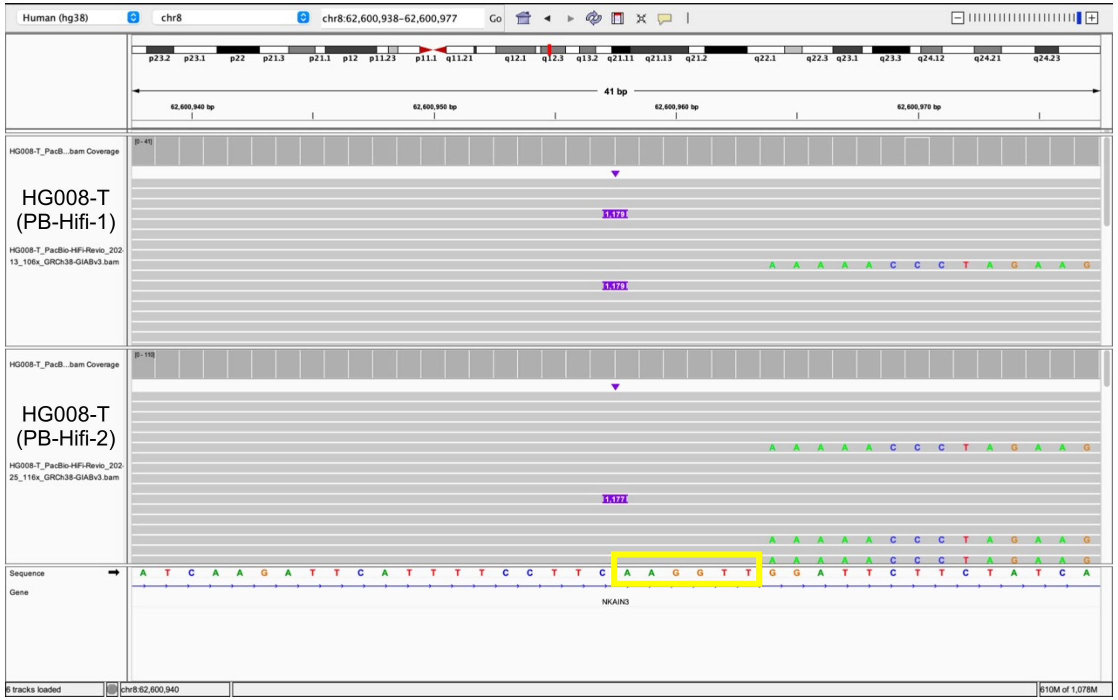


```
Insertion sequence (1179 bp):
5'-AAGGTTAAAAACCCTAGAAGAAAACCTAGGCATTACCATTCAGGACATAGGTGTGGGCAAGGACTTCATG
TCCAAAACACCAAAAGCAATGGCAACAAAAGACAAAATTGACAAATGGGATCTAATTAAACTAAAGAGCTTCT
GCACAGCAAAAGAAACTACCATCAGAGTGAACAGGCAACCTACAACATGGGAGAAAATTTTCGCAACCTACTC
ATCTGACAAAGGGCTAATATCCAGAATCTACAATGAACTCAAACAAATTTACAAGAAAAAAACAAACAACCCC
ATCAAAAAGTGGGCGAAGGACATGAACAGACACTTCTCAAAAGAAGACATTTATGCAGCCAAAAAACACATGA
AGAAATGCTCATCATCACTGGCCATCAGAGAAATGCAAATCAAAACCACTATGAGATATCATCTCACACCAGT
TAGAATGGCAATCATTAAAAAGTCAGGAAACAACAGGTGCTGGAGAGGATGCGGAGAAATAGGAACACTTTTA
CACTGTTGGTGGGACTGTAAACTAGTTCAACCATTGTGGAAGTCAGTGTGGCGATTCCTCAGGGATCTAGAAC
TAGAAATACCATTTGACCCAGCCATCCCATTACTGGGTATATACCCAAATGAGTATAAATCATGCTGCTATAA
AGACACATGCACACGTATGTTTATTGCGGCACTATTCACAATAGCAAAGACTTGGAACCAACCCAAATGTCCA
ACAATGATAGACTGGATTAAGAAAATGTGGCACATATACACCATGGAATACTATGCAGCCATAAAAAATGATG
AGTTCATATCCTTTGTAGGGACATGGATGAAATTGGAAACCATCATTCTCAGTAAACTATCGCAAGAACAAAA
AACCAAACACCGCATATTCTCACTCATAGGTGGGAATTGAACAATGAGATCACATGGACACAGGAAGGGGAAT
ATCACACTCTGGGGACTGTGGTGGGGTGGGGGGAGGGGGGAGGGGATAGCATTGGGAGATATACCTAATGCTA
GATGACACATTAGTGGGTGCAGCGCACCAGCATGGCACATGTATACATATGTAACTAACCTGCACAATGTGCA
CATGTACCCTAAAACTTAGAGTATAATAAAAAAAAAAAAAAAAAAAAAAAAAAAAAAAAAAAAAAAAAAAAAAA
AAAAAAAAAAAAA-3'
```

- **6bp TSD**
- Insertion:7-1163 aligns to L1Hs:4909-6064, **99% identity**

**Range 1: 4909 to 6064** Graphics ▼ Next Match ▲ Previous Match

| Score | Expect | Identities | Gaps | Strand |
|---|---|---|---|---|
| 2080 bits(1126) | 0.0 | 1147/1157(99%) | 1/1157(0%) | Plus/Plus |

- L1 hallmark alleles **ACA, TAG**

```
Query  961   gtggggtggggggaggggggaggggATAGCATTGGGAGATATACCTAATGCTAGATGACA  1020
             ||||||| |||||||||||||| |||||||||||||||||||||||||||||||||||||
Sbjct  5869  GTGGGGTCGGGGGAGGGGGGA-GGGATAGCATTGGGAGATATACCTAATGCTAGATGACA  5927

Query  1021  CATTAGTGGGTGCAGCGCACCAGCATGGCACATGTATACATATGTAACTAACCTGCACAA  1080
             | ||||||||||||||||||||||||||||||||||||||||||||||||||||||||||
Sbjct  5928  CGTTAGTGGGTGCAGCGCACCAGCATGGCACATGTATACATATGTAACTAACCTGCACAA  5987

Query  1081  TGTGCACATGTACCCTAAAACTTAGAGTATAATaaaaaaaaaaaaaaaaaaaaaaaaaaa  1140
             ||||||||||||||||||||||||||||||||||||||||||||||||||||||||||||
Sbjct  5988  TGTGCACATGTACCCTAAAACTTAGAGTATAATAAAAAAAAAAAAAAAAAAAAAAAAAAA  6047
```

### chr8:82189196

3'transduction; 4bp TSD; TAG

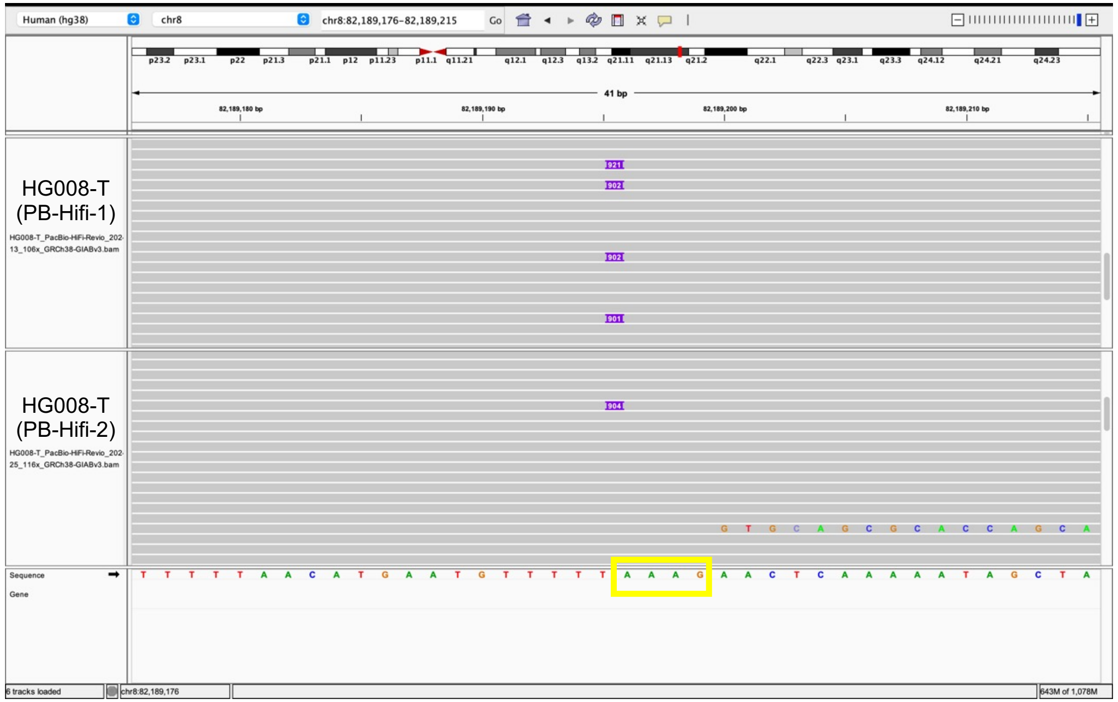


Insertion sequence (921 bp):

5’-AAAGGTGCAGCGCACCAGCATGGCACATGTATACATATGTAACTAACCTGCACAATGTGCACATGTACCC
CTAAAACTTAGAGTATAATAAAAAAAAAAAAAAAAAAAAAAAAAAAAAAAAAAACAAAAATAAGAGAACAGGC
CAGGCATGATGGCTGATGCCTATAGTCTCAGTGCTTTGGGAGGCTGAGATGAGGATTGCTTGAAGCCAGAA
GTTCGAGACCAGCCTGAACAATACCGTGAGACCCTATCTCTACAAACAATAAAAATTATCTGGGTGTGGTGGC
ATATATTTGTAGTACTAGATACTTGAGAGACTGAAGAAGGAGGACCCCTTGAGCCCAGGAGTTTAAGGCTGCA
GTGAGCTATGGTTGTGTCACTGCACTCCAAACAGAGGAAGGTCCTGTCTCTAATAACAATAATAAAAATAGAA
CTATTCAAATCAAGAAATATCAAGTGTTTGTGTGGAATGCAACTTAAGCTCAATATTATAATGTCACACAGAA
TGGCATGAAGGCAATTAGGCCAGGGTCCGGGGGCGAGGGGTGACTTCAGCTAGGCTTTGAAGAATGATTCAGA
TTTGAATTGGTGGAGGAAAAAGGGGAAGATCATTTCAGGCAAAATGAGTAAAAATATGGAAATAGGAAGAATC
CTGGATGGAAAATGAGATGTGGTAATGGGGATATGCTGGAGAAACTAAAGCTTTGTATGGAGAGTAGAACAGC
CTTGGGACAGCTGAGCATTTGGACTGCCTCAAATGAGGAATTTGGAATTTAAAGAAGAAGGTGTTTAATAAA
TATTTGTTGAATGAAAAAAAAAAAAAAAAAAAAAAAAAAAAAAAAAAAAAAAAAAAAAAAAAAAAAAAAAAA
AAAAAAAAAAAAAAAAAAAAAAAAAAAAAAAAAAAAAAAAAAAAAAAA-3’

- **4bp TSD**
- Insertion:5-123 aligns to L1Hs:5937-6054, **99% identity**

**Range 1: 5937 to 6054** Graphics ▼ Next Match ▲ Previous Match

| Score | Expect | Identities | Gaps | Strand |
|---|---|---|---|---|
| 213 bits(115) | 3e-58 | 118/119(99%) | 1/119(0%) | Plus/Plus |

- L1 hallmark alleles **TAG**
- **3’transduction** Insertion:125-814 aligns to chr14:58753667-58754355, adjacent to a full-length L1 in this subject

| SCORE | START | END | QSIZE | IDENTITY | CHROM | STRAND | START | END | SPAN |
|---|---|---|---|---|---|---|---|---|---|
| 687 | 2 | 692 | 798 | 99.3% | chr14 | + | 58753667 | 58754355 | 689 |
| 154 | 23 | 275 | 798 | 82.0% | chr1 | + | 39578610 | 39579305 | 696 |
| 146 | 14 | 268 | 798 | 79.6% | chr4 | – | 164802283 | 164802539 | 257 |
| 136 | 550 | 697 | 798 | 96.0% | chrX | + | 30955387 | 30955534 | 148 |
| 127 | 30 | 268 | 798 | 84.0% | chr17 | + | 34729748 | 34729999 | 252 |

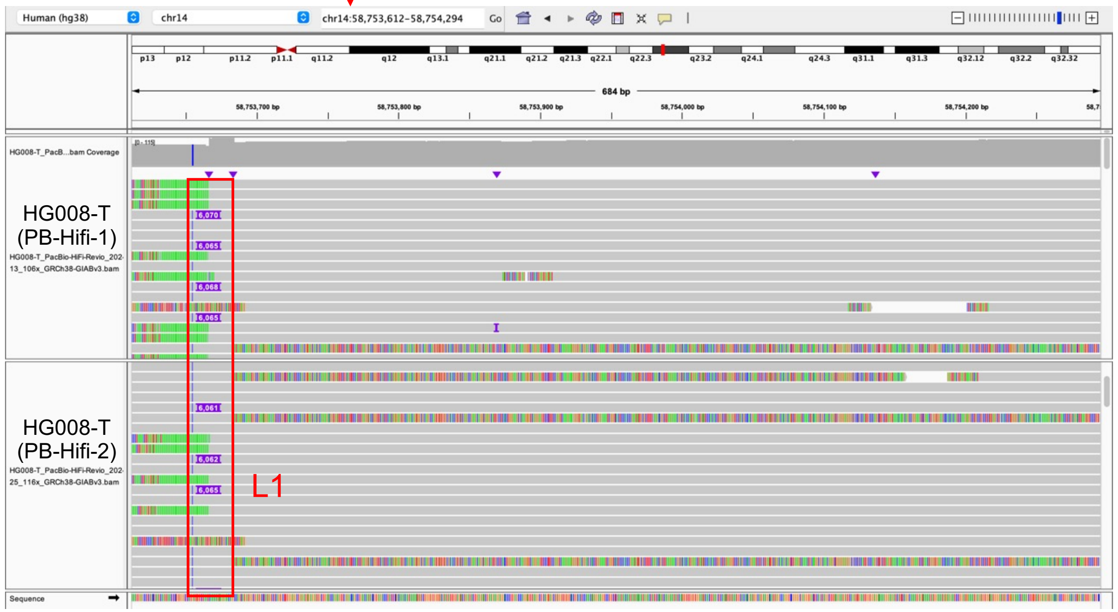


- Illustration of this L1

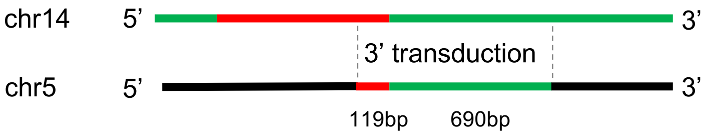

### chr8:104941341

inversion; 14bp TSD; ACA TAG

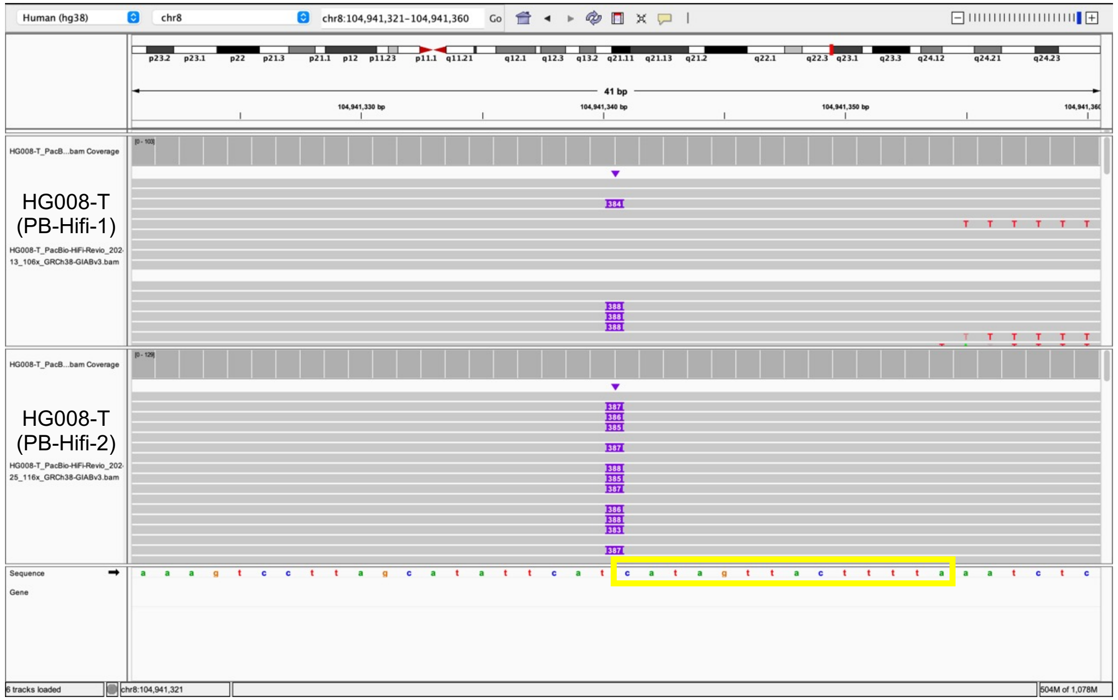


```
Insertion sequence (389 bp):
3’-CATAGTTACTTTTATTTTTTTTTTTTTTTTTTTTTTTTTTTTTTTATTATACTCTAAGTTTTAGGGTACA
TGTGCACATTGTGCAGGTTAGTTACATATGTATACATGTGCCATGCTGGTGCGCTGCACCCACTAATGTGTCA
TCTAGCATTAGGTATATCTCCCAATGCTATCCCTCCCCCCTCCCCCGACCCCACCACAGTCCCCAGAGTGTGA
TATTCCCCTTCCTGTGTCCATGTGATCTCATTGTTCAATTCCCACCTATGAGTGAGAAAATACACCATGGAAT
ACTATGCAGCCATAAAAAATGATGAGTTCATATCCTTTGTAGGGACATGGATGAAATTGGAAACCATCATTCT
CAGTAAACTATCGCAAGAACAAAAAAC-5’
```

- **14bp TSD**

- **5’ inversion**: Two fragments of the sequence aligns to L1Hs:

  (1) Insertion:15-275 aligned to L1Hs:6051-5789, **99% identity**; L1 hallmark alleles **ACA** (TGT reserve complement), **TAG** (CTA reserve complement)

**Range 1: 5789 to 6051** Graphics ▼ Next Match ▲ Previous Match

| Score | Expect | Identities | Gaps | Strand |
|---|---|---|---|---|
| 475 bits(257) | 1e-137 | 261/263(99%) | 0/263(0%) | Plus/Minus |

```
Query  1     tttttttttttttttttttttttttttttttATTATACTCTAAGTTTTAGGGTACATGTG  60
             ||||||||||||||||||||||||||||||||||||||||||||||||||||||||||||
Sbjct  6051  TTTTTTTTTTTTTTTTTTTTTTTTTTTTTTTATTATACTCTAAGTTTTAGGGTACATGTG  5992

Query  61    CACATTGTGCAGGTTAGTTACATATGTATACATGTGCCATGCTGGTGCGCTGCACCCACT  120
             ||||||||||||||||||||||||||||||||||||||||||||||||||||||||||||
Sbjct  5991  CACATTGTGCAGGTTAGTTACATATGTATACATGTGCCATGCTGGTGCGCTGCACCCACT  5932

Query  121   AATGTGTCATCTAGCATTAGGTATATCTCCCAATGCTATccctcccccctcccccgaccc  180
             || |||||||||||||||||||||||||||||||||||||||||||||||||||||||||
Sbjct  5931  AACGTGTCATCTAGCATTAGGTATATCTCCCAATGCTATCCCTCCCCCCTCCCCCGACCC  5872
```

  (2) Insertion:276-389 aligns to L1Hs:5665-5778, 100% identity

**Range 2: 5665 to 5778** Graphics ▼ Next Match ▲ Previous Match

| Score | Expect | Identities | Gaps | Strand |
|---|---|---|---|---|
| 211 bits(114) | 4e-58 | 114/114(100%) | 0/114(0%) | Plus/Plus |

- Illustration of this L1

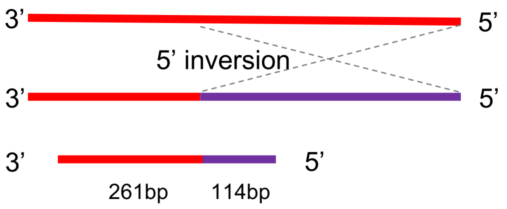

### chr8:107947966

inversion; 16bp TSD; ACA TAG

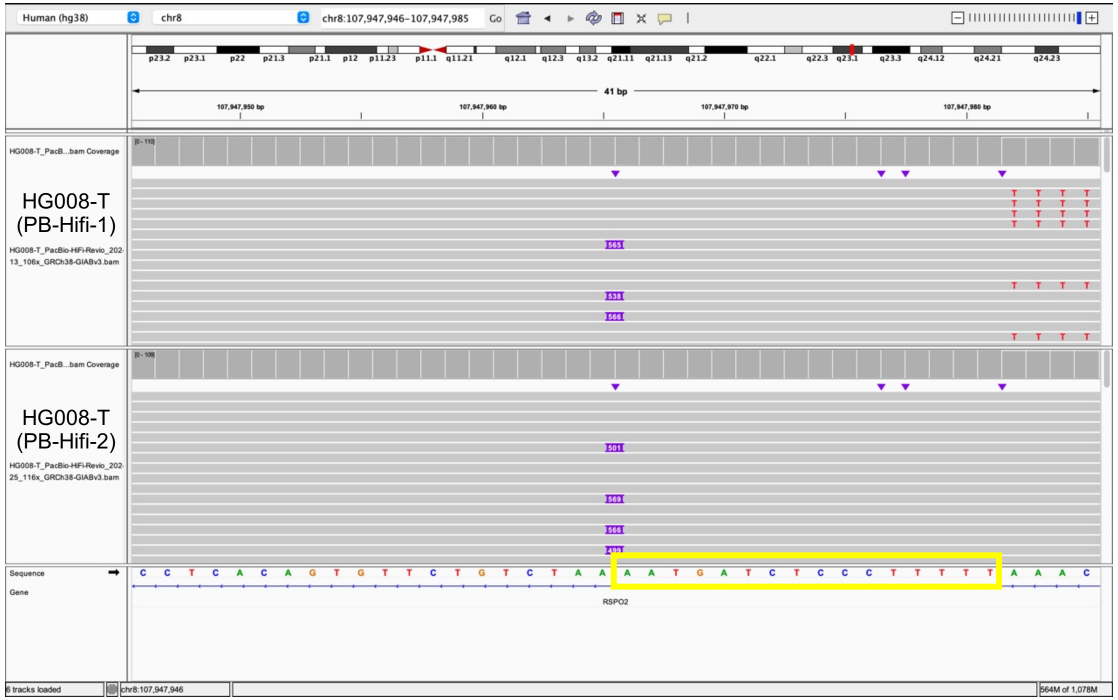


Insertion sequence (565 bp):
3’-AATGATCTCCCTTTTTTTTTTTTTTTTTTTTTTTTTTTTTTTTTTTTTTTTTTTTTTTTTTTTTTTTTTTTTTT
TTTTTTTTTTTTTTTTTTTTTTTTATTATACTCTAAGTTTTAGGGTACATGTGCACATTGTGCAGGTTAGTTA
CATATGTATACATGTGCCATGCTGGTGCACTGCACCCACTAATGTGTCATCTAGCATTAGGTATATCTCCCAA
TGCTATCCCTCCCCCCTCCCCCGACCCCACCACAGTCCCCAGAGTGTGATATTCCCCTTCCTGTGTCCATGTG
ATCTCATTGTTCAATTCCCACCTATGAGTGAGAATATGCGGTGTTTGGTTTTTTGTTCTTGCGATAGTTTACA
AAGGGCTAATATCCAGAATCTACAATGAACTCAAACAAATTTACAAGAAAAAAACAAACAACCCCATCAAAAA
GTGGGCGAAGGACATGAACAGACACTTCTCAAAAGAAGACATTTATGCAGCCAAAAAACACATGAAGAAATGC
TCATCATCACTGGCCATCAGAGAAATGCAAATCAAAACCACTATAAACTACCATCAG-5

- **16bp TSD**
- **5’ inversion**: Two fragments of the sequence aligns to L1Hs:

(1) Insertion:51-361 aligns to L1Hs:6064-5754, **99% identity**; L1 hallmark alleles **ACA** (TGT reserve complement), **TAG** (CTA reserve complement)

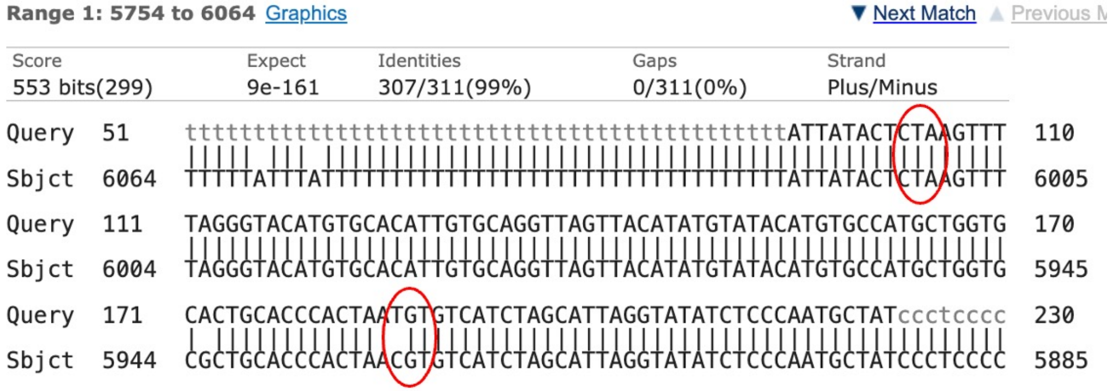


(2) Insertion:360-552 aligns to L1Hs:5124-5316 100% identity

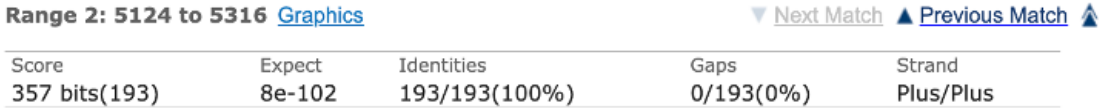


- Illustration of this L1

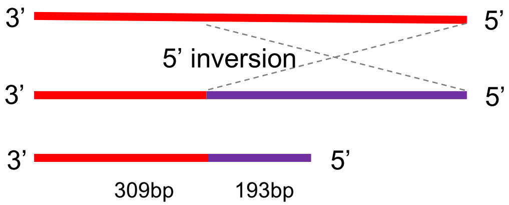

### chr9:104414096

inversion; 14bp TSD; ACA TAG

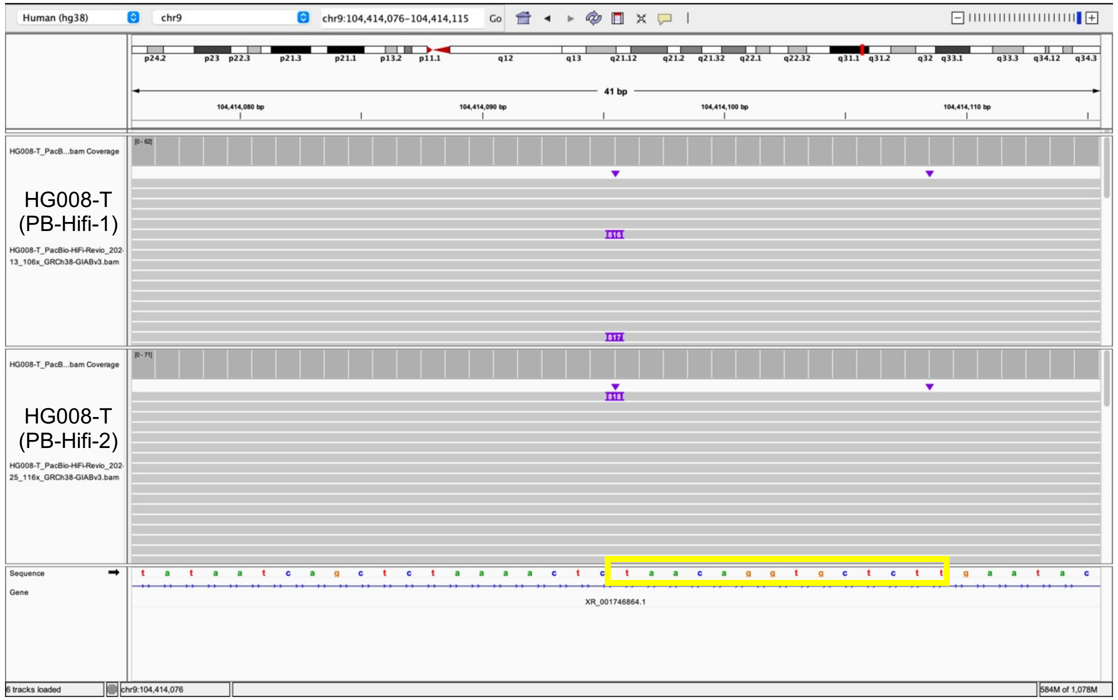


Insertion sequence (816 bp):

3’-TAACAGGTGCTCTTTTTTTTTTTTTTTTTTTTTTTTTTTTTTTTTTTTTTTTTTTTTTTTTTTTTTTTTCTTTTTTCTT
TCTTTTTTTTTTTTTTTTATTATACTCTAAGTTTTAGGGTACATGTGCACATTGTGCAGGTTAGTTACATATGT
ATACATGTGCCATGCTGGTGCGCTGCACCCACTAATGTGTCATCTAGCATTAGGTATATCTCCCAATGCTATC
CCTCCCCCCTCCCCCTACCCCACCACAGTCCCCAGAGTGTGATATTCCCCTTCCTGTGTCCATGTGATCTCAT
TGTTCAATTCCCACCTATGAGTGAGAATATGCGGTGTTTGGTTTTTGTTCTTGCGATAGTTTACTGAGAATGA
TGGTTTCCAATTTCATCCATGTCCCTACAAAGGATATGAACTCATCATTTTTTATGGCTGCATAGTATTCCAT
GGTGTATATGTGCCACATTTTCTTAATCCAGTCTATCATTGTTGGACATTTGGGTTGGTTCCAAGTCTTTGCT
ATTGTGAATAGTGCCGCAATAAACATACGTGTGCATGTGTCTTTATAGCAGCATGATTTATACTCATTTGGGT
ATATACCCAGTAATGGGATGGCTGGGTCAAATGGTATTTCTAGTTCTAGATCCCTGAGGAATCGCCACACTGA
CTTCCACAATGCACATGAAGAAATGCTCATCATCACTGGCCATCAGAGAAATGCAAATCAAAACCACTATGAG
ATATCATCTCACACCAGTTAGAATGGCAATCATTAAAAAGTCAGGAAACAACAGGTGCTGGAGAGGATGCGGA
GAAATAGGAACACTTT-5’

- **14bp TSD**
- **5’ inversion**: Two fragments of the sequence aligns to L1Hs:

(1) Insertion:41-665 aligned to L1Hs:6064-5442, **98% identity**; L1 hallmark alleles **ACA** (TGT reserve complement), **TAG** (CTA reserve complement)

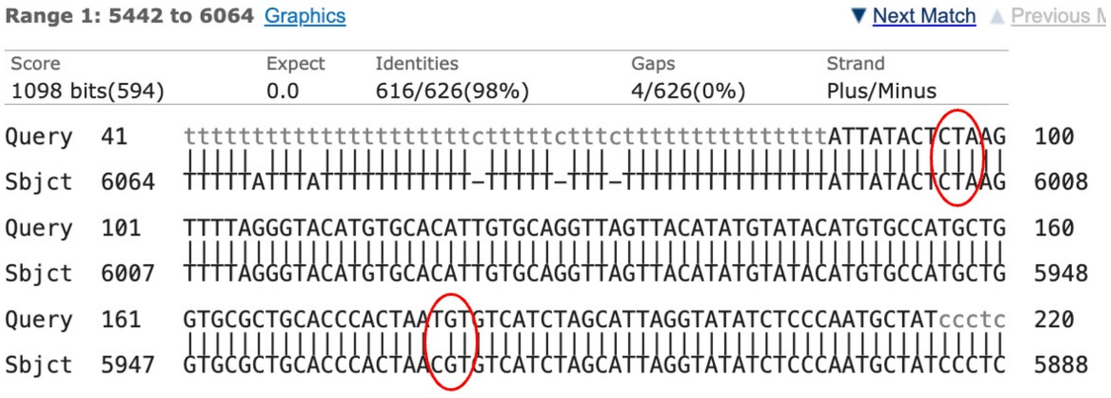


(2) Insertion:666-816 aligns to L1Hs:5258-5408, **99% identity**

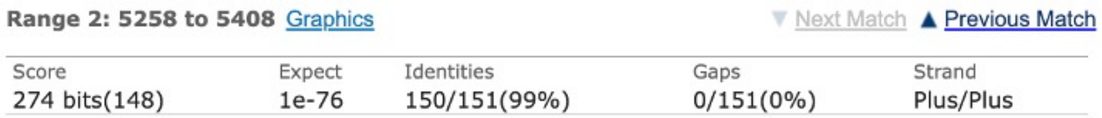


- Illustration of this L1

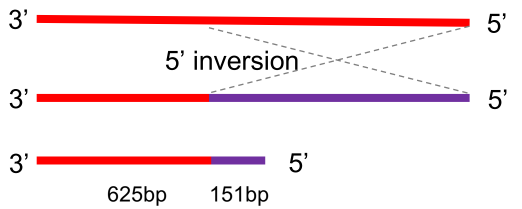

### chr10:25515230

inversion; 3'transduction; 12bp TSD; ACA TAG

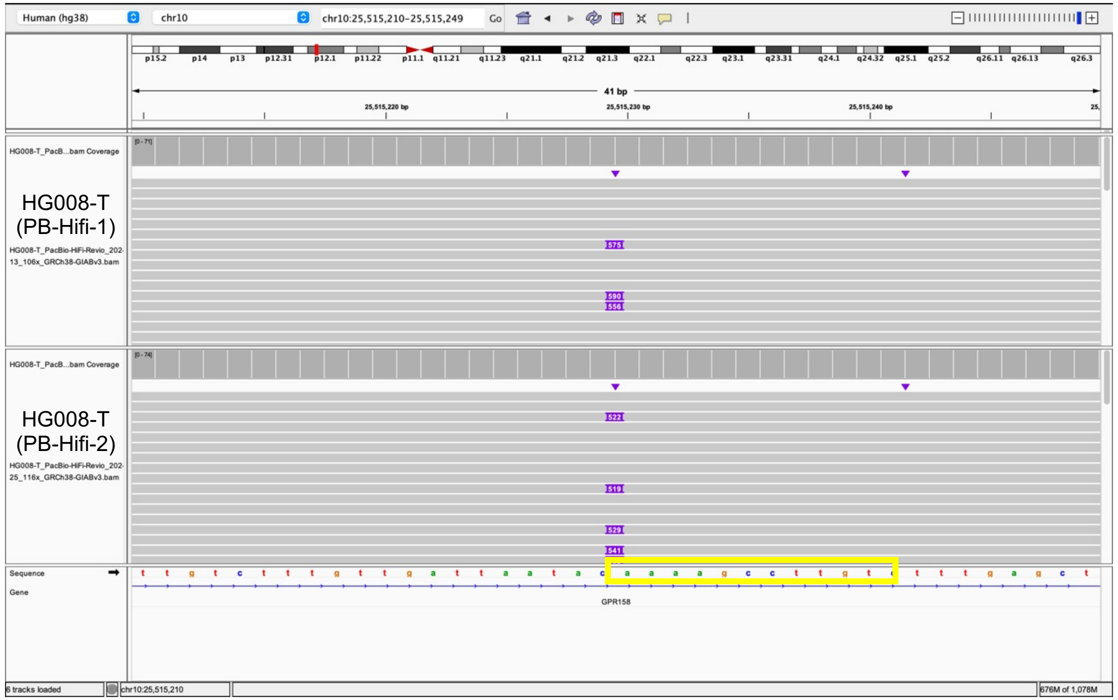


Insertion sequence (575 bp):

5’-AAAAGCCTTGTCATCTAGCATTAGGTATATCTCCCAATGCTATGATGACACATTAGTGGGTGCAGCGCAC
CAGCATGGCACATGTATACATATGTAACTAACCTGCACAATGTGCACATGTACCCTAAAACTTAGAGTATAAT
AAAAAAAAAAAAAAAAAAAAAAAAAAAAAAAAAACAAAAATAAGAGAACAGGCCAGGCATGATGGCTGATGCC
TATAGTCTCAGTGCTTTGGGAGGCTGAGATGAGAGGATTGCTTGAAGCCAGAAGTTCGAGACCAGCCTGAACA
ATACCGTGAGACCCTATCTCTACAAACAATAAAAATTATCTGGGTGTGGTGGCATATATTTGTAGTACTAGAT
ACTTGAGAGACTGAAGAAGGAGGACCCCTTGAGCCCAGGAGTTTAAGGCTGCAGTGAGCTATGGTTGTGTCAC
TGCACTCCAAACAGAGGAAGGTCCTGTCTCTAATAACAATAATAAAAATAGAACTATTCAAATCAAGAAAAAA
AAAAAAAAAAAAAAAAAAAAAAAAAAAAAAAAAAAAAAAAAAAAAAAAAAAAAAAAAAAAAAAAAA-3’

- **12bp TSD**

- **5’ inversion**: Two fragments of the insertion sequence aligns to L1Hs:
  - (1) Insertion:13-43 aligns to L1Hs: 5923-5893, **100% identity**
  - (2) Insertion:44-181 aligned to L1Hs:5921-6058, **99% identity**; L1 hallmark alleles **ACA, TAG**

- **3’transduction:** Insertion:182-506 aligns to chr14:58753671-58753994 (100% identity), with a full-length L1 insertion adjacent to that location in this subject, suggesting a 3’transduction event

| SCORE | START | END | QSIZE | IDENTITY | CHROM | STRAND | START | END | SPAN |
|---|---|---|---|---|---|---|---|---|---|
| 324 | 1 | 324 | 394 | 100.0% | chr14 | + | 58753671 | 58753994 | 324 |
| 152 | 13 | 270 | 394 | 81.9% | chr1 | + | 39579033 | 39579305 | 273 |
| 146 | 9 | 263 | 394 | 79.6% | chr4 | - | 164802283 | 164802539 | 257 |
| 139 | 42 | 270 | 394 | 80.8% | chr6 | - | 154534442 | 154534682 | 241 |
| 138 | 49 | 321 | 394 | 80.9% | chr17 | - | 42176888 | 42177462 | 575 |

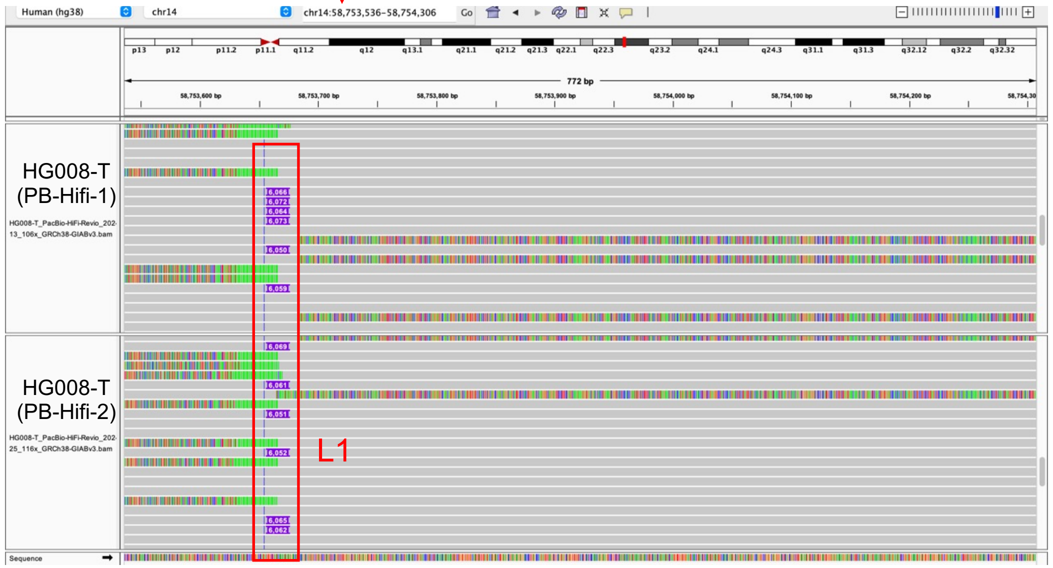


- Illustration of this L1

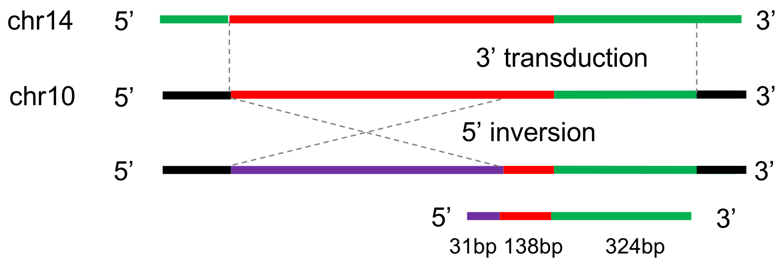

### chr10:84002440

4bp TSD; ACA TAG

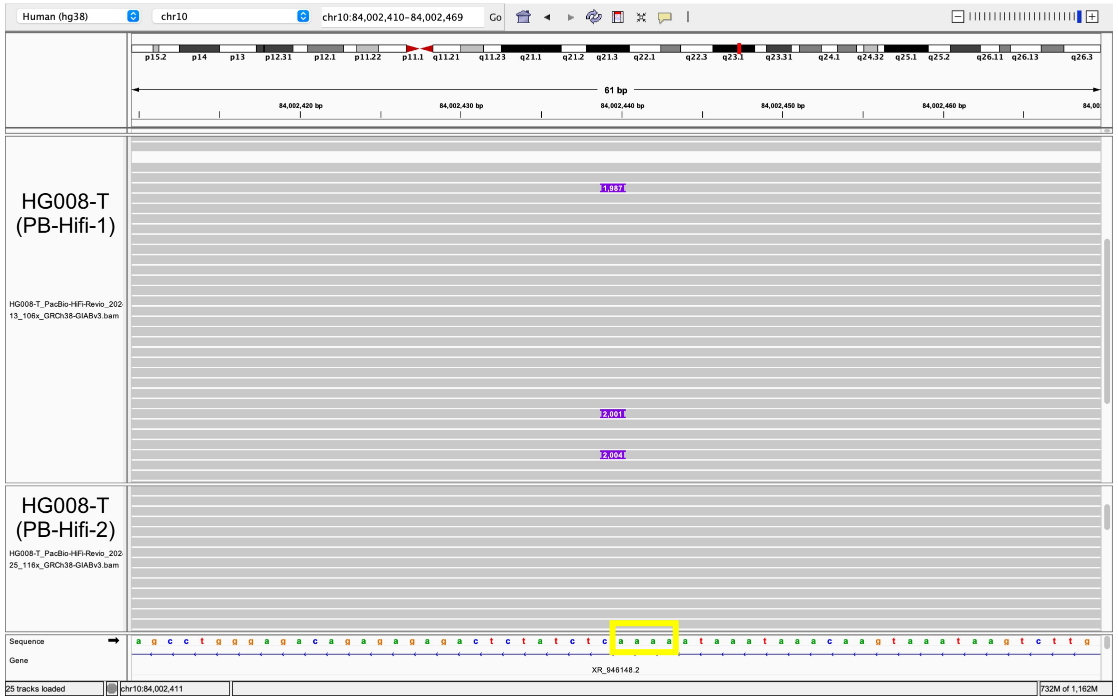


- **4bp TSD**
- Insertion:5-1978 aligns to L1Hs:4091-6064, **99% identity**

**Range 1: 4091 to 6064** Graphics ▼ Next Match ▲ Pre

| Score | Expect | Identities | Gaps | Strand |
|---|---|---|---|---|
| 3568 bits(1932) | 0.0 | 1960/1974(99%) | 0/1974(0%) | Plus/Plus |

- L1 hallmark alleles **ACA, TAG**

```
Query  9608  gggATAGCATTGGGAGATATACCTAATGCTAGATGACACATTAGTGGGTGCAGCGCACCA  9667
Sbjct  5890  GGGATAGCATTGGGAGATATACCTAATGCTAGATGACACGTTAGTGGGTGCAGCGCACCA  5949
Query  9668  GCATGGCACATGTATACATATGTAACTAACCTGCACAATGTGCACATGTACCCTAAAACT  9727
Sbjct  5950  GCATGGCACATGTATACATATGTAACTAACCTGCACAATGTGCACATGTACCCTAAAACT  6009
Query  9728  TAGAGTATAATaaaaaaaaaaaaaaaaaaaaaaaaaaaaaaaaaaaaaaaaaaaa  9782
Sbjct  6010  TAGAGTATAATAAAAAAAAAAAAAAAAAAAAAAAAAAAAAAAAAATAAATAAAAA  6064
```

Insertion sequence (2001 bp):

5’-AAAAGACGACATGATTGTTTATCTAGAAAACCCCATCGTCTCAGCCCAAAATCTCCTTAAGCTGATAAGCAACTTCAGCAAAGTCTCAGGATACAAAATCAATGTACAAAAATCACAAGCATTCTTATACACCAACAACAGACAAACA
GAGAGCCAAATCATGGGTGAACTCCCATTCACAATTGCTTCAAAGAGAATAAAATACCTAGGAATCCAACTTAGAAGGGATGTGAAGGACCTCTTCAAGGAGAACTACAAACCACTGCTCAAGGAAATAAAAGAGGAGACAAACAAATGGA
AGAACATTCCATGCTCATGGGTAGGAAGAATCAATATCGTGAAAATGGCCATACTGCCCAAGGTAATTTACAGATTCAATGCCATCCCCATCAAGCTACCAATGACTTTCTTCACAGAATTGGAAAAAACTACTTTAAAGTTCATATGGAA
CCAAAAAAGAGCCCGCATTGCCAAGTCAATCCTAAGCCAAAAGAACAAAGCTGGAGGCATCACACTACCTGACTTCAAACTATACTACAAGGCTACAGTAACCAAAACAGCATGGTACTGGTACCAAAACAGAGATATAGATCAATGGAAC
AGAACAGAGCCCTCAGAAATAACGCCGCATATCTACAACTATCTGATCTTTGACAAACCTGAGAAAAACAAGCAATGGGGAAAGGATTCCCTATTTAATAAATGGTGCTGGGAAAACTGGCTAGCCATATGTAGAAAGCTGAAACTGGATC
CCTTCCTTACACCTTATACAAAAATCAATTCAAGATGGATTAAAGATTTAAACGTTAAACCTAAAACCATAAAAACCCTAGAAGAAAACCTAGGCATTACCATTCAGGACATAGGTGTGGGCAAGGACTTCATGTCCAAAACACCAAAAGC
AATGGCAACAAAAGACAAAATTGACAAATGGGATCTAATTAAACTAAAGAGCTTCTGCACAGCAAAAGAAACTACCATCAGAGTGAACAGGCAACCTACAACATGGGAGAAAATTTTCGCAACCTACTCATCTGACAAAGGGCTAATATCC
AGAATCTACAATGAACTCAAACAAATTTACAAGAAAAAAACAAACAACCCCATCAAAAAGTGGGCGAAGGACATGAACAGACACTTCTCAAAAGAAGACATTTATGCAGCCAAAAAACACATGAAGAAATGCTCATCATCACTGGCCATCA
GAGAAATGCAAATCAAAACCACTATGAGATATCATCTCACACCAGTTAGAATGGCAATCATTAAAAAGTCAGGAAACAACAGGTGCTGGAGAGGATGCGGAGAAATAGGAACACTTTTACACTGTTGGTGGGACTGTAAACTAGTTCAACC
ATTGTGGAAGTCAGTGTGGCGATTCCTCAGGGATCTAGAACTAGAAATACCATTTGACCCAGCCATCCCATTACTGGGTATATACCCAAATGAGTATAAATCATGCTGCTATAAAGACACATGCACACGTATGTTTATTGCGGCACTATTC
ACAATAGCAAAGACTTGGAACCAACCCAAATGTCCAACAATGATAGACTGGATTAAGAAAATGTGGCACATATACACCATGGAATACTATGCAGCCATAAAAAATGATGAGTTCATATCCTTTGTAGGGACATGGATGAAATTGGAAACCA
TCATTCTCAGTAAACTATCGCAAGAACAAAAAACCAAACACCGCATATTCTCACTCATAGGTGGGAATTGAACAATGAGATCACATGGACACAGGAAGGGGAATATCACACTCTGGGGACTGTGGTGGGGTGGGGGGAGGGGGGAGGGATA
GCATTGGGAGATATACCTAATGCTAGATGACACATTAGTGGGTGCAGCGCACCAGCATGGCACATGTATACATATGTAACTAACCTGCACAATGTGCACATGTACCCTAAAACTTAGAGTATAATAAAAAAAAAAAAAAAAAAAAAAAAAA
AAAAAAAAAAAAAAAAAAAAAAAAAAAAAAAAAAAAAAAAA-3’

### chr12:7380777

no TSD;  ACA TAG

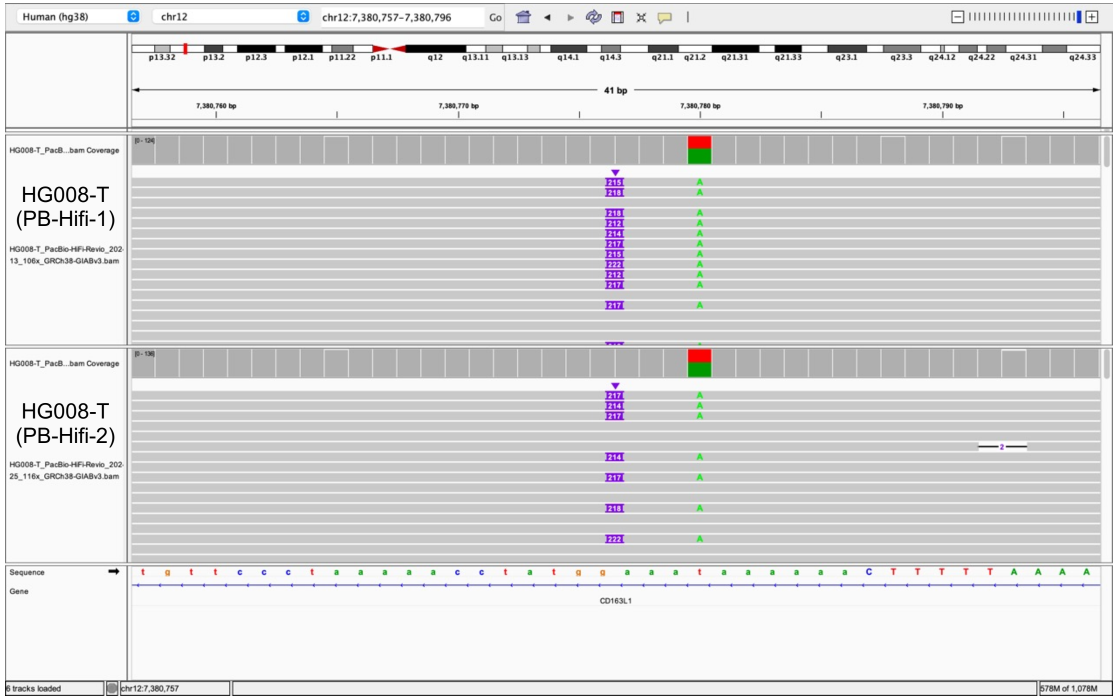


Insertion sequence (215 bp):
5’-GGACTGTGGTGGGGTAGGGGGAGGGGGGAGGGATAGCATTGGGAGATATACCTAATGCTAGATGACACA
TTAGTGGGTGCAGCGCACCAGGCATGGCACATGTATACATATGTAACTAACCTGCACAATGTGCACATGTACC
CTAAAACTTAGAGTATAATAAAAAAAAAAAAAAGAAAGAAAAAGAAAAAAAAAAAAAAAAAAAAAAAAAAAAAAA
-3’

- Insertion:1-208 aligns to L1Hs:5861-6064, **96% identity**

**Range 1: 5861 to 6064** Graphics ▼ Next Match ▲

| Score | Expect | Identities | Gaps | Strand |
|---|---|---|---|---|
| 339 bits(176) | 1e-96 | 197/205(96%) | 1/205(0%) | Plus/Plus |

- L1 hallmark alleles **ACA, TAG**

```
Query  61    GATGACACATTAGTGGGTGCAGCGCACCAGGCATGGCACATGTATACATATGTAACTAAC  120
             |||||||| |||||||||||||||||||| ||||||||||||||||||||||||||||||
Sbjct  5921  GATGACACGTTAGTGGGTGCAGCGCACCA-GCATGGCACATGTATACATATGTAACTAAC  5979

Query  121   CTGCACAATGTGCACATGTACCCTAAAACTTAGAGTATAATaaaaaaaaaaaaagaaaga  180
             |||||||||||||||||||||||||||||||||||||||||||||||||||||| ||| |
Sbjct  5980  CTGCACAATGTGCACATGTACCCTAAAACTTAGAGTATAATAAAAAAAAAAAAA-AAA-A  6037
```

### chr20:54619797

inversion; 18bp TSD; ACA TAG

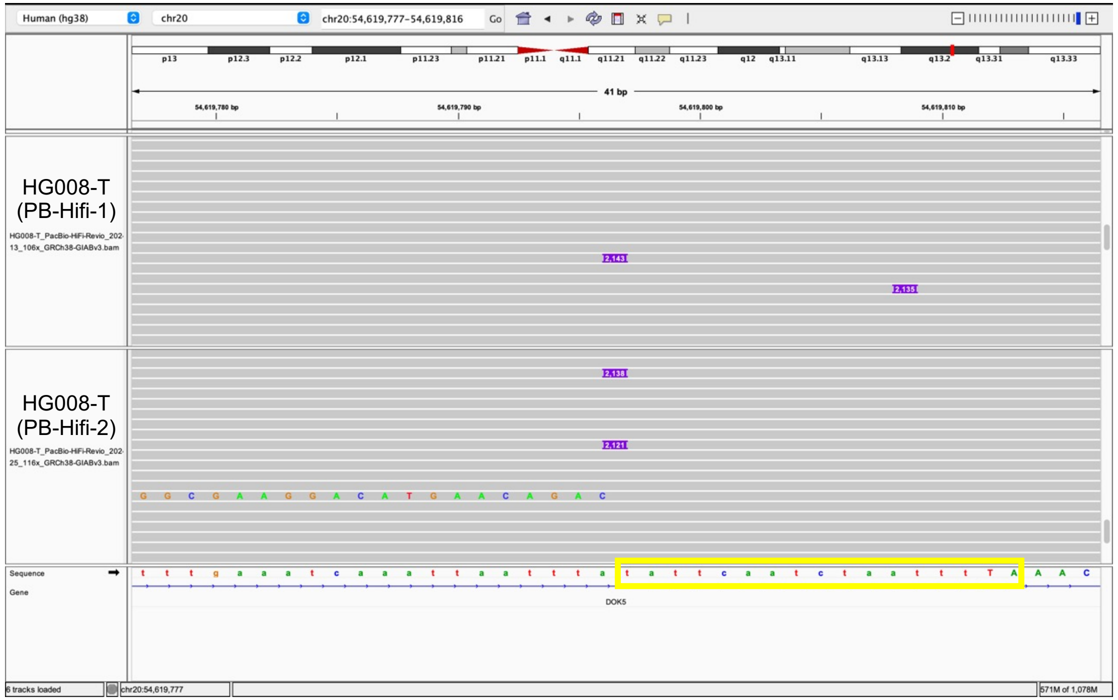


```
Insertion sequence (2124 bp):
3'-TATTCAATCTAATTTTATTTTTTTTTTTTTTTTTTTTTTTTTTTTTTTTTTTATTTATTTATTTATTTATTTT
TTTCCTTTCTTTTTTTTTTTTTTTATTATACTCTAAGTTTTAGGGTACATGTGCACATTGTGCAGGTTAGTTA
CATATGTATACATGTGCCATGCTGGTGCACTGCACCCACTAATGTGTCATCTAGCATTAGGTATATCTCCCAA
TGCTATCCCTCCCCCCTCCCCCCGACCCCACCACAGTCCCCAGAGTGTGATATTCCCCTTCCTGTGTCCATGT
GATCTCATTGTTCAATTCCCACCTATGAGTGAGAATATGCGGTGTTTGGTTTTTTGTTCTTGCGATAGTTTAC
TGAGAATGATGGTTTCCAATTTCATCCATGTCCCTACAAAGGATATGAACTCATCATTTTTTATGGCTGCATA
GTATTCCATGGTGTATATGTGCCACATTTTCTTAATCCAGTCTATCATTGTTGGACATTTGGGTTGGTTCCAA
GTCTTTGCTATTGTGAAATAGTGCCGCAATAAACATACGTGTGCATGTGTCTTTATAGCAGCATGATTTATAG
TCCTTTGGGTATATACCCAGTAATGGGATGGCTGGGTCAAATGGTATTTCTAGTTCTAGATCCTGAGGAATCG
CCACACTGACTTCCACAATGGTTGAACTAGTTTACAGTCCCACCAACAGTGTAAAAGTGTTCCTATTTCTGCA
CATCCTCTCCAGCACCTGTTGTTTCCTGACTTTTTAATGATTGCCATTCTAACTGGTGTGAGATGATATCTCA
TAGTGGTTTTGATTTGCATTTCTCTGATGGCCAGTGATGATGAGCATTTCTTCATGTGTTTTTTGGCTGCATA
AATGGAAGTTCTGGCCAGGGCAATCAGGCAGGAGAAGGAAATAAAGGGTATTCAATTAGGAAAAGAGGAAGTC
AAATTGTCCCT-700bp-CCTACAACATGGGAGAAAATTTCGCAACCTACTCATCTGACAAAGGGCTAATATC
CAGAATCTACAATGAACTCAAACAAATTTACAAGAAAAAAACAAACAACCCCATCAAAAAGTGGGCGAAGGAC
ATGAACAGAC-5'
```

➢ **18bp TSD**

➢ **5' inversion**: Two fragments of the sequence alignes to L1Hs:

(1) Insertion:81-910 aligned to L1Hs:6064-5239, **98% identity**; L1 hallmark alleles **ACA** (TGT reserve complement), **TAG** (CTA reserve complement)

**Range 2: 5239 to 6064** Graphics ▼ Next Match ▲ Previous Match ▲ First

| Score | Expect | Identities | Gaps | Strand |
|---|---|---|---|---|
| 1452 bits(786) | 0.0 | 819/833(98%) | 10/833(1%) | Plus/Minus |

```
Query  81    tttttatttatttatttttttttttttcctttcttttttttttttttttATTATACTCTAA  140
             ||||||||||||| ||| |||||||||  ||| |||||||||||||||||||||||||||
Sbjct  6064  TTTTTATTTATTT-TTT-TTTTTTTTT--TTTTTTTTTTTTTTTTTTTTTATTATACTCTAA  6009

Query  141   GTTTTAGGGTACATGTGCACATTGTGCAGGTTAGTTACATATGTATACATGTGCCATGCT  200
             ||||||||||||||||||||||||||||||||||||||||||||||||||||||||||||
Sbjct  6008  GTTTTAGGGTACATGTGCACATTGTGCAGGTTAGTTACATATGTATACATGTGCCATGCT  5949

Query  201   GGTGCACTGCACCCACTAATGTGTCATCTAGCATTAGGTATATCTCCCAATGCTATccct  260
             ||||| ||||||||||||| ||||||||||||||||||||||||||||||||||||||||
Sbjct  5948  GGTGCGCTGCACCCACTAACGTGTCATCTAGCATTAGGTATATCTCCCAATGCTATCCCT  5889
```

(2) Insertion:909-2124 aligns to L1Hs:4002-5222, **99% identity**

**Range 1: 4002 to 5222** Graphics ▼ Next Match

| Score | Expect | Identities | Gaps | Strand |
|---|---|---|---|---|
| 2200 bits(1191) | 0.0 | 1214/1223(99%) | 9/1223(0%) | Plus/Plus |

- Illustration of this L1

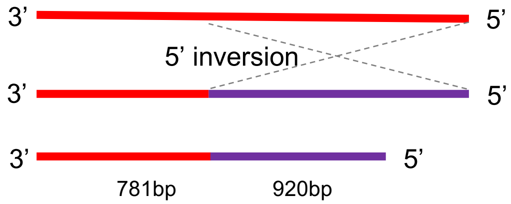

### chrX:69807092

#### 5bp TSD; ACA TAG

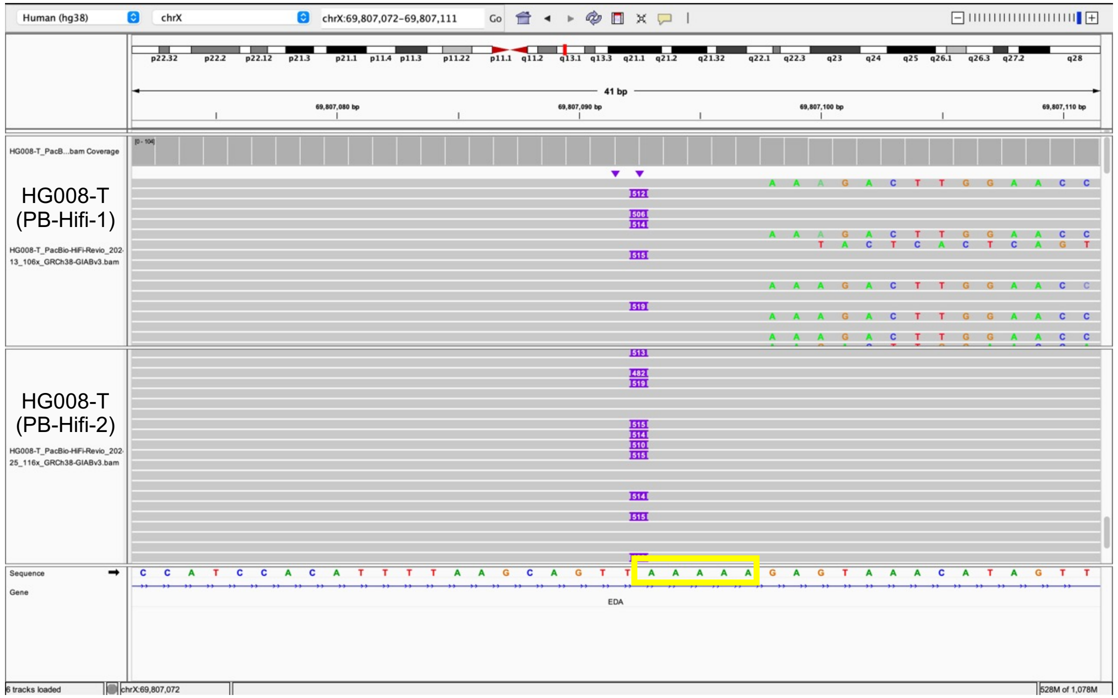


Insertion sequence (512 bp):
5'-AAAAAAAAGACTTGGAACCAACCCAAATGTCCAACAATGATAGACTGGATTAAGAAAATGTGGCACATAT
ACACCATGGAATACTATGCAGCCATAAAAAATGATGAGTTCATATCCTTTGTAGGGACATGGATGAAATTGGA
AACCATCATTCTCAGTAAACTATCGCAAGAACAAAAAACCAAACACCGCATATTCTCACTCATAGGTGGGAAT
TGAACAATGAGATCACATGGACACAGGAAGGGGAATATCACACTCTGGGGACTGTGGTGGGGTCGGGGGAGGG
GGGAGGGATAGCATTGGGAGATATACCTAATGCTAGATGACACATTAGTGGGTGCAGCGCACCAGCATGGCAC
ATGTATACATATGTAACTAACCTGCACAATGTGCACATGTACCCTAAAACTTAGAGTATAATAAAAAAAAAAA
AAAAAAAAAAAAAAAAAAAAAAAACAAAAATAAAAAAAAAAAAAAAAAAAAAAAAAAAAAAAAAAAAAAAAAA
AAAA-3'

- **5bp TSD**
- Insertion:6-470 aligns to L1Hs:5602-6058, **99% identity**

**Range 1: 5602 to 6058** Graphics ▼ Next Match ▲

| Score | Expect | Identities | Gaps | Strand |
|---|---|---|---|---|
| 867 bits(451) | 0.0 | 455/457(99%) | 0/457(0%) | Plus/Plus |

- L1 hallmark alleles **ACA, TAG**

```
Query  306   GGAGATATACCTAATGCTAGATGACACATTAGTGGGTGCAGCGCACCAGCATGGCACATG  365
             ||||||||||||||||||||||||||| ||||||||||||||||||||||||||||||||
Sbjct  5902  GGAGATATACCTAATGCTAGATGACACGTTAGTGGGTGCAGCGCACCAGCATGGCACATG  5961

Query  366   TATACATATGTAACTAACCTGCACAATGTGCACATGTACCCTAAAACTTAGAGTATAATa  425
             ||||||||||||||||||||||||||||||||||||||||||||||||||||||||||||
Sbjct  5962  TATACATATGTAACTAACCTGCACAATGTGCACATGTACCCTAAAACTTAGAGTATAATA  6021
```

### chrX:95987541

3'transduction; no TSD; ACA TAG

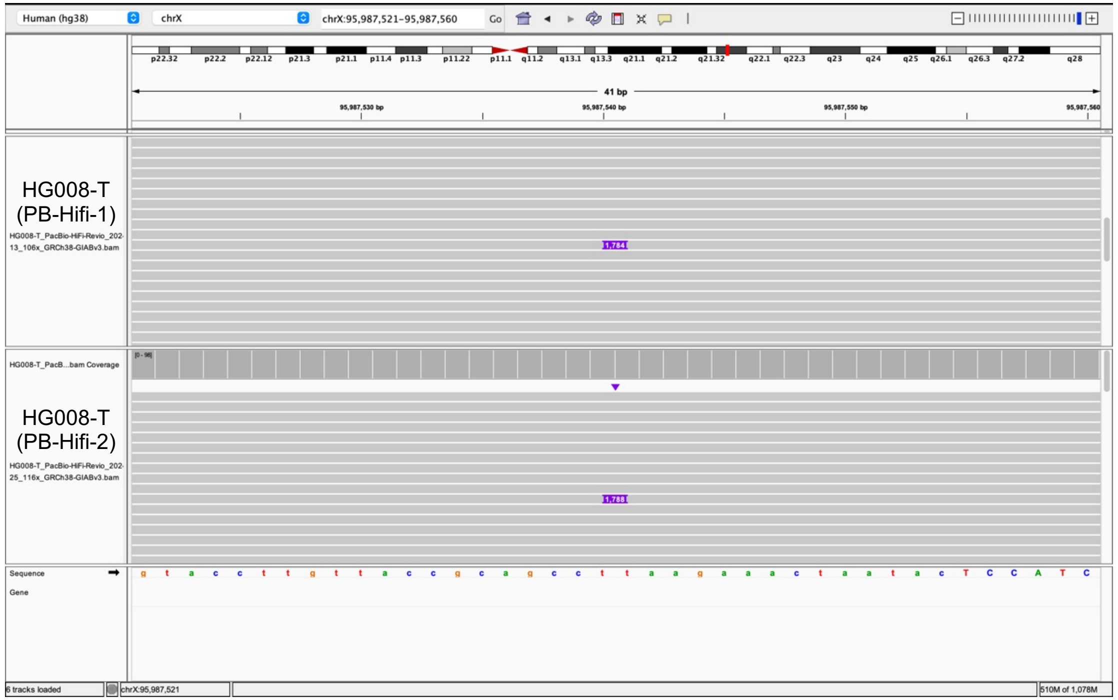


Insertion sequence (1788 bp):

5'-AGGCATTACCATTCAGGACATAGGCGTGGGCAAGGACTTCATGTCCAAAACACCAAAAGCAATGGCAACA
AAAGACAAAATTGACAAATGGGATCTAATTAAACTAAAGAGCTTCTGCACAGCAAAAGAAACTACCATCAGAG
TGAACAGGCAACCTACAACATGGGAGAAAATTTTCGCAACCTACTCATCTGACAAAGGGCTAATATCCAGAAT
CTA-700bp-TCACACTCTGGGGACTGTGGTGGGGTAGGGGGAGGGGGGAGGGATAGCATTGGGAGATATACC
TAATGCTAGATGACACATTAGTGGGTGCAGCGCACCAGCATGGCACATGTATACATATGTAACTAACCTGCAC
AATGTGCACATGTACCCTAAAACTTAGAGTATAATAAAAAAAAAAAAAGAAAGAAAAAGAAACTTCACATGGG
GCCACATGTCTAATAAAGGTGACTGAATCCCAAAATTATATTGGTAACATGGAAATTCCTGATAGGGGAAGAA
TATTCCCAGAAGTGTGACCAGTTAGGGCCAAAATAAAAGGTACAATTTACCCAATTATTATACCTAAGTACCC
GGCCTTATTAAATATATGTTCCTCAATATCAAATTGTACTAAAAATCAGGAAACTAAGCTCCTAGTAGAGAAC
TGGTCAAAGGGAAGATCTTTGTGGAGTCAGCAGTTTCCTCTGTCCTACAATTTGGCAAAATAATCTCATAAGC
ACAAGAAGGTTAATATGCTTGCATTAAAAATGTTAAGTTACTTATAATCCAATTTAAATGTAAGTTCAAATAC
CTTAACATGTTCACATTACTTTTTTAATTCACAAAAACGTTCTTATCTATCTTTATTTTAAGATGTTTTCCAC
ATACATGCTTAGTGAATTTTCTTTTAGGAGATGTAAGAGAGTCTACAAATCTTTATTCTACTATTTCCCAAAA
TATACCCAATTACTGGCAAAAAAAGTTATTATTAAACTGATGAGAAAATATAAAGGTTTATTACTGGATTAAA
AAAAAAAAAAAAAAAAAAAAAAAAAAAAAAAAAAAAAAAAAAAAAAAAAAAAAAAAAAAAAAAAAAAAAAAAA
AAA-3'

- Insertion:1-1117 aligns to L1Hs:4930-6047, **99% identity**

**Range 1: 4930 to 6047** Graphics ▼ Next Match

| Score | Expect | Identities | Gaps | Strand |
|---|---|---|---|---|
| 2008 bits(1087) | 0.0 | 1108/1118(99%) | 1/1118(0%) | Plus/Plus |

- **3'transduction:** 3' of the sequence aligns to chr22:28669303-28669924, adjacent to a L1Hs

| SCORE | START | END | QSIZE | IDENTITY | CHROM | STRAND | START | END | SPAN |
|---|---|---|---|---|---|---|---|---|---|
| 620 | 1 | 622 | 698 | 99.9% | chr22 | + | 28669303 | 28669924 | 622 |
| 40 | 1 | 414 | 698 | 97.7% | chr11 | - | 22493921 | 22494349 | 429 |
| 29 | 357 | 395 | 698 | 73.4% | chr1 | + | 185189536 | 185189565 | 30 |
| 27 | 570 | 603 | 698 | 79.4% | chr1 | + | 187937064 | 187937093 | 30 |
| 26 | 445 | 472 | 698 | 88.9% | chr1 | + | 189859236 | 189859262 | 27 |

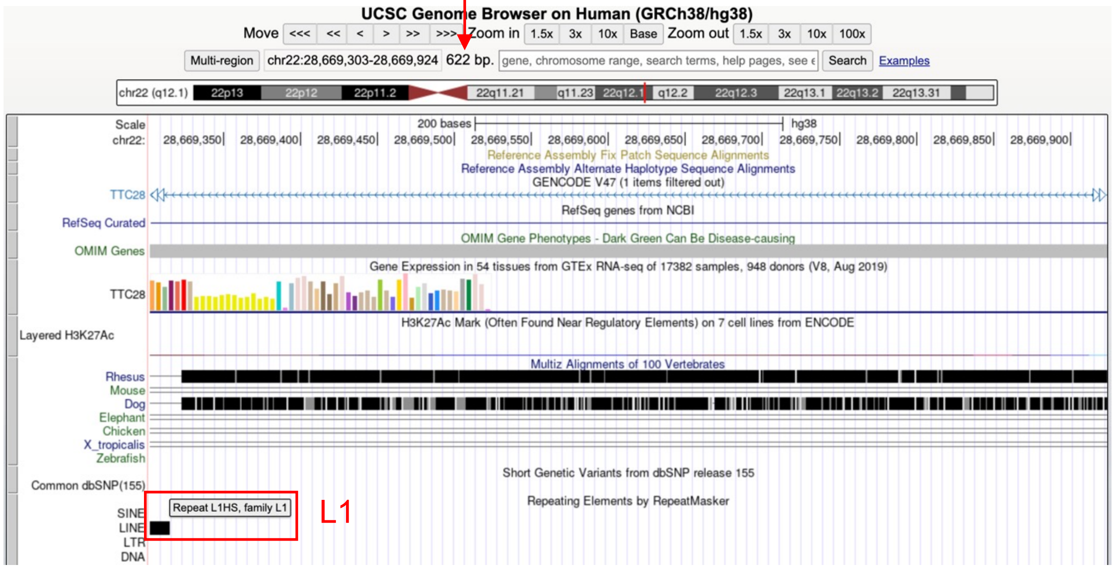


- Illustration of this L1

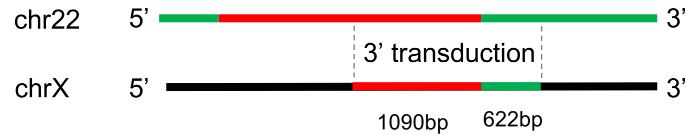

### chrX:143137165

3'transduction; no TSD; TAG

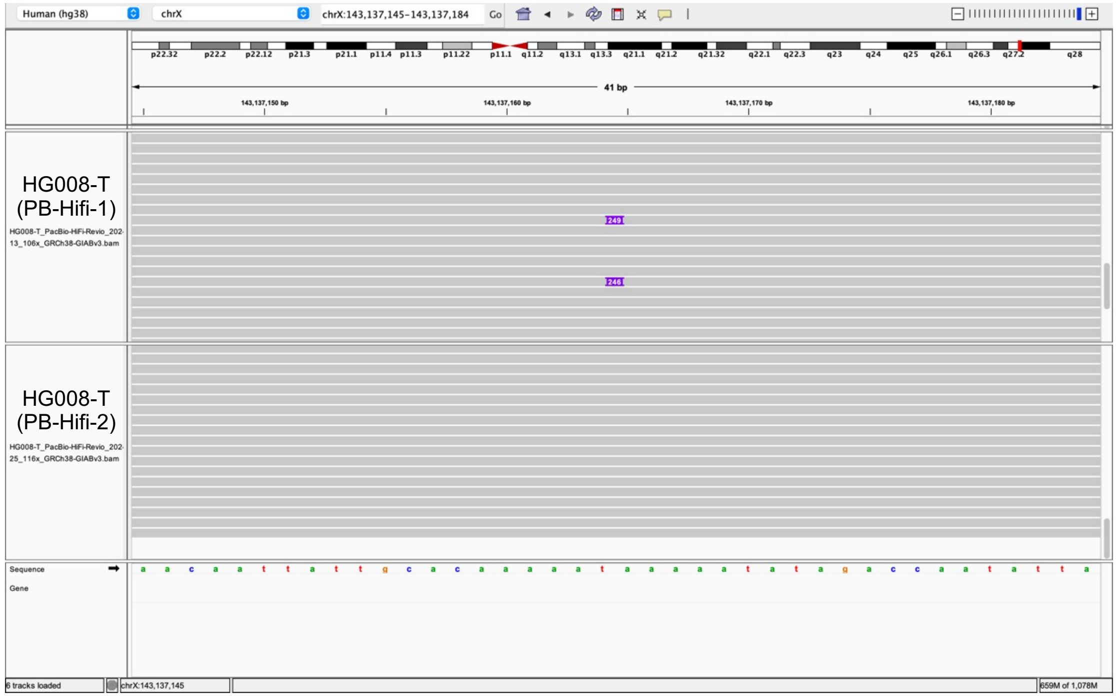


Insertion sequence (249 bp):
5’-GTGCACATGTACCCTAAAACTTAGAGTATAATAAAAAAAAAAAAAAAAAAAAAAAAAAAAAAAAAAGAA
CTGTGCAAAAAAAAAAAAAAAAAAAAAAAAACACTTAGCAATGATACAATGTGCTATCTGAAAAAACTAA
ACTTTTTAAAATATAAATTTGAGGGAAAATGAAAAAAAAAAAAAAAAAAAAAAAAAAAAAAAAAAAAAAAA
AAAAAAAAAAAAAAAAAAAAAAAAAAAAAAAA-3

- Insertion:1-66 aligns to L1Hs:5989-6054 **100% identity**; L1 hallmark alleles **TAG**

**Range 1: 5989 to 6054** Graphics ▼ Next Match ▲ Previous

| Score | Expect | Identities | Gaps | Strand |
|---|---|---|---|---|
| 122 bits(66) | 1e-31 | 66/66(100%) | 0/66(0%) | Plus/Plus |

- **3’transduction:** 3’ of the sequence aligns to chr9:5491407-5491484, adjacent to a full-length L1

| SCORE | START | END | QSIZE | IDENTITY | CHROM | STRAND | START | END | SPAN |
|---|---|---|---|---|---|---|---|---|---|
| 76 | 34 | 111 | 183 | 98.8% | chr9 | + | 5491407 | 5491484 | 78 |
| 36 | 7 | 48 | 183 | 97.4% | chrX | - | 42345341 | 42345383 | 43 |
| 26 | 1 | 26 | 183 | 100.0% | chr10 | - | 19947708 | 19947733 | 26 |
| 23 | 76 | 101 | 183 | 96.2% | chr2 | - | 58018491 | 58018518 | 28 |
| 22 | 26 | 47 | 183 | 100.0% | chr18 | - | 33573296 | 33573317 | 22 |

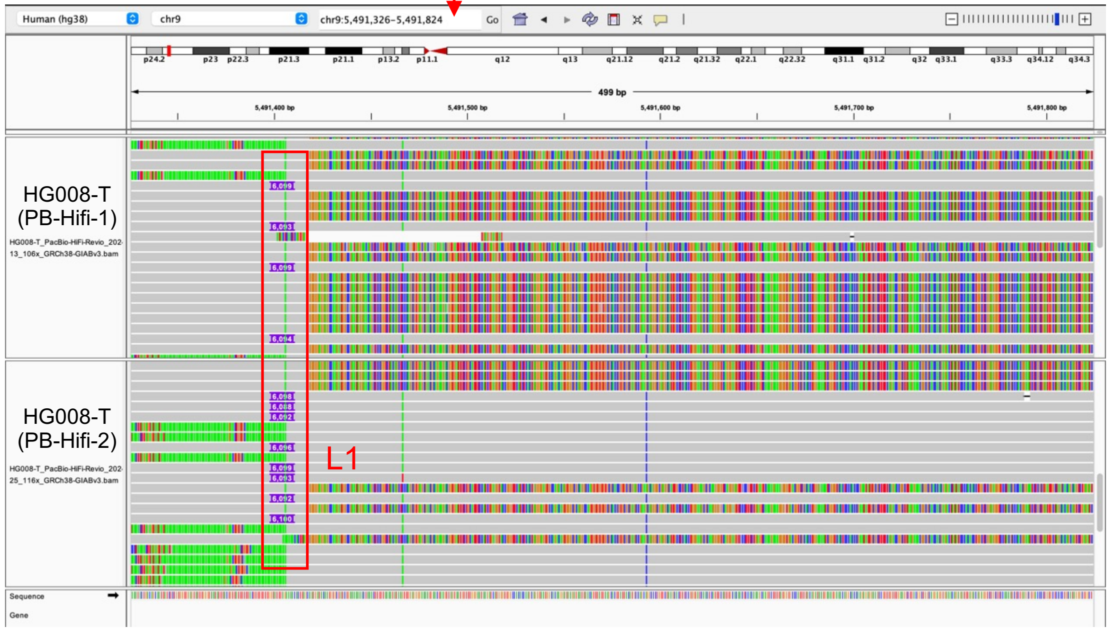


- Illustration of this L1

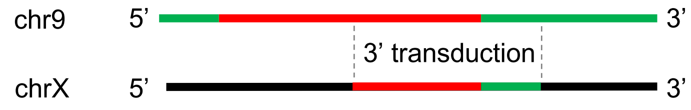
