## Extended Files for "Image-based DNA Sequencing Encoding for Detecting Low-Mosaicism Somatic Mobile Element Insertions": Extended Files 5.pdf

### Benchmark tumor somatic L1 Insertions in HG008-T and their detection by PALMER or xTea_long

(Additional explanation for Supplementary Table 10)

| Chr | Start | End | Supporting Reads | Estimated Mosaicism in PacBio | PALMER detected in PB-Hifi-1 | PALMER detected in PB-Hifi-2 | xTea_long detected in PB-Hifi-1 | xTea_long detected in PB-Hifi-2 | In benchmark list |
|---|---|---|---|---|---|---|---|---|---|
| chr4 | 14331582 | 14331584 | 125 | 100.00% | + | + | + | + | yes |
| chr4 | 112897566 | 112897582 | 89 | 100.00% | | | + | + | yes |
| chr4 | 134389657 | 134389677 | 105 | 100.00% | + | + | + | + | yes |
| chr12 | 7380776 | 7380777 | 134 | 100.00% | + | + | + | + | yes |
| chr8 | 104941340 | 104941341 | 107 | 93.45% | + | + | + | + | yes |
| chrX | 69807091 | 69807093 | 101 | 88.99% | + | + | + | + | yes |
| chr8 | 107947957 | 107947991 | 67 | 72.83% | + | + | + | + | yes |
| chr10 | 25515229 | 25515242 | 35 | 49.30% | + | + | + | + | yes |
| chr5 | 4450306 | 4450341 | 34 | 40.48% | + | | | | yes |
| chr9 | 104414095 | 104414109 | 8 | 12.21% | | | | | yes |
| chr8 | 62600957 | 62600958 | 11 | 9.87% | + | + | + | | yes |
| chr8 | 82189195 | 82189200 | 8 | 8.00% | + | | | | yes |
| chr20 | 54619796 | 54619809 | 10 | 6.25% | + | + | | | yes |
| chr10 | 84002439 | 84002440 | 3 | 4.55% | | | | | yes |
| chr2 | 95103766 | 95103767 | 2 | 2.53% | | | | | yes |
| chrX | 95987540 | 95987541 | 2 | 2.16% | | | | | yes |
| chrX | 143137164 | 143137165 | 2 | 2.16% | | | | | yes |
| chr3 | 175249151 | 175249152 | 2 | 1.79% | | | | | yes |
| chr3 | 137297551 | 137297552 | 2 | 1.68% | | | | | yes |
| chr19 | 27312746 | 27312747 | | | | | + | | no (found in normal) |
| chr20 | 29107410 | 29107411 | | | | | | + | no (found in normal) |
| chr20 | 30145602 | 30145603 | | | | | + | + | no (found in normal) |

**Note:**

A total of 19 benchmark tumor somatic L1 insertions were identified during benchmarking. However, xTea_long identified 3 L1 insertions that not in the benchmark list. Upon manual review using IGV, we confirmed that these 3 insertions were also present in the normal samples, indicating they are not tumor somatic events.

Bam files of the tumor and normal datasets were accessed through the following links:

- HG008-T (PB-Hifi-1): https://42basepairs.com/download/s3/giab/data_somatic/HG008/Liss_lab/BCM_Revio_20240313/HG008-T_PacBio-HiFi-Revio_20240313_106x_GRCh38-GIABv3.bam
- HG008-T (PB-Hifi-2): https://42basepairs.com/download/s3/giab/data_somatic/HG008/Liss_lab/PacBio_Revio_20240125/HG008-T_PacBio-HiFi-Revio_20240125_116x_GRCh38-GIABv3.bam
- HG008-N-D (PB-Hifi-1): https://42basepairs.com/download/s3/giab/data_somatic/HG008/Liss_lab/BCM_Revio_20240313/HG008-N-D_PacBio-HiFi-Revio_20240313_68x_GRCh38-GIABv3.bam
- HG008-N-P (PB-Hifi-2): https://42basepairs.com/download/s3/giab/data_somatic/HG008/Liss_lab/PacBio_Revio_20240125/HG008-N-P_PacBio-HiFi-Revio_20240125_35x_GRCh38-GIABv3.bam

### chr19:27312746

#### Tumor tissue

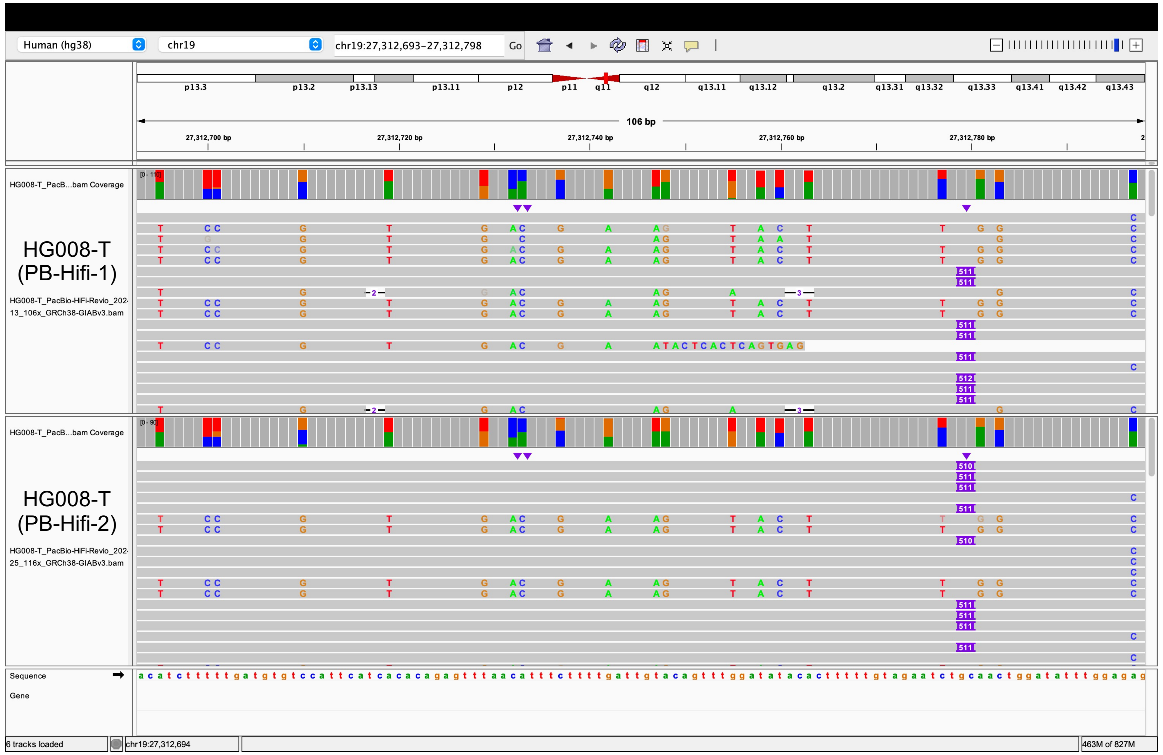


#### Same insertion in normal tissue

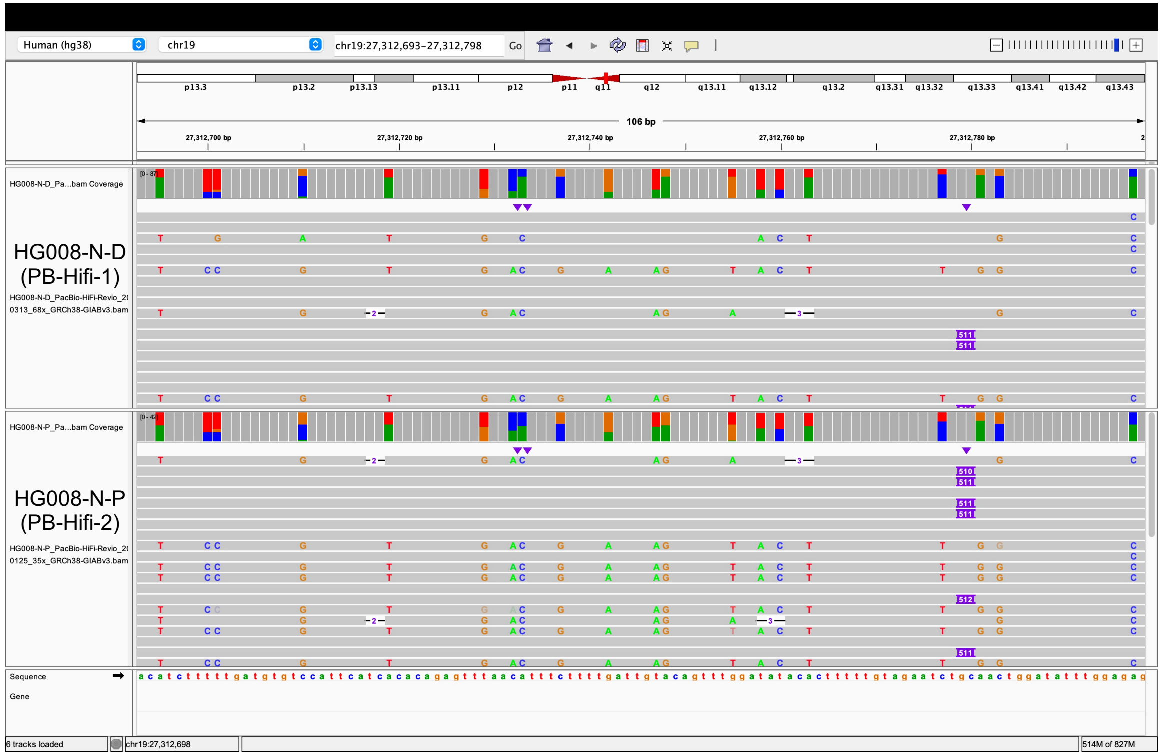

### chr20:29107410

Tumor tissue

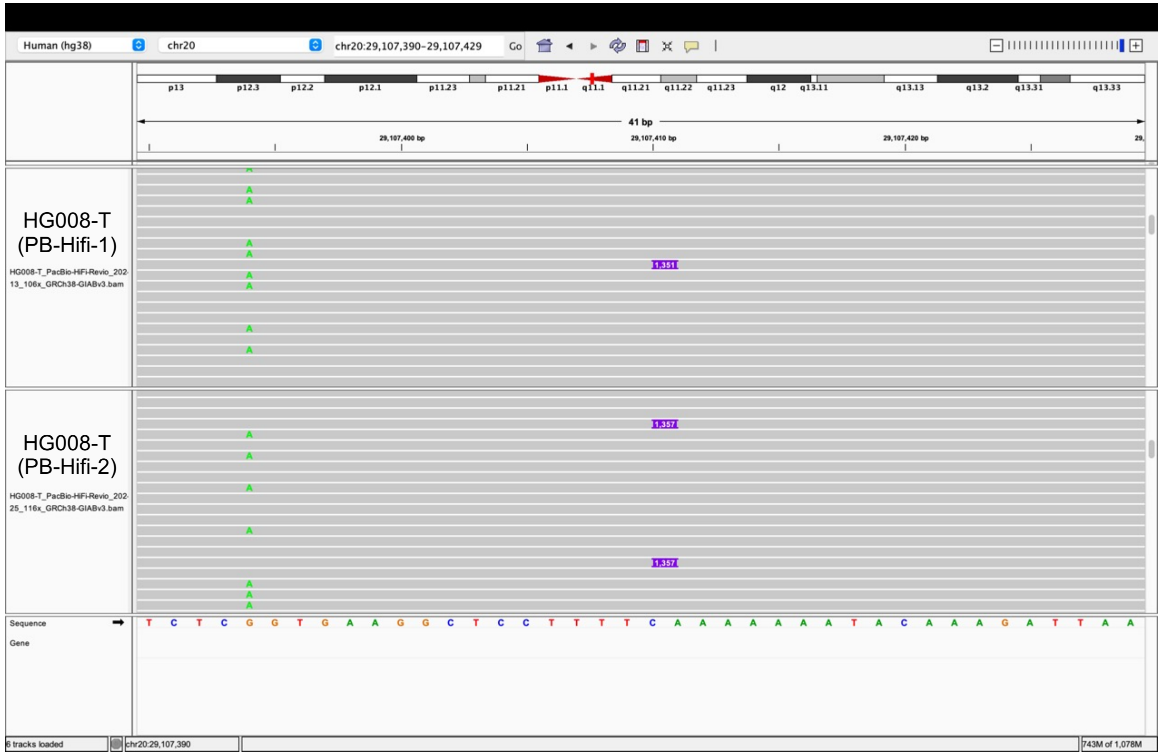


Same insertion in normal tissue

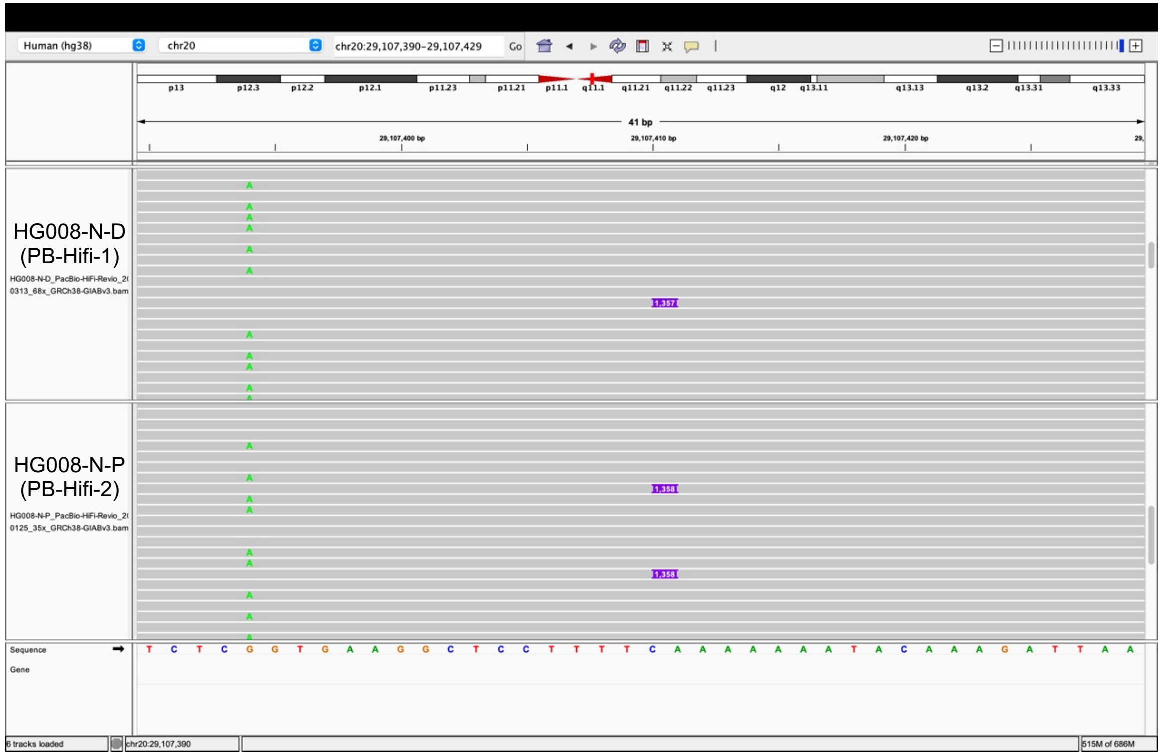

### chr20:30145602

#### Tumor tissue

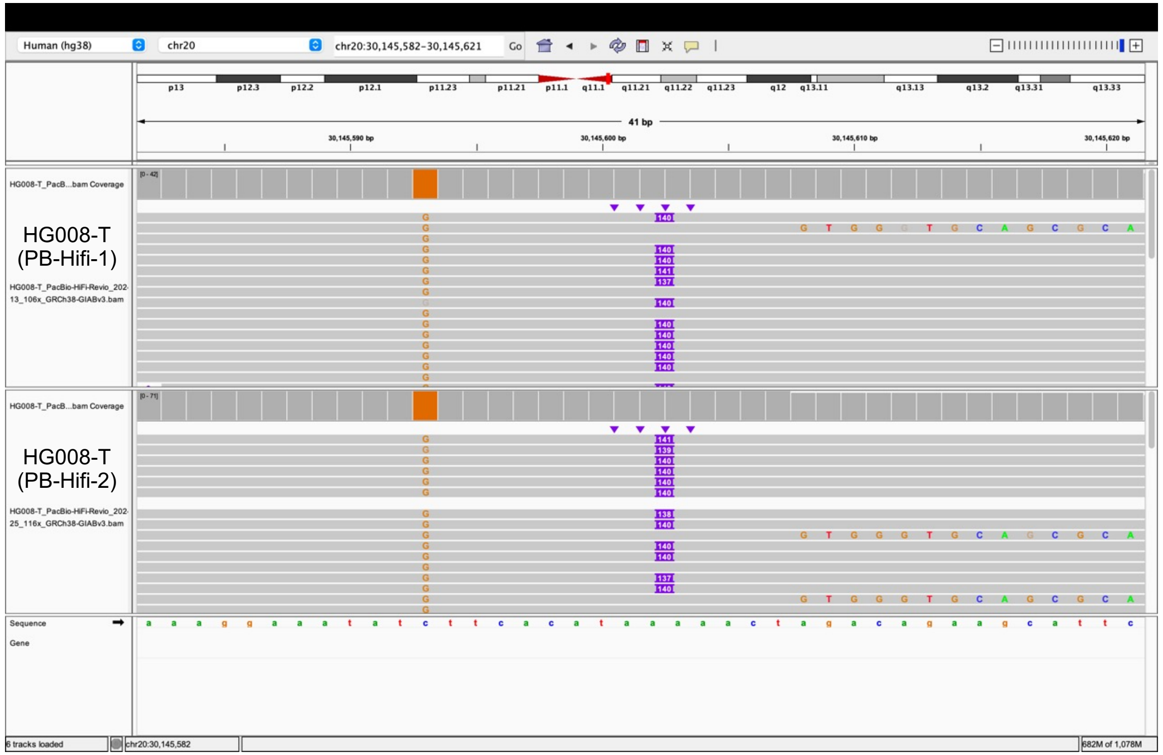


#### Same insertion in normal tissue

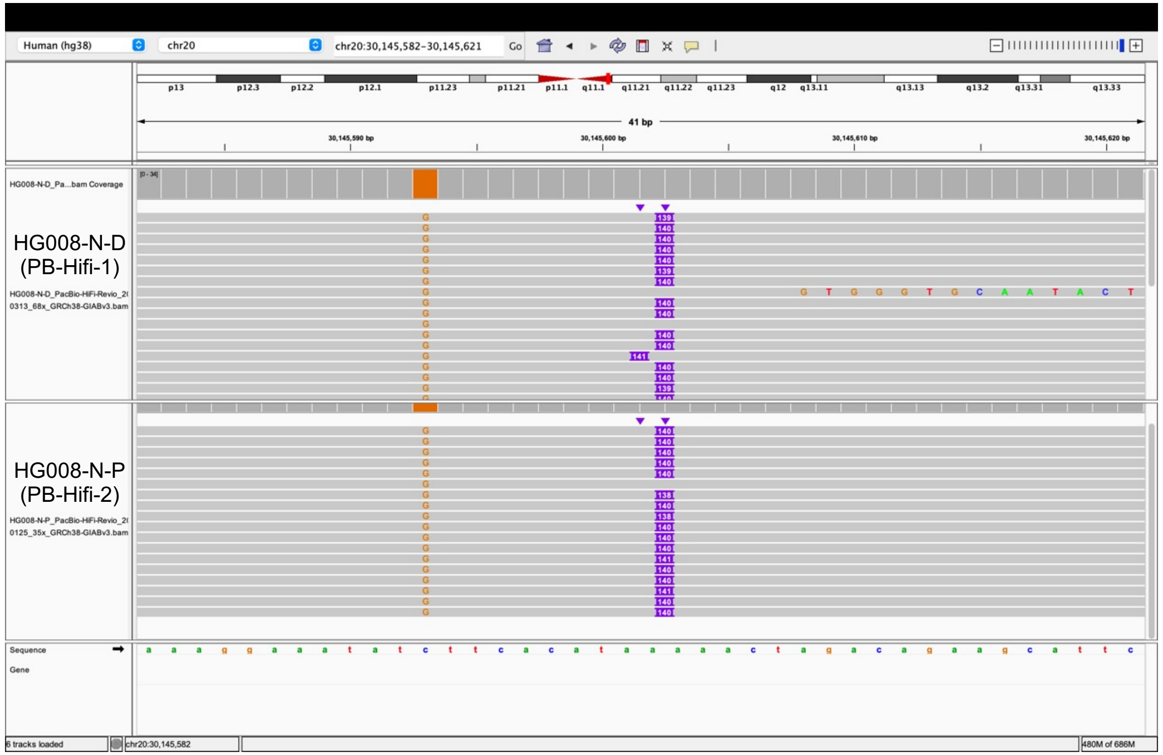
