## Extended Files for "Image-based DNA Sequencing Encoding for Detecting Low-Mosaicism Somatic Mobile Element Insertions": Extended Files 6.pdf

### Tumor somatic *Alu* insertions detected in HG008-T by PALMER or xTea_long

(Additional explanation for Supplementary Table 10)

| Chr | Start | End | PALMER detected in PB-Hifi-1 | PALMER detected in PB-Hifi-2 | xTea_long detected in PB-Hifi-1 | xTea_long detected in PB-Hifi-2 | In benchmark list |
|---|---|---|---|---|---|---|---|
| chr6 | 53132718 | 53132719 | + | | | | no (tandem duplication) |
| chr7 | 7802513 | 7802516 | + | + | | | no (tandem duplication) |
| chr16 | 4484205 | 4484206 | + | + | | | no (tandem duplication) |
| chr19 | 4856353 | 4856353 | | + | | | no (deletion) |
| chr19 | 4860844 | 4860844 | | + | | | no (deletion) |
| chr19 | 8346857 | 8346857 | | + | | | no (found in normal) |
| chr19 | 11217803 | 11217805 | + | + | | | no (inversion; translocation) |
| chrY | 10780622 | 10780622 | + | | | | no (found in normal) |
| chr1 | 748214 | 748215 | | | + | + | no (found in normal) |
| chr1 | 121655868 | 121655869 | | | + | | no (found in normal) |
| chr2 | 91441826 | 91441827 | | | + | + | no (found in normal) |
| chr2 | 242181270 | 242181271 | | | + | + | no (found in normal) |
| chr6 | 32587099 | 32587100 | | | + | + | no (found in normal) |
| chr10 | 28348988 | 28348989 | | | + | | no (found in normal) |
| chr15 | 22584273 | 22584274 | | | + | + | no (found in normal) |
| chr19 | 58605628 | 58605629 | | | | + | no (found in normal) |
| chr22 | 12595451 | 12595452 | | | + | + | no (found in normal) |
| chr22 | 16476005 | 16476006 | | | + | | no (found in normal) |
| chrX | 1067023 | 1067024 | | | | + | no (found in normal) |
| chrX | 321575 | 321576 | | | + | | no (found in normal) |
| chrX | 1334422 | 1334423 | | | + | | no (found in normal) |
| chrX | 1432859 | 1432860 | | | + | + | no (found in normal) |
| chrY | 11349045 | 11349046 | | | + | + | no (found in normal) |

**Note:**

No benchmark tumor somatic *Alu* insertions were identified during benchmarking. However, PALMER and xTea_long erroneously classified 8 and 15 *Alu* insertions, respectively. Upon manual review using IGV, we confirmed that these insertions were structural variants or present in the normal samples, not tumor somatic events.

Bam files of the tumor and normal datasets were accessed through the following links:

- HG008-T (PB-Hifi-1): https://42basepairs.com/download/s3/giab/data_somatic/HG008/Liss_lab/BCM_Revio_20240313/HG008-T_PacBio-HiFi-Revio_20240313_106x_GRCh38-GIABv3.bam
- HG008-T (PB-Hifi-2): https://42basepairs.com/download/s3/giab/data_somatic/HG008/Liss_lab/PacBio_Revio_20240125/HG008-T_PacBio-HiFi-Revio_20240125_116x_GRCh38-GIABv3.bam
- HG008-N-D (PB-Hifi-1): https://42basepairs.com/download/s3/giab/data_somatic/HG008/Liss_lab/BCM_Revio_20240313/HG008-N-D_PacBio-HiFi-Revio_20240313_68x_GRCh38-GIABv3.bam
- HG008-N-P (PB-Hifi-2): https://42basepairs.com/download/s3/giab/data_somatic/HG008/Liss_lab/PacBio_Revio_20240125/HG008-N-P_PacBio-HiFi-Revio_20240125_35x_GRCh38-GIABv3.bam

### chr6:53132718 (tandem duplication)

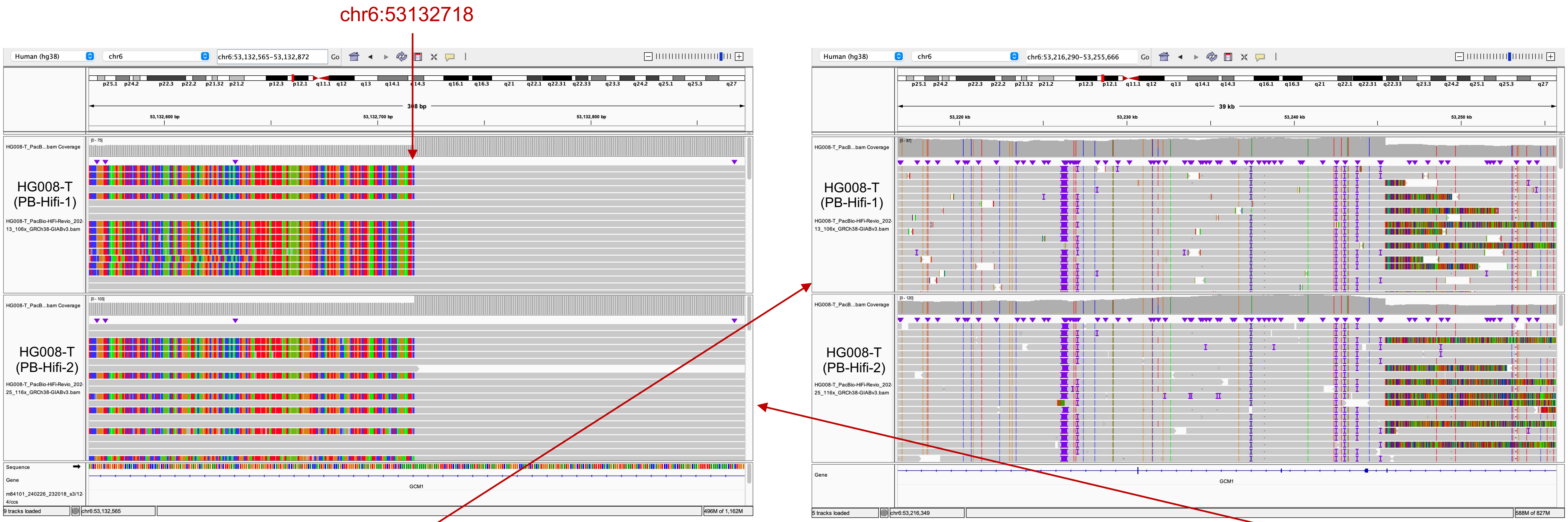


Left-clip aligns to chr6:53226499-53245458

| QUERY | SCORE | START | END | QSIZE | IDENTITY | CHROM | STRAND | START | END | SPAN |
|---|---|---|---|---|---|---|---|---|---|---|
| YourSeq | 18898 | 17 | 18984 | 18989 | 99.9% | chr6 | + | 53226499 | 53245458 | 18960 |

Right-clip aligns to chr6:53132718-53150188

| QUERY | SCORE | START | END | QSIZE | IDENTITY | CHROM | STRAND | START | END | SPAN |
|---|---|---|---|---|---|---|---|---|---|---|
| YourSeq | 17391 | 6 | 17462 | 17462 | 99.9% | chr6 | + | 53132718 | 53150188 | 17471 |

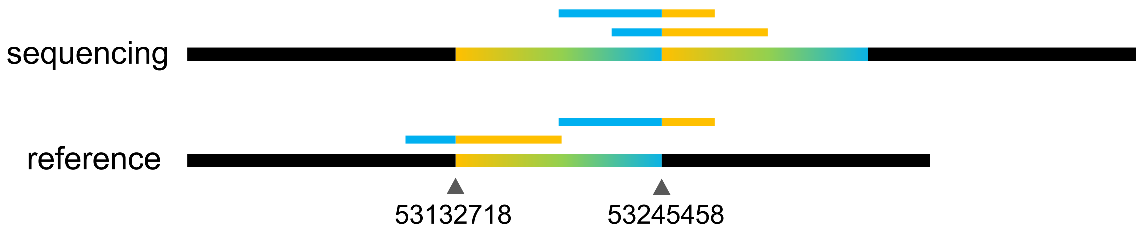

### chr7:7802513 (tandem duplication)

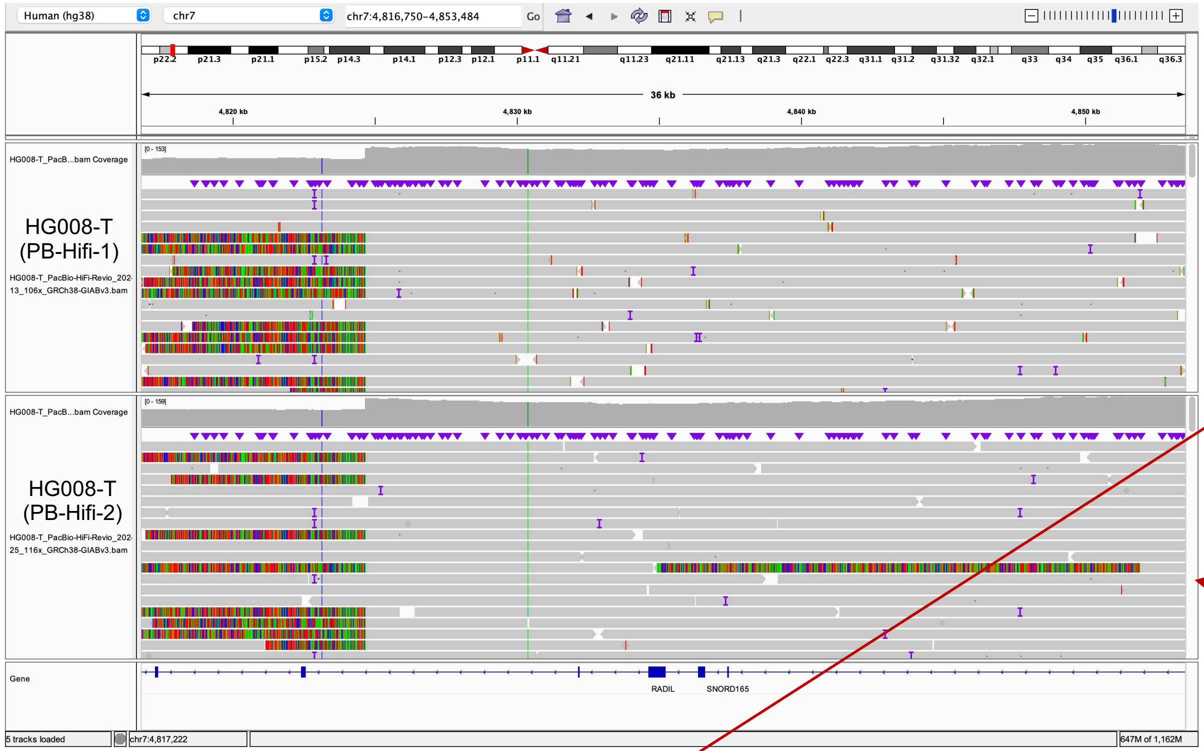


chr7:7802513

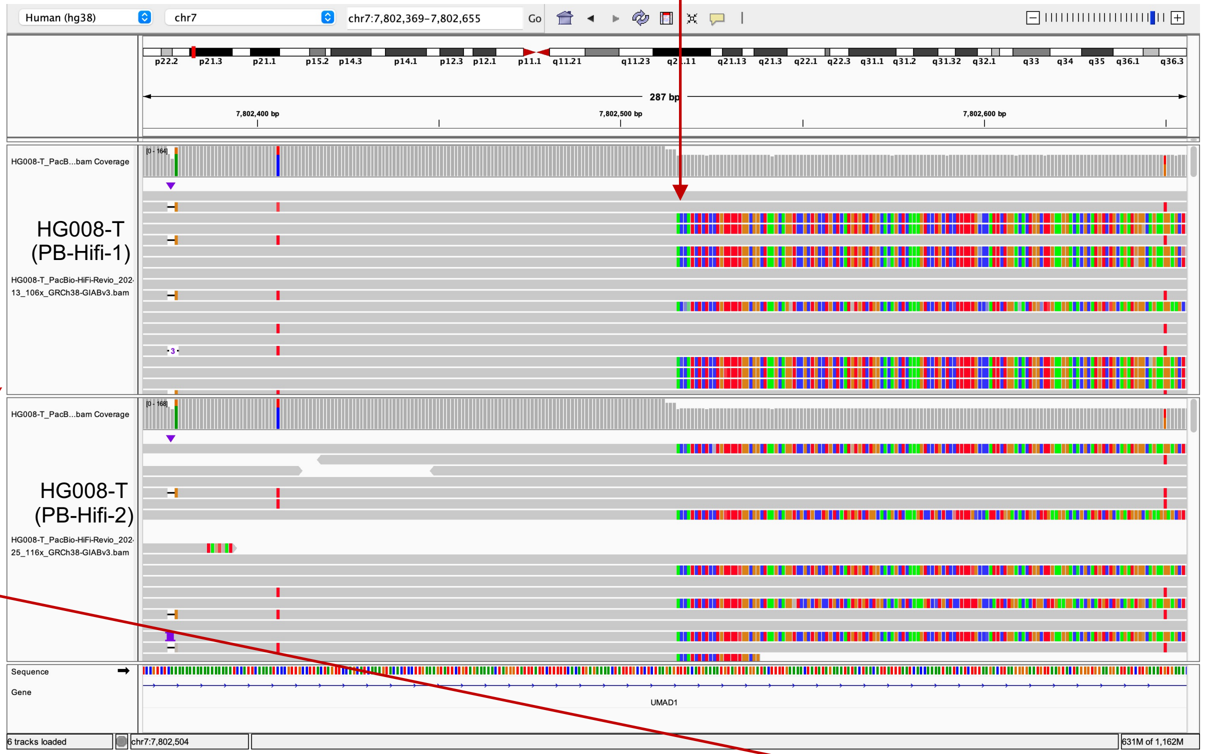


Left-clip aligns to chr17:7794271-7802512

| QUERY | SCORE | START | END | QSIZE | IDENTITY | CHROM | STRAND | START | END | SPAN |
|---|---|---|---|---|---|---|---|---|---|---|
| YourSeq | 8210 | 17 | 8256 | 8256 | 99.9% | chr7 | + | 7794271 | 7802512 | 8242 |

Right-clip aligns to chr7:4824649-4846562

| QUERY | SCORE | START | END | QSIZE | IDENTITY | CHROM | STRAND | START | END | SPAN |
|---|---|---|---|---|---|---|---|---|---|---|
| YourSeq | 11109 | 17 | 21920 | 21920 | 99.6% | chr7 | - | 4824649 | 4846562 | 21914 |

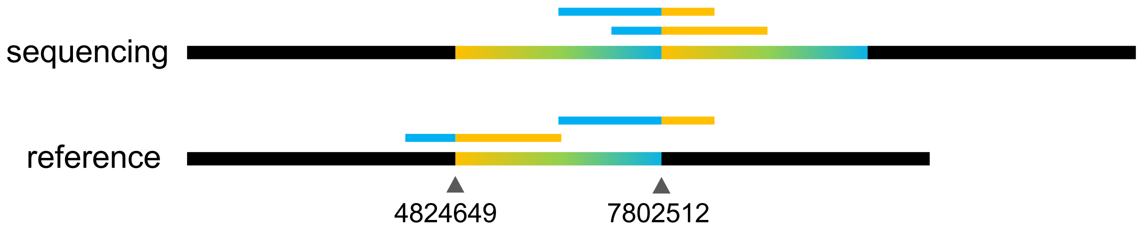

### chr16:4484205 (tandem duplication)

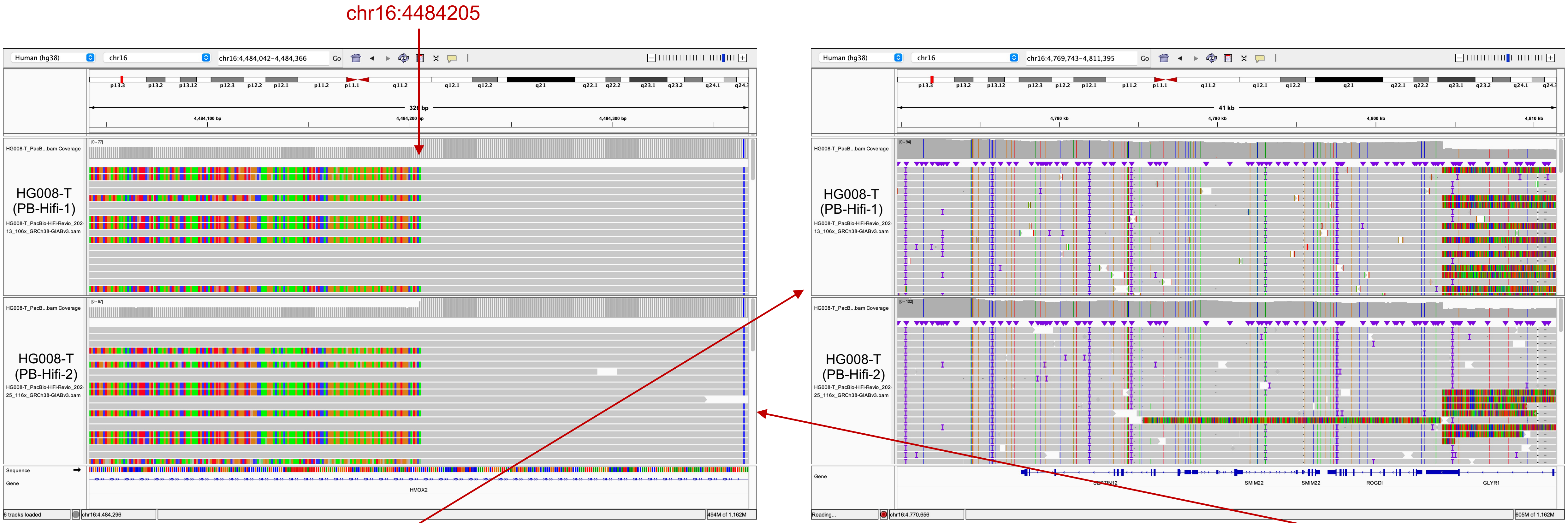


Left-clip aligns to chr6:4776904-4804234

| QUERY | SCORE | START | END | QSIZE | IDENTITY | CHROM | STRAND | START | END | SPAN |
|---|---|---|---|---|---|---|---|---|---|---|
| YourSeq | 21861 | 16 | 27469 | 27469 | 99.5% | chr16 | + | 4776904 | 4804234 | 27331 |

Right-clip aligns to chr7:4484205-4504718

| QUERY | SCORE | START | END | QSIZE | IDENTITY | CHROM | STRAND | START | END | SPAN |
|---|---|---|---|---|---|---|---|---|---|---|
| YourSeq | 13599 | 16 | 20521 | 20521 | 99.7% | chr16 | – | 4484205 | 4504718 | 20514 |

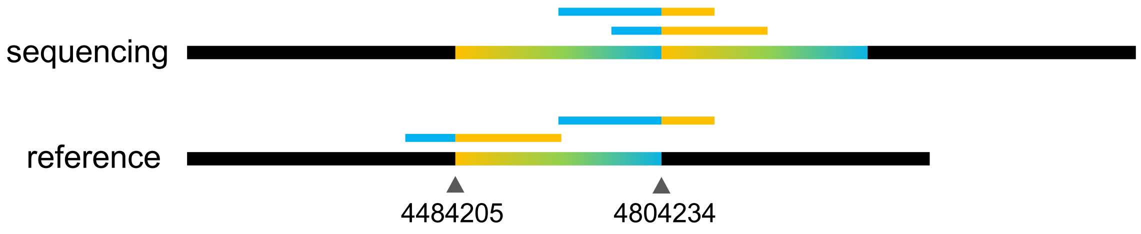

### chr19:4856353 (deletion)

### chr19:4860844 (deletion)

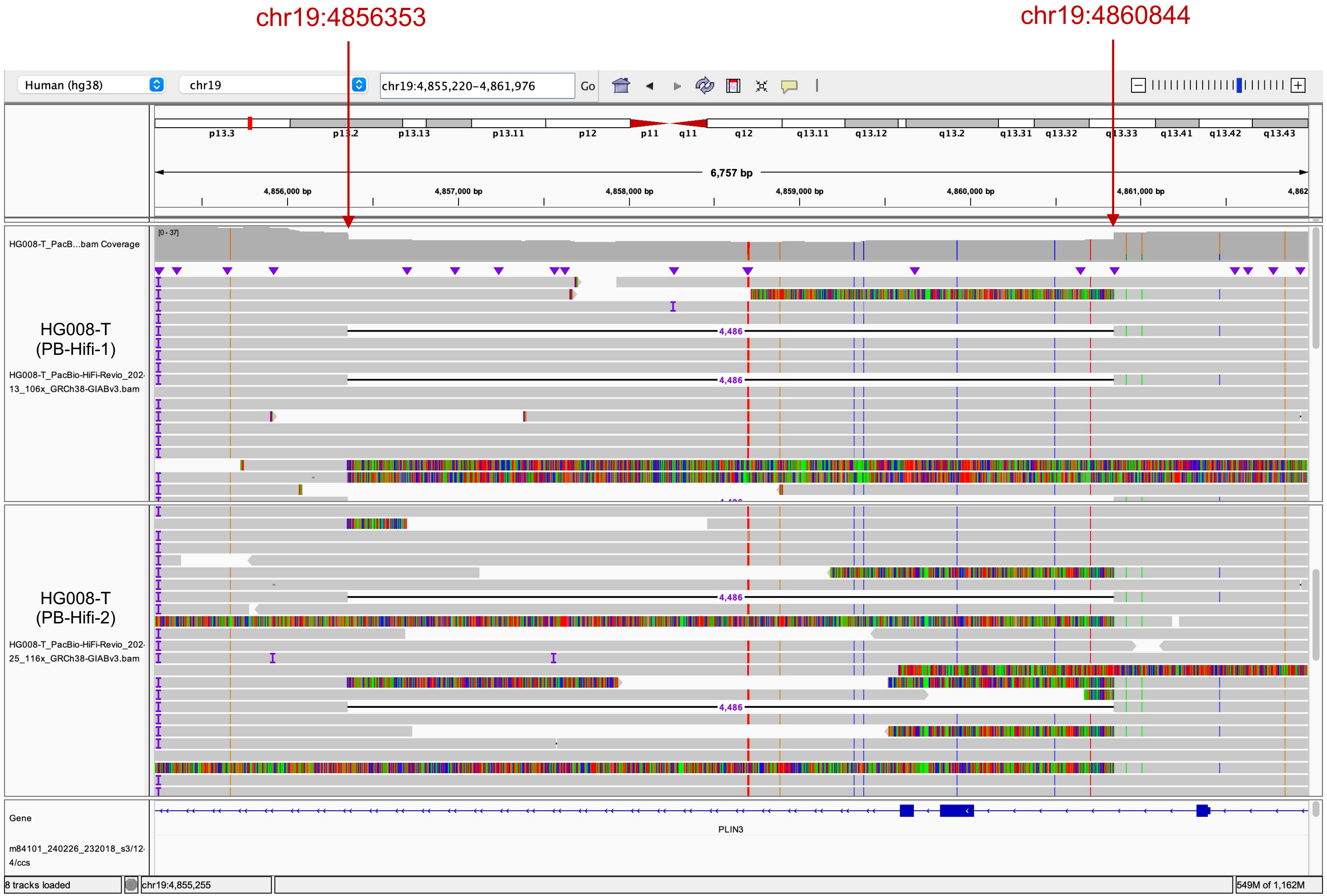

### chr19:8346857 (Found in normal)

#### Tumor tissue

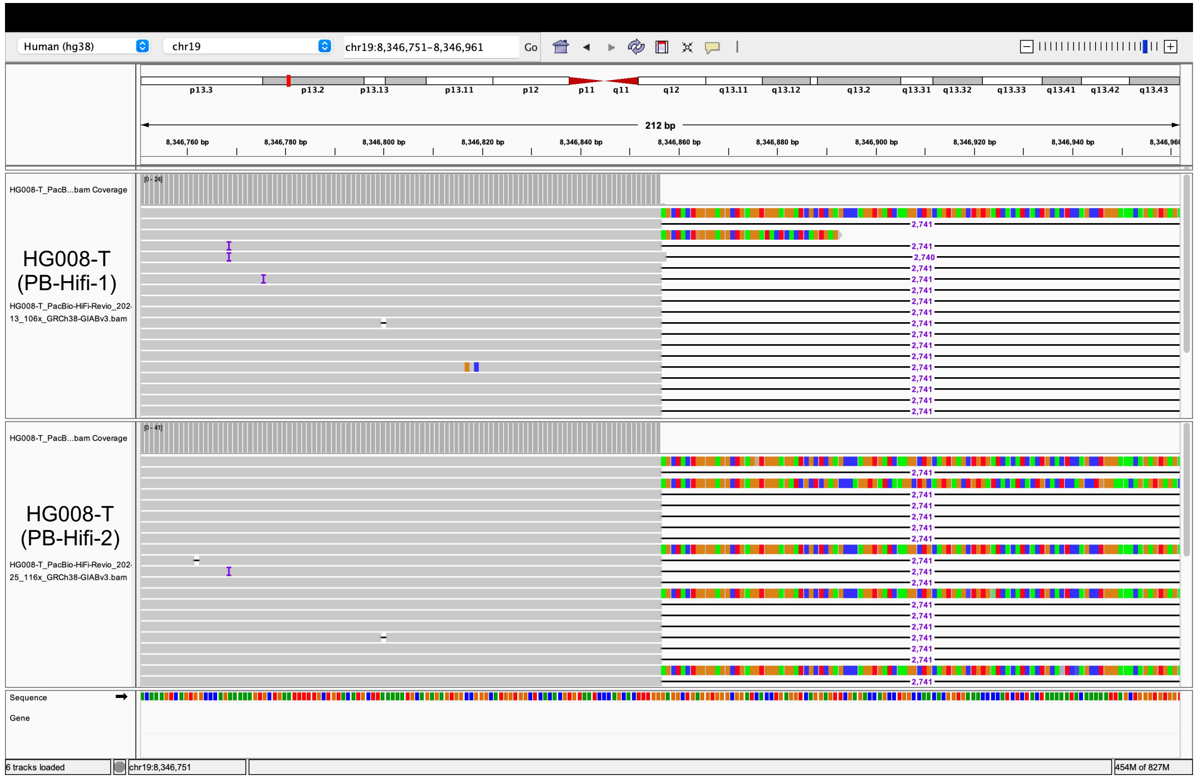


#### Same clipped reads in normal tissue

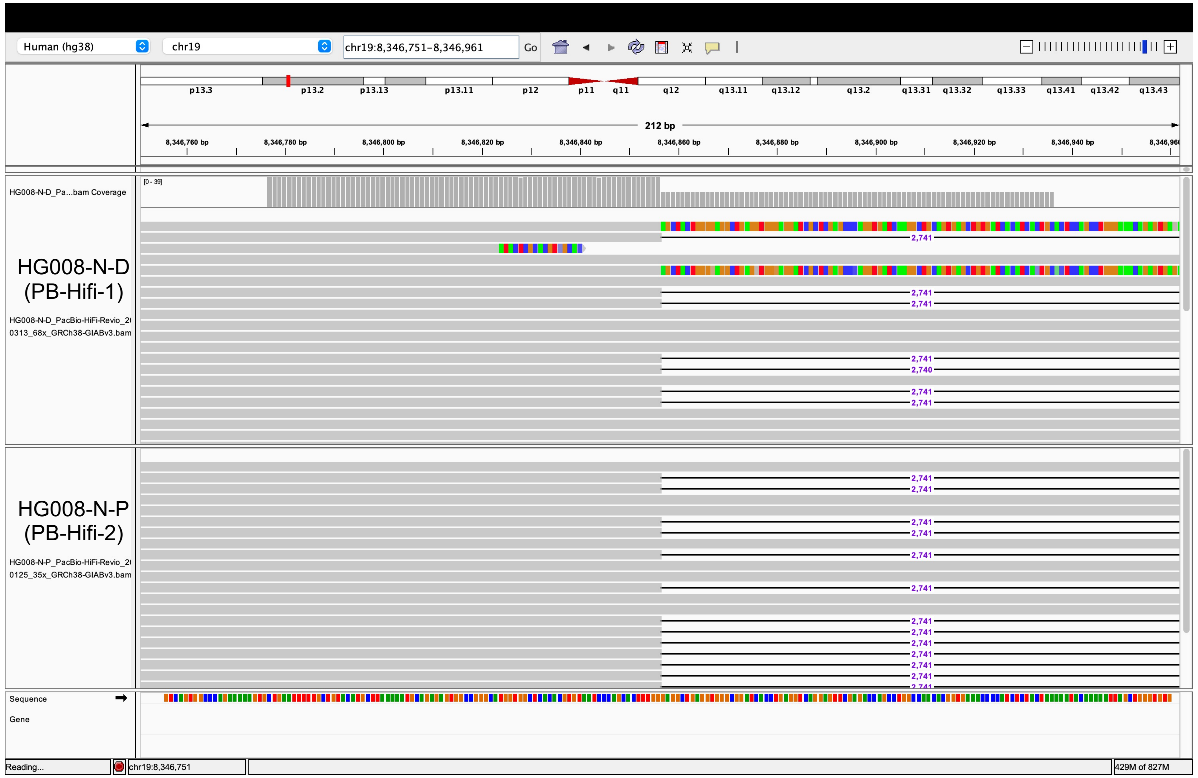

### chr19:11217803 (inversion; translocation)

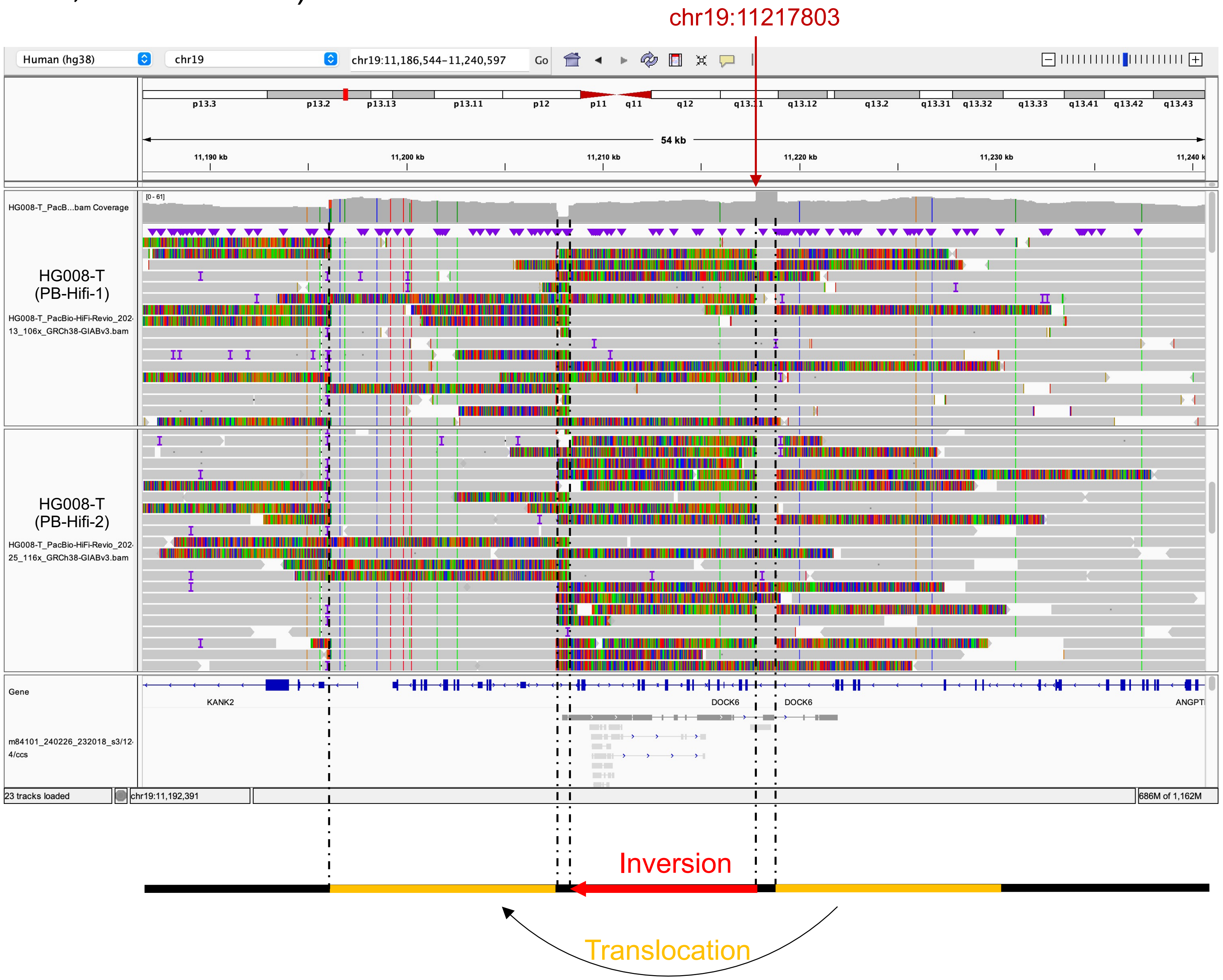

### chrY:10780622 (Found in normal)

#### Tumor tissue

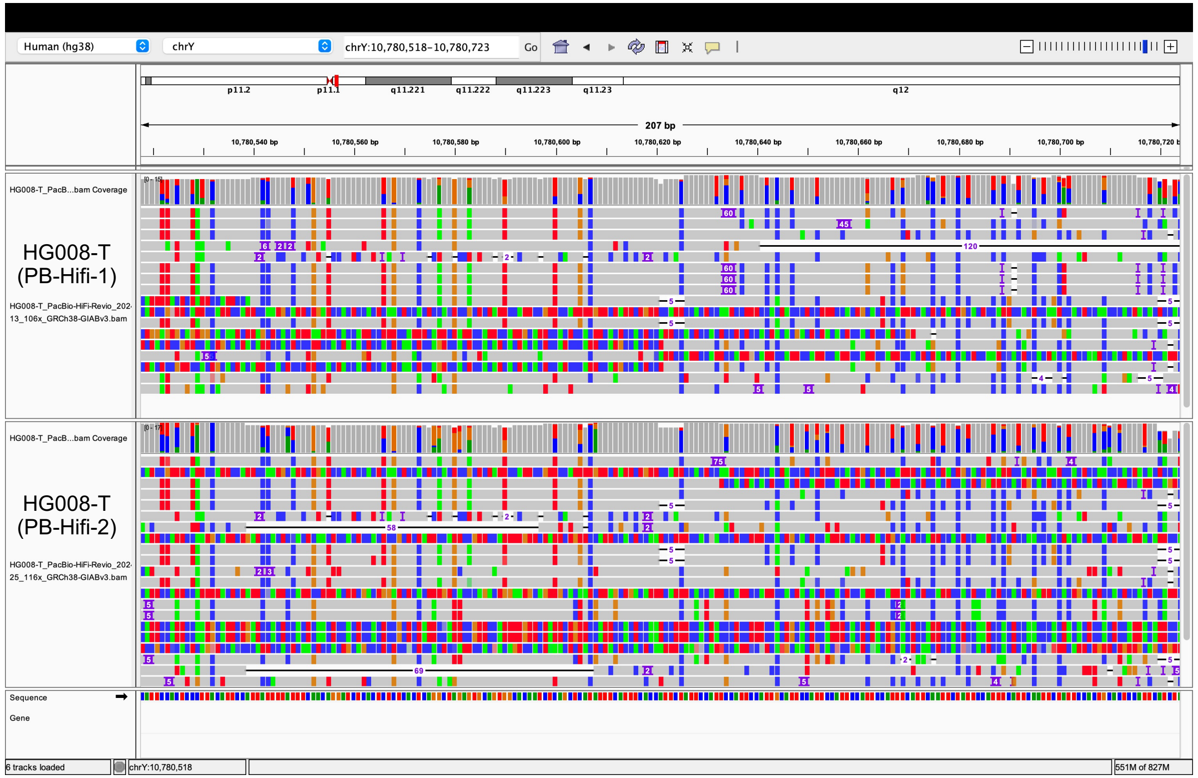


#### Same clipped reads in normal tissue

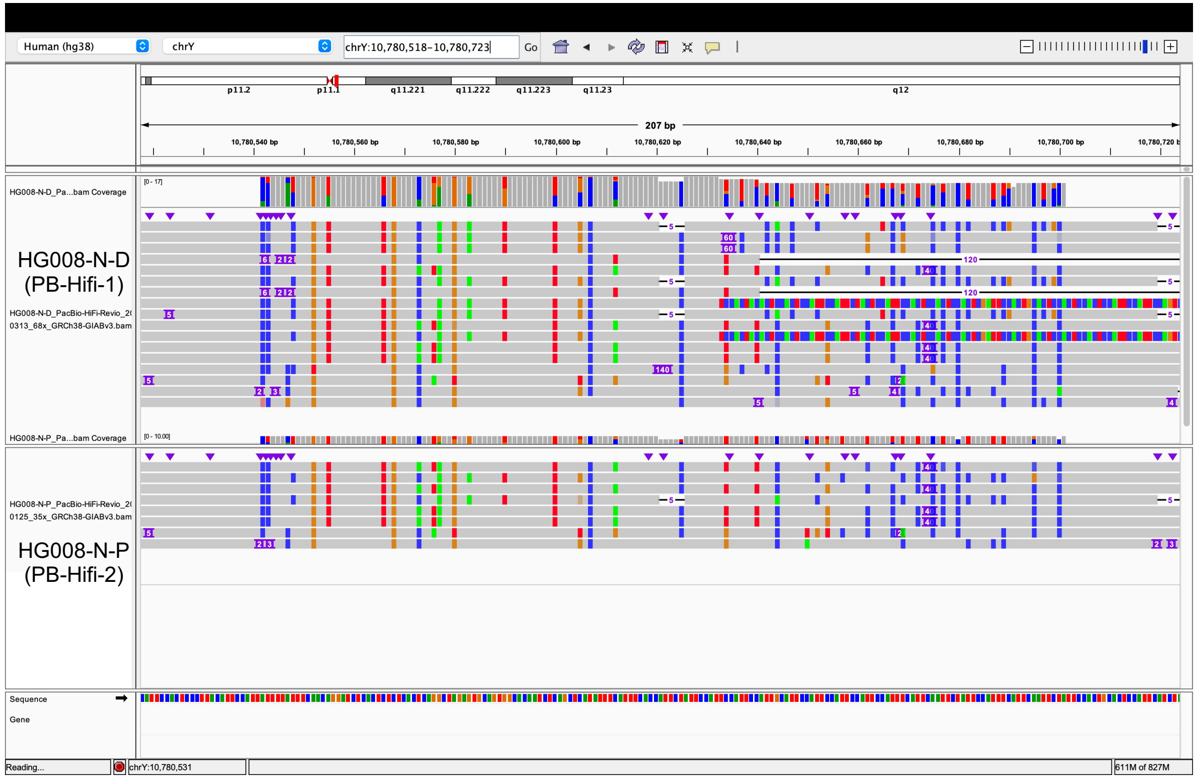

### chr1:748214 (Found in normal)

#### Tumor tissue

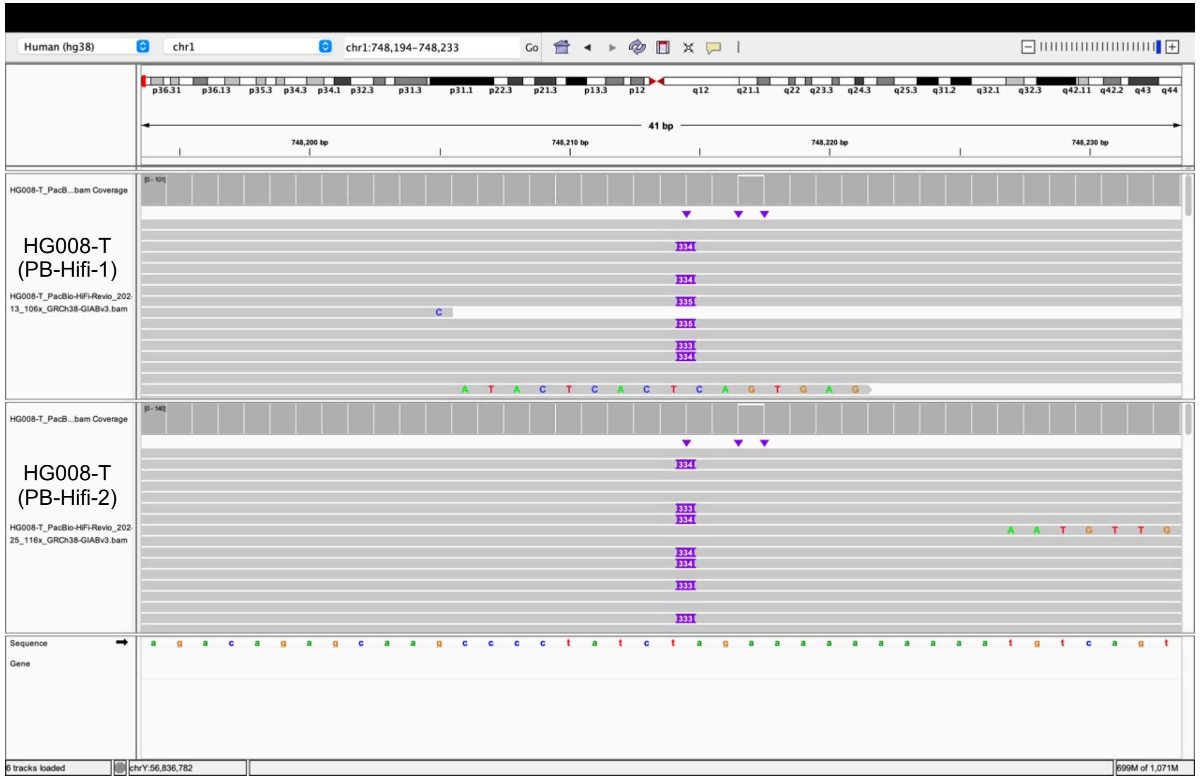


#### Same insertion in normal tissue

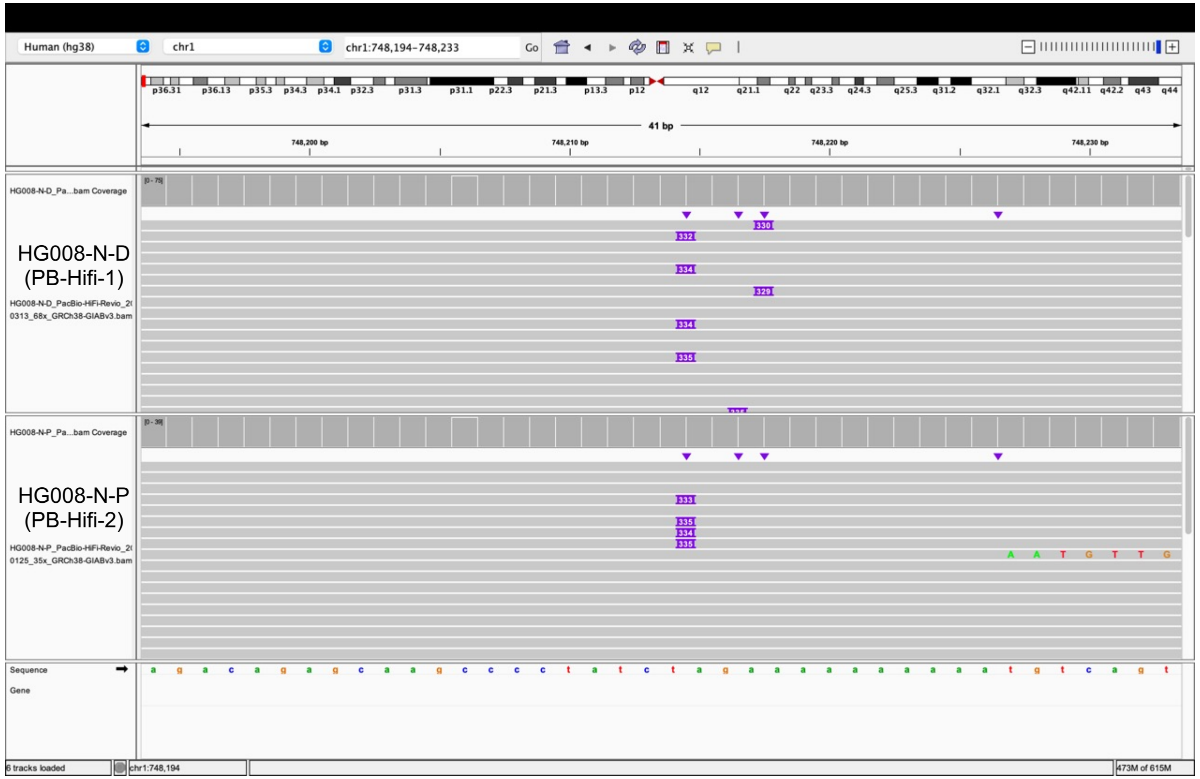

### chr1:121655868 (Found in normal)

Tumor tissue

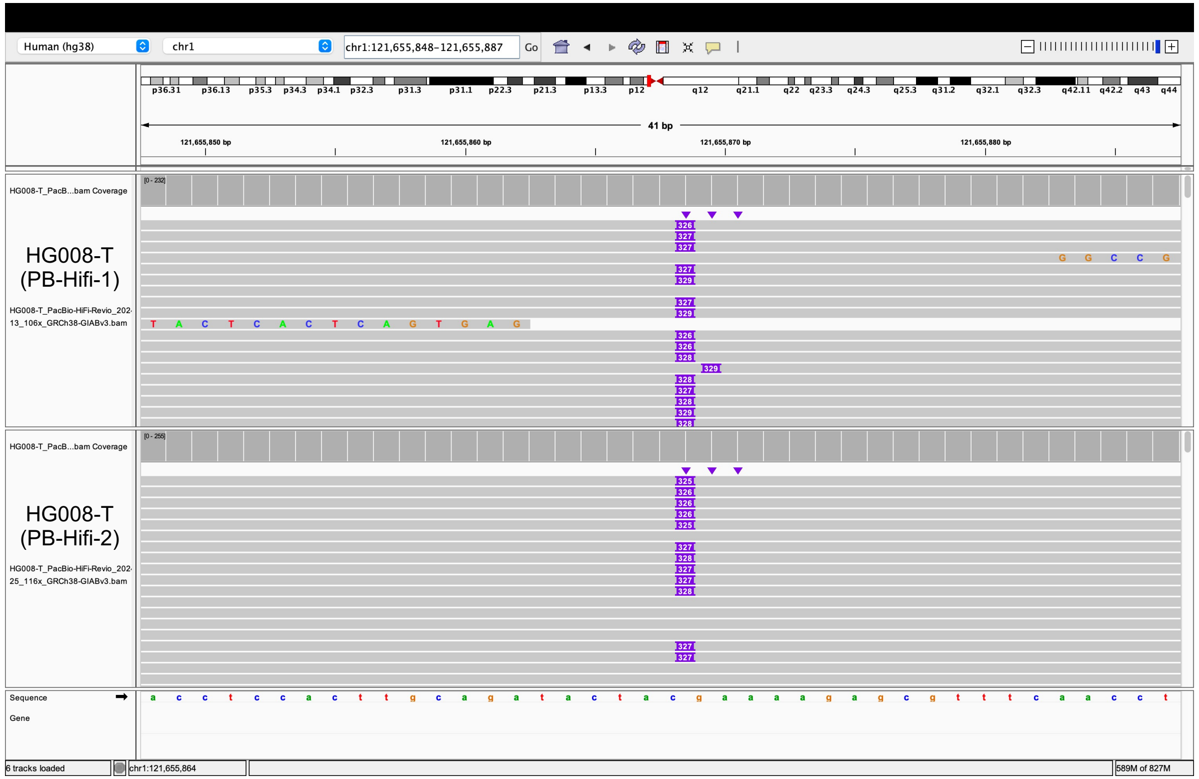


Same insertion in normal tissue

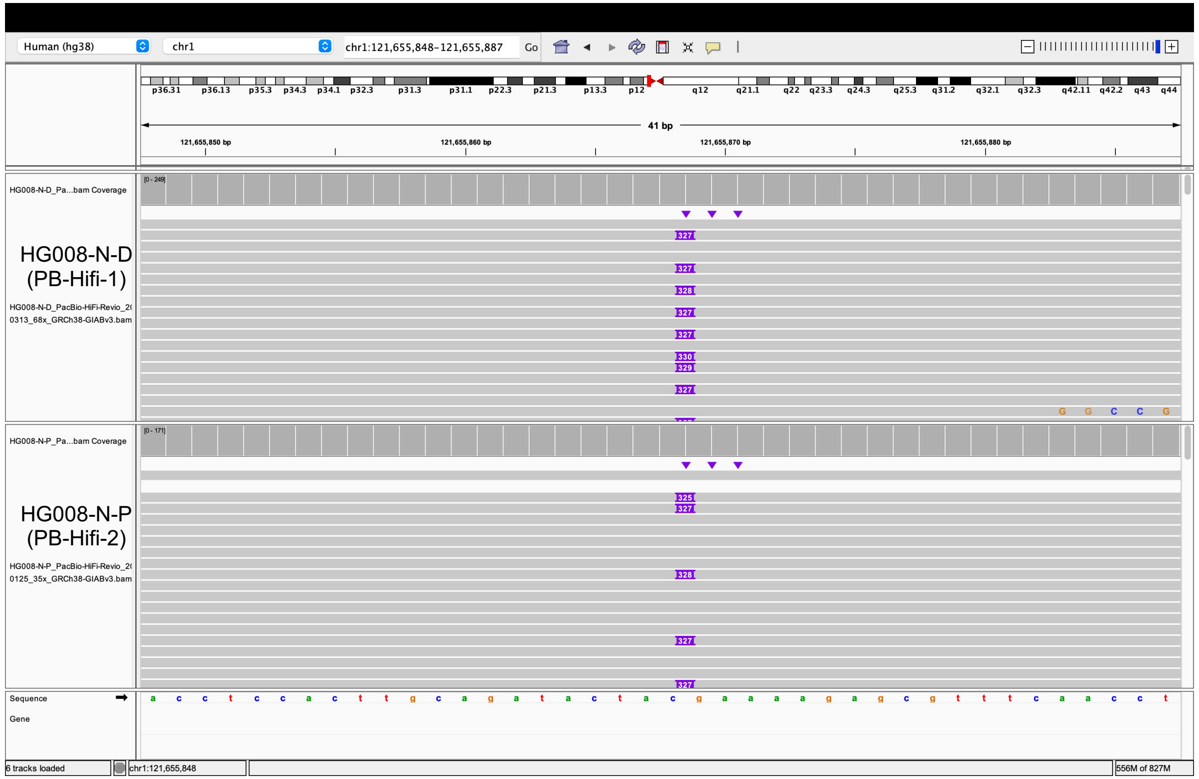

### chr2:91441826 (Found in normal)

Tumor tissue

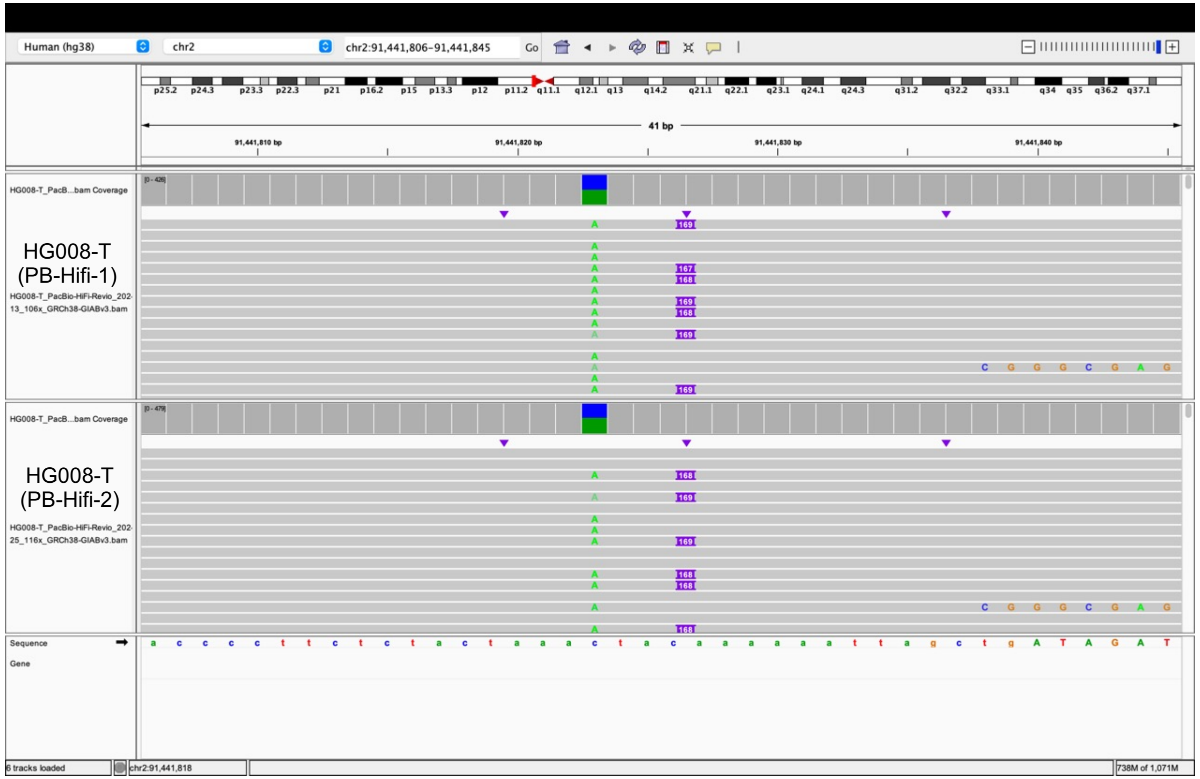


Same insertion in normal tissue

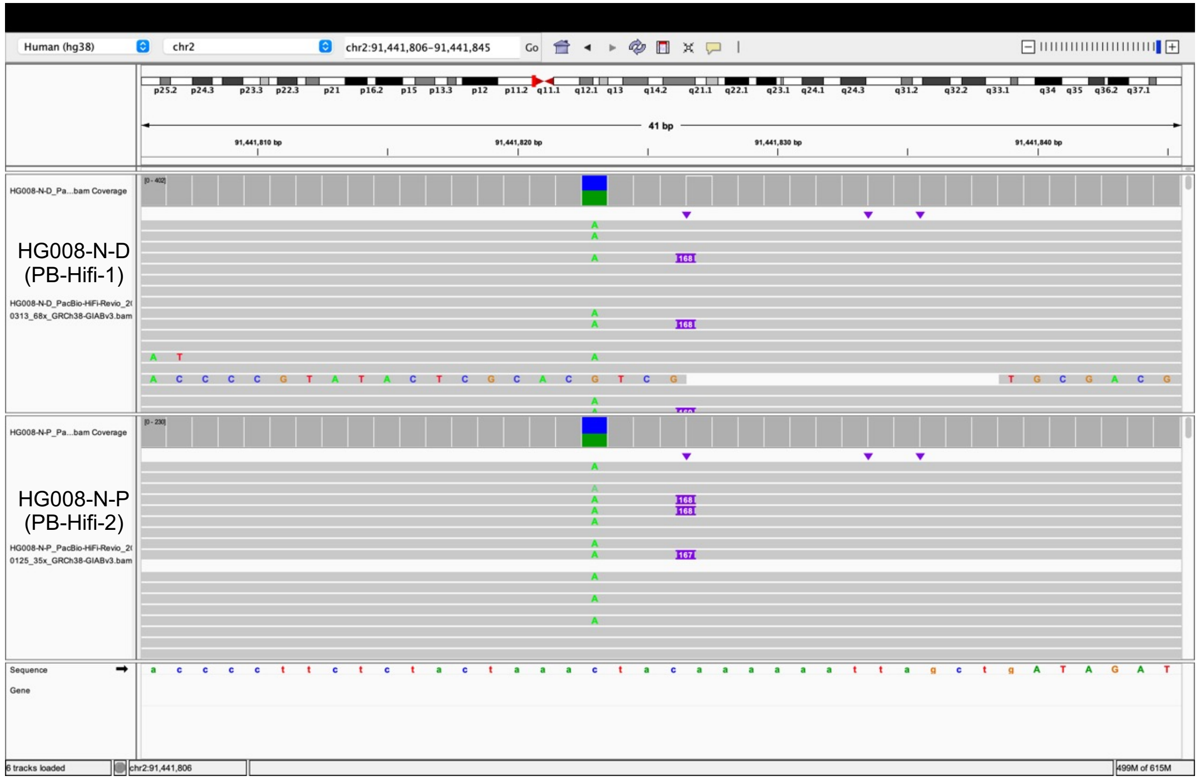

### chr2:242181270 (Found in normal)

Tumor tissue

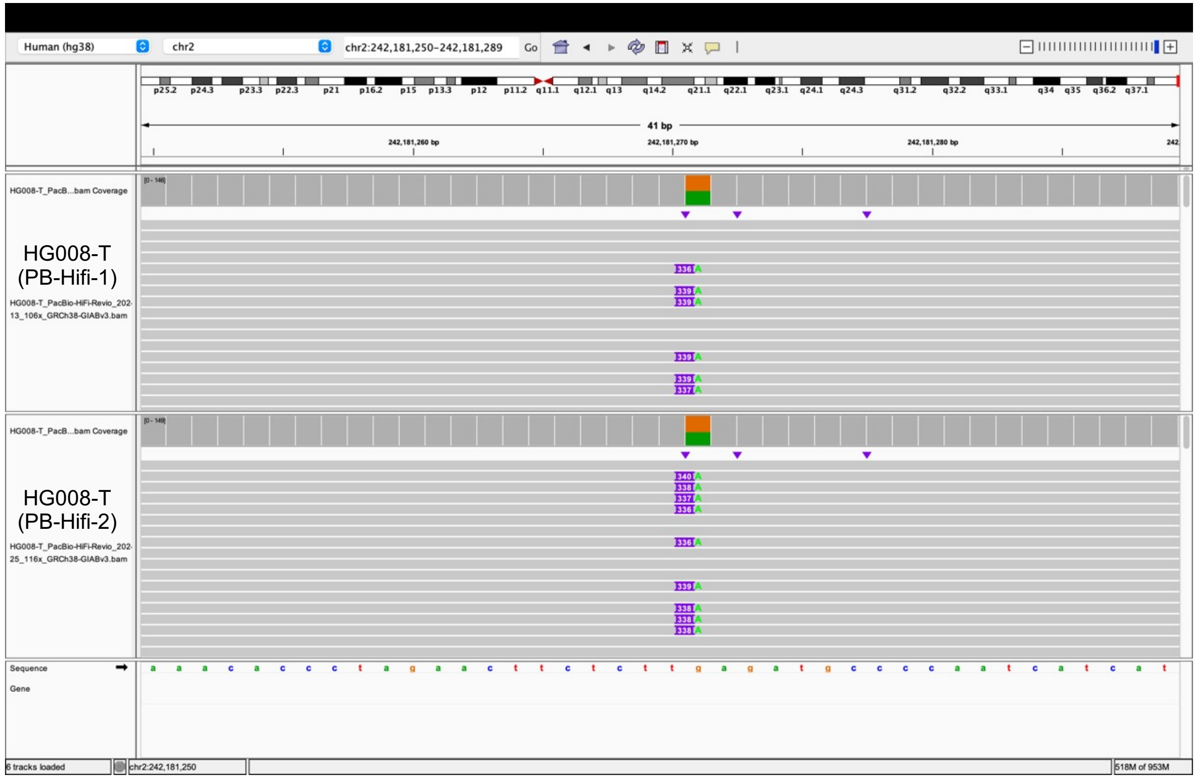


Same insertion in normal tissue

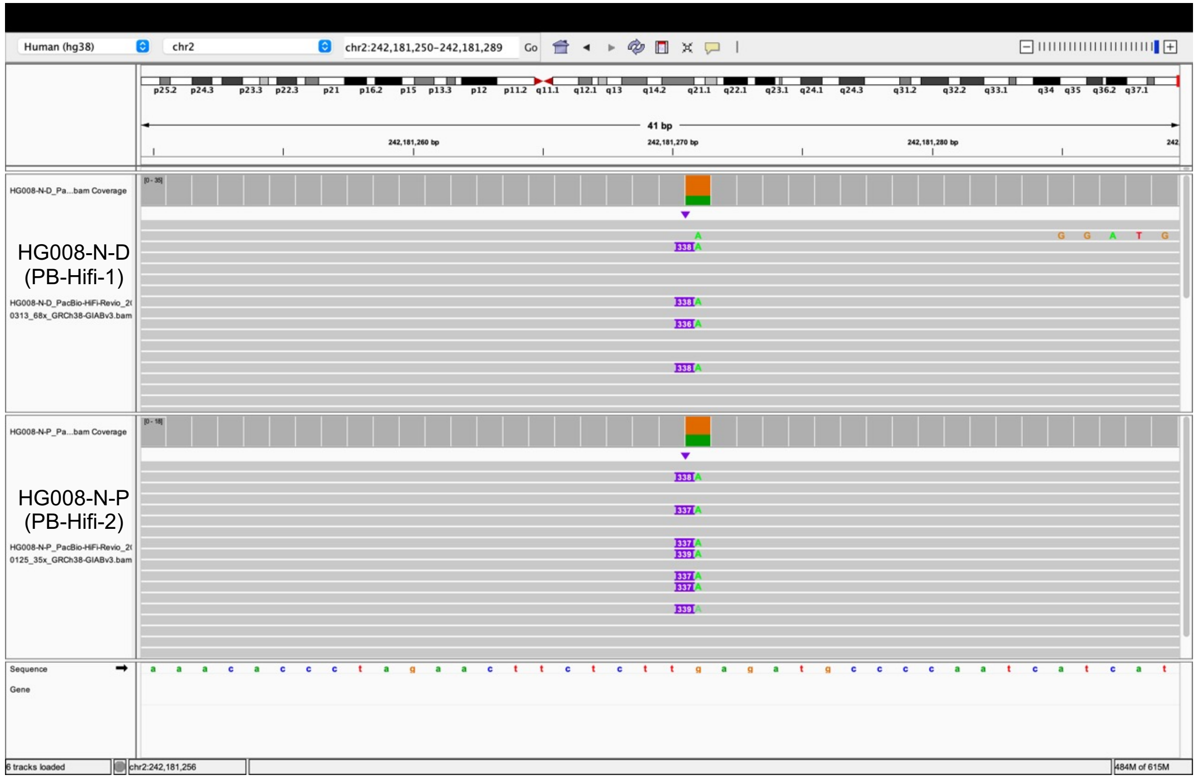

### chr6:32587099 (Found in normal)

Tumor tissue

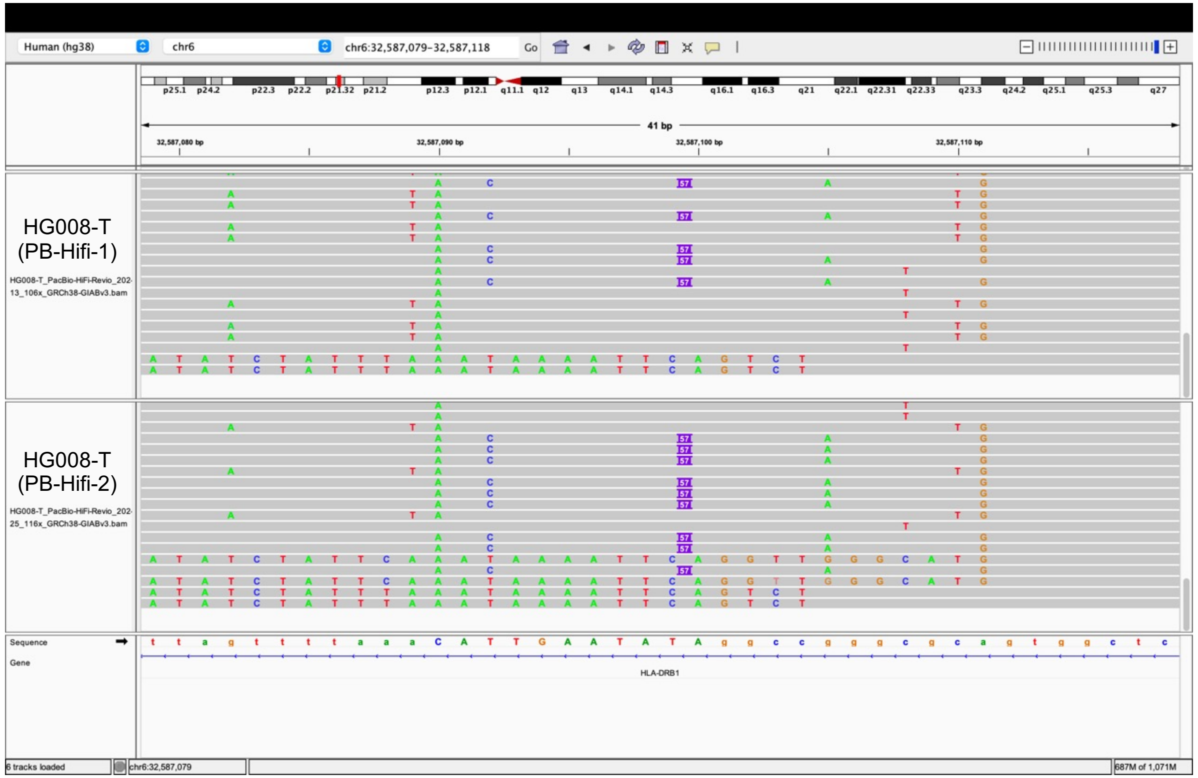


Same insertion in normal tissue

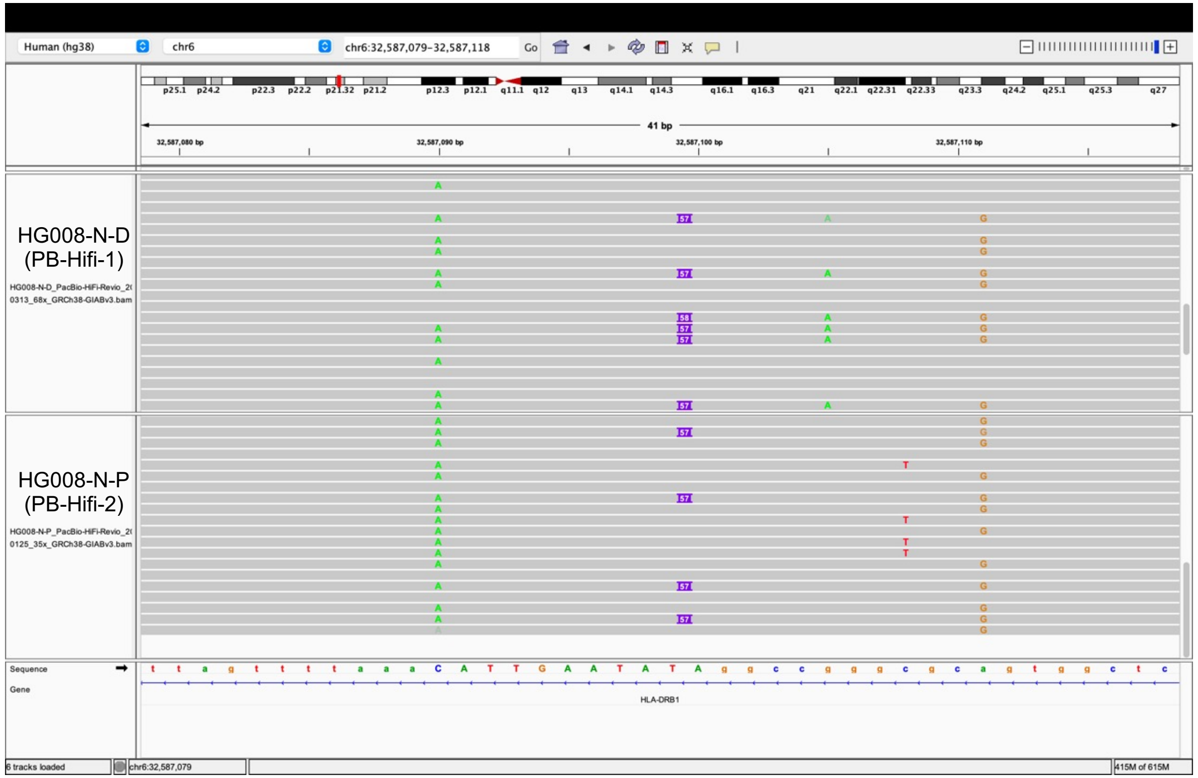

### chr6:28348988 (Found in normal)

Tumor tissue

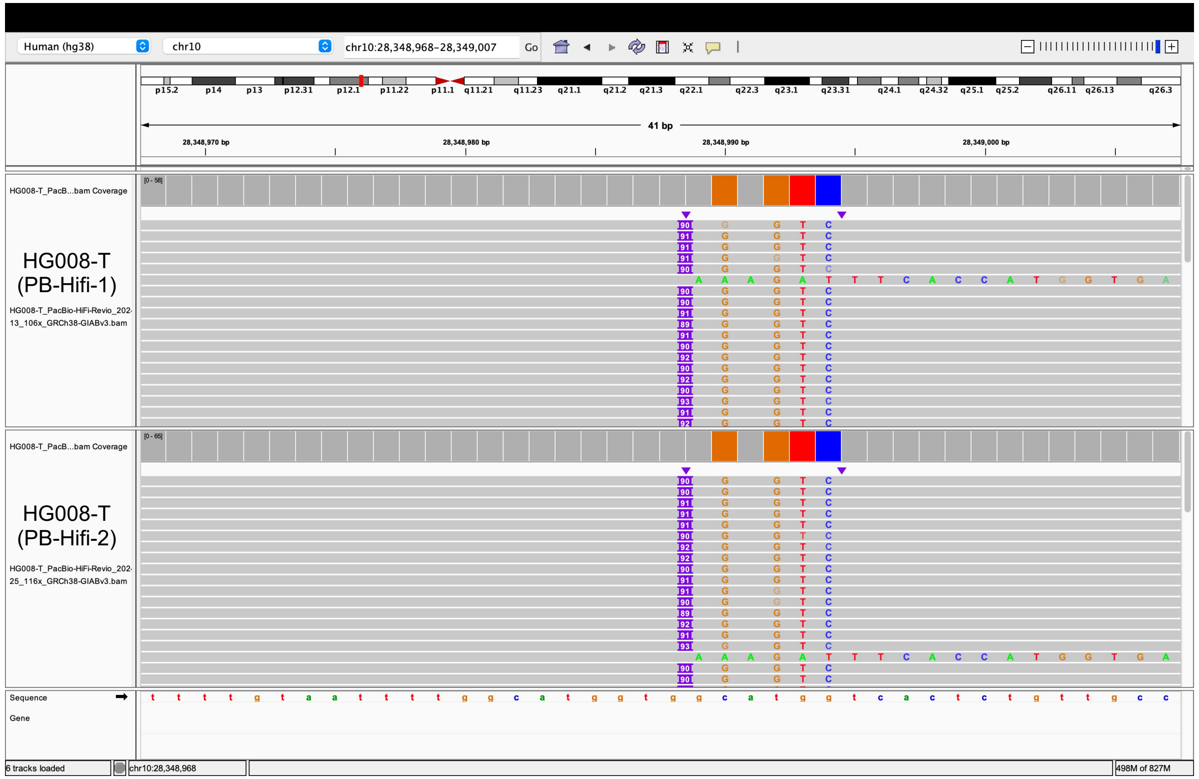


Same insertion in normal tissue

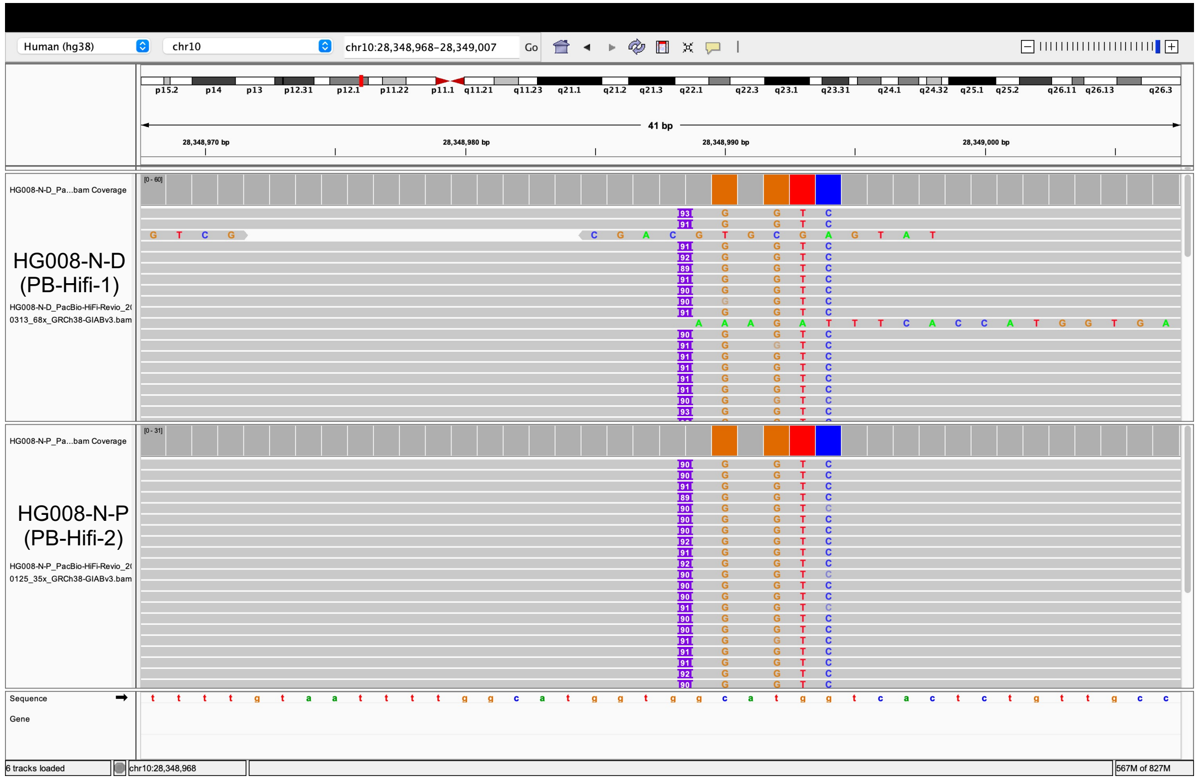

### chr15:22584273 (Found in normal)

Tumor tissue

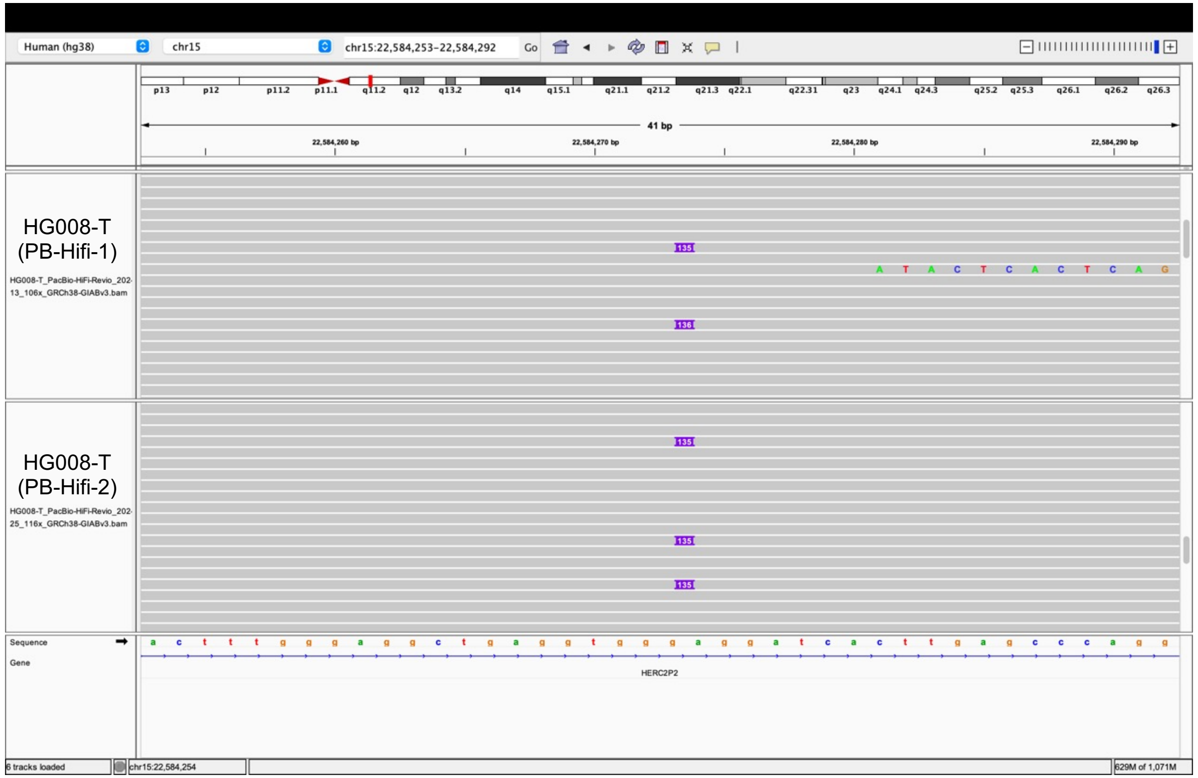


Same insertion in normal tissue

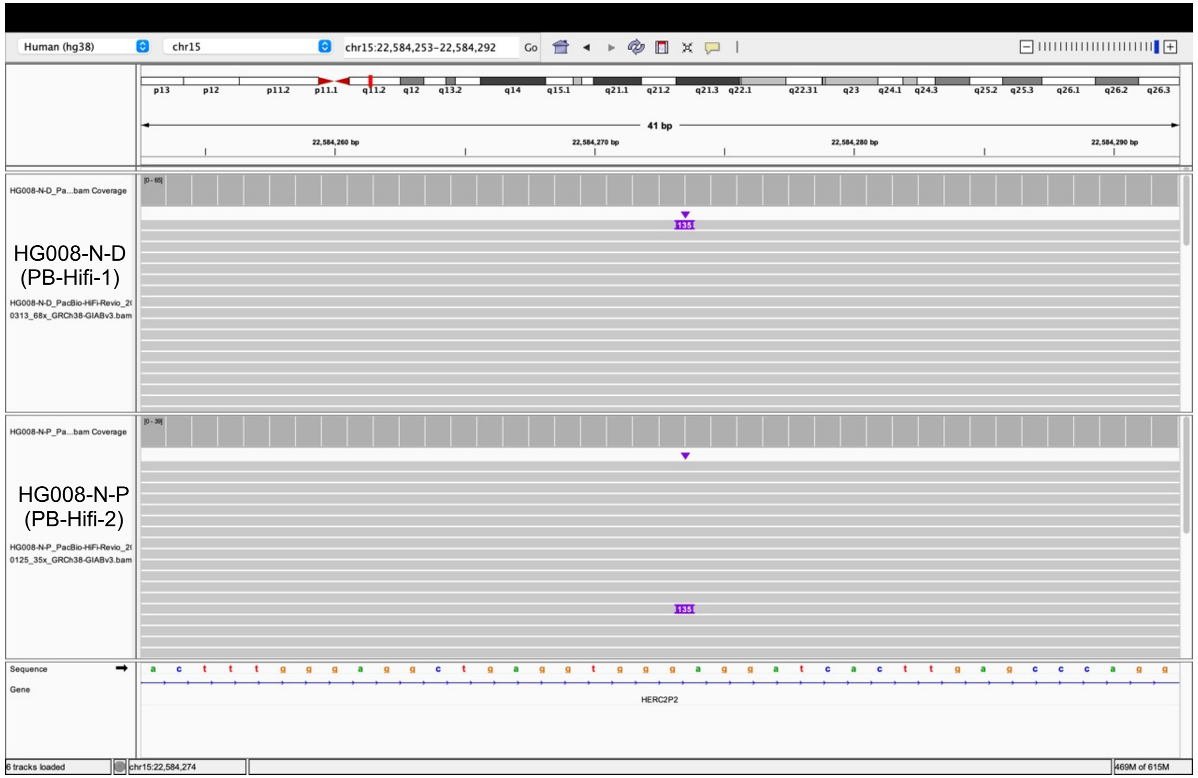

### chr19:58605628 (Found in normal)

Tumor tissue

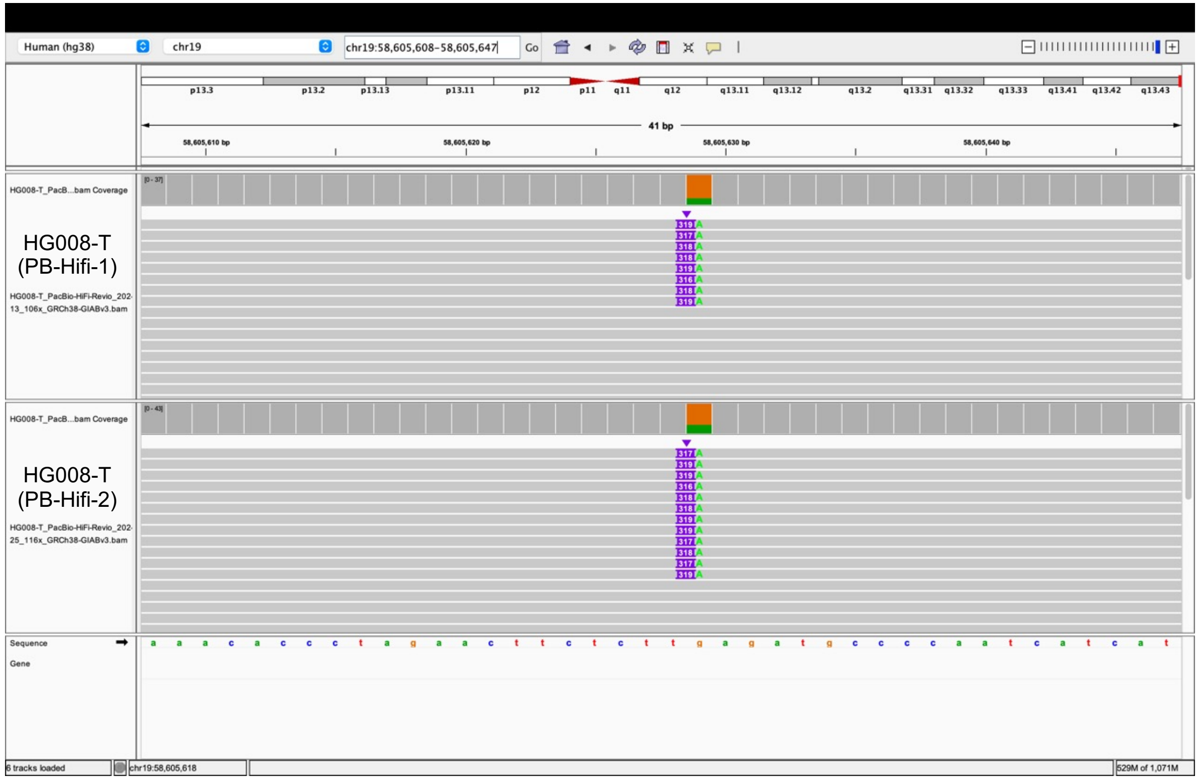


Same insertion in normal tissue

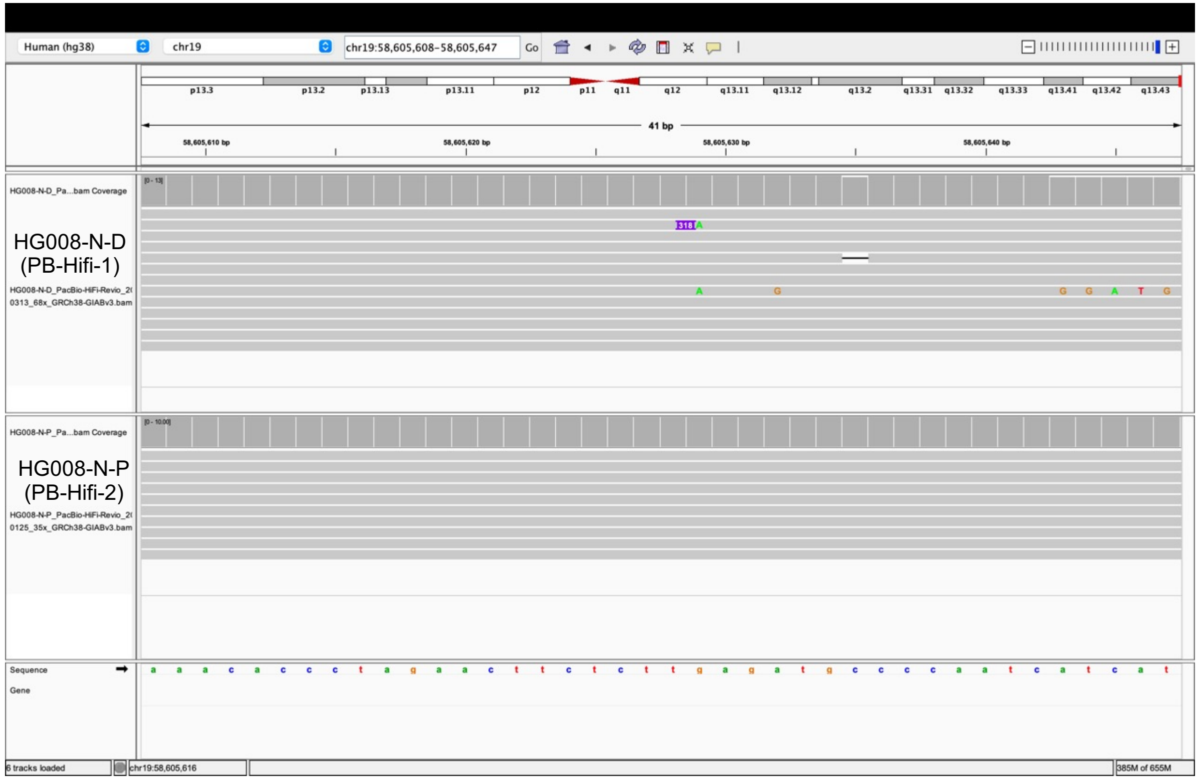

### chr22:12595451 (Found in normal)

#### Tumor tissue

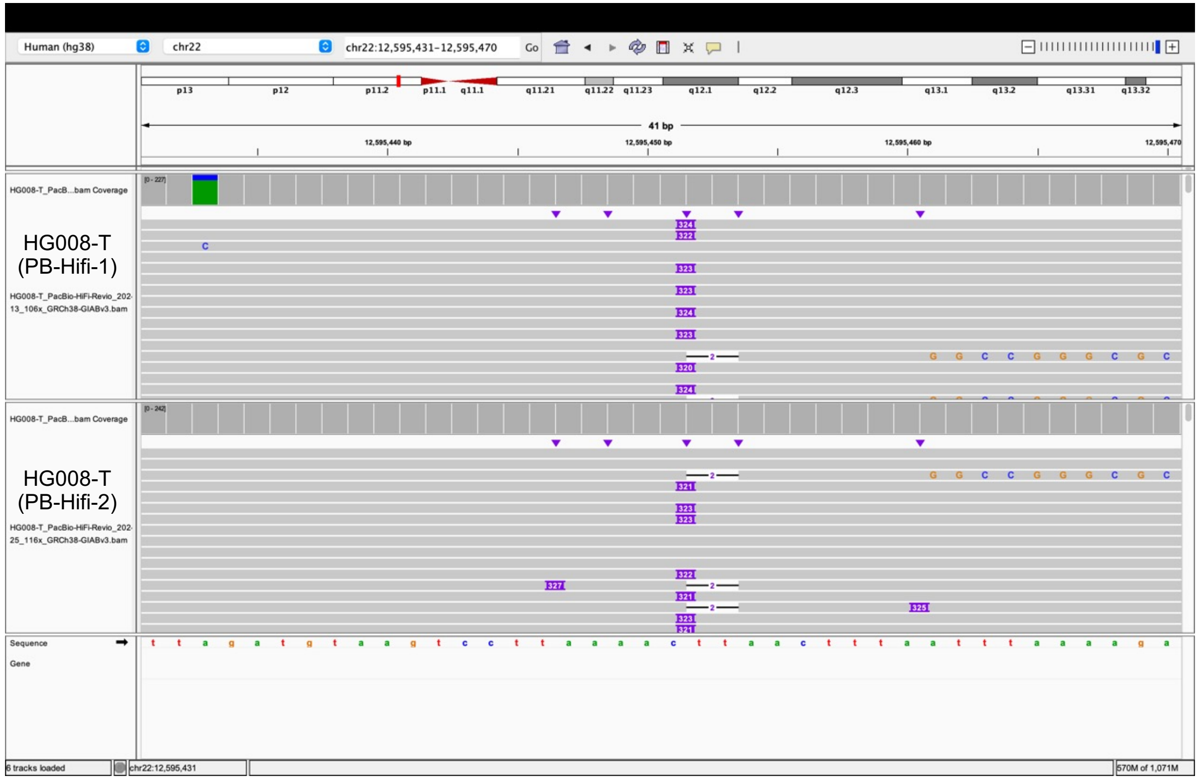


#### Same insertion in normal tissue

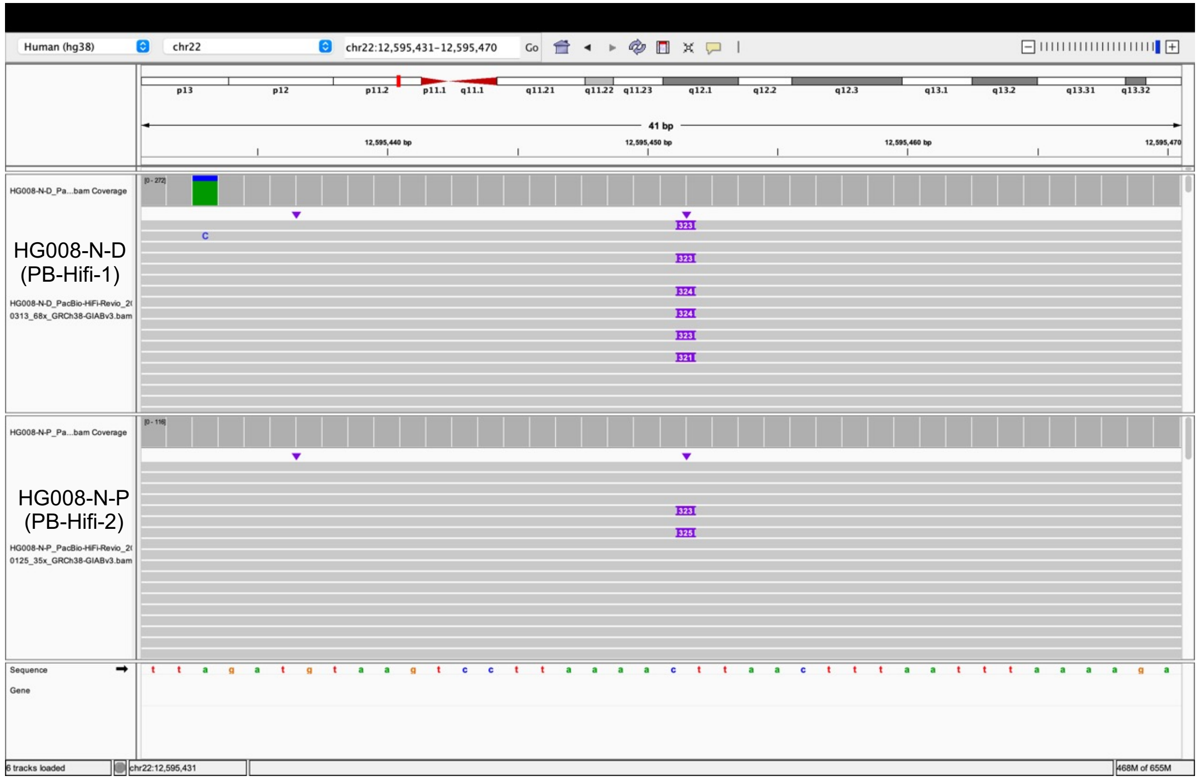

### chr22:16476005 (Found in normal)

#### Tumor tissue

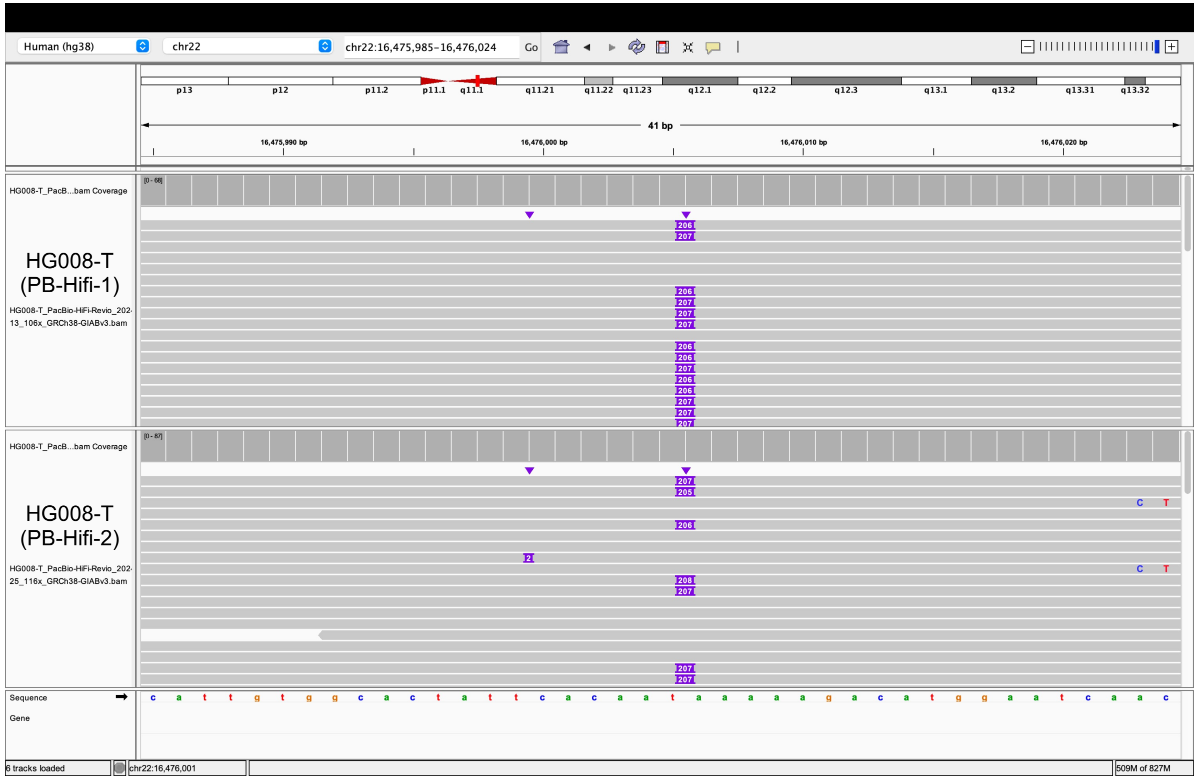


#### Same insertion in normal tissue

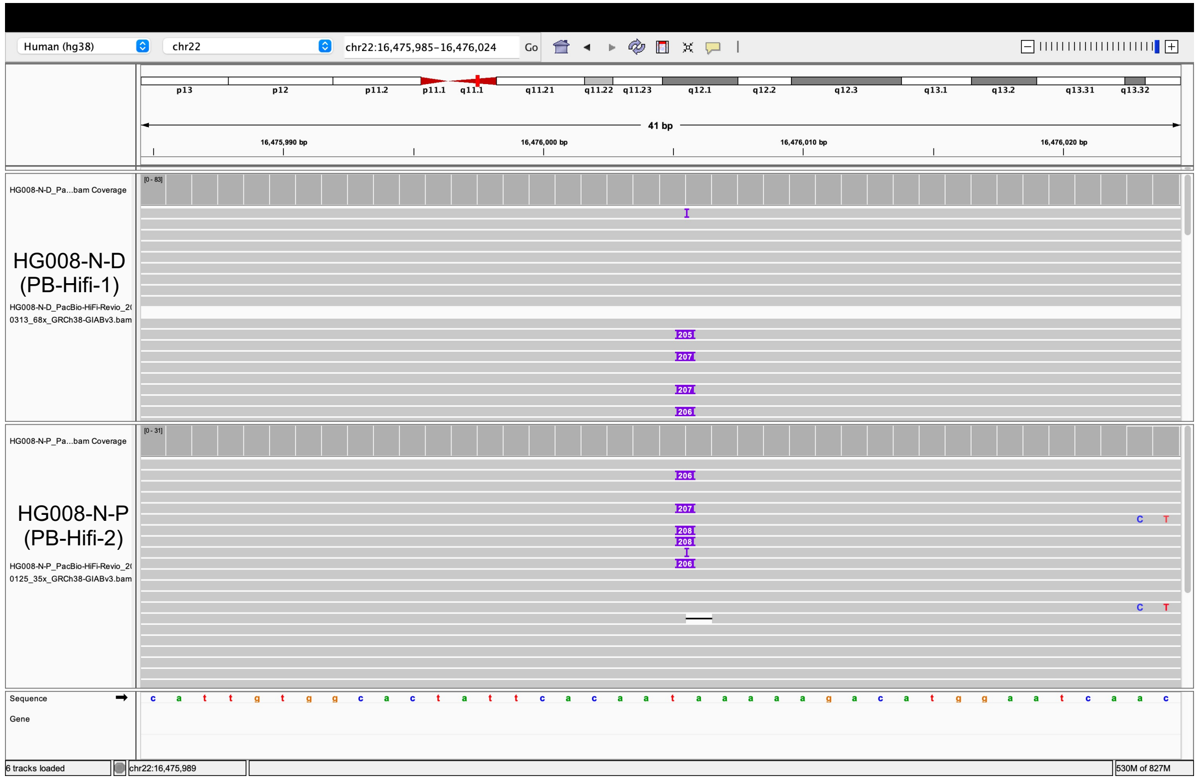

### chrX:1067023 (Found in normal)

#### Tumor tissue

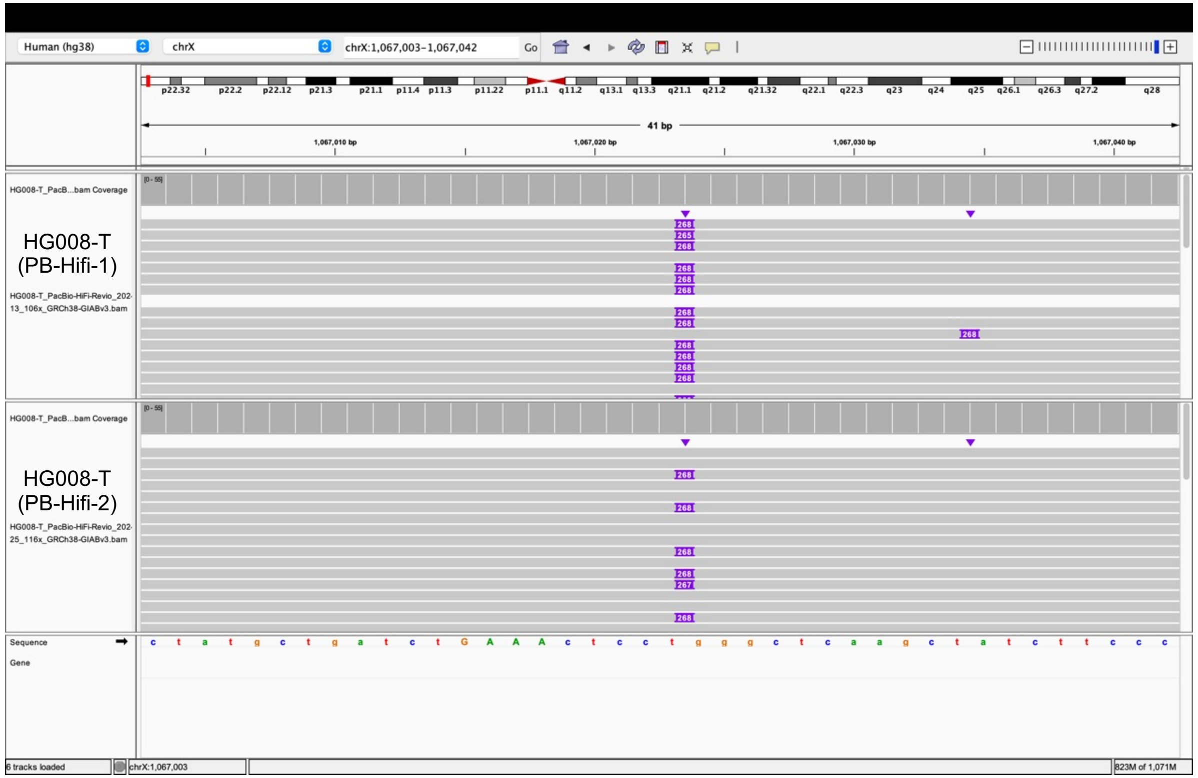


#### Same insertion in normal tissue

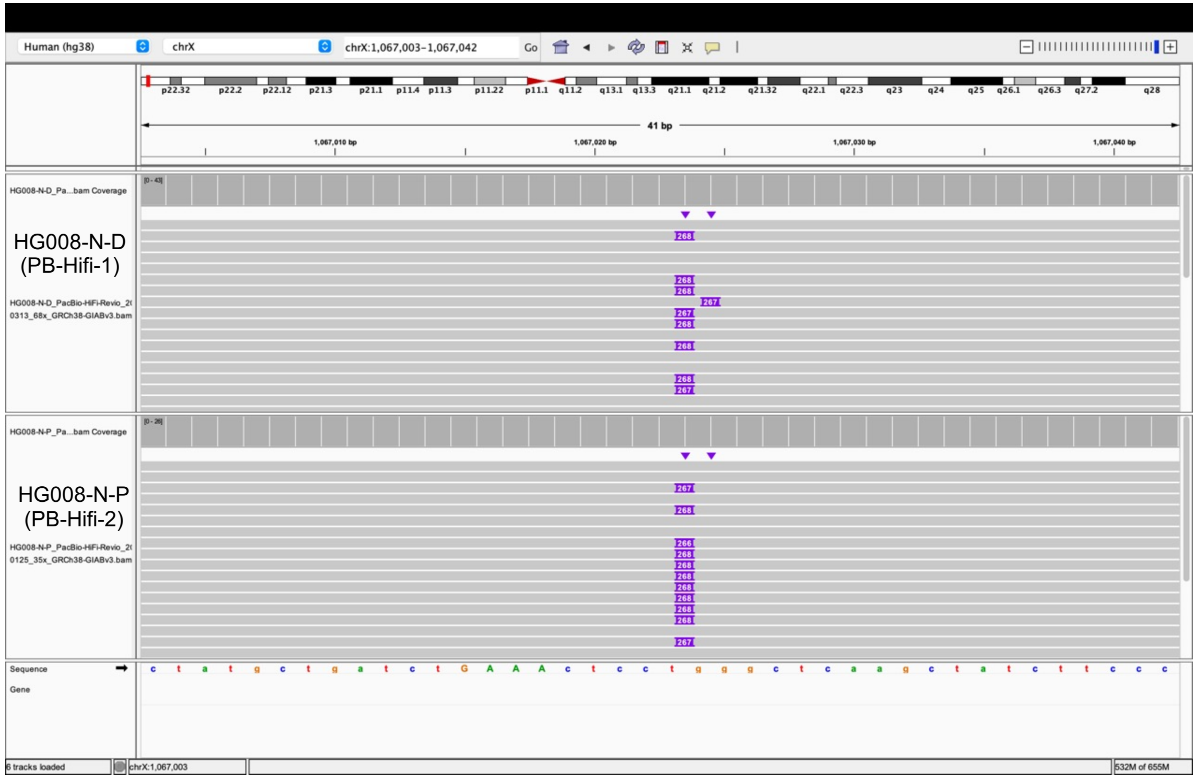

### chrX:321575 (Found in normal)

#### Tumor tissue

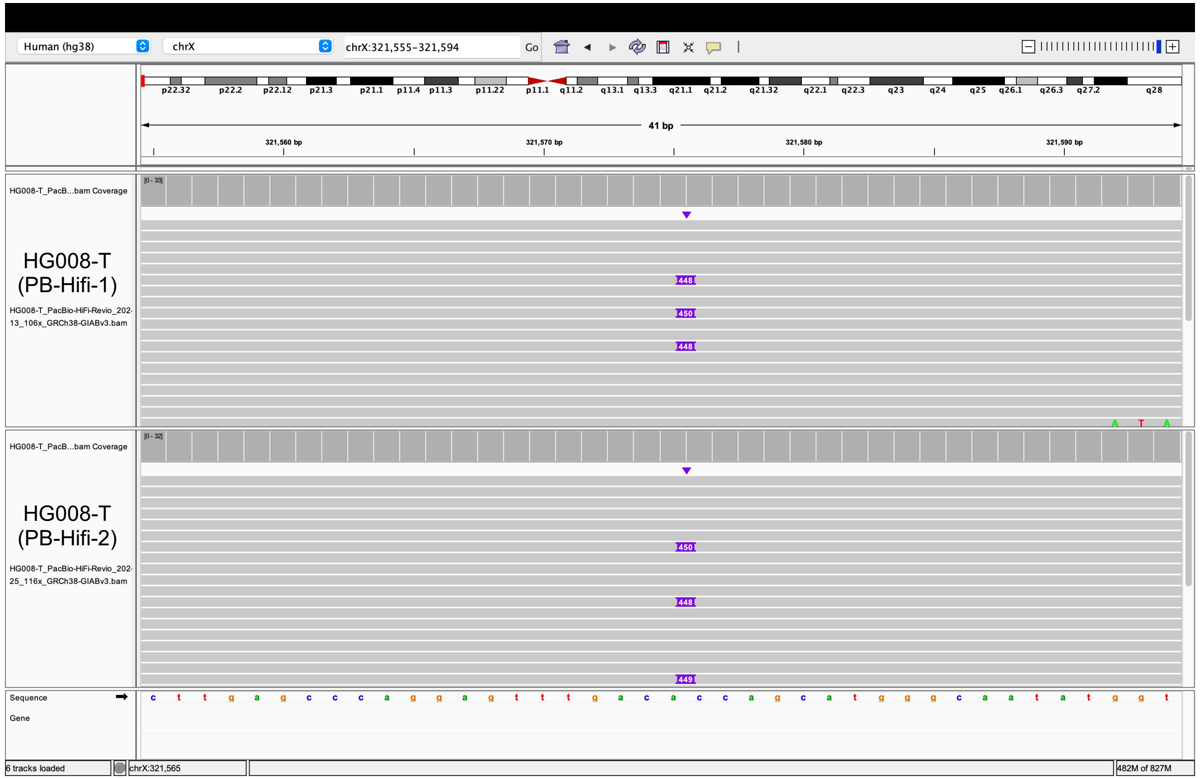


#### Same insertion in normal tissue

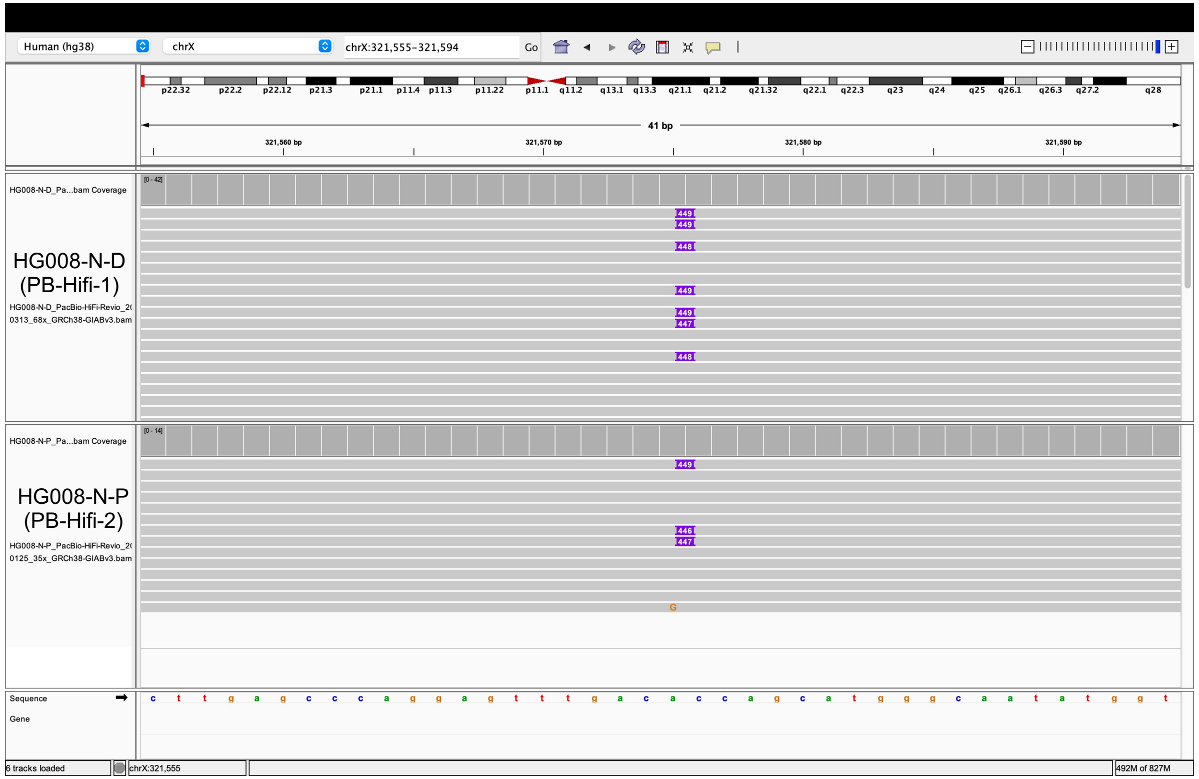

### chrX:1334423 (Found in normal)

#### Tumor tissue

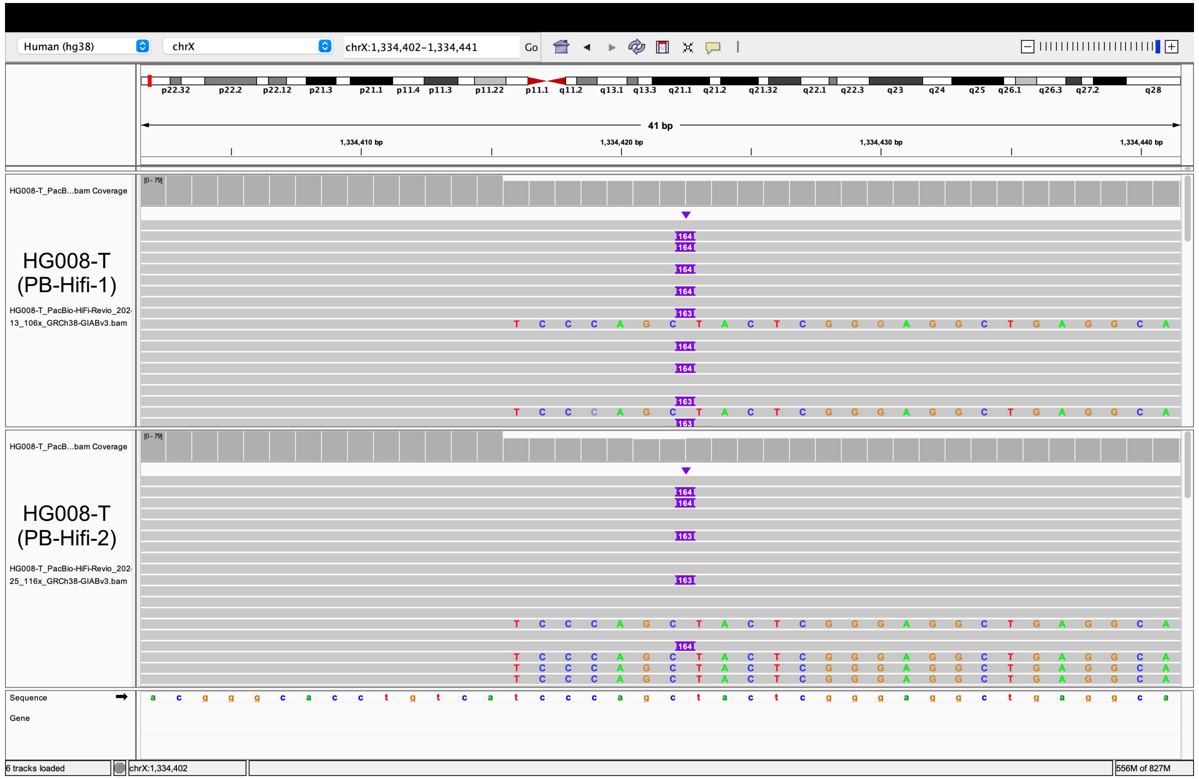


#### Same insertion in normal tissue

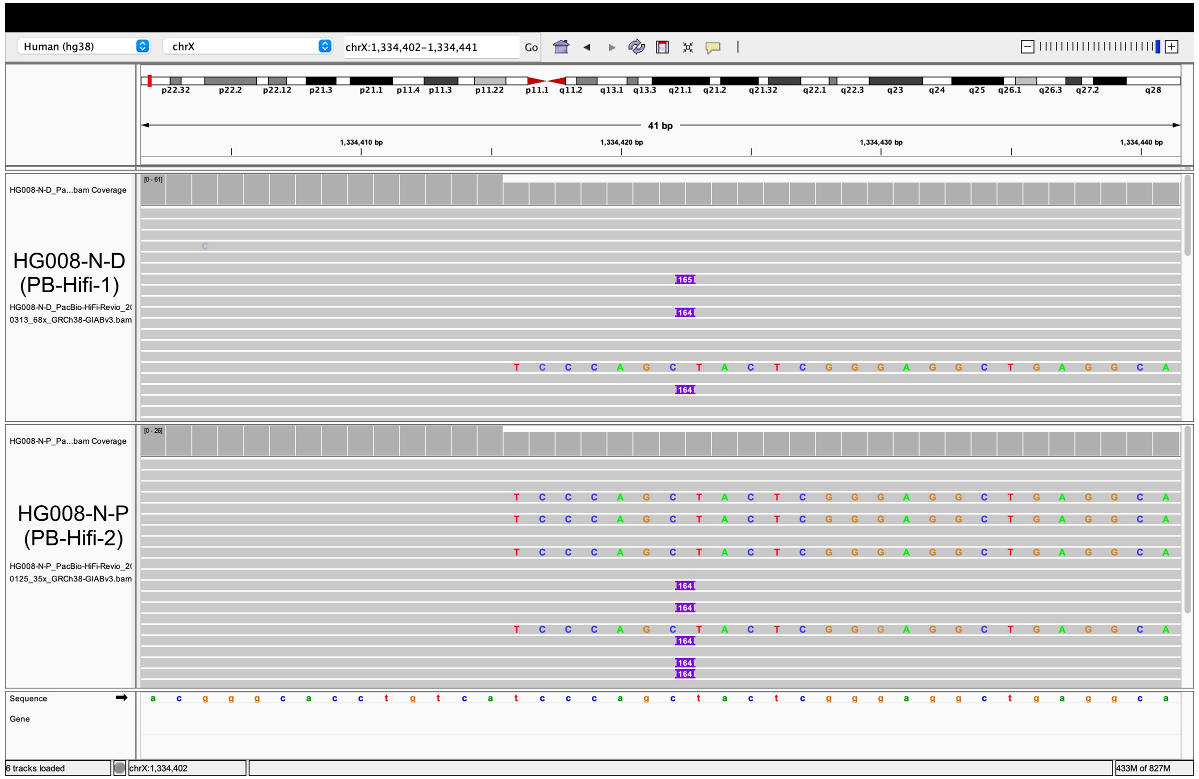

### chrX:1432859 (Found in normal)

#### Tumor tissue

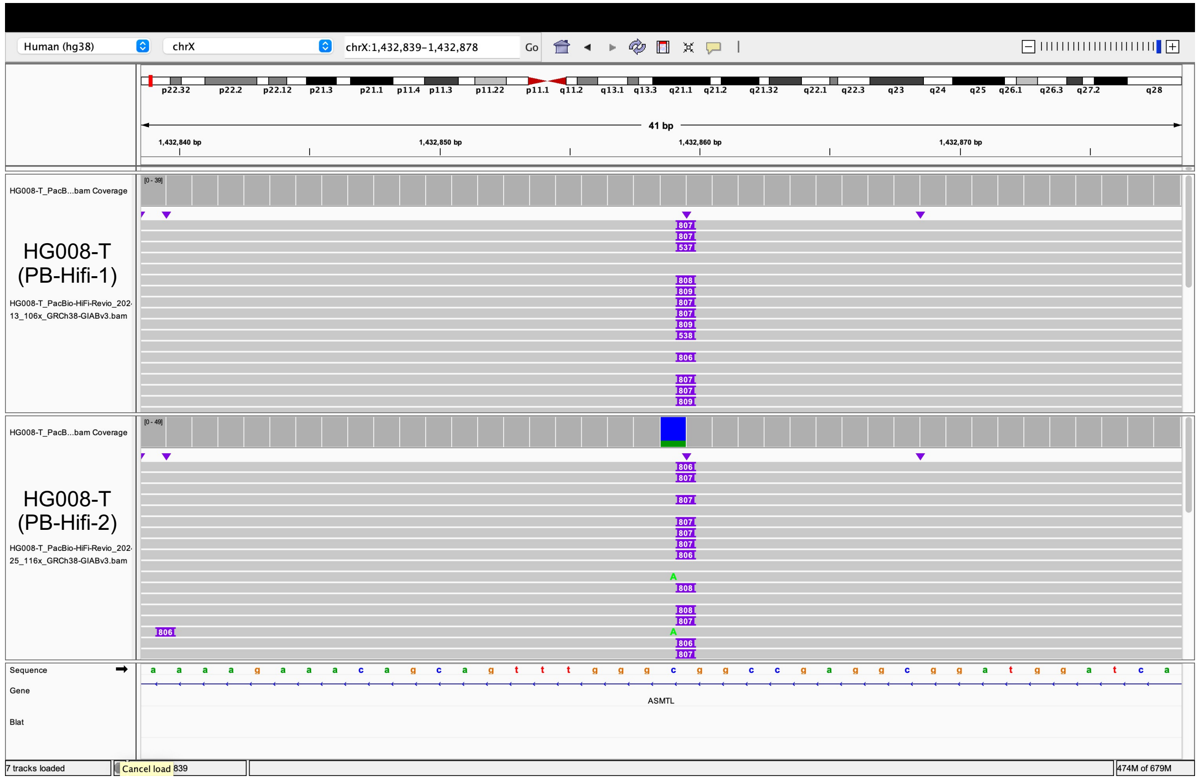


#### Same insertion in normal tissue

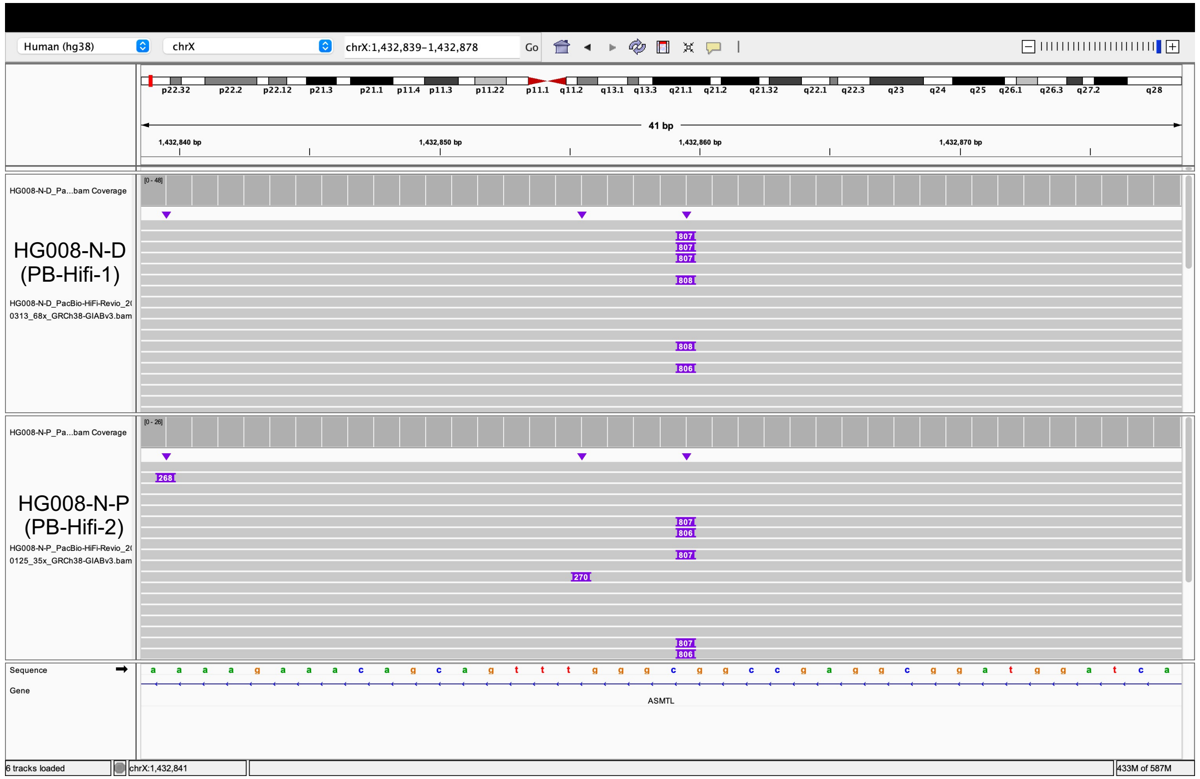

### chrY:11349045 (Found in normal)

Tumor tissue

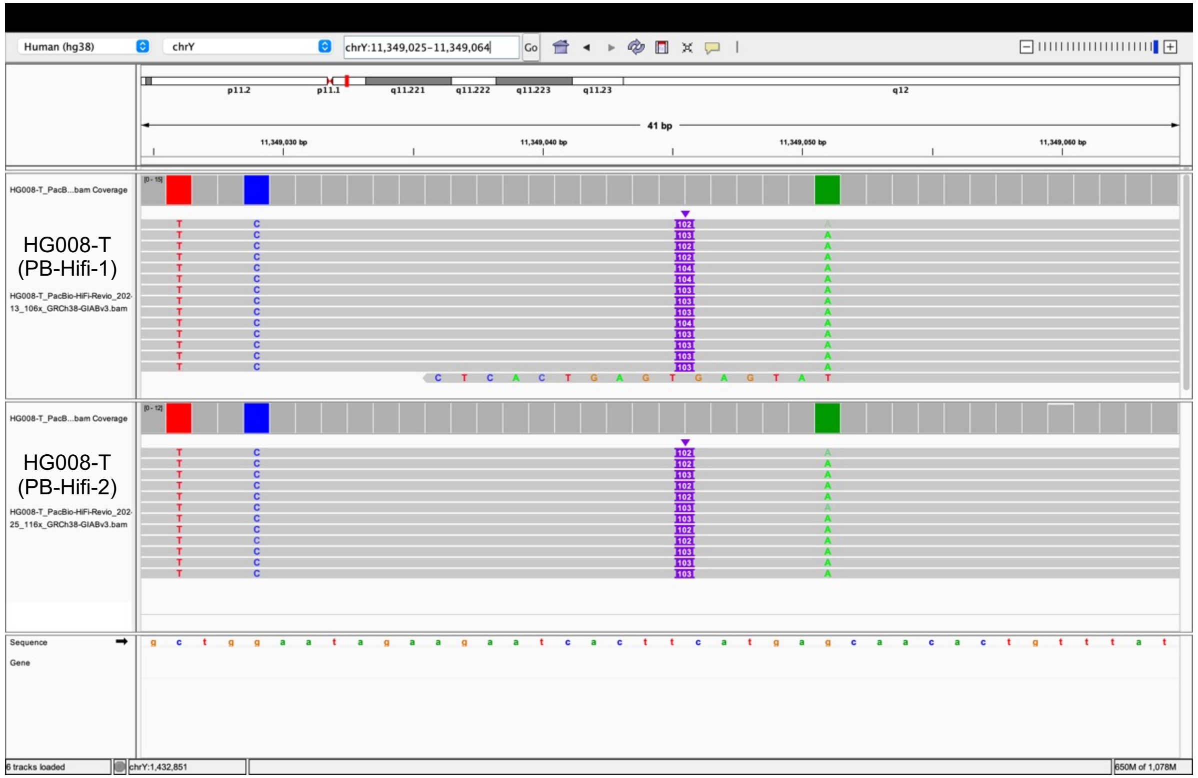


Same insertion in normal tissue

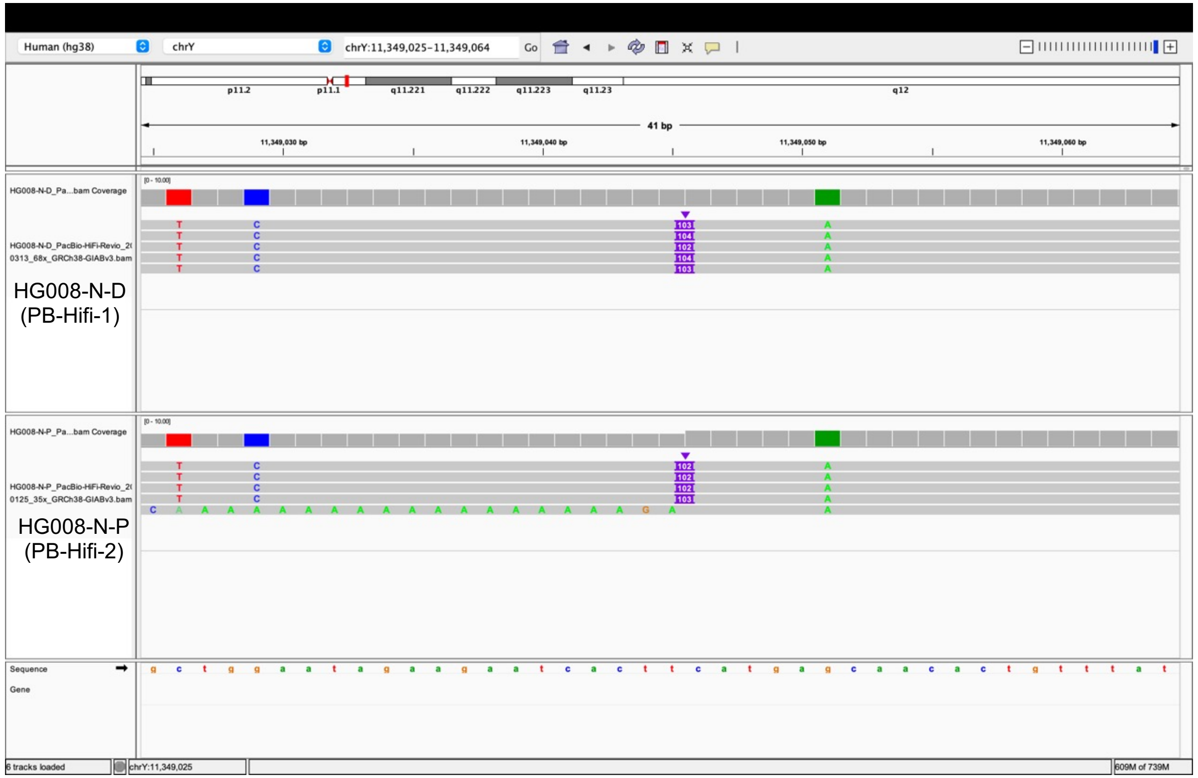
