## Extended Files for "Image-based DNA Sequencing Encoding for Detecting Low-Mosaicism Somatic Mobile Element Insertions": Extended Files 7.pdf

### Tumor somatic SVA Insertions detected in HG008-T by PALMER or xTea_long

(Additional explanation for Supplementary Table 10)

| Chr | Start | End | PALMER detected in PB-Hifi-1 | PALMER detected in PB-Hifi-2 | xTea_long detected in PB-Hifi-1 | xTea_long detected in PB-Hifi-2 | In benchmark list |
|---|---|---|---|---|---|---|---|
| chr1 | 234397802 | 234397803 | | | | + | no (found in normal) |
| chr14 | 90300875 | 90300876 | | | + | + | no (found in normal) |
| chr20 | 30880755 | 30880756 | | | + | + | no (found in normal) |

**Note:**

No benchmark tumor somatic SVA insertions were identified during benchmarking. PALMER also did not identified any SVA insertions. However, xTea_long erroneously classified 3 *Alu* insertions. Upon manual review using IGV, we confirmed that these insertions were also present in the normal samples, indicating they are not tumor somatic events.

Bam files of the tumor and normal datasets were accessed through the following links:

- HG008-T (PB-Hifi-1): https://42basepairs.com/download/s3/giab/data_somatic/HG008/Liss_lab/BCM_Revio_20240313/HG008-T_PacBio-HiFi-Revio_20240313_106x_GRCh38-GIABv3.bam
- HG008-T (PB-Hifi-2): https://42basepairs.com/download/s3/giab/data_somatic/HG008/Liss_lab/PacBio_Revio_20240125/HG008-T_PacBio-HiFi-Revio_20240125_116x_GRCh38-GIABv3.bam
- HG008-N-D (PB-Hifi-1): https://42basepairs.com/download/s3/giab/data_somatic/HG008/Liss_lab/BCM_Revio_20240313/HG008-N-D_PacBio-HiFi-Revio_20240313_68x_GRCh38-GIABv3.bam
- HG008-N-P (PB-Hifi-2): https://42basepairs.com/download/s3/giab/data_somatic/HG008/Liss_lab/PacBio_Revio_20240125/HG008-N-P_PacBio-HiFi-Revio_20240125_35x_GRCh38-GIABv3.bam

### chr1:234397802

Tumor tissue

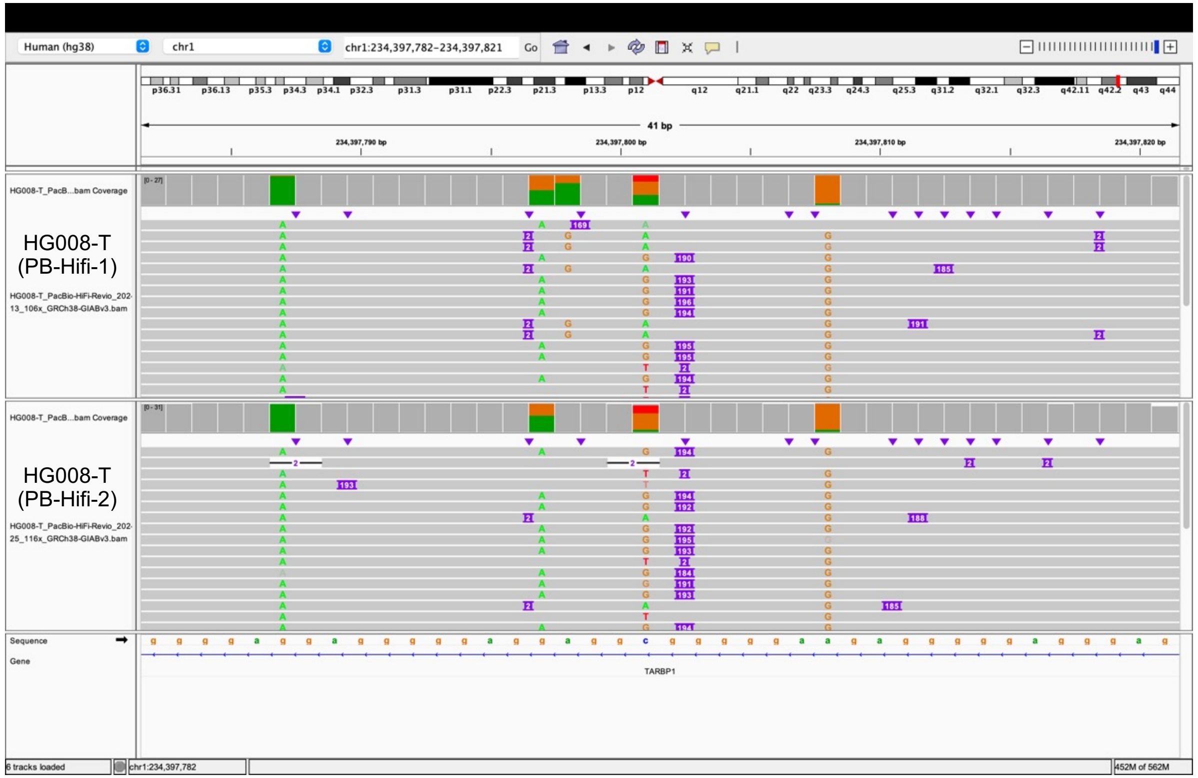


Same insertion in normal tissue

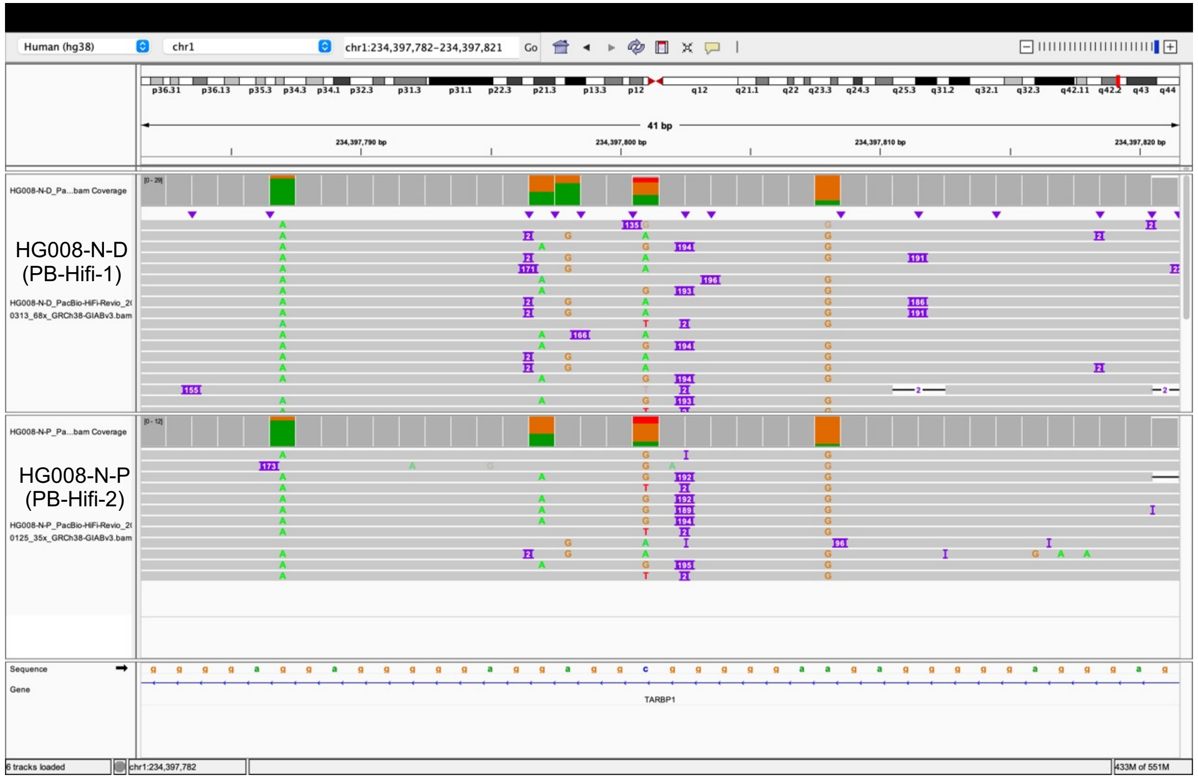

### chr14:90300882

#### Tumor tissue

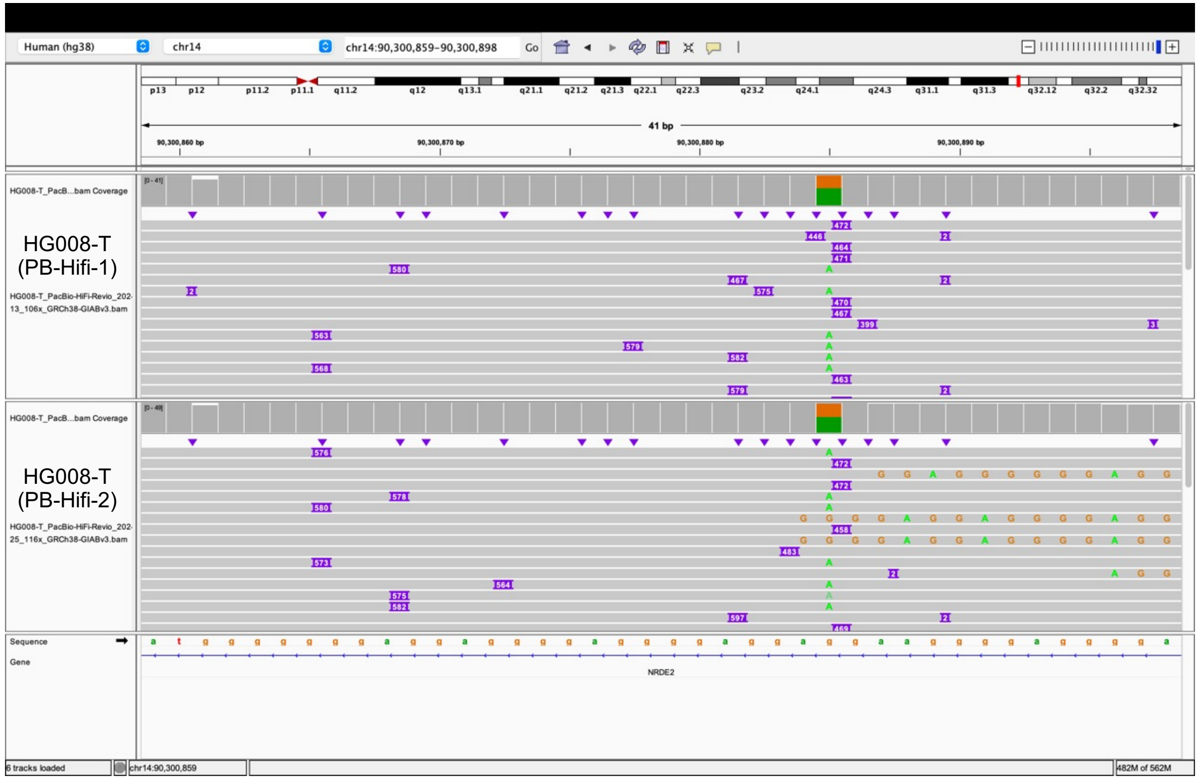


#### Same insertion in normal tissue

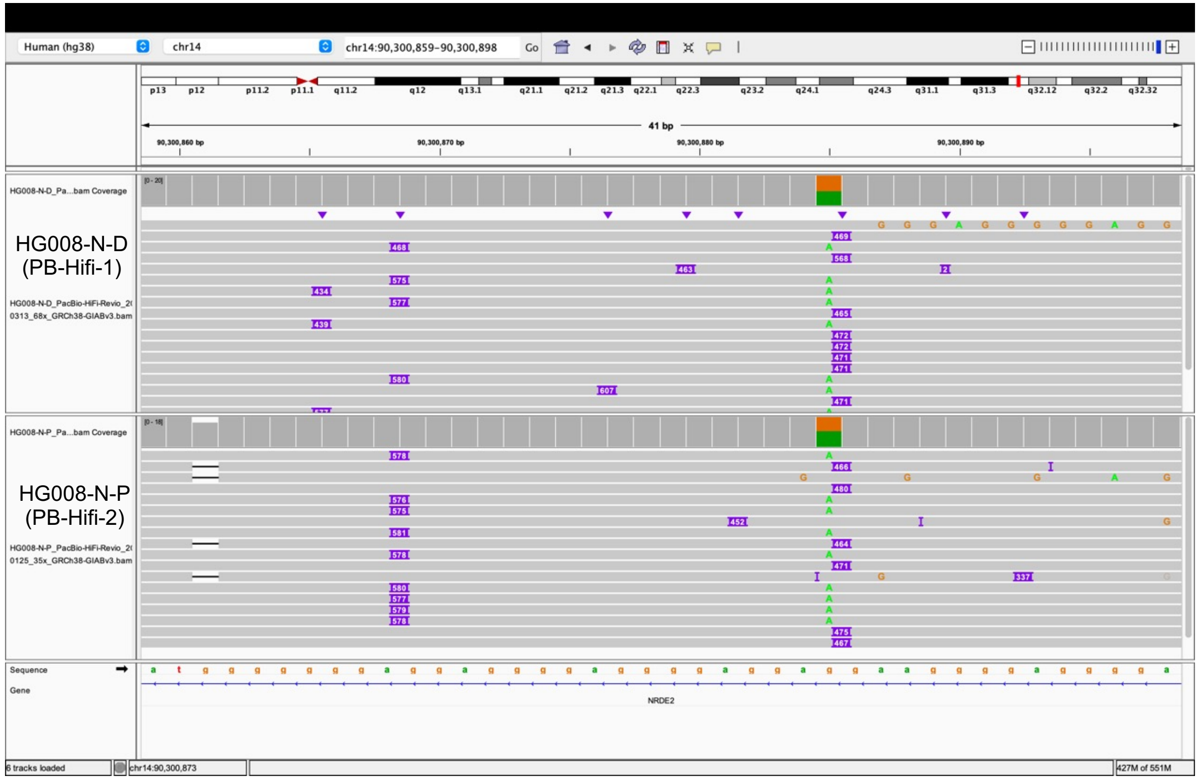

### chr20:30880755

Tumor tissue

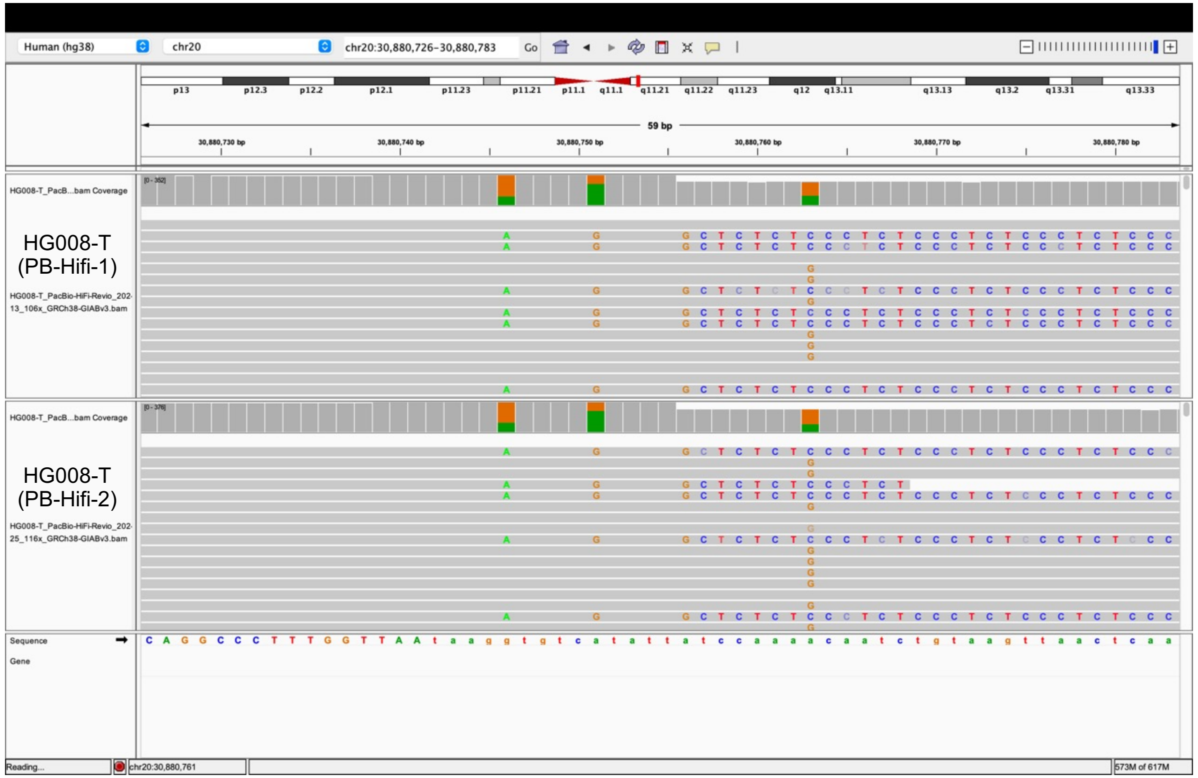


Same insertion in normal tissue

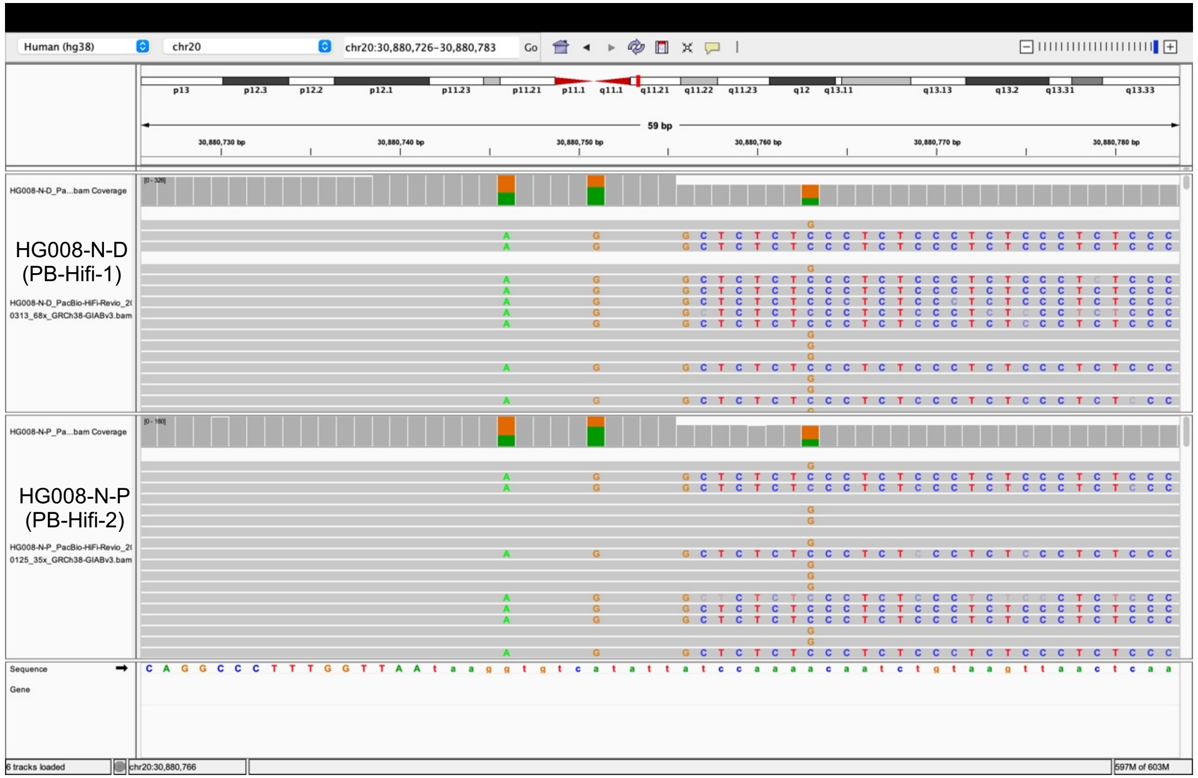
