## Extended Files for "Image-based DNA Sequencing Encoding for Detecting Low-Mosaicism Somatic Mobile Element Insertions": Extended Files 8.pdf

### Detailed visualization of patient DTB-205 cfDNA somatic L1 insertions identified by RetroNet with low mosaicism in the tumor tissue

(Additional explanation for Supplementary Table 7)

| *Chr* | *Start* | *End* | *Supporting Reads (in cfDNA)* | *Depth (in cfDNA)* | *Mosaicism* | *RetroNet (in cfDNA)* | *xTea (in cfDNA)* | *xTea (in tumor)* | *Details* |
|---|---|---|---|---|---|---|---|---|---|
| chr12 | 84700007 | 84700632 | 11 | 198 | 11.09% | + | - | - | Not pass xTea threshold (cfDNA split-reads = 7, Tumor split-reads = 8) |
| chr3 | 16729240 | 16729865 | 11 | 222 | 9.89% | + | - | - | Not pass xTea threshold (cfDNA split-reads = 3, Tumor split-reads = 5) |
| chr6 | 90288299 | 90288337 | 9 | 185 | 9.74% | + | - | - | Not pass xTea threshold (cfDNA split-reads = 4, Tumor split-reads = 0, Tumor discordant pairs = 1) |
| chr6 | 2688545 | 2688595 | 10 | 222 | 9.02% | + | - | - | Not pass xTea threshold (cfDNA split-reads = 6, Tumor split-reads = 4) |
| chr20 | 18942311 | 18942351 | 11 | 247 | 8.91% | + | - | - | Not pass xTea threshold (cfDNA split-reads = 2, Tumor split-reads = 1) |
| chr11 | 86269462 | 86269495 | 10 | 226 | 8.85% | + | - | - | Not pass xTea threshold (cfDNA split-reads = 2, Tumor split-reads = 0, Tumor discordant pairs = 1) |
| chr4 | 54692604 | 54692686 | 6 | 177 | 6.79% | + | - | - | Not pass xTea threshold (cfDNA split-reads = 7, Tumor split-reads = 8) |
| chr1 | 49608812 | 49608854 | 6 | 193 | 6.23% | + | - | - | Not pass xTea threshold (cfDNA split-reads = 2, Tumor split-reads = 0, Tumor discordant pairs = 4) |
| chr10 | 52856193 | 52856226 | 5 | 186 | 5.38% | + | - | - | Not pass xTea threshold (cfDNA split-reads = 0, Tumor split-reads = 0, Tumor discordant pairs = 1) |
| chr3 | 135352916 | 135353541 | 5 | 198 | 5.04% | + | - | - | Not pass xTea threshold (cfDNA split-reads = 3, Tumor split-reads = 0, Tumor discordant pairs = 1) |
| chr4 | 120738758 | 120738760 | 3 | 126 | 4.76% | + | - | - | Not pass xTea threshold (cfDNA split-reads = 3, Tumor split-reads = 11, Tumor discordant pairs = 3) |
| chr11 | 39552615 | 39552698 | 4 | 192 | 4.16% | + | - | - | Not pass xTea threshold (cfDNA split-reads = 0, Tumor split-reads = 0, Tumor discordant pairs = 2) |
| chr5 | 13608580 | 13609205 | 4 | 255 | 3.14% | + | - | - | Not pass xTea threshold (cfDNA split-reads = 0, Tumor split-reads = 0, Tumor discordant pairs = 4) |
| chr4 | 98049814 | 98049851 | 3 | 206 | 2.92% | + | - | - | Not pass xTea threshold (cfDNA split-reads = 0, Tumor split-reads = 0, Tumor discordant pairs = 3) |

### chr12:84700007-84700632

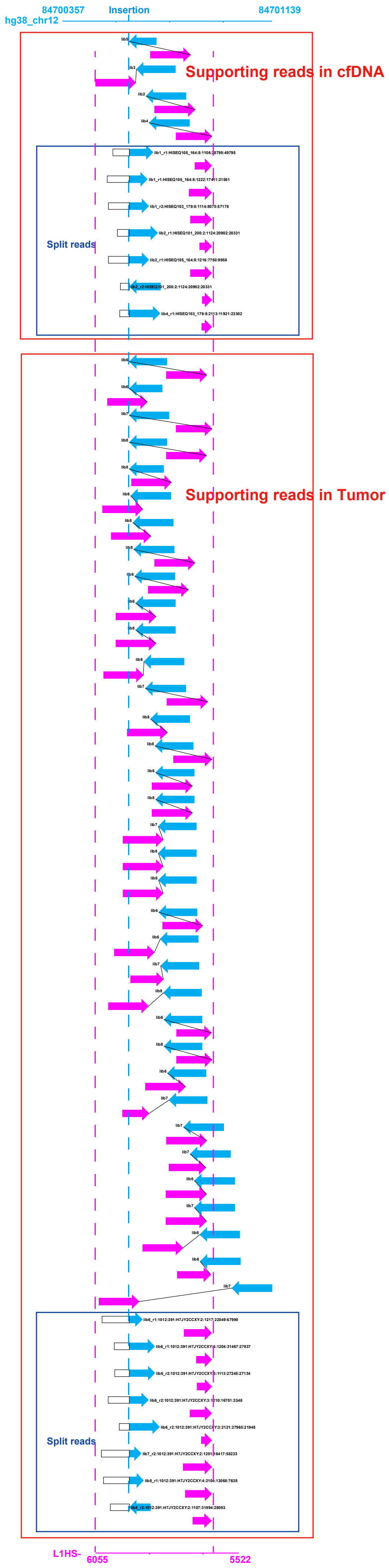

### chr3:16729240-16729865

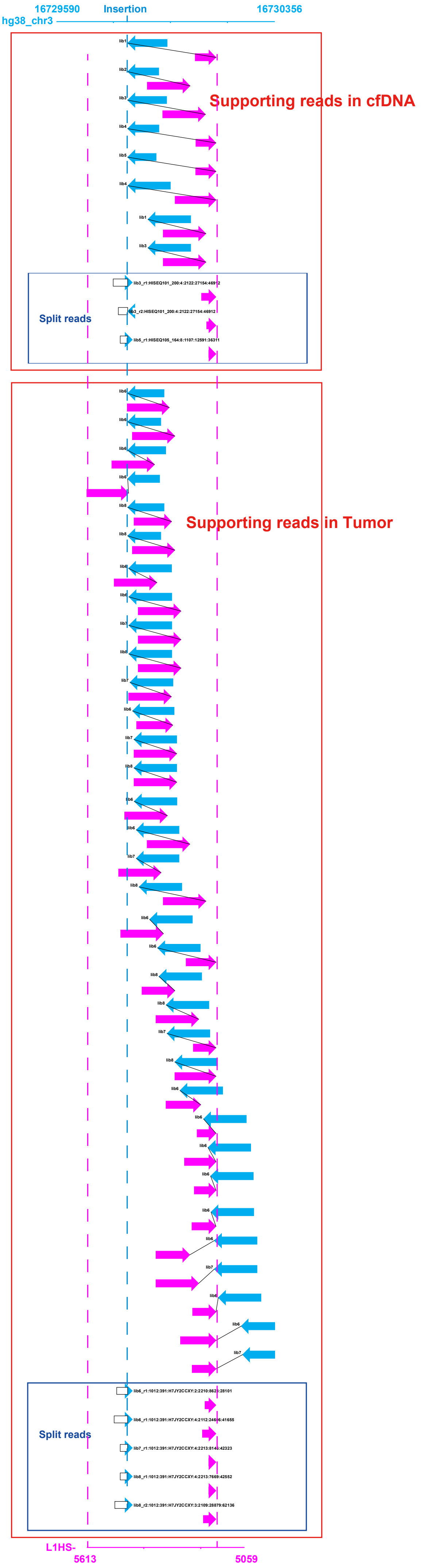

### chr6:90288299-90288337

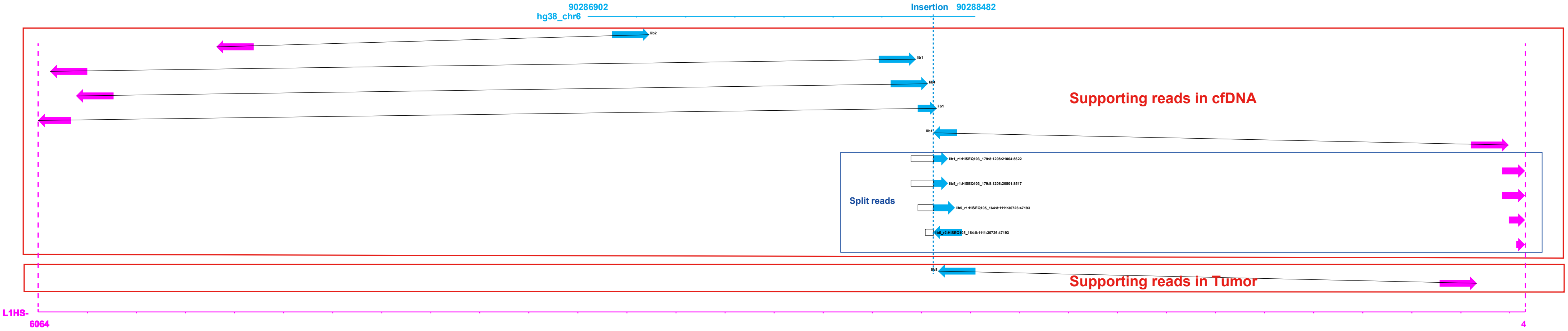

### chr6:2688545-2688595

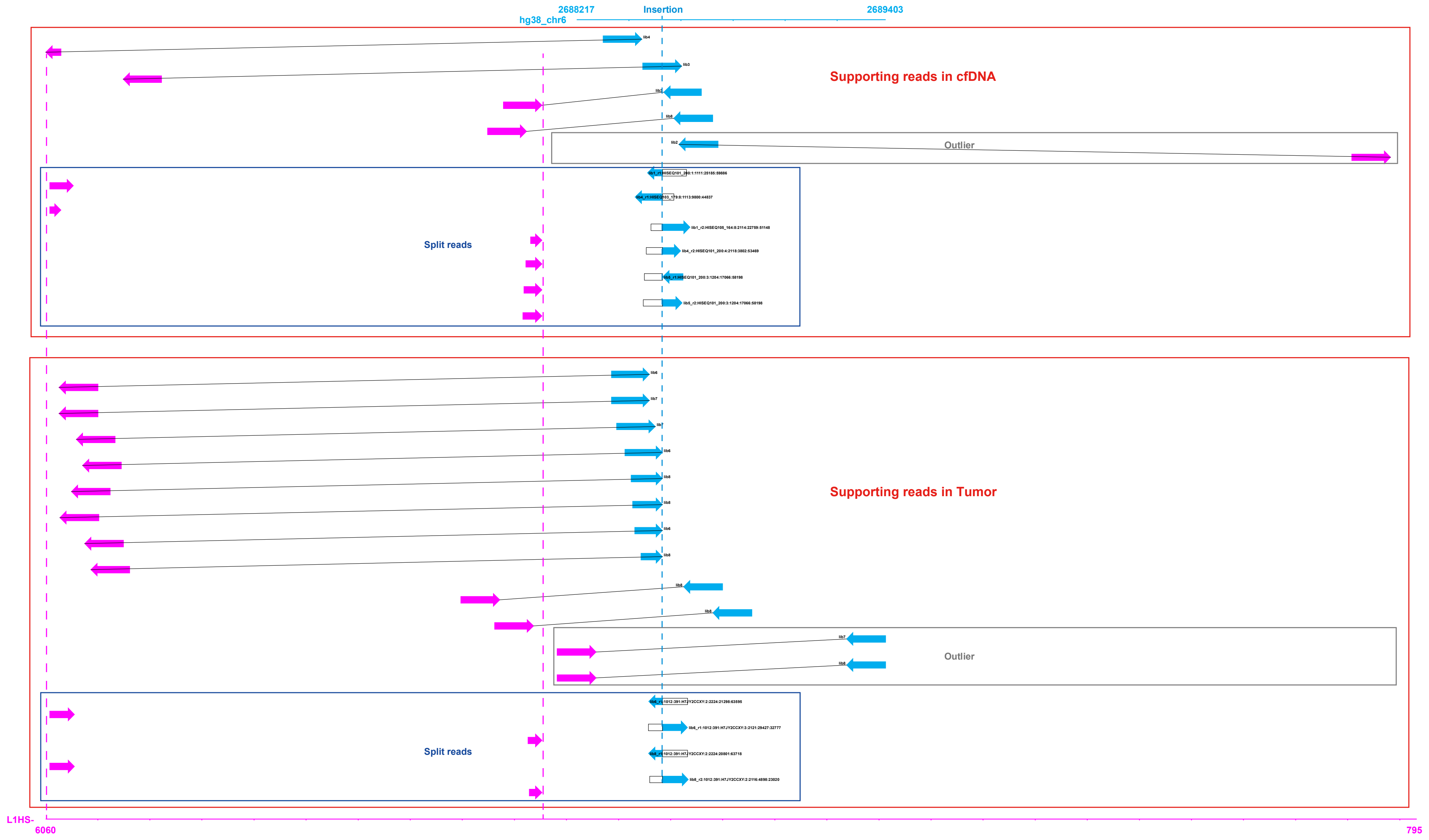

### chr20:18942311-18942351

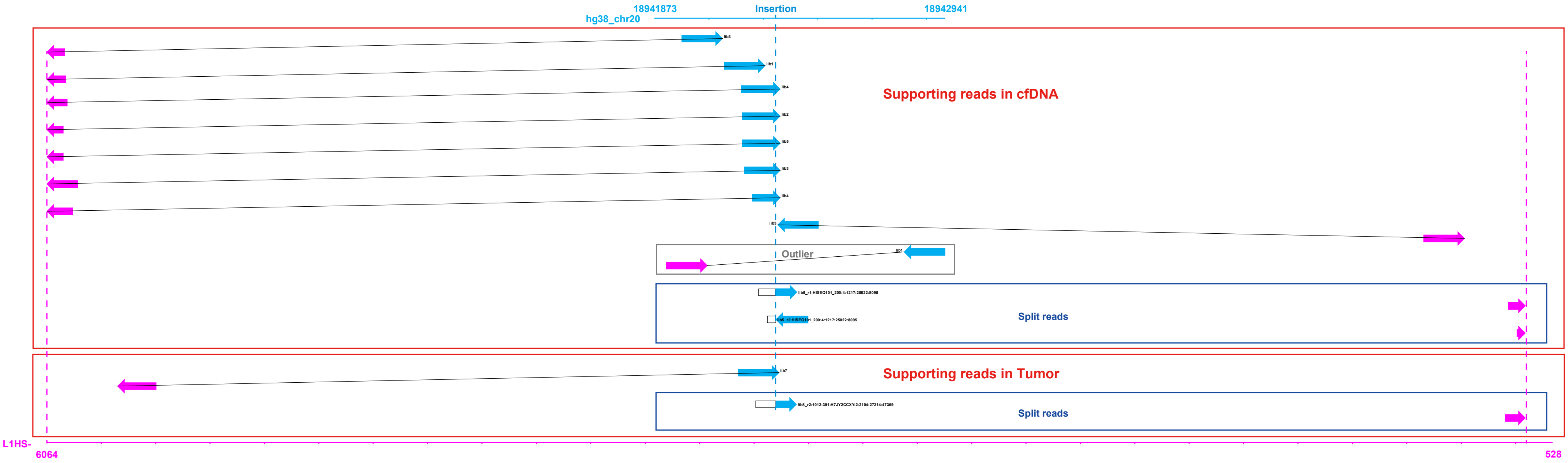

### chr11:86269462-86269495

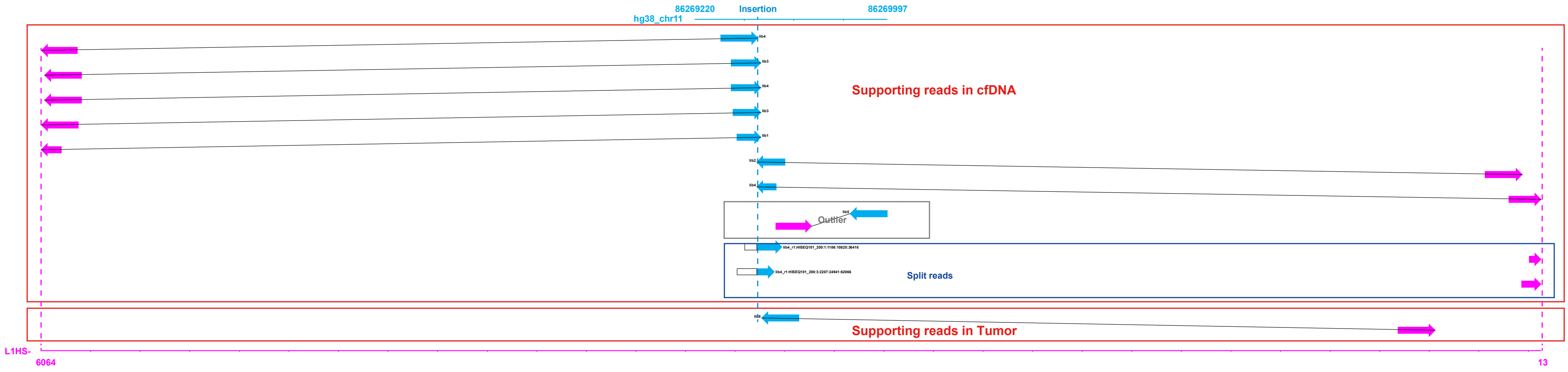

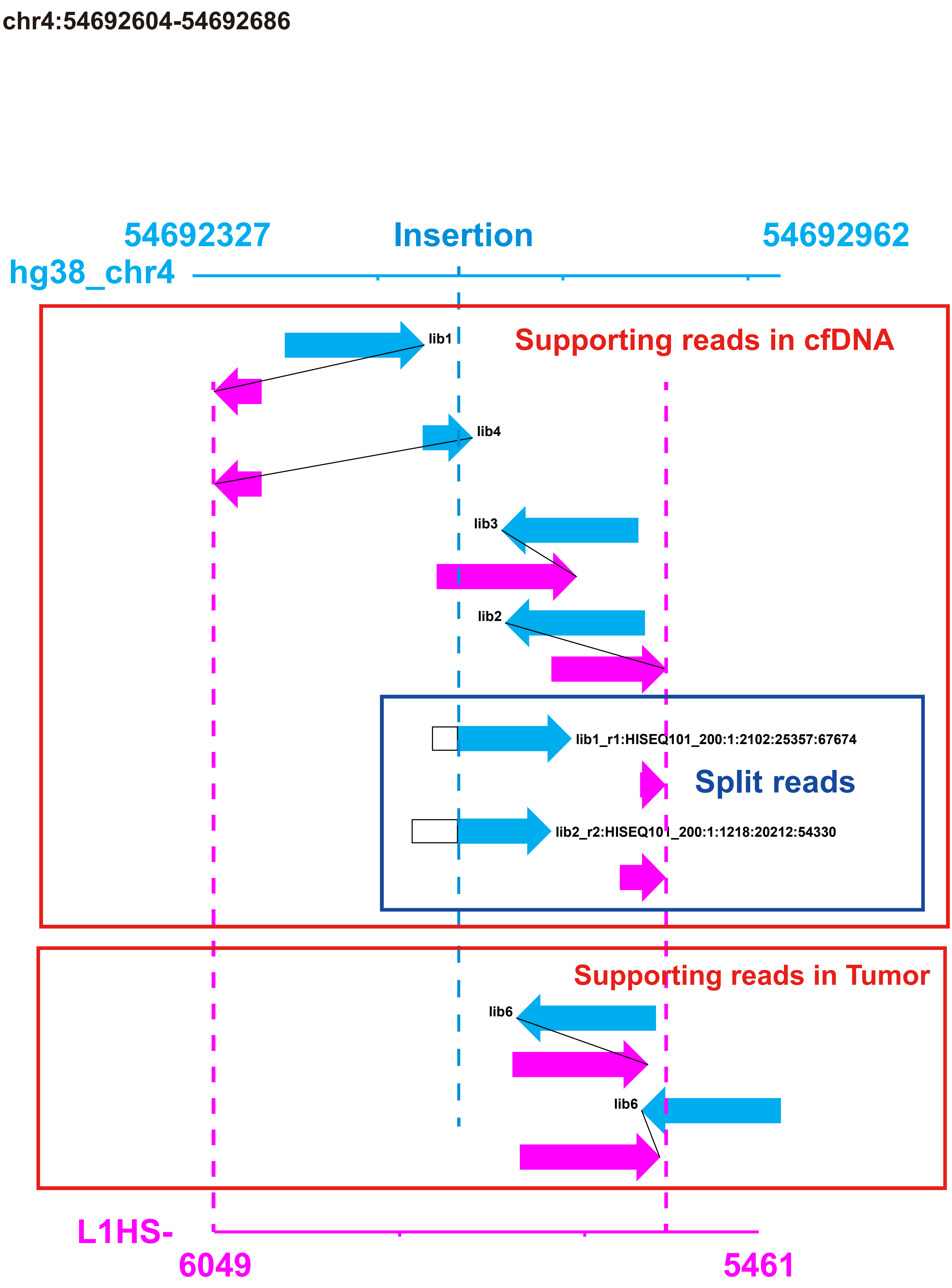
chr4:54692604-54692686
54692327
Insertion
54692962
hg38_chr4
Supporting reads in cfDNA
lib1
lib4
lib3
lib2
lib1_r1:HISEQ101_200:1:2102:25357:67674
Split reads
lib2_r2:HISEQ101_200:1:1218:20212:54330
Supporting reads in Tumor
lib6
lib6
L1HS-
6049
5461

chr1:49608812-49608854

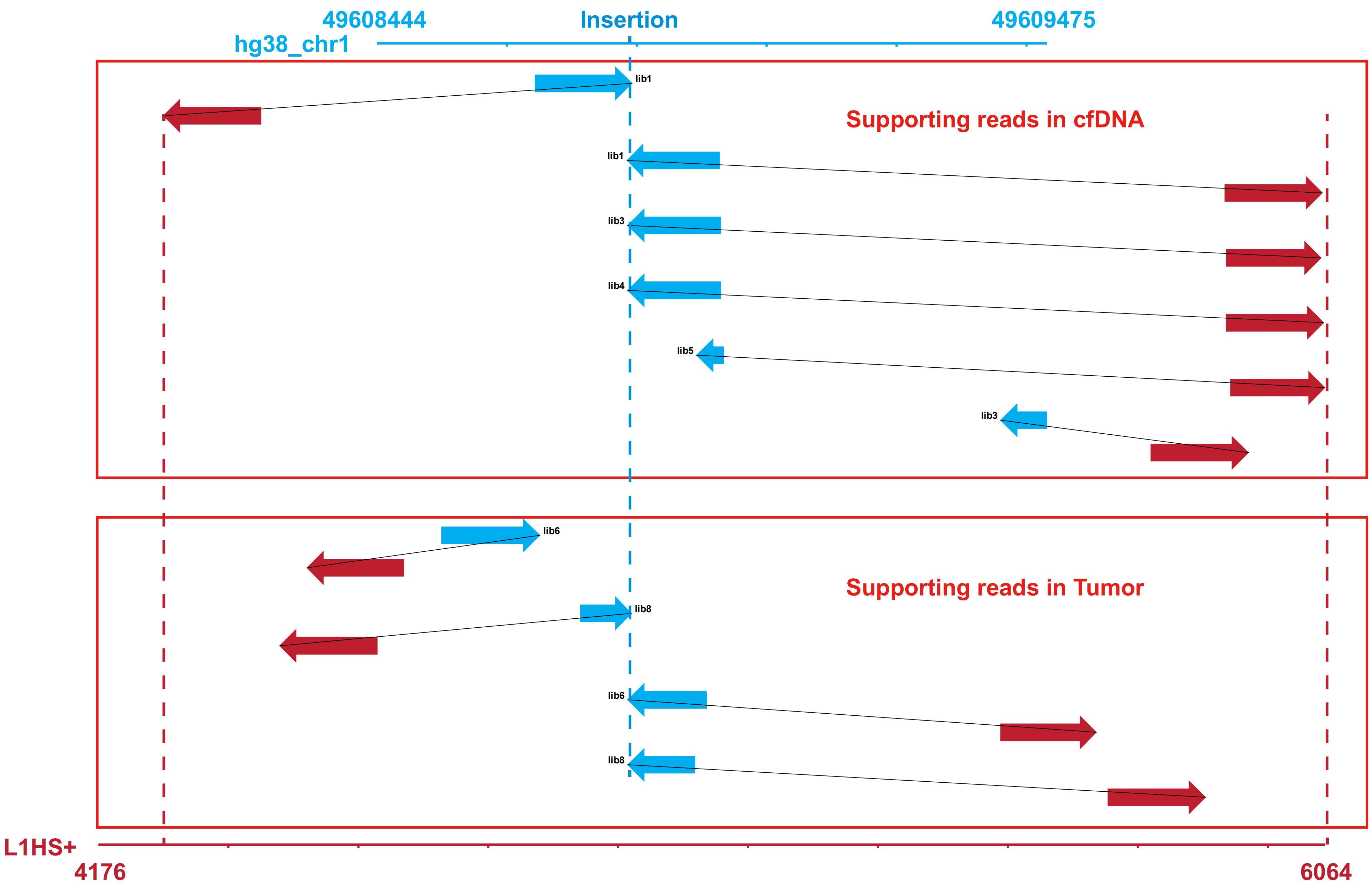

### chr10:52856193-52856226

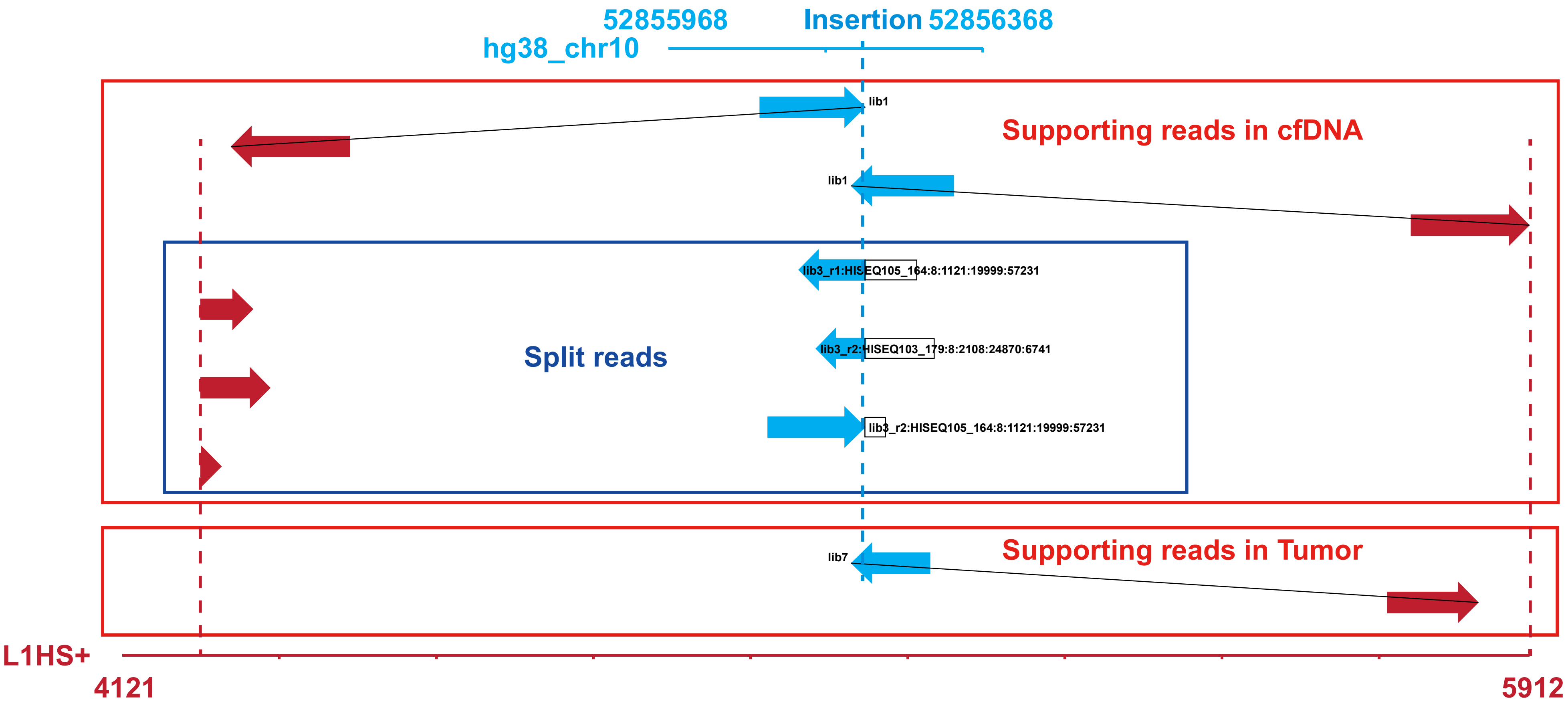

chr3:135352916-135353541

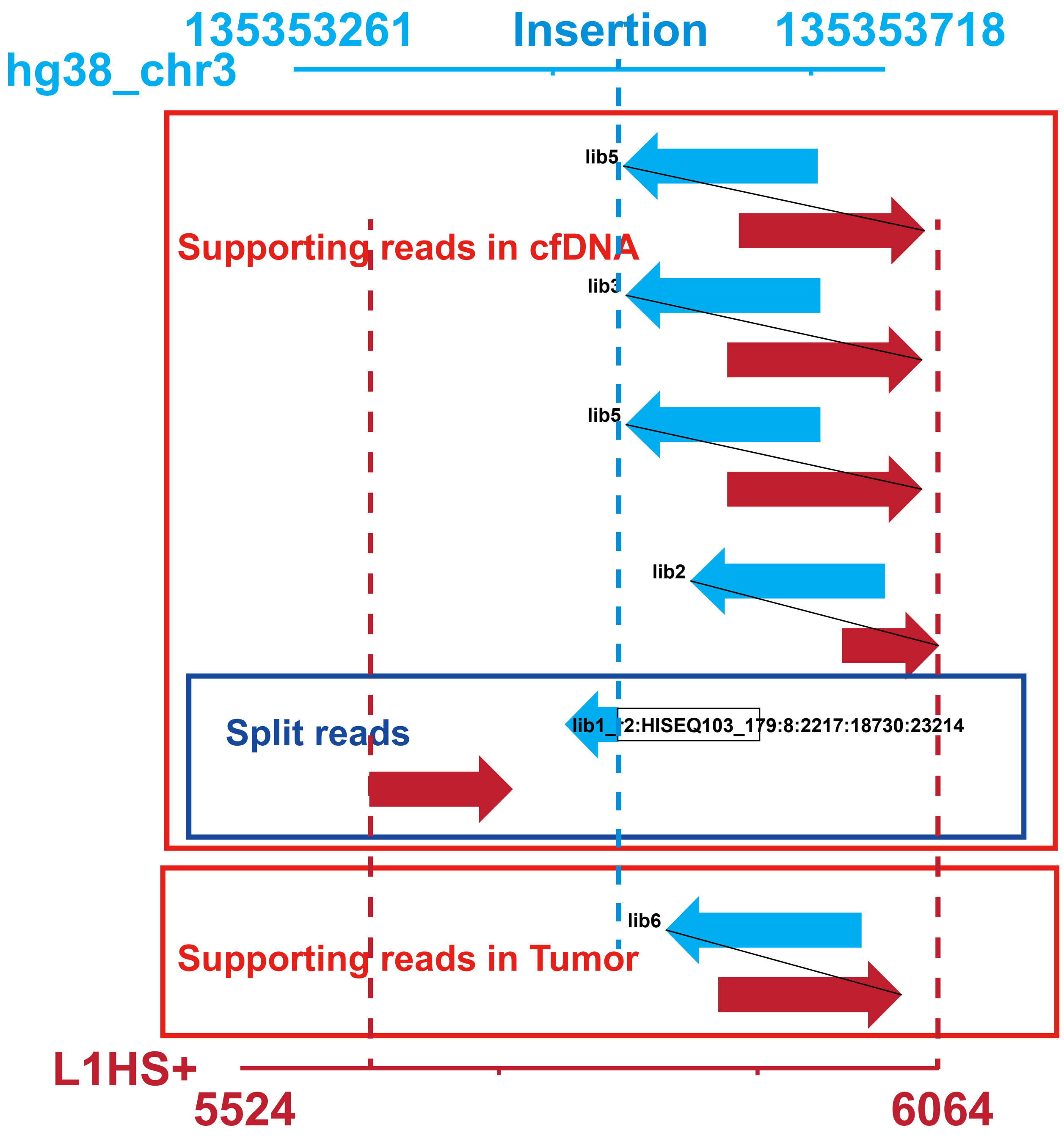

### chr4:120738758-120738760

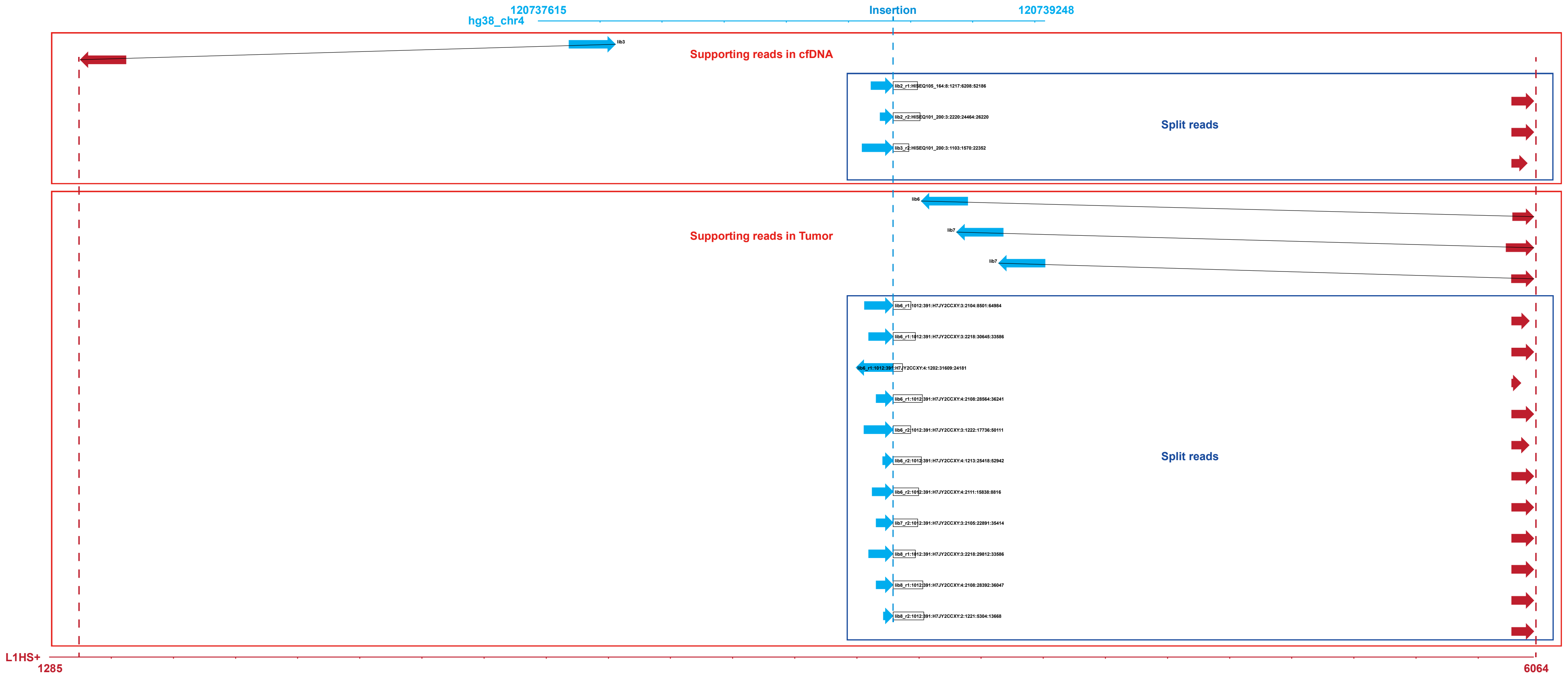

chr11:39552615-39552698

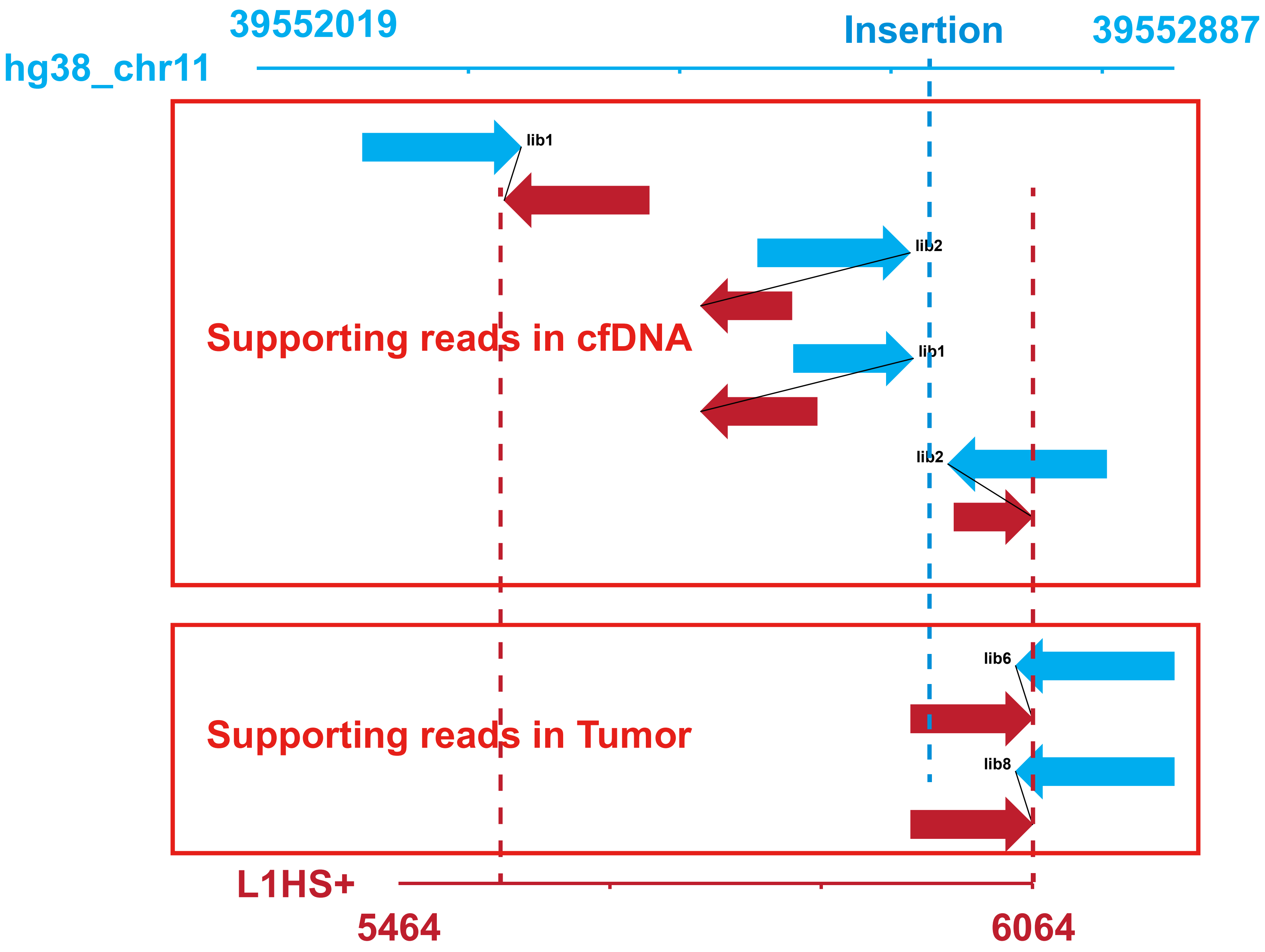

chr5:13608580-13609205

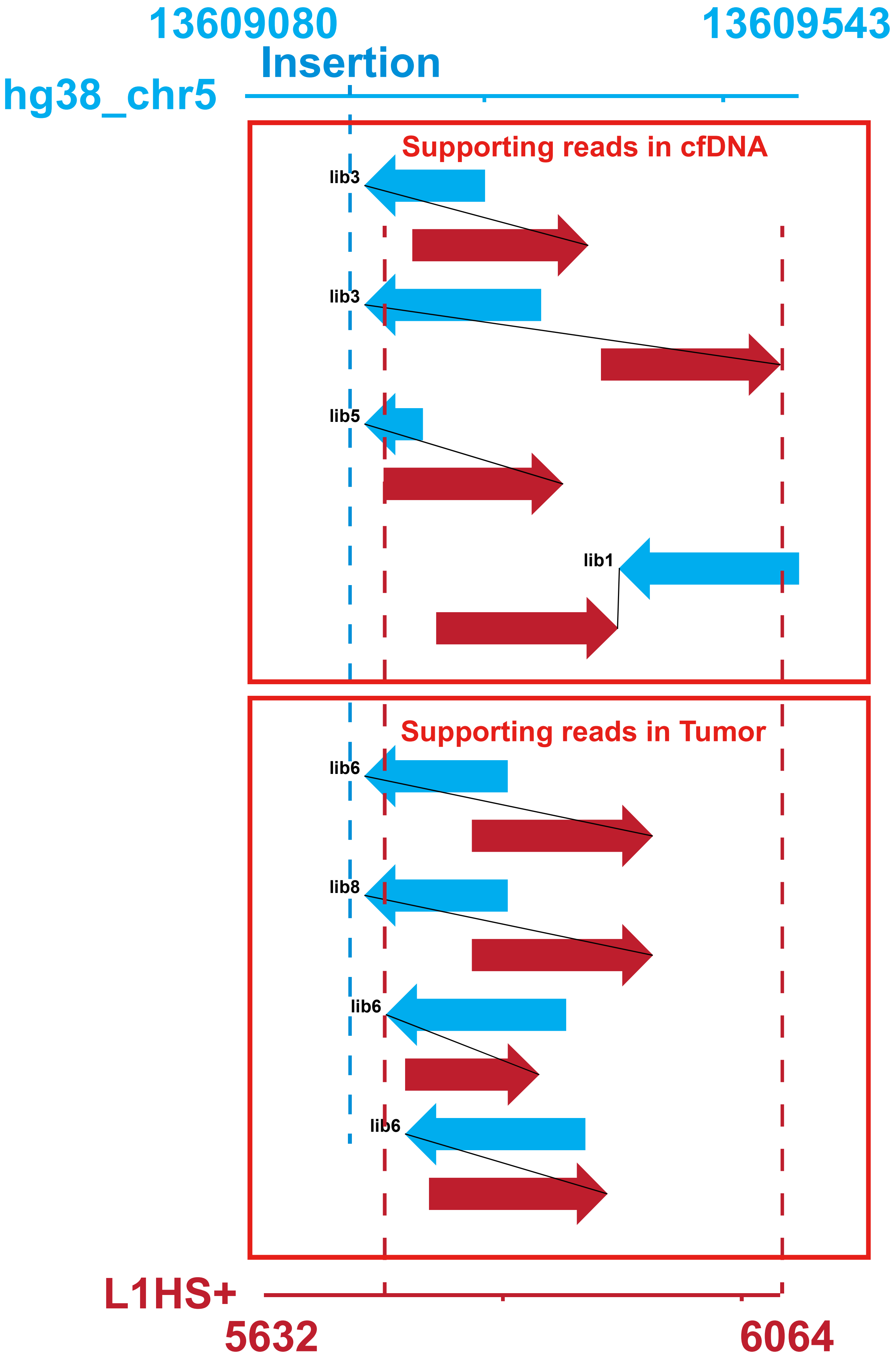

chr4:98049814-98049851

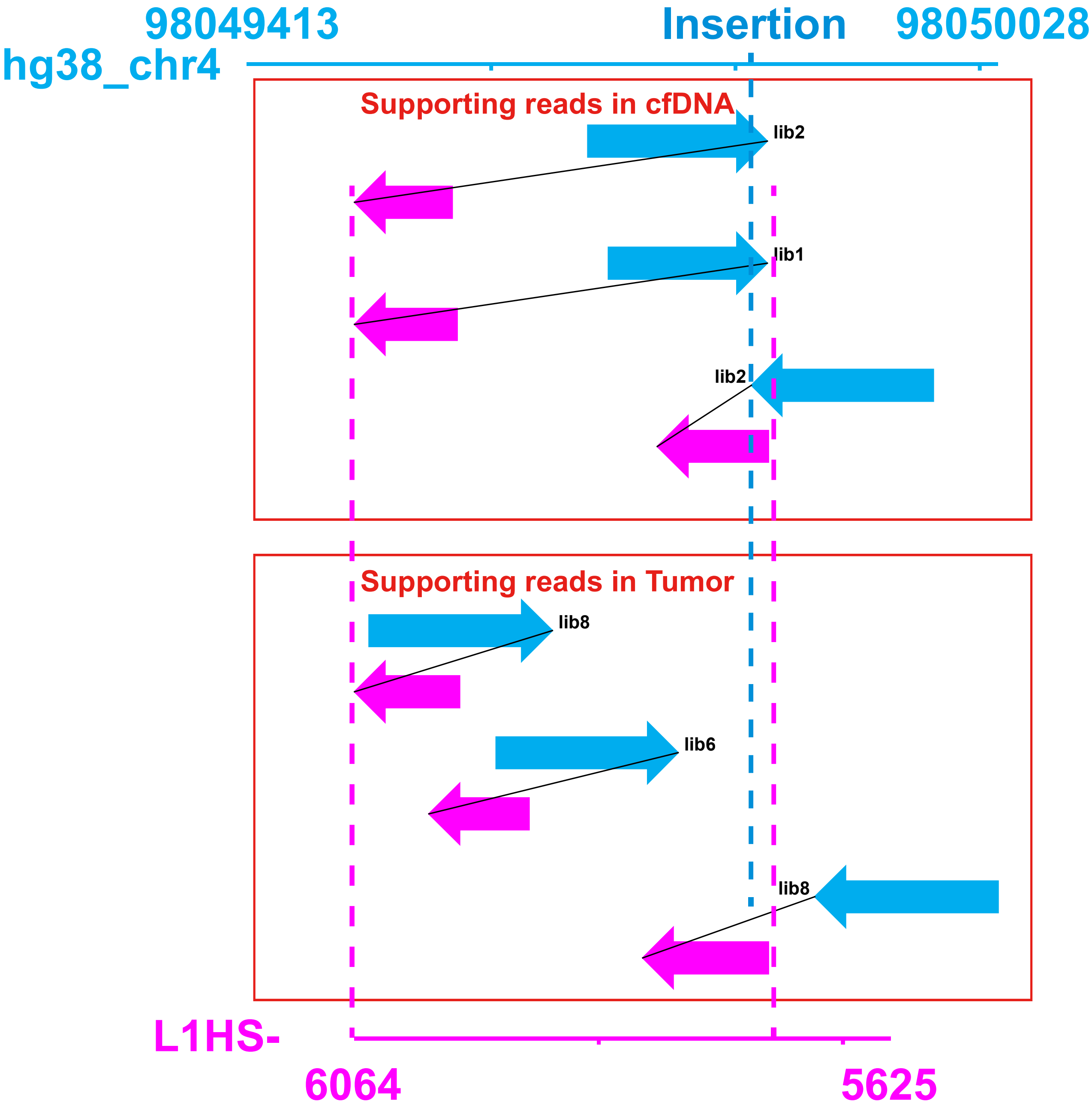
