## Extended Files for "Image-based DNA Sequencing Encoding for Detecting Low-Mosaicism Somatic Mobile Element Insertions": Index of Extended Files.pdf

Extended Files list of article “Image-based DNA Sequencing Encoding for Detecting Low-Mosaicism Somatic Mobile Element Insertions”:

**Extended Files 1**. Manual check of L1 label that were not identified by Pangenome xTea_long PacBio calls.

**Extended Files 2**. Manual check of *Alu* label that were not identified by Pangenome xTea_long PacBio calls.

**Extended Files 3**. Manual check of SVA label that were not identified by Pangenome xTea_long PacBio calls or the Polymorphic SVA dataset.

**Extended Files 4**. Detailed visualization of benchmarked tumor somatic L1 insertions in HG008-T based on PacBio data.

**Extended Files 5**. Benchmark tumor somatic L1 Insertions in HG008-T and their detection by PALMER or xTea_long.

**Extended Files 6**. Tumor somatic *Alu* insertions detected in HG008-T by PALMER or xTea_long.

**Extended Files 7**. Tumor somatic SVA insertions detected in HG008-T by PALMER or xTea_long.

**Extended Files 8**. Detailed visualization of patient DTB-205 cfDNA somatic L1 insertions identified by RetroNet with low mosaicism in the tumor tissue.
